# Adenine DNA methylation, 3D genome organization and gene expression in the parasite *Trichomonas vaginalis*

**DOI:** 10.1101/603894

**Authors:** Ayelen Lizarraga, Zach Klapholz O’Brown, Konstantinos Boulias, Lara Roach, Eric Lieberman Greer, Patricia J. Johnson, Pablo H. Strobl-Mazzulla, Natalia de Miguel

## Abstract

*Trichomonas vaginalis* is a common sexually transmitted parasite that colonizes the human urogenital tract causing infections that range from asymptomatic to highly inflammatory. Recent works have highlighted the importance of histone modifications in the regulation of transcription and parasite pathogenesis. However, the nature of DNA methylation in the parasite remains unexplored. Using a combination of immunological techniques and UHPLC, we analyzed the abundance of DNA methylation in strains with differential pathogenicity demonstrating that N6-methyladenine (6mA), and not 5-methylcytosine (5mC), is the main DNA methylation mark in *T. vaginalis*. Genome-wide distribution of 6mA reveals that this mark is enriched at intergenic regions, with a preference for certain super-families of DNA transposable elements. We show that 6mA in *T. vaginalis* is associated with silencing when present on genes. Interestingly, bioinformatics analysis revealed the presence of transcriptionally active or repressive intervals flanked by 6mA-enriched regions and results from chromatin conformation capture (3C) experiments suggest these 6mA flanked regions are in close spatial proximity. These associations were disrupted when parasites were treated with the demethylation activator ascorbic acid. This finding revealed a new role for 6mA in modulating 3D chromatin structure and gene expression in this divergent member of the Excavata.

**SIGNIFICANCE STATEMENT:** *Trichomonas vaginalis* causes the most common non-viral sexually transmitted infection yet little is known about the regulation of gene expression in this eukaryotic parasite. We demonstrate that N6-methyladenine (6mA) is the main methylation mark in the *T. vaginalis* genome. 6mA is widespread in DNA of eubacterial genera, but uncommon in genomes of most eukaryotes. Examination of the genome-wide distribution of 6mA reveals a preference for intergenic regions. Transcriptionally active or repressive intervals are found to be flanked by 6mA-enriched regions and data suggest 6mA flanked regions are in close 3D spatial proximity. Our findings describe for the first time the presence of DNA methylation in *T. vaginalis* and reveal a new role for 6mA in modulating 3D chromatin architecture.

## INTRODUCTION

*Trichomonas vaginalis* is a flagellated protozoan parasite responsible for trichomoniasis, the most prevalent non-viral, sexually transmitted infection. Although generally asymptomatic (1), *T. vaginalis* infection can cause numerous pathologies such as vaginitis, urethritis, prostatitis and various complications when infection is acquired during pregnancy (2, 3). Chronic infection has been associated with an increased risk of acquiring and transmitting HIV (4) and higher susceptibility to developing cervical or prostate cancer (5, 6).

With ~160Mb distributed over 6 chromosomes, the genome of *T. vaginalis* is the largest protozoan parasite genome sequenced to date (7). To date, the highly repetitive nature of the genome (up to 65% comprised of repetitive elements) prevented the complete assembly of the scaffolds and subsequent studies of *T. vaginalis* chromosome architecture (7). Analysis of the age distribution of gene families coupled with the large genome size and low repeat polymorphism indicates the genome underwent one or more large-scale genome duplication events (7). Intriguingly, only half of the ~46,000 protein-coding genes appear to be expressed (8, 9) and individual members within gene families show differential regulation in response to different stimuli (8–10). These results indicate that extrinsic and intrinsic factors accurately control the fine tuning of gene expression in this extracellular parasite.

Studies on *T. vaginalis* transcriptional regulation mostly focused on identifying cis-regulatory elements and their cognate transcription factors. Few core promoter elements or transcription factors have been identified to date (11). The metazoan-like Initiator (Inr) element (12) present in ~75% of all protein coding genes (7) was found to be the main core promoter element responsible for directing basal transcription in *T. vaginalis*. A novel initiator binding protein, IBP39, was identified and demonstrated to specifically recognize *T. vaginalis* Inr and directly interact with RNAPII (12, 13). Additionally, several conserved motifs that resemble metazoan Myb recognition elements (14) as well as proteins containing a Myb DNA binding domain, like novel transcription factor M3BP (14), have also been found in *T. vaginalis* (11). This Myb recognition elements can act as cis-regulatory elements directing the transcription start site selection (14) and transcription of specific genes (15). However, little is known about other mechanisms of gene regulation in this parasite (11). Epigenetic mechanism regulates gene expression by modifying chromatin accessibility to transcription factors and other components of the transcriptional machinery (16). Specifically, histone modifications have been shown to be of importance in the regulation of gene expression and antigenic variation of other unicellular parasites (17). Recent studies have demonstrated that these modifications are present in *T. vaginalis* and play an essential role in transcriptional regulation and parasite pathogenesis (18, 19).

DNA methylation is one of the major epigenetic mechanisms involved in gene regulation. The most common DNA modification in eukaryotes is C5-methylcytosine (5mC), an epigenetic mark usually associated with gene silencing (20) that participates in a wide variety of processes such as imprinting, X chromosome inactivation, and silencing of transposable elements (21). Though common among higher eukaryotes, 5mC is not universally present in unicellular organisms (17) and thus few protozoan parasites with detectable levels of 5mC in their genome have been identified and its function characterized (17, 22–24). In contrast, N6-methyladenine (6mA) is the most prevalent DNA modification in prokaryotes (25) and early reports debated the existence and abundance of this mark in eukaryotes (26). However, in recent years numerous reports have greatly expanded the list of organisms with 6mA to include flies (27), nematodes (28), green algae (29) and even vertebrates (30–32). The abundance of this mark varies dramatically between species, with levels of 6mA that range from 6-7 parts per million (p.p.m.) of total adenines in mouse embryonic stem cells (30) up to 2.8% of total adenines in early-diverging fungi (33). Likewise, this variation between organisms was also evidenced in their genomic distribution, suggesting that 6mA might have species-specific functions (34). Intriguingly, in some cases the role of 6mA has been associated with active transcription in *Arabidopsis* (35), *Drosophila* (27), *Chlamydomonas* (29) and early-diverging fungi (33). Only two reports describe 6mA as a repressive mark similar to 5mC (30, 36).

Despite its importance in other organisms, the methylation status of *T. vaginalis* DNA remains unexplored. In this work, using a combination of immunological techniques and UHPLC coupled with mass-spectrometry we identify the presence of 6mA and 5mC in the gDNA of *T. vaginalis*. We interrogate the distribution of 6mA across the genome using an adapted methylated DNA immunoprecipitation assay followed by high-throughput sequencing (MeDIP-seq). Interestingly, we found that 6mA is enriched at intergenic regions, with a preference for repetitive elements. Comparison with published RNA-seq data of the same strain (37) suggests 6mA is associated with silenced transcription when found on gene bodies. Finally, results from chromatin conformation capture experiments suggest that 6mA is associated with chromatin loop formation, raising the intriguing possibility that 6mA could have a role in modulating 3D genome architecture. This is the first time DNA methylation is described in *T. vaginalis* and our data highlight a role for 6mA in epigenetic gene regulation in this deep-branching eukaryote.

## RESULTS

### Identification of 6mA and 5mC in *T. vaginalis* genomic DNA

Given the importance of 5mC and 6mA in other organisms, we set out to investigate the presence of these marks in *T. vaginalis* genomic DNA. We first evaluated the presence of 5mC in *T. vaginalis* by a dot blot assay using genomic DNA extracted from two *T. vaginalis* strains, B7268 and G3, with high and low adherence capacity, respectively (Fig. 1A). Unexpectedly, we failed to detect 5mC in either strain, contrary to our chicken gDNA control (Fig. 1A). Likewise, although we were able to detect m5C in the RNA, this mark was undetectable by immunofluorescence in the nucleus of the parasite (Fig. 1B). These data suggest the absence, or very low abundance, of 5mC mark in *T. vaginalis* DNA. On the other hand, we revealed the presence of 6mA mark both in the nucleus and cytoplasm by immunofluorescence (Fig. 1C) using antibodies that specifically recognize methylated adenines (Sup. Fig. 1A). Importantly, parasites treated with RNAse revealed 6mA exclusively detected in the nucleus demonstrating that this epigenetic mark is present in the DNA of *T. vaginalis*. Similarly, using restriction enzymes MboI and DpnI which digest unmethylated and methylated GATC sequence respectively (38), we found that *T. vaginalis* genome possesses both methylated and un-methylated adenines in the GATC context (Fig. 1D). In order to determine whether parasite adherence could be correlated with 6mA abundance, we performed dot blot assays (Fig. 1E) and UHPLC followed by mass spectrometry (Fig. 1F) using gDNA from an adherent (B7268) and a less-adherent (G3) strain. However, we found no difference on the abundance of 6mA levels among them. Importantly, we demonstrated that 6mA is present in 2.5% of all adenines, whereas extremely low levels of 5mC was detected (0.004-0.008% of all cytosines) by UHPLC followed by mass spectrometry (Fig. 1F). To exclude the possibility of signal contamination from *M. hominis*, a common bacterial symbiont of *T. vaginalis* (39), parasite cultures were subjected to antibiotic treatment and checked for the elimination of *M. hominis* prior to the experiments (Sup. Fig. 1B). These data demonstrate the lack of the bacterial symbiont and confirm that the detected 5mC and 6mA is in *T. vaginalis* genomic DNA. Taken together, our results demonstrated that 6mA, but not 5mC, is the predominant DNA methylation mark in *T. vaginalis*.

**Figure 1:**
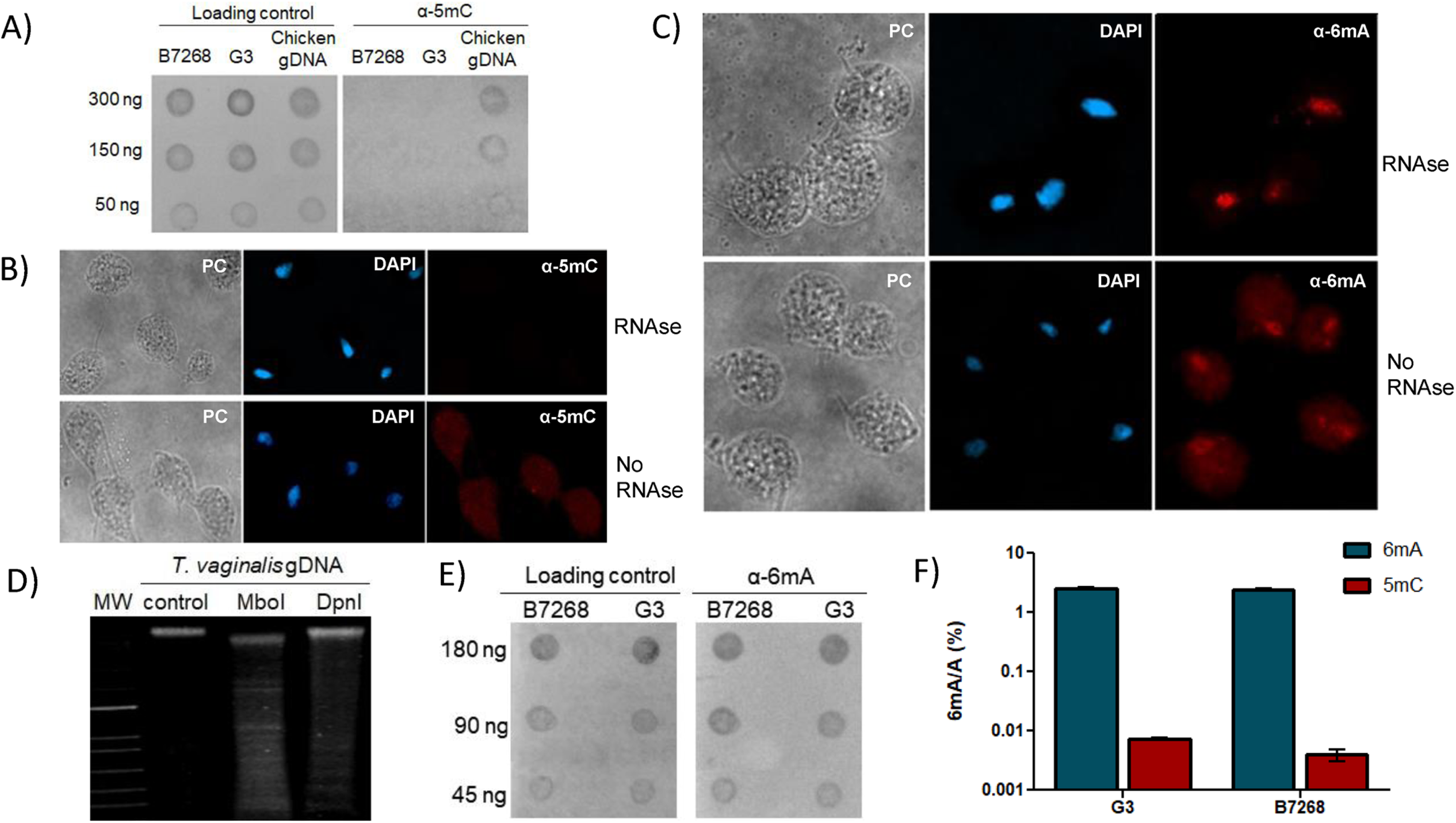
DNA methylation in *T. vaginalis* genome. A) Dot blot assay to detect 5mC shows lack of signal in the genomic DNA of highly-adherent strain B7268 and less-adherent strain G3. Left: gDNA dots stained with methylene blue for loading control. Right: 5mC signal. Chicken gDNA was used as positive control. B) Immunofluorescence assay using specific antibodies detects 5mC in the cytoplasm (RNA) but fails to detect 5mC in *T. vaginalis* nucleus of RNase-treated parasites. Nucleus was stained with DAPI. C) Immunofluorescence assay using anti-6mA specific antibody evidenced this modification in the cytoplasm (RNA) and nucleus (gDNA), or exclusively in the nucleus of RNase-treated parasites. Nucleus was stained with DAPI. D) Methylation sensitive RE assay shows the presence of methylated and un-methylated adenines in *T. vaginalis* gDNA. Lane 1: MW. Lane 2: undigested gDNA. Lane 3: MboI treated gDNA (cuts GATC sites). Lane 4: DpnI treated gDNA (cuts G6mATC sites). E) Anti-6mA dot blot assay of RNA-free gDNA shows similar abundance of 6mA in the genomic DNA of strain B7268 and strain G3. Left: gDNA dots stained with methylene blue for loading control. Right: 6mA signal. F) Quantification of methylated bases in the gDNA of adherent strain B7268 and less-adherent strain G3 using UHPLC followed by mass spectrometry shows high levels of 6mA and low levels of 5mC, but not differences among stains for both modifications. Each column represents the mean and SD of three independent experiments per group. Graphic is shown in logarithmic scale.

To be considered an epigenetic mark, 6mA needs to be maintained after DNA replication. Our observations using hydroxyurea synchronized cultures revealed the expected decrease in 6mA levels in S phase due to DNA replication, which remained during the G2/M phase of the cell cycle, and restored original abundance after cell division (Sup. Fig. 2). These observations support the presence of active methyltransferase(s) responsible for maintaining 6mA levels after cell division.

### Genome-wide mapping of 6mA in *T. vaginalis*

To determine the function of DNA methylation in *T. vaginalis* it is essential to identify its genomic distribution. To this end, we performed an adapted methylated DNA immunoprecipitation technique using specific antibodies followed by high-throughput sequencing (MeDIP-seq) (40) on the adherent strain B7268 (Fig. 2A). Due to the extremely low levels of 5mC in *T. vaginalis*, we focused exclusively on 6mA for further analyses. To this end, two independent high-throughput sequencing experiments, each containing pooled DNA from three independent immunoprecipitation assays, were performed. In order to evaluate specificity and enrichments of 6mA in the immunoprecipitated fraction when using a specific anti-6mA antibody, a dot blot assay was performed using an IgG control antibody (Fig. 2B).

**Figure 2:**
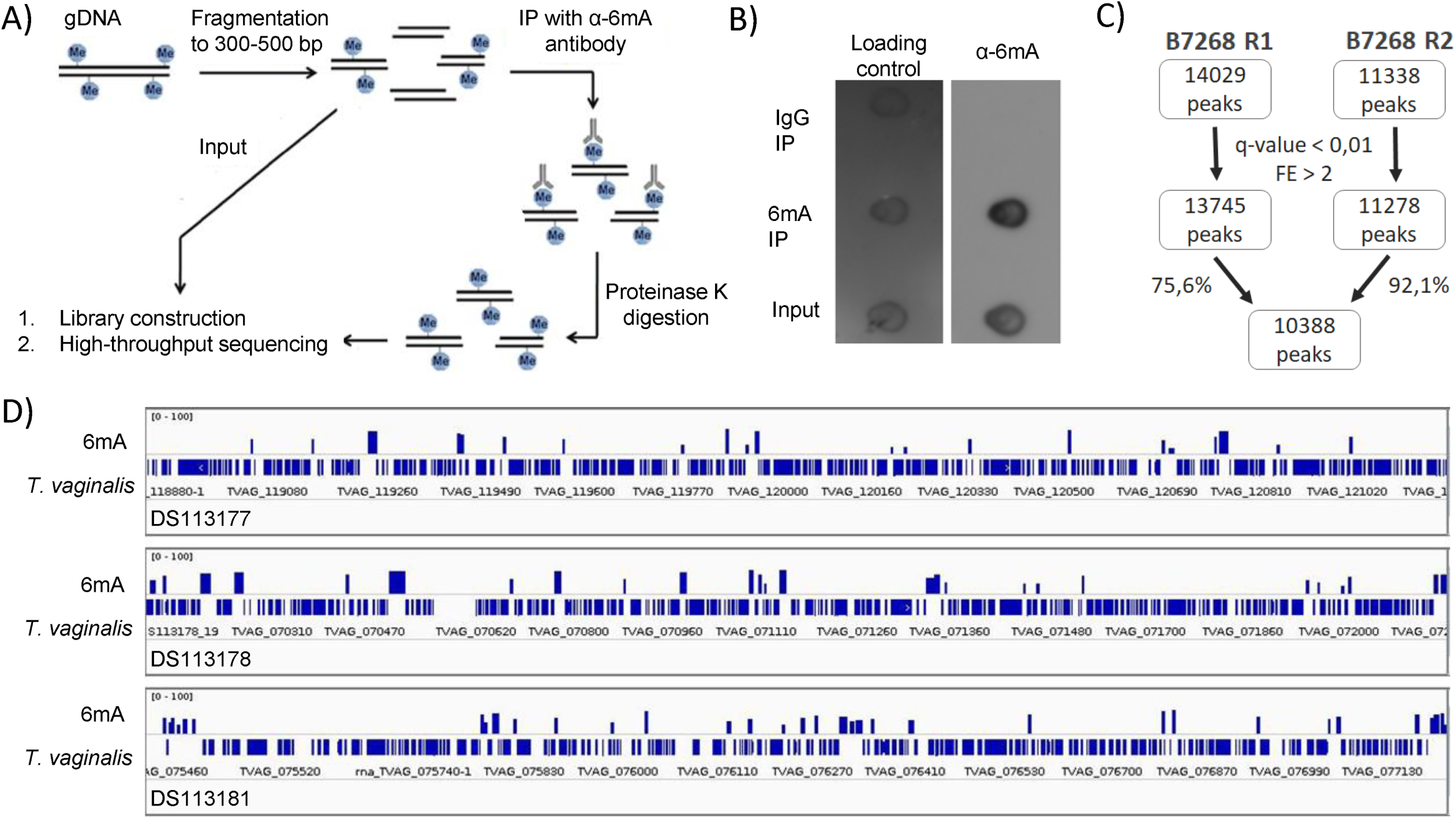
6mA MeDIP-seq of adherent strain B7268. A) Schematic representation of the adapted MeDIP-seq protocol. B) Dot blot assay of the different MeDIP fractions shows enrichment of 6mA-containing fragments in the 6mA-IP fraction compared to the input control. Lack of signal in the IgG-IP fraction shows the specificity of 6mA antibody. C) Workflow showing final peak count obtained from two independent 6mA MeDIP-seq experiments. Peaks were filtered by q-value (<0.01) and fold enrichment (>2). Only peaks with at least 75% overlap between experiments were considered for further study. D) Genome browser visualization of 6mA peaks. Representative image of three different contigs (DS113177, DS113178 and DS113181) shows 6mA possesses a primarily intergenic localization. To row: 6mA peaks. Bottom row: *T. vaginalis* genome

After quality filtering (q-value < 0,01, FE > 2) we obtained a total of 10388 6mA peaks found in both replicates, corresponding to 92,1% and 75,6% of all peaks present in each independent MeDIP-seq experiment (Fig. 2C, Sup. Table 1). In order to validate these results, four genes selected from the MeDIP-seq were analyzed by MeDIP-qPCR demonstrating an enrichment in 6mA immunoprecipated sample compared to sample immunoprecipitated with IgG control antibody (Sup. Fig. 3A). The most prevalent motif, found in 4692 peaks, was CAATACCC (p-value = 1e-1642, Sup. Fig. 3B). Remarkably, peak visualization in a genome browser suggested a clear enrichment of peaks in intergenic regions (Fig. 2D). Indeed, peak annotation revealed that 94% of all MeDIP-seq peaks (9746 6mA peaks) were intergenic (Fig. 3A). Importantly, a shuffled distribution of peaks across *T. vaginalis* genome would result in an even distribution between genes and intergenic regions (Fig. 3A), confirming 6mA is remarkably enriched in intergenic regions.

**Figure 3:**
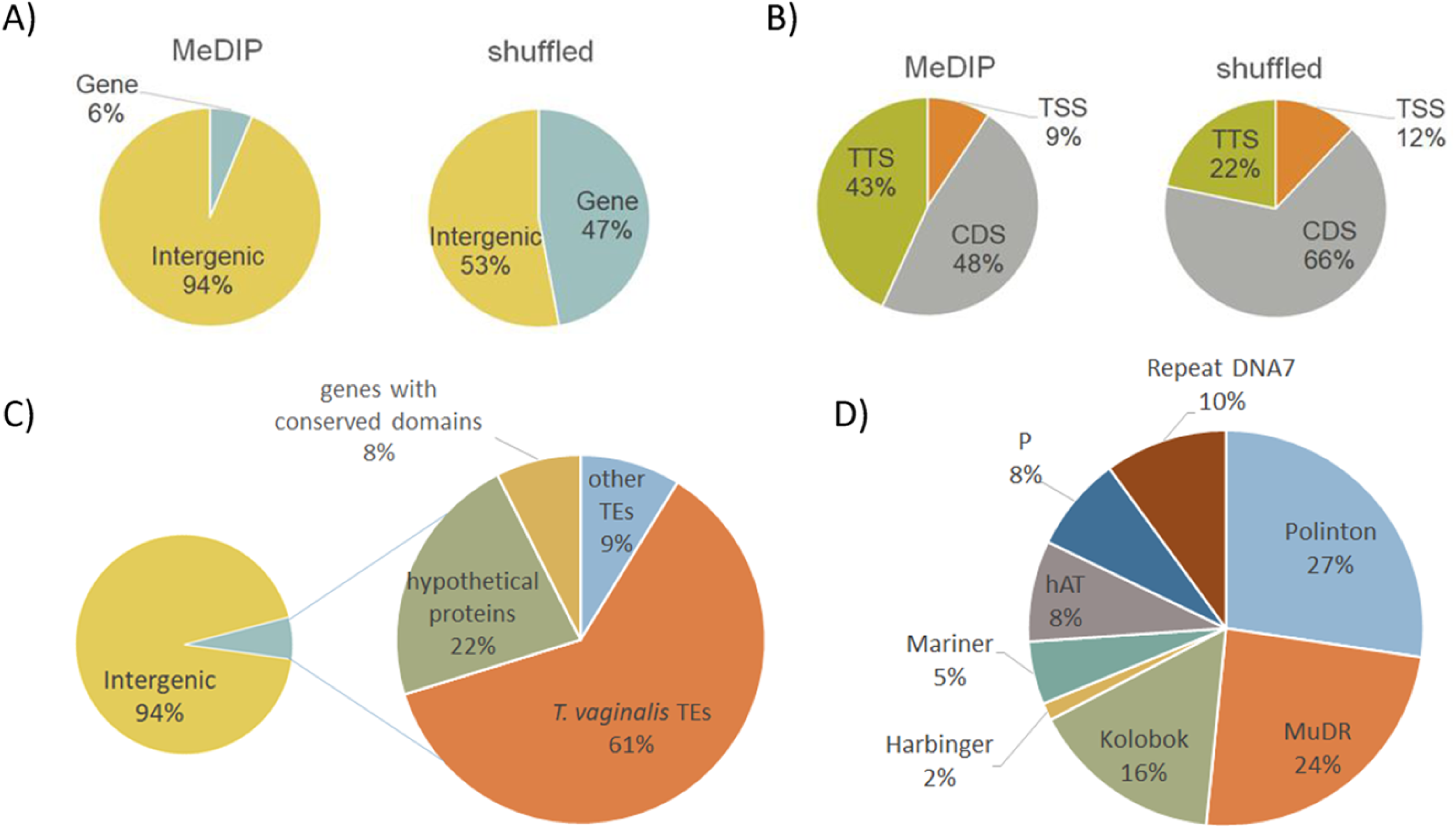
Genome distribution of 6mA peaks. A) 6mA is enriched at intergenic regions. Annotated peaks were grouped according to their genomic localization: peaks that fall within the transcription start sites (TSS), coding region and transcription termination sites (TTS) were considered gene peaks and grouped together, the rest of the peaks were considered intergenic. MeDIP: Percentages of intergenic and gene peaks according MeDIP-seq analysis. Shuffled: MeDIP-seq peaks were randomly distributed across the genome and the percentage of gene and intergenic peaks was calculated. B) Enrichment of 6mA at the TTS region of methylated genes. MeDIP: Percentages of peaks within the TSS, CDS and TTS regions of methylated genes according MeDIP-seq analysis. Shuffled: Peaks randomly distributed across the genome. C) Identity of *T. vaginalis* methylated genes. A total of 642 MeDIP peaks fall within 512 genes: 61% belong to transposable elements described in the parasite (*T. vginalis* TEs), 9% of genes possess sequence similarity to repetitive elements found in other organisms (other TEs), 22% are *T. vaginalis* specific hypothetical proteins with unknown functions (hypothetical proteins), 8% are genes with sequence similarity to genes from other organisms (genes with known domains). D) Identity of *T. vaginalis* methylated repetitive elements. Only intergenic peaks with more than 50% overlap with a *T. vaginalis* RE were considered, obtaining a total of 5596 peaks. Most represented DNA transposon superfamilies included Mutator (MuDR) with 24% of all peaks, Polinton (27%) and Kolobok (16%). Less represented TE superfamilies included hAT (8%), Mariner (5%) and P (8%). 10% of the peaks were found in *T. vaginalis* DNA repeat DNA7.

Among the 642 6mA peaks located in genes, most were distributed between the coding region (48%) and the transcription termination sites (TTS) (43%), with only 9% found in the transcription start sites (TSS) (Fig. 3B). Comparing those numbers to the percentages that would occur if the peaks had a shuffled distribution (22% in the TTS, 60% in the coding region and 12% in the TSS) our results revealed an enrichment of 6mA in the TTS when located in genes (Fig. 3B). In order to understand the functional role of 6mA, we retrieved the IDs of the methylated genes (Sup. Table 2) and found 70% belong to different families of transposable elements, with the majority (61%) belonging to families previously described in *T. vaginalis* according to Repbase (Fig. 3C). The remaining genes could be divided into ones with conserved domains that allow the assignment of a predicted role (8%) and hypothetical proteins with unknown functions (22%) found exclusively in *T. vaginalis* (Fig. 3C). Intriguingly, the observed enrichment of 6mA peaks in the TTS was found exclusively in transposable element (TE) genes, suggesting a particular role for methylation at the 3’ of TE genes. Gene ontology analysis of the methylated genes (excluding genes from TEs) revealed an enrichment of genes involved in protein modification and phosphorylation (p-value < 0.05) (Sup. Table 3). However, it is important to take into account that most of non-TE genes have no identifiable domain (hypothetical proteins) and therefore were not considered in the GO enrichment analysis.

Considering that 6mA has been previously described as an epigenetic mark of transposable elements in other organisms (27, 30) and 70% of *T. vaginalis* methylated genes belong to TEs, we analyzed if intergenic peaks were also located within TEs. Analysis using RepeatMasker revealed that out of 94% of intergenic peaks, 57.4% were found in *T. vaginalis* repetitive elements (RE) previously described in Repbase. Most of the peaks were found in Mutator, Kolobok, and Polinton superfamilies of *T. vaginalis* DNA transposons (Fig. 3D). However, if we consider the numbers of TEs within the genome, repeat element DNA7 and TE super-families P and Kolobok were the most enriched in the MedIP-seq (Sup. Table 4), indicating a preference of intergenic 6mA for certain types of REs and raising the question of whether 6mA could act as a regulator of specific TEs.

### 6mA methylation and gene expression

6mA has been found to be an epigenetic mark of active genes in *Arabidopsis* (35), *Drosophila* (27) and *Chlamydomonas* (29) or repressed genes in mouse embryonic stem cells (30) and rice (36). To examine whether this mark has a role in transcription in *T. vaginalis*, we made use of publicly available RNA-seq data from B7268 strain (37). To test if this data was applicable for our analysis, we divided the genes into low (RPKM = < 1), moderate (1 < RPKM =< 20), and high (RPKM > 20) expression, and selected some well characterized genes from each group to measure their relative expression levels via RT-qPCR. Our results showed the relative expression of each gene was in accordance to what had been described in bibliography (19, 41) and more importantly, we could order the genes based on their relative expression obtained by RT-qPCR similarly as they have been ordered based on their RPKM (Sup. Table 5), validating the use of the RNA-seq in our analysis.

We next looked into the expression levels of the methylated genes using RNA-seq data available and maintaining the same criteria for low, moderate and high expression. Considering that the majority of those genes belong to *T. vaginalis* transposable elements (Fig. 3C), it was not surprising that gene expression analysis revealed over 80% of all methylated genes are poorly expressed, with 14% possessing moderate expression and only a small percentage (4%) highly expressed (Fig. 4A). However, when we considered the expression levels of the methylated genes without taking into account TE genes, poorly expressed genes were still the most abundant group (over 50%, Sup. Table 6), suggesting 6mA could be a sign of repressed expression when found on genes.

**Figure 4:**
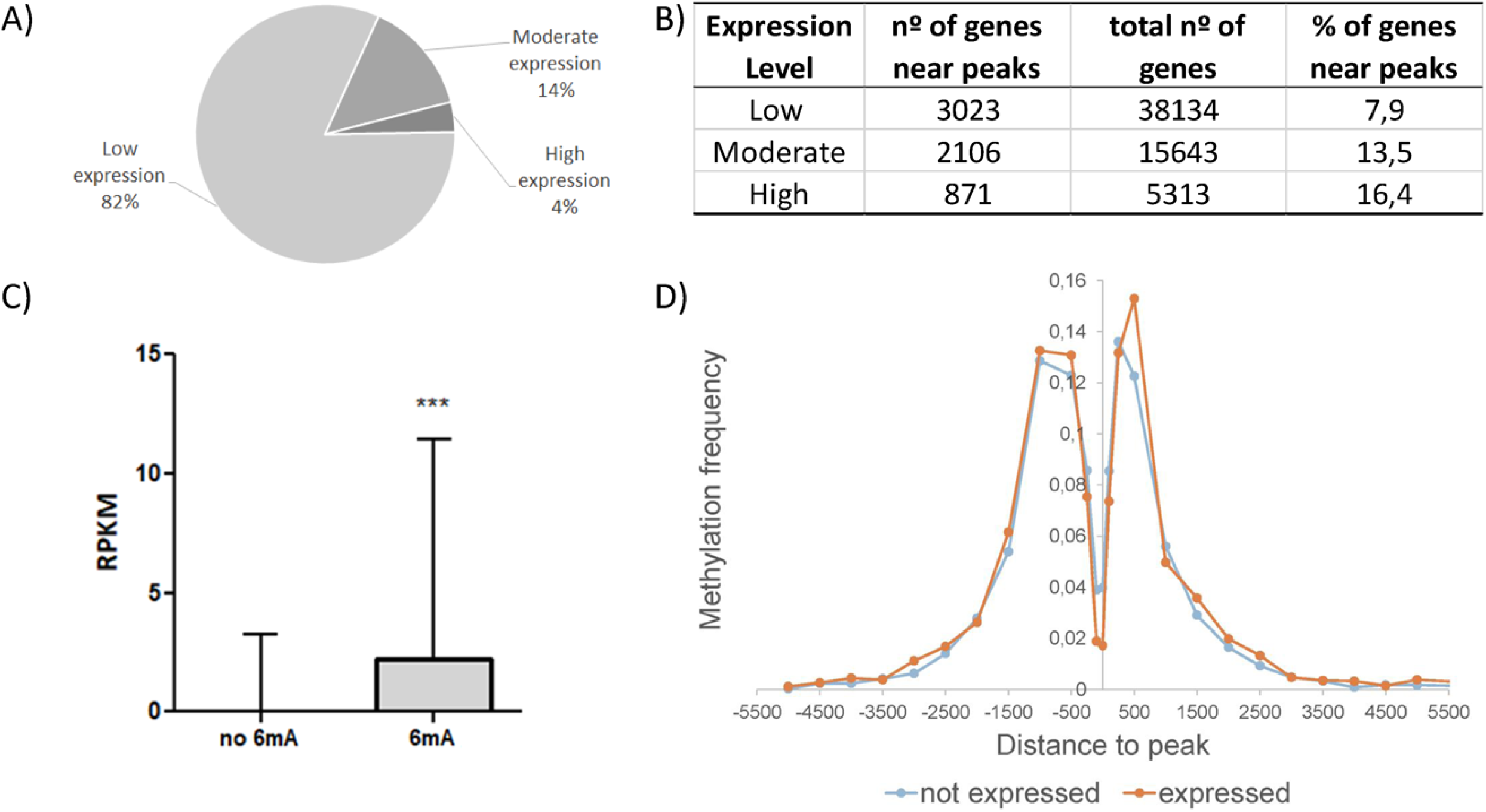
Expression levels of genes closest to 6mA DNA methylation. A) Intragenic 6mA methylation is associated with low expression. 82% of the 512 identified methylated genes are poorly expressed (RPKM =< 1), 14% moderately expressed (1 < RPKM =< 20), and 4% highly expressed (RPKM > 20). B) Percentage of genes with different expression levels near intergenic 6mA peaks. Of the 6000 genes closest to intergenic methylation, 3023 have low expression, 2106 have moderate expression and 871 genes have high expression, corresponding to 7.9%, 13.5% and 16.4% of all *T. vaginalis* poorly, moderately and highly expressed genes respectively. C) Bar plot comparing the median and interquartile range of RPKM expression of genes near intergenic 6mA and genes in non-methylated regions shows genes near intergenic methylation possess a significantly higher expression than genes in non-methylated regions. The p-value was calculated by a two-sided Mann-Whitney test (P<0.0001). D) Distribution of methylation frequency show intergenic 6mA is primarily located between −1500pb and +1000bp of the nearest not expressed (RPKM =< 1) or expressed (RPKM > 1) genes. Methylation frequency of intergenic peaks was calculated and plotted against the distance to the nearest gene, with 0 corresponding to peaks that fall within genes.

As described previously, most of the MeDIP-seq peaks have an intergenic localization (Fig. 3A). In order to evaluate if 6mA could be affecting the expression of adjacent genes as has been described for 6mA in mouse embryonic stem cells (30), we analyzed the expression level of the genes closest to intergenic methylation (Fig. 4B). We obtained a total of 6000 genes found near 6714 peaks. However, due to the fragmented nature of *T. vaginalis* reference genome not all the intergenic peaks could be considered, and it is likely that the real number of genes near intergenic 6mA could be higher. Surprisingly, in contrast with our observations regarding the expression levels of methylated genes (Fig. 4A), we found a higher percentage of expressed genes (13.5% with moderate and 16.4% with high expression) near intergenic methylation compared to poorly expressed genes (Fig. 4B). In fact, genes with intergenic 6mA peaks nearby were expressed at significantly higher levels than genes in unmethylated regions (Fig. 4C). Finally, we examined the association between the 6mA peaks distance and expression level of associated genes. We observed that intergenic 6mA is primarily located between −1500bp and +1000bp of the nearest genes regardless of their expression level (Fig. 4D). Intriguingly, non expressed genes are found significantly closer to their nearest upstream or downstream intergenic peak than expressed genes (Sup. Fig. 3B-C). These observations suggest that the effect of 6mA on gene expression in *T. vaginalis* depends on its relative position to the corresponding gene.

### 6mA delimits chromatin loops

During the analysis of the expression levels of genes near intergenic methylation regions, we noticed various instances of intergenic 6mA peaks flanking sections of the genome where all the genes contained within had similar expression levels. A closer look revealed the presence of 1046 out of 2937 such intervals with more than one gene: 452 containing genes that were not expressed (repressive intervals) and 594 containing expressed genes (active intervals) (Fig. 5A). These intervals varied in the number of genes they contained (from 2 to up to 36 genes) (Fig. 5B) as well as in length, with some spanning over 20kb (Sup. Table 7). As can be observed in Figure 5B, intervals containing a greater number of genes tend to be silenced. Conversely, most of the intervals containing fewer genes are actively transcribed.

**Figure 5:**
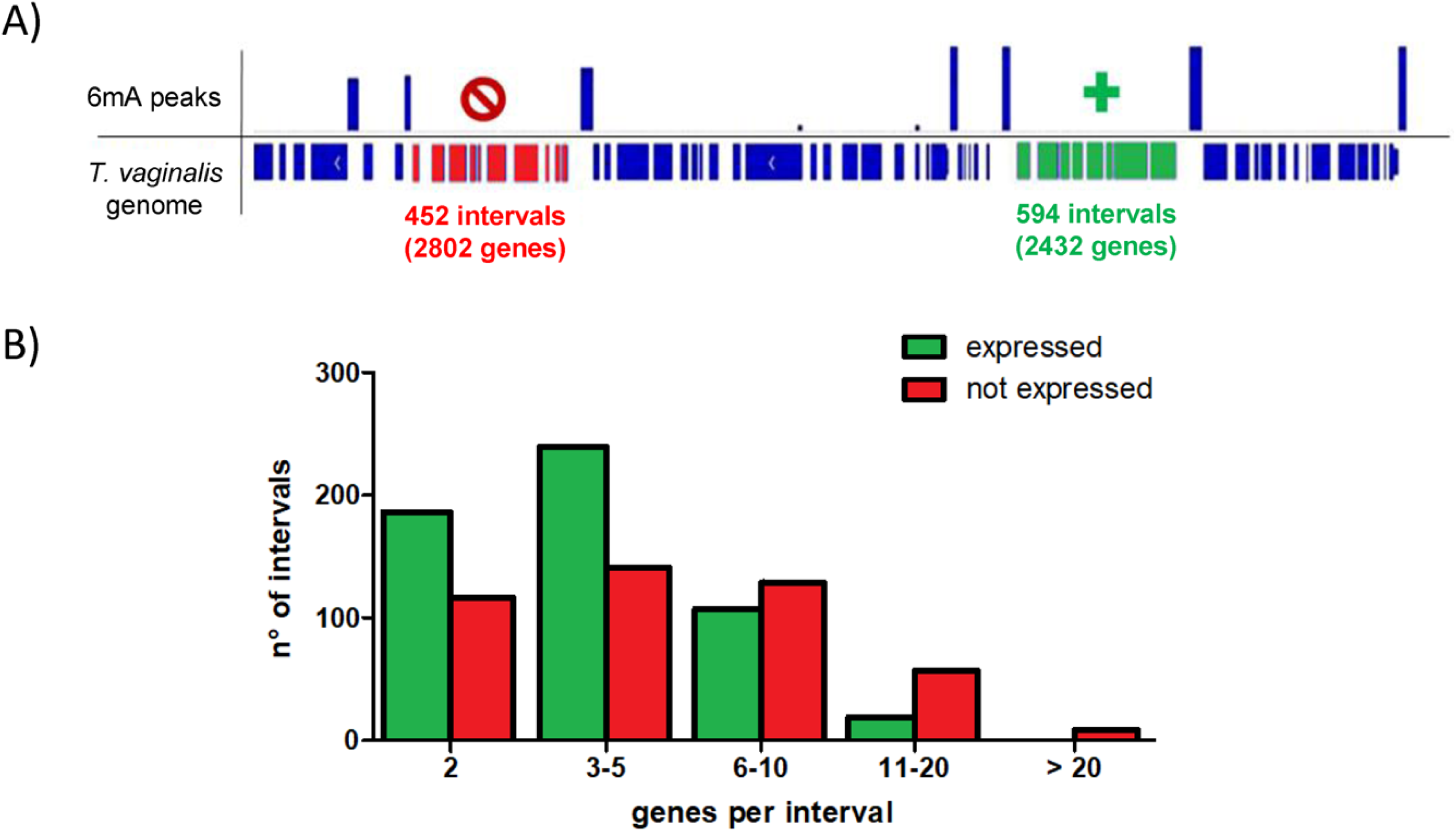
Gene blocks flanked by intergenic 6mA. A) Representation of intervals delimited by 6mA peaks comprised of genes with the similar level of expression. 1046 intervals were found, 452 containing not expressed genes (red) and 592 containing expressed genes (green). B) Quantification of intervals flanked by 6mA containing different numbers of expressed (green) and non-expressed (red) genes.

In other organisms it is well known that the three-dimensional genome organization within the nucleus is an important element in the regulation of gene expression (42). In particular, chromatin looping is a type of intrachromosomal interaction that has been shown to influence gene expression (43). Recent work in mammalian cells demonstrated that 5mC was capable of regulating the expression of two genes 10 kbp apart via chromatin loop formation (44). Based on these antecedents and our bioinformatics observations, we performed a chromatin conformation capture (3C) assay (Fig. 6A) to determine if the observed intervals, delimited by intergenic 6mA, could be forming this type of chromatin architecture. In order to select intervals to analyze by 3C assay, we filtered out all the intervals with fewer than 5 genes, obtaining a list of 187 active and 233 repressive intervals. Surprisingly, we found several intervals containing BspA family members: 2 of the repressive intervals contained 4 BspA genes each, while 6 active intervals contained one BspA gene per interval (Sup. Table 8). In bacteria, the BspA family of cell surface proteins has been shown to be involved in the colonization of the oral mucosa and triggering of host immune response and work on *T. vaginalis* BspAs suggest these proteins might have a similar function in the parasite (45). Therefore, we decided to focus primarily on those active and repressive intervals that contained BspA genes. Unfortunately, the repetitive nature of *T. vaginalis* genome presented a challenge for the design of specific 3C PCR primers, limiting the amount of BspA containing intervals to be considered for the experiment to repressive intervals p5512-5513 and p2296-2297, and active intervals p4130-4131, p560-561 and p271-272. Consequently, we decided to include intervals lacking BspA genes, resulting in four additional active and repressive intervals to be analyzed by 3C assay (Sup. Table 9). Amongst this set of regions, we were unable to amplify only five intervals: BspA-containing intervals p560-561 and p271-272, and intervals p1065-1066 and p228-230. On the other hand, PCR amplification of 3C product using primer pairs for active intervals p456-457, p690-692, p806-807 and p4130-4131 as well as repressive intervals p657-658, p786-787, p1871-1873 and p55125-513 showed bands of the expected size for each interval, suggesting that these separate regions of DNA (Sup. Table 9) were physically linked to each other in the cell. Importantly, these amplification products were either absent or less intense when non-crosslinked control was used as template and completely absent when undigested gDNA control was tested (1-2, Fig. 6B). Additionally, the intensity of these bands was higher than the ones obtained using an alternative set of primers meant to interrogate the interaction between one extreme of each interval and a flanking region located at a greater distance (1-3, Fig. 6B). Furthermore, sequencing of each 3C band verified that the fragments belonged to the expected cross-ligation products (Sup. Fig. 4). Taken together, our results suggest the presence of chromatin looping at these regions. Additionally, we amplified BspA-containing interval p2296-2297 (Sup. Fig. 4I), obtaining a band of the expected size. Although we were not able to achieve reproducible results with this primer pair, band sequencing confirmed that the identity of the fragment was the expected cross-ligation product (Sup. Fig. 4J), suggesting the formation of a chromatin loop.

**Figure 6:**
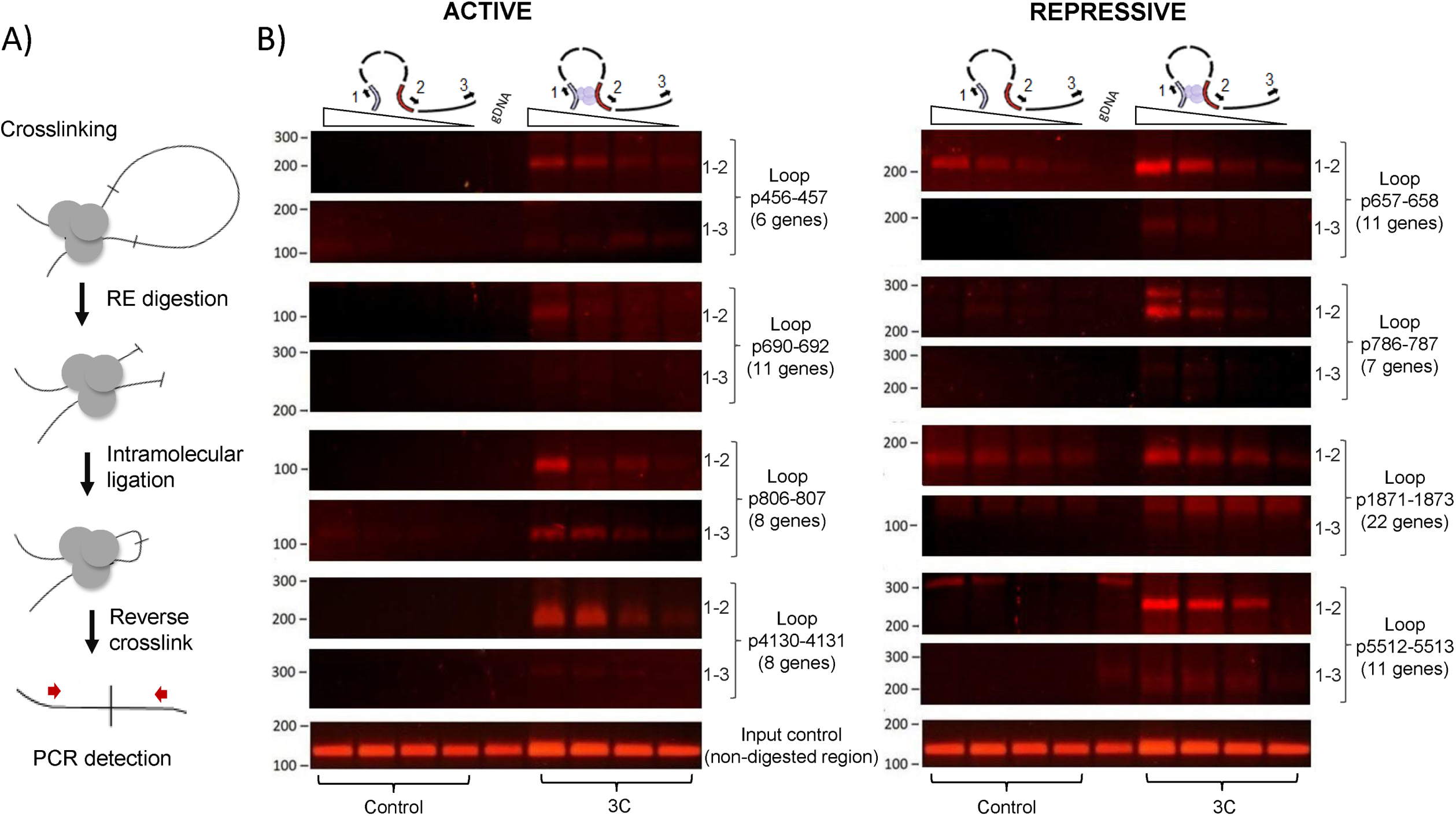
Chromatin conformation capture assay evidence loop formation in *T. vaginalis* genome. A) Schematic representation of the chromatin conformation capture assay (3C). B) 3C assay shows evidence of loop formation in *T. vaginalis* genome. Lanes 1-4: serial dilutions of uncrosslinked control template. Lanes 5: undigested gDNA. Lanes 6-9: serial dilutions of 3C template. Active: amplification of a regions containing a block of transcriptionally-active genes. Repressive: amplification of regions containing a block of transcriptionally-active genes. Non-digested control: amplification of a region without a restriction enzyme site in between the utilized primers.

To examine whether 6mA might be responsible for regulating the formation of these structures, we performed a 3C assay of parasites treated with ascorbic acid (AA), which has been previously shown to favor DNA demethylation in mammalian cells (46). Dot blot assay of parasites treated with AA confirmed a diminution of 6mA mark of parasites treated with 100 mM AA compared to non-treated parasites (Fig. 7A). Excitingly, loop formation as determined by 3C analysis was diminished in parasites treated with AA when compared to the non-treated control (Fig. 7B). Consistent with this finding, our observations suggest that 6mA could indeed be a mark associated with chromatin looping and may regulate the transcriptional activity of gene-associated blocks in *T. vaginalis*.

**Figure 7:**
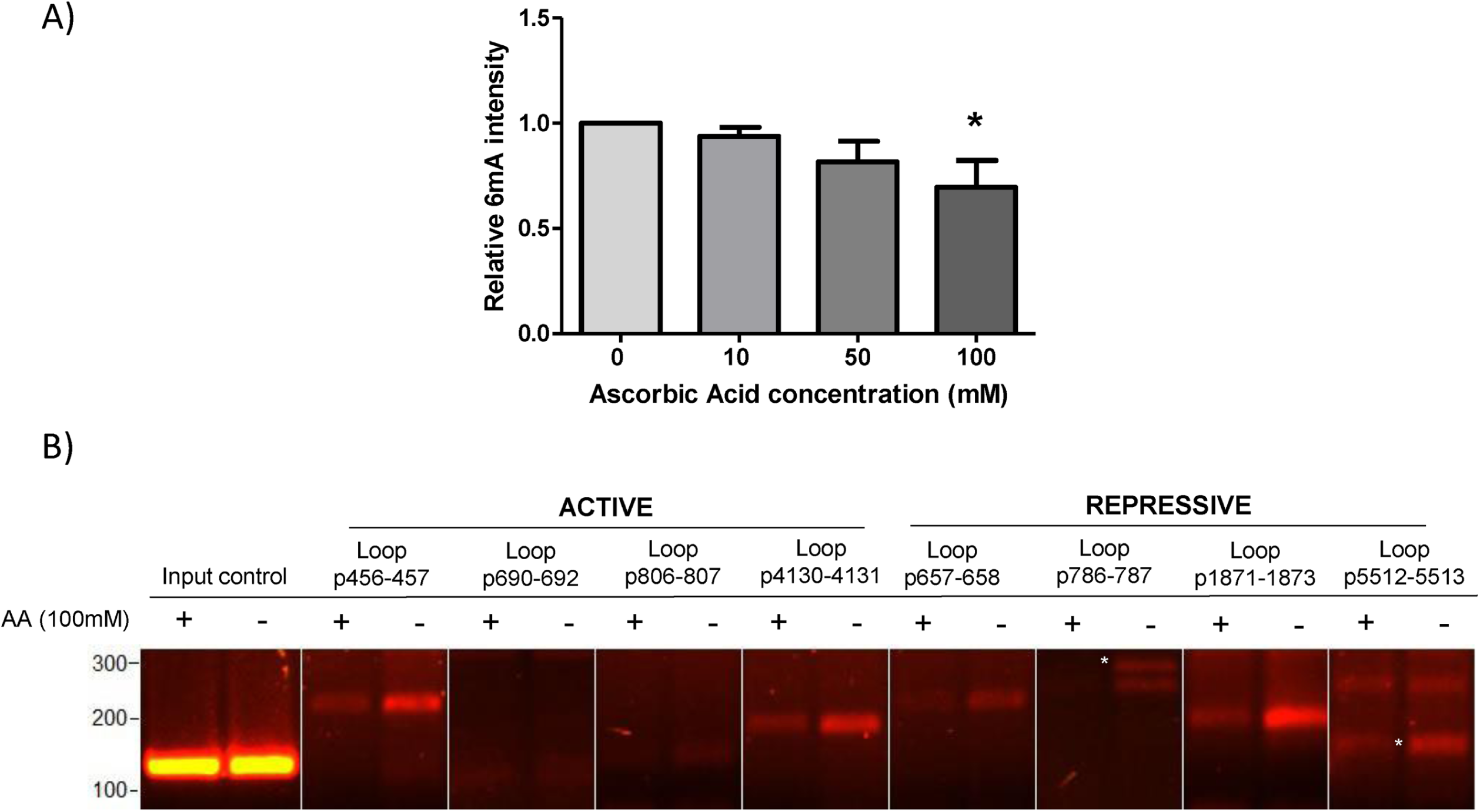
3C assay on parasites treated with ascorbic acid. A) Quantification of 6mA levels of parasites treated with different concentrations of ascorbic acid (AA) shows diminution of 6mA abundance in parasites treated with 100 mM AA compared to untreated parasites. Each column represents the mean and SD of two independent experiments. D) 3C assay of parasites exposed to 100mM of AA (+) shows diminished loop formation compared to non-treated parasites (-). 50 ng of 3C template was used for PCR amplification using primers for each active and repressive loop. Non-digested control: amplification of a region without a restriction enzyme site in between the utilized primers.

## DISCUSSION

Epigenetics studies and its relationship with gene expression in *T. vaginalis* are still in their infancy. To date, only the contribution of histone modifications to the regulation of gene expression in the parasite has been described (18, 19). In the present study, we describe for the first time the presence of 6mA modification in the genome of this unicellular parasite. Here, we present a whole genome analysis of 6mA distribution that provides insight into the possible contributions of this modification to 3D genome architecture and gene expression.

In recent years, the role of 6mA has been expanded from a primarily prokaryotic DNA modification to an important epigenetic mark present in several multicellular and unicellular eukaryotic organisms (34). In this study, we use antibody-dependent and independent methods to demonstrate that 6mA is an abundant DNA modification in *T. vaginalis* genome. 6mA levels in the parasite are comparable to those found in early-diverging fungi (33), making *T. vaginalis* one of the eukaryotic organisms with the highest levels of 6mA reported to date. It is interesting to note that just like early-diverging fungi (33), 5mC in *T. vaginalis* was nearly undetectable. Unfortunately, detection methods that rely on the use of antibodies can give false positives when 5mC levels are low (47), which could explain the low-quality reads obtained from the 5mC MeDIP-seq experiment. These results support our conclusion that 6mA, and not 5mC, is the main DNA modification in *T. vaginalis* genomic DNA.

To be considered an epigenetic mark, 6mA needs to be maintained after DNA replication. In *Tetrahymena*, maintenance methylation occurs quickly after DNA replication (48). Similarly, 6mA in *Chlamydomonas* is installed shortly after DNA replication and is stably maintained during cell proliferation (29). Although the precise moment of 6mA deposition in the newly synthetized DNA strand in the parasite remains elusive, our observations indicate maintenance of 6mA methylation occurs after cell division and supports the presence of active 6mA methyltransferase(s) in the parasite. It should be noted that although phylogenetic analysis revealed the presence of a group of parasite specific 6mA DNA methyltransferases encoded in the genome database (TrichDB) (49), functional analyses had not been performed to validate these candidates yet. On the other hand, while DNA 6mA demethylases have been identified in several organisms (27, 28, 30, 36), no classical TET or AlkB homologs were identified in *T. vaginalis* genome (49). This could be indicative of the presence of non-conserved enzyme, a lack of active 6mA demethylation or the use of an alternative pathway, such as the base excision-repair DNA demethylation pathway found in plants (50). It will be of interest to take a closer look at the DNA glycosylases found in *T. vaginalis* genome database and evaluate if they could be involved in active 6mA DNA demethylation. Identifying the proteins responsible for the regulation of 6mA levels in the parasite will further our understanding of the mechanisms and functions of this mark.

Genome-wide distribution of 6mA varies between species (27–30, 51). An evolutionary conservation for 6mA in unicellular organisms has been proposed due to the similarities in 6mA distribution pattern and function between green algae and *Tetrahymena* (48). However, using a MeDIP-seq approach we found the distribution pattern in *T. vaginalis* resembles more that of 6mA in *Drosophila* and mouse than other unicellular eukaryotes. Methylation peaks in *T. vaginalis* are enriched in intergenic regions and seem to have a preference for certain types of DNA TE superfamilies, reminiscent of the preference for young LINE-1 transposons described in mouse (30). In Drosophila, some 6mA peaks are enriched in the gene bodies of transposons (27). Similarly, we found that the majority of the methylated genes in the parasite belong to transposable elements, with 6mA peaks preferentially located on the CDS or TTS of these TE genes (Sup. Table 10). This predilection for certain types of TEs and the specific localization of peaks in transposon genes raises the question of whether 6mA could act as a regulator of specific TEs in *T. vaginalis*.

Like its genomic distribution, 6mA association with transcription is dependent of the organism in question. It has been demonstrated that 6mA acts as a mark of active transcription in *Arabidopsis* (35), *Drosophila* (27), green algae (29) and fungi (33), while it is a mark of transcriptional repression in mouse (30) and rice (36). In *T. vaginalis*, 6mA seems to be associated with silencing when found on genes. Importantly, we cannot rule out the possibility of 6mA acting in conjunction with other components of the epigenetic code to regulate gene expression in a context-dependent manner. In future studies, it will be interesting to determine whether 6mA binding proteins exist in *T. vaginalis* which could help regulate the response of chromatin to this modification. An integrative analysis is necessary to fully understand how epigenetics shapes the transcriptional landscape of *T. vaginalis*.

Chromatin looping is a common intrachromosomal protein-mediated interaction capable of influencing gene expression. The function of these interactions is determined by the nature of the sequences brought together and the proteins that interact with them (52). For example, promoter-enhancer loops are strongly associated with gene activation (53) and it has been proposed that the presence of chromatin loops may contribute to the co-regulation of specific block of genes by increasing the local concentration of the transcription machinery, resulting in the formation of structures similar to transcription factories (54). Alternatively, chromatin looping has also been implicated in transcriptional repression in *Drosophila* (55) and mammals (56). Interestingly, our bioinformatics analysis revealed the presence of active and repressive intervals of varying length flanked by 6mA. A chromatin conformation capture assay confirmed that the methylated extremes in both transcriptionally active and repressive blocks of genes are in close spatial proximity. Notably, some contained one or more members of the BspA-like gene family of surface proteins which might be involved in parasite pathogenesis (57). In *P. falciparum*, chromosomes containing virulence genes of the *var* family form loops that permit the perinuclear colocalization of the var genes (58), and a recent analysis demonstrated that this clustering of virulent genes is a common feature among other *Plasmodium* species (59). Since clustered organization of virulence genes allows coordination of gene expression in *Plasmodium* (59), future studies are warranted to evaluate if the same is true for *T. vaginalis*.

The association of 6mA with chromatin looping is noteworthy because it points to a novel function for this epigenetic mark in eukaryotes with scarce 5mC levels. Given that this work demonstrated that chromatin structure may regulate, positively or negatively, specific blocks of genes; these finding open new avenues for understanding the role of 6mA writers, erasers and readers that may play an important role on the transcriptional regulation of *T. vaginalis*.

## MATERIALS AND METHODS

### Parasites, cell cultures and media

*Trichomonas vaginalis* strains B7268 (60) and G3 (ATCC PRA-98, Kent, UK) were cultured at 37°C in TYM medium supplemented with 10% horse serum, penicillin and streptomycin (Invitrogen) (61). To obtain *M. hominis* free strains, parasite cultures were treated for two weeks with the recommended working concentration of BM-Cyclin (Roche). *M. hominis* clearance was confirmed by PCR as described previously (62).

### 6mA and 5mC immunolocalization experiments

Parasites in the absence of host cells were incubated at 37°C on glass coverslips as previously described (63) for 4 hours, then fixed and followed by permeabilization in cold methanol for 10 min. Cells were washed with PBS and incubated at 37°C with a 1:100 dilution of RNase A (QIAGEN) in PBS for 3 hours. Non-RNase A treated control parasites were incubated for 3 hours in PBS. Parasites were washed once again, blocked with 5% FBS in PBS for 30 min and incubated overnight with a 1:100 dilution of anti-6mA or anti-5mC (Abcam, USA) primary antibody in PBS containing 2% FBS. Parasites were washed and then incubated with a 1:5000 dilution of Alexa Fluor conjugated secondary antibody (Molecular Probes). Nuclei was stained with DAPI (2 mg/ml) and coverslips were mounted onto microscope slips using Fluoromount Aqueous Mounting Medium (Sigma-Aldrich). All observations were performed on a Zeiss Axio Observer 7 epifluorescence microscope. Adobe Photoshop (Adobe Systems) was used for image processing.

### Genomic DNA extraction

*T. vaginalis* cultures were lysed using a solution of 8 M urea, 2% sarkosyl, 0.15 M NaCl, 0.001 M EDTA, and 0.1 M Tris HCL pH7.5 (64). RNase A was added to a final concentration of 0.1 mg/ml and lysates were incubated at 37°C for 2-3 hours. Total DNA was extracted using phenol/chloroform/isoamyl alcohol (Sigma-Aldrich), precipitated with isopropanol, and resuspended in nuclease-free water.

### Dot Blot Assays

RNA free genomic DNA samples were diluted to the desired concentration and 1 µl of each sample was spotted on Amersham Hybond-N+ membranes (GE Healthcare). For anti-5mC blots, DNA was denatured for 10 minutes at 95°C. Membranes were allowed to air dry and then DNA was crosslinked for 3 minutes twice in a Hoefer UVC 500 Crosslinker (Amersham Biosciences). Membranes were stained with 0.02% methylene blue in 0.3 M NaOAc pH 5.2 solution for DNA loading control and then washed with distilled water. The membranes were blocked for 1 hour in 5% milk TBS-T and incubated overnight at 4°C with 1:1000 anti-6mA primary antibody dilution or 1:500 anti-5mC primary antibody dilution in 5% milk TBS-T. Membranes were washed three times for 10 min with TBS-T and incubated with a 1:10000 dilution of AP-conjugated secondary antibody in 5% milk TBS-T for 1 hour at room temperature. After three washes of 10 min with TBS-T, blots were developed using NBT and BCIP (Roche).

### Restriction enzyme digestion assay

Restriction enzyme digestion was performed by treating 1 µg of genomic DNA with 1 µl of 10 U/µl MboI or DpnI restriction enzyme (Thermo Scientific) at 25°C overnight. Sample with no restriction enzyme was used as control. Enzyme was heat inactivated and the product mixture was separated by agarose gel electrophoresis. Gels were visualized with a UV transilluminator.

### Quantification of 6mA by UHPLC-QQQ-MS/MS Analysis

6mA abundance was quantified as previously described (28, 51). 1.5 µg of genomic DNA in 30 ml ddH2O was digested to free nucleosides using 5 U of DNA Degradase Plus (Zymo Research) in 25 μl reactions incubated for 2 hours at 37°C. The diluted solution was filtered through 0.22 mm filter and 10 ml solution was injected into LC-MS/MS. The nucleosides were separated by reverse phase ultra-high performance liquid chromatography (UHPLC) on a C18 column (Agilent), with online mass spectrometry detection using Agilent 6460 QQQ triple-quadrupole tandem mass spectrometer (MS/MS) set to multiple reaction monitoring (MRM) in positive electrospray ionization mode. Nucleosides were quantified using the nucleoside precursor ion to base ion mass transitions of 266.1-150.0 for 6mA and 252.1-136.0 for A. Quantification of the ratio 6mA/A was performed using the calibration curves obtained from nucleoside standards running at the same time. Measurement of 6mA and dA levels in the DNA degradase enzyme mix alone served as an additional control for background levels of nucleic acid contributed by the digestion enzymes. The low levels of 6mA and dA in the DNA degradase enzyme mix were subtracted from the final measurements. Because *T. vaginalis* have relatively high levels of 6mA, the low levels of methylated and unmethylated DNA in the reaction mix were a negligible fraction of the total digested DNA samples.

### Cell cycle synchronization

Parasites (5 x 105 cells/ml) were incubated overnight at 37°C in the presence of 100 µM hydroxyurea (HU). The following day cells were resuspended in fresh medium without HU, divided into 8 ml aliquots and left to grow at 37°C. An aliquot was taken every two hours and was used for flow cytometry analysis to determine cell cycle stage. The remaining sample was used for genomic DNA extraction to determine abundance of 6mA.

### Flow cytometry

Parasites were treated as described previously (65). Parasites were fixed in 5 ml of ice-cold 100% EtOH, and incubated at 4°C overnight. Each sample was washed once in 1 ml PBS containing 2% v/v bovine serum (BS) and resuspended in 1 ml PBS containing 180 μg/ml RNase A and 2% v/v BS. After 30 min incubation at 37°C samples were stained with 25 μg/ml propidium iodide (PI) solution and incubated an additional 30 min at 37°C. Samples were analyzed using a fluorescence-activated cell sorter (BD FACSCalibur; BD Biosciences, San José, CA, USA) with appropriate filter sets. Data was analyzed using FlowJo 7.6 software.

### MeDIP-seq

20 μg of purified genomic DNA from *T. vaginalis* strain B7268 (60) was diluted in 400 μl 1X IP buffer (10 mM Na-Phosphate pH 7, 140 mM NaCl, 0.05% Triton X-100) and fragmented to 100-500 bp using a water-bath sonicator (Amp: 50%, Cycle: 30 sec ON and 30 sec OFF, Time: 20 min). Sample was divided into aliquots containing 6-7 μg of sonicated DNA and heat-denatured. 1 μg was saved and stored at −20°C to use as input control, the rest was immunoprecipitated overnight at 4°C using 5 μg of anti-6mA antibody (Abcam) bound to protein A magnetic beads (Invitrogen). After three washes with 1X IP buffer, beads containing bound methylated DNA were resuspended in 250 μl digestion buffer (50 mM Tris pH 8, 10 mM EDTA, 0.5% SDS) with 100 μg Proteinase K (Roche) and incubated at 65°C for 4 h. Eluted DNA and input control samples were purified using phenol/choloroform/isoamyl alcohol extraction followed by EtOH precipitation. MeDIP samples were sent to Applied Biological Materials Inc. for library preparation and paired-end sequencing. Two independent high-throughput sequencing experiments, each containing pooled DNA from three independent immunoprecipitation, were performed.

### Bioinformatics analysis of MeDIP-seq data

After the adaptors were trimmed, alignment of high-quality fastq reads to the reference genome (GCF 000002825.2 ASM282v1) was performed using Bowtie2 (66) with –qc-filter parameter and all other parameters at default settings. The BAM files were used as input for MACS2(67), which was run with the –g parameter set at 1.76e8 and all other parameters at default settings. Input reads were used as control sample for peak calling.

Peaks were then filtered by q-value (< 0.01) and fold enrichment (> 2). Only peaks with more than 75% overlap between replicates were considered for further analysis. HOMER2 (68) was used for peak annotation. For repetitive element annotation, RepeatMasker v4.0. (69) was used with a custom library of *T. vaginalis* specific RE taken from Repbase (70). GO enrichment analysis was performed using the Results Analysis feature at TrichDB with default settings.

### Chromatin conformation capture assay

Crosslinking of DNA was achieved by incubation of 50 ml (~1 x 106 cells/ml) of *T. vaginalis* strain B7268 (60) culture with 1% formaldehyde for 15 min at room temperature. Crosslinking was stopped by the addition of 125 mM glycine, followed by 5 min incubation at room temperature and then on ice for 15 min. Cells were harvested, washed in 10 ml PBS/PI and resuspended in 1.4 ml of cold lysis buffer (10 mM Tris-HCl pH 8, 10 mM NaCl, 0.2% NP-40, 1X PI) followed by homogenization with a dounce homogenizer pestle A. After centrifugation, the pellet was washed once with 1.25X Buffer H (Promega) and resuspended in 500 µl of 1.25X Buffer H (Promega). A total of 7.5 μl of SDS 20% was added and incubated for 40 min at 65°C and then for 20 min at 37°C. SDS was quenched with 50 μl of 20% Triton x-100 at 37°C for 1 h. 400U of EcoRI (Promega) was added and incubated overnight at 37°C followed by enzyme heat inactivation with 1% SDS at 65 °C for 20 min. Approximately 30 μl of digested sample was set aside to test restriction enzyme efficiency by electrophoresis and PCR. Digestion product was cooled and transferred to a pre-chilled 15 ml tube containing 4 ml 1.1X Ligase Buffer (Promega) and 1% Triton X-100. After 1 h incubation at 37°C the sample was cooled down and mixed with 30 Weiss Units of T4 DNA Ligase (Promega) and incubated for 4 h in a water bath at 16°C. Crosslinking was reversed overnight at 65 °C in the presence of 300 μg of Proteinase K (Roche). DNA was purified using phenol/choloroform/isoamyl alcohol extraction followed by EtOH precipitation. The sequences of primers used in 3C assays are shown in supplementary table 11. Two-fold serial dilutions starting with 100 ng/µl were used for PCR amplification. 3C PCR fragments were purified and sequenced to verify the identity of the cross-ligation product.

## ACCESSION NUMBERS

All sequencing data that support the findings of this study have been deposited in the NCBI Sequence Read Archive (SRA) and are accessible through the SRA accession number PRJNA526331. All other relevant data are available from the corresponding author on request.

## ACKNOWLEDGEMENTS

We thank Dr. Pablo Manavella and our colleagues in the lab for helpful discussions.

## FUNDING

This research was supported with a grant from the ANPCyT grant BID PICT 2015-2118 and PICT-2018-01892 (NdM). Work in the Greer lab was supported by NIH grants DP2AG055947 and R21HG010066 (ELG) and work in the Johnson lab was supported by R01AI30537. NdM and PHS are researchers from the National Council of Research (CONICET) and UNSAM. AL is a PhD fellow from CONICET. The funders had no role in study design, data collection and analysis, decision to publish, or preparation of the manuscript.

## CONFLICT OF INTEREST

The authors declare no competing interests.

