## Supplemental Information for "Adenine DNA methylation, 3D genome organization and gene expression in the parasite *Trichomonas vaginalis*"

### Supplementary Figure 1

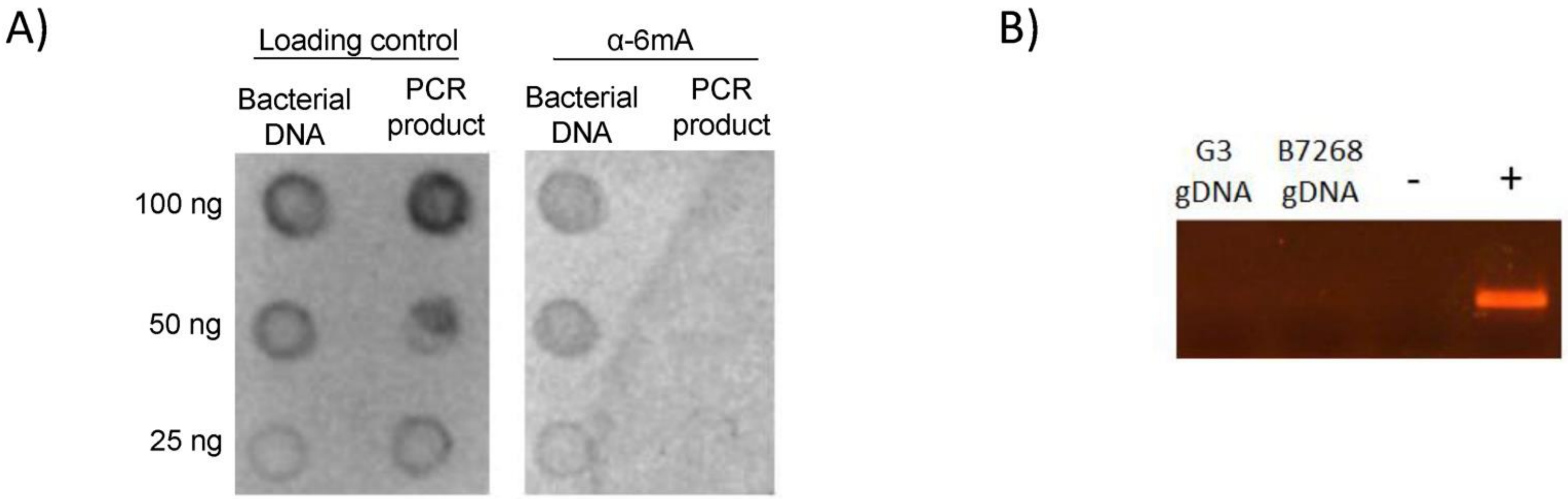


**Supplementary Figure. 1: A) Specificity of α-6mA antibody.** Dot blot assay against 6mA using bacterial DNA as positive control and PCR product as negative control shows the antibody specifically recognizes methylated adenines. Left: gDNA dots stained with methylene blue for loading control. Right: 6mA signal. **B) *T. vaginalis* cultures free of *M. hominis* contamination.** PCR using *M. hominis* specific primers. Lane 1: gDNA of parasites from strain G3 subjected to BM-cyclin treatment. Lane 2: gDNA of parasites from strain B7268 subjected to BM-cyclin treatment. Lane 3: negative control. Lane 4: gDNA of non-treated parasites as positive control of *M. hominis* contamination.

### Supplementary Figure 2

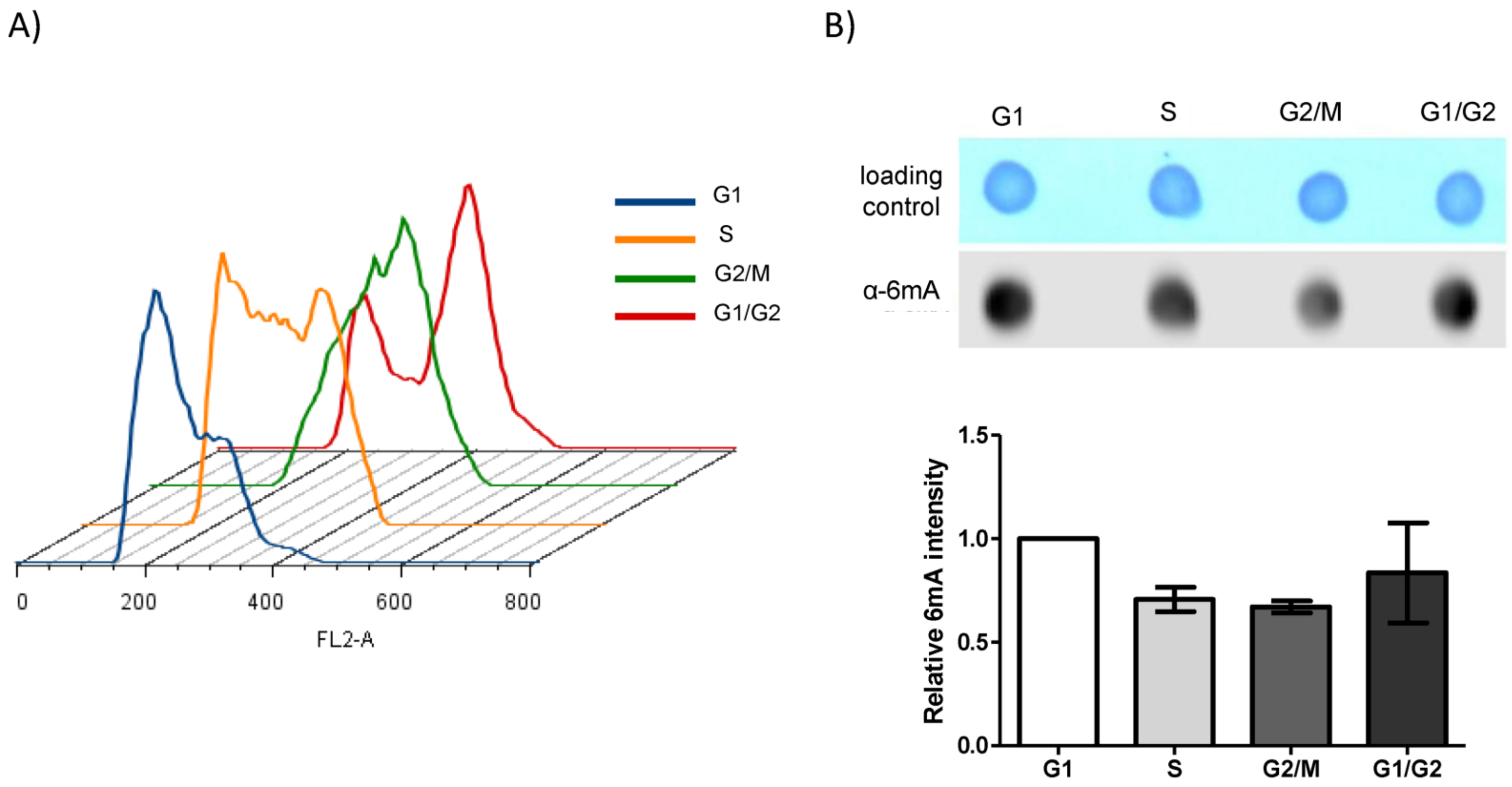


**Supplementary Figure 2: 6mA methylation levels during *T. vaginalis* cell cycle progression.** Samples from a hydroxyurea synchronized culture were taken every two hours post-treatment. Part of the sample was used for cell cycle stage determination by flow cytometry (A), the rest was used for gDNA extraction to measure 6mA levels using a dot blot assay (B). Representative image of two independent experiments is shown. C) Quantification of dot blot signal.

### Supplementary Figure 3

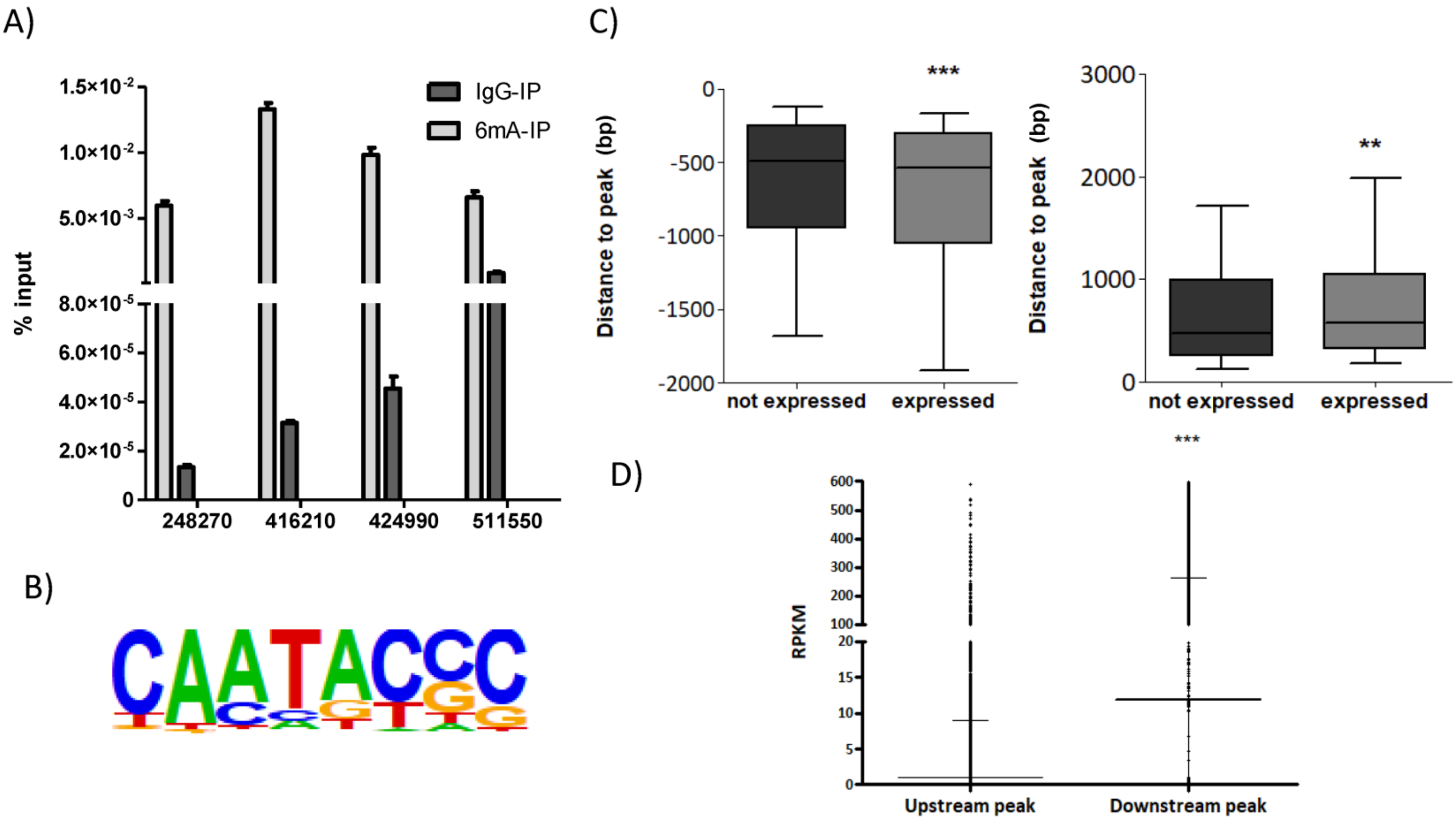


**Supplementary Figure 3: MeDIP-seq analysis.** A) Most prevalent methylation motif identified by HOMER2, found in 4692 peaks (p-value = 1e-1642). B) MeDIP-qPCR of four genes selected from MeDIP-seq demonstrated enrichment in 6mA immunoprecipitated sample (6mA-IP) compared to sample immunoprecipitated with IgG control antibody (Ig-IP). C). Boxplot comparing the median and interquartile range of gene distance to intergenic peaks found at the 5' of expressed and not expressed genes. The p-value was calculated by a two-sided Mann-Whitney test ($P<0.0001$). D) Boxplot comparing the median and interquartile range of gene distance to intergenic peaks found at the 3' of expressed and not expressed genes. The p-value was calculated by a two-tailed unpaired Student's t test ($P < 0.05$).

### Supplementary Figure 4

A)

5′ -CAGCAGCGTCATCAAGAGTCATAGCTGTCTCTCTAAGTTCGTAAA
GTTTGTTGTCTTTAAGGAGCATCAACATTTCGTAAAGAATTTCAATGAT
TTCGTCGTCTTTTTGTGTCTTGTCTGCCAACTTCTTGATTGAAGAAACA
ATCTTCTTAGCGAATTCAGTTTAGCTTCATCGCTGATTTCCGAAGCCCA
GGATGTATTTGCAGCC

B)

5′ -AGCAAAACGAAACCCTTACACAATCATATATGTAGAATTCAACAGT
ATAATGTGACAGGAAAAGAGTTAGTGCATTTTCGCCTTCTTTTGCTTCGA
TACACAAAACGCA

C)

5′ -TCCCTTAGTTATCCCAATCTGAAGATGACGAGGAGTCTGTTTCAT
CATCAGAAGATTGAGCCTTAACTTGGAATTCAAAGAGAACATATGATAT
GCTATTTCCGTATGTGAAGTGCT

D)

5′ -TTGTTACCCAATCCGAGGAGTTTGATGATTGTAGTAGCAATACTT
GCTGTAAATCAATCGCCTTTCGTTGACTTCTGTCCATCAACAAATAATT
TTCTTCTTCTTTCAAGAGGAATTTCATAAGAATTCATCTCTTAAATGCT
GCAAAAAATATTGAAATGCATAACCACGCCATAAGGAAATCC

E)

5′ -TCATAATCAAAGACCTCATAAACGAGAAGAAGAAATTGAAAATAAT
GAAATGGAAATCATTGATGACATTAAAGAGGATGGAATTCCAGCAGATTT
GGCCAAAAAGATCCTACAAAATACGTTAAATCCTCTCAAAGATGATGAAG
TTCATGAAATATTCCAGAAGTTAACTGTAGACGAAGACGGAACAGTACTG
TATAGCGAGCTCG

F)

5′ -TGTATGAGCTGAGTGTAAATGTCTTACAATATTGAATATTTTTAAA
TCTTGATTTATTCAAAAACGTAAGAATAAAAACCGTAAGAATCATACATT
TTTTACTTCATATGTTAAAAAAAATGATACAAAATCGTTCAATATTGTAT
GAATTCAAAATACAAACTTAAAGATTGTATACGTTTAAAATTATTCAGAT
TGTGTTTGATTTCCACTCATTCTGTTTCTTTTGAACTCTAACACTAACAT
TTCATGATGATTACATGATCCATTTGTACATGGTTGAATTGTTCC

G)

5′ -ACCACCCTCACACGTATGAAAACATAGCACAAAAATAATGTTTAT
GTTAGAATGTTAGATCTAACACGTTTTCTAACCTCTTTTCTAACCATGT
TGAATTCTTATTTGATTTTTCATCATAACATGATGATGTTATTCAGATT
TGCATCAATAGCAATCAGATTACC

H)

5′ -CTGATTTCTTTTACTCTCTCGTATATACTAGATTTAAAGAATTCA
CTCACAATCAGTATACTCAATTTTTTGTGCAAAATCTTTTTTATTCTTA
AAATATACTGAATAAATTTACTGCAAAATATGATTTTGTCATGAAACAA
AGAAATTAAATTTTATATATTTTCATTAGTTTTATTTATTTTCCTTGTG
GAGAAAATTTATTAAGAGAATGCCATTCTTGTGCATGAAAATTGGATCT
TTCTGGTTGGGC

I)

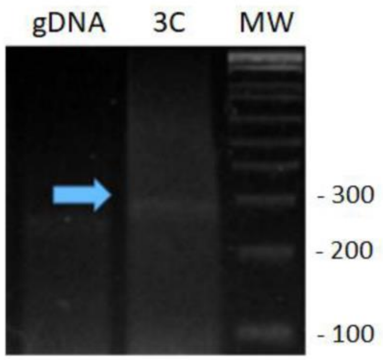


J)

5′ -CTGCGTTCGCTTATCATCTGTAACTCTTCCTCTTTATCTTAACGC
AATATTCGATTTGGCATTTGCCAACACTTCATTAAGAGAAATTAGAGTT
CCATCAGTTGCAGAAGTAATGCCTCATGCCTTTTATTACAATCAAAATC
TTATTAGGGTTATAAACACTGATGTTTTTGAACATGAATTAAGAATATA
CGCTGCTAGCGAATATAGGATTATTACAACATTAGGCAAACAGAATTCT
TTTTTAAAAATCCGAAATTTTCTCATCGTTTTGTATCGTTTCA

**Supplementary Figure 4: Sanger sequencing of 3C PCR fragments.** Amplified 3C fragments of the appropriate length were purified and sequenced to corroborate they belonged to the expected cross-ligation product. Sequencing of active loop p456-457 (A), p690-692 (B), p806-807 (C), and p4130-4131 (D). Sequencing of repressive loop p657-658 (E), p786-787 (F), p1871-1873 (G), and p5512-5513 (H). I) PCR amplification of active loop p2296-2297 shows band of the expected size. Lane 1: undigested gDNA. Lane 2: 3C template. Lane 3: MW. J) Sanger sequencing confirms the amplified fragment belongs to the expected cross-ligation product. EcoRI site is highlighted.

**Supplementary Table 1. Final list of m6A MeDIP-seq peaks**

| Contig | Start | End | Peak name |
|---|---|---|---|
| DS113177 | 47575 | 47989 | BA_peak_1 |
| DS113177 | 100199 | 103541 | BA_peak_3 |
| DS113177 | 140464 | 141180 | BA_peak_4 |
| DS113177 | 142134 | 142779 | BA_peak_5 |
| DS113177 | 161080 | 162108 | BA_peak_6 |
| DS113177 | 241069 | 241899 | BA_peak_8 |
| DS113177 | 260812 | 261777 | BA_peak_9 |
| DS113177 | 269769 | 270312 | BA_peak_10 |
| DS113177 | 274323 | 275003 | BA_peak_11 |
| DS113177 | 335191 | 335599 | BA_peak_12 |
| DS113177 | 369768 | 370319 | BA_peak_14 |
| DS113177 | 414575 | 415401 | BA_peak_15 |
| DS113177 | 456517 | 457230 | BA_peak_16 |
| DS113177 | 480528 | 480926 | BA_peak_18 |
| DS113177 | 482364 | 485673 | BA_peak_19 |
| DS113177 | 540847 | 541265 | BA_peak_21 |
| DS113178 | 2093 | 3515 | BA_peak_22 |
| DS113178 | 7115 | 7672 | BA_peak_23 |
| DS113178 | 21470 | 24803 | BA_peak_24 |
| DS113178 | 34834 | 37625 | BA_peak_25 |
| DS113178 | 77696 | 78451 | BA_peak_26 |
| DS113178 | 94412 | 100115 | BA_peak_27 |
| DS113178 | 141260 | 141946 | BA_peak_28 |
| DS113178 | 158663 | 160517 | BA_peak_29 |
| DS113178 | 185076 | 185402 | BA_peak_30 |
| DS113178 | 206876 | 208991 | BA_peak_31 |
| DS113178 | 233993 | 235339 | BA_peak_32 |
| DS113178 | 237365 | 238187 | BA_peak_33 |
| DS113178 | 239614 | 239975 | BA_peak_34 |
| DS113178 | 245660 | 247503 | BA_peak_35 |
| DS113178 | 302370 | 305320 | BA_peak_36 |
| DS113178 | 309714 | 310056 | BA_peak_38 |
| DS113178 | 362626 | 363053 | BA_peak_41 |
| DS113178 | 449376 | 449735 | BA_peak_42 |
| DS113178 | 453611 | 453963 | BA_peak_43 |
| DS113178 | 465737 | 466264 | BA_peak_45 |
| DS113178 | 498781 | 499718 | BA_peak_46 |
| DS113178 | 501250 | 503713 | BA_peak_47 |
| DS113179 | 42883 | 43985 | BA_peak_48 |
| DS113179 | 60907 | 61444 | BA_peak_49 |
| DS113179 | 73536 | 74985 | BA_peak_50 |
| DS113179 | 89200 | 90406 | BA_peak_51 |
| DS113179 | 111722 | 112418 | BA_peak_52 |
| DS113179 | 126361 | 126759 | BA_peak_53 |

| | | | |
|---|---|---|---|
| DS113179 | 157084 | 157934 | BA_peak_54 |
| DS113179 | 159495 | 159836 | BA_peak_55 |
| DS113179 | 174270 | 174801 | BA_peak_56 |
| DS113179 | 175883 | 176481 | BA_peak_57 |
| DS113179 | 183999 | 184534 | BA_peak_58 |
| DS113179 | 186575 | 186916 | BA_peak_59 |
| DS113179 | 201834 | 202163 | BA_peak_60 |
| DS113179 | 206660 | 207183 | BA_peak_61 |
| DS113179 | 207895 | 208416 | BA_peak_62 |
| DS113179 | 209635 | 210241 | BA_peak_63 |
| DS113179 | 240071 | 240761 | BA_peak_64 |
| DS113179 | 259716 | 260167 | BA_peak_65 |
| DS113179 | 260776 | 261232 | BA_peak_66 |
| DS113179 | 272100 | 272801 | BA_peak_67 |
| DS113179 | 276273 | 277251 | BA_peak_68 |
| DS113179 | 282272 | 282830 | BA_peak_69 |
| DS113179 | 288077 | 288852 | BA_peak_70 |
| DS113179 | 290337 | 291730 | BA_peak_71 |
| DS113179 | 293247 | 294301 | BA_peak_72 |
| DS113179 | 300663 | 301374 | BA_peak_73 |
| DS113179 | 301894 | 304522 | BA_peak_74 |
| DS113179 | 304832 | 305285 | BA_peak_75 |
| DS113179 | 340168 | 340521 | BA_peak_76 |
| DS113179 | 344725 | 345091 | BA_peak_77 |
| DS113179 | 345813 | 346751 | BA_peak_78 |
| DS113179 | 353674 | 355697 | BA_peak_79 |
| DS113179 | 372836 | 373501 | BA_peak_80 |
| DS113179 | 375224 | 375780 | BA_peak_81 |
| DS113179 | 377348 | 378489 | BA_peak_82 |
| DS113179 | 378857 | 379309 | BA_peak_83 |
| DS113179 | 395827 | 396132 | BA_peak_84 |
| DS113179 | 398078 | 399356 | BA_peak_85 |
| DS113179 | 418161 | 421204 | BA_peak_86 |
| DS113179 | 424836 | 425841 | BA_peak_87 |
| DS113179 | 447854 | 448201 | BA_peak_88 |
| DS113179 | 461041 | 461571 | BA_peak_89 |
| DS113179 | 462207 | 462964 | BA_peak_90 |
| DS113179 | 468930 | 469461 | BA_peak_91 |
| DS113179 | 476262 | 476916 | BA_peak_92 |
| DS113179 | 491734 | 494970 | BA_peak_93 |
| DS113180 | 30879 | 31600 | BA_peak_94 |
| DS113180 | 32086 | 32785 | BA_peak_95 |
| DS113180 | 58085 | 58397 | BA_peak_96 |
| DS113180 | 59655 | 60517 | BA_peak_97 |
| DS113180 | 91587 | 91907 | BA_peak_98 |
| DS113180 | 100217 | 100994 | BA_peak_99 |
| DS113180 | 102625 | 103061 | BA_peak_100 |

| | | | |
|---|---|---|---|
| DS113180 | 112035 | 112986 | BA_peak_101 |
| DS113180 | 114076 | 115076 | BA_peak_102 |
| DS113180 | 124577 | 124914 | BA_peak_103 |
| DS113180 | 134701 | 135620 | BA_peak_104 |
| DS113180 | 135975 | 136369 | BA_peak_105 |
| DS113180 | 149520 | 150302 | BA_peak_106 |
| DS113180 | 188635 | 189232 | BA_peak_107 |
| DS113180 | 202424 | 205067 | BA_peak_108 |
| DS113180 | 247366 | 248088 | BA_peak_109 |
| DS113180 | 250564 | 251191 | BA_peak_110 |
| DS113180 | 274467 | 274802 | BA_peak_111 |
| DS113180 | 278173 | 279110 | BA_peak_112 |
| DS113180 | 316235 | 316807 | BA_peak_113 |
| DS113180 | 335828 | 336691 | BA_peak_114 |
| DS113180 | 352653 | 353603 | BA_peak_115 |
| DS113180 | 422947 | 423717 | BA_peak_117 |
| DS113180 | 430788 | 431448 | BA_peak_118 |
| DS113180 | 442841 | 443627 | BA_peak_119 |
| DS113180 | 445367 | 446403 | BA_peak_120 |
| DS113181 | 5987 | 6372 | BA_peak_121 |
| DS113181 | 8149 | 8472 | BA_peak_122 |
| DS113181 | 8969 | 9512 | BA_peak_123 |
| DS113181 | 12746 | 13589 | BA_peak_125 |
| DS113181 | 15947 | 16364 | BA_peak_126 |
| DS113181 | 112950 | 113548 | BA_peak_127 |
| DS113181 | 114447 | 114762 | BA_peak_128 |
| DS113181 | 116647 | 118479 | BA_peak_129 |
| DS113181 | 137796 | 138143 | BA_peak_131 |
| DS113181 | 156397 | 156852 | BA_peak_132 |
| DS113181 | 167867 | 168608 | BA_peak_133 |
| DS113181 | 195582 | 195953 | BA_peak_134 |
| DS113181 | 196368 | 196675 | BA_peak_135 |
| DS113181 | 202833 | 203692 | BA_peak_136 |
| DS113181 | 214018 | 214610 | BA_peak_137 |
| DS113181 | 220292 | 221221 | BA_peak_138 |
| DS113181 | 225142 | 225619 | BA_peak_139 |
| DS113181 | 233323 | 235429 | BA_peak_140 |
| DS113181 | 236978 | 237864 | BA_peak_141 |
| DS113181 | 238940 | 240654 | BA_peak_142 |
| DS113181 | 243950 | 244484 | BA_peak_143 |
| DS113181 | 256474 | 256895 | BA_peak_144 |
| DS113181 | 257710 | 258076 | BA_peak_145 |
| DS113181 | 296696 | 297332 | BA_peak_146 |
| DS113181 | 341678 | 342326 | BA_peak_147 |
| DS113181 | 345349 | 346111 | BA_peak_148 |
| DS113181 | 373583 | 374258 | BA_peak_149 |
| DS113181 | 397414 | 397778 | BA_peak_150 |

| | | | |
|---|---|---|---|
| DS113181 | 400197 | 401215 | BA_peak_151 |
| DS113181 | 426653 | 427447 | BA_peak_152 |
| DS113181 | 432046 | 432659 | BA_peak_153 |
| DS113181 | 432988 | 434218 | BA_peak_154 |
| DS113181 | 435335 | 435956 | BA_peak_155 |
| DS113181 | 436510 | 437199 | BA_peak_156 |
| DS113182 | 15037 | 15368 | BA_peak_157 |
| DS113182 | 22248 | 22610 | BA_peak_158 |
| DS113182 | 44647 | 45841 | BA_peak_159 |
| DS113182 | 91494 | 92251 | BA_peak_160 |
| DS113182 | 98156 | 98824 | BA_peak_161 |
| DS113182 | 121800 | 122931 | BA_peak_162 |
| DS113182 | 148264 | 148854 | BA_peak_163 |
| DS113182 | 173656 | 174198 | BA_peak_164 |
| DS113182 | 195551 | 196178 | BA_peak_165 |
| DS113182 | 236918 | 243600 | BA_peak_166 |
| DS113182 | 252042 | 252370 | BA_peak_167 |
| DS113182 | 253301 | 253776 | BA_peak_168 |
| DS113182 | 305862 | 309150 | BA_peak_169 |
| DS113182 | 309474 | 310117 | BA_peak_170 |
| DS113182 | 312799 | 313490 | BA_peak_171 |
| DS113182 | 318489 | 318934 | BA_peak_172 |
| DS113182 | 320526 | 320965 | BA_peak_174 |
| DS113182 | 322604 | 324291 | BA_peak_175 |
| DS113182 | 328323 | 328866 | BA_peak_176 |
| DS113182 | 334845 | 335277 | BA_peak_177 |
| DS113182 | 354307 | 355579 | BA_peak_179 |
| DS113182 | 418744 | 420087 | BA_peak_180 |
| DS113182 | 422955 | 424074 | BA_peak_181 |
| DS113183 | 1954 | 3460 | BA_peak_183 |
| DS113183 | 4120 | 4487 | BA_peak_184 |
| DS113183 | 14049 | 14606 | BA_peak_185 |
| DS113183 | 29278 | 29899 | BA_peak_186 |
| DS113183 | 129014 | 129930 | BA_peak_187 |
| DS113183 | 173306 | 174013 | BA_peak_188 |
| DS113183 | 197988 | 198620 | BA_peak_189 |
| DS113183 | 205942 | 206711 | BA_peak_190 |
| DS113183 | 227048 | 227369 | BA_peak_191 |
| DS113183 | 232226 | 232936 | BA_peak_192 |
| DS113183 | 240590 | 241157 | BA_peak_193 |
| DS113183 | 259740 | 261221 | BA_peak_194 |
| DS113183 | 262692 | 263210 | BA_peak_195 |
| DS113183 | 265028 | 265862 | BA_peak_196 |
| DS113183 | 298812 | 299426 | BA_peak_197 |
| DS113183 | 325812 | 328080 | BA_peak_198 |
| DS113183 | 345748 | 346469 | BA_peak_199 |
| DS113183 | 347603 | 348016 | BA_peak_200 |

| | | | |
|---|---|---|---|
| DS113183 | 383825 | 384444 | BA_peak_201 |
| DS113183 | 385998 | 386588 | BA_peak_202 |
| DS113183 | 389849 | 390561 | BA_peak_203 |
| DS113183 | 427061 | 427408 | BA_peak_204 |
| DS113184 | 3232 | 3691 | BA_peak_205 |
| DS113184 | 23548 | 24726 | BA_peak_206 |
| DS113184 | 29258 | 29869 | BA_peak_207 |
| DS113184 | 32382 | 35400 | BA_peak_208 |
| DS113184 | 47258 | 47612 | BA_peak_209 |
| DS113184 | 47947 | 51307 | BA_peak_210 |
| DS113184 | 59788 | 60258 | BA_peak_211 |
| DS113184 | 62298 | 63674 | BA_peak_212 |
| DS113184 | 64812 | 65124 | BA_peak_213 |
| DS113184 | 66062 | 66569 | BA_peak_214 |
| DS113184 | 66898 | 67464 | BA_peak_215 |
| DS113184 | 70822 | 71738 | BA_peak_216 |
| DS113184 | 73750 | 74487 | BA_peak_217 |
| DS113184 | 89496 | 91368 | BA_peak_218 |
| DS113184 | 114981 | 115297 | BA_peak_219 |
| DS113184 | 126839 | 127721 | BA_peak_220 |
| DS113184 | 138005 | 138662 | BA_peak_221 |
| DS113184 | 143436 | 144135 | BA_peak_222 |
| DS113184 | 155895 | 156434 | BA_peak_223 |
| DS113184 | 181047 | 181556 | BA_peak_224 |
| DS113184 | 188027 | 188489 | BA_peak_225 |
| DS113184 | 249345 | 249974 | BA_peak_228 |
| DS113184 | 296413 | 296816 | BA_peak_230 |
| DS113184 | 308162 | 308685 | BA_peak_231 |
| DS113184 | 326423 | 327906 | BA_peak_233 |
| DS113184 | 382376 | 382741 | BA_peak_234 |
| DS113184 | 389766 | 390579 | BA_peak_235 |
| DS113184 | 391701 | 392518 | BA_peak_236 |
| DS113184 | 418658 | 419055 | BA_peak_237 |
| DS113184 | 420920 | 421672 | BA_peak_238 |
| DS113185 | 2185 | 2863 | BA_peak_239 |
| DS113185 | 8553 | 9046 | BA_peak_240 |
| DS113185 | 59012 | 59365 | BA_peak_241 |
| DS113185 | 73215 | 73929 | BA_peak_242 |
| DS113185 | 80147 | 84434 | BA_peak_243 |
| DS113185 | 91389 | 92408 | BA_peak_244 |
| DS113185 | 178162 | 178792 | BA_peak_245 |
| DS113185 | 195777 | 196505 | BA_peak_246 |
| DS113185 | 197484 | 198216 | BA_peak_247 |
| DS113185 | 201854 | 202451 | BA_peak_248 |
| DS113185 | 219863 | 220535 | BA_peak_249 |
| DS113185 | 238020 | 239426 | BA_peak_250 |
| DS113185 | 256963 | 257887 | BA_peak_251 |

| | | | |
|---|---|---|---|
| DS113185 | 259013 | 259824 | BA_peak_252 |
| DS113185 | 273252 | 273723 | BA_peak_253 |
| DS113185 | 281668 | 282151 | BA_peak_254 |
| DS113185 | 284603 | 285678 | BA_peak_255 |
| DS113185 | 287033 | 287586 | BA_peak_256 |
| DS113185 | 289462 | 290112 | BA_peak_257 |
| DS113185 | 292104 | 292988 | BA_peak_258 |
| DS113185 | 356887 | 357343 | BA_peak_259 |
| DS113185 | 383023 | 383617 | BA_peak_260 |
| DS113185 | 385727 | 386930 | BA_peak_261 |
| DS113185 | 410884 | 411327 | BA_peak_262 |
| DS113185 | 415096 | 415651 | BA_peak_263 |
| DS113186 | 5404 | 6074 | BA_peak_264 |
| DS113186 | 9237 | 9673 | BA_peak_265 |
| DS113186 | 10746 | 11089 | BA_peak_266 |
| DS113186 | 36028 | 36890 | BA_peak_267 |
| DS113186 | 45271 | 45950 | BA_peak_268 |
| DS113186 | 65380 | 66122 | BA_peak_269 |
| DS113186 | 103874 | 104587 | BA_peak_270 |
| DS113186 | 118481 | 118920 | BA_peak_271 |
| DS113186 | 135214 | 135910 | BA_peak_272 |
| DS113186 | 161502 | 162299 | BA_peak_273 |
| DS113186 | 162863 | 163903 | BA_peak_274 |
| DS113186 | 177240 | 178162 | BA_peak_275 |
| DS113186 | 217773 | 218422 | BA_peak_276 |
| DS113186 | 227604 | 228329 | BA_peak_277 |
| DS113186 | 283033 | 283473 | BA_peak_278 |
| DS113186 | 284013 | 285953 | BA_peak_279 |
| DS113186 | 293315 | 294294 | BA_peak_280 |
| DS113186 | 304567 | 305196 | BA_peak_281 |
| DS113186 | 327020 | 327640 | BA_peak_282 |
| DS113186 | 334675 | 335273 | BA_peak_283 |
| DS113186 | 335847 | 336266 | BA_peak_284 |
| DS113186 | 354947 | 355419 | BA_peak_285 |
| DS113186 | 377563 | 378335 | BA_peak_286 |
| DS113186 | 378791 | 379837 | BA_peak_287 |
| DS113186 | 395033 | 396786 | BA_peak_288 |
| DS113186 | 401638 | 402570 | BA_peak_289 |
| DS113186 | 404971 | 405390 | BA_peak_290 |
| DS113187 | 2116 | 3300 | BA_peak_291 |
| DS113187 | 3694 | 5168 | BA_peak_292 |
| DS113187 | 20346 | 20711 | BA_peak_293 |
| DS113187 | 33022 | 33586 | BA_peak_294 |
| DS113187 | 54081 | 55647 | BA_peak_295 |
| DS113187 | 67598 | 68500 | BA_peak_296 |
| DS113187 | 71908 | 72387 | BA_peak_297 |
| DS113187 | 81468 | 84759 | BA_peak_298 |

| | | | |
|---|---|---|---|
| DS113187 | 106005 | 106910 | BA_peak_299 |
| DS113187 | 120351 | 120749 | BA_peak_300 |
| DS113187 | 131231 | 131952 | BA_peak_301 |
| DS113187 | 140716 | 141140 | BA_peak_302 |
| DS113187 | 141781 | 142551 | BA_peak_303 |
| DS113187 | 143517 | 144148 | BA_peak_304 |
| DS113187 | 212925 | 213806 | BA_peak_305 |
| DS113187 | 232415 | 232945 | BA_peak_306 |
| DS113187 | 234115 | 235079 | BA_peak_307 |
| DS113187 | 236877 | 237570 | BA_peak_308 |
| DS113187 | 244056 | 244527 | BA_peak_309 |
| DS113187 | 250500 | 250939 | BA_peak_310 |
| DS113187 | 265886 | 266407 | BA_peak_311 |
| DS113187 | 306411 | 306955 | BA_peak_313 |
| DS113187 | 363365 | 364030 | BA_peak_314 |
| DS113187 | 379194 | 381294 | BA_peak_315 |
| DS113187 | 381973 | 382358 | BA_peak_316 |
| DS113188 | 5291 | 5918 | BA_peak_317 |
| DS113188 | 12546 | 12992 | BA_peak_318 |
| DS113188 | 16555 | 17484 | BA_peak_319 |
| DS113188 | 17798 | 18418 | BA_peak_320 |
| DS113188 | 49322 | 49783 | BA_peak_322 |
| DS113188 | 52237 | 53093 | BA_peak_323 |
| DS113188 | 55811 | 56660 | BA_peak_324 |
| DS113188 | 57590 | 58762 | BA_peak_325 |
| DS113188 | 61280 | 61768 | BA_peak_326 |
| DS113188 | 62580 | 63889 | BA_peak_327 |
| DS113188 | 98718 | 99810 | BA_peak_328 |
| DS113188 | 125876 | 126212 | BA_peak_330 |
| DS113188 | 144662 | 145586 | BA_peak_331 |
| DS113188 | 183001 | 183797 | BA_peak_332 |
| DS113188 | 224134 | 224806 | BA_peak_333 |
| DS113188 | 232645 | 234710 | BA_peak_334 |
| DS113188 | 252777 | 253450 | BA_peak_335 |
| DS113188 | 253875 | 254681 | BA_peak_336 |
| DS113188 | 277833 | 278596 | BA_peak_337 |
| DS113188 | 278907 | 279324 | BA_peak_338 |
| DS113188 | 279737 | 280589 | BA_peak_339 |
| DS113188 | 298916 | 299880 | BA_peak_340 |
| DS113188 | 320640 | 322455 | BA_peak_341 |
| DS113188 | 324059 | 325348 | BA_peak_342 |
| DS113188 | 328606 | 329411 | BA_peak_343 |
| DS113188 | 334952 | 335389 | BA_peak_344 |
| DS113188 | 349683 | 350293 | BA_peak_345 |
| DS113188 | 354048 | 354513 | BA_peak_346 |
| DS113189 | 15433 | 16485 | BA_peak_347 |
| DS113189 | 18090 | 19448 | BA_peak_348 |

| | | | |
|---|---|---|---|
| DS113189 | 53546 | 53854 | BA_peak_349 |
| DS113189 | 61326 | 62802 | BA_peak_350 |
| DS113189 | 84252 | 84538 | BA_peak_351 |
| DS113189 | 111259 | 111957 | BA_peak_353 |
| DS113189 | 134725 | 135334 | BA_peak_354 |
| DS113189 | 136830 | 137994 | BA_peak_355 |
| DS113189 | 143666 | 144012 | BA_peak_356 |
| DS113189 | 151667 | 152011 | BA_peak_359 |
| DS113189 | 159017 | 159525 | BA_peak_360 |
| DS113189 | 159943 | 160666 | BA_peak_361 |
| DS113189 | 198760 | 199556 | BA_peak_362 |
| DS113189 | 212002 | 213430 | BA_peak_363 |
| DS113189 | 239680 | 241015 | BA_peak_364 |
| DS113189 | 275092 | 275729 | BA_peak_365 |
| DS113189 | 318335 | 319144 | BA_peak_366 |
| DS113189 | 331045 | 331860 | BA_peak_367 |
| DS113189 | 332297 | 334367 | BA_peak_368 |
| DS113189 | 341572 | 342078 | BA_peak_369 |
| DS113189 | 342478 | 343093 | BA_peak_370 |
| DS113189 | 344413 | 344754 | BA_peak_371 |
| DS113190 | 239 | 736 | BA_peak_373 |
| DS113190 | 1297 | 2742 | BA_peak_374 |
| DS113190 | 3499 | 6065 | BA_peak_375 |
| DS113190 | 14033 | 14686 | BA_peak_376 |
| DS113190 | 19227 | 19722 | BA_peak_377 |
| DS113190 | 20671 | 21479 | BA_peak_378 |
| DS113190 | 46702 | 47281 | BA_peak_379 |
| DS113190 | 105883 | 106705 | BA_peak_381 |
| DS113190 | 168756 | 169486 | BA_peak_382 |
| DS113190 | 170131 | 170527 | BA_peak_383 |
| DS113190 | 175969 | 176857 | BA_peak_384 |
| DS113190 | 177914 | 178989 | BA_peak_385 |
| DS113190 | 206935 | 207300 | BA_peak_386 |
| DS113190 | 208793 | 211131 | BA_peak_387 |
| DS113190 | 225072 | 225875 | BA_peak_388 |
| DS113190 | 282131 | 282568 | BA_peak_389 |
| DS113190 | 288250 | 288625 | BA_peak_390 |
| DS113191 | 53387 | 53896 | BA_peak_391 |
| DS113191 | 54482 | 55126 | BA_peak_392 |
| DS113191 | 79510 | 80926 | BA_peak_393 |
| DS113191 | 83644 | 84936 | BA_peak_394 |
| DS113191 | 105418 | 106108 | BA_peak_395 |
| DS113191 | 111030 | 111432 | BA_peak_396 |
| DS113191 | 118082 | 118766 | BA_peak_398 |
| DS113191 | 119076 | 119834 | BA_peak_399 |
| DS113191 | 140013 | 142915 | BA_peak_401 |
| DS113191 | 145977 | 147917 | BA_peak_403 |

| | | | |
|---|---|---|---|
| DS113191 | 156398 | 157306 | BA_peak_404 |
| DS113191 | 182317 | 183087 | BA_peak_405 |
| DS113191 | 189989 | 192256 | BA_peak_406 |
| DS113191 | 198925 | 199327 | BA_peak_407 |
| DS113191 | 217963 | 218560 | BA_peak_408 |
| DS113191 | 221771 | 222974 | BA_peak_410 |
| DS113191 | 273611 | 274050 | BA_peak_411 |
| DS113191 | 278469 | 280614 | BA_peak_412 |
| DS113191 | 290873 | 291555 | BA_peak_413 |
| DS113191 | 309783 | 311131 | BA_peak_414 |
| DS113191 | 311695 | 313422 | BA_peak_415 |
| DS113191 | 320247 | 321112 | BA_peak_416 |
| DS113191 | 321433 | 322908 | BA_peak_417 |
| DS113192 | 26956 | 27608 | BA_peak_419 |
| DS113192 | 59734 | 61848 | BA_peak_420 |
| DS113192 | 132110 | 132879 | BA_peak_421 |
| DS113192 | 139544 | 139949 | BA_peak_422 |
| DS113192 | 161210 | 162629 | BA_peak_423 |
| DS113192 | 214018 | 214494 | BA_peak_424 |
| DS113192 | 278773 | 279608 | BA_peak_425 |
| DS113192 | 284188 | 284938 | BA_peak_426 |
| DS113192 | 314406 | 315916 | BA_peak_427 |
| DS113192 | 317000 | 317777 | BA_peak_428 |
| DS113192 | 323394 | 324065 | BA_peak_429 |
| DS113192 | 325089 | 326232 | BA_peak_430 |
| DS113192 | 343141 | 343460 | BA_peak_431 |
| DS113193 | 10288 | 10945 | BA_peak_432 |
| DS113193 | 35142 | 35825 | BA_peak_433 |
| DS113193 | 45105 | 46095 | BA_peak_435 |
| DS113193 | 65228 | 66326 | BA_peak_438 |
| DS113193 | 76749 | 77364 | BA_peak_439 |
| DS113193 | 105505 | 106216 | BA_peak_440 |
| DS113193 | 106928 | 108287 | BA_peak_441 |
| DS113193 | 133132 | 134448 | BA_peak_442 |
| DS113193 | 135918 | 136230 | BA_peak_443 |
| DS113193 | 301745 | 302333 | BA_peak_445 |
| DS113193 | 315185 | 315836 | BA_peak_446 |
| DS113193 | 326244 | 327026 | BA_peak_447 |
| DS113194 | 40923 | 41387 | BA_peak_448 |
| DS113194 | 42140 | 42569 | BA_peak_449 |
| DS113194 | 69292 | 70139 | BA_peak_450 |
| DS113194 | 143488 | 144247 | BA_peak_451 |
| DS113194 | 157405 | 157822 | BA_peak_452 |
| DS113194 | 177316 | 180655 | BA_peak_453 |
| DS113194 | 187952 | 188530 | BA_peak_454 |
| DS113194 | 207451 | 208295 | BA_peak_455 |
| DS113194 | 238773 | 239442 | BA_peak_456 |

| | | | |
|---|---|---|---|
| DS113194 | 260949 | 264312 | BA_peak_457 |
| DS113194 | 276505 | 277113 | BA_peak_458 |
| DS113194 | 297494 | 298209 | BA_peak_459 |
| DS113195 | 4409 | 4932 | BA_peak_460 |
| DS113195 | 5330 | 5797 | BA_peak_460 |
| DS113195 | 7257 | 9462 | BA_peak_461 |
| DS113195 | 14566 | 15035 | BA_peak_462 |
| DS113195 | 24101 | 27286 | BA_peak_463 |
| DS113195 | 78207 | 78970 | BA_peak_464 |
| DS113195 | 98106 | 100975 | BA_peak_465 |
| DS113195 | 101281 | 101909 | BA_peak_466 |
| DS113195 | 121592 | 122069 | BA_peak_467 |
| DS113195 | 170707 | 171531 | BA_peak_468 |
| DS113195 | 206232 | 207300 | BA_peak_469 |
| DS113195 | 224723 | 226290 | BA_peak_470 |
| DS113195 | 230863 | 231609 | BA_peak_471 |
| DS113195 | 242926 | 243674 | BA_peak_472 |
| DS113195 | 250852 | 251627 | BA_peak_473 |
| DS113195 | 272690 | 274285 | BA_peak_475 |
| DS113195 | 283715 | 284435 | BA_peak_476 |
| DS113195 | 286811 | 287655 | BA_peak_477 |
| DS113195 | 288504 | 289340 | BA_peak_478 |
| DS113195 | 296216 | 297975 | BA_peak_479 |
| DS113196 | 1479 | 1909 | BA_peak_480 |
| DS113196 | 94883 | 95268 | BA_peak_482 |
| DS113196 | 98479 | 99219 | BA_peak_483 |
| DS113196 | 141221 | 141970 | BA_peak_484 |
| DS113196 | 148961 | 149694 | BA_peak_485 |
| DS113196 | 167117 | 167660 | BA_peak_486 |
| DS113196 | 169492 | 171570 | BA_peak_487 |
| DS113196 | 172990 | 174256 | BA_peak_488 |
| DS113196 | 191989 | 192610 | BA_peak_489 |
| DS113196 | 195945 | 197819 | BA_peak_490 |
| DS113196 | 199618 | 200379 | BA_peak_491 |
| DS113196 | 216876 | 217388 | BA_peak_492 |
| DS113196 | 225003 | 225769 | BA_peak_493 |
| DS113196 | 226083 | 226599 | BA_peak_494 |
| DS113196 | 249089 | 252320 | BA_peak_495 |
| DS113196 | 289324 | 290686 | BA_peak_496 |
| DS113197 | 122031 | 122365 | BA_peak_497 |
| DS113197 | 130249 | 130664 | BA_peak_498 |
| DS113197 | 140466 | 141302 | BA_peak_499 |
| DS113197 | 145699 | 146106 | BA_peak_500 |
| DS113197 | 158993 | 159977 | BA_peak_501 |
| DS113197 | 204327 | 205512 | BA_peak_502 |
| DS113197 | 216540 | 216901 | BA_peak_503 |
| DS113197 | 234739 | 235584 | BA_peak_504 |

| | | | |
|---|---|---|---|
| DS113197 | 281106 | 282471 | BA_peak_505 |
| DS113197 | 286432 | 287829 | BA_peak_506 |
| DS113197 | 291045 | 291789 | BA_peak_507 |
| DS113198 | 10426 | 11279 | BA_peak_508 |
| DS113198 | 37883 | 38146 | BA_peak_510 |
| DS113198 | 39063 | 39684 | BA_peak_511 |
| DS113198 | 40055 | 41824 | BA_peak_512 |
| DS113198 | 53347 | 55130 | BA_peak_514 |
| DS113198 | 56695 | 57564 | BA_peak_515 |
| DS113198 | 76329 | 76907 | BA_peak_516 |
| DS113198 | 114315 | 114753 | BA_peak_517 |
| DS113198 | 124444 | 125206 | BA_peak_518 |
| DS113198 | 159809 | 160116 | BA_peak_519 |
| DS113198 | 161167 | 162619 | BA_peak_520 |
| DS113198 | 217212 | 217798 | BA_peak_521 |
| DS113198 | 235248 | 235735 | BA_peak_522 |
| DS113198 | 239093 | 239936 | BA_peak_523 |
| DS113198 | 245354 | 246457 | BA_peak_524 |
| DS113198 | 254664 | 254976 | BA_peak_525 |
| DS113198 | 265049 | 265591 | BA_peak_526 |
| DS113198 | 278907 | 279462 | BA_peak_527 |
| DS113199 | 26358 | 28746 | BA_peak_528 |
| DS113199 | 29087 | 32474 | BA_peak_529 |
| DS113199 | 65685 | 66089 | BA_peak_530 |
| DS113199 | 73562 | 73896 | BA_peak_531 |
| DS113199 | 86494 | 87163 | BA_peak_532 |
| DS113199 | 99924 | 101034 | BA_peak_533 |
| DS113199 | 149210 | 149696 | BA_peak_534 |
| DS113199 | 161987 | 163007 | BA_peak_535 |
| DS113199 | 172929 | 173675 | BA_peak_536 |
| DS113199 | 205154 | 205935 | BA_peak_537 |
| DS113199 | 206460 | 206835 | BA_peak_538 |
| DS113199 | 246627 | 248178 | BA_peak_539 |
| DS113199 | 250687 | 251463 | BA_peak_540 |
| DS113199 | 271624 | 272909 | BA_peak_541 |
| DS113199 | 275094 | 275838 | BA_peak_542 |
| DS113200 | 2635 | 3226 | BA_peak_543 |
| DS113200 | 5894 | 7234 | BA_peak_544 |
| DS113200 | 28880 | 29666 | BA_peak_545 |
| DS113200 | 35924 | 36267 | BA_peak_546 |
| DS113200 | 52850 | 53569 | BA_peak_547 |
| DS113200 | 93960 | 94662 | BA_peak_548 |
| DS113200 | 98742 | 99369 | BA_peak_549 |
| DS113200 | 127072 | 128497 | BA_peak_550 |
| DS113200 | 132878 | 134559 | BA_peak_551 |
| DS113200 | 173385 | 173720 | BA_peak_552 |
| DS113200 | 198305 | 198968 | BA_peak_553 |

| | | | |
|---|---|---|---|
| DS113200 | 213417 | 213911 | BA_peak_554 |
| DS113200 | 222324 | 223035 | BA_peak_555 |
| DS113200 | 223704 | 224333 | BA_peak_556 |
| DS113200 | 249893 | 251541 | BA_peak_557 |
| DS113200 | 253116 | 255291 | BA_peak_558 |
| DS113201 | 44247 | 45049 | BA_peak_559 |
| DS113201 | 72793 | 73771 | BA_peak_560 |
| DS113201 | 111821 | 112325 | BA_peak_561 |
| DS113201 | 138457 | 139345 | BA_peak_562 |
| DS113201 | 163039 | 165288 | BA_peak_563 |
| DS113201 | 173191 | 173885 | BA_peak_564 |
| DS113201 | 194459 | 195236 | BA_peak_565 |
| DS113201 | 211075 | 211587 | BA_peak_566 |
| DS113201 | 215154 | 215908 | BA_peak_567 |
| DS113201 | 216221 | 216658 | BA_peak_568 |
| DS113201 | 223235 | 223962 | BA_peak_569 |
| DS113201 | 224330 | 225242 | BA_peak_570 |
| DS113201 | 244446 | 244853 | BA_peak_572 |
| DS113202 | 2109 | 2942 | BA_peak_573 |
| DS113202 | 12011 | 12518 | BA_peak_574 |
| DS113202 | 12966 | 13399 | BA_peak_575 |
| DS113202 | 29507 | 30938 | BA_peak_576 |
| DS113202 | 46434 | 47664 | BA_peak_577 |
| DS113202 | 79644 | 79992 | BA_peak_578 |
| DS113202 | 85522 | 86357 | BA_peak_579 |
| DS113202 | 87520 | 87925 | BA_peak_580 |
| DS113202 | 99764 | 100546 | BA_peak_581 |
| DS113202 | 101642 | 102235 | BA_peak_582 |
| DS113202 | 119497 | 121148 | BA_peak_583 |
| DS113202 | 145085 | 145902 | BA_peak_584 |
| DS113202 | 158594 | 161822 | BA_peak_585 |
| DS113202 | 165794 | 167582 | BA_peak_586 |
| DS113202 | 183495 | 184862 | BA_peak_587 |
| DS113202 | 187406 | 187726 | BA_peak_588 |
| DS113202 | 191513 | 193017 | BA_peak_589 |
| DS113202 | 194138 | 198661 | BA_peak_590 |
| DS113202 | 206828 | 207494 | BA_peak_591 |
| DS113202 | 209476 | 209876 | BA_peak_592 |
| DS113202 | 236139 | 236588 | BA_peak_594 |
| DS113202 | 241457 | 242584 | BA_peak_595 |
| DS113202 | 244764 | 245237 | BA_peak_596 |
| DS113202 | 247762 | 248225 | BA_peak_597 |
| DS113202 | 251847 | 252972 | BA_peak_598 |
| DS113203 | 1234 | 1547 | BA_peak_599 |
| DS113203 | 11729 | 12711 | BA_peak_600 |
| DS113203 | 29943 | 30315 | BA_peak_601 |
| DS113203 | 72301 | 73158 | BA_peak_602 |

| | | | |
|---|---|---|---|
| DS113203 | 117173 | 117514 | BA_peak_603 |
| DS113203 | 118410 | 119504 | BA_peak_604 |
| DS113203 | 120844 | 121834 | BA_peak_605 |
| DS113203 | 138866 | 139642 | BA_peak_606 |
| DS113203 | 145088 | 145653 | BA_peak_607 |
| DS113203 | 219607 | 220380 | BA_peak_608 |
| DS113203 | 246800 | 247144 | BA_peak_609 |
| DS113203 | 250463 | 251119 | BA_peak_610 |
| DS113203 | 251598 | 252242 | BA_peak_611 |
| DS113204 | 20307 | 20964 | BA_peak_612 |
| DS113204 | 30471 | 30876 | BA_peak_614 |
| DS113204 | 31710 | 32323 | BA_peak_615 |
| DS113204 | 32748 | 33045 | BA_peak_616 |
| DS113204 | 36048 | 36366 | BA_peak_617 |
| DS113204 | 36896 | 37428 | BA_peak_618 |
| DS113204 | 54081 | 54818 | BA_peak_619 |
| DS113204 | 56176 | 57488 | BA_peak_620 |
| DS113204 | 58973 | 59902 | BA_peak_621 |
| DS113204 | 60278 | 61472 | BA_peak_622 |
| DS113204 | 85219 | 85697 | BA_peak_623 |
| DS113204 | 97188 | 97564 | BA_peak_624 |
| DS113204 | 97982 | 98493 | BA_peak_625 |
| DS113204 | 99399 | 99824 | BA_peak_626 |
| DS113204 | 124348 | 124704 | BA_peak_627 |
| DS113204 | 127067 | 127538 | BA_peak_628 |
| DS113204 | 133319 | 133727 | BA_peak_629 |
| DS113204 | 160192 | 161055 | BA_peak_630 |
| DS113204 | 187038 | 187599 | BA_peak_633 |
| DS113204 | 213130 | 216033 | BA_peak_635 |
| DS113204 | 224582 | 225010 | BA_peak_636 |
| DS113204 | 227826 | 228562 | BA_peak_637 |
| DS113204 | 246343 | 246619 | BA_peak_638 |
| DS113205 | 71032 | 71602 | BA_peak_639 |
| DS113205 | 73130 | 73887 | BA_peak_640 |
| DS113205 | 74414 | 76010 | BA_peak_641 |
| DS113205 | 92161 | 92599 | BA_peak_642 |
| DS113205 | 94922 | 96313 | BA_peak_643 |
| DS113205 | 101175 | 101516 | BA_peak_644 |
| DS113205 | 174721 | 175495 | BA_peak_647 |
| DS113205 | 178689 | 179225 | BA_peak_648 |
| DS113205 | 188290 | 188652 | BA_peak_649 |
| DS113205 | 192046 | 192396 | BA_peak_651 |
| DS113205 | 198238 | 199003 | BA_peak_652 |
| DS113205 | 214303 | 214729 | BA_peak_653 |
| DS113205 | 235755 | 236437 | BA_peak_654 |
| DS113206 | 8802 | 9330 | BA_peak_655 |
| DS113206 | 12653 | 12999 | BA_peak_656 |

| | | | |
|---|---|---|---|
| DS113206 | 16237 | 16701 | BA_peak_657 |
| DS113206 | 36345 | 36881 | BA_peak_658 |
| DS113206 | 40590 | 41402 | BA_peak_659 |
| DS113206 | 42215 | 42878 | BA_peak_660 |
| DS113206 | 99602 | 100160 | BA_peak_661 |
| DS113206 | 102670 | 103134 | BA_peak_662 |
| DS113206 | 119087 | 119428 | BA_peak_663 |
| DS113206 | 119876 | 120247 | BA_peak_664 |
| DS113206 | 124810 | 125357 | BA_peak_665 |
| DS113206 | 125983 | 126319 | BA_peak_666 |
| DS113206 | 126857 | 127257 | BA_peak_667 |
| DS113206 | 127745 | 128469 | BA_peak_668 |
| DS113206 | 162789 | 163449 | BA_peak_670 |
| DS113206 | 172110 | 172558 | BA_peak_671 |
| DS113206 | 172949 | 173310 | BA_peak_672 |
| DS113206 | 174396 | 174703 | BA_peak_673 |
| DS113206 | 185569 | 186362 | BA_peak_674 |
| DS113206 | 203831 | 204230 | BA_peak_675 |
| DS113206 | 216093 | 216748 | BA_peak_676 |
| DS113207 | 10295 | 11440 | BA_peak_677 |
| DS113207 | 12522 | 12952 | BA_peak_678 |
| DS113207 | 16968 | 17741 | BA_peak_679 |
| DS113207 | 23577 | 23929 | BA_peak_680 |
| DS113207 | 47941 | 51075 | BA_peak_681 |
| DS113207 | 58944 | 59416 | BA_peak_682 |
| DS113207 | 77425 | 77798 | BA_peak_683 |
| DS113207 | 99831 | 101173 | BA_peak_684 |
| DS113207 | 104305 | 104968 | BA_peak_685 |
| DS113207 | 107674 | 108415 | BA_peak_686 |
| DS113207 | 111207 | 112136 | BA_peak_687 |
| DS113207 | 118480 | 119646 | BA_peak_688 |
| DS113207 | 120677 | 124079 | BA_peak_689 |
| DS113207 | 150691 | 151974 | BA_peak_690 |
| DS113207 | 173066 | 173539 | BA_peak_692 |
| DS113207 | 176546 | 177383 | BA_peak_693 |
| DS113207 | 179409 | 180136 | BA_peak_695 |
| DS113207 | 200617 | 200969 | BA_peak_696 |
| DS113207 | 206402 | 206747 | BA_peak_697 |
| DS113207 | 217518 | 218326 | BA_peak_698 |
| DS113208 | 4438 | 6800 | BA_peak_699 |
| DS113208 | 8696 | 9076 | BA_peak_700 |
| DS113208 | 24318 | 25620 | BA_peak_701 |
| DS113208 | 34312 | 35260 | BA_peak_702 |
| DS113208 | 54391 | 55993 | BA_peak_703 |
| DS113208 | 57373 | 60997 | BA_peak_704 |
| DS113208 | 84777 | 85418 | BA_peak_706 |
| DS113208 | 88106 | 88592 | BA_peak_707 |

| | | | |
|---|---|---|---|
| DS113208 | 92644 | 93064 | BA_peak_708 |
| DS113208 | 111649 | 112172 | BA_peak_709 |
| DS113208 | 116243 | 116933 | BA_peak_710 |
| DS113208 | 126579 | 127162 | BA_peak_711 |
| DS113208 | 131362 | 133484 | BA_peak_712 |
| DS113208 | 137812 | 138811 | BA_peak_713 |
| DS113208 | 146342 | 146991 | BA_peak_714 |
| DS113208 | 163626 | 164961 | BA_peak_715 |
| DS113208 | 170335 | 173674 | BA_peak_716 |
| DS113208 | 179833 | 180589 | BA_peak_717 |
| DS113208 | 194892 | 195719 | BA_peak_718 |
| DS113208 | 198706 | 199097 | BA_peak_719 |
| DS113208 | 212594 | 213279 | BA_peak_720 |
| DS113208 | 217334 | 218276 | BA_peak_722 |
| DS113208 | 236185 | 236844 | BA_peak_723 |
| DS113208 | 237279 | 237683 | BA_peak_724 |
| DS113209 | 10394 | 10857 | BA_peak_725 |
| DS113209 | 26759 | 29977 | BA_peak_726 |
| DS113209 | 41007 | 41546 | BA_peak_727 |
| DS113209 | 60139 | 60527 | BA_peak_728 |
| DS113209 | 79960 | 80600 | BA_peak_729 |
| DS113209 | 98798 | 105733 | BA_peak_730 |
| DS113209 | 117451 | 118125 | BA_peak_731 |
| DS113209 | 125516 | 126569 | BA_peak_732 |
| DS113209 | 139446 | 139890 | BA_peak_733 |
| DS113209 | 187572 | 188267 | BA_peak_734 |
| DS113209 | 209444 | 209910 | BA_peak_735 |
| DS113210 | 12507 | 13208 | BA_peak_736 |
| DS113210 | 58731 | 59433 | BA_peak_737 |
| DS113210 | 110737 | 113161 | BA_peak_738 |
| DS113210 | 135935 | 136540 | BA_peak_739 |
| DS113210 | 137244 | 138058 | BA_peak_740 |
| DS113210 | 138976 | 139379 | BA_peak_741 |
| DS113210 | 140635 | 141170 | BA_peak_742 |
| DS113210 | 141493 | 141826 | BA_peak_743 |
| DS113210 | 159475 | 159899 | BA_peak_744 |
| DS113210 | 161301 | 162012 | BA_peak_745 |
| DS113210 | 163037 | 164011 | BA_peak_746 |
| DS113210 | 165254 | 166396 | BA_peak_747 |
| DS113210 | 168808 | 169815 | BA_peak_749 |
| DS113210 | 170472 | 170787 | BA_peak_750 |
| DS113210 | 188896 | 189343 | BA_peak_751 |
| DS113210 | 210420 | 211241 | BA_peak_752 |
| DS113211 | 50 | 1759 | BA_peak_753 |
| DS113211 | 4535 | 4847 | BA_peak_754 |
| DS113211 | 83453 | 84593 | BA_peak_755 |
| DS113211 | 130158 | 131449 | BA_peak_756 |

| | | | |
|---|---|---|---|
| DS113211 | 168100 | 169000 | BA_peak_757 |
| DS113211 | 173170 | 174133 | BA_peak_758 |
| DS113211 | 202832 | 205017 | BA_peak_759 |
| DS113211 | 207122 | 209338 | BA_peak_760 |
| DS113212 | 4412 | 6019 | BA_peak_761 |
| DS113212 | 40394 | 41157 | BA_peak_762 |
| DS113212 | 47931 | 48770 | BA_peak_763 |
| DS113212 | 59384 | 59629 | BA_peak_764 |
| DS113212 | 61972 | 62626 | BA_peak_765 |
| DS113212 | 66289 | 66674 | BA_peak_766 |
| DS113212 | 68291 | 68792 | BA_peak_767 |
| DS113212 | 115018 | 115397 | BA_peak_768 |
| DS113212 | 116587 | 117937 | BA_peak_769 |
| DS113212 | 125130 | 125433 | BA_peak_770 |
| DS113212 | 159730 | 160164 | BA_peak_771 |
| DS113212 | 185710 | 186340 | BA_peak_772 |
| DS113212 | 197532 | 198327 | BA_peak_773 |
| DS113212 | 208532 | 209081 | BA_peak_774 |
| DS113212 | 224435 | 224846 | BA_peak_775 |
| DS113212 | 226436 | 227167 | BA_peak_776 |
| DS113213 | 1723 | 2382 | BA_peak_777 |
| DS113213 | 5470 | 5827 | BA_peak_778 |
| DS113213 | 62209 | 62663 | BA_peak_779 |
| DS113213 | 70952 | 71548 | BA_peak_780 |
| DS113213 | 78806 | 80007 | BA_peak_781 |
| DS113213 | 80791 | 81716 | BA_peak_782 |
| DS113213 | 86004 | 86394 | BA_peak_783 |
| DS113213 | 94841 | 95423 | BA_peak_784 |
| DS113213 | 112520 | 112855 | BA_peak_785 |
| DS113213 | 124378 | 124735 | BA_peak_786 |
| DS113213 | 141319 | 142018 | BA_peak_787 |
| DS113213 | 142631 | 143028 | BA_peak_788 |
| DS113213 | 172118 | 172524 | BA_peak_789 |
| DS113213 | 176882 | 178230 | BA_peak_790 |
| DS113213 | 190414 | 190787 | BA_peak_791 |
| DS113213 | 221394 | 221872 | BA_peak_792 |
| DS113214 | 8337 | 8899 | BA_peak_793 |
| DS113214 | 26975 | 27312 | BA_peak_794 |
| DS113214 | 70088 | 70530 | BA_peak_795 |
| DS113214 | 73951 | 74284 | BA_peak_796 |
| DS113214 | 75157 | 75675 | BA_peak_797 |
| DS113214 | 93389 | 93996 | BA_peak_798 |
| DS113214 | 107033 | 107957 | BA_peak_799 |
| DS113214 | 108404 | 109785 | BA_peak_800 |
| DS113214 | 116262 | 120658 | BA_peak_801 |
| DS113214 | 142997 | 143580 | BA_peak_802 |
| DS113214 | 159343 | 160115 | BA_peak_803 |

| | | | |
|---|---|---|---|
| DS113214 | 161044 | 161877 | BA_peak_804 |
| DS113214 | 201842 | 205568 | BA_peak_805 |
| DS113215 | 15673 | 17136 | BA_peak_806 |
| DS113215 | 34037 | 35204 | BA_peak_807 |
| DS113215 | 54697 | 55238 | BA_peak_808 |
| DS113215 | 61625 | 62685 | BA_peak_809 |
| DS113215 | 68080 | 69729 | BA_peak_810 |
| DS113215 | 114985 | 115497 | BA_peak_811 |
| DS113215 | 116365 | 116709 | BA_peak_812 |
| DS113215 | 118427 | 118793 | BA_peak_813 |
| DS113215 | 147078 | 147621 | BA_peak_814 |
| DS113215 | 168409 | 168937 | BA_peak_815 |
| DS113215 | 175871 | 176489 | BA_peak_816 |
| DS113215 | 199289 | 199740 | BA_peak_817 |
| DS113215 | 201575 | 201998 | BA_peak_818 |
| DS113216 | 74624 | 76769 | BA_peak_819 |
| DS113216 | 93766 | 94119 | BA_peak_820 |
| DS113216 | 99122 | 99902 | BA_peak_821 |
| DS113216 | 127760 | 131110 | BA_peak_822 |
| DS113216 | 180873 | 181734 | BA_peak_823 |
| DS113216 | 182711 | 184800 | BA_peak_824 |
| DS113216 | 194903 | 199101 | BA_peak_825 |
| DS113216 | 217936 | 218329 | BA_peak_826 |
| DS113216 | 220625 | 221200 | BA_peak_827 |
| DS113217 | 3876 | 4567 | BA_peak_828 |
| DS113217 | 9270 | 10021 | BA_peak_829 |
| DS113217 | 17831 | 18341 | BA_peak_830 |
| DS113217 | 40597 | 41843 | BA_peak_831 |
| DS113217 | 45389 | 46167 | BA_peak_832 |
| DS113217 | 49056 | 49433 | BA_peak_833 |
| DS113217 | 52921 | 54699 | BA_peak_834 |
| DS113217 | 98542 | 99262 | BA_peak_835 |
| DS113217 | 106095 | 106751 | BA_peak_836 |
| DS113217 | 111249 | 111938 | BA_peak_837 |
| DS113217 | 120921 | 121273 | BA_peak_838 |
| DS113217 | 141101 | 141634 | BA_peak_839 |
| DS113217 | 142343 | 143204 | BA_peak_840 |
| DS113217 | 150871 | 152050 | BA_peak_841 |
| DS113217 | 214979 | 215548 | BA_peak_842 |
| DS113217 | 216262 | 216598 | BA_peak_843 |
| DS113217 | 217257 | 218071 | BA_peak_844 |
| DS113218 | 6908 | 7837 | BA_peak_845 |
| DS113218 | 58661 | 60130 | BA_peak_846 |
| DS113218 | 81409 | 81757 | BA_peak_847 |
| DS113218 | 89787 | 90558 | BA_peak_848 |
| DS113218 | 110746 | 111151 | BA_peak_849 |
| DS113218 | 123082 | 123518 | BA_peak_850 |

| | | | |
|---|---|---|---|
| DS113218 | 158067 | 158477 | BA_peak_851 |
| DS113218 | 168518 | 171474 | BA_peak_852 |
| DS113218 | 182404 | 183334 | BA_peak_853 |
| DS113218 | 198124 | 198602 | BA_peak_854 |
| DS113218 | 218007 | 218359 | BA_peak_856 |
| DS113219 | 5647 | 6253 | BA_peak_857 |
| DS113219 | 18771 | 20151 | BA_peak_858 |
| DS113219 | 31870 | 32245 | BA_peak_859 |
| DS113219 | 64040 | 64500 | BA_peak_860 |
| DS113219 | 115515 | 116113 | BA_peak_861 |
| DS113219 | 183903 | 184269 | BA_peak_863 |
| DS113219 | 184590 | 186353 | BA_peak_864 |
| DS113219 | 186981 | 187677 | BA_peak_865 |
| DS113219 | 189560 | 190956 | BA_peak_866 |
| DS113219 | 220032 | 220588 | BA_peak_867 |
| DS113220 | 25566 | 27122 | BA_peak_868 |
| DS113220 | 27615 | 29405 | BA_peak_869 |
| DS113220 | 81579 | 82473 | BA_peak_870 |
| DS113220 | 87709 | 88292 | BA_peak_871 |
| DS113220 | 110413 | 111232 | BA_peak_872 |
| DS113220 | 119799 | 121139 | BA_peak_873 |
| DS113220 | 129188 | 129511 | BA_peak_874 |
| DS113220 | 132345 | 133557 | BA_peak_875 |
| DS113220 | 134181 | 136066 | BA_peak_876 |
| DS113220 | 138509 | 139041 | BA_peak_877 |
| DS113220 | 162974 | 163900 | BA_peak_878 |
| DS113220 | 169113 | 170518 | BA_peak_879 |
| DS113220 | 171067 | 171579 | BA_peak_880 |
| DS113220 | 175874 | 176400 | BA_peak_881 |
| DS113220 | 185817 | 186605 | BA_peak_882 |
| DS113220 | 197088 | 197894 | BA_peak_883 |
| DS113220 | 210073 | 211231 | BA_peak_884 |
| DS113221 | 40965 | 41363 | BA_peak_885 |
| DS113221 | 66424 | 67380 | BA_peak_886 |
| DS113221 | 77110 | 77612 | BA_peak_887 |
| DS113221 | 80679 | 81651 | BA_peak_888 |
| DS113221 | 92721 | 93500 | BA_peak_889 |
| DS113221 | 108448 | 108844 | BA_peak_890 |
| DS113221 | 126618 | 127647 | BA_peak_891 |
| DS113221 | 141213 | 142353 | BA_peak_892 |
| DS113221 | 151684 | 152187 | BA_peak_893 |
| DS113221 | 209795 | 210687 | BA_peak_894 |
| DS113222 | 41712 | 42254 | BA_peak_895 |
| DS113222 | 43328 | 44098 | BA_peak_896 |
| DS113222 | 66656 | 67659 | BA_peak_897 |
| DS113222 | 144191 | 144634 | BA_peak_898 |
| DS113222 | 155820 | 156189 | BA_peak_899 |

| | | | |
|---|---|---|---|
| DS113222 | 169959 | 170471 | BA_peak_900 |
| DS113222 | 170854 | 171675 | BA_peak_901 |
| DS113222 | 177565 | 178223 | BA_peak_902 |
| DS113222 | 178596 | 178945 | BA_peak_903 |
| DS113222 | 180186 | 180814 | BA_peak_904 |
| DS113222 | 188843 | 189552 | BA_peak_905 |
| DS113222 | 196246 | 196890 | BA_peak_906 |
| DS113223 | 14236 | 15157 | BA_peak_907 |
| DS113223 | 15471 | 15856 | BA_peak_908 |
| DS113223 | 76500 | 76848 | BA_peak_909 |
| DS113223 | 88390 | 88774 | BA_peak_910 |
| DS113223 | 130003 | 130574 | BA_peak_911 |
| DS113223 | 148910 | 149642 | BA_peak_912 |
| DS113223 | 160218 | 160656 | BA_peak_914 |
| DS113223 | 161312 | 161691 | BA_peak_915 |
| DS113224 | 6240 | 7122 | BA_peak_916 |
| DS113224 | 101311 | 101759 | BA_peak_917 |
| DS113224 | 108235 | 109614 | BA_peak_918 |
| DS113224 | 132834 | 133460 | BA_peak_919 |
| DS113224 | 153961 | 154631 | BA_peak_920 |
| DS113224 | 166703 | 167701 | BA_peak_921 |
| DS113224 | 171066 | 171817 | BA_peak_922 |
| DS113224 | 179899 | 180458 | BA_peak_923 |
| DS113224 | 186059 | 188333 | BA_peak_924 |
| DS113224 | 210598 | 211094 | BA_peak_925 |
| DS113225 | 23154 | 23694 | BA_peak_926 |
| DS113225 | 26913 | 27294 | BA_peak_927 |
| DS113225 | 29443 | 30219 | BA_peak_928 |
| DS113225 | 36991 | 37343 | BA_peak_929 |
| DS113225 | 49469 | 50104 | BA_peak_930 |
| DS113225 | 60841 | 65037 | BA_peak_931 |
| DS113225 | 67459 | 68152 | BA_peak_932 |
| DS113225 | 107621 | 108196 | BA_peak_933 |
| DS113225 | 137947 | 138737 | BA_peak_934 |
| DS113225 | 139991 | 143354 | BA_peak_935 |
| DS113225 | 197255 | 197864 | BA_peak_936 |
| DS113226 | 27697 | 29572 | BA_peak_937 |
| DS113226 | 35257 | 37570 | BA_peak_938 |
| DS113226 | 45593 | 47014 | BA_peak_939 |
| DS113226 | 98061 | 98511 | BA_peak_940 |
| DS113226 | 105205 | 108752 | BA_peak_941 |
| DS113226 | 149259 | 149595 | BA_peak_942 |
| DS113226 | 183792 | 184908 | BA_peak_943 |
| DS113227 | 10079 | 10488 | BA_peak_945 |
| DS113227 | 65372 | 66025 | BA_peak_946 |
| DS113227 | 67301 | 67766 | BA_peak_947 |
| DS113227 | 86079 | 88189 | BA_peak_948 |

| | | | |
|---|---|---|---|
| DS113227 | 108721 | 109117 | BA_peak_949 |
| DS113227 | 115583 | 115991 | BA_peak_950 |
| DS113227 | 116352 | 116819 | BA_peak_950 |
| DS113227 | 118426 | 118980 | BA_peak_951 |
| DS113227 | 136991 | 137438 | BA_peak_952 |
| DS113227 | 146929 | 147292 | BA_peak_953 |
| DS113227 | 169043 | 169887 | BA_peak_954 |
| DS113227 | 170953 | 174186 | BA_peak_955 |
| DS113228 | 12742 | 13274 | BA_peak_956 |
| DS113228 | 13783 | 14699 | BA_peak_957 |
| DS113228 | 67323 | 68081 | BA_peak_958 |
| DS113228 | 68544 | 69661 | BA_peak_959 |
| DS113228 | 73273 | 74227 | BA_peak_960 |
| DS113228 | 106805 | 108244 | BA_peak_961 |
| DS113228 | 171740 | 173823 | BA_peak_962 |
| DS113228 | 197975 | 199072 | BA_peak_963 |
| DS113228 | 203318 | 203779 | BA_peak_964 |
| DS113229 | 8218 | 8942 | BA_peak_965 |
| DS113229 | 36558 | 37309 | BA_peak_966 |
| DS113229 | 51008 | 51783 | BA_peak_967 |
| DS113229 | 67016 | 70222 | BA_peak_968 |
| DS113229 | 86057 | 86569 | BA_peak_969 |
| DS113229 | 87491 | 88417 | BA_peak_970 |
| DS113229 | 96929 | 97480 | BA_peak_971 |
| DS113229 | 116731 | 117219 | BA_peak_972 |
| DS113229 | 128955 | 132204 | BA_peak_973 |
| DS113229 | 135051 | 135473 | BA_peak_974 |
| DS113229 | 151148 | 151911 | BA_peak_975 |
| DS113229 | 153698 | 154376 | BA_peak_976 |
| DS113229 | 177212 | 177720 | BA_peak_977 |
| DS113229 | 195895 | 196265 | BA_peak_978 |
| DS113229 | 196860 | 197437 | BA_peak_979 |
| DS113229 | 199550 | 199908 | BA_peak_980 |
| DS113229 | 201022 | 202124 | BA_peak_981 |
| DS113230 | 15176 | 16016 | BA_peak_982 |
| DS113230 | 16438 | 17743 | BA_peak_983 |
| DS113230 | 26799 | 27352 | BA_peak_984 |
| DS113230 | 27906 | 28516 | BA_peak_985 |
| DS113230 | 31295 | 31979 | BA_peak_986 |
| DS113230 | 78864 | 79395 | BA_peak_987 |
| DS113230 | 80978 | 81655 | BA_peak_988 |
| DS113230 | 85921 | 86373 | BA_peak_989 |
| DS113230 | 101170 | 102145 | BA_peak_990 |
| DS113230 | 110607 | 111790 | BA_peak_991 |
| DS113230 | 129740 | 130178 | BA_peak_992 |
| DS113230 | 159690 | 160229 | BA_peak_993 |
| DS113230 | 161248 | 161865 | BA_peak_994 |

| | | | |
|---|---|---|---|
| DS113230 | 169575 | 169977 | BA_peak_995 |
| DS113230 | 194743 | 195383 | BA_peak_996 |
| DS113231 | 8762 | 9474 | BA_peak_997 |
| DS113231 | 56636 | 58743 | BA_peak_998 |
| DS113231 | 100148 | 100529 | BA_peak_999 |
| DS113231 | 131395 | 132595 | BA_peak_1000 |
| DS113231 | 135621 | 136306 | BA_peak_1001 |
| DS113231 | 144116 | 144697 | BA_peak_1002 |
| DS113231 | 171442 | 172195 | BA_peak_1003 |
| DS113231 | 172851 | 173413 | BA_peak_1004 |
| DS113231 | 174204 | 174728 | BA_peak_1005 |
| DS113232 | 41098 | 41941 | BA_peak_1006 |
| DS113232 | 45590 | 46638 | BA_peak_1007 |
| DS113232 | 90998 | 91561 | BA_peak_1008 |
| DS113232 | 117063 | 117383 | BA_peak_1009 |
| DS113232 | 143767 | 144598 | BA_peak_1010 |
| DS113232 | 154046 | 154779 | BA_peak_1011 |
| DS113232 | 157581 | 158288 | BA_peak_1012 |
| DS113232 | 190629 | 191145 | BA_peak_1013 |
| DS113232 | 202335 | 202813 | BA_peak_1014 |
| DS113233 | 86505 | 87331 | BA_peak_1015 |
| DS113233 | 95648 | 96477 | BA_peak_1016 |
| DS113233 | 125791 | 126190 | BA_peak_1017 |
| DS113233 | 187937 | 188611 | BA_peak_1018 |
| DS113234 | 5066 | 6412 | BA_peak_1019 |
| DS113234 | 81012 | 81533 | BA_peak_1020 |
| DS113234 | 89121 | 89505 | BA_peak_1021 |
| DS113234 | 92910 | 93514 | BA_peak_1022 |
| DS113234 | 133363 | 134217 | BA_peak_1023 |
| DS113234 | 135005 | 135510 | BA_peak_1024 |
| DS113234 | 140318 | 141465 | BA_peak_1025 |
| DS113234 | 142987 | 143417 | BA_peak_1026 |
| DS113234 | 143900 | 145780 | BA_peak_1027 |
| DS113234 | 154111 | 154700 | BA_peak_1028 |
| DS113234 | 160044 | 160560 | BA_peak_1029 |
| DS113234 | 165481 | 168504 | BA_peak_1030 |
| DS113234 | 182881 | 183309 | BA_peak_1031 |
| DS113234 | 184045 | 184693 | BA_peak_1032 |
| DS113234 | 186698 | 187299 | BA_peak_1033 |
| DS113234 | 188242 | 188612 | BA_peak_1034 |
| DS113235 | 30165 | 30727 | BA_peak_1036 |
| DS113235 | 40163 | 40566 | BA_peak_1037 |
| DS113235 | 81984 | 82562 | BA_peak_1038 |
| DS113235 | 88629 | 90065 | BA_peak_1039 |
| DS113235 | 94020 | 94492 | BA_peak_1040 |
| DS113235 | 96810 | 97559 | BA_peak_1041 |
| DS113235 | 133913 | 134683 | BA_peak_1043 |

| | | | |
|---|---|---|---|
| DS113235 | 143173 | 143923 | BA_peak_1044 |
| DS113235 | 163254 | 164344 | BA_peak_1045 |
| DS113235 | 199905 | 201087 | BA_peak_1046 |
| DS113236 | 13336 | 14139 | BA_peak_1047 |
| DS113236 | 14807 | 15416 | BA_peak_1048 |
| DS113236 | 47633 | 48552 | BA_peak_1049 |
| DS113236 | 71975 | 72345 | BA_peak_1050 |
| DS113236 | 78876 | 79239 | BA_peak_1051 |
| DS113236 | 80158 | 80847 | BA_peak_1052 |
| DS113236 | 89013 | 89632 | BA_peak_1053 |
| DS113236 | 91286 | 93488 | BA_peak_1054 |
| DS113236 | 97308 | 97729 | BA_peak_1055 |
| DS113236 | 103495 | 103862 | BA_peak_1056 |
| DS113236 | 104785 | 105105 | BA_peak_1057 |
| DS113236 | 105764 | 106107 | BA_peak_1058 |
| DS113236 | 135082 | 135476 | BA_peak_1059 |
| DS113236 | 160629 | 161147 | BA_peak_1060 |
| DS113236 | 161651 | 162062 | BA_peak_1061 |
| DS113236 | 163788 | 164308 | BA_peak_1062 |
| DS113236 | 196963 | 197672 | BA_peak_1063 |
| DS113236 | 200947 | 201691 | BA_peak_1064 |
| DS113237 | 16931 | 17463 | BA_peak_1065 |
| DS113237 | 38132 | 38979 | BA_peak_1066 |
| DS113237 | 40793 | 41873 | BA_peak_1067 |
| DS113237 | 48901 | 49226 | BA_peak_1068 |
| DS113237 | 51412 | 54568 | BA_peak_1069 |
| DS113237 | 56538 | 57650 | BA_peak_1070 |
| DS113237 | 63043 | 63530 | BA_peak_1071 |
| DS113237 | 64495 | 64876 | BA_peak_1072 |
| DS113237 | 89623 | 90217 | BA_peak_1073 |
| DS113237 | 139442 | 140110 | BA_peak_1074 |
| DS113238 | 34664 | 35305 | BA_peak_1075 |
| DS113238 | 36345 | 36753 | BA_peak_1076 |
| DS113238 | 39522 | 41075 | BA_peak_1077 |
| DS113238 | 41775 | 43469 | BA_peak_1078 |
| DS113238 | 50832 | 51247 | BA_peak_1079 |
| DS113238 | 52241 | 52765 | BA_peak_1080 |
| DS113238 | 58252 | 58564 | BA_peak_1081 |
| DS113238 | 65221 | 66119 | BA_peak_1082 |
| DS113238 | 90199 | 90648 | BA_peak_1083 |
| DS113238 | 122828 | 123237 | BA_peak_1084 |
| DS113238 | 163109 | 163495 | BA_peak_1086 |
| DS113238 | 177539 | 178276 | BA_peak_1087 |
| DS113238 | 184589 | 185201 | BA_peak_1088 |
| DS113238 | 193457 | 193808 | BA_peak_1089 |
| DS113239 | 29241 | 30203 | BA_peak_1090 |
| DS113239 | 43615 | 44494 | BA_peak_1091 |

| | | | |
|---|---|---|---|
| DS113239 | 45972 | 46771 | BA_peak_1092 |
| DS113239 | 48948 | 49378 | BA_peak_1093 |
| DS113239 | 49937 | 50354 | BA_peak_1094 |
| DS113239 | 59556 | 61045 | BA_peak_1095 |
| DS113239 | 66416 | 69752 | BA_peak_1096 |
| DS113239 | 95261 | 95641 | BA_peak_1098 |
| DS113239 | 121434 | 122083 | BA_peak_1099 |
| DS113239 | 124588 | 124891 | BA_peak_1100 |
| DS113239 | 131053 | 131479 | BA_peak_1101 |
| DS113239 | 139479 | 140024 | BA_peak_1102 |
| DS113239 | 161567 | 162437 | BA_peak_1103 |
| DS113239 | 174059 | 174594 | BA_peak_1104 |
| DS113239 | 175692 | 178908 | BA_peak_1105 |
| DS113240 | 44969 | 45342 | BA_peak_1106 |
| DS113240 | 59864 | 60183 | BA_peak_1107 |
| DS113240 | 90753 | 91229 | BA_peak_1109 |
| DS113240 | 123672 | 125622 | BA_peak_1111 |
| DS113240 | 140611 | 141347 | BA_peak_1112 |
| DS113240 | 147001 | 147327 | BA_peak_1113 |
| DS113240 | 161363 | 163479 | BA_peak_1114 |
| DS113240 | 174236 | 175859 | BA_peak_1115 |
| DS113240 | 176331 | 177414 | BA_peak_1116 |
| DS113241 | 22171 | 22690 | BA_peak_1117 |
| DS113241 | 30074 | 30661 | BA_peak_1118 |
| DS113241 | 72634 | 73594 | BA_peak_1119 |
| DS113241 | 80666 | 81046 | BA_peak_1120 |
| DS113241 | 104227 | 104831 | BA_peak_1121 |
| DS113241 | 106141 | 107280 | BA_peak_1122 |
| DS113242 | 5682 | 6465 | BA_peak_1123 |
| DS113242 | 9898 | 10735 | BA_peak_1124 |
| DS113242 | 14358 | 15109 | BA_peak_1125 |
| DS113242 | 15634 | 16570 | BA_peak_1126 |
| DS113242 | 22264 | 23358 | BA_peak_1127 |
| DS113242 | 36057 | 36492 | BA_peak_1128 |
| DS113242 | 38319 | 38930 | BA_peak_1129 |
| DS113242 | 49333 | 49873 | BA_peak_1130 |
| DS113242 | 54290 | 54593 | BA_peak_1131 |
| DS113242 | 65895 | 66507 | BA_peak_1132 |
| DS113242 | 71711 | 72152 | BA_peak_1133 |
| DS113242 | 86689 | 87253 | BA_peak_1135 |
| DS113242 | 99876 | 100298 | BA_peak_1137 |
| DS113242 | 119917 | 120397 | BA_peak_1138 |
| DS113242 | 130378 | 131068 | BA_peak_1139 |
| DS113242 | 131493 | 132266 | BA_peak_1140 |
| DS113242 | 133093 | 133639 | BA_peak_1141 |
| DS113242 | 147368 | 147966 | BA_peak_1142 |
| DS113242 | 148658 | 150488 | BA_peak_1143 |

| | | | |
|---|---|---|---|
| DS113242 | 151439 | 152711 | BA_peak_1144 |
| DS113242 | 174021 | 175333 | BA_peak_1145 |
| DS113243 | 2858 | 3545 | BA_peak_1146 |
| DS113243 | 4768 | 5197 | BA_peak_1147 |
| DS113243 | 5594 | 5916 | BA_peak_1148 |
| DS113243 | 75891 | 77042 | BA_peak_1149 |
| DS113243 | 87150 | 88236 | BA_peak_1150 |
| DS113243 | 131431 | 132004 | BA_peak_1151 |
| DS113243 | 157173 | 157577 | BA_peak_1152 |
| DS113243 | 158310 | 159017 | BA_peak_1153 |
| DS113243 | 171054 | 171642 | BA_peak_1154 |
| DS113243 | 175466 | 176381 | BA_peak_1155 |
| DS113243 | 189429 | 189838 | BA_peak_1156 |
| DS113244 | 77186 | 77905 | BA_peak_1158 |
| DS113244 | 97459 | 97886 | BA_peak_1159 |
| DS113244 | 99765 | 101041 | BA_peak_1160 |
| DS113244 | 101364 | 101756 | BA_peak_1161 |
| DS113244 | 103060 | 103648 | BA_peak_1162 |
| DS113244 | 104692 | 105444 | BA_peak_1163 |
| DS113244 | 119278 | 120135 | BA_peak_1164 |
| DS113244 | 161275 | 164630 | BA_peak_1165 |
| DS113244 | 168565 | 169455 | BA_peak_1166 |
| DS113244 | 197483 | 198187 | BA_peak_1167 |
| DS113245 | 23809 | 24504 | BA_peak_1168 |
| DS113245 | 39502 | 41053 | BA_peak_1169 |
| DS113245 | 52299 | 53978 | BA_peak_1170 |
| DS113245 | 67131 | 67961 | BA_peak_1171 |
| DS113245 | 71465 | 72271 | BA_peak_1172 |
| DS113245 | 79786 | 80465 | BA_peak_1173 |
| DS113245 | 135652 | 136154 | BA_peak_1175 |
| DS113245 | 144005 | 144827 | BA_peak_1176 |
| DS113245 | 154159 | 154554 | BA_peak_1177 |
| DS113245 | 155098 | 155549 | BA_peak_1178 |
| DS113245 | 157150 | 159016 | BA_peak_1179 |
| DS113245 | 170647 | 171076 | BA_peak_1180 |
| DS113245 | 175567 | 176123 | BA_peak_1182 |
| DS113245 | 191013 | 191323 | BA_peak_1183 |
| DS113245 | 195467 | 196377 | BA_peak_1184 |
| DS113246 | 22677 | 24139 | BA_peak_1185 |
| DS113246 | 42946 | 43689 | BA_peak_1186 |
| DS113246 | 44966 | 45727 | BA_peak_1187 |
| DS113246 | 75449 | 76853 | BA_peak_1188 |
| DS113246 | 100267 | 101093 | BA_peak_1189 |
| DS113246 | 102254 | 102713 | BA_peak_1190 |
| DS113246 | 104706 | 105199 | BA_peak_1191 |
| DS113246 | 133164 | 133577 | BA_peak_1192 |
| DS113246 | 134245 | 134700 | BA_peak_1193 |

| | | | |
|---|---|---|---|
| DS113246 | 184034 | 184496 | BA_peak_1194 |
| DS113246 | 188225 | 188839 | BA_peak_1195 |
| DS113247 | 10994 | 11757 | BA_peak_1196 |
| DS113247 | 49249 | 50155 | BA_peak_1197 |
| DS113247 | 52967 | 53282 | BA_peak_1198 |
| DS113247 | 69521 | 70995 | BA_peak_1199 |
| DS113247 | 81179 | 81457 | BA_peak_1200 |
| DS113247 | 134049 | 134465 | BA_peak_1201 |
| DS113247 | 139339 | 142770 | BA_peak_1202 |
| DS113247 | 149246 | 149836 | BA_peak_1203 |
| DS113248 | 49685 | 50041 | BA_peak_1204 |
| DS113248 | 50631 | 51285 | BA_peak_1205 |
| DS113248 | 67408 | 68026 | BA_peak_1206 |
| DS113248 | 72080 | 72980 | BA_peak_1207 |
| DS113248 | 109448 | 109792 | BA_peak_1208 |
| DS113248 | 121238 | 122236 | BA_peak_1209 |
| DS113248 | 122883 | 123419 | BA_peak_1210 |
| DS113248 | 161428 | 162487 | BA_peak_1211 |
| DS113248 | 175459 | 176798 | BA_peak_1212 |
| DS113248 | 177527 | 178120 | BA_peak_1213 |
| DS113248 | 179036 | 180071 | BA_peak_1214 |
| DS113249 | 20741 | 21175 | BA_peak_1216 |
| DS113249 | 35026 | 35932 | BA_peak_1217 |
| DS113249 | 40367 | 41101 | BA_peak_1218 |
| DS113249 | 111542 | 112119 | BA_peak_1219 |
| DS113249 | 113853 | 114626 | BA_peak_1220 |
| DS113249 | 121598 | 122048 | BA_peak_1221 |
| DS113249 | 127208 | 127756 | BA_peak_1222 |
| DS113249 | 131703 | 132252 | BA_peak_1223 |
| DS113249 | 135792 | 136742 | BA_peak_1224 |
| DS113249 | 150215 | 151310 | BA_peak_1225 |
| DS113250 | 32843 | 33360 | BA_peak_1226 |
| DS113250 | 34427 | 34982 | BA_peak_1227 |
| DS113250 | 41981 | 42480 | BA_peak_1228 |
| DS113250 | 62238 | 62869 | BA_peak_1229 |
| DS113250 | 63898 | 64333 | BA_peak_1230 |
| DS113250 | 115547 | 116414 | BA_peak_1231 |
| DS113250 | 127772 | 128127 | BA_peak_1232 |
| DS113250 | 156272 | 157377 | BA_peak_1233 |
| DS113250 | 186312 | 187338 | BA_peak_1234 |
| DS113250 | 188322 | 189095 | BA_peak_1235 |
| DS113250 | 189649 | 190254 | BA_peak_1236 |
| DS113251 | 23746 | 24146 | BA_peak_1237 |
| DS113251 | 36233 | 38223 | BA_peak_1238 |
| DS113251 | 84161 | 85012 | BA_peak_1239 |
| DS113251 | 86177 | 86896 | BA_peak_1240 |
| DS113251 | 95967 | 96486 | BA_peak_1241 |

| | | | |
|---|---|---|---|
| DS113251 | 114244 | 114596 | BA_peak_1242 |
| DS113251 | 181664 | 182526 | BA_peak_1243 |
| DS113251 | 183250 | 183648 | BA_peak_1244 |
| DS113252 | 21423 | 21783 | BA_peak_1245 |
| DS113252 | 37429 | 37793 | BA_peak_1246 |
| DS113252 | 46277 | 47214 | BA_peak_1247 |
| DS113252 | 50825 | 51372 | BA_peak_1248 |
| DS113252 | 52032 | 52749 | BA_peak_1249 |
| DS113252 | 53675 | 54374 | BA_peak_1250 |
| DS113252 | 64196 | 64504 | BA_peak_1251 |
| DS113252 | 97952 | 98446 | BA_peak_1252 |
| DS113252 | 102149 | 102994 | BA_peak_1253 |
| DS113252 | 125159 | 125560 | BA_peak_1254 |
| DS113252 | 135287 | 136257 | BA_peak_1255 |
| DS113252 | 180283 | 181161 | BA_peak_1256 |
| DS113253 | 1321 | 1792 | BA_peak_1257 |
| DS113253 | 15287 | 19689 | BA_peak_1258 |
| DS113253 | 20013 | 20417 | BA_peak_1259 |
| DS113253 | 20804 | 21824 | BA_peak_1260 |
| DS113253 | 25459 | 26266 | BA_peak_1261 |
| DS113253 | 26623 | 26992 | BA_peak_1262 |
| DS113253 | 63584 | 64236 | BA_peak_1263 |
| DS113253 | 86255 | 86693 | BA_peak_1264 |
| DS113253 | 87754 | 88184 | BA_peak_1265 |
| DS113253 | 88490 | 88824 | BA_peak_1266 |
| DS113253 | 90834 | 91241 | BA_peak_1267 |
| DS113253 | 92152 | 92455 | BA_peak_1268 |
| DS113253 | 155843 | 156150 | BA_peak_1269 |
| DS113253 | 164658 | 165054 | BA_peak_1270 |
| DS113253 | 180601 | 181113 | BA_peak_1271 |
| DS113254 | 2124 | 3005 | BA_peak_1272 |
| DS113254 | 30436 | 31062 | BA_peak_1273 |
| DS113254 | 32216 | 32627 | BA_peak_1274 |
| DS113254 | 41623 | 42218 | BA_peak_1275 |
| DS113254 | 64250 | 64945 | BA_peak_1276 |
| DS113254 | 92559 | 94632 | BA_peak_1277 |
| DS113254 | 112373 | 112899 | BA_peak_1278 |
| DS113254 | 121571 | 122336 | BA_peak_1279 |
| DS113254 | 167864 | 168242 | BA_peak_1280 |
| DS113254 | 173841 | 175198 | BA_peak_1281 |
| DS113255 | 16467 | 17376 | BA_peak_1282 |
| DS113255 | 17820 | 18419 | BA_peak_1283 |
| DS113255 | 18802 | 19924 | BA_peak_1284 |
| DS113255 | 21691 | 22349 | BA_peak_1285 |
| DS113255 | 65166 | 65919 | BA_peak_1286 |
| DS113255 | 66985 | 67937 | BA_peak_1287 |
| DS113255 | 85721 | 86688 | BA_peak_1288 |

| | | | |
|---|---|---|---|
| DS113255 | 119843 | 120423 | BA_peak_1289 |
| DS113256 | 67253 | 67737 | BA_peak_1290 |
| DS113256 | 78735 | 79361 | BA_peak_1291 |
| DS113256 | 85898 | 86650 | BA_peak_1292 |
| DS113256 | 137243 | 137642 | BA_peak_1293 |
| DS113256 | 142806 | 143594 | BA_peak_1294 |
| DS113256 | 147644 | 148005 | BA_peak_1295 |
| DS113256 | 154410 | 155732 | BA_peak_1296 |
| DS113256 | 156506 | 157152 | BA_peak_1297 |
| DS113256 | 182660 | 183030 | BA_peak_1298 |
| DS113256 | 183629 | 184122 | BA_peak_1299 |
| DS113256 | 185941 | 186615 | BA_peak_1300 |
| DS113257 | 241 | 1516 | BA_peak_1301 |
| DS113257 | 51488 | 51923 | BA_peak_1302 |
| DS113257 | 54015 | 54927 | BA_peak_1303 |
| DS113257 | 74416 | 75027 | BA_peak_1304 |
| DS113257 | 80263 | 80609 | BA_peak_1305 |
| DS113257 | 99455 | 100309 | BA_peak_1306 |
| DS113257 | 110828 | 111396 | BA_peak_1307 |
| DS113257 | 115874 | 116262 | BA_peak_1308 |
| DS113257 | 138551 | 139384 | BA_peak_1309 |
| DS113257 | 149803 | 150469 | BA_peak_1310 |
| DS113257 | 181120 | 182542 | BA_peak_1311 |
| DS113258 | 6953 | 7647 | BA_peak_1312 |
| DS113258 | 8935 | 9343 | BA_peak_1313 |
| DS113258 | 52701 | 53311 | BA_peak_1314 |
| DS113258 | 70304 | 70660 | BA_peak_1315 |
| DS113258 | 89212 | 89720 | BA_peak_1316 |
| DS113258 | 135633 | 136478 | BA_peak_1317 |
| DS113258 | 141650 | 142351 | BA_peak_1318 |
| DS113258 | 171880 | 172303 | BA_peak_1319 |
| DS113259 | 16703 | 17368 | BA_peak_1320 |
| DS113259 | 53230 | 54073 | BA_peak_1321 |
| DS113259 | 70530 | 70938 | BA_peak_1322 |
| DS113259 | 75196 | 78286 | BA_peak_1323 |
| DS113259 | 109060 | 112371 | BA_peak_1324 |
| DS113259 | 114926 | 115574 | BA_peak_1325 |
| DS113259 | 117972 | 119354 | BA_peak_1326 |
| DS113259 | 126504 | 127201 | BA_peak_1327 |
| DS113259 | 128360 | 129619 | BA_peak_1328 |
| DS113259 | 130973 | 132129 | BA_peak_1329 |
| DS113259 | 133168 | 134361 | BA_peak_1330 |
| DS113259 | 139718 | 141607 | BA_peak_1331 |
| DS113259 | 144773 | 145548 | BA_peak_1332 |
| DS113259 | 146090 | 148002 | BA_peak_1333 |
| DS113259 | 169126 | 169483 | BA_peak_1334 |
| DS113259 | 181590 | 182314 | BA_peak_1336 |

| | | | |
|---|---|---|---|
| DS113260 | 24272 | 24609 | BA_peak_1337 |
| DS113260 | 37678 | 38113 | BA_peak_1338 |
| DS113260 | 42747 | 43493 | BA_peak_1339 |
| DS113260 | 61929 | 64814 | BA_peak_1340 |
| DS113260 | 120291 | 120715 | BA_peak_1341 |
| DS113260 | 140009 | 140904 | BA_peak_1342 |
| DS113260 | 182243 | 183118 | BA_peak_1343 |
| DS113261 | 7279 | 7912 | BA_peak_1344 |
| DS113261 | 12414 | 14268 | BA_peak_1345 |
| DS113261 | 25993 | 26776 | BA_peak_1346 |
| DS113261 | 43841 | 44183 | BA_peak_1347 |
| DS113261 | 58918 | 59317 | BA_peak_1348 |
| DS113261 | 74924 | 76740 | BA_peak_1349 |
| DS113261 | 102686 | 103939 | BA_peak_1351 |
| DS113261 | 158604 | 158995 | BA_peak_1352 |
| DS113262 | 8736 | 10744 | BA_peak_1353 |
| DS113262 | 13748 | 14392 | BA_peak_1354 |
| DS113262 | 15055 | 15984 | BA_peak_1355 |
| DS113262 | 91291 | 92411 | BA_peak_1356 |
| DS113262 | 101669 | 105542 | BA_peak_1357 |
| DS113262 | 113144 | 115348 | BA_peak_1358 |
| DS113262 | 144476 | 144924 | BA_peak_1359 |
| DS113262 | 167269 | 167681 | BA_peak_1360 |
| DS113262 | 176576 | 177546 | BA_peak_1361 |
| DS113263 | 9284 | 9982 | BA_peak_1362 |
| DS113263 | 20588 | 21489 | BA_peak_1363 |
| DS113263 | 70011 | 71364 | BA_peak_1364 |
| DS113263 | 81908 | 82719 | BA_peak_1365 |
| DS113263 | 97692 | 99767 | BA_peak_1366 |
| DS113263 | 119710 | 120107 | BA_peak_1367 |
| DS113263 | 128335 | 129971 | BA_peak_1368 |
| DS113263 | 130964 | 131266 | BA_peak_1369 |
| DS113264 | 11706 | 12125 | BA_peak_1370 |
| DS113264 | 71250 | 74551 | BA_peak_1371 |
| DS113264 | 126109 | 126915 | BA_peak_1372 |
| DS113264 | 171107 | 172473 | BA_peak_1373 |
| DS113264 | 176485 | 177107 | BA_peak_1374 |
| DS113265 | 27091 | 27447 | BA_peak_1375 |
| DS113265 | 27969 | 29755 | BA_peak_1376 |
| DS113265 | 33557 | 34612 | BA_peak_1377 |
| DS113265 | 31588 | 33177 | BA_peak_1377 |
| DS113265 | 35284 | 35865 | BA_peak_1378 |
| DS113265 | 40554 | 41746 | BA_peak_1379 |
| DS113265 | 42973 | 43958 | BA_peak_1380 |
| DS113265 | 45256 | 45992 | BA_peak_1381 |
| DS113265 | 47316 | 47913 | BA_peak_1382 |
| DS113265 | 48257 | 50111 | BA_peak_1383 |

| | | | |
|---|---|---|---|
| DS113265 | 51602 | 52802 | BA_peak_1384 |
| DS113265 | 62093 | 62593 | BA_peak_1385 |
| DS113265 | 64214 | 64763 | BA_peak_1386 |
| DS113265 | 72627 | 73376 | BA_peak_1387 |
| DS113265 | 105316 | 106132 | BA_peak_1388 |
| DS113265 | 106588 | 108222 | BA_peak_1389 |
| DS113265 | 119982 | 121176 | BA_peak_1390 |
| DS113265 | 121502 | 121823 | BA_peak_1391 |
| DS113265 | 157412 | 158115 | BA_peak_1392 |
| DS113265 | 169668 | 170250 | BA_peak_1393 |
| DS113265 | 171166 | 172473 | BA_peak_1394 |
| DS113266 | 12115 | 13044 | BA_peak_1395 |
| DS113266 | 31065 | 31822 | BA_peak_1396 |
| DS113266 | 93173 | 93507 | BA_peak_1397 |
| DS113266 | 109481 | 109823 | BA_peak_1398 |
| DS113266 | 112331 | 112888 | BA_peak_1399 |
| DS113266 | 138730 | 139588 | BA_peak_1400 |
| DS113266 | 157645 | 158304 | BA_peak_1401 |
| DS113266 | 163471 | 164048 | BA_peak_1402 |
| DS113266 | 172101 | 172718 | BA_peak_1403 |
| DS113267 | 10696 | 11990 | BA_peak_1404 |
| DS113267 | 21023 | 21510 | BA_peak_1405 |
| DS113267 | 45405 | 46392 | BA_peak_1406 |
| DS113267 | 70823 | 71607 | BA_peak_1407 |
| DS113267 | 72601 | 73528 | BA_peak_1408 |
| DS113267 | 84001 | 84411 | BA_peak_1409 |
| DS113267 | 99434 | 100329 | BA_peak_1410 |
| DS113267 | 110864 | 112305 | BA_peak_1411 |
| DS113267 | 112983 | 114026 | BA_peak_1412 |
| DS113267 | 117281 | 118683 | BA_peak_1413 |
| DS113267 | 119070 | 119765 | BA_peak_1414 |
| DS113267 | 143163 | 144932 | BA_peak_1415 |
| DS113267 | 153952 | 154763 | BA_peak_1417 |
| DS113267 | 170671 | 171337 | BA_peak_1418 |
| DS113267 | 171735 | 172467 | BA_peak_1419 |
| DS113267 | 173400 | 175119 | BA_peak_1420 |
| DS113267 | 176062 | 176478 | BA_peak_1421 |
| DS113268 | 10032 | 10646 | BA_peak_1422 |
| DS113268 | 37184 | 38034 | BA_peak_1423 |
| DS113268 | 39877 | 41752 | BA_peak_1424 |
| DS113268 | 52495 | 52835 | BA_peak_1425 |
| DS113268 | 59398 | 59929 | BA_peak_1426 |
| DS113268 | 85069 | 85382 | BA_peak_1427 |
| DS113268 | 95891 | 96276 | BA_peak_1428 |
| DS113268 | 113955 | 114359 | BA_peak_1429 |
| DS113268 | 114941 | 115836 | BA_peak_1430 |
| DS113268 | 116424 | 116861 | BA_peak_1431 |

| | | | |
|---|---|---|---|
| DS113268 | 118235 | 119233 | BA_peak_1432 |
| DS113268 | 132184 | 133318 | BA_peak_1433 |
| DS113268 | 133725 | 135123 | BA_peak_1434 |
| DS113268 | 147044 | 147770 | BA_peak_1436 |
| DS113269 | 3047 | 3739 | BA_peak_1438 |
| DS113269 | 38583 | 40218 | BA_peak_1439 |
| DS113269 | 42987 | 45452 | BA_peak_1440 |
| DS113269 | 64011 | 64426 | BA_peak_1441 |
| DS113269 | 114919 | 115753 | BA_peak_1442 |
| DS113269 | 147703 | 148245 | BA_peak_1443 |
| DS113269 | 149768 | 150303 | BA_peak_1444 |
| DS113269 | 165030 | 165499 | BA_peak_1445 |
| DS113270 | 15415 | 16150 | BA_peak_1446 |
| DS113270 | 44273 | 44675 | BA_peak_1447 |
| DS113270 | 145631 | 145986 | BA_peak_1448 |
| DS113270 | 146506 | 146853 | BA_peak_1449 |
| DS113271 | 9725 | 10219 | BA_peak_1450 |
| DS113271 | 21035 | 21437 | BA_peak_1451 |
| DS113271 | 27987 | 28829 | BA_peak_1452 |
| DS113271 | 31626 | 32137 | BA_peak_1453 |
| DS113271 | 44583 | 44997 | BA_peak_1454 |
| DS113271 | 45719 | 46400 | BA_peak_1455 |
| DS113271 | 46936 | 47426 | BA_peak_1456 |
| DS113271 | 94936 | 95804 | BA_peak_1457 |
| DS113271 | 130552 | 131042 | BA_peak_1458 |
| DS113271 | 132276 | 132928 | BA_peak_1459 |
| DS113271 | 133303 | 134011 | BA_peak_1460 |
| DS113271 | 140509 | 141474 | BA_peak_1461 |
| DS113271 | 145135 | 147101 | BA_peak_1462 |
| DS113271 | 165871 | 166647 | BA_peak_1463 |
| DS113271 | 170482 | 170803 | BA_peak_1464 |
| DS113272 | 18465 | 20388 | BA_peak_1465 |
| DS113272 | 33872 | 34639 | BA_peak_1466 |
| DS113272 | 48441 | 49507 | BA_peak_1467 |
| DS113272 | 50225 | 50824 | BA_peak_1468 |
| DS113272 | 58442 | 59786 | BA_peak_1469 |
| DS113272 | 80027 | 80766 | BA_peak_1470 |
| DS113272 | 110604 | 111259 | BA_peak_1471 |
| DS113272 | 117025 | 117612 | BA_peak_1472 |
| DS113272 | 128239 | 129022 | BA_peak_1473 |
| DS113272 | 129411 | 129932 | BA_peak_1474 |
| DS113272 | 138491 | 139157 | BA_peak_1475 |
| DS113272 | 140945 | 142518 | BA_peak_1476 |
| DS113272 | 142986 | 143705 | BA_peak_1477 |
| DS113272 | 144655 | 145288 | BA_peak_1478 |
| DS113272 | 164276 | 165084 | BA_peak_1479 |
| DS113272 | 165697 | 167798 | BA_peak_1480 |

| | | | |
|---|---|---|---|
| DS113272 | 169960 | 170289 | BA_peak_1481 |
| DS113273 | 7820 | 8450 | BA_peak_1482 |
| DS113273 | 21794 | 22333 | BA_peak_1483 |
| DS113273 | 22774 | 23285 | BA_peak_1484 |
| DS113273 | 37754 | 38573 | BA_peak_1485 |
| DS113273 | 39492 | 40432 | BA_peak_1486 |
| DS113273 | 44763 | 45115 | BA_peak_1487 |
| DS113273 | 53325 | 53919 | BA_peak_1488 |
| DS113273 | 55057 | 56414 | BA_peak_1489 |
| DS113273 | 72787 | 76835 | BA_peak_1490 |
| DS113273 | 164357 | 164996 | BA_peak_1491 |
| DS113274 | 771 | 1212 | BA_peak_1492 |
| DS113274 | 2979 | 3922 | BA_peak_1493 |
| DS113274 | 15574 | 16322 | BA_peak_1494 |
| DS113274 | 42395 | 42773 | BA_peak_1495 |
| DS113274 | 50973 | 53204 | BA_peak_1496 |
| DS113274 | 54091 | 54409 | BA_peak_1497 |
| DS113274 | 79773 | 80738 | BA_peak_1498 |
| DS113274 | 92042 | 92849 | BA_peak_1499 |
| DS113274 | 101738 | 103144 | BA_peak_1500 |
| DS113274 | 113501 | 115010 | BA_peak_1501 |
| DS113274 | 117575 | 118167 | BA_peak_1502 |
| DS113274 | 120327 | 121398 | BA_peak_1503 |
| DS113274 | 139281 | 139648 | BA_peak_1504 |
| DS113274 | 148423 | 148763 | BA_peak_1505 |
| DS113274 | 163879 | 164206 | BA_peak_1506 |
| DS113274 | 164707 | 165034 | BA_peak_1507 |
| DS113275 | 67 | 402 | BA_peak_1508 |
| DS113275 | 6630 | 7587 | BA_peak_1509 |
| DS113275 | 12974 | 13834 | BA_peak_1510 |
| DS113275 | 37443 | 38261 | BA_peak_1511 |
| DS113275 | 123900 | 124846 | BA_peak_1512 |
| DS113275 | 141937 | 142705 | BA_peak_1513 |
| DS113275 | 161495 | 162568 | BA_peak_1514 |
| DS113275 | 167849 | 168152 | BA_peak_1515 |
| DS113276 | 9416 | 10187 | BA_peak_1516 |
| DS113276 | 34109 | 34446 | BA_peak_1517 |
| DS113276 | 38529 | 39162 | BA_peak_1518 |
| DS113276 | 51559 | 52540 | BA_peak_1519 |
| DS113276 | 131735 | 133004 | BA_peak_1520 |
| DS113276 | 133674 | 135234 | BA_peak_1521 |
| DS113277 | 5097 | 5403 | BA_peak_1522 |
| DS113277 | 52684 | 53490 | BA_peak_1523 |
| DS113277 | 55983 | 56753 | BA_peak_1524 |
| DS113277 | 76904 | 77535 | BA_peak_1525 |
| DS113277 | 92423 | 92851 | BA_peak_1526 |
| DS113277 | 112816 | 113764 | BA_peak_1527 |

| | | | |
|---|---|---|---|
| DS113278 | 62063 | 62571 | BA_peak_1528 |
| DS113278 | 90005 | 90618 | BA_peak_1529 |
| DS113278 | 124669 | 125211 | BA_peak_1531 |
| DS113278 | 125925 | 126740 | BA_peak_1532 |
| DS113279 | 10882 | 11273 | BA_peak_1534 |
| DS113279 | 20739 | 21333 | BA_peak_1535 |
| DS113279 | 23153 | 23980 | BA_peak_1536 |
| DS113279 | 26938 | 28896 | BA_peak_1537 |
| DS113279 | 47587 | 49522 | BA_peak_1538 |
| DS113279 | 63441 | 64290 | BA_peak_1539 |
| DS113279 | 66307 | 67103 | BA_peak_1540 |
| DS113279 | 71729 | 72035 | BA_peak_1541 |
| DS113279 | 72661 | 74520 | BA_peak_1542 |
| DS113279 | 77849 | 78495 | BA_peak_1543 |
| DS113279 | 83212 | 84455 | BA_peak_1544 |
| DS113279 | 86097 | 87660 | BA_peak_1545 |
| DS113279 | 93753 | 94016 | BA_peak_1546 |
| DS113279 | 94514 | 95188 | BA_peak_1547 |
| DS113279 | 98099 | 98595 | BA_peak_1548 |
| DS113279 | 100292 | 101235 | BA_peak_1549 |
| DS113279 | 104063 | 104656 | BA_peak_1550 |
| DS113279 | 143500 | 143824 | BA_peak_1551 |
| DS113279 | 160681 | 161048 | BA_peak_1552 |
| DS113279 | 166877 | 168030 | BA_peak_1553 |
| DS113280 | 31053 | 31424 | BA_peak_1554 |
| DS113280 | 47293 | 48202 | BA_peak_1555 |
| DS113280 | 57360 | 61153 | BA_peak_1556 |
| DS113280 | 81969 | 84414 | BA_peak_1557 |
| DS113280 | 88576 | 88936 | BA_peak_1558 |
| DS113280 | 90181 | 90739 | BA_peak_1559 |
| DS113280 | 91221 | 92327 | BA_peak_1560 |
| DS113280 | 103241 | 105249 | BA_peak_1561 |
| DS113280 | 105593 | 106472 | BA_peak_1561 |
| DS113280 | 107515 | 108415 | BA_peak_1562 |
| DS113280 | 119113 | 119873 | BA_peak_1563 |
| DS113280 | 120616 | 121305 | BA_peak_1564 |
| DS113280 | 129968 | 130463 | BA_peak_1565 |
| DS113280 | 135671 | 136362 | BA_peak_1566 |
| DS113280 | 141245 | 142073 | BA_peak_1567 |
| DS113280 | 144753 | 145276 | BA_peak_1568 |
| DS113281 | 3513 | 4417 | BA_peak_1569 |
| DS113281 | 10320 | 10847 | BA_peak_1570 |
| DS113281 | 49166 | 49818 | BA_peak_1572 |
| DS113281 | 50168 | 51121 | BA_peak_1573 |
| DS113281 | 75301 | 76183 | BA_peak_1574 |
| DS113281 | 93519 | 93854 | BA_peak_1575 |
| DS113281 | 94290 | 94657 | BA_peak_1576 |

| | | | |
|---|---|---|---|
| DS113281 | 106072 | 110338 | BA_peak_1577 |
| DS113281 | 135002 | 136120 | BA_peak_1578 |
| DS113281 | 136501 | 137262 | BA_peak_1579 |
| DS113282 | 5118 | 5578 | BA_peak_1580 |
| DS113282 | 10299 | 11036 | BA_peak_1581 |
| DS113282 | 16631 | 17604 | BA_peak_1582 |
| DS113282 | 35124 | 35831 | BA_peak_1583 |
| DS113282 | 43531 | 43987 | BA_peak_1584 |
| DS113282 | 79566 | 80156 | BA_peak_1585 |
| DS113282 | 84941 | 86126 | BA_peak_1586 |
| DS113282 | 116756 | 117181 | BA_peak_1587 |
| DS113282 | 122293 | 124559 | BA_peak_1588 |
| DS113282 | 124959 | 125694 | BA_peak_1589 |
| DS113283 | 23953 | 24762 | BA_peak_1590 |
| DS113283 | 49649 | 50182 | BA_peak_1591 |
| DS113283 | 50753 | 51262 | BA_peak_1592 |
| DS113283 | 65051 | 65436 | BA_peak_1593 |
| DS113283 | 69553 | 69894 | BA_peak_1594 |
| DS113283 | 136673 | 137194 | BA_peak_1595 |
| DS113283 | 145973 | 146765 | BA_peak_1596 |
| DS113284 | 25292 | 25895 | BA_peak_1597 |
| DS113284 | 52703 | 53189 | BA_peak_1598 |
| DS113284 | 66615 | 67188 | BA_peak_1599 |
| DS113284 | 69055 | 69670 | BA_peak_1600 |
| DS113284 | 91191 | 91967 | BA_peak_1601 |
| DS113284 | 95684 | 96873 | BA_peak_1602 |
| DS113284 | 107660 | 108493 | BA_peak_1603 |
| DS113284 | 146462 | 147233 | BA_peak_1604 |
| DS113285 | 37031 | 37775 | BA_peak_1605 |
| DS113285 | 64216 | 64571 | BA_peak_1607 |
| DS113285 | 102469 | 102840 | BA_peak_1608 |
| DS113285 | 122436 | 122771 | BA_peak_1611 |
| DS113285 | 147616 | 147944 | BA_peak_1612 |
| DS113286 | 53846 | 54561 | BA_peak_1613 |
| DS113286 | 75677 | 76572 | BA_peak_1614 |
| DS113286 | 85847 | 86293 | BA_peak_1615 |
| DS113286 | 132806 | 133428 | BA_peak_1616 |
| DS113286 | 142682 | 143155 | BA_peak_1617 |
| DS113286 | 155402 | 156450 | BA_peak_1618 |
| DS113287 | 9344 | 10220 | BA_peak_1619 |
| DS113287 | 16395 | 17018 | BA_peak_1620 |
| DS113287 | 38204 | 38586 | BA_peak_1621 |
| DS113287 | 46785 | 47704 | BA_peak_1622 |
| DS113287 | 76477 | 77528 | BA_peak_1624 |
| DS113287 | 106948 | 107279 | BA_peak_1625 |
| DS113287 | 132272 | 132952 | BA_peak_1627 |
| DS113287 | 141468 | 142546 | BA_peak_1628 |

| | | | |
|---|---|---|---|
| DS113287 | 144641 | 145232 | BA_peak_1629 |
| DS113287 | 152413 | 152842 | BA_peak_1630 |
| DS113288 | 78 | 967 | BA_peak_1631 |
| DS113288 | 4871 | 6661 | BA_peak_1632 |
| DS113288 | 16739 | 17957 | BA_peak_1633 |
| DS113288 | 50163 | 53794 | BA_peak_1634 |
| DS113288 | 54571 | 55092 | BA_peak_1635 |
| DS113288 | 78305 | 80705 | BA_peak_1636 |
| DS113288 | 85405 | 88609 | BA_peak_1637 |
| DS113288 | 107251 | 108783 | BA_peak_1638 |
| DS113288 | 135165 | 135829 | BA_peak_1639 |
| DS113288 | 144587 | 145815 | BA_peak_1640 |
| DS113288 | 150425 | 150783 | BA_peak_1641 |
| DS113289 | 5365 | 5922 | BA_peak_1642 |
| DS113289 | 26312 | 27136 | BA_peak_1643 |
| DS113289 | 29888 | 30689 | BA_peak_1644 |
| DS113289 | 60518 | 62537 | BA_peak_1645 |
| DS113289 | 66348 | 66843 | BA_peak_1646 |
| DS113289 | 88506 | 89645 | BA_peak_1647 |
| DS113289 | 90206 | 90632 | BA_peak_1648 |
| DS113289 | 138467 | 139272 | BA_peak_1650 |
| DS113289 | 156204 | 158155 | BA_peak_1651 |
| DS113290 | 1467 | 2798 | BA_peak_1652 |
| DS113290 | 23766 | 24084 | BA_peak_1654 |
| DS113290 | 33646 | 34033 | BA_peak_1655 |
| DS113290 | 53725 | 54097 | BA_peak_1656 |
| DS113290 | 70543 | 71010 | BA_peak_1658 |
| DS113290 | 87898 | 88777 | BA_peak_1659 |
| DS113290 | 114788 | 115451 | BA_peak_1660 |
| DS113290 | 115767 | 117063 | BA_peak_1661 |
| DS113290 | 117407 | 118588 | BA_peak_1662 |
| DS113290 | 123817 | 126205 | BA_peak_1663 |
| DS113290 | 128859 | 129647 | BA_peak_1664 |
| DS113290 | 133835 | 134614 | BA_peak_1666 |
| DS113290 | 136716 | 137688 | BA_peak_1668 |
| DS113291 | 16312 | 17575 | BA_peak_1669 |
| DS113291 | 19285 | 21065 | BA_peak_1670 |
| DS113291 | 22307 | 22733 | BA_peak_1671 |
| DS113291 | 70433 | 71860 | BA_peak_1672 |
| DS113291 | 74181 | 74653 | BA_peak_1673 |
| DS113291 | 75625 | 76099 | BA_peak_1674 |
| DS113291 | 81043 | 81431 | BA_peak_1675 |
| DS113291 | 92178 | 93361 | BA_peak_1676 |
| DS113291 | 95938 | 96244 | BA_peak_1677 |
| DS113291 | 98432 | 98783 | BA_peak_1678 |
| DS113291 | 123617 | 124369 | BA_peak_1679 |
| DS113291 | 125370 | 126023 | BA_peak_1680 |

| | | | |
|---|---|---|---|
| DS113291 | 127514 | 128041 | BA_peak_1681 |
| DS113291 | 128598 | 129367 | BA_peak_1682 |
| DS113291 | 140465 | 143609 | BA_peak_1683 |
| DS113291 | 146822 | 147350 | BA_peak_1685 |
| DS113292 | 42535 | 43291 | BA_peak_1686 |
| DS113292 | 44082 | 44950 | BA_peak_1687 |
| DS113292 | 123595 | 124007 | BA_peak_1688 |
| DS113293 | 13332 | 14073 | BA_peak_1689 |
| DS113293 | 17710 | 18257 | BA_peak_1690 |
| DS113293 | 53437 | 54408 | BA_peak_1691 |
| DS113293 | 60324 | 60835 | BA_peak_1692 |
| DS113293 | 78374 | 78704 | BA_peak_1693 |
| DS113293 | 142842 | 143227 | BA_peak_1695 |
| DS113293 | 144867 | 145670 | BA_peak_1696 |
| DS113293 | 154634 | 155754 | BA_peak_1697 |
| DS113294 | 76199 | 76811 | BA_peak_1698 |
| DS113294 | 78329 | 79194 | BA_peak_1699 |
| DS113294 | 84813 | 85180 | BA_peak_1700 |
| DS113294 | 86936 | 88159 | BA_peak_1701 |
| DS113294 | 89571 | 90482 | BA_peak_1702 |
| DS113294 | 93876 | 94617 | BA_peak_1703 |
| DS113294 | 108758 | 109109 | BA_peak_1704 |
| DS113294 | 144447 | 145528 | BA_peak_1705 |
| DS113295 | 13910 | 15383 | BA_peak_1706 |
| DS113295 | 16633 | 17182 | BA_peak_1707 |
| DS113295 | 35582 | 36051 | BA_peak_1708 |
| DS113295 | 39238 | 40086 | BA_peak_1710 |
| DS113295 | 43081 | 43593 | BA_peak_1711 |
| DS113295 | 53570 | 54141 | BA_peak_1712 |
| DS113295 | 54892 | 55390 | BA_peak_1713 |
| DS113295 | 120408 | 121906 | BA_peak_1714 |
| DS113296 | 6262 | 6787 | BA_peak_1715 |
| DS113296 | 8509 | 8904 | BA_peak_1716 |
| DS113296 | 29257 | 29655 | BA_peak_1718 |
| DS113296 | 34094 | 34548 | BA_peak_1719 |
| DS113296 | 48215 | 48695 | BA_peak_1720 |
| DS113296 | 49498 | 50170 | BA_peak_1721 |
| DS113296 | 53947 | 54833 | BA_peak_1722 |
| DS113296 | 77497 | 78136 | BA_peak_1723 |
| DS113296 | 93754 | 94119 | BA_peak_1724 |
| DS113296 | 99186 | 99732 | BA_peak_1725 |
| DS113296 | 117524 | 118476 | BA_peak_1726 |
| DS113296 | 121926 | 122734 | BA_peak_1727 |
| DS113296 | 151370 | 152370 | BA_peak_1728 |
| DS113297 | 360 | 1008 | BA_peak_1729 |
| DS113297 | 2674 | 4111 | BA_peak_1730 |
| DS113297 | 4429 | 5787 | BA_peak_1731 |

| | | | |
|---|---|---|---|
| DS113297 | 15322 | 15869 | BA_peak_1732 |
| DS113297 | 50723 | 51073 | BA_peak_1733 |
| DS113297 | 85357 | 85731 | BA_peak_1734 |
| DS113297 | 105294 | 107390 | BA_peak_1735 |
| DS113297 | 113428 | 113809 | BA_peak_1736 |
| DS113297 | 126399 | 127175 | BA_peak_1737 |
| DS113297 | 130138 | 133499 | BA_peak_1738 |
| DS113297 | 146684 | 147594 | BA_peak_1739 |
| DS113297 | 149869 | 150295 | BA_peak_1740 |
| DS113298 | 90 | 488 | BA_peak_1741 |
| DS113298 | 1218 | 2500 | BA_peak_1742 |
| DS113298 | 2935 | 3523 | BA_peak_1743 |
| DS113298 | 10504 | 11175 | BA_peak_1744 |
| DS113298 | 26276 | 29003 | BA_peak_1745 |
| DS113298 | 29582 | 30133 | BA_peak_1746 |
| DS113298 | 56798 | 57197 | BA_peak_1747 |
| DS113298 | 78759 | 79145 | BA_peak_1748 |
| DS113298 | 137222 | 138567 | BA_peak_1749 |
| DS113299 | 3672 | 4060 | BA_peak_1751 |
| DS113299 | 25667 | 26632 | BA_peak_1752 |
| DS113299 | 28623 | 28965 | BA_peak_1753 |
| DS113299 | 63741 | 64536 | BA_peak_1754 |
| DS113299 | 65376 | 66128 | BA_peak_1755 |
| DS113299 | 74713 | 76585 | BA_peak_1756 |
| DS113299 | 130901 | 131650 | BA_peak_1757 |
| DS113299 | 150325 | 151767 | BA_peak_1758 |
| DS113300 | 5856 | 7207 | BA_peak_1759 |
| DS113300 | 86024 | 86687 | BA_peak_1760 |
| DS113300 | 89135 | 89489 | BA_peak_1761 |
| DS113300 | 102654 | 105042 | BA_peak_1763 |
| DS113300 | 112747 | 113422 | BA_peak_1764 |
| DS113301 | 16821 | 17205 | BA_peak_1765 |
| DS113301 | 54074 | 54992 | BA_peak_1766 |
| DS113301 | 77420 | 78034 | BA_peak_1767 |
| DS113302 | 76283 | 77357 | BA_peak_1768 |
| DS113302 | 92995 | 94236 | BA_peak_1770 |
| DS113302 | 128545 | 129225 | BA_peak_1771 |
| DS113303 | 9727 | 10310 | BA_peak_1772 |
| DS113303 | 13093 | 13864 | BA_peak_1773 |
| DS113303 | 23335 | 24108 | BA_peak_1774 |
| DS113303 | 30690 | 32677 | BA_peak_1775 |
| DS113303 | 91550 | 92126 | BA_peak_1777 |
| DS113303 | 93724 | 94209 | BA_peak_1778 |
| DS113303 | 103666 | 104347 | BA_peak_1779 |
| DS113303 | 111812 | 112255 | BA_peak_1780 |
| DS113303 | 142439 | 143309 | BA_peak_1781 |
| DS113304 | 27923 | 28653 | BA_peak_1782 |

| | | | |
|---|---|---|---|
| DS113304 | 58126 | 58558 | BA_peak_1783 |
| DS113304 | 81744 | 82637 | BA_peak_1784 |
| DS113304 | 85368 | 85853 | BA_peak_1785 |
| DS113304 | 89223 | 89957 | BA_peak_1786 |
| DS113304 | 108904 | 109469 | BA_peak_1787 |
| DS113304 | 110369 | 110757 | BA_peak_1788 |
| DS113304 | 129454 | 129828 | BA_peak_1789 |
| DS113305 | 19458 | 20290 | BA_peak_1790 |
| DS113305 | 26434 | 26832 | BA_peak_1791 |
| DS113305 | 28184 | 28591 | BA_peak_1792 |
| DS113305 | 40410 | 41029 | BA_peak_1793 |
| DS113305 | 63261 | 63752 | BA_peak_1794 |
| DS113305 | 69415 | 70567 | BA_peak_1795 |
| DS113305 | 73150 | 73834 | BA_peak_1796 |
| DS113305 | 88506 | 90038 | BA_peak_1797 |
| DS113305 | 90534 | 91089 | BA_peak_1798 |
| DS113305 | 100523 | 100933 | BA_peak_1799 |
| DS113306 | 16395 | 17022 | BA_peak_1800 |
| DS113306 | 40088 | 40672 | BA_peak_1801 |
| DS113306 | 41112 | 41554 | BA_peak_1802 |
| DS113306 | 53782 | 54568 | BA_peak_1803 |
| DS113306 | 64365 | 64934 | BA_peak_1804 |
| DS113306 | 87314 | 87860 | BA_peak_1806 |
| DS113306 | 94778 | 95464 | BA_peak_1807 |
| DS113306 | 99030 | 99668 | BA_peak_1808 |
| DS113306 | 124244 | 124637 | BA_peak_1809 |
| DS113306 | 126156 | 126855 | BA_peak_1810 |
| DS113307 | 5017 | 6512 | BA_peak_1811 |
| DS113307 | 40278 | 40832 | BA_peak_1812 |
| DS113307 | 71154 | 72640 | BA_peak_1814 |
| DS113307 | 91213 | 91618 | BA_peak_1815 |
| DS113307 | 94215 | 94730 | BA_peak_1817 |
| DS113307 | 100313 | 100700 | BA_peak_1818 |
| DS113307 | 101717 | 102155 | BA_peak_1819 |
| DS113307 | 141844 | 142323 | BA_peak_1820 |
| DS113308 | 14040 | 14705 | BA_peak_1821 |
| DS113308 | 26349 | 26941 | BA_peak_1822 |
| DS113308 | 27306 | 27782 | BA_peak_1822 |
| DS113308 | 68214 | 68962 | BA_peak_1823 |
| DS113308 | 69938 | 71198 | BA_peak_1824 |
| DS113308 | 81611 | 82776 | BA_peak_1825 |
| DS113308 | 103949 | 104517 | BA_peak_1826 |
| DS113308 | 105062 | 105987 | BA_peak_1827 |
| DS113308 | 110532 | 110869 | BA_peak_1828 |
| DS113308 | 122714 | 124087 | BA_peak_1829 |
| DS113308 | 132786 | 134852 | BA_peak_1830 |
| DS113309 | 10609 | 12688 | BA_peak_1831 |

| | | | |
|---|---|---|---|
| DS113309 | 13158 | 13926 | BA_peak_1832 |
| DS113309 | 25739 | 28995 | BA_peak_1833 |
| DS113309 | 44139 | 44576 | BA_peak_1834 |
| DS113310 | 26660 | 27467 | BA_peak_1835 |
| DS113310 | 28702 | 29332 | BA_peak_1836 |
| DS113310 | 29944 | 30320 | BA_peak_1837 |
| DS113310 | 37281 | 38069 | BA_peak_1838 |
| DS113310 | 48392 | 48764 | BA_peak_1839 |
| DS113310 | 50815 | 54044 | BA_peak_1840 |
| DS113310 | 89388 | 89958 | BA_peak_1841 |
| DS113310 | 133746 | 134552 | BA_peak_1842 |
| DS113310 | 140807 | 141160 | BA_peak_1843 |
| DS113310 | 141806 | 142448 | BA_peak_1844 |
| DS113311 | 3219 | 3731 | BA_peak_1845 |
| DS113311 | 17748 | 19109 | BA_peak_1846 |
| DS113311 | 20953 | 21554 | BA_peak_1847 |
| DS113311 | 42054 | 43180 | BA_peak_1848 |
| DS113311 | 75027 | 75793 | BA_peak_1849 |
| DS113311 | 76403 | 77080 | BA_peak_1850 |
| DS113311 | 84395 | 84852 | BA_peak_1851 |
| DS113311 | 88398 | 89194 | BA_peak_1852 |
| DS113311 | 94690 | 95043 | BA_peak_1853 |
| DS113311 | 106160 | 106884 | BA_peak_1854 |
| DS113312 | 85834 | 86273 | BA_peak_1856 |
| DS113312 | 87590 | 87979 | BA_peak_1857 |
| DS113312 | 90716 | 91460 | BA_peak_1858 |
| DS113312 | 122648 | 123057 | BA_peak_1859 |
| DS113313 | 20204 | 20602 | BA_peak_1860 |
| DS113313 | 25612 | 25939 | BA_peak_1861 |
| DS113313 | 100054 | 101919 | BA_peak_1862 |
| DS113313 | 108354 | 108682 | BA_peak_1863 |
| DS113313 | 140398 | 140759 | BA_peak_1864 |
| DS113313 | 142376 | 142935 | BA_peak_1865 |
| DS113314 | 7815 | 8205 | BA_peak_1866 |
| DS113314 | 70440 | 70920 | BA_peak_1868 |
| DS113314 | 77839 | 78833 | BA_peak_1869 |
| DS113314 | 88846 | 89452 | BA_peak_1870 |
| DS113314 | 90921 | 91689 | BA_peak_1871 |
| DS113314 | 122957 | 125591 | BA_peak_1873 |
| DS113314 | 138647 | 139359 | BA_peak_1874 |
| DS113315 | 37109 | 37741 | BA_peak_1875 |
| DS113315 | 38695 | 39995 | BA_peak_1876 |
| DS113315 | 66057 | 66519 | BA_peak_1877 |
| DS113315 | 75148 | 75962 | BA_peak_1878 |
| DS113315 | 78627 | 79095 | BA_peak_1880 |
| DS113315 | 84140 | 87473 | BA_peak_1881 |
| DS113315 | 88885 | 90942 | BA_peak_1882 |

| | | | |
|---|---|---|---|
| DS113315 | 102489 | 103003 | BA_peak_1883 |
| DS113315 | 110128 | 112029 | BA_peak_1884 |
| DS113315 | 112503 | 113070 | BA_peak_1885 |
| DS113315 | 117260 | 118489 | BA_peak_1886 |
| DS113315 | 129225 | 130163 | BA_peak_1887 |
| DS113315 | 138513 | 139200 | BA_peak_1888 |
| DS113316 | 4585 | 5670 | BA_peak_1889 |
| DS113316 | 14152 | 15134 | BA_peak_1890 |
| DS113316 | 16807 | 19105 | BA_peak_1891 |
| DS113316 | 27527 | 29203 | BA_peak_1892 |
| DS113316 | 29702 | 30201 | BA_peak_1893 |
| DS113316 | 31853 | 32398 | BA_peak_1894 |
| DS113316 | 40118 | 40863 | BA_peak_1895 |
| DS113316 | 42545 | 45743 | BA_peak_1896 |
| DS113316 | 48833 | 49291 | BA_peak_1897 |
| DS113316 | 50623 | 51314 | BA_peak_1898 |
| DS113316 | 60977 | 61839 | BA_peak_1899 |
| DS113316 | 63073 | 63611 | BA_peak_1900 |
| DS113316 | 65021 | 65688 | BA_peak_1901 |
| DS113316 | 82113 | 82445 | BA_peak_1902 |
| DS113316 | 98193 | 98623 | BA_peak_1903 |
| DS113316 | 139877 | 140529 | BA_peak_1905 |
| DS113317 | 137927 | 138375 | BA_peak_1906 |
| DS113318 | 3927 | 4253 | BA_peak_1907 |
| DS113318 | 6455 | 7835 | BA_peak_1908 |
| DS113318 | 11372 | 12205 | BA_peak_1909 |
| DS113318 | 17159 | 17822 | BA_peak_1910 |
| DS113318 | 41024 | 43478 | BA_peak_1911 |
| DS113318 | 44926 | 45511 | BA_peak_1912 |
| DS113318 | 46209 | 48076 | BA_peak_1913 |
| DS113318 | 48815 | 49753 | BA_peak_1914 |
| DS113318 | 83866 | 84473 | BA_peak_1915 |
| DS113318 | 115247 | 115907 | BA_peak_1916 |
| DS113319 | 41853 | 43661 | BA_peak_1917 |
| DS113319 | 57196 | 59311 | BA_peak_1918 |
| DS113319 | 82718 | 83495 | BA_peak_1919 |
| DS113319 | 122183 | 122746 | BA_peak_1920 |
| DS113319 | 127221 | 127869 | BA_peak_1921 |
| DS113320 | 2698 | 3173 | BA_peak_1922 |
| DS113320 | 11392 | 12212 | BA_peak_1923 |
| DS113320 | 29297 | 30243 | BA_peak_1924 |
| DS113320 | 31443 | 32021 | BA_peak_1925 |
| DS113320 | 98390 | 98731 | BA_peak_1926 |
| DS113320 | 120560 | 121234 | BA_peak_1927 |
| DS113320 | 126562 | 127459 | BA_peak_1928 |
| DS113320 | 132869 | 134661 | BA_peak_1929 |
| DS113321 | 44806 | 45381 | BA_peak_1930 |

| | | | |
|---|---|---|---|
| DS113321 | 47969 | 48354 | BA_peak_1932 |
| DS113321 | 54552 | 54953 | BA_peak_1933 |
| DS113321 | 111775 | 112232 | BA_peak_1934 |
| DS113321 | 127888 | 128289 | BA_peak_1935 |
| DS113321 | 129474 | 130013 | BA_peak_1936 |
| DS113321 | 130485 | 130884 | BA_peak_1937 |
| DS113321 | 134370 | 135195 | BA_peak_1938 |
| DS113322 | 30909 | 32204 | BA_peak_1939 |
| DS113322 | 67267 | 68003 | BA_peak_1940 |
| DS113322 | 78324 | 79725 | BA_peak_1941 |
| DS113322 | 113479 | 114249 | BA_peak_1942 |
| DS113322 | 118976 | 119618 | BA_peak_1943 |
| DS113323 | 2798 | 6076 | BA_peak_1944 |
| DS113323 | 37186 | 37546 | BA_peak_1945 |
| DS113323 | 43468 | 44250 | BA_peak_1947 |
| DS113323 | 69805 | 71269 | BA_peak_1948 |
| DS113323 | 98118 | 99054 | BA_peak_1949 |
| DS113323 | 125012 | 126107 | BA_peak_1950 |
| DS113323 | 129390 | 131176 | BA_peak_1951 |
| DS113323 | 135705 | 136118 | BA_peak_1952 |
| DS113324 | 25728 | 27482 | BA_peak_1953 |
| DS113324 | 41959 | 43016 | BA_peak_1954 |
| DS113324 | 52099 | 52492 | BA_peak_1955 |
| DS113324 | 74647 | 75410 | BA_peak_1956 |
| DS113324 | 99049 | 100619 | BA_peak_1957 |
| DS113324 | 129097 | 129536 | BA_peak_1958 |
| DS113324 | 133359 | 134084 | BA_peak_1959 |
| DS113324 | 135134 | 135756 | BA_peak_1960 |
| DS113324 | 136536 | 137116 | BA_peak_1961 |
| DS113325 | 18150 | 18486 | BA_peak_1962 |
| DS113325 | 35778 | 37057 | BA_peak_1963 |
| DS113325 | 79939 | 81484 | BA_peak_1964 |
| DS113325 | 83752 | 87106 | BA_peak_1965 |
| DS113325 | 90711 | 92489 | BA_peak_1966 |
| DS113325 | 109165 | 109744 | BA_peak_1967 |
| DS113325 | 119519 | 120128 | BA_peak_1968 |
| DS113325 | 125796 | 126382 | BA_peak_1969 |
| DS113325 | 129288 | 129915 | BA_peak_1970 |
| DS113326 | 17963 | 18575 | BA_peak_1972 |
| DS113326 | 19158 | 20263 | BA_peak_1973 |
| DS113326 | 21060 | 22275 | BA_peak_1974 |
| DS113326 | 26370 | 27054 | BA_peak_1975 |
| DS113326 | 27742 | 28322 | BA_peak_1976 |
| DS113326 | 29775 | 31881 | BA_peak_1977 |
| DS113326 | 32840 | 33473 | BA_peak_1978 |
| DS113326 | 38621 | 39452 | BA_peak_1979 |
| DS113326 | 72718 | 73480 | BA_peak_1980 |

| | | | |
|---|---|---|---|
| DS113326 | 83987 | 84652 | BA_peak_1981 |
| DS113326 | 124171 | 124782 | BA_peak_1982 |
| DS113327 | 2829 | 3505 | BA_peak_1983 |
| DS113327 | 22177 | 22605 | BA_peak_1984 |
| DS113327 | 23396 | 24408 | BA_peak_1985 |
| DS113327 | 27452 | 28054 | BA_peak_1986 |
| DS113327 | 29159 | 30394 | BA_peak_1987 |
| DS113327 | 68978 | 69559 | BA_peak_1989 |
| DS113327 | 77373 | 79122 | BA_peak_1990 |
| DS113327 | 87611 | 88055 | BA_peak_1992 |
| DS113327 | 90432 | 90787 | BA_peak_1993 |
| DS113327 | 125024 | 125439 | BA_peak_1994 |
| DS113328 | 16973 | 17606 | BA_peak_1995 |
| DS113328 | 53855 | 55257 | BA_peak_1996 |
| DS113328 | 55928 | 57551 | BA_peak_1997 |
| DS113328 | 81566 | 81965 | BA_peak_1998 |
| DS113328 | 88340 | 89485 | BA_peak_1999 |
| DS113328 | 90271 | 90610 | BA_peak_2000 |
| DS113328 | 91702 | 92145 | BA_peak_2001 |
| DS113328 | 97178 | 97577 | BA_peak_2002 |
| DS113328 | 101963 | 103522 | BA_peak_2003 |
| DS113328 | 104085 | 104521 | BA_peak_2004 |
| DS113328 | 117063 | 117723 | BA_peak_2005 |
| DS113328 | 121057 | 121732 | BA_peak_2006 |
| DS113329 | 28651 | 30759 | BA_peak_2007 |
| DS113329 | 53719 | 54076 | BA_peak_2008 |
| DS113329 | 66848 | 67606 | BA_peak_2009 |
| DS113329 | 68080 | 69214 | BA_peak_2010 |
| DS113329 | 78131 | 83496 | BA_peak_2011 |
| DS113329 | 105395 | 106183 | BA_peak_2012 |
| DS113329 | 124384 | 125028 | BA_peak_2013 |
| DS113330 | 37398 | 37721 | BA_peak_2015 |
| DS113330 | 40185 | 40603 | BA_peak_2016 |
| DS113330 | 47184 | 47759 | BA_peak_2017 |
| DS113330 | 82923 | 84955 | BA_peak_2018 |
| DS113330 | 94680 | 95477 | BA_peak_2019 |
| DS113330 | 107576 | 107977 | BA_peak_2020 |
| DS113330 | 108899 | 109275 | BA_peak_2021 |
| DS113330 | 120100 | 120754 | BA_peak_2023 |
| DS113331 | 5353 | 9813 | BA_peak_2024 |
| DS113331 | 12545 | 13017 | BA_peak_2025 |
| DS113331 | 14177 | 15325 | BA_peak_2026 |
| DS113331 | 27067 | 28056 | BA_peak_2027 |
| DS113331 | 30423 | 31142 | BA_peak_2028 |
| DS113331 | 32073 | 32804 | BA_peak_2029 |
| DS113331 | 76636 | 77050 | BA_peak_2030 |
| DS113331 | 78123 | 79458 | BA_peak_2031 |

| | | | |
|---|---|---|---|
| DS113331 | 81379 | 83914 | BA_peak_2032 |
| DS113331 | 96119 | 97024 | BA_peak_2033 |
| DS113331 | 105932 | 107177 | BA_peak_2034 |
| DS113331 | 108716 | 109488 | BA_peak_2035 |
| DS113331 | 110742 | 111759 | BA_peak_2036 |
| DS113332 | 14583 | 15285 | BA_peak_2037 |
| DS113332 | 20435 | 21416 | BA_peak_2038 |
| DS113332 | 23450 | 24915 | BA_peak_2039 |
| DS113332 | 64850 | 65181 | BA_peak_2040 |
| DS113332 | 80044 | 80830 | BA_peak_2041 |
| DS113332 | 122208 | 122648 | BA_peak_2042 |
| DS113333 | 121 | 1891 | BA_peak_2043 |
| DS113333 | 35984 | 38068 | BA_peak_2044 |
| DS113333 | 52949 | 54326 | BA_peak_2045 |
| DS113333 | 63587 | 64313 | BA_peak_2046 |
| DS113333 | 68928 | 69700 | BA_peak_2048 |
| DS113333 | 85156 | 85613 | BA_peak_2049 |
| DS113333 | 98237 | 99165 | BA_peak_2050 |
| DS113333 | 120159 | 120574 | BA_peak_2051 |
| DS113334 | 11409 | 12230 | BA_peak_2052 |
| DS113334 | 16194 | 16500 | BA_peak_2053 |
| DS113334 | 29512 | 29821 | BA_peak_2054 |
| DS113334 | 47634 | 47972 | BA_peak_2055 |
| DS113334 | 57800 | 58189 | BA_peak_2056 |
| DS113334 | 72285 | 72645 | BA_peak_2059 |
| DS113334 | 73091 | 73819 | BA_peak_2060 |
| DS113334 | 75397 | 76227 | BA_peak_2061 |
| DS113334 | 81164 | 81478 | BA_peak_2062 |
| DS113334 | 82112 | 82939 | BA_peak_2063 |
| DS113334 | 93593 | 94037 | BA_peak_2064 |
| DS113334 | 95857 | 96237 | BA_peak_2065 |
| DS113334 | 105119 | 105663 | BA_peak_2066 |
| DS113334 | 112942 | 113251 | BA_peak_2067 |
| DS113334 | 121894 | 122493 | BA_peak_2068 |
| DS113334 | 127615 | 128461 | BA_peak_2069 |
| DS113334 | 130778 | 131803 | BA_peak_2070 |
| DS113335 | 10780 | 11413 | BA_peak_2071 |
| DS113335 | 11742 | 12124 | BA_peak_2072 |
| DS113335 | 12904 | 13835 | BA_peak_2073 |
| DS113335 | 27712 | 28787 | BA_peak_2074 |
| DS113335 | 40811 | 42272 | BA_peak_2075 |
| DS113335 | 106429 | 106826 | BA_peak_2076 |
| DS113335 | 124727 | 125192 | BA_peak_2077 |
| DS113335 | 126459 | 127438 | BA_peak_2078 |
| DS113336 | 24861 | 25271 | BA_peak_2079 |
| DS113336 | 80665 | 81063 | BA_peak_2080 |
| DS113336 | 84326 | 85098 | BA_peak_2081 |

| | | | |
|---|---|---|---|
| DS113336 | 85873 | 89020 | BA_peak_2082 |
| DS113337 | 2274 | 3010 | BA_peak_2083 |
| DS113337 | 4400 | 4693 | BA_peak_2084 |
| DS113337 | 16340 | 18488 | BA_peak_2085 |
| DS113337 | 20190 | 20779 | BA_peak_2086 |
| DS113337 | 24917 | 25534 | BA_peak_2087 |
| DS113337 | 40268 | 41063 | BA_peak_2088 |
| DS113337 | 67284 | 67673 | BA_peak_2089 |
| DS113337 | 83184 | 84067 | BA_peak_2090 |
| DS113337 | 91811 | 92263 | BA_peak_2091 |
| DS113337 | 105358 | 108639 | BA_peak_2092 |
| DS113337 | 114716 | 116532 | BA_peak_2093 |
| DS113338 | 13624 | 14057 | BA_peak_2095 |
| DS113338 | 51393 | 51707 | BA_peak_2096 |
| DS113338 | 66140 | 69496 | BA_peak_2097 |
| DS113338 | 72335 | 73282 | BA_peak_2098 |
| DS113338 | 76445 | 76789 | BA_peak_2099 |
| DS113338 | 77380 | 78282 | BA_peak_2100 |
| DS113338 | 78643 | 81256 | BA_peak_2101 |
| DS113338 | 92893 | 93526 | BA_peak_2102 |
| DS113338 | 127643 | 128382 | BA_peak_2103 |
| DS113339 | 88648 | 89319 | BA_peak_2104 |
| DS113339 | 93018 | 93503 | BA_peak_2105 |
| DS113339 | 102373 | 104119 | BA_peak_2106 |
| DS113339 | 126302 | 126822 | BA_peak_2107 |
| DS113340 | 30360 | 31213 | BA_peak_2110 |
| DS113340 | 60164 | 60545 | BA_peak_2111 |
| DS113340 | 94001 | 94563 | BA_peak_2112 |
| DS113340 | 95194 | 95928 | BA_peak_2113 |
| DS113340 | 110860 | 111664 | BA_peak_2114 |
| DS113340 | 121106 | 122882 | BA_peak_2115 |
| DS113341 | 468 | 961 | BA_peak_2116 |
| DS113341 | 1967 | 2660 | BA_peak_2117 |
| DS113341 | 4080 | 4764 | BA_peak_2118 |
| DS113341 | 56068 | 56971 | BA_peak_2119 |
| DS113341 | 69091 | 69523 | BA_peak_2120 |
| DS113341 | 97129 | 97474 | BA_peak_2121 |
| DS113341 | 118078 | 119007 | BA_peak_2122 |
| DS113342 | 7068 | 7723 | BA_peak_2123 |
| DS113342 | 8799 | 9118 | BA_peak_2124 |
| DS113342 | 14673 | 17323 | BA_peak_2125 |
| DS113342 | 22752 | 23687 | BA_peak_2126 |
| DS113342 | 29193 | 30081 | BA_peak_2127 |
| DS113342 | 40378 | 41125 | BA_peak_2128 |
| DS113342 | 41562 | 42359 | BA_peak_2129 |
| DS113342 | 92185 | 92987 | BA_peak_2130 |
| DS113342 | 96398 | 96853 | BA_peak_2131 |

| | | | |
|---|---|---|---|
| DS113342 | 121542 | 122109 | BA_peak_2132 |
| DS113342 | 129997 | 130479 | BA_peak_2133 |
| DS113342 | 131317 | 132242 | BA_peak_2134 |
| DS113343 | 30902 | 32830 | BA_peak_2135 |
| DS113343 | 49402 | 50120 | BA_peak_2136 |
| DS113343 | 93616 | 95176 | BA_peak_2138 |
| DS113343 | 102361 | 103907 | BA_peak_2139 |
| DS113343 | 112262 | 113237 | BA_peak_2140 |
| DS113344 | 21671 | 22105 | BA_peak_2141 |
| DS113344 | 26600 | 27415 | BA_peak_2142 |
| DS113344 | 50351 | 51100 | BA_peak_2144 |
| DS113344 | 64871 | 66041 | BA_peak_2145 |
| DS113344 | 77716 | 79592 | BA_peak_2146 |
| DS113344 | 92666 | 92999 | BA_peak_2147 |
| DS113344 | 102978 | 103574 | BA_peak_2148 |
| DS113344 | 107031 | 107966 | BA_peak_2149 |
| DS113344 | 108386 | 109696 | BA_peak_2150 |
| DS113344 | 121348 | 122088 | BA_peak_2151 |
| DS113344 | 122865 | 123486 | BA_peak_2152 |
| DS113344 | 124787 | 125174 | BA_peak_2153 |
| DS113345 | 15236 | 15738 | BA_peak_2154 |
| DS113345 | 26362 | 27799 | BA_peak_2155 |
| DS113345 | 29419 | 30125 | BA_peak_2156 |
| DS113345 | 114581 | 115456 | BA_peak_2158 |
| DS113345 | 123586 | 126480 | BA_peak_2159 |
| DS113345 | 128221 | 129203 | BA_peak_2160 |
| DS113346 | 10441 | 11736 | BA_peak_2161 |
| DS113346 | 17004 | 17842 | BA_peak_2162 |
| DS113346 | 29411 | 30391 | BA_peak_2163 |
| DS113346 | 81036 | 81380 | BA_peak_2164 |
| DS113346 | 86088 | 86436 | BA_peak_2165 |
| DS113346 | 87895 | 88606 | BA_peak_2166 |
| DS113347 | 2964 | 3500 | BA_peak_2167 |
| DS113347 | 4142 | 4562 | BA_peak_2168 |
| DS113347 | 33903 | 34378 | BA_peak_2169 |
| DS113347 | 47338 | 47715 | BA_peak_2170 |
| DS113347 | 48228 | 49310 | BA_peak_2171 |
| DS113347 | 52265 | 53776 | BA_peak_2172 |
| DS113347 | 60398 | 61747 | BA_peak_2173 |
| DS113347 | 64059 | 64832 | BA_peak_2174 |
| DS113347 | 109185 | 109561 | BA_peak_2175 |
| DS113348 | 60 | 451 | BA_peak_2176 |
| DS113348 | 3883 | 4539 | BA_peak_2177 |
| DS113348 | 28849 | 29176 | BA_peak_2178 |
| DS113348 | 35421 | 36218 | BA_peak_2179 |
| DS113348 | 36761 | 37415 | BA_peak_2180 |
| DS113348 | 61267 | 62050 | BA_peak_2181 |

| | | | |
|---|---|---|---|
| DS113348 | 80795 | 81334 | BA_peak_2182 |
| DS113348 | 105964 | 106874 | BA_peak_2183 |
| DS113348 | 113077 | 114159 | BA_peak_2184 |
| DS113348 | 124284 | 125057 | BA_peak_2185 |
| DS113348 | 125533 | 126334 | BA_peak_2186 |
| DS113349 | 4897 | 5353 | BA_peak_2187 |
| DS113349 | 57821 | 58209 | BA_peak_2188 |
| DS113349 | 67634 | 68507 | BA_peak_2189 |
| DS113349 | 69006 | 70064 | BA_peak_2190 |
| DS113349 | 77525 | 78501 | BA_peak_2191 |
| DS113349 | 101220 | 103044 | BA_peak_2192 |
| DS113349 | 116978 | 117395 | BA_peak_2193 |
| DS113349 | 126402 | 126815 | BA_peak_2194 |
| DS113349 | 127320 | 127768 | BA_peak_2195 |
| DS113350 | 27770 | 29815 | BA_peak_2196 |
| DS113350 | 37282 | 37593 | BA_peak_2197 |
| DS113350 | 42805 | 43581 | BA_peak_2198 |
| DS113350 | 70371 | 71036 | BA_peak_2199 |
| DS113350 | 82019 | 82627 | BA_peak_2200 |
| DS113350 | 83171 | 84245 | BA_peak_2201 |
| DS113350 | 91605 | 92511 | BA_peak_2202 |
| DS113350 | 103608 | 104155 | BA_peak_2203 |
| DS113350 | 104696 | 105138 | BA_peak_2204 |
| DS113350 | 109271 | 110295 | BA_peak_2205 |
| DS113350 | 122192 | 124720 | BA_peak_2206 |
| DS113351 | 778 | 1798 | BA_peak_2207 |
| DS113351 | 2162 | 2656 | BA_peak_2208 |
| DS113351 | 14128 | 15440 | BA_peak_2209 |
| DS113351 | 38713 | 39244 | BA_peak_2211 |
| DS113351 | 60366 | 62267 | BA_peak_2213 |
| DS113351 | 65391 | 65826 | BA_peak_2214 |
| DS113351 | 74591 | 75688 | BA_peak_2215 |
| DS113351 | 92183 | 92814 | BA_peak_2216 |
| DS113351 | 105323 | 106136 | BA_peak_2217 |
| DS113351 | 115508 | 116127 | BA_peak_2218 |
| DS113351 | 116735 | 117149 | BA_peak_2219 |
| DS113351 | 122809 | 123284 | BA_peak_2220 |
| DS113352 | 25850 | 26340 | BA_peak_2221 |
| DS113352 | 26697 | 27720 | BA_peak_2222 |
| DS113352 | 32357 | 32930 | BA_peak_2223 |
| DS113352 | 35268 | 36579 | BA_peak_2224 |
| DS113352 | 94703 | 95537 | BA_peak_2225 |
| DS113352 | 101055 | 101523 | BA_peak_2226 |
| DS113352 | 105812 | 106613 | BA_peak_2227 |
| DS113352 | 118331 | 118993 | BA_peak_2228 |
| DS113352 | 121060 | 122335 | BA_peak_2229 |
| DS113352 | 124670 | 125228 | BA_peak_2230 |

| | | | |
|---|---|---|---|
| DS113353 | 14166 | 14600 | BA_peak_2231 |
| DS113353 | 53483 | 55921 | BA_peak_2233 |
| DS113353 | 63224 | 64009 | BA_peak_2234 |
| DS113353 | 72097 | 72488 | BA_peak_2235 |
| DS113353 | 97041 | 97469 | BA_peak_2236 |
| DS113353 | 103542 | 104077 | BA_peak_2237 |
| DS113353 | 114212 | 117523 | BA_peak_2238 |
| DS113354 | 2316 | 2754 | BA_peak_2239 |
| DS113354 | 4409 | 4966 | BA_peak_2240 |
| DS113354 | 5726 | 6485 | BA_peak_2241 |
| DS113354 | 22579 | 23035 | BA_peak_2242 |
| DS113354 | 30219 | 30649 | BA_peak_2243 |
| DS113354 | 49746 | 50697 | BA_peak_2244 |
| DS113354 | 93289 | 93728 | BA_peak_2245 |
| DS113354 | 97117 | 97782 | BA_peak_2246 |
| DS113354 | 99056 | 99649 | BA_peak_2247 |
| DS113354 | 123466 | 123849 | BA_peak_2248 |
| DS113355 | 2123 | 2543 | BA_peak_2249 |
| DS113355 | 8731 | 10164 | BA_peak_2250 |
| DS113355 | 21639 | 22341 | BA_peak_2251 |
| DS113355 | 25017 | 26236 | BA_peak_2252 |
| DS113355 | 27121 | 27924 | BA_peak_2253 |
| DS113355 | 120622 | 121910 | BA_peak_2254 |
| DS113356 | 19832 | 20354 | BA_peak_2255 |
| DS113356 | 25071 | 25451 | BA_peak_2256 |
| DS113356 | 94470 | 95742 | BA_peak_2257 |
| DS113356 | 96530 | 96969 | BA_peak_2258 |
| DS113356 | 123278 | 123727 | BA_peak_2259 |
| DS113357 | 5427 | 6318 | BA_peak_2260 |
| DS113357 | 12106 | 12498 | BA_peak_2261 |
| DS113357 | 15725 | 16494 | BA_peak_2262 |
| DS113357 | 38332 | 39051 | BA_peak_2263 |
| DS113357 | 58216 | 58723 | BA_peak_2264 |
| DS113357 | 119143 | 119853 | BA_peak_2265 |
| DS113357 | 121478 | 123298 | BA_peak_2266 |
| DS113358 | 985 | 1671 | BA_peak_2267 |
| DS113358 | 27516 | 27881 | BA_peak_2268 |
| DS113358 | 28603 | 29019 | BA_peak_2269 |
| DS113358 | 50503 | 51944 | BA_peak_2270 |
| DS113358 | 66546 | 67153 | BA_peak_2271 |
| DS113358 | 68148 | 68954 | BA_peak_2272 |
| DS113358 | 70349 | 71481 | BA_peak_2273 |
| DS113358 | 71994 | 72725 | BA_peak_2274 |
| DS113358 | 73046 | 76681 | BA_peak_2275 |
| DS113358 | 103614 | 104081 | BA_peak_2276 |
| DS113358 | 106021 | 107588 | BA_peak_2277 |
| DS113359 | 23005 | 26731 | BA_peak_2278 |

| | | | |
|---|---|---|---|
| DS113359 | 27222 | 27959 | BA_peak_2278 |
| DS113359 | 112526 | 113108 | BA_peak_2280 |
| DS113360 | 6310 | 6806 | BA_peak_2281 |
| DS113360 | 9148 | 9481 | BA_peak_2282 |
| DS113360 | 14784 | 15385 | BA_peak_2283 |
| DS113360 | 16256 | 16576 | BA_peak_2284 |
| DS113360 | 17292 | 17808 | BA_peak_2285 |
| DS113360 | 18205 | 18852 | BA_peak_2286 |
| DS113360 | 51482 | 51916 | BA_peak_2287 |
| DS113360 | 101950 | 102345 | BA_peak_2288 |
| DS113360 | 112768 | 113881 | BA_peak_2289 |
| DS113360 | 118496 | 118887 | BA_peak_2290 |
| DS113360 | 121182 | 122436 | BA_peak_2291 |
| DS113361 | 3344 | 3755 | BA_peak_2292 |
| DS113361 | 5273 | 5586 | BA_peak_2293 |
| DS113361 | 6029 | 7589 | BA_peak_2294 |
| DS113361 | 79896 | 80241 | BA_peak_2295 |
| DS113361 | 81235 | 81695 | BA_peak_2296 |
| DS113361 | 92773 | 93173 | BA_peak_2297 |
| DS113361 | 93700 | 94852 | BA_peak_2298 |
| DS113361 | 99549 | 99869 | BA_peak_2299 |
| DS113361 | 118120 | 119053 | BA_peak_2300 |
| DS113362 | 71 | 392 | BA_peak_2301 |
| DS113362 | 18493 | 19125 | BA_peak_2302 |
| DS113362 | 19669 | 21743 | BA_peak_2303 |
| DS113362 | 30844 | 31544 | BA_peak_2304 |
| DS113362 | 46164 | 46518 | BA_peak_2305 |
| DS113362 | 48547 | 49174 | BA_peak_2306 |
| DS113362 | 66913 | 68488 | BA_peak_2307 |
| DS113362 | 80951 | 81919 | BA_peak_2308 |
| DS113362 | 87215 | 87877 | BA_peak_2309 |
| DS113362 | 100063 | 100419 | BA_peak_2310 |
| DS113362 | 102701 | 103114 | BA_peak_2311 |
| DS113363 | 44 | 903 | BA_peak_2312 |
| DS113363 | 1352 | 2001 | BA_peak_2313 |
| DS113363 | 6154 | 6537 | BA_peak_2314 |
| DS113363 | 19996 | 20815 | BA_peak_2315 |
| DS113363 | 68024 | 70129 | BA_peak_2316 |
| DS113363 | 71524 | 71883 | BA_peak_2317 |
| DS113363 | 93958 | 94362 | BA_peak_2318 |
| DS113363 | 100871 | 101290 | BA_peak_2319 |
| DS113363 | 102068 | 103503 | BA_peak_2320 |
| DS113363 | 118142 | 118788 | BA_peak_2321 |
| DS113364 | 6718 | 8499 | BA_peak_2322 |
| DS113364 | 41361 | 41681 | BA_peak_2323 |
| DS113364 | 119630 | 120092 | BA_peak_2325 |
| DS113365 | 38124 | 38387 | BA_peak_2326 |

| | | | |
|---|---|---|---|
| DS113365 | 68607 | 69212 | BA_peak_2327 |
| DS113365 | 111040 | 112735 | BA_peak_2328 |
| DS113365 | 113882 | 114300 | BA_peak_2329 |
| DS113366 | 32699 | 33075 | BA_peak_2330 |
| DS113366 | 38033 | 38729 | BA_peak_2331 |
| DS113366 | 47768 | 48123 | BA_peak_2332 |
| DS113366 | 54363 | 55167 | BA_peak_2333 |
| DS113366 | 57669 | 59704 | BA_peak_2334 |
| DS113366 | 62407 | 62858 | BA_peak_2335 |
| DS113366 | 80597 | 81301 | BA_peak_2336 |
| DS113366 | 87034 | 90352 | BA_peak_2337 |
| DS113366 | 105103 | 105597 | BA_peak_2338 |
| DS113366 | 106192 | 108300 | BA_peak_2339 |
| DS113366 | 116410 | 117185 | BA_peak_2340 |
| DS113366 | 117939 | 118387 | BA_peak_2341 |
| DS113367 | 3013 | 3495 | BA_peak_2342 |
| DS113367 | 66952 | 68047 | BA_peak_2343 |
| DS113367 | 85093 | 85608 | BA_peak_2344 |
| DS113367 | 107854 | 108891 | BA_peak_2345 |
| DS113367 | 120959 | 121357 | BA_peak_2346 |
| DS113368 | 14472 | 15797 | BA_peak_2347 |
| DS113368 | 16507 | 17049 | BA_peak_2348 |
| DS113368 | 17920 | 18717 | BA_peak_2349 |
| DS113368 | 19908 | 21017 | BA_peak_2350 |
| DS113368 | 41074 | 43152 | BA_peak_2351 |
| DS113368 | 67136 | 67499 | BA_peak_2352 |
| DS113368 | 68829 | 69603 | BA_peak_2353 |
| DS113368 | 89102 | 89549 | BA_peak_2354 |
| DS113368 | 91220 | 91636 | BA_peak_2355 |
| DS113368 | 92059 | 92596 | BA_peak_2356 |
| DS113368 | 115432 | 116190 | BA_peak_2357 |
| DS113369 | 1884 | 2301 | BA_peak_2358 |
| DS113369 | 16425 | 17015 | BA_peak_2359 |
| DS113369 | 35166 | 35551 | BA_peak_2360 |
| DS113369 | 50140 | 50622 | BA_peak_2361 |
| DS113369 | 114027 | 114469 | BA_peak_2362 |
| DS113369 | 115022 | 115410 | BA_peak_2363 |
| DS113370 | 22893 | 23710 | BA_peak_2364 |
| DS113370 | 24103 | 25042 | BA_peak_2365 |
| DS113370 | 29023 | 29588 | BA_peak_2366 |
| DS113370 | 48974 | 49853 | BA_peak_2367 |
| DS113370 | 87284 | 87975 | BA_peak_2368 |
| DS113370 | 101305 | 102078 | BA_peak_2369 |
| DS113370 | 102542 | 103260 | BA_peak_2370 |
| DS113370 | 115071 | 115407 | BA_peak_2371 |
| DS113371 | 7876 | 8289 | BA_peak_2372 |
| DS113371 | 11847 | 12248 | BA_peak_2373 |

| | | | |
|---|---|---|---|
| DS113371 | 29631 | 29976 | BA_peak_2374 |
| DS113371 | 37987 | 39150 | BA_peak_2376 |
| DS113371 | 57284 | 57949 | BA_peak_2377 |
| DS113371 | 66115 | 66422 | BA_peak_2378 |
| DS113371 | 76023 | 76738 | BA_peak_2379 |
| DS113371 | 90922 | 95012 | BA_peak_2380 |
| DS113371 | 100048 | 100726 | BA_peak_2381 |
| DS113371 | 102957 | 103529 | BA_peak_2382 |
| DS113372 | 18054 | 19398 | BA_peak_2384 |
| DS113373 | 44848 | 45674 | BA_peak_2385 |
| DS113373 | 56152 | 56416 | BA_peak_2387 |
| DS113373 | 56776 | 57110 | BA_peak_2388 |
| DS113373 | 68277 | 69048 | BA_peak_2389 |
| DS113373 | 73823 | 74552 | BA_peak_2390 |
| DS113373 | 75282 | 75818 | BA_peak_2391 |
| DS113373 | 114326 | 116412 | BA_peak_2392 |
| DS113374 | 119 | 625 | BA_peak_2393 |
| DS113374 | 3108 | 3703 | BA_peak_2394 |
| DS113374 | 41792 | 42131 | BA_peak_2395 |
| DS113374 | 50481 | 50946 | BA_peak_2396 |
| DS113374 | 55022 | 55440 | BA_peak_2397 |
| DS113375 | 14522 | 15838 | BA_peak_2398 |
| DS113375 | 41095 | 41601 | BA_peak_2400 |
| DS113375 | 110674 | 113296 | BA_peak_2401 |
| DS113376 | 10357 | 10721 | BA_peak_2402 |
| DS113376 | 40284 | 40625 | BA_peak_2403 |
| DS113376 | 55978 | 56917 | BA_peak_2404 |
| DS113376 | 86463 | 86896 | BA_peak_2406 |
| DS113376 | 112138 | 113136 | BA_peak_2407 |
| DS113377 | 6937 | 7496 | BA_peak_2408 |
| DS113377 | 17621 | 18264 | BA_peak_2409 |
| DS113377 | 51589 | 52707 | BA_peak_2410 |
| DS113377 | 54473 | 55359 | BA_peak_2411 |
| DS113377 | 56959 | 57299 | BA_peak_2412 |
| DS113377 | 59505 | 60218 | BA_peak_2413 |
| DS113377 | 85120 | 85709 | BA_peak_2414 |
| DS113377 | 101362 | 101672 | BA_peak_2415 |
| DS113377 | 106903 | 107683 | BA_peak_2416 |
| DS113378 | 2561 | 3234 | BA_peak_2417 |
| DS113378 | 3635 | 4257 | BA_peak_2418 |
| DS113378 | 5461 | 6077 | BA_peak_2419 |
| DS113378 | 6419 | 7229 | BA_peak_2420 |
| DS113378 | 25731 | 26308 | BA_peak_2421 |
| DS113378 | 53690 | 54107 | BA_peak_2422 |
| DS113378 | 118157 | 118598 | BA_peak_2423 |
| DS113379 | 57111 | 57711 | BA_peak_2424 |
| DS113379 | 67883 | 68687 | BA_peak_2425 |

| | | | |
|---|---|---|---|
| DS113379 | 77840 | 78831 | BA_peak_2426 |
| DS113379 | 87645 | 88119 | BA_peak_2427 |
| DS113379 | 102115 | 103327 | BA_peak_2428 |
| DS113379 | 103643 | 104021 | BA_peak_2429 |
| DS113379 | 117517 | 117864 | BA_peak_2430 |
| DS113379 | 118449 | 118866 | BA_peak_2431 |
| DS113380 | 8187 | 8681 | BA_peak_2433 |
| DS113380 | 15248 | 15874 | BA_peak_2435 |
| DS113380 | 20783 | 21285 | BA_peak_2436 |
| DS113380 | 21692 | 22228 | BA_peak_2437 |
| DS113380 | 29606 | 31309 | BA_peak_2438 |
| DS113380 | 34394 | 35538 | BA_peak_2439 |
| DS113380 | 38894 | 40019 | BA_peak_2440 |
| DS113380 | 50781 | 51119 | BA_peak_2441 |
| DS113380 | 51499 | 53554 | BA_peak_2442 |
| DS113380 | 62622 | 63175 | BA_peak_2443 |
| DS113380 | 68078 | 69050 | BA_peak_2444 |
| DS113380 | 81187 | 81566 | BA_peak_2445 |
| DS113380 | 86929 | 87505 | BA_peak_2446 |
| DS113381 | 22406 | 22809 | BA_peak_2447 |
| DS113381 | 60082 | 60797 | BA_peak_2448 |
| DS113381 | 64562 | 65117 | BA_peak_2449 |
| DS113381 | 65898 | 66592 | BA_peak_2450 |
| DS113381 | 80271 | 83075 | BA_peak_2453 |
| DS113381 | 93807 | 94650 | BA_peak_2454 |
| DS113381 | 102318 | 102904 | BA_peak_2455 |
| DS113382 | 6937 | 7500 | BA_peak_2456 |
| DS113382 | 15810 | 16183 | BA_peak_2457 |
| DS113382 | 25294 | 25997 | BA_peak_2458 |
| DS113382 | 26982 | 28429 | BA_peak_2459 |
| DS113382 | 32666 | 33208 | BA_peak_2460 |
| DS113382 | 44105 | 44735 | BA_peak_2461 |
| DS113382 | 45920 | 46512 | BA_peak_2462 |
| DS113382 | 46817 | 51665 | BA_peak_2463 |
| DS113382 | 58358 | 59026 | BA_peak_2464 |
| DS113382 | 76736 | 77131 | BA_peak_2465 |
| DS113382 | 78018 | 78602 | BA_peak_2466 |
| DS113382 | 80026 | 80546 | BA_peak_2467 |
| DS113382 | 90199 | 90796 | BA_peak_2468 |
| DS113382 | 100481 | 100991 | BA_peak_2469 |
| DS113382 | 107982 | 108403 | BA_peak_2470 |
| DS113383 | 9119 | 10397 | BA_peak_2471 |
| DS113383 | 12434 | 13720 | BA_peak_2472 |
| DS113383 | 14152 | 14550 | BA_peak_2473 |
| DS113383 | 36670 | 37822 | BA_peak_2474 |
| DS113383 | 38241 | 38710 | BA_peak_2475 |
| DS113383 | 39228 | 39611 | BA_peak_2476 |

| | | | |
|---|---|---|---|
| DS113383 | 40231 | 40645 | BA_peak_2477 |
| DS113383 | 44332 | 44796 | BA_peak_2478 |
| DS113383 | 116325 | 117311 | BA_peak_2479 |
| DS113384 | 12458 | 13691 | BA_peak_2480 |
| DS113384 | 33469 | 33874 | BA_peak_2481 |
| DS113384 | 44342 | 45118 | BA_peak_2482 |
| DS113384 | 46358 | 47265 | BA_peak_2483 |
| DS113384 | 48533 | 49563 | BA_peak_2484 |
| DS113384 | 57132 | 58423 | BA_peak_2486 |
| DS113384 | 70672 | 72726 | BA_peak_2487 |
| DS113384 | 113682 | 114245 | BA_peak_2488 |
| DS113385 | 45057 | 45576 | BA_peak_2489 |
| DS113385 | 47740 | 48421 | BA_peak_2490 |
| DS113385 | 86457 | 87123 | BA_peak_2491 |
| DS113385 | 91523 | 92016 | BA_peak_2492 |
| DS113386 | 9843 | 11217 | BA_peak_2493 |
| DS113386 | 27378 | 28280 | BA_peak_2494 |
| DS113386 | 56822 | 58158 | BA_peak_2495 |
| DS113386 | 64366 | 64857 | BA_peak_2496 |
| DS113386 | 85833 | 86399 | BA_peak_2497 |
| DS113386 | 86963 | 87312 | BA_peak_2498 |
| DS113386 | 101934 | 102246 | BA_peak_2499 |
| DS113386 | 102663 | 103672 | BA_peak_2500 |
| DS113386 | 109677 | 110468 | BA_peak_2501 |
| DS113386 | 110781 | 111301 | BA_peak_2502 |
| DS113387 | 19402 | 19948 | BA_peak_2503 |
| DS113387 | 29259 | 29644 | BA_peak_2504 |
| DS113387 | 49728 | 50095 | BA_peak_2505 |
| DS113387 | 51031 | 51510 | BA_peak_2506 |
| DS113387 | 56454 | 57306 | BA_peak_2507 |
| DS113387 | 80955 | 81470 | BA_peak_2508 |
| DS113387 | 81907 | 82306 | BA_peak_2509 |
| DS113387 | 87161 | 87596 | BA_peak_2510 |
| DS113387 | 110318 | 110687 | BA_peak_2511 |
| DS113387 | 113049 | 113425 | BA_peak_2512 |
| DS113387 | 116510 | 116889 | BA_peak_2513 |
| DS113388 | 26957 | 27537 | BA_peak_2515 |
| DS113388 | 32011 | 32457 | BA_peak_2516 |
| DS113388 | 35338 | 36112 | BA_peak_2517 |
| DS113388 | 46364 | 47041 | BA_peak_2518 |
| DS113388 | 47530 | 47867 | BA_peak_2519 |
| DS113388 | 58208 | 58887 | BA_peak_2520 |
| DS113388 | 59586 | 59917 | BA_peak_2521 |
| DS113388 | 68137 | 68892 | BA_peak_2522 |
| DS113388 | 71574 | 71923 | BA_peak_2523 |
| DS113388 | 72616 | 73267 | BA_peak_2524 |
| DS113388 | 80655 | 81245 | BA_peak_2525 |

| | | | |
|---|---|---|---|
| DS113388 | 109618 | 110253 | BA_peak_2526 |
| DS113389 | 3465 | 4869 | BA_peak_2527 |
| DS113389 | 5187 | 5514 | BA_peak_2528 |
| DS113389 | 16564 | 17284 | BA_peak_2529 |
| DS113389 | 20667 | 22968 | BA_peak_2530 |
| DS113389 | 23953 | 25640 | BA_peak_2531 |
| DS113389 | 97224 | 97659 | BA_peak_2532 |
| DS113389 | 110323 | 111207 | BA_peak_2533 |
| DS113389 | 111539 | 111985 | BA_peak_2534 |
| DS113390 | 2486 | 2871 | BA_peak_2535 |
| DS113390 | 31860 | 36272 | BA_peak_2536 |
| DS113390 | 47402 | 48622 | BA_peak_2537 |
| DS113390 | 57220 | 57944 | BA_peak_2538 |
| DS113390 | 71103 | 73688 | BA_peak_2539 |
| DS113390 | 74141 | 75149 | BA_peak_2539 |
| DS113390 | 83555 | 84402 | BA_peak_2540 |
| DS113390 | 86052 | 86536 | BA_peak_2541 |
| DS113390 | 89319 | 89964 | BA_peak_2542 |
| DS113390 | 101649 | 102422 | BA_peak_2543 |
| DS113391 | 5634 | 9815 | BA_peak_2545 |
| DS113391 | 37581 | 38567 | BA_peak_2546 |
| DS113391 | 54451 | 55108 | BA_peak_2547 |
| DS113391 | 56647 | 58526 | BA_peak_2548 |
| DS113391 | 92035 | 92398 | BA_peak_2549 |
| DS113391 | 93056 | 93469 | BA_peak_2550 |
| DS113391 | 93816 | 94438 | BA_peak_2551 |
| DS113392 | 13636 | 14144 | BA_peak_2552 |
| DS113392 | 16916 | 17335 | BA_peak_2553 |
| DS113392 | 33075 | 34414 | BA_peak_2554 |
| DS113392 | 51199 | 52122 | BA_peak_2555 |
| DS113392 | 94564 | 95257 | BA_peak_2556 |
| DS113392 | 96261 | 96864 | BA_peak_2557 |
| DS113393 | 4549 | 6677 | BA_peak_2559 |
| DS113393 | 44373 | 45092 | BA_peak_2560 |
| DS113393 | 54825 | 55335 | BA_peak_2561 |
| DS113393 | 68339 | 68772 | BA_peak_2562 |
| DS113393 | 87175 | 87905 | BA_peak_2563 |
| DS113393 | 113813 | 114265 | BA_peak_2564 |
| DS113394 | 8533 | 9478 | BA_peak_2566 |
| DS113394 | 11343 | 11914 | BA_peak_2567 |
| DS113394 | 83393 | 83814 | BA_peak_2568 |
| DS113394 | 108062 | 111260 | BA_peak_2569 |
| DS113394 | 113350 | 114001 | BA_peak_2570 |
| DS113395 | 8058 | 8748 | BA_peak_2571 |
| DS113395 | 9106 | 9781 | BA_peak_2572 |
| DS113395 | 28126 | 28894 | BA_peak_2573 |
| DS113395 | 36447 | 39596 | BA_peak_2574 |

| | | | |
|---|---|---|---|
| DS113395 | 61174 | 64357 | BA_peak_2575 |
| DS113395 | 71151 | 71805 | BA_peak_2576 |
| DS113395 | 72519 | 73224 | BA_peak_2577 |
| DS113395 | 78747 | 79778 | BA_peak_2578 |
| DS113395 | 84859 | 85566 | BA_peak_2579 |
| DS113395 | 86528 | 86847 | BA_peak_2580 |
| DS113395 | 88663 | 89309 | BA_peak_2581 |
| DS113395 | 104360 | 104803 | BA_peak_2582 |
| DS113395 | 106549 | 108124 | BA_peak_2583 |
| DS113396 | 872 | 3453 | BA_peak_2584 |
| DS113396 | 27342 | 27649 | BA_peak_2585 |
| DS113396 | 36823 | 37355 | BA_peak_2586 |
| DS113396 | 77719 | 78372 | BA_peak_2587 |
| DS113397 | 1716 | 2214 | BA_peak_2588 |
| DS113397 | 11514 | 12320 | BA_peak_2589 |
| DS113397 | 48162 | 50414 | BA_peak_2590 |
| DS113397 | 52062 | 56079 | BA_peak_2591 |
| DS113397 | 65006 | 66104 | BA_peak_2592 |
| DS113397 | 70551 | 70939 | BA_peak_2593 |
| DS113397 | 81989 | 83184 | BA_peak_2594 |
| DS113398 | 27346 | 28279 | BA_peak_2595 |
| DS113398 | 29530 | 30681 | BA_peak_2596 |
| DS113398 | 43276 | 43896 | BA_peak_2597 |
| DS113398 | 54717 | 55364 | BA_peak_2598 |
| DS113398 | 56515 | 57131 | BA_peak_2599 |
| DS113399 | 6 | 471 | BA_peak_2601 |
| DS113399 | 6853 | 7703 | BA_peak_2602 |
| DS113399 | 41472 | 42107 | BA_peak_2603 |
| DS113399 | 67402 | 68284 | BA_peak_2604 |
| DS113399 | 92868 | 93219 | BA_peak_2605 |
| DS113399 | 106337 | 107949 | BA_peak_2606 |
| DS113400 | 12807 | 14376 | BA_peak_2607 |
| DS113400 | 28912 | 29285 | BA_peak_2608 |
| DS113400 | 47756 | 48688 | BA_peak_2609 |
| DS113400 | 49825 | 51007 | BA_peak_2610 |
| DS113400 | 52556 | 53002 | BA_peak_2611 |
| DS113400 | 60119 | 63281 | BA_peak_2612 |
| DS113400 | 81598 | 82491 | BA_peak_2613 |
| DS113400 | 82822 | 83885 | BA_peak_2614 |
| DS113400 | 84678 | 85042 | BA_peak_2615 |
| DS113400 | 85844 | 86183 | BA_peak_2616 |
| DS113401 | 8574 | 9010 | BA_peak_2617 |
| DS113401 | 9328 | 9927 | BA_peak_2618 |
| DS113401 | 10318 | 11002 | BA_peak_2619 |
| DS113401 | 29398 | 29780 | BA_peak_2620 |
| DS113401 | 30184 | 30514 | BA_peak_2621 |
| DS113401 | 37605 | 38591 | BA_peak_2622 |

| | | | |
|---|---|---|---|
| DS113401 | 54926 | 56568 | BA_peak_2623 |
| DS113401 | 93737 | 94354 | BA_peak_2624 |
| DS113401 | 95945 | 96401 | BA_peak_2625 |
| DS113401 | 97088 | 98091 | BA_peak_2626 |
| DS113401 | 100141 | 100714 | BA_peak_2627 |
| DS113402 | 557 | 1279 | BA_peak_2628 |
| DS113402 | 7599 | 8043 | BA_peak_2629 |
| DS113402 | 13651 | 14677 | BA_peak_2630 |
| DS113402 | 16503 | 17351 | BA_peak_2631 |
| DS113402 | 25003 | 28271 | BA_peak_2632 |
| DS113402 | 28699 | 29486 | BA_peak_2633 |
| DS113402 | 43927 | 44403 | BA_peak_2635 |
| DS113402 | 44886 | 45364 | BA_peak_2636 |
| DS113402 | 53913 | 54405 | BA_peak_2637 |
| DS113402 | 105960 | 106418 | BA_peak_2638 |
| DS113402 | 111456 | 112243 | BA_peak_2639 |
| DS113403 | 787 | 1430 | BA_peak_2640 |
| DS113403 | 12680 | 13062 | BA_peak_2641 |
| DS113403 | 19972 | 20612 | BA_peak_2642 |
| DS113403 | 25057 | 25816 | BA_peak_2643 |
| DS113403 | 31301 | 32443 | BA_peak_2644 |
| DS113403 | 93821 | 94989 | BA_peak_2646 |
| DS113403 | 96587 | 97110 | BA_peak_2647 |
| DS113404 | 18568 | 18997 | BA_peak_2648 |
| DS113404 | 37705 | 38427 | BA_peak_2649 |
| DS113404 | 95309 | 96067 | BA_peak_2651 |
| DS113404 | 106884 | 108067 | BA_peak_2652 |
| DS113404 | 108592 | 109183 | BA_peak_2653 |
| DS113405 | 8908 | 10053 | BA_peak_2654 |
| DS113405 | 39849 | 40658 | BA_peak_2655 |
| DS113405 | 96886 | 97239 | BA_peak_2656 |
| DS113405 | 103300 | 104011 | BA_peak_2657 |
| DS113406 | 60275 | 60889 | BA_peak_2658 |
| DS113406 | 105415 | 106311 | BA_peak_2660 |
| DS113407 | 204 | 899 | BA_peak_2661 |
| DS113407 | 54394 | 55646 | BA_peak_2662 |
| DS113407 | 73016 | 75056 | BA_peak_2663 |
| DS113408 | 51412 | 52580 | BA_peak_2664 |
| DS113408 | 62230 | 62878 | BA_peak_2665 |
| DS113408 | 63408 | 65082 | BA_peak_2666 |
| DS113408 | 77965 | 79726 | BA_peak_2667 |
| DS113408 | 97637 | 99672 | BA_peak_2668 |
| DS113409 | 10287 | 10823 | BA_peak_2669 |
| DS113409 | 12240 | 13055 | BA_peak_2670 |
| DS113409 | 13864 | 14788 | BA_peak_2671 |
| DS113409 | 42878 | 43616 | BA_peak_2672 |
| DS113409 | 46071 | 48471 | BA_peak_2673 |

| | | | |
|---|---|---|---|
| DS113409 | 50289 | 50761 | BA_peak_2674 |
| DS113409 | 60127 | 60497 | BA_peak_2675 |
| DS113409 | 91846 | 92517 | BA_peak_2676 |
| DS113409 | 93495 | 94100 | BA_peak_2677 |
| DS113409 | 103957 | 104370 | BA_peak_2678 |
| DS113409 | 104682 | 105684 | BA_peak_2679 |
| DS113409 | 106217 | 107327 | BA_peak_2680 |
| DS113409 | 107997 | 108411 | BA_peak_2681 |
| DS113409 | 108849 | 109260 | BA_peak_2682 |
| DS113410 | 3972 | 4550 | BA_peak_2683 |
| DS113410 | 13364 | 13830 | BA_peak_2684 |
| DS113410 | 27269 | 27911 | BA_peak_2685 |
| DS113410 | 47102 | 47784 | BA_peak_2686 |
| DS113410 | 55691 | 56135 | BA_peak_2687 |
| DS113410 | 98298 | 100412 | BA_peak_2688 |
| DS113410 | 108692 | 109016 | BA_peak_2689 |
| DS113411 | 51351 | 52286 | BA_peak_2690 |
| DS113411 | 58120 | 58725 | BA_peak_2691 |
| DS113411 | 61693 | 62093 | BA_peak_2692 |
| DS113411 | 62932 | 63657 | BA_peak_2693 |
| DS113411 | 64292 | 64953 | BA_peak_2694 |
| DS113411 | 65396 | 65788 | BA_peak_2695 |
| DS113411 | 66118 | 66991 | BA_peak_2696 |
| DS113411 | 67906 | 68319 | BA_peak_2697 |
| DS113411 | 77300 | 80096 | BA_peak_2698 |
| DS113411 | 85756 | 86079 | BA_peak_2699 |
| DS113411 | 86564 | 87037 | BA_peak_2700 |
| DS113412 | 353 | 1106 | BA_peak_2702 |
| DS113412 | 4700 | 5048 | BA_peak_2703 |
| DS113412 | 5747 | 6368 | BA_peak_2704 |
| DS113412 | 28722 | 29117 | BA_peak_2705 |
| DS113412 | 39477 | 40399 | BA_peak_2706 |
| DS113412 | 81864 | 82468 | BA_peak_2707 |
| DS113412 | 83303 | 83796 | BA_peak_2708 |
| DS113412 | 92884 | 93488 | BA_peak_2709 |
| DS113413 | 8530 | 8908 | BA_peak_2710 |
| DS113413 | 38892 | 39594 | BA_peak_2711 |
| DS113413 | 40035 | 40497 | BA_peak_2712 |
| DS113413 | 92989 | 94766 | BA_peak_2713 |
| DS113413 | 104932 | 105967 | BA_peak_2714 |
| DS113414 | 20967 | 21606 | BA_peak_2715 |
| DS113414 | 42871 | 44630 | BA_peak_2716 |
| DS113414 | 54369 | 55781 | BA_peak_2717 |
| DS113414 | 63431 | 64424 | BA_peak_2718 |
| DS113414 | 71237 | 71978 | BA_peak_2719 |
| DS113415 | 64891 | 65268 | BA_peak_2720 |
| DS113415 | 109003 | 109406 | BA_peak_2721 |

| | | | |
|---|---|---|---|
| DS113416 | 1653 | 2019 | BA_peak_2723 |
| DS113416 | 6210 | 7082 | BA_peak_2724 |
| DS113416 | 73846 | 74595 | BA_peak_2725 |
| DS113416 | 82410 | 82733 | BA_peak_2726 |
| DS113417 | 454 | 946 | BA_peak_2727 |
| DS113417 | 13710 | 14147 | BA_peak_2728 |
| DS113417 | 15905 | 17550 | BA_peak_2729 |
| DS113417 | 46333 | 46753 | BA_peak_2730 |
| DS113417 | 106291 | 106604 | BA_peak_2731 |
| DS113418 | 25344 | 25881 | BA_peak_2732 |
| DS113418 | 68289 | 69490 | BA_peak_2733 |
| DS113418 | 70417 | 70819 | BA_peak_2734 |
| DS113418 | 71527 | 72161 | BA_peak_2735 |
| DS113419 | 5975 | 6426 | BA_peak_2736 |
| DS113419 | 26218 | 26997 | BA_peak_2737 |
| DS113419 | 36685 | 37695 | BA_peak_2738 |
| DS113419 | 47779 | 51108 | BA_peak_2739 |
| DS113419 | 53683 | 54503 | BA_peak_2740 |
| DS113419 | 73649 | 74388 | BA_peak_2741 |
| DS113419 | 75069 | 75414 | BA_peak_2742 |
| DS113419 | 88576 | 89012 | BA_peak_2743 |
| DS113419 | 93393 | 94190 | BA_peak_2744 |
| DS113420 | 4919 | 5304 | BA_peak_2745 |
| DS113420 | 25987 | 28661 | BA_peak_2746 |
| DS113421 | 70079 | 71797 | BA_peak_2748 |
| DS113421 | 95891 | 96293 | BA_peak_2749 |
| DS113422 | 7723 | 8263 | BA_peak_2750 |
| DS113422 | 23381 | 24145 | BA_peak_2751 |
| DS113422 | 43162 | 44091 | BA_peak_2752 |
| DS113422 | 50304 | 50694 | BA_peak_2753 |
| DS113422 | 103228 | 104096 | BA_peak_2754 |
| DS113422 | 105224 | 105891 | BA_peak_2755 |
| DS113423 | 2923 | 3717 | BA_peak_2756 |
| DS113423 | 49015 | 50543 | BA_peak_2757 |
| DS113424 | 552 | 950 | BA_peak_2758 |
| DS113424 | 15467 | 15781 | BA_peak_2759 |
| DS113424 | 80706 | 81483 | BA_peak_2760 |
| DS113425 | 1057 | 1397 | BA_peak_2761 |
| DS113425 | 22848 | 24455 | BA_peak_2762 |
| DS113425 | 32298 | 32750 | BA_peak_2763 |
| DS113425 | 59211 | 59721 | BA_peak_2764 |
| DS113426 | 88703 | 90108 | BA_peak_2765 |
| DS113427 | 14808 | 15308 | BA_peak_2767 |
| DS113427 | 18533 | 18917 | BA_peak_2768 |
| DS113427 | 20542 | 20864 | BA_peak_2769 |
| DS113427 | 38494 | 39830 | BA_peak_2770 |
| DS113427 | 40550 | 41182 | BA_peak_2771 |

| | | | |
|---|---|---|---|
| DS113427 | 46671 | 47490 | BA_peak_2772 |
| DS113427 | 59814 | 60472 | BA_peak_2773 |
| DS113427 | 61127 | 61516 | BA_peak_2774 |
| DS113427 | 62053 | 62482 | BA_peak_2775 |
| DS113427 | 79280 | 79648 | BA_peak_2776 |
| DS113427 | 93718 | 94502 | BA_peak_2777 |
| DS113427 | 100113 | 100704 | BA_peak_2778 |
| DS113428 | 10827 | 14060 | BA_peak_2779 |
| DS113428 | 14446 | 15275 | BA_peak_2780 |
| DS113428 | 67819 | 68563 | BA_peak_2781 |
| DS113428 | 73334 | 73700 | BA_peak_2782 |
| DS113428 | 80637 | 81364 | BA_peak_2783 |
| DS113428 | 97800 | 98144 | BA_peak_2784 |
| DS113428 | 98648 | 99981 | BA_peak_2785 |
| DS113429 | 33121 | 34305 | BA_peak_2786 |
| DS113429 | 35412 | 36426 | BA_peak_2787 |
| DS113429 | 73267 | 74551 | BA_peak_2788 |
| DS113429 | 83392 | 84316 | BA_peak_2789 |
| DS113429 | 98362 | 98714 | BA_peak_2790 |
| DS113430 | 3882 | 4263 | BA_peak_2791 |
| DS113430 | 5144 | 5518 | BA_peak_2792 |
| DS113430 | 9314 | 9808 | BA_peak_2794 |
| DS113430 | 21172 | 21623 | BA_peak_2795 |
| DS113430 | 22191 | 22564 | BA_peak_2796 |
| DS113430 | 34352 | 34859 | BA_peak_2798 |
| DS113430 | 37876 | 38705 | BA_peak_2799 |
| DS113430 | 54180 | 55090 | BA_peak_2800 |
| DS113430 | 55652 | 57216 | BA_peak_2801 |
| DS113430 | 57812 | 58359 | BA_peak_2802 |
| DS113430 | 63551 | 65277 | BA_peak_2803 |
| DS113430 | 66751 | 67354 | BA_peak_2804 |
| DS113430 | 70160 | 70491 | BA_peak_2805 |
| DS113430 | 78108 | 78485 | BA_peak_2806 |
| DS113431 | 8115 | 8876 | BA_peak_2807 |
| DS113431 | 10578 | 11093 | BA_peak_2808 |
| DS113431 | 50583 | 51428 | BA_peak_2809 |
| DS113431 | 85208 | 86227 | BA_peak_2810 |
| DS113431 | 87877 | 88271 | BA_peak_2811 |
| DS113431 | 92526 | 92877 | BA_peak_2813 |
| DS113432 | 7159 | 7495 | BA_peak_2814 |
| DS113432 | 40427 | 41905 | BA_peak_2815 |
| DS113432 | 44729 | 45145 | BA_peak_2816 |
| DS113432 | 46304 | 46852 | BA_peak_2817 |
| DS113432 | 57657 | 58068 | BA_peak_2818 |
| DS113433 | 3216 | 3596 | BA_peak_2819 |
| DS113433 | 9327 | 9979 | BA_peak_2820 |
| DS113433 | 29109 | 29658 | BA_peak_2821 |

| | | | |
|---|---|---|---|
| DS113433 | 31982 | 32318 | BA_peak_2823 |
| DS113433 | 34262 | 34701 | BA_peak_2824 |
| DS113433 | 58672 | 59336 | BA_peak_2825 |
| DS113433 | 60155 | 60644 | BA_peak_2826 |
| DS113433 | 61239 | 61906 | BA_peak_2827 |
| DS113433 | 62919 | 63866 | BA_peak_2828 |
| DS113434 | 20377 | 21117 | BA_peak_2829 |
| DS113434 | 29050 | 29429 | BA_peak_2830 |
| DS113434 | 50644 | 51595 | BA_peak_2832 |
| DS113434 | 92709 | 93526 | BA_peak_2833 |
| DS113434 | 94502 | 95572 | BA_peak_2834 |
| DS113434 | 102372 | 102841 | BA_peak_2835 |
| DS113435 | 7709 | 8768 | BA_peak_2836 |
| DS113435 | 19229 | 21758 | BA_peak_2837 |
| DS113435 | 22408 | 22953 | BA_peak_2838 |
| DS113435 | 23880 | 24479 | BA_peak_2839 |
| DS113435 | 26724 | 27850 | BA_peak_2840 |
| DS113435 | 29090 | 29481 | BA_peak_2841 |
| DS113435 | 61084 | 61760 | BA_peak_2842 |
| DS113435 | 70501 | 71020 | BA_peak_2843 |
| DS113436 | 1266 | 1879 | BA_peak_2844 |
| DS113436 | 2724 | 3230 | BA_peak_2845 |
| DS113436 | 50616 | 51108 | BA_peak_2846 |
| DS113436 | 52204 | 52654 | BA_peak_2847 |
| DS113436 | 53105 | 53935 | BA_peak_2848 |
| DS113436 | 55025 | 56637 | BA_peak_2849 |
| DS113436 | 71197 | 72754 | BA_peak_2850 |
| DS113436 | 89580 | 90165 | BA_peak_2851 |
| DS113437 | 15321 | 16515 | BA_peak_2852 |
| DS113437 | 22070 | 23513 | BA_peak_2853 |
| DS113437 | 25154 | 26034 | BA_peak_2854 |
| DS113437 | 27190 | 27836 | BA_peak_2855 |
| DS113437 | 42102 | 42797 | BA_peak_2856 |
| DS113437 | 68777 | 69473 | BA_peak_2857 |
| DS113437 | 91928 | 92464 | BA_peak_2858 |
| DS113437 | 94868 | 95490 | BA_peak_2859 |
| DS113437 | 100150 | 100573 | BA_peak_2860 |
| DS113438 | 26726 | 27444 | BA_peak_2861 |
| DS113438 | 27786 | 28290 | BA_peak_2861 |
| DS113438 | 30185 | 30774 | BA_peak_2862 |
| DS113438 | 36970 | 37798 | BA_peak_2863 |
| DS113438 | 40718 | 41323 | BA_peak_2864 |
| DS113438 | 81438 | 82198 | BA_peak_2865 |
| DS113439 | 88 | 1096 | BA_peak_2866 |
| DS113439 | 8141 | 8978 | BA_peak_2867 |
| DS113439 | 15451 | 15804 | BA_peak_2868 |
| DS113439 | 16189 | 17284 | BA_peak_2869 |

| | | | |
|---|---|---|---|
| DS113439 | 62973 | 63407 | BA_peak_2870 |
| DS113440 | 1096 | 1585 | BA_peak_2871 |
| DS113440 | 18707 | 19407 | BA_peak_2872 |
| DS113440 | 20389 | 22301 | BA_peak_2873 |
| DS113440 | 22721 | 23155 | BA_peak_2874 |
| DS113440 | 25772 | 26165 | BA_peak_2875 |
| DS113440 | 35362 | 36082 | BA_peak_2876 |
| DS113440 | 50576 | 52446 | BA_peak_2877 |
| DS113440 | 80471 | 82480 | BA_peak_2878 |
| DS113440 | 93742 | 94180 | BA_peak_2879 |
| DS113440 | 96410 | 97341 | BA_peak_2880 |
| DS113440 | 98378 | 98932 | BA_peak_2881 |
| DS113440 | 99466 | 100753 | BA_peak_2882 |
| DS113441 | 60526 | 60947 | BA_peak_2883 |
| DS113441 | 65196 | 65640 | BA_peak_2884 |
| DS113442 | 36498 | 37268 | BA_peak_2885 |
| DS113442 | 43725 | 44111 | BA_peak_2886 |
| DS113442 | 45963 | 46345 | BA_peak_2887 |
| DS113442 | 49635 | 50406 | BA_peak_2888 |
| DS113442 | 72264 | 72814 | BA_peak_2889 |
| DS113442 | 86171 | 88245 | BA_peak_2890 |
| DS113442 | 96714 | 97512 | BA_peak_2891 |
| DS113442 | 98688 | 99938 | BA_peak_2892 |
| DS113442 | 102294 | 102856 | BA_peak_2894 |
| DS113443 | 30510 | 30868 | BA_peak_2895 |
| DS113443 | 37314 | 37812 | BA_peak_2896 |
| DS113443 | 40750 | 41315 | BA_peak_2897 |
| DS113443 | 43247 | 45981 | BA_peak_2898 |
| DS113443 | 47297 | 48274 | BA_peak_2899 |
| DS113443 | 54098 | 54634 | BA_peak_2900 |
| DS113443 | 55040 | 55421 | BA_peak_2901 |
| DS113443 | 71430 | 71879 | BA_peak_2902 |
| DS113443 | 74987 | 75716 | BA_peak_2903 |
| DS113443 | 91041 | 91429 | BA_peak_2904 |
| DS113444 | 21810 | 22511 | BA_peak_2905 |
| DS113444 | 40921 | 42441 | BA_peak_2906 |
| DS113444 | 43016 | 43487 | BA_peak_2907 |
| DS113444 | 76340 | 76880 | BA_peak_2908 |
| DS113445 | 3804 | 5008 | BA_peak_2909 |
| DS113445 | 6575 | 8409 | BA_peak_2910 |
| DS113445 | 12131 | 12789 | BA_peak_2911 |
| DS113445 | 50736 | 51914 | BA_peak_2912 |
| DS113445 | 62092 | 62695 | BA_peak_2913 |
| DS113445 | 80867 | 81289 | BA_peak_2914 |
| DS113445 | 97125 | 98023 | BA_peak_2915 |
| DS113446 | 11640 | 12203 | BA_peak_2916 |
| DS113446 | 17568 | 18056 | BA_peak_2917 |

| | | | |
|---|---|---|---|
| DS113446 | 23766 | 24093 | BA_peak_2918 |
| DS113446 | 54972 | 55486 | BA_peak_2919 |
| DS113446 | 90711 | 91093 | BA_peak_2920 |
| DS113447 | 26627 | 27482 | BA_peak_2921 |
| DS113447 | 29620 | 30841 | BA_peak_2922 |
| DS113447 | 31421 | 31999 | BA_peak_2923 |
| DS113447 | 33989 | 34388 | BA_peak_2924 |
| DS113448 | 4118 | 4563 | BA_peak_2925 |
| DS113448 | 9998 | 10744 | BA_peak_2926 |
| DS113448 | 13593 | 14054 | BA_peak_2927 |
| DS113448 | 17216 | 18560 | BA_peak_2928 |
| DS113448 | 31598 | 32245 | BA_peak_2930 |
| DS113448 | 36425 | 36828 | BA_peak_2931 |
| DS113448 | 50411 | 51171 | BA_peak_2932 |
| DS113448 | 76959 | 77400 | BA_peak_2933 |
| DS113448 | 83144 | 83883 | BA_peak_2934 |
| DS113449 | 29061 | 29479 | BA_peak_2935 |
| DS113449 | 55383 | 55996 | BA_peak_2937 |
| DS113449 | 62762 | 63117 | BA_peak_2938 |
| DS113449 | 84793 | 86779 | BA_peak_2939 |
| DS113449 | 91708 | 92487 | BA_peak_2940 |
| DS113450 | 9022 | 9488 | BA_peak_2942 |
| DS113450 | 22337 | 22887 | BA_peak_2943 |
| DS113450 | 23302 | 23918 | BA_peak_2944 |
| DS113450 | 27917 | 28724 | BA_peak_2945 |
| DS113450 | 47381 | 50329 | BA_peak_2946 |
| DS113450 | 54021 | 54565 | BA_peak_2947 |
| DS113450 | 63320 | 63895 | BA_peak_2948 |
| DS113450 | 82884 | 83358 | BA_peak_2949 |
| DS113450 | 83918 | 84694 | BA_peak_2950 |
| DS113450 | 85061 | 86302 | BA_peak_2951 |
| DS113451 | 65975 | 68157 | BA_peak_2952 |
| DS113451 | 79981 | 81323 | BA_peak_2953 |
| DS113451 | 88599 | 89452 | BA_peak_2954 |
| DS113452 | 229 | 592 | BA_peak_2955 |
| DS113452 | 2901 | 3376 | BA_peak_2956 |
| DS113452 | 5266 | 5872 | BA_peak_2957 |
| DS113452 | 16807 | 17574 | BA_peak_2958 |
| DS113452 | 34767 | 35407 | BA_peak_2959 |
| DS113452 | 39018 | 43095 | BA_peak_2960 |
| DS113452 | 72625 | 74422 | BA_peak_2961 |
| DS113452 | 74819 | 76927 | BA_peak_2962 |
| DS113452 | 80916 | 82543 | BA_peak_2963 |
| DS113452 | 87189 | 87957 | BA_peak_2964 |
| DS113453 | 14212 | 14592 | BA_peak_2965 |
| DS113453 | 41696 | 42604 | BA_peak_2966 |
| DS113453 | 44386 | 47750 | BA_peak_2967 |

| | | | |
|---|---|---|---|
| DS113453 | 88152 | 88522 | BA_peak_2968 |
| DS113454 | 2101 | 3160 | BA_peak_2969 |
| DS113454 | 13061 | 13593 | BA_peak_2970 |
| DS113454 | 18503 | 18846 | BA_peak_2971 |
| DS113454 | 22516 | 23226 | BA_peak_2972 |
| DS113454 | 23539 | 25249 | BA_peak_2973 |
| DS113454 | 26726 | 27068 | BA_peak_2974 |
| DS113454 | 46503 | 47032 | BA_peak_2975 |
| DS113454 | 86388 | 87040 | BA_peak_2976 |
| DS113454 | 88207 | 88965 | BA_peak_2977 |
| DS113455 | 33079 | 34068 | BA_peak_2978 |
| DS113455 | 56347 | 56736 | BA_peak_2979 |
| DS113455 | 58417 | 59678 | BA_peak_2980 |
| DS113455 | 83822 | 84659 | BA_peak_2981 |
| DS113455 | 85761 | 86497 | BA_peak_2982 |
| DS113455 | 93364 | 95338 | BA_peak_2983 |
| DS113456 | 32097 | 32711 | BA_peak_2984 |
| DS113456 | 35445 | 35863 | BA_peak_2985 |
| DS113456 | 36310 | 38426 | BA_peak_2985 |
| DS113456 | 55780 | 56392 | BA_peak_2987 |
| DS113456 | 71556 | 71941 | BA_peak_2988 |
| DS113456 | 93796 | 94397 | BA_peak_2989 |
| DS113457 | 18816 | 19162 | BA_peak_2990 |
| DS113457 | 53779 | 54153 | BA_peak_2991 |
| DS113457 | 60283 | 60924 | BA_peak_2992 |
| DS113458 | 9789 | 10183 | BA_peak_2993 |
| DS113458 | 43448 | 43818 | BA_peak_2995 |
| DS113458 | 84856 | 85224 | BA_peak_2996 |
| DS113458 | 94259 | 94617 | BA_peak_2997 |
| DS113459 | 6730 | 7308 | BA_peak_2998 |
| DS113459 | 8527 | 8984 | BA_peak_2999 |
| DS113459 | 12427 | 13562 | BA_peak_3000 |
| DS113459 | 20730 | 21867 | BA_peak_3001 |
| DS113459 | 22182 | 23434 | BA_peak_3002 |
| DS113459 | 50514 | 50824 | BA_peak_3003 |
| DS113460 | 6985 | 7293 | BA_peak_3004 |
| DS113460 | 21608 | 22292 | BA_peak_3006 |
| DS113460 | 23500 | 23937 | BA_peak_3007 |
| DS113460 | 57483 | 59493 | BA_peak_3008 |
| DS113460 | 66829 | 67657 | BA_peak_3009 |
| DS113460 | 71649 | 72261 | BA_peak_3010 |
| DS113460 | 83736 | 84202 | BA_peak_3011 |
| DS113461 | 4324 | 5484 | BA_peak_3012 |
| DS113461 | 10399 | 11750 | BA_peak_3013 |
| DS113461 | 15776 | 17860 | BA_peak_3014 |
| DS113461 | 34446 | 35252 | BA_peak_3015 |
| DS113461 | 55281 | 57042 | BA_peak_3016 |

| | | | |
|---|---|---|---|
| DS113461 | 66024 | 66471 | BA_peak_3017 |
| DS113461 | 75470 | 76588 | BA_peak_3018 |
| DS113461 | 85002 | 85539 | BA_peak_3019 |
| DS113462 | 3544 | 4120 | BA_peak_3020 |
| DS113462 | 4485 | 4828 | BA_peak_3021 |
| DS113462 | 82566 | 82973 | BA_peak_3022 |
| DS113463 | 17215 | 18016 | BA_peak_3023 |
| DS113463 | 75629 | 76096 | BA_peak_3024 |
| DS113463 | 78635 | 79050 | BA_peak_3025 |
| DS113463 | 79712 | 80458 | BA_peak_3026 |
| DS113463 | 88179 | 88644 | BA_peak_3027 |
| DS113464 | 37183 | 37734 | BA_peak_3028 |
| DS113464 | 59256 | 59754 | BA_peak_3029 |
| DS113464 | 65394 | 66010 | BA_peak_3030 |
| DS113465 | 5844 | 6197 | BA_peak_3031 |
| DS113465 | 59377 | 63628 | BA_peak_3032 |
| DS113466 | 4768 | 6242 | BA_peak_3033 |
| DS113466 | 30539 | 31920 | BA_peak_3034 |
| DS113466 | 43141 | 44268 | BA_peak_3035 |
| DS113466 | 62917 | 63498 | BA_peak_3036 |
| DS113466 | 64720 | 65118 | BA_peak_3037 |
| DS113466 | 84599 | 85071 | BA_peak_3038 |
| DS113466 | 93376 | 93685 | BA_peak_3039 |
| DS113467 | 2142 | 3250 | BA_peak_3040 |
| DS113467 | 12838 | 13575 | BA_peak_3041 |
| DS113467 | 17411 | 20751 | BA_peak_3042 |
| DS113467 | 51010 | 54141 | BA_peak_3045 |
| DS113467 | 65855 | 66745 | BA_peak_3046 |
| DS113468 | 3411 | 5597 | BA_peak_3048 |
| DS113468 | 6440 | 6813 | BA_peak_3049 |
| DS113468 | 16014 | 16755 | BA_peak_3050 |
| DS113468 | 17412 | 18169 | BA_peak_3051 |
| DS113468 | 27070 | 27647 | BA_peak_3052 |
| DS113468 | 31122 | 31737 | BA_peak_3053 |
| DS113468 | 32775 | 33475 | BA_peak_3054 |
| DS113468 | 44470 | 44995 | BA_peak_3055 |
| DS113468 | 46587 | 48059 | BA_peak_3056 |
| DS113468 | 49364 | 50398 | BA_peak_3057 |
| DS113468 | 65123 | 65496 | BA_peak_3058 |
| DS113468 | 66592 | 66971 | BA_peak_3059 |
| DS113469 | 40123 | 40491 | BA_peak_3061 |
| DS113469 | 48356 | 48743 | BA_peak_3062 |
| DS113469 | 50424 | 52441 | BA_peak_3063 |
| DS113469 | 53974 | 54845 | BA_peak_3064 |
| DS113469 | 55573 | 56136 | BA_peak_3065 |
| DS113469 | 73103 | 73956 | BA_peak_3066 |
| DS113469 | 80563 | 81059 | BA_peak_3067 |

| | | | |
|---|---|---|---|
| DS113469 | 83034 | 84630 | BA_peak_3068 |
| DS113470 | 32725 | 33344 | BA_peak_3069 |
| DS113470 | 51437 | 52481 | BA_peak_3070 |
| DS113470 | 85396 | 85861 | BA_peak_3071 |
| DS113470 | 88704 | 89268 | BA_peak_3072 |
| DS113470 | 90048 | 90437 | BA_peak_3073 |
| DS113471 | 69986 | 70467 | BA_peak_3075 |
| DS113471 | 70773 | 71229 | BA_peak_3076 |
| DS113471 | 80241 | 81130 | BA_peak_3077 |
| DS113471 | 91410 | 92319 | BA_peak_3078 |
| DS113472 | 3333 | 3978 | BA_peak_3079 |
| DS113472 | 8256 | 8585 | BA_peak_3080 |
| DS113472 | 24899 | 25239 | BA_peak_3082 |
| DS113472 | 88422 | 89103 | BA_peak_3083 |
| DS113473 | 10152 | 11024 | BA_peak_3084 |
| DS113473 | 66463 | 67207 | BA_peak_3085 |
| DS113474 | 3853 | 4228 | BA_peak_3086 |
| DS113474 | 4892 | 5941 | BA_peak_3087 |
| DS113474 | 14250 | 14586 | BA_peak_3088 |
| DS113474 | 45579 | 45928 | BA_peak_3090 |
| DS113474 | 90309 | 90903 | BA_peak_3091 |
| DS113475 | 852 | 1171 | BA_peak_3092 |
| DS113475 | 28261 | 28895 | BA_peak_3094 |
| DS113475 | 41643 | 42173 | BA_peak_3095 |
| DS113475 | 42665 | 44538 | BA_peak_3096 |
| DS113475 | 61312 | 63421 | BA_peak_3097 |
| DS113475 | 79845 | 80327 | BA_peak_3099 |
| DS113476 | 13628 | 14247 | BA_peak_3100 |
| DS113476 | 58595 | 59443 | BA_peak_3101 |
| DS113476 | 77553 | 79082 | BA_peak_3102 |
| DS113477 | 7879 | 9302 | BA_peak_3103 |
| DS113477 | 38327 | 40547 | BA_peak_3104 |
| DS113477 | 41005 | 41323 | BA_peak_3105 |
| DS113477 | 49957 | 50723 | BA_peak_3106 |
| DS113477 | 51558 | 51996 | BA_peak_3107 |
| DS113477 | 62580 | 63565 | BA_peak_3108 |
| DS113477 | 92460 | 93144 | BA_peak_3109 |
| DS113478 | 86334 | 86886 | BA_peak_3110 |
| DS113479 | 2930 | 4341 | BA_peak_3111 |
| DS113479 | 6675 | 7069 | BA_peak_3112 |
| DS113479 | 30252 | 30941 | BA_peak_3113 |
| DS113479 | 56138 | 57360 | BA_peak_3114 |
| DS113479 | 59890 | 60839 | BA_peak_3115 |
| DS113479 | 68248 | 69726 | BA_peak_3116 |
| DS113480 | 6067 | 7550 | BA_peak_3117 |
| DS113480 | 37580 | 38254 | BA_peak_3118 |
| DS113481 | 8644 | 9457 | BA_peak_3120 |

| | | | |
|---|---|---|---|
| DS113481 | 31510 | 32883 | BA_peak_3121 |
| DS113481 | 63416 | 65072 | BA_peak_3122 |
| DS113481 | 81149 | 81939 | BA_peak_3123 |
| DS113482 | 4161 | 4938 | BA_peak_3124 |
| DS113482 | 21528 | 24737 | BA_peak_3125 |
| DS113482 | 49010 | 49614 | BA_peak_3126 |
| DS113482 | 63049 | 64060 | BA_peak_3127 |
| DS113482 | 68110 | 68970 | BA_peak_3128 |
| DS113482 | 83364 | 84056 | BA_peak_3129 |
| DS113482 | 87815 | 89130 | BA_peak_3130 |
| DS113483 | 25628 | 26199 | BA_peak_3131 |
| DS113483 | 27053 | 27610 | BA_peak_3132 |
| DS113483 | 54408 | 55698 | BA_peak_3133 |
| DS113484 | 18548 | 19289 | BA_peak_3134 |
| DS113484 | 41630 | 42417 | BA_peak_3135 |
| DS113484 | 48600 | 49095 | BA_peak_3136 |
| DS113484 | 50644 | 51117 | BA_peak_3137 |
| DS113484 | 64271 | 65035 | BA_peak_3138 |
| DS113484 | 69680 | 70943 | BA_peak_3139 |
| DS113484 | 74265 | 75224 | BA_peak_3140 |
| DS113484 | 78706 | 80068 | BA_peak_3141 |
| DS113485 | 2634 | 3783 | BA_peak_3142 |
| DS113485 | 10453 | 11082 | BA_peak_3143 |
| DS113485 | 38521 | 40551 | BA_peak_3144 |
| DS113485 | 41503 | 42586 | BA_peak_3145 |
| DS113485 | 43526 | 43976 | BA_peak_3146 |
| DS113485 | 52757 | 53402 | BA_peak_3147 |
| DS113485 | 54635 | 54987 | BA_peak_3148 |
| DS113485 | 59565 | 60210 | BA_peak_3149 |
| DS113485 | 66541 | 69823 | BA_peak_3150 |
| DS113485 | 70730 | 71611 | BA_peak_3151 |
| DS113485 | 79449 | 79850 | BA_peak_3152 |
| DS113485 | 87147 | 87671 | BA_peak_3153 |
| DS113485 | 88745 | 89818 | BA_peak_3154 |
| DS113486 | 5315 | 5833 | BA_peak_3155 |
| DS113486 | 13126 | 13537 | BA_peak_3156 |
| DS113486 | 20676 | 21247 | BA_peak_3157 |
| DS113486 | 56221 | 56721 | BA_peak_3158 |
| DS113486 | 64402 | 64815 | BA_peak_3159 |
| DS113486 | 72337 | 73082 | BA_peak_3160 |
| DS113486 | 74038 | 74930 | BA_peak_3161 |
| DS113487 | 17957 | 19890 | BA_peak_3162 |
| DS113488 | 6206 | 6608 | BA_peak_3163 |
| DS113488 | 12363 | 13014 | BA_peak_3164 |
| DS113488 | 22210 | 22717 | BA_peak_3165 |
| DS113488 | 41755 | 42528 | BA_peak_3166 |
| DS113488 | 81560 | 82101 | BA_peak_3167 |

| | | | |
|---|---|---|---|
| DS113489 | 4539 | 5095 | BA_peak_3168 |
| DS113489 | 10765 | 11236 | BA_peak_3169 |
| DS113489 | 35089 | 35799 | BA_peak_3170 |
| DS113489 | 71088 | 72313 | BA_peak_3171 |
| DS113490 | 2134 | 3067 | BA_peak_3172 |
| DS113490 | 8436 | 10436 | BA_peak_3173 |
| DS113490 | 91193 | 92478 | BA_peak_3174 |
| DS113491 | 5672 | 6065 | BA_peak_3175 |
| DS113491 | 6426 | 6849 | BA_peak_3176 |
| DS113491 | 14524 | 15498 | BA_peak_3177 |
| DS113491 | 55591 | 56001 | BA_peak_3178 |
| DS113491 | 58347 | 58710 | BA_peak_3179 |
| DS113492 | 32 | 1951 | BA_peak_3180 |
| DS113492 | 7650 | 8776 | BA_peak_3181 |
| DS113492 | 36666 | 37126 | BA_peak_3182 |
| DS113492 | 71295 | 72460 | BA_peak_3183 |
| DS113492 | 74156 | 75045 | BA_peak_3184 |
| DS113492 | 75811 | 76976 | BA_peak_3185 |
| DS113492 | 79789 | 80428 | BA_peak_3186 |
| DS113492 | 81919 | 82675 | BA_peak_3187 |
| DS113492 | 90045 | 90427 | BA_peak_3188 |
| DS113492 | 91715 | 92728 | BA_peak_3189 |
| DS113494 | 9806 | 10407 | BA_peak_3190 |
| DS113494 | 11744 | 14377 | BA_peak_3191 |
| DS113494 | 14796 | 15475 | BA_peak_3192 |
| DS113494 | 19995 | 20577 | BA_peak_3193 |
| DS113494 | 23835 | 24661 | BA_peak_3194 |
| DS113494 | 37941 | 38705 | BA_peak_3195 |
| DS113494 | 39221 | 39771 | BA_peak_3196 |
| DS113494 | 40099 | 40450 | BA_peak_3197 |
| DS113494 | 54212 | 54618 | BA_peak_3198 |
| DS113494 | 55078 | 55632 | BA_peak_3199 |
| DS113494 | 59595 | 60217 | BA_peak_3200 |
| DS113495 | 34269 | 34596 | BA_peak_3201 |
| DS113495 | 38879 | 39662 | BA_peak_3202 |
| DS113495 | 40975 | 41278 | BA_peak_3203 |
| DS113495 | 41690 | 42575 | BA_peak_3204 |
| DS113495 | 53745 | 54197 | BA_peak_3205 |
| DS113495 | 59130 | 60260 | BA_peak_3206 |
| DS113495 | 65455 | 67071 | BA_peak_3207 |
| DS113495 | 72777 | 74019 | BA_peak_3210 |
| DS113495 | 83010 | 83976 | BA_peak_3211 |
| DS113495 | 84405 | 85168 | BA_peak_3212 |
| DS113496 | 8774 | 9371 | BA_peak_3213 |
| DS113496 | 12573 | 14677 | BA_peak_3214 |
| DS113496 | 20257 | 21567 | BA_peak_3215 |
| DS113496 | 22592 | 23322 | BA_peak_3216 |

| | | | |
|---|---|---|---|
| DS113496 | 29736 | 30104 | BA_peak_3217 |
| DS113496 | 30498 | 30914 | BA_peak_3218 |
| DS113496 | 37930 | 38513 | BA_peak_3219 |
| DS113496 | 44517 | 47663 | BA_peak_3220 |
| DS113496 | 48255 | 48890 | BA_peak_3221 |
| DS113496 | 51011 | 52299 | BA_peak_3222 |
| DS113496 | 82202 | 83326 | BA_peak_3223 |
| DS113496 | 84672 | 85261 | BA_peak_3224 |
| DS113497 | 27667 | 28253 | BA_peak_3226 |
| DS113497 | 29913 | 30685 | BA_peak_3227 |
| DS113497 | 55594 | 56080 | BA_peak_3228 |
| DS113497 | 56409 | 57096 | BA_peak_3229 |
| DS113497 | 62929 | 64193 | BA_peak_3230 |
| DS113497 | 78086 | 79806 | BA_peak_3231 |
| DS113497 | 81526 | 82973 | BA_peak_3232 |
| DS113498 | 3111 | 3440 | BA_peak_3233 |
| DS113498 | 16568 | 17035 | BA_peak_3234 |
| DS113498 | 23018 | 23588 | BA_peak_3235 |
| DS113498 | 38207 | 38551 | BA_peak_3237 |
| DS113498 | 49299 | 49772 | BA_peak_3238 |
| DS113498 | 52760 | 53660 | BA_peak_3239 |
| DS113498 | 63168 | 63806 | BA_peak_3240 |
| DS113498 | 72643 | 73664 | BA_peak_3242 |
| DS113498 | 83181 | 83527 | BA_peak_3243 |
| DS113499 | 58079 | 58860 | BA_peak_3244 |
| DS113500 | 12807 | 13216 | BA_peak_3245 |
| DS113500 | 19822 | 20279 | BA_peak_3246 |
| DS113500 | 32057 | 32705 | BA_peak_3247 |
| DS113500 | 45815 | 47834 | BA_peak_3248 |
| DS113500 | 56574 | 56907 | BA_peak_3251 |
| DS113500 | 66188 | 66639 | BA_peak_3252 |
| DS113500 | 73152 | 74535 | BA_peak_3253 |
| DS113501 | 4474 | 5578 | BA_peak_3255 |
| DS113501 | 69526 | 70285 | BA_peak_3256 |
| DS113501 | 70600 | 71236 | BA_peak_3257 |
| DS113501 | 71703 | 72925 | BA_peak_3258 |
| DS113501 | 83233 | 84658 | BA_peak_3259 |
| DS113502 | 4744 | 5508 | BA_peak_3261 |
| DS113502 | 22957 | 24703 | BA_peak_3262 |
| DS113502 | 58589 | 59453 | BA_peak_3263 |
| DS113502 | 60554 | 62136 | BA_peak_3264 |
| DS113502 | 64326 | 65932 | BA_peak_3265 |
| DS113502 | 74394 | 74798 | BA_peak_3266 |
| DS113503 | 22773 | 23269 | BA_peak_3267 |
| DS113503 | 42686 | 43088 | BA_peak_3268 |
| DS113503 | 63328 | 64559 | BA_peak_3269 |
| DS113503 | 65506 | 65838 | BA_peak_3270 |

| | | | |
|---|---|---|---|
| DS113503 | 85766 | 87270 | BA_peak_3271 |
| DS113504 | 15780 | 16160 | BA_peak_3272 |
| DS113504 | 29890 | 31028 | BA_peak_3273 |
| DS113504 | 31462 | 32126 | BA_peak_3274 |
| DS113504 | 72296 | 73149 | BA_peak_3275 |
| DS113504 | 81339 | 82128 | BA_peak_3276 |
| DS113505 | 7050 | 8592 | BA_peak_3277 |
| DS113505 | 9582 | 10454 | BA_peak_3278 |
| DS113505 | 67436 | 69113 | BA_peak_3279 |
| DS113505 | 69473 | 71435 | BA_peak_3280 |
| DS113505 | 87915 | 88684 | BA_peak_3281 |
| DS113506 | 11437 | 11729 | BA_peak_3282 |
| DS113506 | 15865 | 16277 | BA_peak_3283 |
| DS113506 | 50463 | 50805 | BA_peak_3284 |
| DS113506 | 83755 | 84118 | BA_peak_3285 |
| DS113506 | 85102 | 85666 | BA_peak_3286 |
| DS113507 | 23458 | 24859 | BA_peak_3287 |
| DS113507 | 41728 | 42152 | BA_peak_3288 |
| DS113507 | 43570 | 44835 | BA_peak_3289 |
| DS113507 | 56371 | 56929 | BA_peak_3290 |
| DS113507 | 58624 | 59371 | BA_peak_3291 |
| DS113507 | 66047 | 66443 | BA_peak_3292 |
| DS113507 | 66911 | 68801 | BA_peak_3293 |
| DS113507 | 69237 | 69755 | BA_peak_3294 |
| DS113507 | 79266 | 80490 | BA_peak_3295 |
| DS113507 | 81801 | 82252 | BA_peak_3296 |
| DS113507 | 84170 | 84965 | BA_peak_3297 |
| DS113508 | 11057 | 11445 | BA_peak_3298 |
| DS113508 | 60549 | 60952 | BA_peak_3299 |
| DS113508 | 81780 | 82332 | BA_peak_3300 |
| DS113508 | 83085 | 83440 | BA_peak_3301 |
| DS113509 | 21938 | 22658 | BA_peak_3302 |
| DS113509 | 23460 | 24137 | BA_peak_3303 |
| DS113509 | 24740 | 25063 | BA_peak_3304 |
| DS113509 | 31464 | 32164 | BA_peak_3305 |
| DS113509 | 44727 | 45396 | BA_peak_3306 |
| DS113509 | 77957 | 78747 | BA_peak_3307 |
| DS113509 | 85377 | 86096 | BA_peak_3308 |
| DS113510 | 24733 | 28043 | BA_peak_3309 |
| DS113510 | 44592 | 45955 | BA_peak_3310 |
| DS113510 | 48827 | 49776 | BA_peak_3311 |
| DS113511 | 3693 | 4344 | BA_peak_3312 |
| DS113511 | 21537 | 21848 | BA_peak_3313 |
| DS113511 | 44642 | 44978 | BA_peak_3314 |
| DS113512 | 11450 | 12191 | BA_peak_3315 |
| DS113512 | 28137 | 28501 | BA_peak_3316 |
| DS113512 | 46042 | 46456 | BA_peak_3317 |

| | | | |
|---|---|---|---|
| DS113512 | 54927 | 55561 | BA_peak_3318 |
| DS113512 | 56741 | 57156 | BA_peak_3319 |
| DS113512 | 58217 | 59404 | BA_peak_3320 |
| DS113512 | 67732 | 69063 | BA_peak_3321 |
| DS113512 | 69839 | 70401 | BA_peak_3322 |
| DS113512 | 80380 | 80992 | BA_peak_3323 |
| DS113512 | 81793 | 82501 | BA_peak_3324 |
| DS113513 | 20073 | 20428 | BA_peak_3325 |
| DS113513 | 28355 | 28904 | BA_peak_3326 |
| DS113513 | 33797 | 34277 | BA_peak_3327 |
| DS113513 | 52193 | 52890 | BA_peak_3328 |
| DS113513 | 64355 | 64915 | BA_peak_3329 |
| DS113513 | 78715 | 79144 | BA_peak_3331 |
| DS113514 | 7734 | 8139 | BA_peak_3333 |
| DS113514 | 22712 | 23435 | BA_peak_3334 |
| DS113514 | 27486 | 28063 | BA_peak_3335 |
| DS113514 | 33068 | 33441 | BA_peak_3336 |
| DS113514 | 34587 | 36284 | BA_peak_3337 |
| DS113514 | 39189 | 39998 | BA_peak_3338 |
| DS113514 | 64487 | 66426 | BA_peak_3339 |
| DS113514 | 73805 | 74467 | BA_peak_3340 |
| DS113515 | 596 | 953 | BA_peak_3341 |
| DS113515 | 3827 | 4258 | BA_peak_3342 |
| DS113515 | 44230 | 44593 | BA_peak_3343 |
| DS113515 | 45836 | 47274 | BA_peak_3344 |
| DS113515 | 47626 | 48413 | BA_peak_3345 |
| DS113515 | 48815 | 50192 | BA_peak_3346 |
| DS113515 | 73755 | 75104 | BA_peak_3347 |
| DS113515 | 83612 | 83973 | BA_peak_3348 |
| DS113516 | 7957 | 8478 | BA_peak_3349 |
| DS113516 | 21679 | 22424 | BA_peak_3350 |
| DS113516 | 69278 | 70107 | BA_peak_3351 |
| DS113516 | 71817 | 72214 | BA_peak_3352 |
| DS113516 | 81044 | 81640 | BA_peak_3353 |
| DS113516 | 85463 | 86566 | BA_peak_3354 |
| DS113517 | 23852 | 24415 | BA_peak_3355 |
| DS113517 | 44741 | 46253 | BA_peak_3356 |
| DS113517 | 63274 | 65017 | BA_peak_3358 |
| DS113517 | 85203 | 86015 | BA_peak_3359 |
| DS113518 | 176 | 587 | BA_peak_3360 |
| DS113518 | 7341 | 8586 | BA_peak_3361 |
| DS113518 | 22609 | 23046 | BA_peak_3362 |
| DS113518 | 23794 | 24554 | BA_peak_3363 |
| DS113519 | 15208 | 15756 | BA_peak_3364 |
| DS113519 | 44038 | 45028 | BA_peak_3366 |
| DS113519 | 45861 | 46251 | BA_peak_3367 |
| DS113519 | 77494 | 78139 | BA_peak_3369 |

| | | | |
|---|---|---|---|
| DS113520 | 25920 | 27293 | BA_peak_3372 |
| DS113520 | 38565 | 39792 | BA_peak_3373 |
| DS113520 | 45875 | 46398 | BA_peak_3374 |
| DS113520 | 46861 | 47551 | BA_peak_3375 |
| DS113520 | 64915 | 65874 | BA_peak_3376 |
| DS113520 | 76661 | 77550 | BA_peak_3377 |
| DS113520 | 80103 | 80641 | BA_peak_3378 |
| DS113521 | 7313 | 7798 | BA_peak_3379 |
| DS113521 | 11060 | 11687 | BA_peak_3380 |
| DS113521 | 12180 | 12596 | BA_peak_3381 |
| DS113521 | 13474 | 17130 | BA_peak_3382 |
| DS113521 | 57653 | 57991 | BA_peak_3384 |
| DS113521 | 58573 | 59294 | BA_peak_3385 |
| DS113521 | 60232 | 64320 | BA_peak_3386 |
| DS113522 | 3420 | 4053 | BA_peak_3387 |
| DS113522 | 13075 | 13993 | BA_peak_3388 |
| DS113522 | 37709 | 38058 | BA_peak_3390 |
| DS113522 | 38492 | 39487 | BA_peak_3391 |
| DS113522 | 53715 | 54601 | BA_peak_3392 |
| DS113523 | 30164 | 30952 | BA_peak_3393 |
| DS113523 | 32282 | 32712 | BA_peak_3394 |
| DS113523 | 42937 | 43685 | BA_peak_3395 |
| DS113523 | 69262 | 69577 | BA_peak_3396 |
| DS113523 | 75397 | 76248 | BA_peak_3397 |
| DS113523 | 76679 | 77399 | BA_peak_3398 |
| DS113523 | 84444 | 84866 | BA_peak_3399 |
| DS113524 | 36975 | 37408 | BA_peak_3400 |
| DS113524 | 40276 | 40594 | BA_peak_3401 |
| DS113524 | 45516 | 46685 | BA_peak_3402 |
| DS113524 | 50524 | 51317 | BA_peak_3403 |
| DS113525 | 6009 | 6825 | BA_peak_3404 |
| DS113525 | 14936 | 18727 | BA_peak_3405 |
| DS113525 | 21795 | 22544 | BA_peak_3407 |
| DS113525 | 42495 | 43134 | BA_peak_3408 |
| DS113525 | 45414 | 45927 | BA_peak_3409 |
| DS113525 | 70117 | 70915 | BA_peak_3410 |
| DS113525 | 73585 | 73898 | BA_peak_3411 |
| DS113526 | 75531 | 77629 | BA_peak_3412 |
| DS113527 | 1746 | 2703 | BA_peak_3413 |
| DS113527 | 4113 | 6175 | BA_peak_3414 |
| DS113527 | 13896 | 15671 | BA_peak_3415 |
| DS113527 | 27335 | 28918 | BA_peak_3416 |
| DS113527 | 30604 | 32666 | BA_peak_3417 |
| DS113527 | 50424 | 50961 | BA_peak_3418 |
| DS113527 | 76618 | 77816 | BA_peak_3419 |
| DS113528 | 15523 | 16310 | BA_peak_3420 |
| DS113528 | 38998 | 39390 | BA_peak_3421 |

| | | | |
|---|---|---|---|
| DS113528 | 55917 | 57032 | BA_peak_3422 |
| DS113528 | 77000 | 77827 | BA_peak_3423 |
| DS113529 | 24209 | 24985 | BA_peak_3426 |
| DS113529 | 31975 | 32434 | BA_peak_3427 |
| DS113529 | 75219 | 75750 | BA_peak_3428 |
| DS113530 | 5347 | 7101 | BA_peak_3430 |
| DS113530 | 8334 | 9083 | BA_peak_3431 |
| DS113530 | 74202 | 75293 | BA_peak_3432 |
| DS113531 | 1098 | 1622 | BA_peak_3433 |
| DS113531 | 12101 | 13004 | BA_peak_3434 |
| DS113531 | 35205 | 38552 | BA_peak_3436 |
| DS113531 | 41113 | 41513 | BA_peak_3437 |
| DS113531 | 64647 | 64982 | BA_peak_3438 |
| DS113532 | 31208 | 31621 | BA_peak_3439 |
| DS113532 | 41997 | 42411 | BA_peak_3440 |
| DS113532 | 56011 | 56376 | BA_peak_3441 |
| DS113533 | 19639 | 20271 | BA_peak_3442 |
| DS113533 | 57128 | 58446 | BA_peak_3443 |
| DS113533 | 65557 | 65975 | BA_peak_3444 |
| DS113533 | 80284 | 80632 | BA_peak_3445 |
| DS113534 | 5382 | 6173 | BA_peak_3446 |
| DS113534 | 28474 | 29284 | BA_peak_3447 |
| DS113534 | 41562 | 42286 | BA_peak_3448 |
| DS113534 | 43911 | 44380 | BA_peak_3449 |
| DS113535 | 25669 | 26087 | BA_peak_3450 |
| DS113535 | 32373 | 33154 | BA_peak_3451 |
| DS113535 | 42917 | 44378 | BA_peak_3452 |
| DS113535 | 44795 | 46529 | BA_peak_3453 |
| DS113535 | 63842 | 64270 | BA_peak_3454 |
| DS113535 | 64681 | 65118 | BA_peak_3455 |
| DS113535 | 81985 | 82675 | BA_peak_3456 |
| DS113536 | 10286 | 10592 | BA_peak_3457 |
| DS113536 | 11532 | 12154 | BA_peak_3458 |
| DS113536 | 44887 | 45206 | BA_peak_3459 |
| DS113536 | 65631 | 66103 | BA_peak_3460 |
| DS113536 | 77932 | 78536 | BA_peak_3461 |
| DS113537 | 54267 | 54675 | BA_peak_3462 |
| DS113537 | 64274 | 64639 | BA_peak_3463 |
| DS113537 | 76695 | 77523 | BA_peak_3464 |
| DS113537 | 78114 | 78650 | BA_peak_3465 |
| DS113538 | 1296 | 1737 | BA_peak_3466 |
| DS113538 | 19591 | 19981 | BA_peak_3467 |
| DS113538 | 20542 | 21387 | BA_peak_3468 |
| DS113538 | 24203 | 24629 | BA_peak_3469 |
| DS113538 | 26313 | 26678 | BA_peak_3470 |
| DS113538 | 28306 | 28747 | BA_peak_3471 |
| DS113538 | 30900 | 31500 | BA_peak_3473 |

| | | | |
|---|---|---|---|
| DS113538 | 52364 | 53048 | BA_peak_3475 |
| DS113538 | 56818 | 57688 | BA_peak_3476 |
| DS113538 | 60402 | 61176 | BA_peak_3477 |
| DS113538 | 66695 | 67250 | BA_peak_3478 |
| DS113539 | 8596 | 10070 | BA_peak_3479 |
| DS113539 | 21115 | 21926 | BA_peak_3480 |
| DS113539 | 26135 | 26594 | BA_peak_3481 |
| DS113539 | 55902 | 57419 | BA_peak_3482 |
| DS113539 | 82642 | 83047 | BA_peak_3483 |
| DS113540 | 1104 | 2563 | BA_peak_3484 |
| DS113540 | 14426 | 14879 | BA_peak_3485 |
| DS113540 | 46570 | 47163 | BA_peak_3486 |
| DS113540 | 53546 | 54074 | BA_peak_3487 |
| DS113540 | 54430 | 55802 | BA_peak_3488 |
| DS113540 | 74084 | 74456 | BA_peak_3489 |
| DS113540 | 76570 | 77991 | BA_peak_3490 |
| DS113541 | 10646 | 11402 | BA_peak_3491 |
| DS113541 | 27234 | 27587 | BA_peak_3492 |
| DS113541 | 69689 | 70461 | BA_peak_3493 |
| DS113542 | 2246 | 2804 | BA_peak_3494 |
| DS113542 | 34813 | 35431 | BA_peak_3495 |
| DS113542 | 43948 | 45045 | BA_peak_3496 |
| DS113542 | 46119 | 46905 | BA_peak_3497 |
| DS113542 | 52737 | 53908 | BA_peak_3499 |
| DS113542 | 54918 | 55255 | BA_peak_3500 |
| DS113542 | 56551 | 57138 | BA_peak_3501 |
| DS113543 | 43459 | 44782 | BA_peak_3502 |
| DS113543 | 60520 | 63828 | BA_peak_3503 |
| DS113543 | 75920 | 76297 | BA_peak_3504 |
| DS113544 | 69 | 819 | BA_peak_3505 |
| DS113544 | 30330 | 30844 | BA_peak_3506 |
| DS113544 | 35284 | 35942 | BA_peak_3507 |
| DS113544 | 51163 | 51481 | BA_peak_3509 |
| DS113544 | 52066 | 52869 | BA_peak_3510 |
| DS113544 | 53491 | 54673 | BA_peak_3511 |
| DS113545 | 6898 | 7426 | BA_peak_3512 |
| DS113545 | 9125 | 10762 | BA_peak_3513 |
| DS113545 | 16641 | 17160 | BA_peak_3514 |
| DS113545 | 20048 | 20508 | BA_peak_3515 |
| DS113545 | 21550 | 21959 | BA_peak_3516 |
| DS113545 | 66917 | 67579 | BA_peak_3518 |
| DS113546 | 23 | 434 | BA_peak_3519 |
| DS113546 | 11349 | 11741 | BA_peak_3520 |
| DS113546 | 27749 | 31524 | BA_peak_3522 |
| DS113546 | 31894 | 32223 | BA_peak_3523 |
| DS113546 | 32755 | 33907 | BA_peak_3524 |
| DS113546 | 52117 | 52687 | BA_peak_3525 |

| | | | |
|---|---|---|---|
| DS113546 | 64976 | 66307 | BA_peak_3526 |
| DS113546 | 67878 | 68893 | BA_peak_3527 |
| DS113546 | 71027 | 71394 | BA_peak_3528 |
| DS113547 | 11947 | 12738 | BA_peak_3529 |
| DS113547 | 16168 | 16561 | BA_peak_3530 |
| DS113547 | 22272 | 22576 | BA_peak_3531 |
| DS113547 | 24072 | 24749 | BA_peak_3532 |
| DS113547 | 29143 | 29814 | BA_peak_3533 |
| DS113547 | 39116 | 39455 | BA_peak_3534 |
| DS113547 | 54164 | 54699 | BA_peak_3535 |
| DS113547 | 55705 | 56930 | BA_peak_3536 |
| DS113547 | 57737 | 58110 | BA_peak_3537 |
| DS113547 | 75497 | 76452 | BA_peak_3538 |
| DS113547 | 78231 | 78490 | BA_peak_3539 |
| DS113548 | 30047 | 30468 | BA_peak_3540 |
| DS113548 | 35527 | 36014 | BA_peak_3541 |
| DS113548 | 41377 | 42087 | BA_peak_3542 |
| DS113548 | 43267 | 45862 | BA_peak_3543 |
| DS113548 | 49032 | 49823 | BA_peak_3544 |
| DS113548 | 50796 | 51461 | BA_peak_3545 |
| DS113548 | 52216 | 53631 | BA_peak_3546 |
| DS113548 | 57766 | 58201 | BA_peak_3547 |
| DS113549 | 38788 | 39682 | BA_peak_3548 |
| DS113549 | 54288 | 55243 | BA_peak_3549 |
| DS113549 | 55588 | 58243 | BA_peak_3549 |
| DS113549 | 65770 | 66487 | BA_peak_3551 |
| DS113549 | 73256 | 73972 | BA_peak_3552 |
| DS113550 | 19997 | 20858 | BA_peak_3554 |
| DS113550 | 28787 | 29287 | BA_peak_3555 |
| DS113550 | 31004 | 32252 | BA_peak_3556 |
| DS113550 | 32618 | 33246 | BA_peak_3557 |
| DS113550 | 38268 | 39517 | BA_peak_3558 |
| DS113550 | 41110 | 42906 | BA_peak_3559 |
| DS113550 | 50060 | 50673 | BA_peak_3560 |
| DS113550 | 53041 | 53474 | BA_peak_3561 |
| DS113550 | 62545 | 63171 | BA_peak_3562 |
| DS113551 | 1406 | 2022 | BA_peak_3563 |
| DS113551 | 2326 | 6253 | BA_peak_3564 |
| DS113551 | 15988 | 16693 | BA_peak_3565 |
| DS113552 | 11129 | 11434 | BA_peak_3566 |
| DS113552 | 19645 | 20036 | BA_peak_3567 |
| DS113552 | 20768 | 22513 | BA_peak_3568 |
| DS113552 | 23557 | 24317 | BA_peak_3569 |
| DS113552 | 27297 | 27645 | BA_peak_3570 |
| DS113552 | 29909 | 30553 | BA_peak_3571 |
| DS113552 | 34464 | 35072 | BA_peak_3572 |
| DS113552 | 35487 | 36811 | BA_peak_3573 |

| | | | |
|---|---|---|---|
| DS113552 | 61113 | 61410 | BA_peak_3574 |
| DS113552 | 64431 | 65219 | BA_peak_3575 |
| DS113553 | 43015 | 43957 | BA_peak_3576 |
| DS113553 | 73671 | 74054 | BA_peak_3577 |
| DS113553 | 80365 | 80916 | BA_peak_3578 |
| DS113554 | 16294 | 17653 | BA_peak_3580 |
| DS113554 | 19503 | 20347 | BA_peak_3581 |
| DS113554 | 31039 | 33838 | BA_peak_3582 |
| DS113554 | 43211 | 43838 | BA_peak_3584 |
| DS113554 | 54854 | 55866 | BA_peak_3586 |
| DS113554 | 57843 | 59979 | BA_peak_3587 |
| DS113554 | 60799 | 61858 | BA_peak_3588 |
| DS113554 | 74703 | 75689 | BA_peak_3589 |
| DS113555 | 5865 | 6260 | BA_peak_3590 |
| DS113555 | 8390 | 9606 | BA_peak_3591 |
| DS113555 | 13541 | 14577 | BA_peak_3592 |
| DS113555 | 15199 | 15713 | BA_peak_3593 |
| DS113555 | 80412 | 81099 | BA_peak_3594 |
| DS113556 | 12630 | 13231 | BA_peak_3595 |
| DS113556 | 13779 | 14138 | BA_peak_3596 |
| DS113556 | 14565 | 15215 | BA_peak_3597 |
| DS113556 | 44811 | 45343 | BA_peak_3598 |
| DS113556 | 53315 | 53827 | BA_peak_3599 |
| DS113556 | 73800 | 74168 | BA_peak_3600 |
| DS113556 | 74564 | 75045 | BA_peak_3601 |
| DS113557 | 63913 | 64825 | BA_peak_3602 |
| DS113558 | 1515 | 3732 | BA_peak_3603 |
| DS113558 | 4324 | 5637 | BA_peak_3604 |
| DS113558 | 6141 | 6602 | BA_peak_3605 |
| DS113558 | 11456 | 13111 | BA_peak_3606 |
| DS113558 | 17129 | 18066 | BA_peak_3607 |
| DS113558 | 19739 | 20800 | BA_peak_3608 |
| DS113558 | 22516 | 23500 | BA_peak_3609 |
| DS113558 | 40988 | 41529 | BA_peak_3610 |
| DS113558 | 42240 | 43128 | BA_peak_3611 |
| DS113558 | 61106 | 63588 | BA_peak_3612 |
| DS113558 | 63986 | 64466 | BA_peak_3613 |
| DS113558 | 64968 | 65596 | BA_peak_3614 |
| DS113558 | 73933 | 74466 | BA_peak_3615 |
| DS113559 | 23234 | 23846 | BA_peak_3616 |
| DS113560 | 20401 | 21352 | BA_peak_3619 |
| DS113561 | 2911 | 3385 | BA_peak_3620 |
| DS113561 | 12318 | 13310 | BA_peak_3621 |
| DS113561 | 18358 | 19106 | BA_peak_3624 |
| DS113561 | 21796 | 22219 | BA_peak_3625 |
| DS113561 | 24537 | 25089 | BA_peak_3626 |
| DS113561 | 26277 | 27103 | BA_peak_3627 |

| | | | |
|---|---|---|---|
| DS113561 | 46279 | 49635 | BA_peak_3628 |
| DS113561 | 51355 | 52832 | BA_peak_3629 |
| DS113561 | 53544 | 54680 | BA_peak_3630 |
| DS113561 | 66450 | 67353 | BA_peak_3631 |
| DS113561 | 68615 | 69417 | BA_peak_3632 |
| DS113562 | 8634 | 9278 | BA_peak_3634 |
| DS113562 | 60530 | 62338 | BA_peak_3635 |
| DS113562 | 67702 | 68302 | BA_peak_3636 |
| DS113562 | 76175 | 77653 | BA_peak_3637 |
| DS113564 | 7870 | 8370 | BA_peak_3640 |
| DS113564 | 14755 | 15772 | BA_peak_3641 |
| DS113564 | 29785 | 30813 | BA_peak_3642 |
| DS113564 | 38566 | 38876 | BA_peak_3643 |
| DS113564 | 70494 | 70823 | BA_peak_3644 |
| DS113564 | 74718 | 75624 | BA_peak_3645 |
| DS113564 | 77065 | 78560 | BA_peak_3646 |
| DS113565 | 1444 | 2584 | BA_peak_3647 |
| DS113565 | 4261 | 4756 | BA_peak_3648 |
| DS113565 | 25266 | 25816 | BA_peak_3649 |
| DS113565 | 37757 | 41066 | BA_peak_3650 |
| DS113565 | 44579 | 45348 | BA_peak_3651 |
| DS113565 | 51728 | 53545 | BA_peak_3652 |
| DS113565 | 64804 | 65474 | BA_peak_3653 |
| DS113566 | 2429 | 4683 | BA_peak_3654 |
| DS113566 | 38006 | 38840 | BA_peak_3655 |
| DS113566 | 55287 | 56941 | BA_peak_3656 |
| DS113566 | 64392 | 64725 | BA_peak_3657 |
| DS113567 | 23004 | 23524 | BA_peak_3659 |
| DS113568 | 4793 | 5245 | BA_peak_3660 |
| DS113568 | 7639 | 8312 | BA_peak_3661 |
| DS113568 | 10532 | 10987 | BA_peak_3662 |
| DS113568 | 11490 | 12730 | BA_peak_3662 |
| DS113568 | 21826 | 23433 | BA_peak_3663 |
| DS113568 | 29489 | 30292 | BA_peak_3664 |
| DS113568 | 30923 | 33071 | BA_peak_3665 |
| DS113568 | 66905 | 67379 | BA_peak_3666 |
| DS113569 | 47 | 1738 | BA_peak_3667 |
| DS113569 | 10306 | 10811 | BA_peak_3668 |
| DS113569 | 52122 | 53004 | BA_peak_3669 |
| DS113569 | 53348 | 56006 | BA_peak_3670 |
| DS113569 | 64562 | 65032 | BA_peak_3672 |
| DS113569 | 65736 | 66196 | BA_peak_3673 |
| DS113570 | 197 | 597 | BA_peak_3674 |
| DS113570 | 1028 | 1362 | BA_peak_3675 |
| DS113570 | 60233 | 61513 | BA_peak_3676 |
| DS113570 | 65631 | 68102 | BA_peak_3677 |
| DS113571 | 24005 | 24649 | BA_peak_3678 |

| | | | |
|---|---|---|---|
| DS113571 | 25404 | 26119 | BA_peak_3679 |
| DS113571 | 27073 | 27891 | BA_peak_3680 |
| DS113572 | 101 | 652 | BA_peak_3681 |
| DS113572 | 7881 | 9325 | BA_peak_3683 |
| DS113572 | 38136 | 38592 | BA_peak_3684 |
| DS113572 | 46989 | 48096 | BA_peak_3685 |
| DS113572 | 73501 | 74198 | BA_peak_3686 |
| DS113572 | 74550 | 75176 | BA_peak_3687 |
| DS113573 | 4724 | 5511 | BA_peak_3689 |
| DS113573 | 8923 | 9883 | BA_peak_3690 |
| DS113573 | 19290 | 20063 | BA_peak_3691 |
| DS113573 | 23005 | 23425 | BA_peak_3692 |
| DS113573 | 43875 | 44480 | BA_peak_3693 |
| DS113573 | 64690 | 65450 | BA_peak_3694 |
| DS113574 | 16828 | 17223 | BA_peak_3695 |
| DS113574 | 41958 | 42287 | BA_peak_3696 |
| DS113574 | 49907 | 50338 | BA_peak_3697 |
| DS113574 | 56700 | 61205 | BA_peak_3698 |
| DS113575 | 7299 | 7861 | BA_peak_3699 |
| DS113575 | 37196 | 38012 | BA_peak_3700 |
| DS113575 | 38449 | 38825 | BA_peak_3701 |
| DS113575 | 66062 | 66398 | BA_peak_3702 |
| DS113575 | 66827 | 67219 | BA_peak_3703 |
| DS113575 | 67823 | 68602 | BA_peak_3704 |
| DS113575 | 76292 | 76998 | BA_peak_3705 |
| DS113576 | 14556 | 14890 | BA_peak_3706 |
| DS113576 | 59973 | 60521 | BA_peak_3708 |
| DS113576 | 61164 | 62230 | BA_peak_3709 |
| DS113577 | 7806 | 8439 | BA_peak_3710 |
| DS113577 | 11615 | 12503 | BA_peak_3711 |
| DS113577 | 37882 | 38708 | BA_peak_3712 |
| DS113577 | 50626 | 51890 | BA_peak_3713 |
| DS113577 | 53116 | 53835 | BA_peak_3714 |
| DS113577 | 54796 | 55469 | BA_peak_3715 |
| DS113578 | 29061 | 29533 | BA_peak_3716 |
| DS113578 | 30904 | 31232 | BA_peak_3717 |
| DS113578 | 58058 | 58694 | BA_peak_3718 |
| DS113578 | 59886 | 63217 | BA_peak_3719 |
| DS113578 | 63707 | 64370 | BA_peak_3720 |
| DS113579 | 9713 | 10333 | BA_peak_3721 |
| DS113579 | 27153 | 28057 | BA_peak_3722 |
| DS113579 | 36256 | 36731 | BA_peak_3724 |
| DS113579 | 53585 | 54131 | BA_peak_3725 |
| DS113580 | 14994 | 15422 | BA_peak_3726 |
| DS113580 | 16143 | 16886 | BA_peak_3727 |
| DS113580 | 33915 | 34778 | BA_peak_3728 |
| DS113580 | 38258 | 39054 | BA_peak_3730 |

| | | | |
|---|---|---|---|
| DS113580 | 47637 | 48486 | BA_peak_3731 |
| DS113580 | 53840 | 55405 | BA_peak_3732 |
| DS113580 | 63874 | 64614 | BA_peak_3733 |
| DS113581 | 30529 | 30923 | BA_peak_3734 |
| DS113581 | 58672 | 59186 | BA_peak_3735 |
| DS113581 | 60849 | 61587 | BA_peak_3736 |
| DS113581 | 71486 | 72251 | BA_peak_3737 |
| DS113582 | 17374 | 18153 | BA_peak_3738 |
| DS113582 | 22312 | 22963 | BA_peak_3739 |
| DS113582 | 24593 | 25975 | BA_peak_3740 |
| DS113582 | 30192 | 31238 | BA_peak_3741 |
| DS113582 | 32925 | 34948 | BA_peak_3742 |
| DS113582 | 35425 | 38733 | BA_peak_3743 |
| DS113582 | 65737 | 66312 | BA_peak_3744 |
| DS113582 | 74515 | 74814 | BA_peak_3745 |
| DS113583 | 10046 | 10858 | BA_peak_3746 |
| DS113583 | 23255 | 24129 | BA_peak_3747 |
| DS113583 | 57529 | 58279 | BA_peak_3748 |
| DS113584 | 1592 | 3984 | BA_peak_3749 |
| DS113584 | 25823 | 27098 | BA_peak_3750 |
| DS113584 | 39741 | 40556 | BA_peak_3752 |
| DS113584 | 73328 | 73947 | BA_peak_3753 |
| DS113585 | 21327 | 21860 | BA_peak_3754 |
| DS113585 | 39896 | 40365 | BA_peak_3755 |
| DS113586 | 14691 | 15482 | BA_peak_3756 |
| DS113586 | 33145 | 36421 | BA_peak_3757 |
| DS113586 | 52096 | 52837 | BA_peak_3758 |
| DS113586 | 65449 | 65808 | BA_peak_3759 |
| DS113587 | 5271 | 5891 | BA_peak_3760 |
| DS113587 | 26021 | 26713 | BA_peak_3761 |
| DS113587 | 49818 | 50398 | BA_peak_3762 |
| DS113587 | 53067 | 53468 | BA_peak_3763 |
| DS113588 | 35402 | 35879 | BA_peak_3764 |
| DS113589 | 7898 | 8519 | BA_peak_3765 |
| DS113589 | 16564 | 17549 | BA_peak_3766 |
| DS113589 | 34048 | 34480 | BA_peak_3767 |
| DS113590 | 2754 | 3558 | BA_peak_3768 |
| DS113590 | 36108 | 36482 | BA_peak_3770 |
| DS113590 | 46849 | 47290 | BA_peak_3771 |
| DS113590 | 68922 | 69472 | BA_peak_3772 |
| DS113590 | 70992 | 71368 | BA_peak_3773 |
| DS113591 | 9969 | 10668 | BA_peak_3774 |
| DS113591 | 30707 | 31209 | BA_peak_3775 |
| DS113591 | 62837 | 63354 | BA_peak_3776 |
| DS113591 | 74958 | 75333 | BA_peak_3777 |
| DS113592 | 54076 | 54446 | BA_peak_3778 |
| DS113592 | 55490 | 55843 | BA_peak_3779 |

| | | | |
|---|---|---|---|
| DS113593 | 11392 | 11841 | BA_peak_3781 |
| DS113593 | 29681 | 30214 | BA_peak_3782 |
| DS113593 | 31187 | 32467 | BA_peak_3783 |
| DS113593 | 43562 | 43990 | BA_peak_3784 |
| DS113593 | 44954 | 45432 | BA_peak_3785 |
| DS113593 | 52037 | 52628 | BA_peak_3786 |
| DS113594 | 14213 | 14611 | BA_peak_3787 |
| DS113594 | 17193 | 18002 | BA_peak_3788 |
| DS113594 | 21699 | 22035 | BA_peak_3789 |
| DS113594 | 31391 | 31968 | BA_peak_3790 |
| DS113594 | 47452 | 47772 | BA_peak_3791 |
| DS113594 | 71191 | 71592 | BA_peak_3792 |
| DS113595 | 5088 | 5403 | BA_peak_3793 |
| DS113595 | 69734 | 70359 | BA_peak_3794 |
| DS113596 | 23734 | 24362 | BA_peak_3795 |
| DS113596 | 53537 | 54284 | BA_peak_3796 |
| DS113596 | 60594 | 61183 | BA_peak_3797 |
| DS113597 | 18608 | 19974 | BA_peak_3798 |
| DS113597 | 53656 | 54062 | BA_peak_3800 |
| DS113598 | 2302 | 3381 | BA_peak_3801 |
| DS113598 | 38149 | 41314 | BA_peak_3802 |
| DS113598 | 50020 | 50663 | BA_peak_3803 |
| DS113598 | 60425 | 61193 | BA_peak_3804 |
| DS113598 | 69276 | 69878 | BA_peak_3805 |
| DS113598 | 74165 | 74598 | BA_peak_3807 |
| DS113599 | 17356 | 17845 | BA_peak_3808 |
| DS113599 | 18198 | 19025 | BA_peak_3809 |
| DS113599 | 46890 | 47929 | BA_peak_3810 |
| DS113599 | 74010 | 74801 | BA_peak_3811 |
| DS113600 | 204 | 807 | BA_peak_3812 |
| DS113600 | 16048 | 16523 | BA_peak_3813 |
| DS113600 | 29443 | 30781 | BA_peak_3814 |
| DS113600 | 47102 | 49031 | BA_peak_3815 |
| DS113601 | 34725 | 35362 | BA_peak_3816 |
| DS113601 | 35837 | 38695 | BA_peak_3817 |
| DS113601 | 40213 | 40642 | BA_peak_3818 |
| DS113602 | 8629 | 9001 | BA_peak_3819 |
| DS113602 | 39208 | 40763 | BA_peak_3820 |
| DS113602 | 49730 | 50927 | BA_peak_3821 |
| DS113602 | 57202 | 57870 | BA_peak_3823 |
| DS113602 | 58383 | 59273 | BA_peak_3824 |
| DS113602 | 64252 | 64743 | BA_peak_3825 |
| DS113603 | 101 | 785 | BA_peak_3826 |
| DS113603 | 1956 | 5221 | BA_peak_3827 |
| DS113603 | 26922 | 30264 | BA_peak_3828 |
| DS113603 | 30713 | 32181 | BA_peak_3829 |
| DS113603 | 44316 | 45043 | BA_peak_3830 |

| | | | |
|---|---|---|---|
| DS113603 | 51217 | 51564 | BA_peak_3831 |
| DS113603 | 59508 | 59962 | BA_peak_3832 |
| DS113603 | 69658 | 70258 | BA_peak_3833 |
| DS113603 | 71098 | 72840 | BA_peak_3834 |
| DS113604 | 7042 | 7485 | BA_peak_3835 |
| DS113604 | 68088 | 68645 | BA_peak_3836 |
| DS113605 | 468 | 1066 | BA_peak_3837 |
| DS113605 | 4282 | 4636 | BA_peak_3838 |
| DS113605 | 16191 | 17709 | BA_peak_3839 |
| DS113605 | 18025 | 20187 | BA_peak_3840 |
| DS113605 | 32815 | 33768 | BA_peak_3841 |
| DS113605 | 36683 | 37249 | BA_peak_3842 |
| DS113605 | 73398 | 74239 | BA_peak_3843 |
| DS113606 | 723 | 1281 | BA_peak_3844 |
| DS113606 | 8046 | 8602 | BA_peak_3845 |
| DS113606 | 42266 | 42653 | BA_peak_3846 |
| DS113606 | 70134 | 71339 | BA_peak_3847 |
| DS113606 | 73365 | 73991 | BA_peak_3848 |
| DS113607 | 1926 | 2497 | BA_peak_3849 |
| DS113607 | 3997 | 5049 | BA_peak_3850 |
| DS113607 | 11739 | 12334 | BA_peak_3851 |
| DS113608 | 23041 | 25300 | BA_peak_3852 |
| DS113608 | 26103 | 26768 | BA_peak_3853 |
| DS113608 | 35497 | 36308 | BA_peak_3854 |
| DS113608 | 58856 | 60122 | BA_peak_3855 |
| DS113609 | 1782 | 2133 | BA_peak_3856 |
| DS113609 | 11543 | 12335 | BA_peak_3857 |
| DS113609 | 23770 | 24311 | BA_peak_3858 |
| DS113609 | 25435 | 25970 | BA_peak_3859 |
| DS113609 | 43428 | 43749 | BA_peak_3860 |
| DS113609 | 46573 | 50526 | BA_peak_3862 |
| DS113609 | 57277 | 58039 | BA_peak_3864 |
| DS113609 | 59413 | 60365 | BA_peak_3865 |
| DS113609 | 62049 | 63023 | BA_peak_3866 |
| DS113610 | 30976 | 31307 | BA_peak_3868 |
| DS113611 | 7488 | 8515 | BA_peak_3869 |
| DS113611 | 14042 | 14476 | BA_peak_3870 |
| DS113611 | 66843 | 68739 | BA_peak_3871 |
| DS113612 | 929 | 1431 | BA_peak_3872 |
| DS113612 | 51630 | 52750 | BA_peak_3873 |
| DS113612 | 72780 | 73176 | BA_peak_3874 |
| DS113613 | 31635 | 32126 | BA_peak_3875 |
| DS113613 | 37391 | 38219 | BA_peak_3876 |
| DS113613 | 61095 | 61435 | BA_peak_3878 |
| DS113613 | 65059 | 65428 | BA_peak_3879 |
| DS113614 | 19592 | 20343 | BA_peak_3880 |
| DS113615 | 9410 | 10181 | BA_peak_3881 |

| | | | |
|---|---|---|---|
| DS113615 | 17724 | 18198 | BA_peak_3882 |
| DS113615 | 31882 | 32384 | BA_peak_3883 |
| DS113615 | 39898 | 40269 | BA_peak_3884 |
| DS113615 | 44651 | 45041 | BA_peak_3885 |
| DS113615 | 69851 | 70236 | BA_peak_3886 |
| DS113616 | 6609 | 7413 | BA_peak_3887 |
| DS113616 | 8621 | 9144 | BA_peak_3888 |
| DS113617 | 30138 | 30628 | BA_peak_3889 |
| DS113617 | 32836 | 33931 | BA_peak_3890 |
| DS113617 | 48050 | 48862 | BA_peak_3892 |
| DS113617 | 52790 | 53306 | BA_peak_3893 |
| DS113617 | 68883 | 69683 | BA_peak_3894 |
| DS113618 | 39007 | 39610 | BA_peak_3895 |
| DS113618 | 51575 | 52183 | BA_peak_3897 |
| DS113618 | 53604 | 54894 | BA_peak_3898 |
| DS113618 | 63488 | 63956 | BA_peak_3899 |
| DS113619 | 19044 | 19606 | BA_peak_3900 |
| DS113619 | 36377 | 37206 | BA_peak_3901 |
| DS113620 | 6723 | 8716 | BA_peak_3902 |
| DS113620 | 13425 | 14045 | BA_peak_3903 |
| DS113620 | 50112 | 50536 | BA_peak_3905 |
| DS113621 | 13903 | 14572 | BA_peak_3906 |
| DS113621 | 21259 | 21641 | BA_peak_3907 |
| DS113621 | 26542 | 27355 | BA_peak_3908 |
| DS113621 | 31967 | 32717 | BA_peak_3909 |
| DS113621 | 43994 | 46377 | BA_peak_3910 |
| DS113621 | 46925 | 50287 | BA_peak_3911 |
| DS113621 | 69458 | 69855 | BA_peak_3912 |
| DS113621 | 71738 | 72065 | BA_peak_3913 |
| DS113622 | 32 | 1722 | BA_peak_3914 |
| DS113622 | 13673 | 14455 | BA_peak_3915 |
| DS113622 | 16566 | 17060 | BA_peak_3916 |
| DS113622 | 49272 | 49952 | BA_peak_3917 |
| DS113622 | 51710 | 52695 | BA_peak_3918 |
| DS113623 | 46859 | 47699 | BA_peak_3920 |
| DS113623 | 56188 | 57382 | BA_peak_3921 |
| DS113623 | 67839 | 68409 | BA_peak_3922 |
| DS113624 | 20254 | 20707 | BA_peak_3923 |
| DS113624 | 36431 | 39998 | BA_peak_3924 |
| DS113624 | 42359 | 46519 | BA_peak_3925 |
| DS113624 | 65826 | 67483 | BA_peak_3926 |
| DS113625 | 50227 | 51005 | BA_peak_3927 |
| DS113625 | 65934 | 66679 | BA_peak_3928 |
| DS113625 | 70400 | 71983 | BA_peak_3929 |
| DS113626 | 36 | 459 | BA_peak_3930 |
| DS113626 | 3871 | 4662 | BA_peak_3931 |
| DS113626 | 30257 | 30689 | BA_peak_3932 |

| | | | |
|---|---|---|---|
| DS113626 | 46957 | 47658 | BA_peak_3933 |
| DS113626 | 50431 | 51292 | BA_peak_3934 |
| DS113626 | 51851 | 52458 | BA_peak_3935 |
| DS113627 | 53727 | 54581 | BA_peak_3936 |
| DS113627 | 70940 | 71226 | BA_peak_3937 |
| DS113628 | 63785 | 64477 | BA_peak_3938 |
| DS113629 | 35334 | 38573 | BA_peak_3939 |
| DS113629 | 51573 | 51963 | BA_peak_3940 |
| DS113629 | 55430 | 56216 | BA_peak_3941 |
| DS113630 | 1943 | 2652 | BA_peak_3943 |
| DS113630 | 3946 | 4630 | BA_peak_3944 |
| DS113630 | 6083 | 6922 | BA_peak_3945 |
| DS113630 | 25826 | 26600 | BA_peak_3946 |
| DS113630 | 27598 | 28292 | BA_peak_3947 |
| DS113630 | 34652 | 35053 | BA_peak_3948 |
| DS113630 | 35686 | 37105 | BA_peak_3949 |
| DS113630 | 45097 | 45752 | BA_peak_3950 |
| DS113630 | 57579 | 58240 | BA_peak_3951 |
| DS113631 | 12747 | 13521 | BA_peak_3952 |
| DS113631 | 19698 | 20469 | BA_peak_3953 |
| DS113631 | 28855 | 29172 | BA_peak_3954 |
| DS113631 | 30602 | 31577 | BA_peak_3955 |
| DS113631 | 34304 | 34886 | BA_peak_3956 |
| DS113631 | 37211 | 40493 | BA_peak_3957 |
| DS113632 | 7199 | 7516 | BA_peak_3958 |
| DS113632 | 7860 | 8629 | BA_peak_3959 |
| DS113632 | 18238 | 19104 | BA_peak_3960 |
| DS113632 | 25276 | 25718 | BA_peak_3961 |
| DS113632 | 67367 | 67987 | BA_peak_3962 |
| DS113633 | 174 | 1857 | BA_peak_3963 |
| DS113633 | 20394 | 21185 | BA_peak_3964 |
| DS113633 | 35607 | 35939 | BA_peak_3965 |
| DS113633 | 37564 | 38371 | BA_peak_3966 |
| DS113633 | 50306 | 50883 | BA_peak_3967 |
| DS113634 | 62674 | 63511 | BA_peak_3968 |
| DS113634 | 65129 | 65973 | BA_peak_3969 |
| DS113634 | 66708 | 68068 | BA_peak_3970 |
| DS113634 | 69604 | 70464 | BA_peak_3971 |
| DS113635 | 27693 | 28828 | BA_peak_3972 |
| DS113635 | 29628 | 30099 | BA_peak_3973 |
| DS113635 | 31472 | 32204 | BA_peak_3974 |
| DS113635 | 33188 | 33687 | BA_peak_3975 |
| DS113635 | 58944 | 59821 | BA_peak_3976 |
| DS113636 | 6387 | 6747 | BA_peak_3977 |
| DS113636 | 34056 | 35411 | BA_peak_3978 |
| DS113637 | 65983 | 66415 | BA_peak_3979 |
| DS113638 | 44877 | 45314 | BA_peak_3981 |

| | | | |
|---|---|---|---|
| DS113638 | 56667 | 57424 | BA_peak_3982 |
| DS113639 | 764 | 1375 | BA_peak_3983 |
| DS113639 | 6829 | 7654 | BA_peak_3984 |
| DS113639 | 8673 | 9018 | BA_peak_3985 |
| DS113639 | 11887 | 13442 | BA_peak_3986 |
| DS113639 | 15706 | 16134 | BA_peak_3987 |
| DS113639 | 16452 | 17039 | BA_peak_3988 |
| DS113639 | 46651 | 47004 | BA_peak_3989 |
| DS113639 | 48693 | 49456 | BA_peak_3990 |
| DS113639 | 56338 | 56660 | BA_peak_3991 |
| DS113640 | 14075 | 14432 | BA_peak_3992 |
| DS113640 | 15314 | 15668 | BA_peak_3993 |
| DS113640 | 22888 | 23269 | BA_peak_3994 |
| DS113640 | 57856 | 59204 | BA_peak_3995 |
| DS113641 | 7780 | 8538 | BA_peak_3996 |
| DS113641 | 9580 | 10223 | BA_peak_3997 |
| DS113641 | 23289 | 26534 | BA_peak_3998 |
| DS113641 | 34471 | 35004 | BA_peak_3999 |
| DS113641 | 62389 | 62814 | BA_peak_4000 |
| DS113641 | 66094 | 66484 | BA_peak_4001 |
| DS113642 | 66338 | 66798 | BA_peak_4002 |
| DS113643 | 13432 | 13823 | BA_peak_4003 |
| DS113644 | 11213 | 12354 | BA_peak_4004 |
| DS113644 | 18454 | 21732 | BA_peak_4005 |
| DS113645 | 66989 | 68254 | BA_peak_4006 |
| DS113646 | 19500 | 20197 | BA_peak_4007 |
| DS113646 | 21113 | 21850 | BA_peak_4008 |
| DS113646 | 49543 | 49889 | BA_peak_4009 |
| DS113647 | 6452 | 7240 | BA_peak_4010 |
| DS113647 | 11048 | 11507 | BA_peak_4012 |
| DS113647 | 11976 | 13042 | BA_peak_4013 |
| DS113647 | 15397 | 16217 | BA_peak_4014 |
| DS113647 | 59234 | 59619 | BA_peak_4015 |
| DS113648 | 41309 | 42124 | BA_peak_4017 |
| DS113649 | 6237 | 7430 | BA_peak_4018 |
| DS113649 | 44102 | 44737 | BA_peak_4019 |
| DS113649 | 46165 | 46601 | BA_peak_4020 |
| DS113649 | 59798 | 62520 | BA_peak_4021 |
| DS113650 | 26893 | 27229 | BA_peak_4022 |
| DS113650 | 27911 | 28574 | BA_peak_4023 |
| DS113650 | 36163 | 36982 | BA_peak_4024 |
| DS113650 | 42710 | 43432 | BA_peak_4025 |
| DS113650 | 57957 | 58715 | BA_peak_4026 |
| DS113650 | 60075 | 61398 | BA_peak_4027 |
| DS113651 | 15259 | 15616 | BA_peak_4028 |
| DS113651 | 56811 | 57288 | BA_peak_4029 |
| DS113652 | 45195 | 45590 | BA_peak_4031 |

| | | | |
|---|---|---|---|
| DS113653 | 5506 | 5916 | BA_peak_4032 |
| DS113653 | 20143 | 20454 | BA_peak_4033 |
| DS113653 | 49027 | 51321 | BA_peak_4034 |
| DS113653 | 55813 | 58630 | BA_peak_4035 |
| DS113654 | 9762 | 10914 | BA_peak_4036 |
| DS113654 | 15154 | 18690 | BA_peak_4037 |
| DS113654 | 20679 | 23609 | BA_peak_4038 |
| DS113654 | 42530 | 43327 | BA_peak_4039 |
| DS113654 | 59836 | 61210 | BA_peak_4040 |
| DS113655 | 22736 | 23297 | BA_peak_4041 |
| DS113655 | 49247 | 50642 | BA_peak_4042 |
| DS113655 | 59604 | 60568 | BA_peak_4043 |
| DS113656 | 12625 | 12967 | BA_peak_4044 |
| DS113656 | 25213 | 25987 | BA_peak_4045 |
| DS113656 | 62554 | 62949 | BA_peak_4046 |
| DS113657 | 2998 | 4060 | BA_peak_4048 |
| DS113657 | 6056 | 6678 | BA_peak_4049 |
| DS113657 | 13921 | 15055 | BA_peak_4050 |
| DS113657 | 16672 | 19118 | BA_peak_4051 |
| DS113657 | 67905 | 68408 | BA_peak_4053 |
| DS113658 | 3860 | 4351 | BA_peak_4054 |
| DS113658 | 26739 | 27642 | BA_peak_4055 |
| DS113658 | 39059 | 39378 | BA_peak_4056 |
| DS113658 | 52406 | 52952 | BA_peak_4057 |
| DS113659 | 4724 | 5117 | BA_peak_4059 |
| DS113659 | 5427 | 5830 | BA_peak_4060 |
| DS113659 | 10209 | 10618 | BA_peak_4061 |
| DS113659 | 12260 | 12643 | BA_peak_4062 |
| DS113659 | 13197 | 13685 | BA_peak_4063 |
| DS113659 | 42886 | 43283 | BA_peak_4064 |
| DS113659 | 49292 | 49691 | BA_peak_4065 |
| DS113659 | 57624 | 59361 | BA_peak_4066 |
| DS113659 | 62779 | 64021 | BA_peak_4067 |
| DS113660 | 191 | 888 | BA_peak_4068 |
| DS113660 | 39648 | 40071 | BA_peak_4070 |
| DS113660 | 40546 | 42201 | BA_peak_4071 |
| DS113660 | 44635 | 45186 | BA_peak_4072 |
| DS113660 | 46375 | 47093 | BA_peak_4073 |
| DS113660 | 62108 | 62995 | BA_peak_4074 |
| DS113661 | 115 | 952 | BA_peak_4075 |
| DS113661 | 3951 | 4344 | BA_peak_4076 |
| DS113661 | 8936 | 9514 | BA_peak_4077 |
| DS113661 | 22236 | 22700 | BA_peak_4078 |
| DS113661 | 23292 | 23781 | BA_peak_4079 |
| DS113661 | 25558 | 25922 | BA_peak_4080 |
| DS113661 | 52685 | 53095 | BA_peak_4081 |
| DS113661 | 53813 | 54485 | BA_peak_4082 |

| | | | |
|---|---|---|---|
| DS113663 | 5827 | 6465 | BA_peak_4084 |
| DS113663 | 44530 | 45465 | BA_peak_4085 |
| DS113664 | 5821 | 7553 | BA_peak_4086 |
| DS113664 | 34322 | 35318 | BA_peak_4087 |
| DS113665 | 323 | 989 | BA_peak_4088 |
| DS113665 | 1925 | 2805 | BA_peak_4089 |
| DS113665 | 32216 | 32887 | BA_peak_4092 |
| DS113665 | 62295 | 64974 | BA_peak_4093 |
| DS113666 | 24099 | 24578 | BA_peak_4094 |
| DS113667 | 1351 | 1718 | BA_peak_4095 |
| DS113667 | 14619 | 15237 | BA_peak_4096 |
| DS113667 | 19223 | 20070 | BA_peak_4097 |
| DS113667 | 20947 | 21616 | BA_peak_4098 |
| DS113667 | 56034 | 56572 | BA_peak_4099 |
| DS113667 | 57794 | 58281 | BA_peak_4100 |
| DS113668 | 39790 | 40230 | BA_peak_4101 |
| DS113668 | 66515 | 66822 | BA_peak_4103 |
| DS113669 | 54397 | 55004 | BA_peak_4104 |
| DS113669 | 58576 | 59223 | BA_peak_4105 |
| DS113670 | 33587 | 33883 | BA_peak_4106 |
| DS113670 | 36034 | 36351 | BA_peak_4107 |
| DS113671 | 19184 | 19647 | BA_peak_4109 |
| DS113671 | 44770 | 45161 | BA_peak_4111 |
| DS113672 | 22033 | 23279 | BA_peak_4112 |
| DS113672 | 34870 | 36454 | BA_peak_4113 |
| DS113673 | 15892 | 16678 | BA_peak_4114 |
| DS113673 | 50417 | 50843 | BA_peak_4115 |
| DS113673 | 63212 | 64241 | BA_peak_4116 |
| DS113673 | 66080 | 66945 | BA_peak_4117 |
| DS113674 | 11796 | 12529 | BA_peak_4118 |
| DS113674 | 44063 | 47100 | BA_peak_4119 |
| DS113674 | 53436 | 56139 | BA_peak_4120 |
| DS113674 | 63159 | 64906 | BA_peak_4121 |
| DS113675 | 58160 | 59000 | BA_peak_4122 |
| DS113676 | 14811 | 15278 | BA_peak_4123 |
| DS113676 | 17162 | 17955 | BA_peak_4124 |
| DS113676 | 18880 | 19382 | BA_peak_4125 |
| DS113676 | 32067 | 33812 | BA_peak_4126 |
| DS113676 | 34191 | 35605 | BA_peak_4127 |
| DS113677 | 7790 | 9283 | BA_peak_4128 |
| DS113677 | 25578 | 26004 | BA_peak_4129 |
| DS113677 | 50938 | 51759 | BA_peak_4130 |
| DS113677 | 62428 | 62874 | BA_peak_4131 |
| DS113677 | 65013 | 65375 | BA_peak_4132 |
| DS113678 | 27034 | 27679 | BA_peak_4134 |
| DS113678 | 28754 | 29312 | BA_peak_4135 |
| DS113678 | 48707 | 49466 | BA_peak_4136 |

| | | | |
|---|---|---|---|
| DS113679 | 18688 | 19268 | BA_peak_4137 |
| DS113679 | 19629 | 20051 | BA_peak_4138 |
| DS113679 | 24004 | 24699 | BA_peak_4139 |
| DS113679 | 26207 | 26551 | BA_peak_4140 |
| DS113679 | 58972 | 63064 | BA_peak_4141 |
| DS113679 | 63566 | 64305 | BA_peak_4142 |
| DS113679 | 64676 | 65134 | BA_peak_4143 |
| DS113680 | 8078 | 8394 | BA_peak_4144 |
| DS113680 | 21567 | 22196 | BA_peak_4145 |
| DS113680 | 32023 | 32927 | BA_peak_4146 |
| DS113680 | 39641 | 40055 | BA_peak_4147 |
| DS113680 | 65019 | 65580 | BA_peak_4148 |
| DS113681 | 5680 | 6880 | BA_peak_4149 |
| DS113681 | 18103 | 18643 | BA_peak_4150 |
| DS113681 | 39027 | 39727 | BA_peak_4151 |
| DS113681 | 57084 | 57686 | BA_peak_4152 |
| DS113682 | 229 | 596 | BA_peak_4153 |
| DS113682 | 41899 | 42342 | BA_peak_4154 |
| DS113682 | 43845 | 44348 | BA_peak_4155 |
| DS113682 | 58509 | 59085 | BA_peak_4156 |
| DS113683 | 5512 | 6272 | BA_peak_4157 |
| DS113683 | 25603 | 26030 | BA_peak_4158 |
| DS113683 | 47391 | 47831 | BA_peak_4159 |
| DS113683 | 55017 | 55428 | BA_peak_4160 |
| DS113684 | 1087 | 1501 | BA_peak_4161 |
| DS113684 | 2329 | 2939 | BA_peak_4162 |
| DS113684 | 28979 | 29409 | BA_peak_4163 |
| DS113684 | 29779 | 30170 | BA_peak_4164 |
| DS113684 | 62022 | 62680 | BA_peak_4165 |
| DS113684 | 63120 | 64060 | BA_peak_4166 |
| DS113685 | 7695 | 8191 | BA_peak_4167 |
| DS113685 | 55957 | 56463 | BA_peak_4168 |
| DS113685 | 56776 | 57152 | BA_peak_4169 |
| DS113686 | 12485 | 12817 | BA_peak_4170 |
| DS113686 | 57197 | 57812 | BA_peak_4171 |
| DS113687 | 8828 | 10094 | BA_peak_4172 |
| DS113687 | 37418 | 37723 | BA_peak_4173 |
| DS113687 | 47428 | 50715 | BA_peak_4174 |
| DS113688 | 1458 | 2807 | BA_peak_4175 |
| DS113688 | 13685 | 14365 | BA_peak_4176 |
| DS113688 | 14937 | 16372 | BA_peak_4177 |
| DS113688 | 30865 | 32039 | BA_peak_4178 |
| DS113688 | 40521 | 40865 | BA_peak_4179 |
| DS113688 | 58330 | 58896 | BA_peak_4180 |
| DS113688 | 62739 | 63784 | BA_peak_4181 |
| DS113689 | 28911 | 33218 | BA_peak_4183 |
| DS113690 | 21091 | 21758 | BA_peak_4184 |

| | | | |
|---|---|---|---|
| DS113690 | 22519 | 22896 | BA_peak_4185 |
| DS113690 | 23312 | 23773 | BA_peak_4186 |
| DS113690 | 28798 | 29340 | BA_peak_4187 |
| DS113690 | 58434 | 59047 | BA_peak_4189 |
| DS113691 | 40433 | 41207 | BA_peak_4190 |
| DS113691 | 58216 | 59081 | BA_peak_4191 |
| DS113691 | 63376 | 63781 | BA_peak_4192 |
| DS113692 | 4456 | 4892 | BA_peak_4193 |
| DS113692 | 13989 | 14331 | BA_peak_4194 |
| DS113692 | 21833 | 22327 | BA_peak_4195 |
| DS113692 | 38673 | 39305 | BA_peak_4196 |
| DS113692 | 52914 | 53620 | BA_peak_4197 |
| DS113692 | 54019 | 54783 | BA_peak_4198 |
| DS113693 | 3061 | 4037 | BA_peak_4199 |
| DS113693 | 5990 | 6386 | BA_peak_4200 |
| DS113693 | 9881 | 10454 | BA_peak_4201 |
| DS113693 | 13880 | 14229 | BA_peak_4202 |
| DS113693 | 27432 | 27885 | BA_peak_4203 |
| DS113693 | 34996 | 35835 | BA_peak_4204 |
| DS113693 | 36142 | 37950 | BA_peak_4205 |
| DS113694 | 25630 | 26499 | BA_peak_4206 |
| DS113694 | 47822 | 48311 | BA_peak_4207 |
| DS113694 | 56914 | 57239 | BA_peak_4209 |
| DS113694 | 62337 | 63091 | BA_peak_4210 |
| DS113695 | 25524 | 25939 | BA_peak_4211 |
| DS113695 | 30039 | 32618 | BA_peak_4212 |
| DS113695 | 33267 | 34085 | BA_peak_4213 |
| DS113695 | 43083 | 43687 | BA_peak_4214 |
| DS113695 | 48893 | 49355 | BA_peak_4215 |
| DS113695 | 49843 | 50261 | BA_peak_4216 |
| DS113695 | 52853 | 53640 | BA_peak_4217 |
| DS113696 | 1342 | 1818 | BA_peak_4218 |
| DS113696 | 3174 | 3710 | BA_peak_4219 |
| DS113696 | 44152 | 44641 | BA_peak_4220 |
| DS113696 | 47431 | 49240 | BA_peak_4221 |
| DS113696 | 49712 | 51063 | BA_peak_4222 |
| DS113696 | 51753 | 52717 | BA_peak_4223 |
| DS113697 | 14278 | 16952 | BA_peak_4224 |
| DS113697 | 28401 | 28988 | BA_peak_4225 |
| DS113697 | 39558 | 40945 | BA_peak_4226 |
| DS113697 | 41481 | 41921 | BA_peak_4227 |
| DS113698 | 250 | 683 | BA_peak_4228 |
| DS113698 | 5517 | 7058 | BA_peak_4230 |
| DS113698 | 16905 | 17638 | BA_peak_4231 |
| DS113698 | 38086 | 41435 | BA_peak_4232 |
| DS113698 | 44377 | 44837 | BA_peak_4233 |
| DS113698 | 61244 | 61932 | BA_peak_4234 |

| | | | |
|---|---|---|---|
| DS113699 | 6492 | 7110 | BA_peak_4235 |
| DS113699 | 14406 | 15317 | BA_peak_4236 |
| DS113699 | 61836 | 63514 | BA_peak_4237 |
| DS113700 | 28012 | 28742 | BA_peak_4238 |
| DS113701 | 12368 | 13430 | BA_peak_4239 |
| DS113701 | 13777 | 14147 | BA_peak_4239 |
| DS113701 | 20899 | 21705 | BA_peak_4240 |
| DS113701 | 26059 | 26417 | BA_peak_4241 |
| DS113701 | 58942 | 59338 | BA_peak_4242 |
| DS113701 | 62435 | 63355 | BA_peak_4243 |
| DS113702 | 1008 | 1556 | BA_peak_4244 |
| DS113702 | 5975 | 6350 | BA_peak_4245 |
| DS113702 | 37083 | 37576 | BA_peak_4246 |
| DS113702 | 38753 | 40321 | BA_peak_4247 |
| DS113703 | 120 | 951 | BA_peak_4248 |
| DS113704 | 11490 | 11842 | BA_peak_4249 |
| DS113704 | 32912 | 36757 | BA_peak_4250 |
| DS113704 | 43687 | 44184 | BA_peak_4251 |
| DS113704 | 57572 | 58027 | BA_peak_4252 |
| DS113705 | 29644 | 30042 | BA_peak_4253 |
| DS113706 | 4065 | 5284 | BA_peak_4254 |
| DS113706 | 8002 | 8898 | BA_peak_4255 |
| DS113706 | 34363 | 35330 | BA_peak_4256 |
| DS113707 | 3823 | 4171 | BA_peak_4257 |
| DS113707 | 30175 | 32744 | BA_peak_4259 |
| DS113707 | 50332 | 50952 | BA_peak_4260 |
| DS113707 | 62112 | 62586 | BA_peak_4261 |
| DS113708 | 1457 | 1782 | BA_peak_4262 |
| DS113708 | 4382 | 4930 | BA_peak_4263 |
| DS113708 | 12022 | 12695 | BA_peak_4265 |
| DS113708 | 31854 | 32198 | BA_peak_4266 |
| DS113708 | 32742 | 33242 | BA_peak_4267 |
| DS113708 | 34345 | 36163 | BA_peak_4268 |
| DS113708 | 43584 | 44371 | BA_peak_4269 |
| DS113709 | 177 | 3627 | BA_peak_4270 |
| DS113709 | 4087 | 5030 | BA_peak_4271 |
| DS113709 | 10093 | 10512 | BA_peak_4272 |
| DS113709 | 11020 | 11990 | BA_peak_4273 |
| DS113709 | 60333 | 61131 | BA_peak_4274 |
| DS113710 | 3057 | 5106 | BA_peak_4275 |
| DS113710 | 6944 | 7704 | BA_peak_4276 |
| DS113710 | 47697 | 48985 | BA_peak_4277 |
| DS113710 | 49725 | 50648 | BA_peak_4278 |
| DS113710 | 55508 | 55832 | BA_peak_4279 |
| DS113711 | 16625 | 17748 | BA_peak_4280 |
| DS113711 | 39371 | 39755 | BA_peak_4281 |
| DS113711 | 59790 | 60472 | BA_peak_4282 |

| | | | |
|---|---|---|---|
| DS113711 | 60925 | 62188 | BA_peak_4283 |
| DS113712 | 13179 | 13851 | BA_peak_4284 |
| DS113712 | 18356 | 19059 | BA_peak_4285 |
| DS113712 | 19480 | 20324 | BA_peak_4286 |
| DS113712 | 20771 | 21130 | BA_peak_4287 |
| DS113712 | 29840 | 33033 | BA_peak_4288 |
| DS113712 | 38642 | 40004 | BA_peak_4289 |
| DS113712 | 42417 | 43025 | BA_peak_4290 |
| DS113712 | 50467 | 50806 | BA_peak_4291 |
| DS113713 | 9177 | 9835 | BA_peak_4292 |
| DS113713 | 10904 | 11248 | BA_peak_4293 |
| DS113713 | 11930 | 15467 | BA_peak_4294 |
| DS113713 | 41676 | 42062 | BA_peak_4295 |
| DS113713 | 43014 | 43984 | BA_peak_4296 |
| DS113713 | 44458 | 45123 | BA_peak_4297 |
| DS113713 | 51875 | 52641 | BA_peak_4298 |
| DS113713 | 53093 | 53559 | BA_peak_4299 |
| DS113714 | 13915 | 14266 | BA_peak_4300 |
| DS113714 | 18449 | 18961 | BA_peak_4301 |
| DS113714 | 29157 | 30384 | BA_peak_4302 |
| DS113714 | 30696 | 31040 | BA_peak_4303 |
| DS113714 | 31436 | 32633 | BA_peak_4304 |
| DS113714 | 34448 | 34943 | BA_peak_4305 |
| DS113714 | 40084 | 40458 | BA_peak_4306 |
| DS113714 | 41144 | 41724 | BA_peak_4307 |
| DS113714 | 42352 | 43379 | BA_peak_4308 |
| DS113714 | 54618 | 55394 | BA_peak_4309 |
| DS113715 | 24303 | 25043 | BA_peak_4310 |
| DS113715 | 56766 | 57328 | BA_peak_4311 |
| DS113716 | 45821 | 46375 | BA_peak_4312 |
| DS113718 | 18743 | 19123 | BA_peak_4313 |
| DS113718 | 22997 | 23684 | BA_peak_4314 |
| DS113718 | 29263 | 29941 | BA_peak_4315 |
| DS113719 | 8472 | 9319 | BA_peak_4316 |
| DS113719 | 39050 | 39475 | BA_peak_4317 |
| DS113720 | 6132 | 8043 | BA_peak_4318 |
| DS113720 | 8439 | 8792 | BA_peak_4319 |
| DS113720 | 33723 | 34087 | BA_peak_4320 |
| DS113720 | 38715 | 39888 | BA_peak_4321 |
| DS113721 | 25304 | 25655 | BA_peak_4322 |
| DS113721 | 39488 | 39890 | BA_peak_4323 |
| DS113721 | 50131 | 50802 | BA_peak_4324 |
| DS113721 | 59013 | 59820 | BA_peak_4325 |
| DS113722 | 27241 | 28522 | BA_peak_4326 |
| DS113722 | 41150 | 41871 | BA_peak_4327 |
| DS113722 | 47282 | 47952 | BA_peak_4328 |
| DS113722 | 60489 | 60901 | BA_peak_4329 |

| | | | |
|---|---|---|---|
| DS113723 | 1147 | 1717 | BA_peak_4330 |
| DS113723 | 26572 | 27086 | BA_peak_4331 |
| DS113723 | 42719 | 43141 | BA_peak_4332 |
| DS113723 | 55216 | 55601 | BA_peak_4333 |
| DS113724 | 8840 | 9223 | BA_peak_4335 |
| DS113724 | 23315 | 23702 | BA_peak_4336 |
| DS113724 | 24302 | 24620 | BA_peak_4337 |
| DS113725 | 3056 | 3678 | BA_peak_4338 |
| DS113725 | 12233 | 12615 | BA_peak_4339 |
| DS113725 | 21060 | 21650 | BA_peak_4340 |
| DS113725 | 22594 | 23192 | BA_peak_4341 |
| DS113725 | 23953 | 24344 | BA_peak_4342 |
| DS113725 | 28120 | 28811 | BA_peak_4343 |
| DS113725 | 54685 | 55528 | BA_peak_4344 |
| DS113725 | 56110 | 56646 | BA_peak_4345 |
| DS113726 | 5589 | 6785 | BA_peak_4346 |
| DS113727 | 4904 | 5713 | BA_peak_4347 |
| DS113727 | 6662 | 7093 | BA_peak_4348 |
| DS113727 | 7514 | 8872 | BA_peak_4349 |
| DS113727 | 14319 | 15135 | BA_peak_4350 |
| DS113727 | 49692 | 50144 | BA_peak_4351 |
| DS113729 | 11337 | 11938 | BA_peak_4352 |
| DS113729 | 19660 | 20415 | BA_peak_4353 |
| DS113729 | 41208 | 41635 | BA_peak_4354 |
| DS113730 | 13035 | 13815 | BA_peak_4355 |
| DS113731 | 56 | 855 | BA_peak_4358 |
| DS113731 | 2089 | 2493 | BA_peak_4359 |
| DS113731 | 24807 | 25177 | BA_peak_4360 |
| DS113731 | 33940 | 34416 | BA_peak_4361 |
| DS113731 | 38539 | 38905 | BA_peak_4362 |
| DS113731 | 42291 | 42763 | BA_peak_4363 |
| DS113732 | 35533 | 35920 | BA_peak_4365 |
| DS113732 | 38551 | 39220 | BA_peak_4366 |
| DS113732 | 44911 | 46109 | BA_peak_4367 |
| DS113732 | 52125 | 52510 | BA_peak_4368 |
| DS113732 | 55135 | 55623 | BA_peak_4369 |
| DS113733 | 18966 | 20430 | BA_peak_4370 |
| DS113733 | 48801 | 49815 | BA_peak_4372 |
| DS113733 | 50860 | 51534 | BA_peak_4373 |
| DS113733 | 53966 | 55334 | BA_peak_4374 |
| DS113734 | 5039 | 6504 | BA_peak_4375 |
| DS113734 | 33176 | 36517 | BA_peak_4376 |
| DS113734 | 44287 | 44668 | BA_peak_4377 |
| DS113734 | 55761 | 56268 | BA_peak_4378 |
| DS113736 | 17423 | 18442 | BA_peak_4379 |
| DS113737 | 355 | 714 | BA_peak_4380 |
| DS113737 | 2629 | 5858 | BA_peak_4381 |

| | | | |
|---|---|---|---|
| DS113737 | 51263 | 51835 | BA_peak_4382 |
| DS113737 | 52161 | 53269 | BA_peak_4383 |
| DS113737 | 54144 | 55645 | BA_peak_4384 |
| DS113737 | 56996 | 57683 | BA_peak_4385 |
| DS113737 | 58884 | 59264 | BA_peak_4386 |
| DS113738 | 82 | 932 | BA_peak_4387 |
| DS113738 | 30946 | 31440 | BA_peak_4388 |
| DS113738 | 34124 | 36048 | BA_peak_4389 |
| DS113738 | 42098 | 42965 | BA_peak_4390 |
| DS113738 | 43406 | 43841 | BA_peak_4391 |
| DS113738 | 57296 | 57664 | BA_peak_4392 |
| DS113739 | 12524 | 13156 | BA_peak_4394 |
| DS113739 | 14055 | 15262 | BA_peak_4395 |
| DS113739 | 17353 | 18082 | BA_peak_4396 |
| DS113739 | 25279 | 25612 | BA_peak_4397 |
| DS113739 | 27481 | 28674 | BA_peak_4398 |
| DS113740 | 98 | 1067 | BA_peak_4400 |
| DS113740 | 1480 | 2091 | BA_peak_4401 |
| DS113740 | 3003 | 3550 | BA_peak_4402 |
| DS113740 | 13392 | 14144 | BA_peak_4403 |
| DS113740 | 25692 | 26368 | BA_peak_4404 |
| DS113740 | 42695 | 43058 | BA_peak_4405 |
| DS113741 | 46486 | 47188 | BA_peak_4406 |
| DS113741 | 53734 | 54452 | BA_peak_4407 |
| DS113742 | 34825 | 36148 | BA_peak_4408 |
| DS113742 | 50333 | 51402 | BA_peak_4409 |
| DS113743 | 7942 | 9642 | BA_peak_4410 |
| DS113743 | 9953 | 11424 | BA_peak_4411 |
| DS113743 | 14384 | 15928 | BA_peak_4412 |
| DS113743 | 17867 | 18976 | BA_peak_4413 |
| DS113743 | 36293 | 36918 | BA_peak_4414 |
| DS113743 | 47091 | 48009 | BA_peak_4416 |
| DS113744 | 6093 | 6449 | BA_peak_4417 |
| DS113744 | 24363 | 25168 | BA_peak_4418 |
| DS113744 | 28182 | 28957 | BA_peak_4419 |
| DS113744 | 36272 | 37474 | BA_peak_4420 |
| DS113744 | 48484 | 49223 | BA_peak_4421 |
| DS113744 | 56055 | 56401 | BA_peak_4422 |
| DS113745 | 6484 | 7141 | BA_peak_4423 |
| DS113745 | 47154 | 49414 | BA_peak_4424 |
| DS113745 | 54339 | 55354 | BA_peak_4425 |
| DS113745 | 56030 | 57430 | BA_peak_4426 |
| DS113746 | 20518 | 21189 | BA_peak_4427 |
| DS113746 | 36609 | 36948 | BA_peak_4428 |
| DS113746 | 47324 | 47924 | BA_peak_4429 |
| DS113746 | 49296 | 50559 | BA_peak_4430 |
| DS113747 | 908 | 1893 | BA_peak_4431 |

| | | | |
|---|---|---|---|
| DS113747 | 3479 | 4403 | BA_peak_4432 |
| DS113747 | 9899 | 10379 | BA_peak_4433 |
| DS113747 | 12483 | 13580 | BA_peak_4434 |
| DS113748 | 1820 | 2193 | BA_peak_4435 |
| DS113748 | 29383 | 32692 | BA_peak_4436 |
| DS113748 | 42464 | 43944 | BA_peak_4437 |
| DS113749 | 5704 | 6349 | BA_peak_4438 |
| DS113749 | 9196 | 10039 | BA_peak_4439 |
| DS113749 | 30208 | 30913 | BA_peak_4440 |
| DS113749 | 31679 | 32257 | BA_peak_4441 |
| DS113749 | 33221 | 33827 | BA_peak_4442 |
| DS113749 | 54040 | 54946 | BA_peak_4443 |
| DS113750 | 9899 | 10732 | BA_peak_4444 |
| DS113751 | 345 | 711 | BA_peak_4445 |
| DS113751 | 5856 | 6357 | BA_peak_4446 |
| DS113751 | 12386 | 12916 | BA_peak_4448 |
| DS113751 | 15943 | 16735 | BA_peak_4449 |
| DS113751 | 53716 | 54439 | BA_peak_4450 |
| DS113752 | 22361 | 23605 | BA_peak_4451 |
| DS113752 | 24150 | 25614 | BA_peak_4452 |
| DS113752 | 32805 | 34048 | BA_peak_4453 |
| DS113752 | 43884 | 44728 | BA_peak_4454 |
| DS113752 | 47236 | 47943 | BA_peak_4455 |
| DS113752 | 48384 | 48776 | BA_peak_4456 |
| DS113752 | 49823 | 50256 | BA_peak_4457 |
| DS113753 | 2814 | 4721 | BA_peak_4458 |
| DS113753 | 6638 | 7132 | BA_peak_4459 |
| DS113753 | 7667 | 8439 | BA_peak_4460 |
| DS113753 | 8822 | 9688 | BA_peak_4461 |
| DS113753 | 26879 | 27351 | BA_peak_4462 |
| DS113753 | 39066 | 40237 | BA_peak_4464 |
| DS113753 | 41735 | 42049 | BA_peak_4465 |
| DS113753 | 45851 | 46823 | BA_peak_4466 |
| DS113753 | 53830 | 54230 | BA_peak_4467 |
| DS113754 | 57151 | 57830 | BA_peak_4469 |
| DS113755 | 7228 | 8976 | BA_peak_4470 |
| DS113755 | 30839 | 31655 | BA_peak_4471 |
| DS113755 | 32493 | 33159 | BA_peak_4472 |
| DS113755 | 38544 | 39304 | BA_peak_4473 |
| DS113756 | 10482 | 10942 | BA_peak_4474 |
| DS113756 | 35958 | 36425 | BA_peak_4475 |
| DS113757 | 12794 | 13176 | BA_peak_4477 |
| DS113757 | 38877 | 42400 | BA_peak_4478 |
| DS113758 | 2182 | 2752 | BA_peak_4479 |
| DS113758 | 5059 | 5763 | BA_peak_4480 |
| DS113758 | 28722 | 29131 | BA_peak_4482 |
| DS113758 | 30533 | 31065 | BA_peak_4483 |

| | | | |
|---|---|---|---|
| DS113758 | 33600 | 34337 | BA_peak_4484 |
| DS113758 | 38524 | 39515 | BA_peak_4485 |
| DS113758 | 45069 | 45430 | BA_peak_4486 |
| DS113759 | 10472 | 10790 | BA_peak_4487 |
| DS113759 | 17263 | 17949 | BA_peak_4488 |
| DS113761 | 17510 | 18367 | BA_peak_4490 |
| DS113761 | 42140 | 42901 | BA_peak_4491 |
| DS113762 | 20261 | 21273 | BA_peak_4492 |
| DS113762 | 41251 | 41957 | BA_peak_4493 |
| DS113762 | 56759 | 57358 | BA_peak_4494 |
| DS113763 | 18195 | 18728 | BA_peak_4496 |
| DS113763 | 37733 | 38237 | BA_peak_4497 |
| DS113763 | 53688 | 55467 | BA_peak_4498 |
| DS113764 | 36589 | 37128 | BA_peak_4499 |
| DS113764 | 41236 | 41909 | BA_peak_4500 |
| DS113764 | 42436 | 42915 | BA_peak_4501 |
| DS113764 | 43307 | 43794 | BA_peak_4502 |
| DS113765 | 1734 | 5142 | BA_peak_4503 |
| DS113765 | 5768 | 6895 | BA_peak_4504 |
| DS113765 | 20020 | 20719 | BA_peak_4505 |
| DS113765 | 22782 | 23117 | BA_peak_4506 |
| DS113765 | 25552 | 26123 | BA_peak_4507 |
| DS113765 | 47157 | 48745 | BA_peak_4508 |
| DS113766 | 27705 | 28234 | BA_peak_4509 |
| DS113766 | 31010 | 33544 | BA_peak_4510 |
| DS113767 | 2361 | 2770 | BA_peak_4512 |
| DS113767 | 3165 | 3594 | BA_peak_4513 |
| DS113767 | 20176 | 21512 | BA_peak_4514 |
| DS113767 | 49048 | 49412 | BA_peak_4515 |
| DS113767 | 51002 | 54268 | BA_peak_4516 |
| DS113767 | 56494 | 56897 | BA_peak_4517 |
| DS113768 | 8880 | 9840 | BA_peak_4518 |
| DS113768 | 47154 | 47685 | BA_peak_4519 |
| DS113770 | 26112 | 26623 | BA_peak_4520 |
| DS113770 | 29302 | 30119 | BA_peak_4521 |
| DS113770 | 47160 | 47509 | BA_peak_4522 |
| DS113770 | 48655 | 49396 | BA_peak_4523 |
| DS113771 | 24994 | 27648 | BA_peak_4524 |
| DS113771 | 29258 | 30555 | BA_peak_4525 |
| DS113771 | 39118 | 39700 | BA_peak_4526 |
| DS113771 | 40229 | 40647 | BA_peak_4527 |
| DS113772 | 6000 | 6552 | BA_peak_4528 |
| DS113772 | 20238 | 20866 | BA_peak_4529 |
| DS113772 | 21466 | 22363 | BA_peak_4530 |
| DS113772 | 22990 | 24316 | BA_peak_4531 |
| DS113772 | 41586 | 42613 | BA_peak_4532 |
| DS113772 | 43595 | 44633 | BA_peak_4533 |

| | | | |
|---|---|---|---|
| DS113773 | 2295 | 3145 | BA_peak_4534 |
| DS113774 | 14932 | 15820 | BA_peak_4535 |
| DS113774 | 18020 | 19109 | BA_peak_4536 |
| DS113774 | 21843 | 22230 | BA_peak_4537 |
| DS113774 | 29594 | 32964 | BA_peak_4538 |
| DS113774 | 33327 | 34175 | BA_peak_4539 |
| DS113774 | 34744 | 35474 | BA_peak_4540 |
| DS113774 | 37131 | 37566 | BA_peak_4541 |
| DS113775 | 11343 | 12104 | BA_peak_4542 |
| DS113775 | 44762 | 45667 | BA_peak_4543 |
| DS113775 | 48042 | 49687 | BA_peak_4545 |
| DS113776 | 31319 | 31997 | BA_peak_4546 |
| DS113776 | 42511 | 42900 | BA_peak_4547 |
| DS113776 | 43638 | 45445 | BA_peak_4548 |
| DS113776 | 52258 | 52614 | BA_peak_4549 |
| DS113777 | 1069 | 1792 | BA_peak_4550 |
| DS113777 | 5457 | 5883 | BA_peak_4551 |
| DS113777 | 27835 | 28461 | BA_peak_4552 |
| DS113777 | 55388 | 55833 | BA_peak_4553 |
| DS113778 | 14710 | 16602 | BA_peak_4554 |
| DS113778 | 28352 | 28801 | BA_peak_4555 |
| DS113778 | 29117 | 30380 | BA_peak_4556 |
| DS113778 | 33309 | 35369 | BA_peak_4557 |
| DS113779 | 6025 | 8101 | BA_peak_4558 |
| DS113779 | 24441 | 25647 | BA_peak_4560 |
| DS113779 | 36010 | 36839 | BA_peak_4561 |
| DS113779 | 49022 | 49761 | BA_peak_4563 |
| DS113780 | 1813 | 3156 | BA_peak_4564 |
| DS113780 | 6150 | 6553 | BA_peak_4565 |
| DS113780 | 27492 | 28112 | BA_peak_4566 |
| DS113780 | 55008 | 55828 | BA_peak_4567 |
| DS113781 | 6486 | 7880 | BA_peak_4568 |
| DS113781 | 21235 | 22718 | BA_peak_4569 |
| DS113781 | 50283 | 50887 | BA_peak_4570 |
| DS113782 | 29889 | 30281 | BA_peak_4572 |
| DS113782 | 39084 | 39745 | BA_peak_4573 |
| DS113782 | 40802 | 41480 | BA_peak_4574 |
| DS113782 | 44330 | 45275 | BA_peak_4575 |
| DS113782 | 48784 | 49279 | BA_peak_4576 |
| DS113783 | 260 | 1970 | BA_peak_4577 |
| DS113783 | 5176 | 7836 | BA_peak_4578 |
| DS113783 | 14196 | 14592 | BA_peak_4579 |
| DS113783 | 16253 | 16701 | BA_peak_4580 |
| DS113783 | 43159 | 43688 | BA_peak_4581 |
| DS113784 | 19703 | 19964 | BA_peak_4582 |
| DS113784 | 38183 | 41171 | BA_peak_4583 |
| DS113785 | 13896 | 14995 | BA_peak_4584 |

| | | | |
|---|---|---|---|
| DS113785 | 41264 | 41594 | BA_peak_4585 |
| DS113785 | 50415 | 50941 | BA_peak_4586 |
| DS113786 | 15224 | 16060 | BA_peak_4587 |
| DS113786 | 31236 | 32194 | BA_peak_4588 |
| DS113787 | 6195 | 9077 | BA_peak_4589 |
| DS113787 | 15098 | 15716 | BA_peak_4590 |
| DS113787 | 29377 | 29768 | BA_peak_4591 |
| DS113787 | 36442 | 36923 | BA_peak_4592 |
| DS113788 | 2661 | 3477 | BA_peak_4593 |
| DS113788 | 46056 | 47224 | BA_peak_4594 |
| DS113788 | 47645 | 48104 | BA_peak_4595 |
| DS113788 | 48411 | 49764 | BA_peak_4596 |
| DS113789 | 21163 | 21521 | BA_peak_4597 |
| DS113789 | 30587 | 31394 | BA_peak_4598 |
| DS113789 | 46763 | 47581 | BA_peak_4599 |
| DS113790 | 11549 | 11982 | BA_peak_4600 |
| DS113791 | 42218 | 43011 | BA_peak_4601 |
| DS113791 | 45147 | 47666 | BA_peak_4602 |
| DS113791 | 48565 | 48977 | BA_peak_4603 |
| DS113791 | 49668 | 50411 | BA_peak_4604 |
| DS113791 | 50803 | 51138 | BA_peak_4605 |
| DS113791 | 52706 | 54033 | BA_peak_4606 |
| DS113792 | 6163 | 6843 | BA_peak_4607 |
| DS113792 | 16181 | 17487 | BA_peak_4608 |
| DS113792 | 17917 | 18630 | BA_peak_4609 |
| DS113792 | 20032 | 21068 | BA_peak_4610 |
| DS113792 | 30708 | 31551 | BA_peak_4611 |
| DS113793 | 28904 | 29676 | BA_peak_4612 |
| DS113793 | 54300 | 54978 | BA_peak_4613 |
| DS113795 | 11862 | 12252 | BA_peak_4614 |
| DS113795 | 20000 | 22035 | BA_peak_4615 |
| DS113795 | 29564 | 30285 | BA_peak_4616 |
| DS113795 | 49255 | 49755 | BA_peak_4617 |
| DS113795 | 50180 | 50522 | BA_peak_4618 |
| DS113795 | 51053 | 51887 | BA_peak_4619 |
| DS113796 | 7742 | 8177 | BA_peak_4620 |
| DS113796 | 13199 | 14884 | BA_peak_4621 |
| DS113796 | 24721 | 25399 | BA_peak_4622 |
| DS113796 | 26044 | 26697 | BA_peak_4623 |
| DS113796 | 38047 | 38468 | BA_peak_4624 |
| DS113797 | 70 | 1454 | BA_peak_4626 |
| DS113797 | 9140 | 9576 | BA_peak_4627 |
| DS113797 | 17948 | 18316 | BA_peak_4628 |
| DS113797 | 37200 | 37639 | BA_peak_4629 |
| DS113798 | 12840 | 13162 | BA_peak_4630 |
| DS113798 | 13827 | 14608 | BA_peak_4631 |
| DS113798 | 50109 | 50546 | BA_peak_4632 |

| | | | |
|---|---|---|---|
| DS113798 | 51114 | 53574 | BA_peak_4633 |
| DS113799 | 66 | 1777 | BA_peak_4634 |
| DS113799 | 30728 | 31527 | BA_peak_4635 |
| DS113799 | 32531 | 32838 | BA_peak_4636 |
| DS113799 | 40207 | 41568 | BA_peak_4637 |
| DS113799 | 46247 | 46583 | BA_peak_4638 |
| DS113800 | 18978 | 19342 | BA_peak_4639 |
| DS113800 | 25745 | 26517 | BA_peak_4640 |
| DS113800 | 49555 | 49868 | BA_peak_4642 |
| DS113801 | 22496 | 22949 | BA_peak_4643 |
| DS113801 | 35071 | 35567 | BA_peak_4644 |
| DS113801 | 46144 | 46864 | BA_peak_4645 |
| DS113802 | 6387 | 6926 | BA_peak_4646 |
| DS113802 | 32115 | 32630 | BA_peak_4647 |
| DS113802 | 53459 | 53917 | BA_peak_4648 |
| DS113803 | 267 | 598 | BA_peak_4649 |
| DS113803 | 2023 | 2550 | BA_peak_4650 |
| DS113803 | 13524 | 14500 | BA_peak_4651 |
| DS113803 | 15171 | 15576 | BA_peak_4652 |
| DS113803 | 29350 | 29851 | BA_peak_4653 |
| DS113803 | 30589 | 32155 | BA_peak_4654 |
| DS113803 | 48325 | 48683 | BA_peak_4655 |
| DS113803 | 53372 | 53754 | BA_peak_4656 |
| DS113806 | 16202 | 16976 | BA_peak_4657 |
| DS113806 | 17554 | 18211 | BA_peak_4658 |
| DS113806 | 19399 | 19756 | BA_peak_4659 |
| DS113807 | 22174 | 24253 | BA_peak_4660 |
| DS113808 | 25169 | 26080 | BA_peak_4661 |
| DS113808 | 37604 | 37953 | BA_peak_4662 |
| DS113809 | 16055 | 16443 | BA_peak_4664 |
| DS113809 | 35430 | 36103 | BA_peak_4665 |
| DS113809 | 38948 | 39319 | BA_peak_4666 |
| DS113810 | 14883 | 15204 | BA_peak_4667 |
| DS113810 | 28294 | 29076 | BA_peak_4668 |
| DS113810 | 30033 | 30803 | BA_peak_4669 |
| DS113810 | 46314 | 46721 | BA_peak_4670 |
| DS113810 | 47085 | 47901 | BA_peak_4670 |
| DS113810 | 49320 | 49817 | BA_peak_4671 |
| DS113810 | 52298 | 52771 | BA_peak_4672 |
| DS113811 | 432 | 896 | BA_peak_4673 |
| DS113811 | 44998 | 47525 | BA_peak_4675 |
| DS113812 | 21120 | 22990 | BA_peak_4676 |
| DS113813 | 1537 | 1910 | BA_peak_4677 |
| DS113813 | 4080 | 5036 | BA_peak_4678 |
| DS113813 | 43603 | 46938 | BA_peak_4679 |
| DS113813 | 49876 | 50406 | BA_peak_4680 |
| DS113814 | 8391 | 8982 | BA_peak_4681 |

| | | | |
|---|---|---|---|
| DS113814 | 23542 | 23907 | BA_peak_4682 |
| DS113814 | 42723 | 43217 | BA_peak_4683 |
| DS113815 | 37377 | 37723 | BA_peak_4684 |
| DS113815 | 47163 | 47810 | BA_peak_4685 |
| DS113815 | 48483 | 49473 | BA_peak_4686 |
| DS113816 | 38727 | 39641 | BA_peak_4689 |
| DS113817 | 29 | 442 | BA_peak_4690 |
| DS113817 | 1263 | 2595 | BA_peak_4691 |
| DS113817 | 30685 | 31419 | BA_peak_4692 |
| DS113818 | 5622 | 6060 | BA_peak_4693 |
| DS113818 | 11647 | 13442 | BA_peak_4694 |
| DS113818 | 25404 | 25725 | BA_peak_4695 |
| DS113818 | 43173 | 43838 | BA_peak_4696 |
| DS113819 | 171 | 965 | BA_peak_4697 |
| DS113819 | 12913 | 16162 | BA_peak_4698 |
| DS113819 | 30738 | 31037 | BA_peak_4699 |
| DS113819 | 41281 | 41617 | BA_peak_4700 |
| DS113820 | 23159 | 23475 | BA_peak_4701 |
| DS113820 | 24890 | 25849 | BA_peak_4702 |
| DS113820 | 26862 | 27459 | BA_peak_4703 |
| DS113821 | 40340 | 40783 | BA_peak_4704 |
| DS113821 | 41592 | 42132 | BA_peak_4705 |
| DS113821 | 45587 | 46184 | BA_peak_4706 |
| DS113821 | 46669 | 47327 | BA_peak_4707 |
| DS113821 | 49071 | 49405 | BA_peak_4708 |
| DS113821 | 50709 | 51591 | BA_peak_4709 |
| DS113822 | 9446 | 10578 | BA_peak_4710 |
| DS113823 | 3755 | 4268 | BA_peak_4711 |
| DS113823 | 39252 | 40628 | BA_peak_4712 |
| DS113823 | 45603 | 46147 | BA_peak_4713 |
| DS113824 | 7175 | 7853 | BA_peak_4714 |
| DS113824 | 19530 | 20189 | BA_peak_4715 |
| DS113824 | 48620 | 50318 | BA_peak_4716 |
| DS113825 | 42461 | 42819 | BA_peak_4717 |
| DS113826 | 879 | 3135 | BA_peak_4718 |
| DS113826 | 4163 | 4510 | BA_peak_4719 |
| DS113826 | 5492 | 6192 | BA_peak_4720 |
| DS113826 | 37720 | 38106 | BA_peak_4721 |
| DS113827 | 985 | 1524 | BA_peak_4722 |
| DS113827 | 9700 | 11015 | BA_peak_4723 |
| DS113828 | 3304 | 3675 | BA_peak_4724 |
| DS113828 | 51260 | 51954 | BA_peak_4725 |
| DS113829 | 17 | 1106 | BA_peak_4726 |
| DS113829 | 3909 | 4763 | BA_peak_4727 |
| DS113829 | 45684 | 47162 | BA_peak_4728 |
| DS113830 | 6535 | 8212 | BA_peak_4729 |
| DS113830 | 20683 | 21243 | BA_peak_4730 |

| | | | |
|---|---|---|---|
| DS113830 | 22621 | 22963 | BA_peak_4731 |
| DS113830 | 41780 | 42117 | BA_peak_4732 |
| DS113831 | 8196 | 8950 | BA_peak_4733 |
| DS113831 | 10069 | 10799 | BA_peak_4734 |
| DS113832 | 19405 | 19766 | BA_peak_4735 |
| DS113832 | 44629 | 45331 | BA_peak_4736 |
| DS113833 | 24522 | 24910 | BA_peak_4738 |
| DS113834 | 40591 | 41291 | BA_peak_4739 |
| DS113835 | 16057 | 17274 | BA_peak_4740 |
| DS113835 | 33185 | 33852 | BA_peak_4741 |
| DS113836 | 12763 | 13915 | BA_peak_4742 |
| DS113836 | 16372 | 17330 | BA_peak_4743 |
| DS113836 | 20722 | 21116 | BA_peak_4744 |
| DS113836 | 24357 | 25201 | BA_peak_4745 |
| DS113836 | 43292 | 43654 | BA_peak_4746 |
| DS113836 | 44135 | 45066 | BA_peak_4747 |
| DS113836 | 45482 | 47666 | BA_peak_4748 |
| DS113837 | 4876 | 5367 | BA_peak_4749 |
| DS113837 | 20494 | 22530 | BA_peak_4750 |
| DS113837 | 26712 | 29961 | BA_peak_4751 |
| DS113837 | 32009 | 32324 | BA_peak_4752 |
| DS113837 | 36247 | 36513 | BA_peak_4753 |
| DS113838 | 2831 | 3624 | BA_peak_4755 |
| DS113838 | 26994 | 27602 | BA_peak_4756 |
| DS113838 | 32504 | 33289 | BA_peak_4757 |
| DS113838 | 35194 | 35973 | BA_peak_4758 |
| DS113839 | 9267 | 9939 | BA_peak_4759 |
| DS113839 | 43276 | 44183 | BA_peak_4760 |
| DS113840 | 3491 | 3834 | BA_peak_4761 |
| DS113840 | 8841 | 9171 | BA_peak_4762 |
| DS113840 | 13034 | 14314 | BA_peak_4763 |
| DS113841 | 4135 | 4905 | BA_peak_4765 |
| DS113841 | 42147 | 42965 | BA_peak_4767 |
| DS113842 | 5069 | 5422 | BA_peak_4768 |
| DS113842 | 7003 | 7627 | BA_peak_4769 |
| DS113842 | 35975 | 37345 | BA_peak_4770 |
| DS113843 | 13907 | 14536 | BA_peak_4771 |
| DS113843 | 26353 | 26760 | BA_peak_4773 |
| DS113843 | 43726 | 44778 | BA_peak_4774 |
| DS113843 | 45162 | 46171 | BA_peak_4774 |
| DS113844 | 9147 | 10779 | BA_peak_4775 |
| DS113844 | 23559 | 24332 | BA_peak_4776 |
| DS113844 | 45668 | 46272 | BA_peak_4777 |
| DS113844 | 46821 | 47772 | BA_peak_4778 |
| DS113844 | 49946 | 50748 | BA_peak_4779 |
| DS113845 | 37478 | 37888 | BA_peak_4781 |
| DS113845 | 40981 | 41285 | BA_peak_4782 |

| | | | |
|---|---|---|---|
| DS113845 | 42706 | 43363 | BA_peak_4783 |
| DS113846 | 7791 | 8566 | BA_peak_4784 |
| DS113846 | 9411 | 9764 | BA_peak_4785 |
| DS113846 | 15711 | 18913 | BA_peak_4786 |
| DS113846 | 19311 | 20209 | BA_peak_4787 |
| DS113846 | 21659 | 22095 | BA_peak_4788 |
| DS113846 | 39953 | 40369 | BA_peak_4789 |
| DS113847 | 33381 | 33996 | BA_peak_4790 |
| DS113847 | 41481 | 42751 | BA_peak_4791 |
| DS113847 | 45413 | 45773 | BA_peak_4792 |
| DS113848 | 7396 | 7864 | BA_peak_4793 |
| DS113848 | 10319 | 10745 | BA_peak_4794 |
| DS113848 | 18442 | 20239 | BA_peak_4795 |
| DS113848 | 21548 | 22551 | BA_peak_4796 |
| DS113848 | 31249 | 31969 | BA_peak_4797 |
| DS113848 | 38800 | 39351 | BA_peak_4798 |
| DS113849 | 2771 | 4313 | BA_peak_4800 |
| DS113849 | 5632 | 7450 | BA_peak_4801 |
| DS113849 | 23191 | 25746 | BA_peak_4802 |
| DS113849 | 31855 | 32669 | BA_peak_4803 |
| DS113849 | 47067 | 48130 | BA_peak_4804 |
| DS113849 | 48570 | 50256 | BA_peak_4805 |
| DS113850 | 4127 | 7557 | BA_peak_4806 |
| DS113850 | 26391 | 27111 | BA_peak_4807 |
| DS113851 | 3694 | 4539 | BA_peak_4808 |
| DS113851 | 6506 | 6875 | BA_peak_4809 |
| DS113851 | 11786 | 12493 | BA_peak_4810 |
| DS113851 | 49185 | 50246 | BA_peak_4811 |
| DS113852 | 9734 | 11061 | BA_peak_4812 |
| DS113852 | 43530 | 44013 | BA_peak_4813 |
| DS113852 | 46072 | 46422 | BA_peak_4814 |
| DS113853 | 6063 | 6871 | BA_peak_4815 |
| DS113853 | 10808 | 11172 | BA_peak_4816 |
| DS113853 | 46057 | 46982 | BA_peak_4817 |
| DS113853 | 47359 | 47774 | BA_peak_4817 |
| DS113854 | 16011 | 16933 | BA_peak_4818 |
| DS113854 | 48648 | 49581 | BA_peak_4819 |
| DS113855 | 25215 | 25969 | BA_peak_4820 |
| DS113855 | 44301 | 44843 | BA_peak_4821 |
| DS113856 | 17483 | 17742 | BA_peak_4822 |
| DS113857 | 22103 | 22458 | BA_peak_4823 |
| DS113857 | 30505 | 31175 | BA_peak_4824 |
| DS113858 | 682 | 1003 | BA_peak_4825 |
| DS113858 | 12489 | 12807 | BA_peak_4826 |
| DS113858 | 19439 | 20018 | BA_peak_4827 |
| DS113858 | 24422 | 25020 | BA_peak_4828 |
| DS113859 | 21245 | 21632 | BA_peak_4829 |

| | | | |
|---|---|---|---|
| DS113859 | 30783 | 31640 | BA_peak_4830 |
| DS113860 | 40422 | 40984 | BA_peak_4831 |
| DS113861 | 11608 | 12234 | BA_peak_4832 |
| DS113861 | 14744 | 15268 | BA_peak_4833 |
| DS113861 | 16518 | 16845 | BA_peak_4834 |
| DS113861 | 40833 | 41175 | BA_peak_4835 |
| DS113862 | 7116 | 8471 | BA_peak_4836 |
| DS113862 | 20789 | 21344 | BA_peak_4837 |
| DS113862 | 22615 | 22967 | BA_peak_4838 |
| DS113862 | 24846 | 25614 | BA_peak_4839 |
| DS113862 | 48132 | 49248 | BA_peak_4840 |
| DS113863 | 8073 | 8445 | BA_peak_4841 |
| DS113863 | 15883 | 16269 | BA_peak_4842 |
| DS113863 | 22193 | 22639 | BA_peak_4843 |
| DS113863 | 37759 | 39476 | BA_peak_4844 |
| DS113864 | 1129 | 1753 | BA_peak_4845 |
| DS113864 | 34203 | 35024 | BA_peak_4846 |
| DS113864 | 35698 | 36066 | BA_peak_4847 |
| DS113865 | 12700 | 13079 | BA_peak_4848 |
| DS113865 | 21604 | 22184 | BA_peak_4849 |
| DS113865 | 30724 | 31592 | BA_peak_4850 |
| DS113865 | 38608 | 38960 | BA_peak_4851 |
| DS113865 | 39961 | 40912 | BA_peak_4852 |
| DS113865 | 44291 | 45173 | BA_peak_4853 |
| DS113866 | 2754 | 3913 | BA_peak_4854 |
| DS113866 | 17403 | 18323 | BA_peak_4855 |
| DS113866 | 19809 | 21086 | BA_peak_4856 |
| DS113866 | 39118 | 40180 | BA_peak_4857 |
| DS113867 | 3472 | 3945 | BA_peak_4858 |
| DS113867 | 12218 | 13273 | BA_peak_4860 |
| DS113867 | 28903 | 29470 | BA_peak_4861 |
| DS113867 | 33768 | 34299 | BA_peak_4862 |
| DS113868 | 7396 | 8178 | BA_peak_4863 |
| DS113868 | 18273 | 18618 | BA_peak_4864 |
| DS113868 | 46688 | 47035 | BA_peak_4865 |
| DS113869 | 19785 | 23138 | BA_peak_4866 |
| DS113869 | 38140 | 39025 | BA_peak_4867 |
| DS113869 | 45754 | 46987 | BA_peak_4868 |
| DS113870 | 24344 | 25048 | BA_peak_4869 |
| DS113870 | 30752 | 31384 | BA_peak_4870 |
| DS113870 | 32370 | 32945 | BA_peak_4871 |
| DS113870 | 33977 | 34297 | BA_peak_4872 |
| DS113870 | 40320 | 40791 | BA_peak_4873 |
| DS113870 | 43485 | 44300 | BA_peak_4874 |
| DS113870 | 44697 | 45820 | BA_peak_4875 |
| DS113871 | 935 | 1318 | BA_peak_4876 |
| DS113871 | 1986 | 3490 | BA_peak_4877 |

| | | | |
|---|---|---|---|
| DS113871 | 45321 | 46524 | BA_peak_4878 |
| DS113872 | 9891 | 10471 | BA_peak_4879 |
| DS113873 | 1143 | 2179 | BA_peak_4880 |
| DS113873 | 7748 | 8334 | BA_peak_4881 |
| DS113873 | 21742 | 22344 | BA_peak_4882 |
| DS113873 | 41323 | 41800 | BA_peak_4883 |
| DS113873 | 42243 | 43893 | BA_peak_4884 |
| DS113874 | 14259 | 14595 | BA_peak_4887 |
| DS113874 | 19787 | 20634 | BA_peak_4888 |
| DS113874 | 31456 | 32549 | BA_peak_4889 |
| DS113874 | 33966 | 35064 | BA_peak_4890 |
| DS113874 | 38576 | 39677 | BA_peak_4891 |
| DS113874 | 41470 | 42364 | BA_peak_4892 |
| DS113875 | 10626 | 12119 | BA_peak_4893 |
| DS113875 | 18888 | 19448 | BA_peak_4894 |
| DS113875 | 32457 | 32940 | BA_peak_4895 |
| DS113875 | 41465 | 42028 | BA_peak_4896 |
| DS113875 | 43840 | 44392 | BA_peak_4897 |
| DS113877 | 2868 | 3172 | BA_peak_4898 |
| DS113877 | 5778 | 6231 | BA_peak_4899 |
| DS113877 | 18673 | 19518 | BA_peak_4900 |
| DS113877 | 27775 | 28572 | BA_peak_4901 |
| DS113878 | 29298 | 29983 | BA_peak_4902 |
| DS113878 | 40815 | 41685 | BA_peak_4903 |
| DS113879 | 11638 | 12318 | BA_peak_4904 |
| DS113879 | 26755 | 27561 | BA_peak_4905 |
| DS113880 | 961 | 1303 | BA_peak_4906 |
| DS113880 | 13285 | 14322 | BA_peak_4907 |
| DS113880 | 14801 | 15156 | BA_peak_4908 |
| DS113880 | 28080 | 28680 | BA_peak_4909 |
| DS113880 | 29621 | 30674 | BA_peak_4910 |
| DS113881 | 35684 | 36146 | BA_peak_4911 |
| DS113881 | 40494 | 40999 | BA_peak_4912 |
| DS113881 | 43248 | 43561 | BA_peak_4913 |
| DS113882 | 39174 | 39810 | BA_peak_4915 |
| DS113882 | 44407 | 44723 | BA_peak_4916 |
| DS113882 | 46580 | 47242 | BA_peak_4917 |
| DS113883 | 1181 | 2322 | BA_peak_4918 |
| DS113883 | 5069 | 5974 | BA_peak_4919 |
| DS113883 | 37195 | 37951 | BA_peak_4921 |
| DS113883 | 40835 | 41750 | BA_peak_4922 |
| DS113884 | 3027 | 3342 | BA_peak_4923 |
| DS113884 | 30276 | 30930 | BA_peak_4924 |
| DS113885 | 15332 | 16097 | BA_peak_4925 |
| DS113885 | 17324 | 17848 | BA_peak_4926 |
| DS113887 | 3722 | 4746 | BA_peak_4928 |
| DS113887 | 32534 | 32980 | BA_peak_4929 |

| | | | |
|---|---|---|---|
| DS113887 | 33446 | 34367 | BA_peak_4930 |
| DS113887 | 35136 | 35590 | BA_peak_4931 |
| DS113887 | 42176 | 42798 | BA_peak_4932 |
| DS113887 | 43724 | 44303 | BA_peak_4933 |
| DS113888 | 13407 | 14054 | BA_peak_4935 |
| DS113888 | 26917 | 27306 | BA_peak_4936 |
| DS113888 | 38093 | 39461 | BA_peak_4937 |
| DS113888 | 43450 | 43755 | BA_peak_4938 |
| DS113889 | 4146 | 4463 | BA_peak_4939 |
| DS113889 | 22970 | 23674 | BA_peak_4940 |
| DS113889 | 25383 | 25792 | BA_peak_4941 |
| DS113889 | 28155 | 28541 | BA_peak_4942 |
| DS113889 | 30044 | 30974 | BA_peak_4943 |
| DS113889 | 37318 | 37985 | BA_peak_4944 |
| DS113889 | 40869 | 41394 | BA_peak_4945 |
| DS113890 | 6370 | 8929 | BA_peak_4946 |
| DS113890 | 18418 | 19193 | BA_peak_4947 |
| DS113890 | 34151 | 35456 | BA_peak_4948 |
| DS113891 | 30377 | 30860 | BA_peak_4949 |
| DS113892 | 3547 | 6326 | BA_peak_4950 |
| DS113893 | 11450 | 11885 | BA_peak_4951 |
| DS113893 | 19591 | 20023 | BA_peak_4952 |
| DS113893 | 43525 | 44089 | BA_peak_4953 |
| DS113895 | 157 | 726 | BA_peak_4954 |
| DS113895 | 19407 | 19815 | BA_peak_4955 |
| DS113895 | 42794 | 43147 | BA_peak_4956 |
| DS113896 | 24095 | 24658 | BA_peak_4957 |
| DS113896 | 30008 | 30406 | BA_peak_4958 |
| DS113897 | 20368 | 20761 | BA_peak_4961 |
| DS113897 | 28443 | 28861 | BA_peak_4962 |
| DS113898 | 1702 | 2089 | BA_peak_4963 |
| DS113898 | 26355 | 27225 | BA_peak_4964 |
| DS113898 | 31943 | 32288 | BA_peak_4965 |
| DS113899 | 2698 | 3041 | BA_peak_4966 |
| DS113899 | 6692 | 7279 | BA_peak_4967 |
| DS113899 | 13719 | 14171 | BA_peak_4968 |
| DS113899 | 24586 | 26471 | BA_peak_4969 |
| DS113899 | 35549 | 36114 | BA_peak_4970 |
| DS113899 | 45432 | 45986 | BA_peak_4971 |
| DS113900 | 9681 | 10064 | BA_peak_4972 |
| DS113900 | 11198 | 11769 | BA_peak_4973 |
| DS113901 | 33852 | 34432 | BA_peak_4974 |
| DS113901 | 37167 | 37794 | BA_peak_4975 |
| DS113902 | 4137 | 5509 | BA_peak_4976 |
| DS113902 | 10304 | 10954 | BA_peak_4977 |
| DS113902 | 28539 | 29166 | BA_peak_4978 |
| DS113903 | 8442 | 8867 | BA_peak_4979 |

| | | | |
|---|---|---|---|
| DS113903 | 12531 | 12868 | BA_peak_4980 |
| DS113903 | 35879 | 37624 | BA_peak_4981 |
| DS113904 | 1315 | 2777 | BA_peak_4983 |
| DS113904 | 5245 | 6049 | BA_peak_4984 |
| DS113904 | 10162 | 10505 | BA_peak_4985 |
| DS113904 | 42191 | 42874 | BA_peak_4986 |
| DS113904 | 43680 | 45034 | BA_peak_4987 |
| DS113905 | 14906 | 15274 | BA_peak_4988 |
| DS113905 | 45130 | 45867 | BA_peak_4989 |
| DS113906 | 38 | 1140 | BA_peak_4990 |
| DS113906 | 4398 | 5450 | BA_peak_4991 |
| DS113906 | 5800 | 7506 | BA_peak_4992 |
| DS113906 | 8711 | 9148 | BA_peak_4993 |
| DS113906 | 11949 | 12681 | BA_peak_4994 |
| DS113906 | 24566 | 25236 | BA_peak_4995 |
| DS113906 | 26265 | 26710 | BA_peak_4996 |
| DS113906 | 27351 | 28157 | BA_peak_4997 |
| DS113906 | 37570 | 38173 | BA_peak_4998 |
| DS113907 | 5331 | 5942 | BA_peak_4999 |
| DS113907 | 8635 | 9020 | BA_peak_5000 |
| DS113907 | 45162 | 45758 | BA_peak_5001 |
| DS113908 | 8857 | 9655 | BA_peak_5002 |
| DS113908 | 21890 | 23575 | BA_peak_5003 |
| DS113909 | 10270 | 10812 | BA_peak_5004 |
| DS113910 | 7707 | 8537 | BA_peak_5005 |
| DS113910 | 12162 | 12862 | BA_peak_5006 |
| DS113910 | 31699 | 32605 | BA_peak_5007 |
| DS113910 | 35428 | 35949 | BA_peak_5008 |
| DS113910 | 40709 | 41465 | BA_peak_5009 |
| DS113911 | 1761 | 2374 | BA_peak_5010 |
| DS113911 | 9500 | 11028 | BA_peak_5011 |
| DS113911 | 11963 | 12303 | BA_peak_5012 |
| DS113911 | 20439 | 22526 | BA_peak_5013 |
| DS113912 | 119 | 618 | BA_peak_5014 |
| DS113913 | 24314 | 24715 | BA_peak_5015 |
| DS113913 | 28993 | 29818 | BA_peak_5016 |
| DS113914 | 6010 | 6666 | BA_peak_5017 |
| DS113914 | 16178 | 16950 | BA_peak_5018 |
| DS113914 | 22982 | 24646 | BA_peak_5019 |
| DS113914 | 29361 | 29694 | BA_peak_5020 |
| DS113915 | 1728 | 2434 | BA_peak_5021 |
| DS113915 | 4852 | 6611 | BA_peak_5022 |
| DS113915 | 7454 | 7890 | BA_peak_5023 |
| DS113915 | 12966 | 14182 | BA_peak_5024 |
| DS113915 | 33071 | 33622 | BA_peak_5025 |
| DS113916 | 14953 | 15488 | BA_peak_5027 |
| DS113917 | 18184 | 18915 | BA_peak_5028 |

| | | | |
|---|---|---|---|
| DS113917 | 32220 | 33050 | BA_peak_5029 |
| DS113919 | 33584 | 34198 | BA_peak_5030 |
| DS113919 | 42743 | 43386 | BA_peak_5031 |
| DS113920 | 15779 | 17128 | BA_peak_5032 |
| DS113920 | 21976 | 22761 | BA_peak_5033 |
| DS113920 | 25769 | 26709 | BA_peak_5034 |
| DS113920 | 31020 | 31814 | BA_peak_5035 |
| DS113920 | 32194 | 32668 | BA_peak_5036 |
| DS113921 | 3243 | 3636 | BA_peak_5037 |
| DS113921 | 18989 | 20487 | BA_peak_5038 |
| DS113921 | 31908 | 32898 | BA_peak_5039 |
| DS113922 | 3777 | 5129 | BA_peak_5040 |
| DS113923 | 772 | 1144 | BA_peak_5041 |
| DS113923 | 28325 | 28848 | BA_peak_5042 |
| DS113923 | 35339 | 35676 | BA_peak_5043 |
| DS113923 | 41437 | 42833 | BA_peak_5044 |
| DS113925 | 1055 | 1643 | BA_peak_5045 |
| DS113925 | 4940 | 5289 | BA_peak_5046 |
| DS113925 | 16343 | 17030 | BA_peak_5047 |
| DS113925 | 36528 | 37836 | BA_peak_5048 |
| DS113926 | 1465 | 3421 | BA_peak_5049 |
| DS113926 | 4960 | 6395 | BA_peak_5050 |
| DS113926 | 43492 | 44100 | BA_peak_5052 |
| DS113927 | 11537 | 13746 | BA_peak_5053 |
| DS113927 | 22048 | 23127 | BA_peak_5054 |
| DS113927 | 41580 | 42061 | BA_peak_5055 |
| DS113927 | 43766 | 44137 | BA_peak_5056 |
| DS113930 | 4071 | 4894 | BA_peak_5057 |
| DS113930 | 15343 | 15890 | BA_peak_5058 |
| DS113930 | 34316 | 34637 | BA_peak_5059 |
| DS113931 | 17790 | 18120 | BA_peak_5060 |
| DS113931 | 18698 | 19433 | BA_peak_5061 |
| DS113931 | 33554 | 34575 | BA_peak_5062 |
| DS113932 | 10741 | 11462 | BA_peak_5063 |
| DS113932 | 20729 | 21617 | BA_peak_5064 |
| DS113932 | 42601 | 43453 | BA_peak_5065 |
| DS113933 | 18615 | 19020 | BA_peak_5066 |
| DS113933 | 38653 | 39557 | BA_peak_5067 |
| DS113935 | 42440 | 43085 | BA_peak_5068 |
| DS113936 | 12757 | 13396 | BA_peak_5069 |
| DS113936 | 33373 | 33831 | BA_peak_5070 |
| DS113936 | 34324 | 34698 | BA_peak_5071 |
| DS113937 | 10840 | 11691 | BA_peak_5073 |
| DS113937 | 13547 | 13916 | BA_peak_5074 |
| DS113938 | 302 | 606 | BA_peak_5075 |
| DS113938 | 3604 | 5064 | BA_peak_5076 |
| DS113938 | 5584 | 6170 | BA_peak_5077 |

| | | | |
|---|---|---|---|
| DS113938 | 14794 | 15124 | BA_peak_5078 |
| DS113938 | 23035 | 26378 | BA_peak_5079 |
| DS113938 | 41981 | 43275 | BA_peak_5080 |
| DS113939 | 1648 | 2218 | BA_peak_5081 |
| DS113939 | 2603 | 4084 | BA_peak_5082 |
| DS113939 | 14506 | 14864 | BA_peak_5083 |
| DS113940 | 21067 | 22222 | BA_peak_5084 |
| DS113940 | 32421 | 33121 | BA_peak_5085 |
| DS113940 | 35989 | 36479 | BA_peak_5086 |
| DS113941 | 3960 | 5206 | BA_peak_5087 |
| DS113941 | 15596 | 16282 | BA_peak_5088 |
| DS113941 | 32654 | 33194 | BA_peak_5089 |
| DS113942 | 6864 | 8154 | BA_peak_5090 |
| DS113942 | 24096 | 24764 | BA_peak_5091 |
| DS113943 | 20944 | 21268 | BA_peak_5092 |
| DS113944 | 10068 | 10449 | BA_peak_5093 |
| DS113944 | 11570 | 13184 | BA_peak_5094 |
| DS113944 | 13977 | 14632 | BA_peak_5095 |
| DS113945 | 4360 | 4733 | BA_peak_5096 |
| DS113945 | 15133 | 15552 | BA_peak_5097 |
| DS113945 | 17230 | 18190 | BA_peak_5098 |
| DS113945 | 21827 | 22287 | BA_peak_5099 |
| DS113946 | 8546 | 9144 | BA_peak_5100 |
| DS113946 | 20318 | 20922 | BA_peak_5101 |
| DS113946 | 36389 | 36743 | BA_peak_5102 |
| DS113947 | 1478 | 2114 | BA_peak_5103 |
| DS113947 | 30911 | 31273 | BA_peak_5104 |
| DS113947 | 34043 | 35250 | BA_peak_5105 |
| DS113948 | 431 | 985 | BA_peak_5106 |
| DS113948 | 7041 | 9150 | BA_peak_5107 |
| DS113948 | 13842 | 14478 | BA_peak_5108 |
| DS113948 | 27991 | 28453 | BA_peak_5109 |
| DS113948 | 35993 | 38280 | BA_peak_5110 |
| DS113949 | 16573 | 17108 | BA_peak_5111 |
| DS113949 | 32898 | 34004 | BA_peak_5112 |
| DS113949 | 34714 | 35392 | BA_peak_5113 |
| DS113949 | 36351 | 36733 | BA_peak_5114 |
| DS113950 | 31297 | 32265 | BA_peak_5115 |
| DS113950 | 32938 | 33352 | BA_peak_5116 |
| DS113951 | 3739 | 4065 | BA_peak_5117 |
| DS113951 | 6357 | 6744 | BA_peak_5118 |
| DS113951 | 7055 | 7379 | BA_peak_5119 |
| DS113951 | 16108 | 16620 | BA_peak_5120 |
| DS113951 | 22676 | 23045 | BA_peak_5121 |
| DS113951 | 29117 | 29507 | BA_peak_5122 |
| DS113951 | 35139 | 35444 | BA_peak_5124 |
| DS113952 | 136 | 1736 | BA_peak_5125 |

| | | | |
|---|---|---|---|
| DS113952 | 31470 | 32297 | BA_peak_5126 |
| DS113952 | 35283 | 36221 | BA_peak_5127 |
| DS113953 | 2559 | 3811 | BA_peak_5128 |
| DS113954 | 948 | 1429 | BA_peak_5130 |
| DS113954 | 9174 | 10290 | BA_peak_5131 |
| DS113954 | 13738 | 14561 | BA_peak_5132 |
| DS113955 | 26364 | 26788 | BA_peak_5133 |
| DS113955 | 40919 | 41853 | BA_peak_5134 |
| DS113956 | 7897 | 8201 | BA_peak_5135 |
| DS113956 | 20841 | 22231 | BA_peak_5138 |
| DS113956 | 23636 | 24301 | BA_peak_5139 |
| DS113958 | 4628 | 5521 | BA_peak_5140 |
| DS113958 | 14992 | 15399 | BA_peak_5141 |
| DS113960 | 3462 | 3875 | BA_peak_5142 |
| DS113960 | 4996 | 5956 | BA_peak_5143 |
| DS113960 | 6527 | 7133 | BA_peak_5144 |
| DS113960 | 18284 | 18751 | BA_peak_5145 |
| DS113960 | 38618 | 38996 | BA_peak_5146 |
| DS113960 | 39569 | 39899 | BA_peak_5147 |
| DS113961 | 3708 | 5571 | BA_peak_5148 |
| DS113961 | 6091 | 6470 | BA_peak_5149 |
| DS113961 | 13506 | 15286 | BA_peak_5150 |
| DS113961 | 18385 | 18838 | BA_peak_5151 |
| DS113961 | 16159 | 17434 | BA_peak_5151 |
| DS113961 | 20208 | 20521 | BA_peak_5152 |
| DS113961 | 37510 | 37906 | BA_peak_5153 |
| DS113961 | 39598 | 39937 | BA_peak_5154 |
| DS113962 | 9859 | 10338 | BA_peak_5155 |
| DS113962 | 10677 | 11163 | BA_peak_5156 |
| DS113963 | 39848 | 40544 | BA_peak_5158 |
| DS113964 | 8519 | 9213 | BA_peak_5159 |
| DS113964 | 17608 | 18559 | BA_peak_5160 |
| DS113964 | 22983 | 23939 | BA_peak_5161 |
| DS113964 | 25849 | 26444 | BA_peak_5162 |
| DS113964 | 27543 | 28154 | BA_peak_5163 |
| DS113965 | 24495 | 25117 | BA_peak_5164 |
| DS113966 | 2087 | 2593 | BA_peak_5165 |
| DS113966 | 18045 | 18962 | BA_peak_5166 |
| DS113966 | 34326 | 34914 | BA_peak_5167 |
| DS113967 | 17036 | 17857 | BA_peak_5168 |
| DS113968 | 6663 | 7311 | BA_peak_5169 |
| DS113968 | 11805 | 12204 | BA_peak_5170 |
| DS113969 | 3228 | 3648 | BA_peak_5171 |
| DS113970 | 565 | 926 | BA_peak_5172 |
| DS113970 | 30367 | 30828 | BA_peak_5173 |
| DS113971 | 6369 | 7034 | BA_peak_5174 |
| DS113971 | 7610 | 7960 | BA_peak_5175 |

| | | | |
|---|---|---|---|
| DS113971 | 24245 | 25298 | BA_peak_5176 |
| DS113971 | 25831 | 27710 | BA_peak_5177 |
| DS113971 | 33899 | 34277 | BA_peak_5178 |
| DS113971 | 34870 | 36835 | BA_peak_5179 |
| DS113972 | 11326 | 11762 | BA_peak_5180 |
| DS113972 | 20836 | 21559 | BA_peak_5181 |
| DS113972 | 25436 | 25966 | BA_peak_5182 |
| DS113972 | 30013 | 31084 | BA_peak_5183 |
| DS113972 | 38217 | 38846 | BA_peak_5185 |
| DS113974 | 32315 | 33410 | BA_peak_5186 |
| DS113974 | 34111 | 35379 | BA_peak_5187 |
| DS113974 | 37060 | 37604 | BA_peak_5188 |
| DS113975 | 8251 | 8586 | BA_peak_5189 |
| DS113975 | 8928 | 9262 | BA_peak_5190 |
| DS113975 | 10962 | 11463 | BA_peak_5191 |
| DS113975 | 28712 | 29119 | BA_peak_5192 |
| DS113975 | 29455 | 30242 | BA_peak_5193 |
| DS113975 | 34110 | 34776 | BA_peak_5194 |
| DS113975 | 36377 | 36724 | BA_peak_5195 |
| DS113975 | 40046 | 40349 | BA_peak_5196 |
| DS113976 | 7091 | 8462 | BA_peak_5197 |
| DS113976 | 10834 | 11195 | BA_peak_5198 |
| DS113976 | 19918 | 20279 | BA_peak_5199 |
| DS113976 | 22412 | 23741 | BA_peak_5200 |
| DS113976 | 25912 | 27866 | BA_peak_5201 |
| DS113976 | 28196 | 29276 | BA_peak_5202 |
| DS113976 | 31999 | 32861 | BA_peak_5203 |
| DS113977 | 5353 | 6005 | BA_peak_5204 |
| DS113977 | 6317 | 6836 | BA_peak_5205 |
| DS113977 | 10229 | 10589 | BA_peak_5206 |
| DS113977 | 11075 | 11788 | BA_peak_5207 |
| DS113977 | 37803 | 38554 | BA_peak_5208 |
| DS113978 | 8188 | 8650 | BA_peak_5209 |
| DS113979 | 14759 | 15321 | BA_peak_5210 |
| DS113979 | 16209 | 16613 | BA_peak_5211 |
| DS113979 | 20579 | 21243 | BA_peak_5213 |
| DS113979 | 23453 | 23865 | BA_peak_5214 |
| DS113979 | 24274 | 24656 | BA_peak_5215 |
| DS113979 | 30344 | 30773 | BA_peak_5216 |
| DS113979 | 33507 | 34362 | BA_peak_5218 |
| DS113979 | 38940 | 39287 | BA_peak_5219 |
| DS113980 | 22745 | 23259 | BA_peak_5220 |
| DS113980 | 23577 | 24015 | BA_peak_5221 |
| DS113980 | 30907 | 31414 | BA_peak_5222 |
| DS113980 | 32222 | 34555 | BA_peak_5223 |
| DS113980 | 35923 | 36465 | BA_peak_5224 |
| DS113981 | 22550 | 23429 | BA_peak_5225 |

| | | | |
|---|---|---|---|
| DS113981 | 27028 | 27389 | BA_peak_5226 |
| DS113981 | 32221 | 34453 | BA_peak_5227 |
| DS113982 | 23714 | 24233 | BA_peak_5228 |
| DS113982 | 25946 | 26677 | BA_peak_5229 |
| DS113982 | 27506 | 27859 | BA_peak_5230 |
| DS113983 | 121 | 946 | BA_peak_5231 |
| DS113983 | 10601 | 12124 | BA_peak_5232 |
| DS113983 | 16833 | 17640 | BA_peak_5233 |
| DS113983 | 25380 | 26033 | BA_peak_5234 |
| DS113983 | 30386 | 31024 | BA_peak_5235 |
| DS113984 | 5151 | 5548 | BA_peak_5236 |
| DS113984 | 14922 | 15483 | BA_peak_5237 |
| DS113985 | 3986 | 5122 | BA_peak_5238 |
| DS113985 | 6633 | 7461 | BA_peak_5239 |
| DS113985 | 14647 | 15379 | BA_peak_5240 |
| DS113985 | 25631 | 27173 | BA_peak_5241 |
| DS113985 | 33680 | 34095 | BA_peak_5242 |
| DS113986 | 24912 | 26453 | BA_peak_5243 |
| DS113987 | 2273 | 2931 | BA_peak_5244 |
| DS113987 | 3412 | 4115 | BA_peak_5245 |
| DS113987 | 20243 | 21669 | BA_peak_5246 |
| DS113987 | 23516 | 23990 | BA_peak_5247 |
| DS113987 | 24401 | 24933 | BA_peak_5248 |
| DS113988 | 14728 | 15338 | BA_peak_5249 |
| DS113988 | 22510 | 23279 | BA_peak_5250 |
| DS113988 | 23586 | 25157 | BA_peak_5251 |
| DS113988 | 31193 | 31763 | BA_peak_5252 |
| DS113989 | 7653 | 8035 | BA_peak_5254 |
| DS113990 | 12175 | 12964 | BA_peak_5255 |
| DS113990 | 15833 | 16289 | BA_peak_5256 |
| DS113990 | 29431 | 30214 | BA_peak_5257 |
| DS113991 | 17554 | 18237 | BA_peak_5258 |
| DS113991 | 23539 | 24278 | BA_peak_5259 |
| DS113991 | 25157 | 26573 | BA_peak_5260 |
| DS113991 | 31279 | 32692 | BA_peak_5261 |
| DS113991 | 33080 | 33598 | BA_peak_5262 |
| DS113992 | 11807 | 12531 | BA_peak_5263 |
| DS113992 | 25551 | 26089 | BA_peak_5264 |
| DS113992 | 27844 | 28167 | BA_peak_5265 |
| DS113992 | 29291 | 29950 | BA_peak_5266 |
| DS113993 | 11028 | 11645 | BA_peak_5267 |
| DS113994 | 25316 | 26114 | BA_peak_5269 |
| DS113994 | 33339 | 33810 | BA_peak_5270 |
| DS113994 | 36645 | 36955 | BA_peak_5271 |
| DS113995 | 18186 | 18530 | BA_peak_5272 |
| DS113996 | 3146 | 3468 | BA_peak_5273 |
| DS113996 | 5005 | 5908 | BA_peak_5274 |

| | | | |
|---|---|---|---|
| DS113996 | 6246 | 6733 | BA_peak_5275 |
| DS113996 | 27331 | 28045 | BA_peak_5276 |
| DS113997 | 19385 | 20206 | BA_peak_5277 |
| DS113997 | 24123 | 24906 | BA_peak_5278 |
| DS113997 | 25341 | 26237 | BA_peak_5279 |
| DS113997 | 29770 | 30205 | BA_peak_5280 |
| DS113998 | 3453 | 4331 | BA_peak_5281 |
| DS113998 | 7794 | 8503 | BA_peak_5282 |
| DS113998 | 28574 | 29112 | BA_peak_5283 |
| DS113999 | 25346 | 26154 | BA_peak_5284 |
| DS114001 | 14352 | 14717 | BA_peak_5285 |
| DS114001 | 15134 | 15502 | BA_peak_5285 |
| DS114002 | 3693 | 4176 | BA_peak_5286 |
| DS114002 | 5523 | 6341 | BA_peak_5287 |
| DS114002 | 14931 | 15474 | BA_peak_5288 |
| DS114002 | 34100 | 34905 | BA_peak_5289 |
| DS114004 | 13923 | 14784 | BA_peak_5290 |
| DS114004 | 19283 | 19985 | BA_peak_5291 |
| DS114004 | 26219 | 26831 | BA_peak_5292 |
| DS114004 | 27375 | 27813 | BA_peak_5293 |
| DS114004 | 29163 | 30211 | BA_peak_5294 |
| DS114004 | 36023 | 36632 | BA_peak_5295 |
| DS114005 | 8606 | 9216 | BA_peak_5296 |
| DS114005 | 12862 | 13456 | BA_peak_5297 |
| DS114005 | 13973 | 14411 | BA_peak_5298 |
| DS114006 | 23302 | 24202 | BA_peak_5300 |
| DS114007 | 2069 | 2631 | BA_peak_5301 |
| DS114007 | 32923 | 33485 | BA_peak_5302 |
| DS114007 | 36023 | 36462 | BA_peak_5303 |
| DS114008 | 26015 | 26347 | BA_peak_5304 |
| DS114009 | 9971 | 10366 | BA_peak_5305 |
| DS114009 | 11320 | 11858 | BA_peak_5306 |
| DS114009 | 12529 | 12927 | BA_peak_5307 |
| DS114009 | 16545 | 17216 | BA_peak_5308 |
| DS114009 | 19165 | 19529 | BA_peak_5309 |
| DS114009 | 19897 | 20705 | BA_peak_5310 |
| DS114010 | 20003 | 20398 | BA_peak_5311 |
| DS114012 | 5050 | 6062 | BA_peak_5312 |
| DS114012 | 9758 | 10437 | BA_peak_5313 |
| DS114013 | 4500 | 5858 | BA_peak_5315 |
| DS114014 | 16015 | 16974 | BA_peak_5316 |
| DS114014 | 21433 | 22290 | BA_peak_5317 |
| DS114014 | 23316 | 24111 | BA_peak_5318 |
| DS114014 | 24766 | 27517 | BA_peak_5319 |
| DS114015 | 425 | 1368 | BA_peak_5320 |
| DS114015 | 16594 | 17861 | BA_peak_5321 |
| DS114015 | 23619 | 24346 | BA_peak_5322 |

| | | | |
|---|---|---|---|
| DS114015 | 26090 | 26810 | BA_peak_5323 |
| DS114015 | 27946 | 30036 | BA_peak_5324 |
| DS114016 | 3706 | 4080 | BA_peak_5325 |
| DS114016 | 30897 | 31714 | BA_peak_5326 |
| DS114017 | 3158 | 3646 | BA_peak_5327 |
| DS114017 | 19644 | 20493 | BA_peak_5328 |
| DS114017 | 27971 | 28316 | BA_peak_5329 |
| DS114017 | 34455 | 35499 | BA_peak_5330 |
| DS114019 | 733 | 1418 | BA_peak_5331 |
| DS114019 | 11236 | 11905 | BA_peak_5332 |
| DS114019 | 12604 | 15040 | BA_peak_5333 |
| DS114020 | 1832 | 2294 | BA_peak_5334 |
| DS114021 | 1684 | 4378 | BA_peak_5335 |
| DS114021 | 4820 | 5472 | BA_peak_5336 |
| DS114021 | 5893 | 6698 | BA_peak_5337 |
| DS114021 | 16145 | 18039 | BA_peak_5338 |
| DS114021 | 20886 | 21193 | BA_peak_5339 |
| DS114021 | 30145 | 31364 | BA_peak_5340 |
| DS114022 | 601 | 952 | BA_peak_5341 |
| DS114022 | 3789 | 4328 | BA_peak_5342 |
| DS114022 | 32152 | 32493 | BA_peak_5343 |
| DS114022 | 32983 | 33477 | BA_peak_5344 |
| DS114022 | 35403 | 36000 | BA_peak_5345 |
| DS114023 | 34131 | 34910 | BA_peak_5346 |
| DS114023 | 35466 | 36157 | BA_peak_5347 |
| DS114024 | 652 | 991 | BA_peak_5348 |
| DS114025 | 33861 | 34315 | BA_peak_5349 |
| DS114026 | 609 | 1160 | BA_peak_5350 |
| DS114026 | 11342 | 11861 | BA_peak_5351 |
| DS114026 | 24707 | 25333 | BA_peak_5352 |
| DS114027 | 5590 | 6131 | BA_peak_5353 |
| DS114027 | 13938 | 14357 | BA_peak_5354 |
| DS114028 | 4400 | 4867 | BA_peak_5355 |
| DS114028 | 5379 | 7220 | BA_peak_5355 |
| DS114028 | 33744 | 34988 | BA_peak_5356 |
| DS114030 | 780 | 1087 | BA_peak_5358 |
| DS114030 | 14530 | 15017 | BA_peak_5359 |
| DS114030 | 15388 | 16242 | BA_peak_5360 |
| DS114031 | 28023 | 28401 | BA_peak_5361 |
| DS114032 | 10078 | 11441 | BA_peak_5362 |
| DS114032 | 27864 | 28791 | BA_peak_5363 |
| DS114032 | 29167 | 29473 | BA_peak_5364 |
| DS114032 | 31780 | 33037 | BA_peak_5365 |
| DS114033 | 4808 | 5605 | BA_peak_5366 |
| DS114033 | 6766 | 7452 | BA_peak_5367 |
| DS114033 | 16335 | 17162 | BA_peak_5368 |
| DS114033 | 18545 | 20573 | BA_peak_5369 |

| | | | |
|---|---|---|---|
| DS114033 | 20877 | 21686 | BA_peak_5370 |
| DS114033 | 26020 | 26526 | BA_peak_5371 |
| DS114034 | 3898 | 4733 | BA_peak_5372 |
| DS114034 | 6256 | 7418 | BA_peak_5373 |
| DS114035 | 19825 | 20187 | BA_peak_5374 |
| DS114036 | 1380 | 1773 | BA_peak_5376 |
| DS114037 | 3816 | 4711 | BA_peak_5377 |
| DS114037 | 8395 | 9921 | BA_peak_5378 |
| DS114037 | 14594 | 16643 | BA_peak_5379 |
| DS114037 | 28980 | 29777 | BA_peak_5380 |
| DS114038 | 26275 | 26713 | BA_peak_5381 |
| DS114039 | 92 | 613 | BA_peak_5382 |
| DS114039 | 2294 | 2899 | BA_peak_5383 |
| DS114039 | 5895 | 6283 | BA_peak_5384 |
| DS114039 | 29956 | 30676 | BA_peak_5385 |
| DS114040 | 30193 | 30971 | BA_peak_5386 |
| DS114041 | 4106 | 5329 | BA_peak_5387 |
| DS114041 | 5674 | 7313 | BA_peak_5387 |
| DS114041 | 7787 | 8468 | BA_peak_5388 |
| DS114042 | 3308 | 5343 | BA_peak_5389 |
| DS114042 | 5729 | 6548 | BA_peak_5390 |
| DS114043 | 13694 | 14107 | BA_peak_5391 |
| DS114043 | 27365 | 27692 | BA_peak_5392 |
| DS114043 | 32200 | 32786 | BA_peak_5393 |
| DS114044 | 9273 | 9642 | BA_peak_5394 |
| DS114045 | 26575 | 28337 | BA_peak_5398 |
| DS114045 | 28776 | 29348 | BA_peak_5399 |
| DS114046 | 9729 | 10152 | BA_peak_5400 |
| DS114046 | 10933 | 11736 | BA_peak_5401 |
| DS114046 | 18032 | 18688 | BA_peak_5402 |
| DS114046 | 23087 | 26023 | BA_peak_5403 |
| DS114047 | 26878 | 27222 | BA_peak_5404 |
| DS114048 | 11728 | 13277 | BA_peak_5405 |
| DS114049 | 9143 | 10076 | BA_peak_5406 |
| DS114049 | 21048 | 21422 | BA_peak_5407 |
| DS114050 | 20720 | 21511 | BA_peak_5408 |
| DS114051 | 18537 | 18888 | BA_peak_5409 |
| DS114052 | 8834 | 9905 | BA_peak_5410 |
| DS114052 | 21333 | 22037 | BA_peak_5411 |
| DS114054 | 32611 | 33196 | BA_peak_5413 |
| DS114056 | 6069 | 6962 | BA_peak_5414 |
| DS114056 | 7816 | 8490 | BA_peak_5415 |
| DS114056 | 8906 | 9908 | BA_peak_5416 |
| DS114056 | 12936 | 13673 | BA_peak_5418 |
| DS114056 | 14245 | 14665 | BA_peak_5419 |
| DS114056 | 25555 | 26509 | BA_peak_5420 |
| DS114057 | 6170 | 6512 | BA_peak_5421 |

| | | | |
|---|---|---|---|
| DS114057 | 15411 | 17906 | BA_peak_5422 |
| DS114058 | 9465 | 10527 | BA_peak_5423 |
| DS114058 | 14226 | 14990 | BA_peak_5424 |
| DS114058 | 15914 | 16320 | BA_peak_5425 |
| DS114058 | 29476 | 30843 | BA_peak_5426 |
| DS114060 | 12306 | 12963 | BA_peak_5427 |
| DS114060 | 24789 | 25471 | BA_peak_5428 |
| DS114060 | 27188 | 27767 | BA_peak_5429 |
| DS114061 | 250 | 528 | BA_peak_5430 |
| DS114061 | 975 | 1591 | BA_peak_5431 |
| DS114061 | 19666 | 20042 | BA_peak_5432 |
| DS114061 | 26842 | 27212 | BA_peak_5433 |
| DS114061 | 27557 | 27943 | BA_peak_5434 |
| DS114062 | 1866 | 2478 | BA_peak_5435 |
| DS114062 | 26724 | 27266 | BA_peak_5437 |
| DS114063 | 8720 | 9287 | BA_peak_5438 |
| DS114063 | 10698 | 11489 | BA_peak_5439 |
| DS114063 | 12449 | 13058 | BA_peak_5440 |
| DS114063 | 13833 | 14239 | BA_peak_5441 |
| DS114063 | 22050 | 22820 | BA_peak_5442 |
| DS114064 | 18150 | 18579 | BA_peak_5443 |
| DS114064 | 23776 | 24172 | BA_peak_5444 |
| DS114064 | 24497 | 25442 | BA_peak_5445 |
| DS114064 | 25750 | 26918 | BA_peak_5446 |
| DS114065 | 2893 | 3777 | BA_peak_5447 |
| DS114066 | 1917 | 3757 | BA_peak_5449 |
| DS114066 | 9607 | 10757 | BA_peak_5450 |
| DS114068 | 6599 | 7435 | BA_peak_5452 |
| DS114069 | 1948 | 3074 | BA_peak_5453 |
| DS114069 | 6671 | 7309 | BA_peak_5454 |
| DS114069 | 10476 | 11399 | BA_peak_5455 |
| DS114069 | 11802 | 12265 | BA_peak_5456 |
| DS114069 | 15038 | 15610 | BA_peak_5457 |
| DS114069 | 25670 | 27103 | BA_peak_5458 |
| DS114071 | 50 | 530 | BA_peak_5459 |
| DS114071 | 1189 | 1925 | BA_peak_5460 |
| DS114071 | 26343 | 26970 | BA_peak_5461 |
| DS114071 | 27643 | 28699 | BA_peak_5462 |
| DS114071 | 31755 | 32212 | BA_peak_5463 |
| DS114073 | 11668 | 12214 | BA_peak_5464 |
| DS114073 | 16542 | 17088 | BA_peak_5465 |
| DS114074 | 21569 | 22208 | BA_peak_5466 |
| DS114074 | 22842 | 23231 | BA_peak_5467 |
| DS114074 | 30817 | 31933 | BA_peak_5468 |
| DS114075 | 26633 | 27432 | BA_peak_5469 |
| DS114075 | 29688 | 30099 | BA_peak_5470 |
| DS114075 | 30528 | 31003 | BA_peak_5471 |

| | | | |
|---|---|---|---|
| DS114076 | 5930 | 6272 | BA_peak_5472 |
| DS114076 | 20724 | 22201 | BA_peak_5473 |
| DS114076 | 23884 | 25637 | BA_peak_5474 |
| DS114076 | 26366 | 26854 | BA_peak_5475 |
| DS114076 | 31413 | 31734 | BA_peak_5476 |
| DS114077 | 164 | 664 | BA_peak_5477 |
| DS114077 | 32062 | 32539 | BA_peak_5478 |
| DS114078 | 15999 | 16566 | BA_peak_5479 |
| DS114079 | 5959 | 6944 | BA_peak_5481 |
| DS114079 | 18556 | 20226 | BA_peak_5483 |
| DS114080 | 19079 | 19493 | BA_peak_5484 |
| DS114080 | 22828 | 23648 | BA_peak_5485 |
| DS114081 | 2759 | 3202 | BA_peak_5486 |
| DS114081 | 19821 | 20431 | BA_peak_5487 |
| DS114082 | 31549 | 32533 | BA_peak_5488 |
| DS114083 | 10442 | 11727 | BA_peak_5489 |
| DS114083 | 18955 | 19511 | BA_peak_5490 |
| DS114083 | 19825 | 21044 | BA_peak_5491 |
| DS114084 | 5425 | 5893 | BA_peak_5492 |
| DS114084 | 8327 | 8815 | BA_peak_5493 |
| DS114084 | 27079 | 28302 | BA_peak_5494 |
| DS114085 | 4239 | 4571 | BA_peak_5496 |
| DS114085 | 14348 | 14655 | BA_peak_5497 |
| DS114086 | 4888 | 5461 | BA_peak_5498 |
| DS114087 | 29670 | 30364 | BA_peak_5499 |
| DS114087 | 31489 | 32372 | BA_peak_5500 |
| DS114088 | 11805 | 12630 | BA_peak_5501 |
| DS114088 | 30017 | 30633 | BA_peak_5502 |
| DS114088 | 31148 | 31632 | BA_peak_5503 |
| DS114090 | 40 | 3085 | BA_peak_5504 |
| DS114090 | 5656 | 6590 | BA_peak_5505 |
| DS114090 | 16421 | 18198 | BA_peak_5506 |
| DS114091 | 28203 | 28741 | BA_peak_5507 |
| DS114092 | 3138 | 5116 | BA_peak_5508 |
| DS114092 | 5923 | 6878 | BA_peak_5509 |
| DS114092 | 7538 | 7842 | BA_peak_5510 |
| DS114092 | 20431 | 21105 | BA_peak_5511 |
| DS114093 | 2932 | 3275 | BA_peak_5512 |
| DS114093 | 19458 | 19929 | BA_peak_5513 |
| DS114094 | 149 | 639 | BA_peak_5514 |
| DS114094 | 1973 | 2407 | BA_peak_5515 |
| DS114094 | 27280 | 27651 | BA_peak_5516 |
| DS114095 | 6752 | 7373 | BA_peak_5517 |
| DS114097 | 12809 | 17393 | BA_peak_5518 |
| DS114097 | 23824 | 24434 | BA_peak_5519 |
| DS114098 | 20097 | 20781 | BA_peak_5520 |
| DS114099 | 30201 | 31238 | BA_peak_5521 |

| | | | |
|---|---|---|---|
| DS114100 | 8886 | 9313 | BA_peak_5522 |
| DS114101 | 7805 | 8874 | BA_peak_5523 |
| DS114102 | 326 | 1198 | BA_peak_5524 |
| DS114102 | 14353 | 15038 | BA_peak_5525 |
| DS114103 | 758 | 1189 | BA_peak_5526 |
| DS114104 | 11531 | 11912 | BA_peak_5527 |
| DS114104 | 24195 | 24730 | BA_peak_5528 |
| DS114105 | 101 | 571 | BA_peak_5529 |
| DS114105 | 6907 | 8654 | BA_peak_5530 |
| DS114105 | 12517 | 14357 | BA_peak_5531 |
| DS114105 | 15293 | 16354 | BA_peak_5532 |
| DS114105 | 27585 | 28838 | BA_peak_5533 |
| DS114106 | 4805 | 6288 | BA_peak_5534 |
| DS114106 | 9813 | 10380 | BA_peak_5535 |
| DS114109 | 18157 | 18778 | BA_peak_5536 |
| DS114109 | 24949 | 25722 | BA_peak_5537 |
| DS114110 | 12226 | 12619 | BA_peak_5538 |
| DS114110 | 14449 | 15408 | BA_peak_5539 |
| DS114112 | 5100 | 5441 | BA_peak_5540 |
| DS114114 | 7164 | 8037 | BA_peak_5541 |
| DS114114 | 17498 | 18341 | BA_peak_5542 |
| DS114114 | 19142 | 19646 | BA_peak_5543 |
| DS114115 | 1938 | 2326 | BA_peak_5544 |
| DS114115 | 25718 | 26540 | BA_peak_5545 |
| DS114116 | 11047 | 11585 | BA_peak_5546 |
| DS114116 | 24511 | 25753 | BA_peak_5547 |
| DS114116 | 26627 | 26963 | BA_peak_5548 |
| DS114117 | 349 | 743 | BA_peak_5549 |
| DS114117 | 1349 | 1669 | BA_peak_5550 |
| DS114117 | 14993 | 15388 | BA_peak_5551 |
| DS114118 | 23194 | 24285 | BA_peak_5552 |
| DS114120 | 19521 | 20495 | BA_peak_5553 |
| DS114121 | 5987 | 7228 | BA_peak_5554 |
| DS114121 | 26548 | 27435 | BA_peak_5555 |
| DS114122 | 3989 | 4297 | BA_peak_5556 |
| DS114123 | 2514 | 3238 | BA_peak_5557 |
| DS114123 | 5017 | 6558 | BA_peak_5559 |
| DS114123 | 7432 | 8725 | BA_peak_5560 |
| DS114125 | 9107 | 9917 | BA_peak_5562 |
| DS114126 | 227 | 1972 | BA_peak_5563 |
| DS114127 | 115 | 730 | BA_peak_5564 |
| DS114127 | 3372 | 4013 | BA_peak_5565 |
| DS114127 | 8079 | 9240 | BA_peak_5566 |
| DS114127 | 20719 | 21494 | BA_peak_5567 |
| DS114127 | 24906 | 28123 | BA_peak_5568 |
| DS114128 | 147 | 1576 | BA_peak_5569 |
| DS114128 | 13595 | 13997 | BA_peak_5570 |

| | | | |
|---|---|---|---|
| DS114128 | 14606 | 15530 | BA_peak_5571 |
| DS114129 | 20051 | 20356 | BA_peak_5572 |
| DS114129 | 23878 | 24213 | BA_peak_5573 |
| DS114130 | 1910 | 2818 | BA_peak_5574 |
| DS114130 | 4512 | 4890 | BA_peak_5575 |
| DS114130 | 28556 | 28976 | BA_peak_5576 |
| DS114132 | 16429 | 16817 | BA_peak_5578 |
| DS114132 | 18680 | 19880 | BA_peak_5579 |
| DS114133 | 13226 | 14026 | BA_peak_5580 |
| DS114133 | 20338 | 20662 | BA_peak_5581 |
| DS114133 | 28929 | 29650 | BA_peak_5582 |
| DS114134 | 7917 | 9095 | BA_peak_5583 |
| DS114134 | 20468 | 21375 | BA_peak_5584 |
| DS114136 | 5961 | 6315 | BA_peak_5585 |
| DS114136 | 22281 | 22913 | BA_peak_5586 |
| DS114136 | 26120 | 26534 | BA_peak_5587 |
| DS114136 | 28593 | 29823 | BA_peak_5588 |
| DS114137 | 6757 | 7543 | BA_peak_5589 |
| DS114138 | 4873 | 5245 | BA_peak_5590 |
| DS114138 | 11153 | 11658 | BA_peak_5591 |
| DS114138 | 13002 | 13709 | BA_peak_5592 |
| DS114138 | 24539 | 25192 | BA_peak_5593 |
| DS114138 | 29287 | 29627 | BA_peak_5594 |
| DS114140 | 16673 | 17077 | BA_peak_5595 |
| DS114141 | 186 | 539 | BA_peak_5596 |
| DS114141 | 25313 | 25954 | BA_peak_5598 |
| DS114143 | 17400 | 18789 | BA_peak_5599 |
| DS114143 | 22744 | 23160 | BA_peak_5600 |
| DS114144 | 12988 | 16645 | BA_peak_5601 |
| DS114145 | 6181 | 7205 | BA_peak_5602 |
| DS114146 | 12806 | 13351 | BA_peak_5603 |
| DS114146 | 18764 | 19150 | BA_peak_5604 |
| DS114146 | 19646 | 20172 | BA_peak_5605 |
| DS114147 | 16346 | 16812 | BA_peak_5606 |
| DS114149 | 13073 | 13676 | BA_peak_5607 |
| DS114149 | 21854 | 22281 | BA_peak_5608 |
| DS114149 | 26130 | 27127 | BA_peak_5609 |
| DS114150 | 168 | 474 | BA_peak_5610 |
| DS114150 | 5440 | 5886 | BA_peak_5611 |
| DS114150 | 7143 | 8053 | BA_peak_5612 |
| DS114150 | 10682 | 11839 | BA_peak_5613 |
| DS114150 | 28171 | 28711 | BA_peak_5614 |
| DS114151 | 23344 | 24269 | BA_peak_5616 |
| DS114152 | 3664 | 4942 | BA_peak_5617 |
| DS114152 | 5352 | 6586 | BA_peak_5618 |
| DS114153 | 685 | 1178 | BA_peak_5619 |
| DS114153 | 1517 | 2981 | BA_peak_5620 |

| | | | |
|---|---|---|---|
| DS114155 | 6079 | 6671 | BA_peak_5621 |
| DS114155 | 19037 | 19456 | BA_peak_5622 |
| DS114156 | 17338 | 17736 | BA_peak_5623 |
| DS114157 | 24833 | 26163 | BA_peak_5624 |
| DS114158 | 4210 | 6636 | BA_peak_5625 |
| DS114158 | 7565 | 7961 | BA_peak_5626 |
| DS114158 | 8980 | 9397 | BA_peak_5627 |
| DS114158 | 10852 | 11620 | BA_peak_5628 |
| DS114159 | 15897 | 16250 | BA_peak_5629 |
| DS114159 | 25401 | 26333 | BA_peak_5630 |
| DS114159 | 26676 | 27145 | BA_peak_5631 |
| DS114160 | 18594 | 19141 | BA_peak_5632 |
| DS114161 | 1358 | 1847 | BA_peak_5633 |
| DS114161 | 12542 | 13060 | BA_peak_5634 |
| DS114161 | 25368 | 25897 | BA_peak_5635 |
| DS114162 | 351 | 705 | BA_peak_5636 |
| DS114162 | 8722 | 13344 | BA_peak_5637 |
| DS114162 | 20674 | 20983 | BA_peak_5638 |
| DS114164 | 4044 | 6143 | BA_peak_5639 |
| DS114164 | 7724 | 8525 | BA_peak_5640 |
| DS114164 | 19783 | 20383 | BA_peak_5641 |
| DS114165 | 4081 | 4666 | BA_peak_5642 |
| DS114165 | 13313 | 14075 | BA_peak_5643 |
| DS114165 | 21027 | 21707 | BA_peak_5644 |
| DS114165 | 24370 | 25201 | BA_peak_5645 |
| DS114166 | 16159 | 16736 | BA_peak_5646 |
| DS114166 | 18056 | 18494 | BA_peak_5647 |
| DS114166 | 18842 | 19344 | BA_peak_5648 |
| DS114167 | 17922 | 18399 | BA_peak_5649 |
| DS114169 | 7193 | 7690 | BA_peak_5652 |
| DS114169 | 21925 | 22574 | BA_peak_5653 |
| DS114170 | 16931 | 17449 | BA_peak_5655 |
| DS114170 | 21374 | 22702 | BA_peak_5656 |
| DS114170 | 27149 | 27582 | BA_peak_5658 |
| DS114171 | 81 | 653 | BA_peak_5659 |
| DS114171 | 2942 | 3652 | BA_peak_5660 |
| DS114171 | 4152 | 4828 | BA_peak_5661 |
| DS114172 | 11812 | 13653 | BA_peak_5662 |
| DS114172 | 13966 | 14979 | BA_peak_5663 |
| DS114172 | 20041 | 20332 | BA_peak_5664 |
| DS114173 | 11220 | 11967 | BA_peak_5665 |
| DS114174 | 90 | 418 | BA_peak_5666 |
| DS114174 | 939 | 1839 | BA_peak_5667 |
| DS114174 | 7111 | 7631 | BA_peak_5668 |
| DS114174 | 8354 | 8819 | BA_peak_5669 |
| DS114174 | 22923 | 24050 | BA_peak_5670 |
| DS114176 | 6043 | 7310 | BA_peak_5672 |

| | | | |
|---|---|---|---|
| DS114176 | 20023 | 20641 | BA_peak_5673 |
| DS114176 | 23677 | 25459 | BA_peak_5674 |
| DS114176 | 26158 | 27709 | BA_peak_5675 |
| DS114177 | 7308 | 8530 | BA_peak_5676 |
| DS114178 | 13636 | 14554 | BA_peak_5677 |
| DS114178 | 26708 | 27271 | BA_peak_5678 |
| DS114179 | 8495 | 9150 | BA_peak_5679 |
| DS114180 | 11794 | 13240 | BA_peak_5680 |
| DS114180 | 13854 | 14875 | BA_peak_5681 |
| DS114181 | 6452 | 6969 | BA_peak_5682 |
| DS114182 | 6735 | 7262 | BA_peak_5684 |
| DS114182 | 18036 | 18451 | BA_peak_5685 |
| DS114183 | 11451 | 12142 | BA_peak_5686 |
| DS114183 | 12659 | 14539 | BA_peak_5687 |
| DS114183 | 14970 | 15419 | BA_peak_5688 |
| DS114184 | 158 | 1443 | BA_peak_5690 |
| DS114185 | 2651 | 4223 | BA_peak_5691 |
| DS114185 | 14568 | 15108 | BA_peak_5692 |
| DS114185 | 16354 | 16850 | BA_peak_5693 |
| DS114186 | 94 | 2086 | BA_peak_5694 |
| DS114186 | 17187 | 17719 | BA_peak_5695 |
| DS114186 | 26127 | 26811 | BA_peak_5696 |
| DS114187 | 1315 | 1789 | BA_peak_5697 |
| DS114187 | 14116 | 14756 | BA_peak_5698 |
| DS114187 | 21351 | 22710 | BA_peak_5699 |
| DS114187 | 25154 | 25668 | BA_peak_5700 |
| DS114188 | 2044 | 2632 | BA_peak_5701 |
| DS114188 | 19968 | 20707 | BA_peak_5702 |
| DS114190 | 10053 | 10728 | BA_peak_5703 |
| DS114190 | 17762 | 18146 | BA_peak_5704 |
| DS114191 | 9219 | 10027 | BA_peak_5705 |
| DS114192 | 2891 | 3578 | BA_peak_5706 |
| DS114192 | 22629 | 22962 | BA_peak_5707 |
| DS114193 | 9445 | 10304 | BA_peak_5708 |
| DS114193 | 19382 | 19905 | BA_peak_5709 |
| DS114196 | 2528 | 2967 | BA_peak_5710 |
| DS114196 | 8310 | 8814 | BA_peak_5711 |
| DS114197 | 5928 | 7079 | BA_peak_5712 |
| DS114197 | 17889 | 18394 | BA_peak_5713 |
| DS114199 | 7083 | 7703 | BA_peak_5714 |
| DS114200 | 16473 | 17215 | BA_peak_5715 |
| DS114200 | 19699 | 21178 | BA_peak_5716 |
| DS114200 | 26218 | 26750 | BA_peak_5717 |
| DS114201 | 1445 | 2035 | BA_peak_5718 |
| DS114201 | 14249 | 14652 | BA_peak_5719 |
| DS114202 | 16137 | 16523 | BA_peak_5720 |
| DS114202 | 21909 | 22283 | BA_peak_5721 |

| | | | |
|---|---|---|---|
| DS114203 | 5170 | 5620 | BA_peak_5722 |
| DS114203 | 6947 | 7415 | BA_peak_5723 |
| DS114204 | 11470 | 11869 | BA_peak_5724 |
| DS114205 | 23151 | 24166 | BA_peak_5725 |
| DS114205 | 25262 | 25680 | BA_peak_5726 |
| DS114206 | 10984 | 12418 | BA_peak_5727 |
| DS114207 | 16902 | 17704 | BA_peak_5728 |
| DS114207 | 15061 | 16558 | BA_peak_5728 |
| DS114207 | 26076 | 26460 | BA_peak_5729 |
| DS114208 | 6012 | 6902 | BA_peak_5730 |
| DS114209 | 25957 | 26466 | BA_peak_5731 |
| DS114210 | 1271 | 1784 | BA_peak_5732 |
| DS114210 | 20473 | 20877 | BA_peak_5733 |
| DS114212 | 13850 | 14126 | BA_peak_5734 |
| DS114212 | 14794 | 15520 | BA_peak_5735 |
| DS114213 | 5236 | 5988 | BA_peak_5736 |
| DS114213 | 13337 | 14284 | BA_peak_5737 |
| DS114213 | 15525 | 16104 | BA_peak_5738 |
| DS114214 | 15377 | 15804 | BA_peak_5739 |
| DS114215 | 2749 | 3404 | BA_peak_5740 |
| DS114215 | 11124 | 11803 | BA_peak_5741 |
| DS114215 | 17145 | 17933 | BA_peak_5742 |
| DS114217 | 20793 | 21384 | BA_peak_5743 |
| DS114218 | 1255 | 1855 | BA_peak_5744 |
| DS114220 | 4300 | 5541 | BA_peak_5745 |
| DS114220 | 6096 | 6719 | BA_peak_5746 |
| DS114220 | 18940 | 19421 | BA_peak_5747 |
| DS114221 | 52 | 526 | BA_peak_5748 |
| DS114221 | 2036 | 4179 | BA_peak_5749 |
| DS114221 | 5127 | 6357 | BA_peak_5750 |
| DS114221 | 9680 | 10333 | BA_peak_5751 |
| DS114221 | 17559 | 18038 | BA_peak_5752 |
| DS114222 | 23326 | 23803 | BA_peak_5753 |
| DS114223 | 1707 | 3151 | BA_peak_5754 |
| DS114223 | 9074 | 9385 | BA_peak_5755 |
| DS114224 | 11611 | 12125 | BA_peak_5756 |
| DS114225 | 18697 | 19362 | BA_peak_5757 |
| DS114226 | 4383 | 5775 | BA_peak_5759 |
| DS114226 | 6096 | 6693 | BA_peak_5760 |
| DS114226 | 7115 | 8444 | BA_peak_5761 |
| DS114227 | 1075 | 2584 | BA_peak_5763 |
| DS114227 | 2974 | 3336 | BA_peak_5764 |
| DS114229 | 2169 | 3550 | BA_peak_5765 |
| DS114230 | 3091 | 3504 | BA_peak_5766 |
| DS114230 | 9123 | 9980 | BA_peak_5767 |
| DS114231 | 70 | 547 | BA_peak_5768 |
| DS114231 | 4011 | 5620 | BA_peak_5769 |

| | | | |
|---|---|---|---|
| DS114231 | 9209 | 9571 | BA_peak_5770 |
| DS114232 | 122 | 507 | BA_peak_5771 |
| DS114232 | 1925 | 2352 | BA_peak_5772 |
| DS114232 | 4554 | 5796 | BA_peak_5773 |
| DS114232 | 6589 | 7377 | BA_peak_5774 |
| DS114232 | 17120 | 18383 | BA_peak_5775 |
| DS114233 | 326 | 2164 | BA_peak_5776 |
| DS114233 | 10265 | 11601 | BA_peak_5777 |
| DS114233 | 22654 | 23078 | BA_peak_5778 |
| DS114234 | 5108 | 5870 | BA_peak_5779 |
| DS114234 | 24501 | 24951 | BA_peak_5780 |
| DS114236 | 22637 | 23049 | BA_peak_5782 |
| DS114237 | 1215 | 1573 | BA_peak_5783 |
| DS114237 | 2324 | 2920 | BA_peak_5784 |
| DS114237 | 22664 | 23924 | BA_peak_5785 |
| DS114239 | 4124 | 8302 | BA_peak_5786 |
| DS114239 | 21000 | 23149 | BA_peak_5787 |
| DS114239 | 24357 | 24903 | BA_peak_5788 |
| DS114240 | 14113 | 14536 | BA_peak_5789 |
| DS114240 | 15408 | 16354 | BA_peak_5790 |
| DS114240 | 22025 | 22705 | BA_peak_5791 |
| DS114242 | 217 | 633 | BA_peak_5792 |
| DS114242 | 11851 | 12205 | BA_peak_5793 |
| DS114243 | 4665 | 5165 | BA_peak_5794 |
| DS114243 | 21108 | 21784 | BA_peak_5795 |
| DS114244 | 3482 | 5159 | BA_peak_5796 |
| DS114244 | 15055 | 15949 | BA_peak_5798 |
| DS114245 | 5016 | 5690 | BA_peak_5800 |
| DS114245 | 17897 | 19560 | BA_peak_5801 |
| DS114245 | 20072 | 20722 | BA_peak_5802 |
| DS114246 | 18648 | 19078 | BA_peak_5804 |
| DS114246 | 19546 | 19976 | BA_peak_5805 |
| DS114247 | 19024 | 19709 | BA_peak_5806 |
| DS114247 | 21007 | 21456 | BA_peak_5807 |
| DS114248 | 77 | 525 | BA_peak_5808 |
| DS114248 | 1005 | 1981 | BA_peak_5809 |
| DS114248 | 2848 | 3967 | BA_peak_5810 |
| DS114248 | 13160 | 13593 | BA_peak_5811 |
| DS114248 | 18119 | 18709 | BA_peak_5812 |
| DS114248 | 20191 | 20806 | BA_peak_5813 |
| DS114249 | 11027 | 11560 | BA_peak_5814 |
| DS114249 | 21928 | 22283 | BA_peak_5815 |
| DS114249 | 22902 | 24518 | BA_peak_5816 |
| DS114250 | 1067 | 1761 | BA_peak_5817 |
| DS114251 | 4025 | 5043 | BA_peak_5818 |
| DS114251 | 6035 | 7522 | BA_peak_5819 |
| DS114251 | 8036 | 9061 | BA_peak_5820 |

| | | | |
|---|---|---|---|
| DS114251 | 19757 | 20169 | BA_peak_5821 |
| DS114252 | 95 | 786 | BA_peak_5822 |
| DS114252 | 22640 | 23299 | BA_peak_5823 |
| DS114252 | 23796 | 24414 | BA_peak_5824 |
| DS114253 | 21409 | 22454 | BA_peak_5825 |
| DS114255 | 7042 | 7811 | BA_peak_5826 |
| DS114257 | 2660 | 4196 | BA_peak_5827 |
| DS114257 | 12812 | 13382 | BA_peak_5828 |
| DS114257 | 14663 | 16148 | BA_peak_5829 |
| DS114258 | 8733 | 9172 | BA_peak_5830 |
| DS114258 | 12102 | 12603 | BA_peak_5831 |
| DS114261 | 102 | 824 | BA_peak_5832 |
| DS114261 | 2309 | 2774 | BA_peak_5833 |
| DS114261 | 21179 | 21619 | BA_peak_5834 |
| DS114262 | 3390 | 4078 | BA_peak_5835 |
| DS114262 | 9708 | 10845 | BA_peak_5836 |
| DS114262 | 19237 | 19581 | BA_peak_5837 |
| DS114263 | 8459 | 9432 | BA_peak_5838 |
| DS114264 | 8404 | 10246 | BA_peak_5839 |
| DS114265 | 15437 | 15786 | BA_peak_5840 |
| DS114265 | 21482 | 22549 | BA_peak_5841 |
| DS114266 | 17433 | 20886 | BA_peak_5842 |
| DS114267 | 23514 | 23842 | BA_peak_5843 |
| DS114268 | 1159 | 1645 | BA_peak_5844 |
| DS114269 | 9064 | 10073 | BA_peak_5845 |
| DS114269 | 13933 | 14409 | BA_peak_5846 |
| DS114270 | 725 | 1660 | BA_peak_5847 |
| DS114270 | 11027 | 12265 | BA_peak_5848 |
| DS114270 | 13178 | 13498 | BA_peak_5849 |
| DS114270 | 14284 | 14696 | BA_peak_5850 |
| DS114271 | 4612 | 5480 | BA_peak_5852 |
| DS114272 | 1488 | 2925 | BA_peak_5854 |
| DS114273 | 1368 | 3010 | BA_peak_5855 |
| DS114273 | 16248 | 16802 | BA_peak_5856 |
| DS114275 | 9326 | 9749 | BA_peak_5857 |
| DS114276 | 4949 | 5629 | BA_peak_5858 |
| DS114276 | 18599 | 19095 | BA_peak_5859 |
| DS114277 | 2208 | 4290 | BA_peak_5860 |
| DS114277 | 16364 | 16742 | BA_peak_5861 |
| DS114277 | 17083 | 18404 | BA_peak_5862 |
| DS114278 | 7380 | 9423 | BA_peak_5863 |
| DS114278 | 13913 | 15219 | BA_peak_5864 |
| DS114279 | 6768 | 7248 | BA_peak_5865 |
| DS114279 | 20245 | 20745 | BA_peak_5866 |
| DS114280 | 20283 | 21764 | BA_peak_5867 |
| DS114282 | 15435 | 16772 | BA_peak_5868 |
| DS114282 | 21914 | 22919 | BA_peak_5869 |

| | | | |
|---|---|---|---|
| DS114283 | 3212 | 3829 | BA_peak_5870 |
| DS114284 | 16064 | 18438 | BA_peak_5871 |
| DS114284 | 18911 | 19327 | BA_peak_5872 |
| DS114284 | 19782 | 21993 | BA_peak_5873 |
| DS114285 | 8270 | 8927 | BA_peak_5874 |
| DS114285 | 20314 | 20969 | BA_peak_5875 |
| DS114286 | 5958 | 6524 | BA_peak_5876 |
| DS114286 | 17167 | 19624 | BA_peak_5877 |
| DS114288 | 20028 | 20777 | BA_peak_5879 |
| DS114290 | 11547 | 11870 | BA_peak_5880 |
| DS114290 | 12992 | 13926 | BA_peak_5881 |
| DS114290 | 19598 | 20352 | BA_peak_5882 |
| DS114291 | 9369 | 10539 | BA_peak_5883 |
| DS114293 | 9242 | 10104 | BA_peak_5885 |
| DS114293 | 12824 | 13204 | BA_peak_5886 |
| DS114293 | 14596 | 14965 | BA_peak_5887 |
| DS114294 | 929 | 2183 | BA_peak_5888 |
| DS114294 | 12248 | 15596 | BA_peak_5889 |
| DS114294 | 19681 | 20323 | BA_peak_5890 |
| DS114295 | 982 | 1414 | BA_peak_5891 |
| DS114295 | 16651 | 17256 | BA_peak_5892 |
| DS114295 | 17739 | 18168 | BA_peak_5893 |
| DS114295 | 18836 | 19245 | BA_peak_5894 |
| DS114296 | 14244 | 15082 | BA_peak_5896 |
| DS114296 | 15960 | 16311 | BA_peak_5897 |
| DS114301 | 97 | 632 | BA_peak_5898 |
| DS114302 | 2111 | 2487 | BA_peak_5899 |
| DS114302 | 2951 | 3545 | BA_peak_5900 |
| DS114304 | 14073 | 15239 | BA_peak_5901 |
| DS114304 | 15694 | 16300 | BA_peak_5902 |
| DS114304 | 18479 | 19189 | BA_peak_5903 |
| DS114305 | 5826 | 6241 | BA_peak_5904 |
| DS114306 | 31 | 3213 | BA_peak_5905 |
| DS114306 | 4429 | 4846 | BA_peak_5906 |
| DS114307 | 1660 | 2257 | BA_peak_5907 |
| DS114308 | 2184 | 2577 | BA_peak_5908 |
| DS114308 | 20501 | 22213 | BA_peak_5909 |
| DS114309 | 16048 | 16448 | BA_peak_5910 |
| DS114310 | 13994 | 14485 | BA_peak_5911 |
| DS114311 | 6942 | 7821 | BA_peak_5912 |
| DS114311 | 13882 | 14328 | BA_peak_5913 |
| DS114312 | 21236 | 21583 | BA_peak_5914 |
| DS114313 | 16349 | 17188 | BA_peak_5915 |
| DS114317 | 3299 | 3797 | BA_peak_5916 |
| DS114317 | 5762 | 6505 | BA_peak_5917 |
| DS114317 | 7599 | 8052 | BA_peak_5918 |
| DS114317 | 9703 | 10480 | BA_peak_5919 |

| | | | |
|---|---|---|---|
| DS114317 | 11125 | 12780 | BA_peak_5920 |
| DS114320 | 21433 | 21883 | BA_peak_5921 |
| DS114321 | 2205 | 2793 | BA_peak_5922 |
| DS114321 | 17981 | 19992 | BA_peak_5923 |
| DS114322 | 7067 | 8259 | BA_peak_5924 |
| DS114322 | 9000 | 9826 | BA_peak_5925 |
| DS114322 | 10227 | 11120 | BA_peak_5926 |
| DS114323 | 16566 | 16955 | BA_peak_5927 |
| DS114323 | 20613 | 21149 | BA_peak_5928 |
| DS114326 | 6320 | 6776 | BA_peak_5929 |
| DS114327 | 3060 | 3491 | BA_peak_5930 |
| DS114327 | 17494 | 18585 | BA_peak_5931 |
| DS114328 | 14672 | 15196 | BA_peak_5932 |
| DS114328 | 15669 | 16002 | BA_peak_5933 |
| DS114329 | 9388 | 10775 | BA_peak_5934 |
| DS114331 | 4247 | 5658 | BA_peak_5935 |
| DS114331 | 11658 | 16559 | BA_peak_5936 |
| DS114332 | 13094 | 14212 | BA_peak_5938 |
| DS114332 | 15042 | 15867 | BA_peak_5939 |
| DS114332 | 17592 | 20160 | BA_peak_5940 |
| DS114333 | 712 | 1537 | BA_peak_5941 |
| DS114333 | 6570 | 8217 | BA_peak_5942 |
| DS114333 | 20673 | 21269 | BA_peak_5943 |
| DS114334 | 7552 | 7967 | BA_peak_5944 |
| DS114334 | 18241 | 18546 | BA_peak_5945 |
| DS114334 | 20691 | 21378 | BA_peak_5946 |
| DS114335 | 6726 | 9755 | BA_peak_5947 |
| DS114335 | 16980 | 17352 | BA_peak_5948 |
| DS114335 | 17999 | 18843 | BA_peak_5949 |
| DS114335 | 19389 | 20277 | BA_peak_5950 |
| DS114336 | 1648 | 1984 | BA_peak_5952 |
| DS114336 | 19861 | 20602 | BA_peak_5953 |
| DS114337 | 14095 | 14531 | BA_peak_5954 |
| DS114338 | 155 | 1632 | BA_peak_5955 |
| DS114339 | 988 | 1636 | BA_peak_5956 |
| DS114339 | 2200 | 3942 | BA_peak_5957 |
| DS114339 | 15027 | 15419 | BA_peak_5958 |
| DS114340 | 1207 | 1597 | BA_peak_5959 |
| DS114340 | 8477 | 9429 | BA_peak_5960 |
| DS114343 | 3059 | 3685 | BA_peak_5961 |
| DS114344 | 19708 | 20242 | BA_peak_5963 |
| DS114345 | 1972 | 3053 | BA_peak_5964 |
| DS114345 | 7277 | 7877 | BA_peak_5965 |
| DS114346 | 3471 | 3783 | BA_peak_5966 |
| DS114347 | 6906 | 7482 | BA_peak_5968 |
| DS114350 | 11750 | 12115 | BA_peak_5969 |
| DS114350 | 18116 | 20650 | BA_peak_5970 |

| | | | |
|---|---|---|---|
| DS114352 | 10593 | 11455 | BA_peak_5971 |
| DS114353 | 8502 | 9003 | BA_peak_5972 |
| DS114355 | 3583 | 4902 | BA_peak_5973 |
| DS114355 | 6807 | 7277 | BA_peak_5974 |
| DS114355 | 15216 | 15956 | BA_peak_5975 |
| DS114355 | 16380 | 17101 | BA_peak_5976 |
| DS114355 | 18405 | 18788 | BA_peak_5977 |
| DS114356 | 12628 | 13240 | BA_peak_5978 |
| DS114356 | 15058 | 16426 | BA_peak_5979 |
| DS114356 | 17407 | 18020 | BA_peak_5980 |
| DS114356 | 18790 | 19353 | BA_peak_5981 |
| DS114357 | 13589 | 14364 | BA_peak_5982 |
| DS114358 | 268 | 1372 | BA_peak_5983 |
| DS114358 | 6104 | 6407 | BA_peak_5984 |
| DS114360 | 662 | 1327 | BA_peak_5986 |
| DS114362 | 16492 | 17287 | BA_peak_5987 |
| DS114363 | 19534 | 19877 | BA_peak_5988 |
| DS114365 | 241 | 866 | BA_peak_5989 |
| DS114367 | 4606 | 5319 | BA_peak_5990 |
| DS114367 | 8295 | 8677 | BA_peak_5991 |
| DS114367 | 18461 | 18796 | BA_peak_5992 |
| DS114368 | 13609 | 14034 | BA_peak_5993 |
| DS114369 | 125 | 686 | BA_peak_5994 |
| DS114370 | 371 | 739 | BA_peak_5995 |
| DS114370 | 16809 | 17451 | BA_peak_5996 |
| DS114371 | 7692 | 8193 | BA_peak_5997 |
| DS114372 | 2165 | 2543 | BA_peak_5999 |
| DS114372 | 10119 | 10827 | BA_peak_6000 |
| DS114372 | 16813 | 17280 | BA_peak_6001 |
| DS114373 | 11718 | 12106 | BA_peak_6003 |
| DS114374 | 14287 | 14624 | BA_peak_6004 |
| DS114374 | 16420 | 16813 | BA_peak_6005 |
| DS114375 | 11784 | 12187 | BA_peak_6006 |
| DS114376 | 7844 | 8492 | BA_peak_6007 |
| DS114377 | 8146 | 8527 | BA_peak_6008 |
| DS114378 | 3093 | 3744 | BA_peak_6009 |
| DS114378 | 4756 | 5353 | BA_peak_6010 |
| DS114378 | 7488 | 8542 | BA_peak_6011 |
| DS114380 | 15459 | 17103 | BA_peak_6012 |
| DS114382 | 5018 | 5482 | BA_peak_6013 |
| DS114382 | 6184 | 7437 | BA_peak_6014 |
| DS114383 | 8907 | 9682 | BA_peak_6015 |
| DS114383 | 14985 | 15660 | BA_peak_6016 |
| DS114384 | 1922 | 2252 | BA_peak_6017 |
| DS114384 | 3153 | 4383 | BA_peak_6018 |
| DS114384 | 5016 | 5644 | BA_peak_6019 |
| DS114385 | 10929 | 12548 | BA_peak_6020 |

| | | | |
|---|---|---|---|
| DS114386 | 4424 | 7668 | BA_peak_6021 |
| DS114386 | 17051 | 18975 | BA_peak_6022 |
| DS114387 | 3526 | 3887 | BA_peak_6023 |
| DS114388 | 5383 | 5986 | BA_peak_6025 |
| DS114390 | 37 | 1228 | BA_peak_6026 |
| DS114390 | 4651 | 5299 | BA_peak_6027 |
| DS114390 | 13083 | 13520 | BA_peak_6028 |
| DS114392 | 3235 | 3734 | BA_peak_6029 |
| DS114393 | 2843 | 3315 | BA_peak_6030 |
| DS114393 | 7683 | 8074 | BA_peak_6031 |
| DS114394 | 762 | 1105 | BA_peak_6032 |
| DS114394 | 7121 | 7402 | BA_peak_6033 |
| DS114395 | 10848 | 11340 | BA_peak_6034 |
| DS114398 | 6086 | 6419 | BA_peak_6035 |
| DS114398 | 8778 | 9113 | BA_peak_6036 |
| DS114400 | 67 | 1137 | BA_peak_6037 |
| DS114400 | 8566 | 8947 | BA_peak_6038 |
| DS114400 | 9653 | 10127 | BA_peak_6039 |
| DS114400 | 13543 | 14200 | BA_peak_6040 |
| DS114400 | 18367 | 18937 | BA_peak_6041 |
| DS114401 | 3073 | 4675 | BA_peak_6042 |
| DS114401 | 12357 | 12729 | BA_peak_6043 |
| DS114403 | 13572 | 14149 | BA_peak_6044 |
| DS114405 | 7043 | 7430 | BA_peak_6045 |
| DS114406 | 17106 | 17734 | BA_peak_6047 |
| DS114407 | 235 | 643 | BA_peak_6048 |
| DS114407 | 8790 | 9949 | BA_peak_6049 |
| DS114407 | 10356 | 10804 | BA_peak_6050 |
| DS114408 | 16008 | 16291 | BA_peak_6052 |
| DS114409 | 92 | 1454 | BA_peak_6053 |
| DS114411 | 12658 | 13655 | BA_peak_6054 |
| DS114411 | 14230 | 14603 | BA_peak_6055 |
| DS114412 | 11967 | 12436 | BA_peak_6056 |
| DS114414 | 92 | 616 | BA_peak_6057 |
| DS114414 | 1378 | 1801 | BA_peak_6058 |
| DS114414 | 2549 | 3081 | BA_peak_6059 |
| DS114414 | 3504 | 4326 | BA_peak_6060 |
| DS114415 | 13028 | 13776 | BA_peak_6061 |
| DS114415 | 17582 | 17949 | BA_peak_6062 |
| DS114418 | 317 | 776 | BA_peak_6063 |
| DS114419 | 4774 | 5242 | BA_peak_6064 |
| DS114420 | 7053 | 7680 | BA_peak_6065 |
| DS114421 | 15617 | 16453 | BA_peak_6066 |
| DS114421 | 17295 | 17855 | BA_peak_6067 |
| DS114423 | 13997 | 14303 | BA_peak_6069 |
| DS114425 | 865 | 1653 | BA_peak_6070 |
| DS114425 | 7045 | 7592 | BA_peak_6071 |

| | | | |
|---|---|---|---|
| DS114425 | 12071 | 12625 | BA_peak_6072 |
| DS114426 | 8597 | 9451 | BA_peak_6073 |
| DS114426 | 17321 | 17735 | BA_peak_6074 |
| DS114428 | 3820 | 4575 | BA_peak_6075 |
| DS114429 | 4357 | 4874 | BA_peak_6076 |
| DS114429 | 11591 | 12823 | BA_peak_6077 |
| DS114430 | 392 | 798 | BA_peak_6078 |
| DS114430 | 1629 | 1961 | BA_peak_6079 |
| DS114430 | 11099 | 11543 | BA_peak_6080 |
| DS114430 | 15251 | 15568 | BA_peak_6081 |
| DS114430 | 16683 | 17540 | BA_peak_6082 |
| DS114431 | 840 | 1492 | BA_peak_6083 |
| DS114431 | 8842 | 9495 | BA_peak_6084 |
| DS114432 | 9588 | 10033 | BA_peak_6085 |
| DS114432 | 12007 | 12685 | BA_peak_6086 |
| DS114433 | 16188 | 17527 | BA_peak_6087 |
| DS114434 | 12617 | 13008 | BA_peak_6088 |
| DS114435 | 1569 | 4554 | BA_peak_6089 |
| DS114435 | 16132 | 16894 | BA_peak_6090 |
| DS114437 | 14238 | 14570 | BA_peak_6091 |
| DS114438 | 15876 | 17119 | BA_peak_6092 |
| DS114439 | 17009 | 17426 | BA_peak_6093 |
| DS114440 | 7505 | 7984 | BA_peak_6094 |
| DS114440 | 15994 | 17253 | BA_peak_6095 |
| DS114441 | 6437 | 6843 | BA_peak_6096 |
| DS114441 | 14780 | 15612 | BA_peak_6097 |
| DS114442 | 15606 | 17273 | BA_peak_6098 |
| DS114443 | 7942 | 8789 | BA_peak_6099 |
| DS114446 | 2207 | 2853 | BA_peak_6100 |
| DS114446 | 7536 | 7921 | BA_peak_6101 |
| DS114446 | 8334 | 8639 | BA_peak_6102 |
| DS114447 | 3244 | 3713 | BA_peak_6103 |
| DS114448 | 3435 | 3883 | BA_peak_6104 |
| DS114449 | 13348 | 13682 | BA_peak_6105 |
| DS114449 | 14358 | 15706 | BA_peak_6106 |
| DS114454 | 111 | 867 | BA_peak_6108 |
| DS114454 | 5038 | 5384 | BA_peak_6109 |
| DS114454 | 11800 | 12306 | BA_peak_6110 |
| DS114455 | 26 | 557 | BA_peak_6111 |
| DS114455 | 4221 | 4926 | BA_peak_6112 |
| DS114455 | 5391 | 7158 | BA_peak_6113 |
| DS114456 | 12970 | 13741 | BA_peak_6115 |
| DS114457 | 7564 | 8200 | BA_peak_6116 |
| DS114457 | 15390 | 16824 | BA_peak_6117 |
| DS114462 | 13790 | 14587 | BA_peak_6119 |
| DS114470 | 6937 | 7411 | BA_peak_6121 |
| DS114471 | 780 | 1144 | BA_peak_6122 |

| | | | |
|---|---|---|---|
| DS114472 | 7788 | 8295 | BA_peak_6124 |
| DS114473 | 4387 | 5494 | BA_peak_6125 |
| DS114473 | 7629 | 9036 | BA_peak_6126 |
| DS114473 | 14208 | 15459 | BA_peak_6127 |
| DS114477 | 640 | 1817 | BA_peak_6128 |
| DS114477 | 12969 | 13360 | BA_peak_6129 |
| DS114477 | 14922 | 15461 | BA_peak_6130 |
| DS114478 | 221 | 889 | BA_peak_6131 |
| DS114478 | 1275 | 1855 | BA_peak_6132 |
| DS114479 | 7888 | 8197 | BA_peak_6134 |
| DS114479 | 15598 | 16166 | BA_peak_6135 |
| DS114480 | 11566 | 12192 | BA_peak_6136 |
| DS114481 | 4147 | 4625 | BA_peak_6137 |
| DS114481 | 10398 | 10890 | BA_peak_6138 |
| DS114482 | 12046 | 12588 | BA_peak_6139 |
| DS114482 | 13485 | 13941 | BA_peak_6140 |
| DS114483 | 5606 | 6231 | BA_peak_6141 |
| DS114484 | 286 | 738 | BA_peak_6142 |
| DS114488 | 14195 | 15603 | BA_peak_6143 |
| DS114489 | 4024 | 4852 | BA_peak_6144 |
| DS114490 | 4285 | 4894 | BA_peak_6145 |
| DS114491 | 15233 | 15685 | BA_peak_6146 |
| DS114493 | 3078 | 4701 | BA_peak_6147 |
| DS114493 | 14799 | 15787 | BA_peak_6148 |
| DS114494 | 3707 | 4713 | BA_peak_6149 |
| DS114495 | 8755 | 10051 | BA_peak_6150 |
| DS114495 | 15295 | 15649 | BA_peak_6151 |
| DS114496 | 1878 | 2449 | BA_peak_6152 |
| DS114496 | 13482 | 13895 | BA_peak_6153 |
| DS114497 | 1945 | 2271 | BA_peak_6154 |
| DS114497 | 5871 | 6776 | BA_peak_6155 |
| DS114497 | 13022 | 14058 | BA_peak_6156 |
| DS114498 | 14130 | 15519 | BA_peak_6157 |
| DS114499 | 8546 | 9245 | BA_peak_6158 |
| DS114499 | 11555 | 12152 | BA_peak_6159 |
| DS114499 | 12473 | 12808 | BA_peak_6160 |
| DS114500 | 7963 | 8348 | BA_peak_6161 |
| DS114501 | 1125 | 2223 | BA_peak_6162 |
| DS114501 | 14365 | 14836 | BA_peak_6163 |
| DS114502 | 2518 | 4718 | BA_peak_6164 |
| DS114502 | 14259 | 14896 | BA_peak_6165 |
| DS114503 | 1508 | 1965 | BA_peak_6166 |
| DS114504 | 13755 | 14430 | BA_peak_6167 |
| DS114507 | 27 | 511 | BA_peak_6169 |
| DS114507 | 2090 | 2631 | BA_peak_6170 |
| DS114507 | 3008 | 3741 | BA_peak_6171 |
| DS114508 | 14354 | 15026 | BA_peak_6172 |

| | | | |
|---|---|---|---|
| DS114509 | 13602 | 14125 | BA_peak_6173 |
| DS114510 | 9752 | 10970 | BA_peak_6174 |
| DS114510 | 11323 | 12637 | BA_peak_6175 |
| DS114510 | 13256 | 13610 | BA_peak_6176 |
| DS114511 | 4075 | 4392 | BA_peak_6177 |
| DS114512 | 20 | 420 | BA_peak_6178 |
| DS114514 | 13516 | 14307 | BA_peak_6179 |
| DS114516 | 10985 | 11575 | BA_peak_6180 |
| DS114523 | 8454 | 9021 | BA_peak_6185 |
| DS114524 | 10897 | 11877 | BA_peak_6187 |
| DS114525 | 9660 | 9996 | BA_peak_6189 |
| DS114526 | 5380 | 5783 | BA_peak_6190 |
| DS114527 | 12274 | 13545 | BA_peak_6191 |
| DS114529 | 13464 | 14475 | BA_peak_6192 |
| DS114530 | 6790 | 7578 | BA_peak_6193 |
| DS114531 | 1939 | 2483 | BA_peak_6194 |
| DS114532 | 3824 | 4192 | BA_peak_6195 |
| DS114532 | 7047 | 7650 | BA_peak_6196 |
| DS114533 | 13232 | 13923 | BA_peak_6197 |
| DS114534 | 3139 | 4164 | BA_peak_6198 |
| DS114535 | 786 | 1105 | BA_peak_6200 |
| DS114535 | 10008 | 10345 | BA_peak_6201 |
| DS114536 | 3266 | 3605 | BA_peak_6202 |
| DS114536 | 8124 | 8439 | BA_peak_6203 |
| DS114537 | 41 | 569 | BA_peak_6204 |
| DS114537 | 8779 | 9243 | BA_peak_6205 |
| DS114538 | 236 | 663 | BA_peak_6206 |
| DS114540 | 6615 | 6967 | BA_peak_6207 |
| DS114541 | 10033 | 11015 | BA_peak_6208 |
| DS114542 | 2855 | 3607 | BA_peak_6209 |
| DS114542 | 10461 | 10914 | BA_peak_6210 |
| DS114542 | 11698 | 12526 | BA_peak_6211 |
| DS114543 | 72 | 753 | BA_peak_6212 |
| DS114544 | 2067 | 4557 | BA_peak_6213 |
| DS114545 | 3201 | 3674 | BA_peak_6214 |
| DS114545 | 6133 | 7333 | BA_peak_6215 |
| DS114548 | 10415 | 10840 | BA_peak_6217 |
| DS114549 | 13408 | 13744 | BA_peak_6218 |
| DS114551 | 122 | 494 | BA_peak_6219 |
| DS114552 | 3734 | 4057 | BA_peak_6221 |
| DS114552 | 6685 | 7382 | BA_peak_6223 |
| DS114552 | 12573 | 13735 | BA_peak_6224 |
| DS114553 | 4702 | 6055 | BA_peak_6225 |
| DS114553 | 12300 | 13785 | BA_peak_6226 |
| DS114555 | 172 | 1365 | BA_peak_6227 |
| DS114555 | 3677 | 4376 | BA_peak_6228 |
| DS114558 | 12228 | 13018 | BA_peak_6230 |

| | | | |
|---|---|---|---|
| DS114559 | 70 | 373 | BA_peak_6231 |
| DS114560 | 10411 | 10850 | BA_peak_6233 |
| DS114561 | 243 | 655 | BA_peak_6234 |
| DS114561 | 13257 | 13637 | BA_peak_6235 |
| DS114562 | 7232 | 8196 | BA_peak_6236 |
| DS114563 | 4499 | 4977 | BA_peak_6237 |
| DS114563 | 5419 | 5838 | BA_peak_6238 |
| DS114564 | 1277 | 2188 | BA_peak_6239 |
| DS114564 | 10227 | 10535 | BA_peak_6240 |
| DS114565 | 4703 | 5311 | BA_peak_6242 |
| DS114566 | 5973 | 6265 | BA_peak_6243 |
| DS114566 | 6927 | 7292 | BA_peak_6244 |
| DS114566 | 8864 | 9439 | BA_peak_6245 |
| DS114568 | 99 | 866 | BA_peak_6246 |
| DS114568 | 2938 | 3316 | BA_peak_6247 |
| DS114568 | 9381 | 9714 | BA_peak_6248 |
| DS114571 | 574 | 1598 | BA_peak_6249 |
| DS114571 | 2907 | 4203 | BA_peak_6250 |
| DS114575 | 4925 | 5831 | BA_peak_6251 |
| DS114577 | 9419 | 9742 | BA_peak_6252 |
| DS114577 | 11143 | 11776 | BA_peak_6253 |
| DS114580 | 11761 | 13137 | BA_peak_6255 |
| DS114582 | 8742 | 9250 | BA_peak_6256 |
| DS114583 | 9546 | 10167 | BA_peak_6257 |
| DS114588 | 6281 | 6860 | BA_peak_6258 |
| DS114589 | 11036 | 11576 | BA_peak_6259 |
| DS114590 | 9237 | 9636 | BA_peak_6260 |
| DS114594 | 1565 | 2758 | BA_peak_6261 |
| DS114594 | 6659 | 7189 | BA_peak_6262 |
| DS114595 | 12253 | 12884 | BA_peak_6263 |
| DS114596 | 6550 | 6911 | BA_peak_6264 |
| DS114599 | 4991 | 5449 | BA_peak_6266 |
| DS114600 | 66 | 572 | BA_peak_6267 |
| DS114600 | 12497 | 12841 | BA_peak_6268 |
| DS114601 | 9267 | 9610 | BA_peak_6269 |
| DS114602 | 453 | 834 | BA_peak_6270 |
| DS114602 | 11486 | 12477 | BA_peak_6271 |
| DS114603 | 6795 | 7167 | BA_peak_6272 |
| DS114604 | 1018 | 1326 | BA_peak_6273 |
| DS114604 | 9970 | 11217 | BA_peak_6274 |
| DS114605 | 2446 | 2843 | BA_peak_6275 |
| DS114606 | 5654 | 6330 | BA_peak_6276 |
| DS114606 | 6741 | 7174 | BA_peak_6277 |
| DS114610 | 5127 | 5639 | BA_peak_6279 |
| DS114611 | 302 | 1085 | BA_peak_6280 |
| DS114613 | 2723 | 3099 | BA_peak_6281 |
| DS114614 | 1120 | 1564 | BA_peak_6282 |

| | | | |
|---|---|---|---|
| DS114614 | 2946 | 3339 | BA_peak_6283 |
| DS114620 | 3449 | 4280 | BA_peak_6284 |
| DS114622 | 6242 | 6595 | BA_peak_6286 |
| DS114622 | 9909 | 10810 | BA_peak_6287 |
| DS114623 | 11762 | 12374 | BA_peak_6288 |
| DS114624 | 1642 | 2313 | BA_peak_6289 |
| DS114624 | 2737 | 4345 | BA_peak_6290 |
| DS114624 | 4974 | 5314 | BA_peak_6291 |
| DS114624 | 5706 | 7131 | BA_peak_6292 |
| DS114627 | 6320 | 7325 | BA_peak_6293 |
| DS114627 | 7972 | 8320 | BA_peak_6294 |
| DS114629 | 2923 | 3624 | BA_peak_6295 |
| DS114629 | 11795 | 12209 | BA_peak_6296 |
| DS114630 | 49 | 602 | BA_peak_6297 |
| DS114633 | 6798 | 7405 | BA_peak_6298 |
| DS114633 | 8766 | 9443 | BA_peak_6299 |
| DS114636 | 70 | 1466 | BA_peak_6300 |
| DS114636 | 5780 | 6223 | BA_peak_6301 |
| DS114642 | 5055 | 5956 | BA_peak_6303 |
| DS114642 | 7497 | 8639 | BA_peak_6304 |
| DS114642 | 10961 | 11872 | BA_peak_6305 |
| DS114645 | 10748 | 11340 | BA_peak_6306 |
| DS114647 | 227 | 1048 | BA_peak_6307 |
| DS114648 | 2883 | 3925 | BA_peak_6308 |
| DS114648 | 5585 | 6464 | BA_peak_6309 |
| DS114648 | 7088 | 7677 | BA_peak_6310 |
| DS114650 | 9420 | 10322 | BA_peak_6311 |
| DS114658 | 3869 | 4597 | BA_peak_6313 |
| DS114659 | 2582 | 3222 | BA_peak_6315 |
| DS114660 | 1050 | 2756 | BA_peak_6316 |
| DS114660 | 11034 | 11651 | BA_peak_6317 |
| DS114662 | 46 | 1228 | BA_peak_6318 |
| DS114662 | 1633 | 1921 | BA_peak_6318 |
| DS114664 | 2721 | 3274 | BA_peak_6319 |
| DS114665 | 801 | 1608 | BA_peak_6320 |
| DS114666 | 10141 | 11025 | BA_peak_6321 |
| DS114667 | 9 | 410 | BA_peak_6322 |
| DS114667 | 908 | 1946 | BA_peak_6323 |
| DS114670 | 5616 | 6258 | BA_peak_6324 |
| DS114672 | 10524 | 10900 | BA_peak_6325 |
| DS114676 | 6226 | 6602 | BA_peak_6326 |
| DS114678 | 5197 | 5540 | BA_peak_6328 |
| DS114678 | 10882 | 11332 | BA_peak_6329 |
| DS114679 | 822 | 1398 | BA_peak_6330 |
| DS114681 | 1481 | 2233 | BA_peak_6331 |
| DS114681 | 2608 | 3011 | BA_peak_6332 |
| DS114683 | 4910 | 5517 | BA_peak_6333 |

| | | | |
|---|---|---|---|
| DS114684 | 485 | 781 | BA_peak_6334 |
| DS114684 | 1403 | 1817 | BA_peak_6335 |
| DS114687 | 4614 | 5054 | BA_peak_6336 |
| DS114698 | 2550 | 3175 | BA_peak_6338 |
| DS114698 | 3801 | 4415 | BA_peak_6339 |
| DS114700 | 788 | 1253 | BA_peak_6341 |
| DS114700 | 6203 | 6585 | BA_peak_6342 |
| DS114700 | 10403 | 10751 | BA_peak_6343 |
| DS114701 | 9302 | 9706 | BA_peak_6344 |
| DS114702 | 7186 | 7695 | BA_peak_6345 |
| DS114702 | 8299 | 8741 | BA_peak_6346 |
| DS114702 | 9445 | 9980 | BA_peak_6347 |
| DS114703 | 6023 | 6434 | BA_peak_6348 |
| DS114703 | 6777 | 7180 | BA_peak_6349 |
| DS114703 | 8149 | 8947 | BA_peak_6350 |
| DS114706 | 192 | 860 | BA_peak_6351 |
| DS114706 | 9213 | 10654 | BA_peak_6352 |
| DS114707 | 8346 | 10011 | BA_peak_6353 |
| DS114710 | 7872 | 8306 | BA_peak_6354 |
| DS114715 | 4499 | 4963 | BA_peak_6355 |
| DS114715 | 6453 | 6955 | BA_peak_6356 |
| DS114717 | 8003 | 8629 | BA_peak_6357 |
| DS114717 | 8962 | 9722 | BA_peak_6358 |
| DS114718 | 132 | 558 | BA_peak_6359 |
| DS114718 | 3485 | 3888 | BA_peak_6360 |
| DS114718 | 6624 | 7199 | BA_peak_6361 |
| DS114720 | 3721 | 4034 | BA_peak_6362 |
| DS114726 | 6377 | 6858 | BA_peak_6364 |
| DS114727 | 5710 | 6171 | BA_peak_6365 |
| DS114727 | 9724 | 10313 | BA_peak_6366 |
| DS114728 | 3198 | 3902 | BA_peak_6367 |
| DS114733 | 6522 | 7380 | BA_peak_6368 |
| DS114737 | 85 | 502 | BA_peak_6369 |
| DS114737 | 2130 | 3128 | BA_peak_6370 |
| DS114739 | 523 | 893 | BA_peak_6371 |
| DS114740 | 5056 | 5437 | BA_peak_6372 |
| DS114741 | 6704 | 7029 | BA_peak_6374 |
| DS114741 | 7625 | 8055 | BA_peak_6375 |
| DS114743 | 2204 | 3362 | BA_peak_6376 |
| DS114743 | 8323 | 8644 | BA_peak_6377 |
| DS114744 | 5131 | 5454 | BA_peak_6378 |
| DS114746 | 318 | 1012 | BA_peak_6379 |
| DS114746 | 1341 | 1882 | BA_peak_6380 |
| DS114748 | 65 | 478 | BA_peak_6381 |
| DS114751 | 6765 | 7072 | BA_peak_6382 |
| DS114751 | 9124 | 9712 | BA_peak_6383 |
| DS114752 | 20 | 487 | BA_peak_6384 |

| | | | |
|---|---|---|---|
| DS114755 | 8289 | 8783 | BA_peak_6385 |
| DS114756 | 1222 | 1563 | BA_peak_6387 |
| DS114756 | 3348 | 4028 | BA_peak_6388 |
| DS114759 | 75 | 1148 | BA_peak_6389 |
| DS114759 | 4093 | 4741 | BA_peak_6390 |
| DS114763 | 1094 | 1778 | BA_peak_6391 |
| DS114768 | 5598 | 6439 | BA_peak_6393 |
| DS114768 | 7639 | 8134 | BA_peak_6394 |
| DS114773 | 1063 | 1735 | BA_peak_6396 |
| DS114773 | 8488 | 9515 | BA_peak_6398 |
| DS114775 | 6965 | 7544 | BA_peak_6399 |
| DS114779 | 5599 | 6795 | BA_peak_6400 |
| DS114780 | 1512 | 1913 | BA_peak_6401 |
| DS114783 | 30 | 1000 | BA_peak_6402 |
| DS114783 | 7453 | 8331 | BA_peak_6403 |
| DS114783 | 9149 | 9461 | BA_peak_6404 |
| DS114784 | 6154 | 7560 | BA_peak_6405 |
| DS114786 | 186 | 959 | BA_peak_6406 |
| DS114786 | 1319 | 1624 | BA_peak_6407 |
| DS114786 | 2359 | 3136 | BA_peak_6408 |
| DS114791 | 5576 | 6868 | BA_peak_6409 |
| DS114791 | 7935 | 8288 | BA_peak_6410 |
| DS114791 | 8606 | 9203 | BA_peak_6411 |
| DS114795 | 5698 | 6213 | BA_peak_6412 |
| DS114796 | 795 | 1189 | BA_peak_6413 |
| DS114796 | 8606 | 9136 | BA_peak_6414 |
| DS114798 | 7368 | 7882 | BA_peak_6415 |
| DS114800 | 787 | 2176 | BA_peak_6416 |
| DS114801 | 964 | 1599 | BA_peak_6417 |
| DS114801 | 2426 | 2870 | BA_peak_6418 |
| DS114802 | 64 | 701 | BA_peak_6419 |
| DS114805 | 8529 | 8887 | BA_peak_6420 |
| DS114810 | 1140 | 1956 | BA_peak_6421 |
| DS114810 | 2879 | 3657 | BA_peak_6422 |
| DS114811 | 2112 | 3368 | BA_peak_6423 |
| DS114814 | 1259 | 2031 | BA_peak_6424 |
| DS114814 | 4703 | 5080 | BA_peak_6425 |
| DS114814 | 6044 | 7006 | BA_peak_6426 |
| DS114815 | 4255 | 5808 | BA_peak_6427 |
| DS114816 | 4501 | 5568 | BA_peak_6428 |
| DS114816 | 6862 | 7909 | BA_peak_6429 |
| DS114818 | 2802 | 3798 | BA_peak_6430 |
| DS114818 | 8573 | 8973 | BA_peak_6431 |
| DS114819 | 1869 | 2558 | BA_peak_6432 |
| DS114820 | 8408 | 8741 | BA_peak_6433 |
| DS114822 | 3469 | 4341 | BA_peak_6434 |
| DS114823 | 6129 | 6542 | BA_peak_6435 |

| | | | |
|---|---|---|---|
| DS114825 | 8276 | 8798 | BA_peak_6436 |
| DS114826 | 8135 | 8747 | BA_peak_6437 |
| DS114827 | 5674 | 6336 | BA_peak_6438 |
| DS114828 | 179 | 1419 | BA_peak_6439 |
| DS114832 | 3881 | 5777 | BA_peak_6440 |
| DS114836 | 1337 | 2331 | BA_peak_6441 |
| DS114837 | 6178 | 7354 | BA_peak_6442 |
| DS114841 | 7563 | 8757 | BA_peak_6443 |
| DS114850 | 7135 | 7684 | BA_peak_6447 |
| DS114850 | 7993 | 8660 | BA_peak_6448 |
| DS114854 | 38 | 866 | BA_peak_6449 |
| DS114854 | 1253 | 1607 | BA_peak_6450 |
| DS114854 | 3523 | 3929 | BA_peak_6451 |
| DS114857 | 5272 | 6666 | BA_peak_6452 |
| DS114859 | 4674 | 5847 | BA_peak_6453 |
| DS114863 | 2491 | 3038 | BA_peak_6454 |
| DS114866 | 772 | 3314 | BA_peak_6455 |
| DS114866 | 3972 | 4549 | BA_peak_6456 |
| DS114868 | 38 | 737 | BA_peak_6457 |
| DS114869 | 5166 | 5471 | BA_peak_6458 |
| DS114870 | 6310 | 7257 | BA_peak_6459 |
| DS114871 | 7947 | 8288 | BA_peak_6460 |
| DS114874 | 210 | 674 | BA_peak_6461 |
| DS114874 | 1704 | 2015 | BA_peak_6462 |
| DS114874 | 4089 | 4498 | BA_peak_6463 |
| DS114874 | 5255 | 6250 | BA_peak_6464 |
| DS114874 | 8112 | 8447 | BA_peak_6465 |
| DS114876 | 7762 | 8381 | BA_peak_6466 |
| DS114877 | 5680 | 7287 | BA_peak_6467 |
| DS114877 | 7910 | 8241 | BA_peak_6468 |
| DS114878 | 7799 | 8290 | BA_peak_6469 |
| DS114881 | 2987 | 3867 | BA_peak_6470 |
| DS114883 | 7694 | 8251 | BA_peak_6471 |
| DS114884 | 1340 | 1767 | BA_peak_6472 |
| DS114886 | 6179 | 6508 | BA_peak_6473 |
| DS114890 | 3018 | 3498 | BA_peak_6474 |
| DS114892 | 6722 | 7786 | BA_peak_6475 |
| DS114893 | 901 | 1362 | BA_peak_6476 |
| DS114894 | 285 | 998 | BA_peak_6477 |
| DS114894 | 4268 | 5013 | BA_peak_6478 |
| DS114897 | 2504 | 3406 | BA_peak_6479 |
| DS114897 | 4425 | 4863 | BA_peak_6480 |
| DS114897 | 6212 | 6694 | BA_peak_6481 |
| DS114901 | 3230 | 3702 | BA_peak_6482 |
| DS114908 | 1035 | 2011 | BA_peak_6483 |
| DS114908 | 7162 | 7658 | BA_peak_6484 |
| DS114909 | 5830 | 6551 | BA_peak_6485 |

| | | | |
|---|---|---|---|
| DS114910 | 65 | 513 | BA_peak_6486 |
| DS114910 | 2121 | 2473 | BA_peak_6487 |
| DS114911 | 257 | 870 | BA_peak_6488 |
| DS114913 | 774 | 1157 | BA_peak_6489 |
| DS114913 | 1663 | 2500 | BA_peak_6490 |
| DS114914 | 5947 | 6434 | BA_peak_6491 |
| DS114914 | 7131 | 7534 | BA_peak_6492 |
| DS114917 | 1418 | 2916 | BA_peak_6493 |
| DS114920 | 4051 | 4706 | BA_peak_6494 |
| DS114921 | 7271 | 7696 | BA_peak_6495 |
| DS114922 | 6537 | 7729 | BA_peak_6496 |
| DS114924 | 956 | 1287 | BA_peak_6497 |
| DS114929 | 138 | 776 | BA_peak_6498 |
| DS114930 | 7324 | 7687 | BA_peak_6499 |
| DS114931 | 841 | 1482 | BA_peak_6500 |
| DS114931 | 6279 | 7504 | BA_peak_6501 |
| DS114932 | 6477 | 7136 | BA_peak_6502 |
| DS114933 | 2981 | 3856 | BA_peak_6504 |
| DS114934 | 2227 | 2958 | BA_peak_6505 |
| DS114934 | 6989 | 7489 | BA_peak_6506 |
| DS114936 | 441 | 1534 | BA_peak_6507 |
| DS114936 | 5830 | 6358 | BA_peak_6508 |
| DS114939 | 4714 | 5100 | BA_peak_6509 |
| DS114940 | 4350 | 5751 | BA_peak_6510 |
| DS114940 | 6358 | 6750 | BA_peak_6511 |
| DS114942 | 3719 | 4431 | BA_peak_6512 |
| DS114942 | 5051 | 6414 | BA_peak_6513 |
| DS114946 | 789 | 1233 | BA_peak_6514 |
| DS114946 | 3356 | 3843 | BA_peak_6515 |
| DS114950 | 6698 | 7208 | BA_peak_6516 |
| DS114960 | 13 | 452 | BA_peak_6517 |
| DS114961 | 868 | 1309 | BA_peak_6518 |
| DS114963 | 1017 | 1838 | BA_peak_6519 |
| DS114965 | 2502 | 2878 | BA_peak_6520 |
| DS114966 | 1459 | 2201 | BA_peak_6521 |
| DS114966 | 6918 | 7479 | BA_peak_6522 |
| DS114967 | 2000 | 2480 | BA_peak_6523 |
| DS114967 | 5924 | 7351 | BA_peak_6524 |
| DS114968 | 4825 | 5170 | BA_peak_6525 |
| DS114968 | 6830 | 7385 | BA_peak_6526 |
| DS114970 | 66 | 768 | BA_peak_6527 |
| DS114970 | 6893 | 7411 | BA_peak_6528 |
| DS114973 | 6681 | 7042 | BA_peak_6529 |
| DS114975 | 6805 | 7229 | BA_peak_6530 |
| DS114976 | 44 | 440 | BA_peak_6531 |
| DS114977 | 4160 | 4560 | BA_peak_6532 |
| DS114980 | 519 | 1000 | BA_peak_6533 |

| | | | |
|---|---|---|---|
| DS114984 | 4746 | 5300 | BA_peak_6534 |
| DS114984 | 5771 | 6070 | BA_peak_6535 |
| DS114992 | 149 | 858 | BA_peak_6536 |
| DS114992 | 2174 | 2768 | BA_peak_6537 |
| DS114995 | 6117 | 6924 | BA_peak_6538 |
| DS114996 | 1124 | 2360 | BA_peak_6539 |
| DS115004 | 98 | 403 | BA_peak_6541 |
| DS115011 | 2108 | 2434 | BA_peak_6542 |
| DS115011 | 5375 | 6374 | BA_peak_6543 |
| DS115013 | 1531 | 2155 | BA_peak_6544 |
| DS115015 | 141 | 501 | BA_peak_6545 |
| DS115026 | 5887 | 6218 | BA_peak_6546 |
| DS115031 | 34 | 2577 | BA_peak_6547 |
| DS115033 | 5888 | 6613 | BA_peak_6549 |
| DS115034 | 5277 | 5832 | BA_peak_6550 |
| DS115037 | 5021 | 5485 | BA_peak_6551 |
| DS115041 | 1044 | 1595 | BA_peak_6552 |
| DS115042 | 5413 | 6253 | BA_peak_6553 |
| DS115049 | 799 | 1629 | BA_peak_6557 |
| DS115049 | 3654 | 4069 | BA_peak_6558 |
| DS115055 | 3514 | 3857 | BA_peak_6559 |
| DS115056 | 3934 | 4382 | BA_peak_6560 |
| DS115058 | 1042 | 1764 | BA_peak_6561 |
| DS115060 | 5776 | 6169 | BA_peak_6562 |
| DS115061 | 247 | 1659 | BA_peak_6563 |
| DS115069 | 2884 | 3336 | BA_peak_6564 |
| DS115071 | 908 | 1217 | BA_peak_6565 |
| DS115074 | 3375 | 4442 | BA_peak_6566 |
| DS115087 | 305 | 869 | BA_peak_6567 |
| DS115089 | 2478 | 4268 | BA_peak_6568 |
| DS115090 | 102 | 442 | BA_peak_6569 |
| DS115091 | 4384 | 5473 | BA_peak_6571 |
| DS115093 | 2353 | 3246 | BA_peak_6572 |
| DS115094 | 343 | 879 | BA_peak_6573 |
| DS115094 | 5715 | 6231 | BA_peak_6574 |
| DS115096 | 3497 | 4237 | BA_peak_6575 |
| DS115102 | 775 | 2528 | BA_peak_6578 |
| DS115104 | 69 | 669 | BA_peak_6579 |
| DS115104 | 2461 | 2900 | BA_peak_6580 |
| DS115104 | 3799 | 4347 | BA_peak_6581 |
| DS115110 | 55 | 1304 | BA_peak_6583 |
| DS115112 | 4684 | 5346 | BA_peak_6585 |
| DS115115 | 5740 | 6221 | BA_peak_6586 |
| DS115121 | 5515 | 6173 | BA_peak_6587 |
| DS115132 | 128 | 494 | BA_peak_6589 |
| DS115132 | 5599 | 6078 | BA_peak_6590 |
| DS115134 | 5588 | 6181 | BA_peak_6591 |

| | | | |
|---|---|---|---|
| DS115136 | 40 | 837 | BA_peak_6592 |
| DS115137 | 4763 | 6187 | BA_peak_6593 |
| DS115138 | 87 | 1317 | BA_peak_6594 |
| DS115141 | 813 | 1134 | BA_peak_6595 |
| DS115141 | 1441 | 1954 | BA_peak_6596 |
| DS115145 | 48 | 1380 | BA_peak_6597 |
| DS115148 | 96 | 384 | BA_peak_6598 |
| DS115148 | 705 | 1471 | BA_peak_6599 |
| DS115149 | 939 | 2087 | BA_peak_6600 |
| DS115155 | 206 | 1289 | BA_peak_6602 |
| DS115159 | 94 | 741 | BA_peak_6604 |
| DS115159 | 2109 | 3228 | BA_peak_6605 |
| DS115163 | 1712 | 2223 | BA_peak_6606 |
| DS115164 | 1798 | 3154 | BA_peak_6607 |
| DS115166 | 342 | 873 | BA_peak_6608 |
| DS115169 | 1620 | 1938 | BA_peak_6609 |
| DS115171 | 2701 | 3174 | BA_peak_6610 |
| DS115171 | 3905 | 4319 | BA_peak_6611 |
| DS115174 | 1270 | 1787 | BA_peak_6612 |
| DS115174 | 2450 | 3188 | BA_peak_6613 |
| DS115174 | 5496 | 5876 | BA_peak_6614 |
| DS115179 | 5305 | 5947 | BA_peak_6615 |
| DS115180 | 5237 | 5739 | BA_peak_6616 |
| DS115182 | 4074 | 4458 | BA_peak_6617 |
| DS115186 | 3740 | 4101 | BA_peak_6618 |
| DS115190 | 2984 | 3512 | BA_peak_6619 |
| DS115194 | 3295 | 4000 | BA_peak_6620 |
| DS115196 | 569 | 1181 | BA_peak_6621 |
| DS115200 | 4362 | 5033 | BA_peak_6622 |
| DS115201 | 5058 | 5791 | BA_peak_6623 |
| DS115209 | 1990 | 2394 | BA_peak_6624 |
| DS115210 | 5285 | 5778 | BA_peak_6625 |
| DS115214 | 431 | 949 | BA_peak_6626 |
| DS115221 | 1040 | 1511 | BA_peak_6627 |
| DS115224 | 4904 | 5441 | BA_peak_6628 |
| DS115225 | 3293 | 3659 | BA_peak_6629 |
| DS115232 | 35 | 1867 | BA_peak_6630 |
| DS115234 | 1491 | 1821 | BA_peak_6631 |
| DS115237 | 4897 | 5508 | BA_peak_6632 |
| DS115243 | 4132 | 4549 | BA_peak_6633 |
| DS115251 | 18 | 464 | BA_peak_6634 |
| DS115251 | 2338 | 2766 | BA_peak_6635 |
| DS115253 | 5151 | 5506 | BA_peak_6636 |
| DS115254 | 1107 | 1464 | BA_peak_6637 |
| DS115254 | 1830 | 2137 | BA_peak_6638 |
| DS115257 | 392 | 801 | BA_peak_6639 |
| DS115257 | 1806 | 2434 | BA_peak_6640 |

| | | | |
|---|---|---|---|
| DS115260 | 4325 | 4793 | BA_peak_6641 |
| DS115261 | 2038 | 2645 | BA_peak_6642 |
| DS115261 | 3148 | 4065 | BA_peak_6643 |
| DS115265 | 177 | 772 | BA_peak_6644 |
| DS115266 | 173 | 640 | BA_peak_6645 |
| DS115273 | 123 | 1042 | BA_peak_6646 |
| DS115278 | 3978 | 4366 | BA_peak_6647 |
| DS115278 | 4759 | 5252 | BA_peak_6648 |
| DS115281 | 2107 | 2874 | BA_peak_6649 |
| DS115283 | 235 | 903 | BA_peak_6650 |
| DS115284 | 54 | 1372 | BA_peak_6651 |
| DS115286 | 4574 | 5162 | BA_peak_6652 |
| DS115292 | 3382 | 4413 | BA_peak_6653 |
| DS115294 | 213 | 753 | BA_peak_6654 |
| DS115294 | 3585 | 4046 | BA_peak_6655 |
| DS115297 | 286 | 1377 | BA_peak_6656 |
| DS115315 | 4836 | 5186 | BA_peak_6657 |
| DS115317 | 1876 | 2640 | BA_peak_6658 |
| DS115317 | 3440 | 3789 | BA_peak_6659 |
| DS115317 | 4865 | 5191 | BA_peak_6660 |
| DS115334 | 2150 | 2731 | BA_peak_6661 |
| DS115336 | 351 | 1001 | BA_peak_6662 |
| DS115336 | 2421 | 2791 | BA_peak_6663 |
| DS115339 | 2898 | 4271 | BA_peak_6664 |
| DS115340 | 105 | 965 | BA_peak_6665 |
| DS115342 | 3105 | 3756 | BA_peak_6666 |
| DS115354 | 2871 | 3399 | BA_peak_6668 |
| DS115356 | 3599 | 4307 | BA_peak_6669 |
| DS115359 | 3652 | 4263 | BA_peak_6670 |
| DS115363 | 3409 | 3949 | BA_peak_6671 |
| DS115364 | 4673 | 5076 | BA_peak_6672 |
| DS115366 | 593 | 2317 | BA_peak_6673 |
| DS115366 | 3940 | 4255 | BA_peak_6674 |
| DS115368 | 1487 | 1852 | BA_peak_6675 |
| DS115369 | 6 | 541 | BA_peak_6676 |
| DS115369 | 864 | 2495 | BA_peak_6677 |
| DS115370 | 79 | 398 | BA_peak_6678 |
| DS115379 | 2939 | 3408 | BA_peak_6679 |
| DS115385 | 269 | 702 | BA_peak_6680 |
| DS115385 | 1173 | 2037 | BA_peak_6681 |
| DS115385 | 4108 | 4463 | BA_peak_6683 |
| DS115388 | 4415 | 4930 | BA_peak_6684 |
| DS115392 | 887 | 1399 | BA_peak_6685 |
| DS115399 | 2727 | 3059 | BA_peak_6686 |
| DS115401 | 3086 | 4953 | BA_peak_6687 |
| DS115403 | 86 | 592 | BA_peak_6688 |
| DS115405 | 3708 | 4837 | BA_peak_6690 |

| | | | |
|---|---|---|---|
| DS115407 | 4461 | 4939 | BA_peak_6691 |
| DS115408 | 332 | 686 | BA_peak_6692 |
| DS115408 | 2522 | 3155 | BA_peak_6693 |
| DS115408 | 3548 | 4788 | BA_peak_6694 |
| DS115411 | 1922 | 2881 | BA_peak_6695 |
| DS115411 | 4332 | 4751 | BA_peak_6696 |
| DS115416 | 4121 | 4912 | BA_peak_6697 |
| DS115433 | 4132 | 4619 | BA_peak_6698 |
| DS115438 | 226 | 577 | BA_peak_6700 |
| DS115439 | 296 | 615 | BA_peak_6701 |
| DS115449 | 921 | 1461 | BA_peak_6702 |
| DS115454 | 2682 | 3029 | BA_peak_6703 |
| DS115464 | 846 | 1370 | BA_peak_6704 |
| DS115465 | 3616 | 4038 | BA_peak_6705 |
| DS115466 | 1592 | 1918 | BA_peak_6706 |
| DS115470 | 302 | 628 | BA_peak_6707 |
| DS115486 | 1809 | 2155 | BA_peak_6709 |
| DS115506 | 82 | 643 | BA_peak_6710 |
| DS115517 | 3264 | 3893 | BA_peak_6712 |
| DS115520 | 2643 | 3048 | BA_peak_6713 |
| DS115523 | 992 | 1920 | BA_peak_6714 |
| DS115531 | 3810 | 4142 | BA_peak_6715 |
| DS115534 | 2167 | 2695 | BA_peak_6716 |
| DS115538 | 125 | 551 | BA_peak_6717 |
| DS115539 | 2907 | 3583 | BA_peak_6718 |
| DS115542 | 115 | 1363 | BA_peak_6719 |
| DS115542 | 4053 | 4549 | BA_peak_6720 |
| DS115551 | 200 | 675 | BA_peak_6721 |
| DS115551 | 3781 | 4127 | BA_peak_6722 |
| DS115554 | 3812 | 4450 | BA_peak_6723 |
| DS115555 | 1718 | 2049 | BA_peak_6724 |
| DS115563 | 142 | 480 | BA_peak_6725 |
| DS115563 | 1472 | 2695 | BA_peak_6726 |
| DS115563 | 3029 | 3441 | BA_peak_6727 |
| DS115563 | 3940 | 4233 | BA_peak_6728 |
| DS115572 | 47 | 548 | BA_peak_6730 |
| DS115572 | 1108 | 1654 | BA_peak_6731 |
| DS115575 | 1751 | 2401 | BA_peak_6732 |
| DS115576 | 1139 | 1739 | BA_peak_6733 |
| DS115585 | 723 | 1173 | BA_peak_6734 |
| DS115594 | 1352 | 1835 | BA_peak_6735 |
| DS115599 | 864 | 1475 | BA_peak_6736 |
| DS115600 | 54 | 1019 | BA_peak_6737 |
| DS115607 | 2957 | 3685 | BA_peak_6738 |
| DS115615 | 128 | 813 | BA_peak_6739 |
| DS115620 | 488 | 1666 | BA_peak_6740 |
| DS115620 | 2865 | 3432 | BA_peak_6741 |

| | | | |
|---|---|---|---|
| DS115623 | 1921 | 4341 | BA_peak_6742 |
| DS115624 | 3310 | 3623 | BA_peak_6743 |
| DS115625 | 1819 | 2369 | BA_peak_6744 |
| DS115629 | 2897 | 3564 | BA_peak_6745 |
| DS115635 | 545 | 948 | BA_peak_6746 |
| DS115639 | 1226 | 1575 | BA_peak_6747 |
| DS115640 | 2658 | 3055 | BA_peak_6748 |
| DS115642 | 3039 | 4266 | BA_peak_6750 |
| DS115644 | 104 | 535 | BA_peak_6751 |
| DS115655 | 293 | 638 | BA_peak_6752 |
| DS115657 | 1147 | 2354 | BA_peak_6753 |
| DS115662 | 211 | 751 | BA_peak_6754 |
| DS115664 | 5 | 3107 | BA_peak_6756 |
| DS115669 | 1178 | 1608 | BA_peak_6757 |
| DS115681 | 1498 | 1865 | BA_peak_6758 |
| DS115681 | 3686 | 4138 | BA_peak_6759 |
| DS115690 | 562 | 1471 | BA_peak_6760 |
| DS115690 | 3600 | 3931 | BA_peak_6761 |
| DS115692 | 196 | 718 | BA_peak_6762 |
| DS115706 | 41 | 555 | BA_peak_6763 |
| DS115712 | 3444 | 4129 | BA_peak_6766 |
| DS115726 | 3669 | 4012 | BA_peak_6767 |
| DS115732 | 849 | 1554 | BA_peak_6768 |
| DS115738 | 1853 | 2688 | BA_peak_6769 |
| DS115739 | 2815 | 3271 | BA_peak_6770 |
| DS115740 | 1522 | 2055 | BA_peak_6771 |
| DS115742 | 3426 | 4089 | BA_peak_6772 |
| DS115743 | 3329 | 3860 | BA_peak_6773 |
| DS115762 | 290 | 703 | BA_peak_6774 |
| DS115764 | 2155 | 3230 | BA_peak_6775 |
| DS115770 | 2674 | 3041 | BA_peak_6776 |
| DS115778 | 1172 | 1988 | BA_peak_6777 |
| DS115778 | 2638 | 3789 | BA_peak_6778 |
| DS115780 | 3314 | 3874 | BA_peak_6779 |
| DS115789 | 1067 | 1441 | BA_peak_6780 |
| DS115790 | 179 | 551 | BA_peak_6781 |
| DS115794 | 2972 | 3399 | BA_peak_6782 |
| DS115795 | 268 | 702 | BA_peak_6783 |
| DS115806 | 194 | 845 | BA_peak_6784 |
| DS115808 | 3102 | 3416 | BA_peak_6785 |
| DS115816 | 1317 | 2263 | BA_peak_6786 |
| DS115819 | 2048 | 3931 | BA_peak_6787 |
| DS115833 | 2933 | 3580 | BA_peak_6788 |
| DS115835 | 344 | 692 | BA_peak_6789 |
| DS115837 | 1721 | 2106 | BA_peak_6790 |
| DS115840 | 3245 | 3871 | BA_peak_6792 |
| DS115847 | 361 | 735 | BA_peak_6793 |

| | | | |
|---|---|---|---|
| DS115847 | 2985 | 3363 | BA_peak_6794 |
| DS115849 | 3195 | 3611 | BA_peak_6795 |
| DS115856 | 728 | 1502 | BA_peak_6796 |
| DS115864 | 2912 | 3847 | BA_peak_6798 |
| DS115865 | 2605 | 3765 | BA_peak_6799 |
| DS115875 | 114 | 610 | BA_peak_6801 |
| DS115877 | 1659 | 2945 | BA_peak_6802 |
| DS115877 | 3429 | 3827 | BA_peak_6803 |
| DS115879 | 100 | 427 | BA_peak_6804 |
| DS115887 | 151 | 684 | BA_peak_6805 |
| DS115894 | 116 | 718 | BA_peak_6806 |
| DS115900 | 2058 | 3544 | BA_peak_6807 |
| DS115907 | 265 | 861 | BA_peak_6808 |
| DS115916 | 3355 | 3697 | BA_peak_6809 |
| DS115929 | 34 | 1329 | BA_peak_6811 |
| DS115933 | 345 | 710 | BA_peak_6812 |
| DS115933 | 1472 | 2233 | BA_peak_6813 |
| DS115933 | 3160 | 3644 | BA_peak_6814 |
| DS115936 | 2627 | 3249 | BA_peak_6815 |
| DS115941 | 386 | 2819 | BA_peak_6816 |
| DS115943 | 669 | 1420 | BA_peak_6817 |
| DS115948 | 238 | 1791 | BA_peak_6818 |
| DS115949 | 184 | 640 | BA_peak_6819 |
| DS115949 | 1233 | 1578 | BA_peak_6820 |
| DS115954 | 612 | 1386 | BA_peak_6821 |
| DS115954 | 1949 | 2680 | BA_peak_6822 |
| DS115956 | 256 | 814 | BA_peak_6823 |
| DS115956 | 1440 | 2164 | BA_peak_6824 |
| DS115959 | 431 | 912 | BA_peak_6825 |
| DS115959 | 2277 | 2781 | BA_peak_6826 |
| DS115970 | 357 | 990 | BA_peak_6827 |
| DS115973 | 1906 | 2422 | BA_peak_6828 |
| DS115978 | 1111 | 2790 | BA_peak_6830 |
| DS115988 | 58 | 635 | BA_peak_6831 |
| DS115988 | 2943 | 3569 | BA_peak_6832 |
| DS116011 | 206 | 1118 | BA_peak_6833 |
| DS116019 | 401 | 1662 | BA_peak_6834 |
| DS116026 | 281 | 833 | BA_peak_6835 |
| DS116027 | 97 | 462 | BA_peak_6836 |
| DS116028 | 53 | 539 | BA_peak_6838 |
| DS116028 | 2476 | 2815 | BA_peak_6839 |
| DS116031 | 132 | 778 | BA_peak_6841 |
| DS116038 | 214 | 792 | BA_peak_6842 |
| DS116041 | 905 | 1293 | BA_peak_6845 |
| DS116041 | 2889 | 3521 | BA_peak_6846 |
| DS116055 | 95 | 609 | BA_peak_6847 |
| DS116069 | 1699 | 2053 | BA_peak_6848 |

| | | | |
|---|---|---|---|
| DS116078 | 2243 | 2622 | BA_peak_6849 |
| DS116080 | 43 | 400 | BA_peak_6850 |
| DS116080 | 2225 | 3419 | BA_peak_6851 |
| DS116094 | 1530 | 2429 | BA_peak_6852 |
| DS116112 | 2358 | 3332 | BA_peak_6853 |
| DS116117 | 3111 | 3495 | BA_peak_6854 |
| DS116122 | 1722 | 2167 | BA_peak_6855 |
| DS116125 | 89 | 633 | BA_peak_6856 |
| DS116125 | 2944 | 3402 | BA_peak_6857 |
| DS116127 | 2466 | 2716 | BA_peak_6858 |
| DS116132 | 1720 | 2758 | BA_peak_6859 |
| DS116133 | 48 | 471 | BA_peak_6860 |
| DS116134 | 1986 | 3375 | BA_peak_6861 |
| DS116143 | 2749 | 3441 | BA_peak_6862 |
| DS116147 | 383 | 744 | BA_peak_6863 |
| DS116147 | 1365 | 2026 | BA_peak_6864 |
| DS116147 | 2794 | 3409 | BA_peak_6865 |
| DS116159 | 125 | 490 | BA_peak_6866 |
| DS116159 | 1562 | 2183 | BA_peak_6867 |
| DS116175 | 877 | 1353 | BA_peak_6868 |
| DS116177 | 295 | 654 | BA_peak_6869 |
| DS116177 | 1370 | 2986 | BA_peak_6870 |
| DS116181 | 2510 | 3255 | BA_peak_6871 |
| DS116195 | 2166 | 3043 | BA_peak_6872 |
| DS116203 | 902 | 1832 | BA_peak_6873 |
| DS116210 | 1841 | 2408 | BA_peak_6874 |
| DS116221 | 1450 | 1799 | BA_peak_6875 |
| DS116221 | 2408 | 3326 | BA_peak_6876 |
| DS116222 | 136 | 851 | BA_peak_6877 |
| DS116229 | 2799 | 3304 | BA_peak_6879 |
| DS116233 | 2657 | 3274 | BA_peak_6880 |
| DS116241 | 2674 | 3172 | BA_peak_6881 |
| DS116244 | 61 | 1474 | BA_peak_6882 |
| DS116248 | 861 | 1347 | BA_peak_6883 |
| DS116254 | 210 | 612 | BA_peak_6884 |
| DS116257 | 77 | 898 | BA_peak_6886 |
| DS116258 | 1133 | 2026 | BA_peak_6887 |
| DS116267 | 2839 | 3170 | BA_peak_6888 |
| DS116269 | 977 | 1471 | BA_peak_6889 |
| DS116273 | 432 | 757 | BA_peak_6890 |
| DS116276 | 1960 | 2324 | BA_peak_6891 |
| DS116277 | 73 | 1684 | BA_peak_6892 |
| DS116283 | 1033 | 2501 | BA_peak_6893 |
| DS116286 | 1491 | 2036 | BA_peak_6894 |
| DS116288 | 351 | 951 | BA_peak_6895 |
| DS116293 | 2234 | 2597 | BA_peak_6896 |
| DS116295 | 2534 | 3100 | BA_peak_6897 |

| | | | |
|---|---|---|---|
| DS116300 | 2373 | 3161 | BA_peak_6898 |
| DS116302 | 2686 | 3012 | BA_peak_6899 |
| DS116310 | 791 | 1969 | BA_peak_6900 |
| DS116311 | 2395 | 2871 | BA_peak_6901 |
| DS116313 | 1098 | 1416 | BA_peak_6902 |
| DS116319 | 412 | 863 | BA_peak_6903 |
| DS116319 | 2101 | 2438 | BA_peak_6904 |
| DS116321 | 1252 | 2291 | BA_peak_6905 |
| DS116322 | 86 | 864 | BA_peak_6906 |
| DS116322 | 2007 | 2621 | BA_peak_6907 |
| DS116335 | 352 | 985 | BA_peak_6909 |
| DS116335 | 2467 | 3173 | BA_peak_6910 |
| DS116337 | 1325 | 2487 | BA_peak_6911 |
| DS116350 | 1098 | 1618 | BA_peak_6913 |
| DS116368 | 640 | 1097 | BA_peak_6915 |
| DS116369 | 408 | 826 | BA_peak_6916 |
| DS116369 | 2683 | 3161 | BA_peak_6917 |
| DS116371 | 2711 | 3028 | BA_peak_6918 |
| DS116375 | 727 | 1443 | BA_peak_6919 |
| DS116385 | 1794 | 2178 | BA_peak_6920 |
| DS116386 | 544 | 890 | BA_peak_6921 |
| DS116386 | 1919 | 2650 | BA_peak_6922 |
| DS116399 | 2597 | 3116 | BA_peak_6923 |
| DS116402 | 67 | 601 | BA_peak_6924 |
| DS116406 | 1671 | 2095 | BA_peak_6925 |
| DS116407 | 857 | 1241 | BA_peak_6926 |
| DS116409 | 599 | 1014 | BA_peak_6927 |
| DS116414 | 28 | 546 | BA_peak_6928 |
| DS116416 | 182 | 500 | BA_peak_6929 |
| DS116419 | 1087 | 1562 | BA_peak_6930 |
| DS116425 | 1590 | 2430 | BA_peak_6931 |
| DS116433 | 1450 | 2108 | BA_peak_6932 |
| DS116443 | 211 | 1117 | BA_peak_6933 |
| DS116447 | 2415 | 2998 | BA_peak_6935 |
| DS116448 | 12 | 342 | BA_peak_6936 |
| DS116448 | 995 | 1464 | BA_peak_6937 |
| DS116450 | 2501 | 3083 | BA_peak_6938 |
| DS116453 | 239 | 734 | BA_peak_6939 |
| DS116457 | 1334 | 1841 | BA_peak_6940 |
| DS116467 | 2426 | 2977 | BA_peak_6941 |
| DS116470 | 164 | 850 | BA_peak_6942 |
| DS116470 | 1586 | 2402 | BA_peak_6943 |
| DS116493 | 102 | 554 | BA_peak_6945 |
| DS116497 | 1759 | 2552 | BA_peak_6948 |
| DS116499 | 2315 | 2976 | BA_peak_6949 |
| DS116503 | 2033 | 2359 | BA_peak_6950 |
| DS116503 | 2693 | 2999 | BA_peak_6951 |

| | | | |
|---|---|---|---|
| DS116504 | 888 | 1496 | BA_peak_6952 |
| DS116513 | 2636 | 3026 | BA_peak_6953 |
| DS116525 | 2447 | 2964 | BA_peak_6956 |
| DS116528 | 1640 | 2419 | BA_peak_6957 |
| DS116530 | 2649 | 3014 | BA_peak_6958 |
| DS116534 | 2273 | 2707 | BA_peak_6959 |
| DS116539 | 1792 | 2328 | BA_peak_6960 |
| DS116540 | 51 | 696 | BA_peak_6961 |
| DS116540 | 1066 | 1790 | BA_peak_6962 |
| DS116541 | 1652 | 2083 | BA_peak_6963 |
| DS116554 | 2546 | 2891 | BA_peak_6965 |
| DS116557 | 2294 | 2929 | BA_peak_6966 |
| DS116568 | 2356 | 2921 | BA_peak_6967 |
| DS116569 | 2496 | 2811 | BA_peak_6968 |
| DS116576 | 1195 | 2006 | BA_peak_6969 |
| DS116577 | 682 | 1051 | BA_peak_6970 |
| DS116581 | 2207 | 2740 | BA_peak_6971 |
| DS116588 | 91 | 2606 | BA_peak_6972 |
| DS116591 | 376 | 1205 | BA_peak_6973 |
| DS116594 | 2301 | 2827 | BA_peak_6974 |
| DS116605 | 645 | 1578 | BA_peak_6975 |
| DS116605 | 2159 | 2901 | BA_peak_6976 |
| DS116613 | 148 | 576 | BA_peak_6977 |
| DS116620 | 141 | 620 | BA_peak_6978 |
| DS116621 | 853 | 1235 | BA_peak_6979 |
| DS116633 | 416 | 1192 | BA_peak_6980 |
| DS116635 | 743 | 1508 | BA_peak_6981 |
| DS116642 | 2467 | 2948 | BA_peak_6982 |
| DS116646 | 994 | 1485 | BA_peak_6983 |
| DS116664 | 2315 | 2911 | BA_peak_6984 |
| DS116669 | 807 | 1188 | BA_peak_6985 |
| DS116674 | 2167 | 2850 | BA_peak_6986 |
| DS116679 | 184 | 488 | BA_peak_6987 |
| DS116682 | 112 | 1211 | BA_peak_6988 |
| DS116682 | 2153 | 2761 | BA_peak_6989 |
| DS116686 | 1171 | 1617 | BA_peak_6990 |
| DS116693 | 61 | 568 | BA_peak_6991 |
| DS116693 | 1209 | 1924 | BA_peak_6992 |
| DS116697 | 444 | 1096 | BA_peak_6993 |
| DS116700 | 118 | 629 | BA_peak_6994 |
| DS116700 | 1127 | 2232 | BA_peak_6995 |
| DS116708 | 817 | 1292 | BA_peak_6996 |
| DS116716 | 1022 | 1811 | BA_peak_6997 |
| DS116716 | 2332 | 2804 | BA_peak_6998 |
| DS116720 | 61 | 639 | BA_peak_6999 |
| DS116722 | 2025 | 2714 | BA_peak_7000 |
| DS116724 | 1479 | 2615 | BA_peak_7001 |

| | | | |
|---|---|---|---|
| DS116725 | 1428 | 1796 | BA_peak_7003 |
| DS116731 | 1185 | 1536 | BA_peak_7005 |
| DS116738 | 129 | 1322 | BA_peak_7006 |
| DS116739 | 2007 | 2758 | BA_peak_7007 |
| DS116747 | 521 | 1266 | BA_peak_7008 |
| DS116748 | 100 | 419 | BA_peak_7009 |
| DS116749 | 1770 | 2800 | BA_peak_7010 |
| DS116753 | 1444 | 1810 | BA_peak_7011 |
| DS116757 | 218 | 937 | BA_peak_7012 |
| DS116757 | 2357 | 2688 | BA_peak_7013 |
| DS116761 | 285 | 735 | BA_peak_7014 |
| DS116770 | 443 | 1096 | BA_peak_7015 |
| DS116779 | 216 | 821 | BA_peak_7017 |
| DS116780 | 54 | 1635 | BA_peak_7018 |
| DS116802 | 2132 | 2713 | BA_peak_7020 |
| DS116814 | 1255 | 1720 | BA_peak_7021 |
| DS116817 | 145 | 2683 | BA_peak_7022 |
| DS116823 | 45 | 1459 | BA_peak_7023 |
| DS116843 | 177 | 487 | BA_peak_7025 |
| DS116867 | 846 | 1672 | BA_peak_7026 |
| DS116868 | 2218 | 2710 | BA_peak_7027 |
| DS116872 | 900 | 1657 | BA_peak_7028 |
| DS116876 | 1048 | 2661 | BA_peak_7029 |
| DS116877 | 1736 | 2680 | BA_peak_7030 |
| DS116891 | 1943 | 2613 | BA_peak_7031 |
| DS116896 | 22 | 398 | BA_peak_7032 |
| DS116899 | 1716 | 2366 | BA_peak_7033 |
| DS116917 | 485 | 1057 | BA_peak_7034 |
| DS116938 | 1012 | 1553 | BA_peak_7036 |
| DS116942 | 37 | 446 | BA_peak_7037 |
| DS116948 | 2035 | 2506 | BA_peak_7038 |
| DS116955 | 2128 | 2622 | BA_peak_7039 |
| DS116956 | 58 | 504 | BA_peak_7040 |
| DS116956 | 1106 | 1822 | BA_peak_7041 |
| DS116959 | 134 | 573 | BA_peak_7042 |
| DS116974 | 1370 | 1852 | BA_peak_7043 |
| DS116986 | 1890 | 2510 | BA_peak_7044 |
| DS116988 | 1845 | 2328 | BA_peak_7045 |
| DS117014 | 1848 | 2242 | BA_peak_7046 |
| DS117016 | 139 | 648 | BA_peak_7047 |
| DS117016 | 2088 | 2549 | BA_peak_7048 |
| DS117017 | 784 | 1324 | BA_peak_7049 |
| DS117022 | 846 | 1493 | BA_peak_7050 |
| DS117029 | 138 | 799 | BA_peak_7051 |
| DS117034 | 610 | 1007 | BA_peak_7052 |
| DS117052 | 205 | 512 | BA_peak_7053 |
| DS117063 | 1839 | 2337 | BA_peak_7054 |

| | | | |
|---|---|---|---|
| DS117069 | 619 | 950 | BA_peak_7055 |
| DS117071 | 113 | 849 | BA_peak_7056 |
| DS117076 | 1993 | 2485 | BA_peak_7057 |
| DS117077 | 47 | 634 | BA_peak_7058 |
| DS117081 | 725 | 1228 | BA_peak_7059 |
| DS117081 | 1672 | 2048 | BA_peak_7060 |
| DS117088 | 63 | 553 | BA_peak_7061 |
| DS117094 | 188 | 2121 | BA_peak_7062 |
| DS117108 | 1940 | 2480 | BA_peak_7063 |
| DS117120 | 749 | 1089 | BA_peak_7065 |
| DS117125 | 1627 | 2238 | BA_peak_7066 |
| DS117132 | 838 | 1625 | BA_peak_7067 |
| DS117145 | 301 | 833 | BA_peak_7068 |
| DS117155 | 269 | 835 | BA_peak_7069 |
| DS117155 | 1467 | 2405 | BA_peak_7070 |
| DS117165 | 391 | 1179 | BA_peak_7071 |
| DS117177 | 290 | 660 | BA_peak_7072 |
| DS117177 | 2149 | 2520 | BA_peak_7073 |
| DS117181 | 713 | 1046 | BA_peak_7074 |
| DS117186 | 394 | 2326 | BA_peak_7075 |
| DS117187 | 507 | 2272 | BA_peak_7076 |
| DS117195 | 337 | 2187 | BA_peak_7077 |
| DS117207 | 107 | 1050 | BA_peak_7078 |
| DS117215 | 265 | 1236 | BA_peak_7079 |
| DS117220 | 112 | 1413 | BA_peak_7080 |
| DS117221 | 1728 | 2174 | BA_peak_7081 |
| DS117229 | 1722 | 2381 | BA_peak_7083 |
| DS117231 | 142 | 462 | BA_peak_7084 |
| DS117231 | 851 | 1395 | BA_peak_7085 |
| DS117232 | 776 | 1158 | BA_peak_7086 |
| DS117232 | 1496 | 1927 | BA_peak_7087 |
| DS117241 | 928 | 1419 | BA_peak_7088 |
| DS117251 | 299 | 1451 | BA_peak_7089 |
| DS117251 | 1921 | 2430 | BA_peak_7090 |
| DS117252 | 378 | 869 | BA_peak_7091 |
| DS117252 | 1272 | 1779 | BA_peak_7092 |
| DS117256 | 25 | 532 | BA_peak_7093 |
| DS117271 | 129 | 517 | BA_peak_7095 |
| DS117281 | 44 | 636 | BA_peak_7096 |
| DS117288 | 1901 | 2436 | BA_peak_7097 |
| DS117289 | 197 | 604 | BA_peak_7098 |
| DS117302 | 195 | 688 | BA_peak_7099 |
| DS117305 | 922 | 1543 | BA_peak_7100 |
| DS117306 | 125 | 573 | BA_peak_7101 |
| DS117313 | 38 | 714 | BA_peak_7102 |
| DS117316 | 174 | 539 | BA_peak_7103 |
| DS117316 | 1039 | 1368 | BA_peak_7104 |

| | | | |
|---|---|---|---|
| DS117316 | 1740 | 2109 | BA_peak_7105 |
| DS117317 | 1881 | 2350 | BA_peak_7106 |
| DS117320 | 990 | 1546 | BA_peak_7107 |
| DS117325 | 72 | 1283 | BA_peak_7108 |
| DS117355 | 1754 | 2399 | BA_peak_7109 |
| DS117357 | 108 | 478 | BA_peak_7111 |
| DS117362 | 195 | 746 | BA_peak_7112 |
| DS117364 | 760 | 1521 | BA_peak_7113 |
| DS117365 | 186 | 551 | BA_peak_7114 |
| DS117366 | 1628 | 2418 | BA_peak_7115 |
| DS117383 | 129 | 758 | BA_peak_7116 |
| DS117387 | 821 | 1227 | BA_peak_7117 |
| DS117410 | 32 | 477 | BA_peak_7119 |
| DS117414 | 135 | 623 | BA_peak_7121 |
| DS117418 | 1057 | 2262 | BA_peak_7122 |
| DS117436 | 711 | 1178 | BA_peak_7123 |
| DS117446 | 113 | 1058 | BA_peak_7124 |
| DS117452 | 889 | 1622 | BA_peak_7125 |
| DS117453 | 1310 | 1831 | BA_peak_7126 |
| DS117477 | 679 | 1352 | BA_peak_7127 |
| DS117484 | 1900 | 2303 | BA_peak_7128 |
| DS117487 | 545 | 897 | BA_peak_7129 |
| DS117510 | 1647 | 2084 | BA_peak_7131 |
| DS117515 | 1195 | 1661 | BA_peak_7132 |
| DS117516 | 181 | 918 | BA_peak_7133 |
| DS117528 | 107 | 681 | BA_peak_7134 |
| DS117531 | 255 | 1371 | BA_peak_7135 |
| DS117546 | 927 | 1525 | BA_peak_7136 |
| DS117549 | 951 | 2242 | BA_peak_7137 |
| DS117563 | 1794 | 2143 | BA_peak_7138 |
| DS117564 | 492 | 1250 | BA_peak_7139 |
| DS117564 | 1584 | 2297 | BA_peak_7140 |
| DS117569 | 1754 | 2197 | BA_peak_7142 |
| DS117572 | 1872 | 2280 | BA_peak_7143 |
| DS117577 | 784 | 1464 | BA_peak_7144 |
| DS117584 | 136 | 468 | BA_peak_7145 |
| DS117589 | 739 | 1135 | BA_peak_7146 |
| DS117589 | 1628 | 2122 | BA_peak_7147 |
| DS117590 | 990 | 1467 | BA_peak_7148 |
| DS117591 | 1021 | 1435 | BA_peak_7149 |
| DS117592 | 1744 | 2126 | BA_peak_7150 |
| DS117611 | 95 | 1678 | BA_peak_7151 |
| DS117621 | 201 | 888 | BA_peak_7152 |
| DS117625 | 31 | 704 | BA_peak_7153 |
| DS117636 | 175 | 714 | BA_peak_7154 |
| DS117639 | 1721 | 2155 | BA_peak_7155 |
| DS117644 | 1455 | 1972 | BA_peak_7156 |

| | | | |
|---|---|---|---|
| DS117649 | 529 | 841 | BA_peak_7157 |
| DS117649 | 1705 | 2159 | BA_peak_7158 |
| DS117650 | 1556 | 1850 | BA_peak_7159 |
| DS117654 | 180 | 793 | BA_peak_7160 |
| DS117660 | 1541 | 2184 | BA_peak_7161 |
| DS117674 | 128 | 656 | BA_peak_7162 |
| DS117678 | 873 | 1370 | BA_peak_7163 |
| DS117682 | 76 | 413 | BA_peak_7164 |
| DS117688 | 1496 | 1917 | BA_peak_7165 |
| DS117689 | 123 | 554 | BA_peak_7166 |
| DS117721 | 646 | 2168 | BA_peak_7167 |
| DS117776 | 40 | 2194 | BA_peak_7168 |
| DS117777 | 1342 | 2059 | BA_peak_7169 |
| DS117779 | 1486 | 2193 | BA_peak_7171 |
| DS117784 | 1513 | 2021 | BA_peak_7172 |
| DS117789 | 910 | 1370 | BA_peak_7173 |
| DS117798 | 109 | 548 | BA_peak_7174 |
| DS117815 | 459 | 856 | BA_peak_7175 |
| DS117821 | 1554 | 2086 | BA_peak_7176 |
| DS117826 | 141 | 1287 | BA_peak_7177 |
| DS117832 | 1344 | 2008 | BA_peak_7178 |
| DS117834 | 1542 | 1903 | BA_peak_7179 |
| DS117852 | 966 | 1503 | BA_peak_7180 |
| DS117855 | 282 | 692 | BA_peak_7181 |
| DS117859 | 56 | 482 | BA_peak_7182 |
| DS117859 | 1693 | 2039 | BA_peak_7183 |
| DS117866 | 60 | 740 | BA_peak_7184 |
| DS117866 | 1405 | 2133 | BA_peak_7185 |
| DS117874 | 752 | 1332 | BA_peak_7186 |
| DS117879 | 110 | 1219 | BA_peak_7187 |
| DS117880 | 178 | 725 | BA_peak_7188 |
| DS117893 | 852 | 1258 | BA_peak_7189 |
| DS117897 | 72 | 415 | BA_peak_7190 |
| DS117897 | 774 | 2005 | BA_peak_7191 |
| DS117900 | 430 | 838 | BA_peak_7192 |
| DS117900 | 1403 | 2154 | BA_peak_7193 |
| DS117902 | 1593 | 2135 | BA_peak_7194 |
| DS117904 | 124 | 633 | BA_peak_7195 |
| DS117907 | 1578 | 2048 | BA_peak_7196 |
| DS117911 | 1614 | 2103 | BA_peak_7197 |
| DS117915 | 1416 | 1989 | BA_peak_7198 |
| DS117924 | 915 | 1424 | BA_peak_7199 |
| DS117943 | 1633 | 1998 | BA_peak_7200 |
| DS117950 | 1323 | 1842 | BA_peak_7201 |
| DS117957 | 22 | 410 | BA_peak_7202 |
| DS117963 | 1353 | 2072 | BA_peak_7203 |
| DS117964 | 1241 | 2109 | BA_peak_7204 |

| | | | |
|---|---|---|---|
| DS117982 | 1646 | 2092 | BA_peak_7205 |
| DS117991 | 313 | 867 | BA_peak_7206 |
| DS117992 | 126 | 452 | BA_peak_7207 |
| DS117992 | 1081 | 1708 | BA_peak_7208 |
| DS118001 | 151 | 893 | BA_peak_7209 |
| DS118001 | 1331 | 1736 | BA_peak_7210 |
| DS118035 | 71 | 880 | BA_peak_7212 |
| DS118053 | 144 | 542 | BA_peak_7213 |
| DS118058 | 639 | 1292 | BA_peak_7216 |
| DS118058 | 1606 | 2002 | BA_peak_7217 |
| DS118075 | 553 | 925 | BA_peak_7218 |
| DS118076 | 109 | 630 | BA_peak_7219 |
| DS118077 | 1447 | 1928 | BA_peak_7220 |
| DS118081 | 488 | 2091 | BA_peak_7221 |
| DS118097 | 1390 | 1784 | BA_peak_7222 |
| DS118109 | 178 | 592 | BA_peak_7223 |
| DS118109 | 1697 | 2016 | BA_peak_7224 |
| DS118122 | 92 | 625 | BA_peak_7225 |
| DS118124 | 197 | 1183 | BA_peak_7226 |
| DS118133 | 293 | 772 | BA_peak_7227 |
| DS118134 | 41 | 884 | BA_peak_7228 |
| DS118148 | 333 | 937 | BA_peak_7229 |
| DS118153 | 155 | 1842 | BA_peak_7230 |
| DS118154 | 630 | 2030 | BA_peak_7231 |
| DS118167 | 185 | 683 | BA_peak_7232 |
| DS118184 | 593 | 2034 | BA_peak_7234 |
| DS118187 | 35 | 769 | BA_peak_7235 |
| DS118189 | 670 | 1619 | BA_peak_7236 |
| DS118197 | 1604 | 2017 | BA_peak_7238 |
| DS118201 | 118 | 679 | BA_peak_7239 |
| DS118207 | 76 | 544 | BA_peak_7241 |
| DS118207 | 970 | 1859 | BA_peak_7242 |
| DS118210 | 1111 | 1832 | BA_peak_7243 |
| DS118213 | 108 | 417 | BA_peak_7244 |
| DS118225 | 102 | 529 | BA_peak_7245 |
| DS118225 | 1543 | 1876 | BA_peak_7246 |
| DS118228 | 109 | 424 | BA_peak_7247 |
| DS118231 | 141 | 621 | BA_peak_7248 |
| DS118234 | 1505 | 2010 | BA_peak_7249 |
| DS118255 | 41 | 599 | BA_peak_7250 |
| DS118258 | 158 | 634 | BA_peak_7251 |
| DS118258 | 1343 | 1897 | BA_peak_7252 |
| DS118271 | 1296 | 1904 | BA_peak_7253 |
| DS118278 | 1276 | 1890 | BA_peak_7254 |
| DS118284 | 1392 | 1824 | BA_peak_7255 |
| DS118285 | 1423 | 1756 | BA_peak_7256 |
| DS118314 | 973 | 1944 | BA_peak_7257 |

| | | | |
|---|---|---|---|
| DS118327 | 102 | 890 | BA_peak_7258 |
| DS118345 | 154 | 539 | BA_peak_7259 |
| DS118363 | 44 | 584 | BA_peak_7262 |
| DS118368 | 831 | 1100 | BA_peak_7263 |
| DS118385 | 53 | 897 | BA_peak_7264 |
| DS118400 | 1140 | 1471 | BA_peak_7265 |
| DS118404 | 1449 | 1783 | BA_peak_7266 |
| DS118409 | 85 | 677 | BA_peak_7267 |
| DS118436 | 1276 | 1857 | BA_peak_7269 |
| DS118442 | 55 | 1195 | BA_peak_7270 |
| DS118442 | 1607 | 1973 | BA_peak_7271 |
| DS118454 | 231 | 962 | BA_peak_7272 |
| DS118470 | 1004 | 1660 | BA_peak_7274 |
| DS118472 | 827 | 1785 | BA_peak_7275 |
| DS118478 | 1033 | 1966 | BA_peak_7276 |
| DS118485 | 202 | 849 | BA_peak_7277 |
| DS118485 | 1623 | 1939 | BA_peak_7278 |
| DS118501 | 1535 | 1885 | BA_peak_7279 |
| DS118507 | 1470 | 1928 | BA_peak_7280 |
| DS118512 | 41 | 842 | BA_peak_7281 |
| DS118513 | 200 | 1828 | BA_peak_7282 |
| DS118519 | 23 | 579 | BA_peak_7283 |
| DS118525 | 1293 | 1958 | BA_peak_7284 |
| DS118541 | 1510 | 1891 | BA_peak_7285 |
| DS118545 | 96 | 417 | BA_peak_7287 |
| DS118552 | 1273 | 1773 | BA_peak_7288 |
| DS118554 | 149 | 1677 | BA_peak_7289 |
| DS118570 | 1272 | 1741 | BA_peak_7290 |
| DS118587 | 557 | 1517 | BA_peak_7291 |
| DS118598 | 1187 | 1820 | BA_peak_7292 |
| DS118602 | 1353 | 1890 | BA_peak_7293 |
| DS118630 | 27 | 711 | BA_peak_7294 |
| DS118634 | 70 | 1796 | BA_peak_7295 |
| DS118682 | 85 | 481 | BA_peak_7297 |
| DS118684 | 454 | 1169 | BA_peak_7298 |
| DS118693 | 1382 | 1828 | BA_peak_7299 |
| DS118697 | 1349 | 1769 | BA_peak_7300 |
| DS118703 | 159 | 1097 | BA_peak_7301 |
| DS118708 | 896 | 1243 | BA_peak_7302 |
| DS118709 | 393 | 735 | BA_peak_7303 |
| DS118720 | 177 | 512 | BA_peak_7304 |
| DS118735 | 187 | 568 | BA_peak_7305 |
| DS118738 | 591 | 1002 | BA_peak_7306 |
| DS118746 | 66 | 810 | BA_peak_7307 |
| DS118746 | 1154 | 1794 | BA_peak_7308 |
| DS118756 | 166 | 697 | BA_peak_7310 |
| DS118756 | 1182 | 1832 | BA_peak_7311 |

| | | | |
|---|---|---|---|
| DS118767 | 227 | 740 | BA_peak_7312 |
| DS118772 | 1397 | 1829 | BA_peak_7313 |
| DS118774 | 159 | 505 | BA_peak_7314 |
| DS118789 | 1497 | 1856 | BA_peak_7316 |
| DS118814 | 165 | 794 | BA_peak_7318 |
| DS118820 | 103 | 907 | BA_peak_7319 |
| DS118828 | 60 | 408 | BA_peak_7320 |
| DS118847 | 159 | 789 | BA_peak_7321 |
| DS118869 | 1441 | 1751 | BA_peak_7322 |
| DS118874 | 4 | 421 | BA_peak_7323 |
| DS118897 | 980 | 1510 | BA_peak_7324 |
| DS118924 | 80 | 782 | BA_peak_7325 |
| DS118930 | 1480 | 1860 | BA_peak_7326 |
| DS118931 | 1270 | 1844 | BA_peak_7327 |
| DS118936 | 379 | 881 | BA_peak_7328 |
| DS118936 | 1384 | 1725 | BA_peak_7329 |
| DS118940 | 321 | 881 | BA_peak_7331 |
| DS118951 | 180 | 1253 | BA_peak_7332 |
| DS118957 | 1010 | 1327 | BA_peak_7333 |
| DS118990 | 1043 | 1463 | BA_peak_7335 |
| DS118993 | 199 | 1204 | BA_peak_7336 |
| DS119000 | 723 | 1855 | BA_peak_7337 |
| DS119025 | 1070 | 1659 | BA_peak_7338 |
| DS119036 | 1272 | 1815 | BA_peak_7339 |
| DS119042 | 669 | 1842 | BA_peak_7340 |
| DS119051 | 108 | 447 | BA_peak_7341 |
| DS119060 | 223 | 677 | BA_peak_7342 |
| DS119062 | 701 | 1073 | BA_peak_7343 |
| DS119067 | 1399 | 1779 | BA_peak_7344 |
| DS119076 | 353 | 667 | BA_peak_7345 |
| DS119076 | 1046 | 1568 | BA_peak_7346 |
| DS119081 | 40 | 1051 | BA_peak_7347 |
| DS119091 | 1280 | 1828 | BA_peak_7348 |
| DS119095 | 286 | 1804 | BA_peak_7349 |
| DS119102 | 710 | 1858 | BA_peak_7350 |
| DS119108 | 26 | 1859 | BA_peak_7351 |
| DS119133 | 328 | 1031 | BA_peak_7352 |
| DS119137 | 486 | 844 | BA_peak_7353 |
| DS119144 | 270 | 588 | BA_peak_7354 |
| DS119162 | 183 | 522 | BA_peak_7355 |
| DS119167 | 1180 | 1601 | BA_peak_7356 |
| DS119171 | 924 | 1468 | BA_peak_7357 |
| DS119183 | 1420 | 1850 | BA_peak_7358 |
| DS119195 | 214 | 532 | BA_peak_7359 |
| DS119197 | 1405 | 1752 | BA_peak_7360 |
| DS119200 | 1462 | 1850 | BA_peak_7361 |
| DS119218 | 155 | 605 | BA_peak_7363 |

| | | | |
|---|---|---|---|
| DS119231 | 88 | 538 | BA_peak_7364 |
| DS119238 | 725 | 1645 | BA_peak_7365 |
| DS119242 | 86 | 501 | BA_peak_7366 |
| DS119254 | 281 | 798 | BA_peak_7367 |
| DS119261 | 1235 | 1800 | BA_peak_7368 |
| DS119270 | 56 | 1701 | BA_peak_7369 |
| DS119271 | 1260 | 1714 | BA_peak_7370 |
| DS119282 | 982 | 1446 | BA_peak_7371 |
| DS119288 | 128 | 583 | BA_peak_7372 |
| DS119326 | 941 | 1754 | BA_peak_7374 |
| DS119339 | 46 | 386 | BA_peak_7375 |
| DS119339 | 765 | 1096 | BA_peak_7376 |
| DS119341 | 102 | 482 | BA_peak_7377 |
| DS119360 | 1157 | 1655 | BA_peak_7379 |
| DS119367 | 1240 | 1733 | BA_peak_7380 |
| DS119372 | 1106 | 1534 | BA_peak_7381 |
| DS119374 | 25 | 481 | BA_peak_7382 |
| DS119377 | 47 | 435 | BA_peak_7383 |
| DS119378 | 855 | 1328 | BA_peak_7384 |
| DS119401 | 31 | 468 | BA_peak_7385 |
| DS119403 | 1346 | 1710 | BA_peak_7386 |
| DS119420 | 118 | 587 | BA_peak_7387 |
| DS119433 | 941 | 1416 | BA_peak_7388 |
| DS119438 | 1147 | 1457 | BA_peak_7389 |
| DS119449 | 270 | 670 | BA_peak_7391 |
| DS119456 | 193 | 553 | BA_peak_7392 |
| DS119470 | 1054 | 1510 | BA_peak_7393 |
| DS119476 | 1289 | 1697 | BA_peak_7394 |
| DS119481 | 782 | 1246 | BA_peak_7395 |
| DS119482 | 16 | 437 | BA_peak_7396 |
| DS119515 | 1167 | 1453 | BA_peak_7398 |
| DS119521 | 111 | 728 | BA_peak_7399 |
| DS119528 | 47 | 1797 | BA_peak_7400 |
| DS119530 | 155 | 577 | BA_peak_7401 |
| DS119566 | 27 | 450 | BA_peak_7402 |
| DS119567 | 73 | 479 | BA_peak_7403 |
| DS119603 | 1372 | 1697 | BA_peak_7405 |
| DS119608 | 22 | 494 | BA_peak_7406 |
| DS119610 | 1123 | 1530 | BA_peak_7407 |
| DS119611 | 125 | 924 | BA_peak_7408 |
| DS119622 | 127 | 659 | BA_peak_7410 |
| DS119630 | 117 | 1146 | BA_peak_7411 |
| DS119634 | 87 | 390 | BA_peak_7412 |
| DS119635 | 1481 | 1785 | BA_peak_7413 |
| DS119650 | 1097 | 1409 | BA_peak_7414 |
| DS119654 | 774 | 1630 | BA_peak_7415 |
| DS119657 | 81 | 525 | BA_peak_7416 |

| | | | |
|---|---|---|---|
| DS119662 | 207 | 538 | BA_peak_7417 |
| DS119666 | 69 | 476 | BA_peak_7418 |
| DS119668 | 33 | 482 | BA_peak_7420 |
| DS119674 | 93 | 415 | BA_peak_7422 |
| DS119674 | 941 | 1468 | BA_peak_7423 |
| DS119690 | 236 | 590 | BA_peak_7424 |
| DS119690 | 1291 | 1707 | BA_peak_7425 |
| DS119717 | 1166 | 1571 | BA_peak_7428 |
| DS119767 | 1296 | 1744 | BA_peak_7430 |
| DS119772 | 1002 | 1736 | BA_peak_7431 |
| DS119781 | 922 | 1378 | BA_peak_7432 |
| DS119792 | 155 | 723 | BA_peak_7433 |
| DS119806 | 493 | 1569 | BA_peak_7434 |
| DS119810 | 50 | 939 | BA_peak_7435 |
| DS119819 | 179 | 774 | BA_peak_7436 |
| DS119832 | 1022 | 1636 | BA_peak_7437 |
| DS119834 | 1208 | 1590 | BA_peak_7438 |
| DS119837 | 984 | 1703 | BA_peak_7439 |
| DS119867 | 575 | 1665 | BA_peak_7440 |
| DS119878 | 190 | 837 | BA_peak_7441 |
| DS119887 | 1167 | 1490 | BA_peak_7442 |
| DS119892 | 298 | 653 | BA_peak_7443 |
| DS119892 | 1142 | 1621 | BA_peak_7444 |
| DS119894 | 37 | 565 | BA_peak_7445 |
| DS119896 | 214 | 540 | BA_peak_7446 |
| DS119896 | 1216 | 1533 | BA_peak_7447 |
| DS119899 | 35 | 1467 | BA_peak_7448 |
| DS119914 | 83 | 1131 | BA_peak_7449 |
| DS119953 | 738 | 1606 | BA_peak_7450 |
| DS119958 | 135 | 508 | BA_peak_7451 |
| DS119963 | 177 | 669 | BA_peak_7452 |
| DS119966 | 89 | 566 | BA_peak_7453 |
| DS119966 | 962 | 1614 | BA_peak_7454 |
| DS119973 | 1199 | 1588 | BA_peak_7455 |
| DS119986 | 774 | 1267 | BA_peak_7456 |
| DS119992 | 34 | 694 | BA_peak_7457 |
| DS120007 | 42 | 658 | BA_peak_7458 |
| DS120007 | 1193 | 1678 | BA_peak_7459 |
| DS120010 | 74 | 560 | BA_peak_7460 |
| DS120028 | 1278 | 1658 | BA_peak_7461 |
| DS120080 | 64 | 650 | BA_peak_7462 |
| DS120089 | 1198 | 1561 | BA_peak_7463 |
| DS120091 | 101 | 1482 | BA_peak_7464 |
| DS120099 | 1215 | 1617 | BA_peak_7465 |
| DS120105 | 159 | 477 | BA_peak_7466 |
| DS120115 | 1029 | 1527 | BA_peak_7467 |
| DS120136 | 201 | 611 | BA_peak_7469 |

| | | | |
|---|---|---|---|
| DS120139 | 482 | 967 | BA_peak_7470 |
| DS120150 | 42 | 1122 | BA_peak_7473 |
| DS120163 | 233 | 596 | BA_peak_7475 |
| DS120174 | 1165 | 1626 | BA_peak_7476 |
| DS120176 | 358 | 738 | BA_peak_7477 |
| DS120184 | 46 | 525 | BA_peak_7478 |
| DS120198 | 191 | 626 | BA_peak_7479 |
| DS120206 | 782 | 1696 | BA_peak_7480 |
| DS120210 | 89 | 685 | BA_peak_7481 |
| DS120215 | 164 | 635 | BA_peak_7482 |
| DS120216 | 1320 | 1650 | BA_peak_7483 |
| DS120222 | 46 | 1721 | BA_peak_7484 |
| DS120249 | 1036 | 1632 | BA_peak_7485 |
| DS120252 | 365 | 748 | BA_peak_7486 |
| DS120269 | 1142 | 1643 | BA_peak_7487 |
| DS120281 | 128 | 534 | BA_peak_7488 |
| DS120281 | 898 | 1503 | BA_peak_7489 |
| DS120283 | 224 | 631 | BA_peak_7490 |
| DS120322 | 101 | 606 | BA_peak_7491 |
| DS120332 | 973 | 1498 | BA_peak_7492 |
| DS120337 | 1156 | 1669 | BA_peak_7493 |
| DS120338 | 133 | 548 | BA_peak_7494 |
| DS120342 | 37 | 528 | BA_peak_7495 |
| DS120348 | 137 | 759 | BA_peak_7496 |
| DS120362 | 31 | 1616 | BA_peak_7498 |
| DS120400 | 924 | 1294 | BA_peak_7499 |
| DS120413 | 1000 | 1369 | BA_peak_7500 |
| DS120418 | 996 | 1563 | BA_peak_7501 |
| DS120433 | 1128 | 1592 | BA_peak_7502 |
| DS120453 | 167 | 484 | BA_peak_7503 |
| DS120458 | 152 | 557 | BA_peak_7504 |
| DS120458 | 870 | 1289 | BA_peak_7505 |
| DS120498 | 166 | 678 | BA_peak_7507 |
| DS120524 | 32 | 766 | BA_peak_7508 |
| DS120553 | 506 | 1480 | BA_peak_7509 |
| DS120555 | 786 | 1544 | BA_peak_7510 |
| DS120565 | 507 | 1480 | BA_peak_7511 |
| DS120576 | 1301 | 1641 | BA_peak_7512 |
| DS120577 | 263 | 753 | BA_peak_7513 |
| DS120581 | 1044 | 1465 | BA_peak_7514 |
| DS120585 | 133 | 643 | BA_peak_7515 |
| DS120587 | 270 | 702 | BA_peak_7516 |
| DS120590 | 792 | 1542 | BA_peak_7517 |
| DS120594 | 303 | 610 | BA_peak_7518 |
| DS120620 | 1234 | 1654 | BA_peak_7519 |
| DS120623 | 1234 | 1674 | BA_peak_7520 |
| DS120629 | 126 | 616 | BA_peak_7521 |

| | | | |
|---|---|---|---|
| DS120632 | 65 | 649 | BA_peak_7522 |
| DS120642 | 241 | 688 | BA_peak_7524 |
| DS120682 | 112 | 767 | BA_peak_7526 |
| DS120687 | 13 | 660 | BA_peak_7527 |
| DS120702 | 35 | 376 | BA_peak_7528 |
| DS120713 | 180 | 706 | BA_peak_7530 |
| DS120714 | 468 | 948 | BA_peak_7531 |
| DS120717 | 102 | 644 | BA_peak_7532 |
| DS120725 | 233 | 620 | BA_peak_7533 |
| DS120734 | 1186 | 1509 | BA_peak_7534 |
| DS120735 | 73 | 551 | BA_peak_7535 |
| DS120736 | 902 | 1607 | BA_peak_7536 |
| DS120738 | 203 | 675 | BA_peak_7537 |
| DS120758 | 77 | 574 | BA_peak_7538 |
| DS120762 | 24 | 432 | BA_peak_7539 |
| DS120763 | 59 | 581 | BA_peak_7540 |
| DS120776 | 274 | 1690 | BA_peak_7541 |
| DS120798 | 974 | 1344 | BA_peak_7543 |
| DS120810 | 913 | 1439 | BA_peak_7544 |
| DS120813 | 208 | 593 | BA_peak_7545 |
| DS120816 | 27 | 464 | BA_peak_7546 |
| DS120839 | 1180 | 1483 | BA_peak_7547 |
| DS120841 | 1132 | 1566 | BA_peak_7548 |
| DS120852 | 919 | 1559 | BA_peak_7549 |
| DS120856 | 1130 | 1666 | BA_peak_7550 |
| DS120887 | 120 | 792 | BA_peak_7551 |
| DS120891 | 865 | 1387 | BA_peak_7552 |
| DS120899 | 99 | 523 | BA_peak_7553 |
| DS120916 | 916 | 1435 | BA_peak_7554 |
| DS120917 | 1286 | 1608 | BA_peak_7555 |
| DS120923 | 118 | 1180 | BA_peak_7556 |
| DS120930 | 843 | 1284 | BA_peak_7557 |
| DS120933 | 37 | 894 | BA_peak_7558 |
| DS120942 | 209 | 599 | BA_peak_7559 |
| DS120945 | 1169 | 1509 | BA_peak_7560 |
| DS120949 | 141 | 982 | BA_peak_7561 |
| DS120964 | 120 | 450 | BA_peak_7562 |
| DS120977 | 988 | 1660 | BA_peak_7563 |
| DS120985 | 178 | 501 | BA_peak_7564 |
| DS120986 | 962 | 1448 | BA_peak_7566 |
| DS120997 | 65 | 664 | BA_peak_7567 |
| DS121013 | 303 | 687 | BA_peak_7568 |
| DS121021 | 515 | 969 | BA_peak_7570 |
| DS121026 | 64 | 816 | BA_peak_7571 |
| DS121036 | 1107 | 1567 | BA_peak_7572 |
| DS121065 | 889 | 1426 | BA_peak_7574 |
| DS121072 | 79 | 1656 | BA_peak_7575 |

| | | | |
|---|---|---|---|
| DS121081 | 791 | 1510 | BA_peak_7578 |
| DS121084 | 128 | 624 | BA_peak_7579 |
| DS121087 | 879 | 1234 | BA_peak_7580 |
| DS121100 | 115 | 759 | BA_peak_7581 |
| DS121111 | 195 | 1521 | BA_peak_7582 |
| DS121148 | 1199 | 1645 | BA_peak_7584 |
| DS121162 | 263 | 850 | BA_peak_7587 |
| DS121181 | 869 | 1256 | BA_peak_7588 |
| DS121185 | 960 | 1638 | BA_peak_7589 |
| DS121203 | 1267 | 1640 | BA_peak_7590 |
| DS121227 | 1173 | 1526 | BA_peak_7591 |
| DS121228 | 1072 | 1431 | BA_peak_7592 |
| DS121232 | 1031 | 1539 | BA_peak_7593 |
| DS121249 | 1015 | 1648 | BA_peak_7595 |
| DS121266 | 758 | 1273 | BA_peak_7596 |
| DS121268 | 85 | 850 | BA_peak_7597 |
| DS121297 | 1164 | 1474 | BA_peak_7598 |
| DS121300 | 136 | 608 | BA_peak_7599 |
| DS121306 | 90 | 657 | BA_peak_7600 |
| DS121316 | 186 | 629 | BA_peak_7601 |
| DS121321 | 653 | 1147 | BA_peak_7602 |
| DS121330 | 812 | 1476 | BA_peak_7603 |
| DS121333 | 1023 | 1504 | BA_peak_7604 |
| DS121341 | 277 | 663 | BA_peak_7605 |
| DS121348 | 449 | 892 | BA_peak_7606 |
| DS121365 | 42 | 807 | BA_peak_7607 |
| DS121367 | 189 | 525 | BA_peak_7608 |
| DS121367 | 851 | 1606 | BA_peak_7609 |
| DS121368 | 34 | 375 | BA_peak_7610 |
| DS121378 | 26 | 694 | BA_peak_7611 |
| DS121394 | 109 | 660 | BA_peak_7612 |
| DS121397 | 193 | 530 | BA_peak_7613 |
| DS121420 | 851 | 1619 | BA_peak_7614 |
| DS121425 | 94 | 433 | BA_peak_7615 |
| DS121425 | 1002 | 1572 | BA_peak_7616 |
| DS121435 | 930 | 1507 | BA_peak_7617 |
| DS121447 | 109 | 623 | BA_peak_7618 |
| DS121462 | 75 | 473 | BA_peak_7619 |
| DS121465 | 156 | 585 | BA_peak_7620 |
| DS121471 | 952 | 1464 | BA_peak_7621 |
| DS121484 | 67 | 1235 | BA_peak_7622 |
| DS121494 | 1125 | 1458 | BA_peak_7623 |
| DS121499 | 204 | 824 | BA_peak_7624 |
| DS121507 | 975 | 1307 | BA_peak_7625 |
| DS121516 | 890 | 1402 | BA_peak_7626 |
| DS121539 | 105 | 1128 | BA_peak_7628 |
| DS121543 | 907 | 1615 | BA_peak_7629 |

| | | | |
|---|---|---|---|
| DS121550 | 1002 | 1350 | BA_peak_7630 |
| DS121560 | 1240 | 1583 | BA_peak_7631 |
| DS121587 | 938 | 1391 | BA_peak_7632 |
| DS121590 | 768 | 1597 | BA_peak_7633 |
| DS121600 | 77 | 585 | BA_peak_7634 |
| DS121634 | 74 | 567 | BA_peak_7636 |
| DS121637 | 51 | 481 | BA_peak_7637 |
| DS121647 | 58 | 468 | BA_peak_7638 |
| DS121661 | 171 | 620 | BA_peak_7639 |
| DS121661 | 1025 | 1556 | BA_peak_7640 |
| DS121664 | 194 | 639 | BA_peak_7641 |
| DS121664 | 1133 | 1581 | BA_peak_7642 |
| DS121669 | 898 | 1601 | BA_peak_7643 |
| DS121672 | 1057 | 1466 | BA_peak_7644 |
| DS121676 | 436 | 800 | BA_peak_7645 |
| DS121677 | 153 | 489 | BA_peak_7646 |
| DS121681 | 1161 | 1551 | BA_peak_7647 |
| DS121700 | 235 | 538 | BA_peak_7648 |
| DS121720 | 888 | 1313 | BA_peak_7650 |
| DS121725 | 467 | 1540 | BA_peak_7651 |
| DS121737 | 1178 | 1552 | BA_peak_7652 |
| DS121747 | 285 | 785 | BA_peak_7653 |
| DS121757 | 898 | 1554 | BA_peak_7654 |
| DS121790 | 275 | 642 | BA_peak_7655 |
| DS121800 | 689 | 1337 | BA_peak_7656 |
| DS121803 | 1030 | 1462 | BA_peak_7657 |
| DS121815 | 924 | 1611 | BA_peak_7658 |
| DS121824 | 127 | 1579 | BA_peak_7659 |
| DS121825 | 737 | 1544 | BA_peak_7660 |
| DS121828 | 34 | 518 | BA_peak_7661 |
| DS121848 | 1281 | 1600 | BA_peak_7662 |
| DS121857 | 846 | 1560 | BA_peak_7663 |
| DS121858 | 61 | 1119 | BA_peak_7664 |
| DS121859 | 1040 | 1414 | BA_peak_7665 |
| DS121876 | 841 | 1544 | BA_peak_7666 |
| DS121887 | 270 | 619 | BA_peak_7667 |
| DS121900 | 123 | 524 | BA_peak_7668 |
| DS121917 | 214 | 1527 | BA_peak_7669 |
| DS121945 | 446 | 795 | BA_peak_7670 |
| DS121951 | 1188 | 1559 | BA_peak_7671 |
| DS121963 | 174 | 726 | BA_peak_7672 |
| DS121974 | 645 | 1588 | BA_peak_7673 |
| DS122001 | 349 | 751 | BA_peak_7674 |
| DS122013 | 216 | 1387 | BA_peak_7676 |
| DS122018 | 78 | 1014 | BA_peak_7677 |
| DS122026 | 857 | 1528 | BA_peak_7679 |
| DS122028 | 1057 | 1582 | BA_peak_7680 |

| | | | |
|---|---|---|---|
| DS122036 | 804 | 1381 | BA_peak_7681 |
| DS122047 | 862 | 1555 | BA_peak_7682 |
| DS122064 | 168 | 521 | BA_peak_7683 |
| DS122067 | 830 | 1327 | BA_peak_7684 |
| DS122082 | 902 | 1349 | BA_peak_7686 |
| DS122094 | 467 | 1013 | BA_peak_7688 |
| DS122113 | 185 | 691 | BA_peak_7690 |
| DS122115 | 995 | 1353 | BA_peak_7691 |
| DS122121 | 766 | 1380 | BA_peak_7692 |
| DS122123 | 766 | 1342 | BA_peak_7693 |
| DS122134 | 556 | 1387 | BA_peak_7694 |
| DS122140 | 26 | 575 | BA_peak_7695 |
| DS122142 | 1217 | 1532 | BA_peak_7696 |
| DS122153 | 117 | 1560 | BA_peak_7697 |
| DS122159 | 251 | 785 | BA_peak_7698 |
| DS122165 | 213 | 612 | BA_peak_7699 |
| DS122175 | 287 | 1493 | BA_peak_7700 |
| DS122182 | 1183 | 1575 | BA_peak_7701 |
| DS122189 | 210 | 697 | BA_peak_7702 |
| DS122194 | 822 | 1366 | BA_peak_7703 |
| DS122214 | 1171 | 1572 | BA_peak_7704 |
| DS122234 | 68 | 1518 | BA_peak_7705 |
| DS122244 | 235 | 542 | BA_peak_7706 |
| DS122251 | 223 | 509 | BA_peak_7707 |
| DS122251 | 1035 | 1550 | BA_peak_7708 |
| DS122266 | 220 | 1378 | BA_peak_7709 |
| DS122267 | 206 | 552 | BA_peak_7710 |
| DS122283 | 72 | 430 | BA_peak_7711 |
| DS122283 | 779 | 1442 | BA_peak_7712 |
| DS122294 | 66 | 583 | BA_peak_7713 |
| DS122294 | 1059 | 1425 | BA_peak_7714 |
| DS122308 | 1175 | 1542 | BA_peak_7715 |
| DS122312 | 927 | 1566 | BA_peak_7716 |
| DS122337 | 1094 | 1457 | BA_peak_7717 |
| DS122358 | 199 | 516 | BA_peak_7718 |
| DS122370 | 57 | 548 | BA_peak_7719 |
| DS122383 | 214 | 1497 | BA_peak_7720 |
| DS122387 | 252 | 1194 | BA_peak_7721 |
| DS122392 | 1021 | 1554 | BA_peak_7723 |
| DS122401 | 1055 | 1536 | BA_peak_7724 |
| DS122408 | 131 | 1457 | BA_peak_7725 |
| DS122412 | 1058 | 1354 | BA_peak_7726 |
| DS122420 | 84 | 1457 | BA_peak_7727 |
| DS122425 | 700 | 1126 | BA_peak_7728 |
| DS122429 | 937 | 1352 | BA_peak_7729 |
| DS122440 | 106 | 548 | BA_peak_7730 |
| DS122449 | 43 | 1462 | BA_peak_7731 |

| | | | |
|---|---|---|---|
| DS122471 | 926 | 1427 | BA_peak_7734 |
| DS122474 | 42 | 807 | BA_peak_7735 |
| DS122490 | 85 | 992 | BA_peak_7736 |
| DS122493 | 287 | 629 | BA_peak_7737 |
| DS122506 | 298 | 1275 | BA_peak_7738 |
| DS122511 | 952 | 1424 | BA_peak_7739 |
| DS122512 | 1112 | 1453 | BA_peak_7740 |
| DS122522 | 39 | 398 | BA_peak_7742 |
| DS122580 | 57 | 1489 | BA_peak_7745 |
| DS122595 | 1089 | 1435 | BA_peak_7746 |
| DS122601 | 834 | 1539 | BA_peak_7747 |
| DS122617 | 80 | 484 | BA_peak_7749 |
| DS122632 | 187 | 593 | BA_peak_7750 |
| DS122645 | 114 | 520 | BA_peak_7751 |
| DS122655 | 86 | 469 | BA_peak_7752 |
| DS122657 | 210 | 685 | BA_peak_7753 |
| DS122682 | 45 | 404 | BA_peak_7754 |
| DS122689 | 1068 | 1476 | BA_peak_7755 |
| DS122719 | 1117 | 1540 | BA_peak_7756 |
| DS122729 | 1085 | 1547 | BA_peak_7757 |
| DS122734 | 58 | 708 | BA_peak_7758 |
| DS122734 | 1016 | 1515 | BA_peak_7759 |
| DS122745 | 743 | 1453 | BA_peak_7760 |
| DS122761 | 50 | 581 | BA_peak_7761 |
| DS122776 | 440 | 1411 | BA_peak_7762 |
| DS122786 | 993 | 1504 | BA_peak_7763 |
| DS122806 | 198 | 542 | BA_peak_7764 |
| DS122824 | 395 | 1522 | BA_peak_7765 |
| DS122834 | 892 | 1360 | BA_peak_7766 |
| DS122866 | 92 | 477 | BA_peak_7767 |
| DS122881 | 148 | 1173 | BA_peak_7768 |
| DS122886 | 91 | 632 | BA_peak_7770 |
| DS122904 | 43 | 1525 | BA_peak_7771 |
| DS122915 | 40 | 528 | BA_peak_7772 |
| DS122939 | 175 | 1207 | BA_peak_7773 |
| DS122944 | 997 | 1460 | BA_peak_7774 |
| DS122953 | 188 | 610 | BA_peak_7775 |
| DS122954 | 1137 | 1542 | BA_peak_7776 |
| DS122960 | 204 | 625 | BA_peak_7777 |
| DS122965 | 933 | 1326 | BA_peak_7778 |
| DS122993 | 886 | 1254 | BA_peak_7779 |
| DS122996 | 933 | 1506 | BA_peak_7780 |
| DS123002 | 949 | 1269 | BA_peak_7781 |
| DS123021 | 219 | 676 | BA_peak_7782 |
| DS123028 | 155 | 1279 | BA_peak_7783 |
| DS123046 | 211 | 646 | BA_peak_7784 |
| DS123048 | 695 | 1328 | BA_peak_7785 |

| | | | |
|---|---|---|---|
| DS123049 | 44 | 592 | BA_peak_7786 |
| DS123055 | 998 | 1432 | BA_peak_7787 |
| DS123064 | 44 | 543 | BA_peak_7788 |
| DS123065 | 1072 | 1528 | BA_peak_7789 |
| DS123086 | 1007 | 1469 | BA_peak_7790 |
| DS123088 | 948 | 1360 | BA_peak_7792 |
| DS123094 | 144 | 698 | BA_peak_7793 |
| DS123095 | 1201 | 1524 | BA_peak_7794 |
| DS123104 | 123 | 624 | BA_peak_7795 |
| DS123117 | 6 | 449 | BA_peak_7796 |
| DS123124 | 34 | 1472 | BA_peak_7797 |
| DS123125 | 31 | 398 | BA_peak_7798 |
| DS123132 | 518 | 1149 | BA_peak_7799 |
| DS123153 | 1127 | 1546 | BA_peak_7801 |
| DS123158 | 936 | 1495 | BA_peak_7802 |
| DS123172 | 1103 | 1543 | BA_peak_7803 |
| DS123187 | 203 | 1040 | BA_peak_7804 |
| DS123189 | 22 | 640 | BA_peak_7805 |
| DS123199 | 167 | 603 | BA_peak_7806 |
| DS123249 | 79 | 1177 | BA_peak_7807 |
| DS123263 | 866 | 1198 | BA_peak_7808 |
| DS123270 | 57 | 1253 | BA_peak_7810 |
| DS123313 | 304 | 727 | BA_peak_7812 |
| DS123331 | 744 | 1322 | BA_peak_7813 |
| DS123337 | 43 | 922 | BA_peak_7814 |
| DS123369 | 16 | 373 | BA_peak_7816 |
| DS123369 | 873 | 1480 | BA_peak_7817 |
| DS123377 | 836 | 1444 | BA_peak_7818 |
| DS123398 | 1103 | 1429 | BA_peak_7819 |
| DS123405 | 1073 | 1513 | BA_peak_7820 |
| DS123407 | 157 | 527 | BA_peak_7821 |
| DS123425 | 917 | 1351 | BA_peak_7822 |
| DS123476 | 1126 | 1440 | BA_peak_7823 |
| DS123500 | 481 | 807 | BA_peak_7824 |
| DS123515 | 922 | 1507 | BA_peak_7825 |
| DS123548 | 1025 | 1456 | BA_peak_7826 |
| DS123574 | 301 | 623 | BA_peak_7828 |
| DS123595 | 915 | 1485 | BA_peak_7829 |
| DS123606 | 837 | 1487 | BA_peak_7830 |
| DS123623 | 692 | 1120 | BA_peak_7831 |
| DS123635 | 823 | 1432 | BA_peak_7833 |
| DS123651 | 130 | 1329 | BA_peak_7834 |
| DS123680 | 111 | 629 | BA_peak_7835 |
| DS123686 | 808 | 1261 | BA_peak_7836 |
| DS123688 | 595 | 996 | BA_peak_7837 |
| DS123689 | 80 | 438 | BA_peak_7838 |
| DS123693 | 299 | 828 | BA_peak_7839 |

| | | | |
|---|---|---|---|
| DS123694 | 789 | 1454 | BA_peak_7840 |
| DS123729 | 194 | 1467 | BA_peak_7841 |
| DS123738 | 147 | 1413 | BA_peak_7843 |
| DS123745 | 764 | 1200 | BA_peak_7845 |
| DS123750 | 981 | 1405 | BA_peak_7846 |
| DS123766 | 76 | 849 | BA_peak_7847 |
| DS123777 | 829 | 1276 | BA_peak_7848 |
| DS123792 | 270 | 607 | BA_peak_7849 |
| DS123818 | 18 | 609 | BA_peak_7851 |
| DS123818 | 976 | 1434 | BA_peak_7852 |
| DS123833 | 273 | 658 | BA_peak_7853 |
| DS123838 | 465 | 1394 | BA_peak_7854 |
| DS123849 | 153 | 795 | BA_peak_7855 |
| DS123854 | 13 | 711 | BA_peak_7856 |
| DS123860 | 122 | 663 | BA_peak_7857 |
| DS123861 | 850 | 1371 | BA_peak_7858 |
| DS123868 | 90 | 547 | BA_peak_7859 |
| DS123876 | 707 | 1400 | BA_peak_7860 |
| DS123885 | 50 | 430 | BA_peak_7861 |
| DS123892 | 36 | 1499 | BA_peak_7862 |
| DS123900 | 888 | 1387 | BA_peak_7863 |
| DS123907 | 921 | 1313 | BA_peak_7864 |
| DS123943 | 980 | 1337 | BA_peak_7866 |
| DS123945 | 376 | 801 | BA_peak_7867 |
| DS123965 | 529 | 1104 | BA_peak_7868 |
| DS123967 | 474 | 834 | BA_peak_7869 |
| DS123970 | 63 | 812 | BA_peak_7870 |
| DS123971 | 587 | 1082 | BA_peak_7871 |
| DS123981 | 328 | 1331 | BA_peak_7872 |
| DS123985 | 75 | 671 | BA_peak_7873 |
| DS124004 | 130 | 631 | BA_peak_7874 |
| DS124005 | 1135 | 1459 | BA_peak_7875 |
| DS124014 | 203 | 509 | BA_peak_7876 |
| DS124020 | 17 | 1399 | BA_peak_7877 |
| DS124073 | 942 | 1270 | BA_peak_7878 |
| DS124102 | 1044 | 1463 | BA_peak_7880 |
| DS124122 | 1002 | 1400 | BA_peak_7881 |
| DS124126 | 42 | 379 | BA_peak_7882 |
| DS124126 | 970 | 1346 | BA_peak_7883 |
| DS124147 | 898 | 1229 | BA_peak_7884 |
| DS124184 | 304 | 620 | BA_peak_7885 |
| DS124236 | 1162 | 1479 | BA_peak_7886 |
| DS124253 | 867 | 1469 | BA_peak_7887 |
| DS124261 | 235 | 762 | BA_peak_7888 |
| DS124284 | 880 | 1254 | BA_peak_7889 |
| DS124298 | 82 | 620 | BA_peak_7890 |
| DS124299 | 62 | 493 | BA_peak_7891 |

| | | | |
|---|---|---|---|
| DS124319 | 651 | 1057 | BA_peak_7892 |
| DS124327 | 670 | 1148 | BA_peak_7893 |
| DS124337 | 194 | 706 | BA_peak_7894 |
| DS124345 | 82 | 682 | BA_peak_7895 |
| DS124362 | 1076 | 1413 | BA_peak_7897 |
| DS124367 | 781 | 1432 | BA_peak_7898 |
| DS124374 | 1083 | 1470 | BA_peak_7899 |
| DS124379 | 328 | 750 | BA_peak_7900 |
| DS124392 | 15 | 453 | BA_peak_7901 |
| DS124412 | 716 | 1328 | BA_peak_7902 |
| DS124414 | 90 | 648 | BA_peak_7903 |
| DS124433 | 1049 | 1442 | BA_peak_7904 |
| DS124444 | 223 | 576 | BA_peak_7905 |
| DS124448 | 1087 | 1355 | BA_peak_7906 |
| DS124477 | 19 | 458 | BA_peak_7907 |
| DS124478 | 970 | 1449 | BA_peak_7908 |
| DS124481 | 884 | 1422 | BA_peak_7909 |
| DS124515 | 611 | 1064 | BA_peak_7911 |
| DS124520 | 474 | 827 | BA_peak_7912 |
| DS124533 | 207 | 609 | BA_peak_7913 |
| DS124575 | 872 | 1235 | BA_peak_7914 |
| DS124579 | 102 | 591 | BA_peak_7915 |
| DS124617 | 460 | 1069 | BA_peak_7916 |
| DS124624 | 877 | 1323 | BA_peak_7917 |
| DS124640 | 123 | 621 | BA_peak_7918 |
| DS124641 | 388 | 840 | BA_peak_7919 |
| DS124662 | 290 | 610 | BA_peak_7920 |
| DS124699 | 362 | 894 | BA_peak_7921 |
| DS124705 | 89 | 570 | BA_peak_7922 |
| DS124706 | 866 | 1425 | BA_peak_7923 |
| DS124730 | 1036 | 1418 | BA_peak_7924 |
| DS124731 | 845 | 1454 | BA_peak_7925 |
| DS124733 | 31 | 390 | BA_peak_7926 |
| DS124740 | 75 | 1271 | BA_peak_7927 |
| DS124742 | 462 | 1455 | BA_peak_7928 |
| DS124747 | 42 | 1380 | BA_peak_7929 |
| DS124749 | 901 | 1359 | BA_peak_7931 |
| DS124782 | 952 | 1384 | BA_peak_7932 |
| DS124786 | 724 | 1259 | BA_peak_7933 |
| DS124795 | 933 | 1327 | BA_peak_7934 |
| DS124810 | 64 | 447 | BA_peak_7935 |
| DS124810 | 886 | 1378 | BA_peak_7936 |
| DS124852 | 1102 | 1437 | BA_peak_7937 |
| DS124871 | 1003 | 1306 | BA_peak_7939 |
| DS124879 | 486 | 1430 | BA_peak_7940 |
| DS124892 | 189 | 752 | BA_peak_7942 |
| DS124897 | 106 | 1137 | BA_peak_7943 |

| | | | |
|---|---|---|---|
| DS124911 | 105 | 659 | BA_peak_7945 |
| DS124917 | 786 | 1406 | BA_peak_7946 |
| DS124923 | 1006 | 1409 | BA_peak_7947 |
| DS124934 | 105 | 453 | BA_peak_7948 |
| DS124961 | 726 | 1375 | BA_peak_7949 |
| DS124969 | 755 | 1049 | BA_peak_7950 |
| DS125001 | 756 | 1368 | BA_peak_7951 |
| DS125013 | 686 | 1081 | BA_peak_7953 |
| DS125015 | 497 | 1374 | BA_peak_7954 |
| DS125033 | 990 | 1403 | BA_peak_7955 |
| DS125034 | 27 | 480 | BA_peak_7956 |
| DS125051 | 82 | 570 | BA_peak_7957 |
| DS125054 | 24 | 382 | BA_peak_7958 |
| DS125100 | 334 | 1225 | BA_peak_7960 |
| DS125104 | 800 | 1389 | BA_peak_7961 |
| DS125111 | 333 | 841 | BA_peak_7963 |
| DS125139 | 277 | 1267 | BA_peak_7964 |
| DS125144 | 175 | 1233 | BA_peak_7965 |
| DS125147 | 482 | 1413 | BA_peak_7966 |
| DS125162 | 89 | 1398 | BA_peak_7968 |
| DS125171 | 206 | 536 | BA_peak_7969 |
| DS125181 | 80 | 409 | BA_peak_7970 |
| DS125211 | 735 | 1154 | BA_peak_7971 |
| DS125220 | 176 | 1220 | BA_peak_7972 |
| DS125222 | 354 | 930 | BA_peak_7973 |
| DS125241 | 969 | 1353 | BA_peak_7974 |
| DS125244 | 118 | 1148 | BA_peak_7975 |
| DS125249 | 199 | 1349 | BA_peak_7976 |
| DS125251 | 760 | 1161 | BA_peak_7977 |
| DS125256 | 762 | 1420 | BA_peak_7978 |
| DS125267 | 959 | 1414 | BA_peak_7979 |
| DS125273 | 904 | 1423 | BA_peak_7980 |
| DS125299 | 881 | 1279 | BA_peak_7982 |
| DS125304 | 48 | 704 | BA_peak_7983 |
| DS125320 | 533 | 1415 | BA_peak_7985 |
| DS125321 | 188 | 895 | BA_peak_7986 |
| DS125327 | 26 | 535 | BA_peak_7987 |
| DS125344 | 892 | 1283 | BA_peak_7988 |
| DS125356 | 276 | 873 | BA_peak_7989 |
| DS125360 | 25 | 423 | BA_peak_7990 |
| DS125390 | 902 | 1309 | BA_peak_7991 |
| DS125391 | 102 | 1253 | BA_peak_7992 |
| DS125397 | 133 | 1432 | BA_peak_7993 |
| DS125431 | 1021 | 1409 | BA_peak_7994 |
| DS125467 | 111 | 750 | BA_peak_7995 |
| DS125471 | 29 | 457 | BA_peak_7996 |
| DS125476 | 1002 | 1368 | BA_peak_7997 |

| | | | |
|---|---|---|---|
| DS125477 | 121 | 836 | BA_peak_7998 |
| DS125484 | 165 | 489 | BA_peak_7999 |
| DS125492 | 626 | 1419 | BA_peak_8000 |
| DS125495 | 150 | 637 | BA_peak_8001 |
| DS125498 | 355 | 1131 | BA_peak_8002 |
| DS125502 | 709 | 1190 | BA_peak_8003 |
| DS125514 | 288 | 597 | BA_peak_8004 |
| DS125545 | 63 | 555 | BA_peak_8005 |
| DS125545 | 1008 | 1423 | BA_peak_8006 |
| DS125563 | 277 | 929 | BA_peak_8008 |
| DS125579 | 124 | 610 | BA_peak_8010 |
| DS125587 | 189 | 612 | BA_peak_8011 |
| DS125596 | 128 | 514 | BA_peak_8012 |
| DS125598 | 593 | 1353 | BA_peak_8014 |
| DS125626 | 642 | 1271 | BA_peak_8016 |
| DS125631 | 879 | 1278 | BA_peak_8017 |
| DS125634 | 665 | 1342 | BA_peak_8018 |
| DS125640 | 98 | 1356 | BA_peak_8019 |
| DS125672 | 525 | 1308 | BA_peak_8020 |
| DS125678 | 285 | 1378 | BA_peak_8021 |
| DS125698 | 159 | 1365 | BA_peak_8022 |
| DS125702 | 718 | 1402 | BA_peak_8023 |
| DS125717 | 26 | 617 | BA_peak_8025 |
| DS125725 | 38 | 505 | BA_peak_8026 |
| DS125753 | 1009 | 1333 | BA_peak_8027 |
| DS125759 | 1024 | 1334 | BA_peak_8028 |
| DS125764 | 708 | 1364 | BA_peak_8029 |
| DS125773 | 67 | 1284 | BA_peak_8030 |
| DS125783 | 689 | 1113 | BA_peak_8031 |
| DS125784 | 681 | 1256 | BA_peak_8032 |
| DS125799 | 809 | 1378 | BA_peak_8033 |
| DS125807 | 547 | 1310 | BA_peak_8034 |
| DS125825 | 300 | 603 | BA_peak_8035 |
| DS125855 | 994 | 1372 | BA_peak_8037 |
| DS125860 | 409 | 1127 | BA_peak_8038 |
| DS125872 | 246 | 785 | BA_peak_8039 |
| DS125912 | 565 | 1203 | BA_peak_8040 |
| DS125915 | 76 | 643 | BA_peak_8041 |
| DS125985 | 662 | 1177 | BA_peak_8044 |
| DS125996 | 944 | 1409 | BA_peak_8045 |
| DS126013 | 340 | 688 | BA_peak_8046 |
| DS126020 | 145 | 557 | BA_peak_8047 |
| DS126036 | 76 | 693 | BA_peak_8048 |
| DS126042 | 85 | 631 | BA_peak_8049 |
| DS126061 | 25 | 404 | BA_peak_8050 |
| DS126065 | 662 | 1284 | BA_peak_8051 |
| DS126079 | 125 | 1317 | BA_peak_8052 |

| | | | |
|---|---|---|---|
| DS126080 | 684 | 1375 | BA_peak_8053 |
| DS126108 | 324 | 729 | BA_peak_8054 |
| DS126112 | 886 | 1235 | BA_peak_8055 |
| DS126152 | 39 | 666 | BA_peak_8057 |
| DS126178 | 71 | 1339 | BA_peak_8058 |
| DS126184 | 107 | 1373 | BA_peak_8059 |
| DS126186 | 258 | 660 | BA_peak_8060 |
| DS126190 | 103 | 503 | BA_peak_8061 |
| DS126203 | 107 | 713 | BA_peak_8062 |
| DS126215 | 89 | 633 | BA_peak_8063 |
| DS126215 | 981 | 1373 | BA_peak_8064 |
| DS126218 | 46 | 655 | BA_peak_8065 |
| DS126237 | 84 | 664 | BA_peak_8066 |
| DS126241 | 158 | 1212 | BA_peak_8067 |
| DS126244 | 781 | 1304 | BA_peak_8068 |
| DS126251 | 502 | 960 | BA_peak_8069 |
| DS126253 | 120 | 929 | BA_peak_8070 |
| DS126259 | 737 | 1207 | BA_peak_8071 |
| DS126279 | 920 | 1292 | BA_peak_8073 |
| DS126321 | 825 | 1249 | BA_peak_8074 |
| DS126408 | 181 | 510 | BA_peak_8078 |
| DS126430 | 90 | 553 | BA_peak_8079 |
| DS126435 | 495 | 832 | BA_peak_8080 |
| DS126442 | 150 | 873 | BA_peak_8081 |
| DS126473 | 870 | 1202 | BA_peak_8082 |
| DS126477 | 360 | 1155 | BA_peak_8083 |
| DS126482 | 54 | 486 | BA_peak_8084 |
| DS126498 | 720 | 1091 | BA_peak_8086 |
| DS126502 | 131 | 558 | BA_peak_8087 |
| DS126503 | 211 | 1081 | BA_peak_8088 |
| DS126535 | 240 | 755 | BA_peak_8089 |
| DS126546 | 138 | 1393 | BA_peak_8090 |
| DS126557 | 737 | 1379 | BA_peak_8091 |
| DS126559 | 1014 | 1387 | BA_peak_8092 |
| DS126573 | 177 | 1337 | BA_peak_8094 |
| DS126574 | 744 | 1355 | BA_peak_8095 |
| DS126576 | 654 | 1303 | BA_peak_8096 |
| DS126607 | 719 | 1285 | BA_peak_8097 |
| DS126619 | 542 | 1031 | BA_peak_8098 |
| DS126629 | 167 | 524 | BA_peak_8099 |
| DS126635 | 894 | 1309 | BA_peak_8100 |
| DS126682 | 867 | 1218 | BA_peak_8102 |
| DS126693 | 148 | 1197 | BA_peak_8103 |
| DS126694 | 737 | 1283 | BA_peak_8104 |
| DS126699 | 185 | 535 | BA_peak_8105 |
| DS126699 | 941 | 1319 | BA_peak_8106 |
| DS126730 | 33 | 591 | BA_peak_8107 |

| | | | |
|---|---|---|---|
| DS126730 | 942 | 1336 | BA_peak_8108 |
| DS126745 | 115 | 515 | BA_peak_8109 |
| DS126757 | 818 | 1311 | BA_peak_8110 |
| DS126780 | 179 | 1172 | BA_peak_8111 |
| DS126792 | 371 | 1268 | BA_peak_8112 |
| DS126797 | 704 | 1219 | BA_peak_8113 |
| DS126814 | 486 | 1330 | BA_peak_8114 |
| DS126848 | 236 | 1366 | BA_peak_8115 |
| DS126871 | 168 | 540 | BA_peak_8117 |
| DS126896 | 632 | 964 | BA_peak_8118 |
| DS126904 | 86 | 1364 | BA_peak_8119 |
| DS126929 | 839 | 1164 | BA_peak_8120 |
| DS126932 | 700 | 1338 | BA_peak_8121 |
| DS126934 | 988 | 1324 | BA_peak_8122 |
| DS126941 | 530 | 939 | BA_peak_8123 |
| DS126991 | 88 | 1316 | BA_peak_8124 |
| DS126999 | 296 | 643 | BA_peak_8125 |
| DS127018 | 226 | 707 | BA_peak_8126 |
| DS127023 | 81 | 1235 | BA_peak_8127 |
| DS127025 | 46 | 484 | BA_peak_8128 |
| DS127050 | 272 | 662 | BA_peak_8129 |
| DS127057 | 70 | 492 | BA_peak_8130 |
| DS127059 | 295 | 1215 | BA_peak_8131 |
| DS127066 | 452 | 1304 | BA_peak_8132 |
| DS127103 | 827 | 1197 | BA_peak_8133 |
| DS127109 | 113 | 527 | BA_peak_8134 |
| DS127112 | 173 | 635 | BA_peak_8135 |
| DS127142 | 127 | 530 | BA_peak_8136 |
| DS127142 | 1014 | 1369 | BA_peak_8137 |
| DS127153 | 657 | 1289 | BA_peak_8138 |
| DS127177 | 768 | 1150 | BA_peak_8139 |
| DS127184 | 816 | 1200 | BA_peak_8140 |
| DS127197 | 31 | 1290 | BA_peak_8141 |
| DS127211 | 844 | 1306 | BA_peak_8142 |
| DS127223 | 886 | 1271 | BA_peak_8143 |
| DS127243 | 676 | 1206 | BA_peak_8144 |
| DS127273 | 742 | 1301 | BA_peak_8145 |
| DS127296 | 764 | 1210 | BA_peak_8146 |
| DS127302 | 75 | 1165 | BA_peak_8148 |
| DS127371 | 575 | 1355 | BA_peak_8150 |
| DS127390 | 43 | 685 | BA_peak_8151 |
| DS127404 | 131 | 488 | BA_peak_8152 |
| DS127419 | 590 | 1199 | BA_peak_8153 |
| DS127442 | 890 | 1263 | BA_peak_8155 |
| DS127468 | 233 | 771 | BA_peak_8156 |
| DS127472 | 761 | 1326 | BA_peak_8157 |
| DS127478 | 477 | 1044 | BA_peak_8158 |

| | | | |
|---|---|---|---|
| DS127508 | 774 | 1231 | BA_peak_8159 |
| DS127510 | 873 | 1221 | BA_peak_8160 |
| DS127559 | 476 | 886 | BA_peak_8161 |
| DS127563 | 223 | 1319 | BA_peak_8162 |
| DS127594 | 917 | 1238 | BA_peak_8164 |
| DS127603 | 146 | 664 | BA_peak_8165 |
| DS127604 | 571 | 896 | BA_peak_8166 |
| DS127606 | 324 | 1273 | BA_peak_8167 |
| DS127611 | 40 | 927 | BA_peak_8168 |
| DS127644 | 920 | 1329 | BA_peak_8169 |
| DS127645 | 856 | 1333 | BA_peak_8170 |
| DS127659 | 474 | 819 | BA_peak_8171 |
| DS127663 | 108 | 558 | BA_peak_8172 |
| DS127671 | 85 | 1182 | BA_peak_8173 |
| DS127683 | 849 | 1172 | BA_peak_8174 |
| DS127688 | 8 | 578 | BA_peak_8175 |
| DS127697 | 198 | 883 | BA_peak_8176 |
| DS127710 | 902 | 1231 | BA_peak_8177 |
| DS127719 | 51 | 1198 | BA_peak_8179 |
| DS127741 | 196 | 688 | BA_peak_8180 |
| DS127746 | 368 | 1322 | BA_peak_8181 |
| DS127794 | 18 | 1316 | BA_peak_8183 |
| DS127798 | 172 | 1211 | BA_peak_8184 |
| DS127803 | 742 | 1246 | BA_peak_8185 |
| DS127808 | 253 | 1203 | BA_peak_8186 |
| DS127822 | 68 | 534 | BA_peak_8188 |
| DS127822 | 925 | 1350 | BA_peak_8189 |
| DS127888 | 134 | 1215 | BA_peak_8190 |
| DS127892 | 906 | 1279 | BA_peak_8191 |
| DS127896 | 131 | 498 | BA_peak_8192 |
| DS127924 | 763 | 1347 | BA_peak_8193 |
| DS127929 | 535 | 956 | BA_peak_8194 |
| DS127935 | 84 | 854 | BA_peak_8195 |
| DS127936 | 117 | 1219 | BA_peak_8196 |
| DS127945 | 743 | 1302 | BA_peak_8197 |
| DS127960 | 865 | 1197 | BA_peak_8198 |
| DS127974 | 790 | 1146 | BA_peak_8199 |
| DS127975 | 767 | 1179 | BA_peak_8200 |
| DS127981 | 102 | 462 | BA_peak_8201 |
| DS127981 | 951 | 1313 | BA_peak_8202 |
| DS127994 | 376 | 1329 | BA_peak_8203 |
| DS128007 | 97 | 599 | BA_peak_8205 |
| DS128021 | 221 | 603 | BA_peak_8206 |
| DS128045 | 148 | 466 | BA_peak_8207 |
| DS128045 | 828 | 1176 | BA_peak_8208 |
| DS128088 | 95 | 759 | BA_peak_8209 |
| DS128089 | 52 | 950 | BA_peak_8210 |

| | | | |
|---|---|---|---|
| DS128093 | 218 | 1323 | BA_peak_8211 |
| DS128108 | 331 | 665 | BA_peak_8212 |
| DS128180 | 901 | 1286 | BA_peak_8214 |
| DS128190 | 229 | 801 | BA_peak_8215 |
| DS128191 | 43 | 1181 | BA_peak_8216 |
| DS128207 | 73 | 423 | BA_peak_8217 |
| DS128211 | 149 | 1269 | BA_peak_8218 |
| DS128230 | 113 | 1036 | BA_peak_8219 |
| DS128235 | 925 | 1270 | BA_peak_8221 |
| DS128241 | 57 | 1247 | BA_peak_8222 |
| DS128258 | 809 | 1295 | BA_peak_8223 |
| DS128269 | 242 | 576 | BA_peak_8224 |
| DS128283 | 27 | 404 | BA_peak_8225 |
| DS128285 | 119 | 487 | BA_peak_8226 |
| DS128297 | 17 | 1235 | BA_peak_8227 |
| DS128301 | 28 | 495 | BA_peak_8228 |
| DS128309 | 380 | 693 | BA_peak_8229 |
| DS128350 | 51 | 727 | BA_peak_8232 |
| DS128353 | 60 | 1297 | BA_peak_8233 |
| DS128356 | 113 | 1198 | BA_peak_8234 |
| DS128367 | 15 | 437 | BA_peak_8237 |
| DS128389 | 241 | 591 | BA_peak_8238 |
| DS128419 | 31 | 826 | BA_peak_8239 |
| DS128445 | 232 | 607 | BA_peak_8240 |
| DS128454 | 365 | 779 | BA_peak_8241 |
| DS128471 | 204 | 561 | BA_peak_8242 |
| DS128478 | 46 | 838 | BA_peak_8243 |
| DS128483 | 140 | 959 | BA_peak_8244 |
| DS128492 | 338 | 1290 | BA_peak_8245 |
| DS128503 | 444 | 1278 | BA_peak_8246 |
| DS128504 | 100 | 1154 | BA_peak_8247 |
| DS128517 | 756 | 1142 | BA_peak_8248 |
| DS128583 | 950 | 1328 | BA_peak_8249 |
| DS128591 | 436 | 1233 | BA_peak_8250 |
| DS128595 | 701 | 1235 | BA_peak_8251 |
| DS128603 | 75 | 500 | BA_peak_8252 |
| DS128616 | 48 | 1318 | BA_peak_8253 |
| DS128617 | 140 | 1154 | BA_peak_8254 |
| DS128657 | 75 | 624 | BA_peak_8255 |
| DS128679 | 271 | 685 | BA_peak_8256 |
| DS128681 | 661 | 1001 | BA_peak_8257 |
| DS128705 | 741 | 1229 | BA_peak_8258 |
| DS128713 | 153 | 1164 | BA_peak_8259 |
| DS128725 | 10 | 414 | BA_peak_8260 |
| DS128757 | 26 | 802 | BA_peak_8261 |
| DS128772 | 775 | 1161 | BA_peak_8262 |
| DS128781 | 247 | 680 | BA_peak_8264 |

| | | | |
|---|---|---|---|
| DS128796 | 686 | 1023 | BA_peak_8266 |
| DS128797 | 650 | 1250 | BA_peak_8267 |
| DS128827 | 64 | 1093 | BA_peak_8268 |
| DS128828 | 455 | 1205 | BA_peak_8269 |
| DS128850 | 654 | 1044 | BA_peak_8270 |
| DS128876 | 171 | 1053 | BA_peak_8271 |
| DS128891 | 856 | 1228 | BA_peak_8272 |
| DS128896 | 651 | 1101 | BA_peak_8273 |
| DS128910 | 560 | 1180 | BA_peak_8274 |
| DS128930 | 851 | 1244 | BA_peak_8275 |
| DS128958 | 276 | 873 | BA_peak_8276 |
| DS128960 | 830 | 1139 | BA_peak_8277 |
| DS128979 | 128 | 431 | BA_peak_8279 |
| DS128990 | 183 | 1253 | BA_peak_8280 |
| DS129005 | 965 | 1303 | BA_peak_8282 |
| DS129041 | 195 | 546 | BA_peak_8283 |
| DS129050 | 384 | 1094 | BA_peak_8284 |
| DS129079 | 133 | 1147 | BA_peak_8286 |
| DS129080 | 172 | 1275 | BA_peak_8287 |
| DS129083 | 219 | 830 | BA_peak_8288 |
| DS129084 | 882 | 1194 | BA_peak_8289 |
| DS129118 | 60 | 875 | BA_peak_8290 |
| DS129163 | 897 | 1221 | BA_peak_8291 |
| DS129170 | 9 | 431 | BA_peak_8293 |
| DS129181 | 122 | 862 | BA_peak_8294 |
| DS129201 | 332 | 917 | BA_peak_8295 |
| DS129205 | 108 | 939 | BA_peak_8296 |
| DS129215 | 762 | 1103 | BA_peak_8297 |
| DS129266 | 249 | 658 | BA_peak_8299 |
| DS129276 | 597 | 1284 | BA_peak_8300 |
| DS129279 | 77 | 1267 | BA_peak_8301 |
| DS129291 | 536 | 1136 | BA_peak_8302 |
| DS129314 | 333 | 941 | BA_peak_8303 |
| DS129324 | 825 | 1115 | BA_peak_8304 |
| DS129393 | 718 | 1265 | BA_peak_8307 |
| DS129417 | 846 | 1155 | BA_peak_8308 |
| DS129445 | 511 | 932 | BA_peak_8309 |
| DS129451 | 117 | 861 | BA_peak_8310 |
| DS129458 | 604 | 1193 | BA_peak_8311 |
| DS129462 | 899 | 1218 | BA_peak_8312 |
| DS129494 | 217 | 535 | BA_peak_8314 |
| DS129496 | 619 | 1276 | BA_peak_8315 |
| DS129507 | 64 | 544 | BA_peak_8316 |
| DS129532 | 163 | 454 | BA_peak_8318 |
| DS129564 | 789 | 1259 | BA_peak_8319 |
| DS129565 | 18 | 1124 | BA_peak_8320 |
| DS129583 | 670 | 1227 | BA_peak_8321 |

| | | | |
|---|---|---|---|
| DS129603 | 821 | 1231 | BA_peak_8323 |
| DS129613 | 34 | 513 | BA_peak_8324 |
| DS129613 | 845 | 1243 | BA_peak_8325 |
| DS129617 | 20 | 369 | BA_peak_8326 |
| DS129617 | 799 | 1201 | BA_peak_8327 |
| DS129623 | 667 | 1090 | BA_peak_8328 |
| DS129654 | 91 | 657 | BA_peak_8329 |
| DS129663 | 587 | 1127 | BA_peak_8330 |
| DS129665 | 739 | 1234 | BA_peak_8331 |
| DS129677 | 923 | 1287 | BA_peak_8332 |
| DS129694 | 39 | 1209 | BA_peak_8333 |
| DS129729 | 485 | 1197 | BA_peak_8334 |
| DS129731 | 244 | 814 | BA_peak_8335 |
| DS129745 | 58 | 635 | BA_peak_8336 |
| DS129748 | 485 | 1028 | BA_peak_8337 |
| DS129749 | 19 | 639 | BA_peak_8338 |
| DS129766 | 22 | 1219 | BA_peak_8339 |
| DS129767 | 87 | 463 | BA_peak_8340 |
| DS129804 | 872 | 1281 | BA_peak_8341 |
| DS129805 | 314 | 633 | BA_peak_8342 |
| DS129817 | 155 | 564 | BA_peak_8343 |
| DS129821 | 519 | 1239 | BA_peak_8344 |
| DS129856 | 316 | 1235 | BA_peak_8345 |
| DS129880 | 230 | 903 | BA_peak_8346 |
| DS129889 | 648 | 1171 | BA_peak_8347 |
| DS129895 | 359 | 966 | BA_peak_8348 |
| DS129896 | 509 | 1257 | BA_peak_8349 |
| DS129906 | 569 | 1192 | BA_peak_8350 |
| DS129917 | 451 | 876 | BA_peak_8351 |
| DS129942 | 43 | 407 | BA_peak_8353 |
| DS129947 | 813 | 1208 | BA_peak_8354 |
| DS129950 | 491 | 1190 | BA_peak_8355 |
| DS129959 | 630 | 1272 | BA_peak_8356 |
| DS129988 | 233 | 982 | BA_peak_8357 |
| DS130016 | 527 | 1256 | BA_peak_8358 |
| DS130045 | 336 | 832 | BA_peak_8359 |
| DS130048 | 551 | 883 | BA_peak_8360 |
| DS130050 | 76 | 1213 | BA_peak_8361 |
| DS130068 | 65 | 370 | BA_peak_8362 |
| DS130094 | 14 | 992 | BA_peak_8363 |
| DS130102 | 316 | 688 | BA_peak_8364 |
| DS130122 | 171 | 1054 | BA_peak_8365 |
| DS130133 | 29 | 1198 | BA_peak_8366 |
| DS130135 | 194 | 506 | BA_peak_8367 |
| DS130142 | 845 | 1180 | BA_peak_8368 |
| DS130146 | 85 | 1204 | BA_peak_8369 |
| DS130161 | 107 | 498 | BA_peak_8370 |

| | | | |
|---|---|---|---|
| DS130161 | 847 | 1202 | BA_peak_8371 |
| DS130162 | 408 | 997 | BA_peak_8372 |
| DS130166 | 190 | 596 | BA_peak_8373 |
| DS130201 | 138 | 1091 | BA_peak_8374 |
| DS130207 | 193 | 1253 | BA_peak_8375 |
| DS130217 | 799 | 1135 | BA_peak_8376 |
| DS130221 | 87 | 672 | BA_peak_8377 |
| DS130232 | 178 | 609 | BA_peak_8378 |
| DS130241 | 643 | 1257 | BA_peak_8379 |
| DS130281 | 274 | 695 | BA_peak_8382 |
| DS130301 | 68 | 1119 | BA_peak_8383 |
| DS130304 | 131 | 1002 | BA_peak_8384 |
| DS130399 | 27 | 600 | BA_peak_8386 |
| DS130403 | 154 | 515 | BA_peak_8387 |
| DS130410 | 643 | 1098 | BA_peak_8388 |
| DS130417 | 773 | 1091 | BA_peak_8390 |
| DS130479 | 34 | 1074 | BA_peak_8391 |
| DS130488 | 362 | 1136 | BA_peak_8392 |
| DS130494 | 69 | 453 | BA_peak_8393 |
| DS130496 | 65 | 480 | BA_peak_8394 |
| DS130544 | 735 | 1078 | BA_peak_8395 |
| DS130549 | 149 | 958 | BA_peak_8396 |
| DS130590 | 152 | 781 | BA_peak_8397 |
| DS130601 | 726 | 1120 | BA_peak_8398 |
| DS130602 | 61 | 418 | BA_peak_8399 |
| DS130608 | 52 | 417 | BA_peak_8400 |
| DS130615 | 747 | 1242 | BA_peak_8401 |
| DS130625 | 60 | 430 | BA_peak_8403 |
| DS130631 | 315 | 1154 | BA_peak_8404 |
| DS130635 | 44 | 457 | BA_peak_8405 |
| DS130640 | 146 | 1147 | BA_peak_8406 |
| DS130643 | 830 | 1174 | BA_peak_8407 |
| DS130669 | 529 | 1137 | BA_peak_8408 |
| DS130686 | 172 | 1222 | BA_peak_8409 |
| DS130702 | 44 | 449 | BA_peak_8410 |
| DS130713 | 769 | 1264 | BA_peak_8411 |
| DS130742 | 18 | 1185 | BA_peak_8413 |
| DS130801 | 828 | 1255 | BA_peak_8415 |
| DS130803 | 550 | 1252 | BA_peak_8416 |
| DS130814 | 476 | 1152 | BA_peak_8417 |
| DS130831 | 650 | 1194 | BA_peak_8418 |
| DS130855 | 198 | 506 | BA_peak_8419 |
| DS130873 | 108 | 539 | BA_peak_8420 |
| DS130876 | 525 | 1169 | BA_peak_8421 |
| DS130882 | 133 | 611 | BA_peak_8422 |
| DS130892 | 496 | 888 | BA_peak_8423 |
| DS130914 | 92 | 442 | BA_peak_8424 |

| | | | |
|---|---|---|---|
| DS130943 | 111 | 1199 | BA_peak_8425 |
| DS130945 | 882 | 1224 | BA_peak_8426 |
| DS130963 | 654 | 1181 | BA_peak_8427 |
| DS130964 | 70 | 1118 | BA_peak_8428 |
| DS130984 | 700 | 1051 | BA_peak_8429 |
| DS130991 | 51 | 550 | BA_peak_8430 |
| DS131017 | 14 | 422 | BA_peak_8431 |
| DS131043 | 696 | 1217 | BA_peak_8433 |
| DS131050 | 196 | 1237 | BA_peak_8434 |
| DS131077 | 552 | 1097 | BA_peak_8435 |
| DS131078 | 822 | 1193 | BA_peak_8436 |
| DS131079 | 59 | 484 | BA_peak_8437 |
| DS131128 | 690 | 1065 | BA_peak_8438 |
| DS131191 | 27 | 359 | BA_peak_8439 |
| DS131199 | 42 | 387 | BA_peak_8440 |
| DS131210 | 621 | 1182 | BA_peak_8441 |
| DS131213 | 825 | 1232 | BA_peak_8442 |
| DS131225 | 47 | 664 | BA_peak_8443 |
| DS131282 | 462 | 890 | BA_peak_8444 |
| DS131291 | 122 | 1211 | BA_peak_8445 |
| DS131337 | 353 | 691 | BA_peak_8447 |
| DS131384 | 107 | 923 | BA_peak_8448 |
| DS131417 | 751 | 1232 | BA_peak_8449 |
| DS131421 | 172 | 1033 | BA_peak_8450 |
| DS131422 | 50 | 481 | BA_peak_8451 |
| DS131425 | 31 | 557 | BA_peak_8452 |
| DS131446 | 795 | 1174 | BA_peak_8453 |
| DS131458 | 59 | 1232 | BA_peak_8454 |
| DS131473 | 270 | 1189 | BA_peak_8455 |
| DS131487 | 731 | 1202 | BA_peak_8456 |
| DS131499 | 312 | 963 | BA_peak_8457 |
| DS131504 | 746 | 1203 | BA_peak_8458 |
| DS131575 | 74 | 572 | BA_peak_8459 |
| DS131605 | 466 | 1005 | BA_peak_8460 |
| DS131624 | 131 | 1119 | BA_peak_8462 |
| DS131628 | 659 | 1123 | BA_peak_8463 |
| DS131629 | 279 | 591 | BA_peak_8464 |
| DS131643 | 635 | 930 | BA_peak_8465 |
| DS131671 | 114 | 546 | BA_peak_8467 |
| DS131682 | 865 | 1212 | BA_peak_8469 |
| DS131687 | 95 | 582 | BA_peak_8470 |
| DS131745 | 557 | 1165 | BA_peak_8471 |
| DS131770 | 8 | 439 | BA_peak_8472 |
| DS131775 | 194 | 737 | BA_peak_8473 |
| DS131786 | 40 | 437 | BA_peak_8474 |
| DS131789 | 243 | 1069 | BA_peak_8475 |
| DS131839 | 52 | 956 | BA_peak_8476 |

| | | | |
|---|---|---|---|
| DS131880 | 148 | 1060 | BA_peak_8477 |
| DS131890 | 470 | 868 | BA_peak_8478 |
| DS131903 | 214 | 1157 | BA_peak_8480 |
| DS131905 | 23 | 406 | BA_peak_8481 |
| DS131919 | 145 | 615 | BA_peak_8482 |
| DS131920 | 209 | 541 | BA_peak_8483 |
| DS131961 | 32 | 408 | BA_peak_8484 |
| DS131970 | 161 | 1210 | BA_peak_8485 |
| DS131977 | 258 | 1225 | BA_peak_8486 |
| DS131982 | 21 | 448 | BA_peak_8487 |
| DS131988 | 254 | 1037 | BA_peak_8488 |
| DS131991 | 188 | 1061 | BA_peak_8489 |
| DS132006 | 246 | 1192 | BA_peak_8490 |
| DS132016 | 547 | 969 | BA_peak_8491 |
| DS132017 | 336 | 610 | BA_peak_8492 |
| DS132024 | 19 | 1037 | BA_peak_8493 |
| DS132038 | 44 | 418 | BA_peak_8494 |
| DS132043 | 29 | 1189 | BA_peak_8495 |
| DS132098 | 34 | 1176 | BA_peak_8496 |
| DS132130 | 114 | 687 | BA_peak_8498 |
| DS132131 | 155 | 543 | BA_peak_8499 |
| DS132150 | 198 | 916 | BA_peak_8500 |
| DS132161 | 520 | 1150 | BA_peak_8501 |
| DS132165 | 27 | 1209 | BA_peak_8502 |
| DS132186 | 596 | 1173 | BA_peak_8503 |
| DS132193 | 65 | 446 | BA_peak_8504 |
| DS132255 | 207 | 541 | BA_peak_8505 |
| DS132269 | 109 | 558 | BA_peak_8506 |
| DS132285 | 734 | 1116 | BA_peak_8507 |
| DS132299 | 387 | 998 | BA_peak_8508 |
| DS132311 | 311 | 1152 | BA_peak_8509 |
| DS132324 | 468 | 1152 | BA_peak_8510 |
| DS132330 | 782 | 1177 | BA_peak_8511 |
| DS132340 | 101 | 582 | BA_peak_8512 |
| DS132360 | 35 | 671 | BA_peak_8513 |
| DS132415 | 704 | 1017 | BA_peak_8514 |
| DS132416 | 44 | 348 | BA_peak_8515 |
| DS132425 | 123 | 431 | BA_peak_8516 |
| DS132463 | 61 | 1005 | BA_peak_8519 |
| DS132472 | 472 | 1193 | BA_peak_8521 |
| DS132489 | 174 | 1048 | BA_peak_8522 |
| DS132501 | 45 | 486 | BA_peak_8523 |
| DS132512 | 675 | 1101 | BA_peak_8524 |
| DS132514 | 151 | 579 | BA_peak_8525 |
| DS132567 | 432 | 1033 | BA_peak_8526 |
| DS132568 | 658 | 1071 | BA_peak_8527 |
| DS132603 | 807 | 1117 | BA_peak_8528 |

| | | | |
|---|---|---|---|
| DS132628 | 56 | 1125 | BA_peak_8529 |
| DS132642 | 660 | 1045 | BA_peak_8530 |
| DS132654 | 432 | 914 | BA_peak_8532 |
| DS132684 | 181 | 1159 | BA_peak_8533 |
| DS132723 | 67 | 596 | BA_peak_8534 |
| DS132732 | 50 | 409 | BA_peak_8535 |
| DS132742 | 91 | 454 | BA_peak_8536 |
| DS132756 | 496 | 1177 | BA_peak_8537 |
| DS132764 | 745 | 1171 | BA_peak_8538 |
| DS132795 | 133 | 524 | BA_peak_8539 |
| DS132822 | 312 | 1049 | BA_peak_8541 |
| DS132834 | 99 | 569 | BA_peak_8542 |
| DS132857 | 357 | 1024 | BA_peak_8543 |
| DS132898 | 47 | 378 | BA_peak_8545 |
| DS132902 | 138 | 976 | BA_peak_8546 |
| DS132933 | 230 | 872 | BA_peak_8547 |
| DS132936 | 52 | 1168 | BA_peak_8548 |
| DS132937 | 160 | 603 | BA_peak_8549 |
| DS132951 | 234 | 1202 | BA_peak_8550 |
| DS132952 | 255 | 728 | BA_peak_8551 |
| DS132957 | 672 | 1167 | BA_peak_8552 |
| DS132962 | 286 | 1123 | BA_peak_8553 |
| DS132986 | 469 | 922 | BA_peak_8554 |
| DS133012 | 164 | 1199 | BA_peak_8555 |
| DS133026 | 185 | 596 | BA_peak_8556 |
| DS133031 | 13 | 696 | BA_peak_8557 |
| DS133051 | 702 | 1156 | BA_peak_8558 |
| DS133083 | 115 | 474 | BA_peak_8559 |
| DS133097 | 192 | 922 | BA_peak_8560 |
| DS133121 | 139 | 512 | BA_peak_8561 |
| DS133126 | 433 | 1135 | BA_peak_8562 |
| DS133158 | 425 | 1040 | BA_peak_8563 |
| DS133168 | 92 | 1158 | BA_peak_8564 |
| DS133185 | 86 | 678 | BA_peak_8565 |
| DS133196 | 63 | 878 | BA_peak_8566 |
| DS133215 | 80 | 495 | BA_peak_8567 |
| DS133229 | 84 | 1052 | BA_peak_8568 |
| DS133243 | 143 | 901 | BA_peak_8569 |
| DS133250 | 739 | 1129 | BA_peak_8570 |
| DS133278 | 244 | 1003 | BA_peak_8572 |
| DS133297 | 531 | 884 | BA_peak_8573 |
| DS133303 | 71 | 428 | BA_peak_8574 |
| DS133326 | 734 | 1092 | BA_peak_8575 |
| DS133329 | 72 | 546 | BA_peak_8576 |
| DS133337 | 168 | 424 | BA_peak_8577 |
| DS133347 | 394 | 690 | BA_peak_8578 |
| DS133375 | 110 | 943 | BA_peak_8580 |

| | | | |
|---|---|---|---|
| DS133377 | 253 | 1095 | BA_peak_8581 |
| DS133404 | 53 | 1008 | BA_peak_8582 |
| DS133406 | 35 | 513 | BA_peak_8583 |
| DS133442 | 150 | 683 | BA_peak_8584 |
| DS133444 | 419 | 940 | BA_peak_8585 |
| DS133462 | 507 | 1165 | BA_peak_8586 |
| DS133470 | 84 | 533 | BA_peak_8587 |
| DS133501 | 618 | 1080 | BA_peak_8588 |
| DS133509 | 672 | 1148 | BA_peak_8589 |
| DS133521 | 123 | 489 | BA_peak_8590 |
| DS133528 | 190 | 1113 | BA_peak_8591 |
| DS133530 | 330 | 1082 | BA_peak_8592 |
| DS133551 | 243 | 933 | BA_peak_8593 |
| DS133596 | 171 | 489 | BA_peak_8594 |
| DS133616 | 161 | 1162 | BA_peak_8595 |
| DS133624 | 761 | 1115 | BA_peak_8596 |
| DS133638 | 267 | 1134 | BA_peak_8597 |
| DS133711 | 237 | 1013 | BA_peak_8598 |
| DS133715 | 131 | 571 | BA_peak_8599 |
| DS133742 | 296 | 1128 | BA_peak_8600 |
| DS133768 | 154 | 478 | BA_peak_8601 |
| DS133771 | 216 | 1135 | BA_peak_8602 |
| DS133774 | 186 | 509 | BA_peak_8603 |
| DS133804 | 45 | 1170 | BA_peak_8604 |
| DS133881 | 547 | 951 | BA_peak_8606 |
| DS133894 | 666 | 1135 | BA_peak_8607 |
| DS133949 | 82 | 633 | BA_peak_8609 |
| DS133957 | 231 | 654 | BA_peak_8610 |
| DS134040 | 104 | 655 | BA_peak_8613 |
| DS134074 | 276 | 1118 | BA_peak_8614 |
| DS134093 | 22 | 372 | BA_peak_8615 |
| DS134100 | 24 | 588 | BA_peak_8616 |
| DS134135 | 304 | 874 | BA_peak_8617 |
| DS134140 | 750 | 1173 | BA_peak_8618 |
| DS134181 | 138 | 958 | BA_peak_8619 |
| DS134182 | 601 | 1057 | BA_peak_8620 |
| DS134191 | 128 | 1038 | BA_peak_8621 |
| DS134224 | 208 | 698 | BA_peak_8623 |
| DS134231 | 398 | 1144 | BA_peak_8624 |
| DS134244 | 513 | 1036 | BA_peak_8625 |
| DS134249 | 44 | 520 | BA_peak_8626 |
| DS134270 | 139 | 528 | BA_peak_8628 |
| DS134316 | 183 | 506 | BA_peak_8629 |
| DS134320 | 199 | 546 | BA_peak_8630 |
| DS134355 | 594 | 1077 | BA_peak_8631 |
| DS134359 | 182 | 1143 | BA_peak_8632 |
| DS134369 | 146 | 1176 | BA_peak_8633 |

| | | | |
|---|---|---|---|
| DS134372 | 685 | 1096 | BA_peak_8634 |
| DS134401 | 544 | 1044 | BA_peak_8635 |
| DS134421 | 26 | 1182 | BA_peak_8636 |
| DS134423 | 626 | 1130 | BA_peak_8637 |
| DS134449 | 62 | 940 | BA_peak_8638 |
| DS134483 | 689 | 1156 | BA_peak_8639 |
| DS134532 | 44 | 557 | BA_peak_8642 |
| DS134540 | 163 | 515 | BA_peak_8643 |
| DS134560 | 232 | 760 | BA_peak_8644 |
| DS134576 | 232 | 762 | BA_peak_8645 |
| DS134584 | 109 | 1108 | BA_peak_8646 |
| DS134592 | 330 | 1106 | BA_peak_8647 |
| DS134595 | 379 | 719 | BA_peak_8648 |
| DS134606 | 407 | 828 | BA_peak_8649 |
| DS134612 | 166 | 976 | BA_peak_8650 |
| DS134631 | 76 | 1127 | BA_peak_8651 |
| DS134654 | 101 | 1105 | BA_peak_8652 |
| DS134655 | 295 | 1103 | BA_peak_8653 |
| DS134660 | 51 | 395 | BA_peak_8654 |
| DS134664 | 647 | 1145 | BA_peak_8655 |
| DS134666 | 425 | 1137 | BA_peak_8656 |
| DS134690 | 407 | 1060 | BA_peak_8657 |
| DS134699 | 701 | 1021 | BA_peak_8658 |
| DS134712 | 163 | 578 | BA_peak_8659 |
| DS134747 | 75 | 702 | BA_peak_8660 |
| DS134748 | 381 | 911 | BA_peak_8661 |
| DS134781 | 693 | 1105 | BA_peak_8662 |
| DS134801 | 646 | 1048 | BA_peak_8664 |
| DS134804 | 106 | 571 | BA_peak_8665 |
| DS134809 | 457 | 857 | BA_peak_8666 |
| DS134832 | 144 | 523 | BA_peak_8667 |
| DS134849 | 30 | 460 | BA_peak_8669 |
| DS134866 | 371 | 679 | BA_peak_8670 |
| DS134867 | 176 | 804 | BA_peak_8671 |
| DS134941 | 219 | 977 | BA_peak_8672 |
| DS134943 | 794 | 1141 | BA_peak_8673 |
| DS134974 | 197 | 551 | BA_peak_8675 |
| DS134977 | 230 | 752 | BA_peak_8676 |
| DS134988 | 752 | 1061 | BA_peak_8677 |
| DS135006 | 32 | 1075 | BA_peak_8678 |
| DS135016 | 146 | 491 | BA_peak_8679 |
| DS135051 | 147 | 1029 | BA_peak_8680 |
| DS135052 | 212 | 893 | BA_peak_8681 |
| DS135079 | 175 | 784 | BA_peak_8682 |
| DS135165 | 20 | 571 | BA_peak_8684 |
| DS135166 | 80 | 762 | BA_peak_8685 |
| DS135174 | 61 | 714 | BA_peak_8686 |

| | | | |
|---|---|---|---|
| DS135198 | 279 | 790 | BA_peak_8688 |
| DS135222 | 179 | 617 | BA_peak_8689 |
| DS135265 | 41 | 1134 | BA_peak_8690 |
| DS135297 | 441 | 993 | BA_peak_8691 |
| DS135332 | 477 | 1134 | BA_peak_8692 |
| DS135335 | 130 | 732 | BA_peak_8693 |
| DS135337 | 604 | 916 | BA_peak_8694 |
| DS135354 | 124 | 499 | BA_peak_8695 |
| DS135355 | 43 | 1127 | BA_peak_8696 |
| DS135364 | 319 | 979 | BA_peak_8697 |
| DS135372 | 64 | 431 | BA_peak_8698 |
| DS135385 | 364 | 1086 | BA_peak_8699 |
| DS135412 | 547 | 869 | BA_peak_8700 |
| DS135416 | 263 | 764 | BA_peak_8701 |
| DS135423 | 142 | 691 | BA_peak_8702 |
| DS135435 | 662 | 975 | BA_peak_8703 |
| DS135453 | 459 | 908 | BA_peak_8704 |
| DS135460 | 16 | 390 | BA_peak_8705 |
| DS135483 | 741 | 1080 | BA_peak_8706 |
| DS135519 | 157 | 568 | BA_peak_8707 |
| DS135560 | 107 | 887 | BA_peak_8708 |
| DS135580 | 447 | 1014 | BA_peak_8709 |
| DS135586 | 673 | 1127 | BA_peak_8710 |
| DS135599 | 577 | 1075 | BA_peak_8712 |
| DS135606 | 114 | 965 | BA_peak_8713 |
| DS135611 | 148 | 764 | BA_peak_8714 |
| DS135642 | 125 | 543 | BA_peak_8715 |
| DS135652 | 408 | 742 | BA_peak_8716 |
| DS135662 | 248 | 762 | BA_peak_8717 |
| DS135702 | 108 | 606 | BA_peak_8718 |
| DS135717 | 19 | 886 | BA_peak_8719 |
| DS135723 | 408 | 813 | BA_peak_8720 |
| DS135741 | 62 | 552 | BA_peak_8721 |
| DS135755 | 304 | 639 | BA_peak_8722 |
| DS135761 | 651 | 1078 | BA_peak_8723 |
| DS135786 | 635 | 1132 | BA_peak_8724 |
| DS135804 | 716 | 1068 | BA_peak_8725 |
| DS135812 | 608 | 1039 | BA_peak_8726 |
| DS135837 | 116 | 1085 | BA_peak_8727 |
| DS135838 | 141 | 466 | BA_peak_8728 |
| DS135840 | 104 | 458 | BA_peak_8729 |
| DS135876 | 41 | 370 | BA_peak_8730 |
| DS135881 | 320 | 820 | BA_peak_8731 |
| DS135889 | 275 | 745 | BA_peak_8732 |
| DS135934 | 40 | 1092 | BA_peak_8733 |
| DS135981 | 706 | 1114 | BA_peak_8734 |
| DS135995 | 288 | 1021 | BA_peak_8735 |

| | | | |
|---|---|---|---|
| DS135996 | 161 | 1026 | BA_peak_8736 |
| DS136013 | 124 | 1086 | BA_peak_8737 |
| DS136107 | 91 | 1118 | BA_peak_8738 |
| DS136146 | 621 | 1074 | BA_peak_8740 |
| DS136147 | 130 | 1120 | BA_peak_8741 |
| DS136176 | 527 | 1111 | BA_peak_8742 |
| DS136188 | 35 | 936 | BA_peak_8744 |
| DS136254 | 103 | 1101 | BA_peak_8745 |
| DS136283 | 504 | 1091 | BA_peak_8746 |
| DS136287 | 106 | 499 | BA_peak_8747 |
| DS136310 | 101 | 594 | BA_peak_8748 |
| DS136325 | 129 | 420 | BA_peak_8749 |
| DS136340 | 244 | 1030 | BA_peak_8750 |
| DS136345 | 384 | 895 | BA_peak_8751 |
| DS136353 | 676 | 992 | BA_peak_8752 |
| DS136389 | 293 | 748 | BA_peak_8753 |
| DS136392 | 287 | 897 | BA_peak_8754 |
| DS136414 | 365 | 1090 | BA_peak_8755 |
| DS136426 | 115 | 785 | BA_peak_8756 |
| DS136433 | 37 | 391 | BA_peak_8757 |
| DS136436 | 259 | 991 | BA_peak_8758 |
| DS136446 | 500 | 919 | BA_peak_8759 |
| DS136493 | 119 | 1015 | BA_peak_8761 |
| DS136494 | 130 | 720 | BA_peak_8762 |
| DS136499 | 251 | 1073 | BA_peak_8763 |
| DS136551 | 247 | 614 | BA_peak_8764 |
| DS136562 | 227 | 833 | BA_peak_8766 |
| DS136569 | 355 | 934 | BA_peak_8767 |
| DS136577 | 445 | 929 | BA_peak_8768 |
| DS136589 | 440 | 960 | BA_peak_8769 |
| DS136596 | 417 | 1078 | BA_peak_8770 |
| DS136605 | 124 | 459 | BA_peak_8771 |
| DS136646 | 164 | 878 | BA_peak_8772 |
| DS136669 | 690 | 1041 | BA_peak_8775 |
| DS136673 | 115 | 668 | BA_peak_8776 |
| DS136693 | 141 | 623 | BA_peak_8777 |
| DS136704 | 585 | 1127 | BA_peak_8778 |
| DS136722 | 354 | 1106 | BA_peak_8779 |
| DS136762 | 317 | 1051 | BA_peak_8781 |
| DS136768 | 59 | 776 | BA_peak_8782 |
| DS136771 | 50 | 539 | BA_peak_8783 |
| DS136775 | 487 | 1063 | BA_peak_8784 |
| DS136781 | 309 | 1004 | BA_peak_8785 |
| DS136796 | 771 | 1100 | BA_peak_8786 |
| DS136818 | 17 | 376 | BA_peak_8787 |
| DS136836 | 22 | 1073 | BA_peak_8788 |
| DS136842 | 461 | 998 | BA_peak_8789 |

| | | | |
|---|---|---|---|
| DS136854 | 89 | 1024 | BA_peak_8790 |
| DS136864 | 202 | 783 | BA_peak_8791 |
| DS136872 | 134 | 1076 | BA_peak_8792 |
| DS136892 | 444 | 819 | BA_peak_8793 |
| DS136931 | 454 | 1017 | BA_peak_8796 |
| DS136934 | 210 | 930 | BA_peak_8797 |
| DS136987 | 727 | 1117 | BA_peak_8798 |
| DS136992 | 69 | 461 | BA_peak_8799 |
| DS137028 | 39 | 531 | BA_peak_8800 |
| DS137031 | 155 | 1089 | BA_peak_8801 |
| DS137033 | 441 | 844 | BA_peak_8802 |
| DS137035 | 366 | 942 | BA_peak_8803 |
| DS137052 | 500 | 831 | BA_peak_8804 |
| DS137071 | 9 | 602 | BA_peak_8805 |
| DS137080 | 330 | 1056 | BA_peak_8806 |
| DS137140 | 531 | 952 | BA_peak_8809 |
| DS137146 | 83 | 996 | BA_peak_8810 |
| DS137221 | 188 | 820 | BA_peak_8812 |
| DS137239 | 142 | 524 | BA_peak_8813 |
| DS137251 | 551 | 1065 | BA_peak_8814 |
| DS137258 | 236 | 789 | BA_peak_8815 |
| DS137272 | 713 | 1063 | BA_peak_8816 |
| DS137279 | 626 | 956 | BA_peak_8817 |
| DS137307 | 158 | 984 | BA_peak_8818 |
| DS137310 | 515 | 1103 | BA_peak_8819 |
| DS137314 | 66 | 827 | BA_peak_8820 |
| DS137352 | 55 | 1010 | BA_peak_8821 |
| DS137354 | 144 | 1062 | BA_peak_8822 |
| DS137355 | 21 | 1080 | BA_peak_8823 |
| DS137356 | 73 | 535 | BA_peak_8824 |
| DS137377 | 358 | 1033 | BA_peak_8825 |
| DS137384 | 83 | 1089 | BA_peak_8826 |
| DS137414 | 442 | 990 | BA_peak_8828 |
| DS137416 | 54 | 1023 | BA_peak_8829 |
| DS137417 | 166 | 634 | BA_peak_8830 |
| DS137447 | 39 | 603 | BA_peak_8831 |
| DS137453 | 574 | 1111 | BA_peak_8832 |
| DS137490 | 197 | 778 | BA_peak_8834 |
| DS137497 | 131 | 1080 | BA_peak_8835 |
| DS137542 | 134 | 593 | BA_peak_8837 |
| DS137547 | 481 | 912 | BA_peak_8838 |
| DS137560 | 40 | 509 | BA_peak_8839 |
| DS137589 | 467 | 876 | BA_peak_8841 |
| DS137617 | 16 | 1072 | BA_peak_8842 |
| DS137650 | 218 | 795 | BA_peak_8843 |
| DS137668 | 389 | 986 | BA_peak_8844 |
| DS137699 | 721 | 1033 | BA_peak_8846 |

| | | | |
|---|---|---|---|
| DS137707 | 291 | 769 | BA_peak_8847 |
| DS137714 | 191 | 544 | BA_peak_8848 |
| DS137719 | 89 | 898 | BA_peak_8849 |
| DS137739 | 294 | 779 | BA_peak_8850 |
| DS137764 | 86 | 649 | BA_peak_8851 |
| DS137784 | 137 | 678 | BA_peak_8852 |
| DS137802 | 572 | 881 | BA_peak_8853 |
| DS137803 | 611 | 1038 | BA_peak_8854 |
| DS137825 | 410 | 941 | BA_peak_8855 |
| DS137829 | 112 | 1060 | BA_peak_8856 |
| DS137832 | 17 | 1014 | BA_peak_8857 |
| DS137855 | 219 | 878 | BA_peak_8859 |
| DS137946 | 496 | 972 | BA_peak_8861 |
| DS137955 | 29 | 690 | BA_peak_8862 |
| DS137960 | 65 | 474 | BA_peak_8863 |
| DS137964 | 130 | 786 | BA_peak_8864 |
| DS138000 | 165 | 668 | BA_peak_8865 |
| DS138019 | 728 | 1075 | BA_peak_8866 |
| DS138030 | 81 | 518 | BA_peak_8867 |
| DS138079 | 42 | 786 | BA_peak_8868 |
| DS138125 | 97 | 826 | BA_peak_8869 |
| DS138139 | 57 | 722 | BA_peak_8870 |
| DS138148 | 120 | 541 | BA_peak_8871 |
| DS138159 | 26 | 1098 | BA_peak_8872 |
| DS138203 | 116 | 707 | BA_peak_8873 |
| DS138213 | 300 | 609 | BA_peak_8874 |
| DS138227 | 126 | 501 | BA_peak_8875 |
| DS138260 | 62 | 381 | BA_peak_8877 |
| DS138271 | 40 | 978 | BA_peak_8879 |
| DS138289 | 92 | 978 | BA_peak_8880 |
| DS138295 | 256 | 710 | BA_peak_8881 |
| DS138300 | 112 | 1002 | BA_peak_8882 |
| DS138305 | 353 | 761 | BA_peak_8883 |
| DS138312 | 530 | 1008 | BA_peak_8884 |
| DS138316 | 411 | 850 | BA_peak_8885 |
| DS138329 | 337 | 631 | BA_peak_8886 |
| DS138375 | 26 | 941 | BA_peak_8887 |
| DS138435 | 547 | 1032 | BA_peak_8888 |
| DS138467 | 641 | 1053 | BA_peak_8889 |
| DS138516 | 120 | 958 | BA_peak_8890 |
| DS138566 | 96 | 1055 | BA_peak_8892 |
| DS138570 | 37 | 547 | BA_peak_8893 |
| DS138577 | 133 | 1003 | BA_peak_8894 |
| DS138591 | 444 | 832 | BA_peak_8895 |
| DS138596 | 600 | 963 | BA_peak_8896 |
| DS138601 | 261 | 810 | BA_peak_8897 |
| DS138617 | 321 | 729 | BA_peak_8898 |

| | | | |
|---|---|---|---|
| DS138651 | 297 | 914 | BA_peak_8899 |
| DS138659 | 259 | 1026 | BA_peak_8900 |
| DS138670 | 303 | 780 | BA_peak_8901 |
| DS138717 | 97 | 1071 | BA_peak_8902 |
| DS138720 | 356 | 980 | BA_peak_8903 |
| DS138721 | 97 | 421 | BA_peak_8904 |
| DS138756 | 31 | 1095 | BA_peak_8906 |
| DS138767 | 41 | 673 | BA_peak_8907 |
| DS138777 | 24 | 394 | BA_peak_8908 |
| DS138780 | 515 | 992 | BA_peak_8909 |
| DS138793 | 424 | 996 | BA_peak_8910 |
| DS138803 | 62 | 1010 | BA_peak_8911 |
| DS138827 | 350 | 1073 | BA_peak_8913 |
| DS138854 | 148 | 1039 | BA_peak_8914 |
| DS138856 | 240 | 660 | BA_peak_8915 |
| DS138871 | 329 | 931 | BA_peak_8916 |
| DS138912 | 324 | 1018 | BA_peak_8917 |
| DS138932 | 153 | 983 | BA_peak_8918 |
| DS138944 | 108 | 877 | BA_peak_8919 |
| DS138959 | 346 | 883 | BA_peak_8920 |
| DS138975 | 42 | 703 | BA_peak_8921 |
| DS139031 | 66 | 861 | BA_peak_8922 |
| DS139032 | 366 | 707 | BA_peak_8923 |
| DS139074 | 84 | 780 | BA_peak_8925 |
| DS139084 | 267 | 634 | BA_peak_8926 |
| DS139129 | 116 | 1025 | BA_peak_8928 |
| DS139136 | 216 | 667 | BA_peak_8929 |
| DS139155 | 631 | 965 | BA_peak_8930 |
| DS139176 | 619 | 1069 | BA_peak_8931 |
| DS139243 | 247 | 907 | BA_peak_8932 |
| DS139262 | 432 | 1031 | BA_peak_8933 |
| DS139280 | 143 | 812 | BA_peak_8934 |
| DS139282 | 250 | 703 | BA_peak_8935 |
| DS139316 | 89 | 853 | BA_peak_8936 |
| DS139331 | 200 | 528 | BA_peak_8937 |
| DS139344 | 174 | 1042 | BA_peak_8939 |
| DS139376 | 400 | 831 | BA_peak_8940 |
| DS139413 | 128 | 1048 | BA_peak_8941 |
| DS139416 | 399 | 1046 | BA_peak_8942 |
| DS139425 | 62 | 345 | BA_peak_8943 |
| DS139428 | 203 | 785 | BA_peak_8944 |
| DS139462 | 110 | 636 | BA_peak_8945 |
| DS139466 | 20 | 957 | BA_peak_8946 |
| DS139475 | 69 | 938 | BA_peak_8947 |
| DS139580 | 545 | 896 | BA_peak_8949 |
| DS139586 | 132 | 695 | BA_peak_8950 |
| DS139624 | 65 | 610 | BA_peak_8952 |

| | | | |
|---|---|---|---|
| DS139659 | 115 | 985 | BA_peak_8955 |
| DS139674 | 257 | 607 | BA_peak_8956 |
| DS139722 | 179 | 1041 | BA_peak_8958 |
| DS139739 | 619 | 966 | BA_peak_8959 |
| DS139768 | 248 | 638 | BA_peak_8960 |
| DS139770 | 196 | 635 | BA_peak_8961 |
| DS139781 | 202 | 778 | BA_peak_8962 |
| DS139785 | 192 | 959 | BA_peak_8963 |
| DS139810 | 128 | 937 | BA_peak_8964 |
| DS139811 | 50 | 387 | BA_peak_8965 |
| DS139821 | 92 | 524 | BA_peak_8966 |
| DS139835 | 181 | 1069 | BA_peak_8967 |
| DS139877 | 184 | 939 | BA_peak_8969 |
| DS139906 | 26 | 1058 | BA_peak_8970 |
| DS139922 | 64 | 660 | BA_peak_8971 |
| DS139941 | 517 | 933 | BA_peak_8973 |
| DS139944 | 97 | 514 | BA_peak_8974 |
| DS139947 | 39 | 724 | BA_peak_8975 |
| DS139959 | 296 | 804 | BA_peak_8976 |
| DS139989 | 13 | 943 | BA_peak_8977 |
| DS140030 | 234 | 1030 | BA_peak_8979 |
| DS140032 | 25 | 483 | BA_peak_8980 |
| DS140039 | 60 | 1027 | BA_peak_8981 |
| DS140045 | 113 | 907 | BA_peak_8982 |
| DS140046 | 44 | 392 | BA_peak_8983 |
| DS140048 | 27 | 985 | BA_peak_8984 |
| DS140159 | 96 | 663 | BA_peak_8986 |
| DS140169 | 321 | 752 | BA_peak_8987 |
| DS140178 | 268 | 577 | BA_peak_8988 |
| DS140195 | 22 | 663 | BA_peak_8989 |
| DS140214 | 285 | 633 | BA_peak_8990 |
| DS140230 | 75 | 1046 | BA_peak_8991 |
| DS140258 | 173 | 1001 | BA_peak_8992 |
| DS140358 | 284 | 870 | BA_peak_8993 |
| DS140368 | 144 | 503 | BA_peak_8994 |
| DS140392 | 104 | 615 | BA_peak_8995 |
| DS140468 | 460 | 1012 | BA_peak_8997 |
| DS140492 | 377 | 935 | BA_peak_8998 |
| DS140504 | 55 | 455 | BA_peak_8999 |
| DS140508 | 34 | 765 | BA_peak_9000 |
| DS140518 | 199 | 695 | BA_peak_9001 |
| DS140558 | 72 | 883 | BA_peak_9002 |
| DS140567 | 191 | 959 | BA_peak_9003 |
| DS140575 | 284 | 877 | BA_peak_9004 |
| DS140580 | 342 | 905 | BA_peak_9005 |
| DS140581 | 229 | 839 | BA_peak_9006 |
| DS140591 | 30 | 681 | BA_peak_9007 |

| | | | |
|---|---|---|---|
| DS140620 | 145 | 682 | BA_peak_9008 |
| DS140622 | 179 | 918 | BA_peak_9009 |
| DS140635 | 511 | 879 | BA_peak_9010 |
| DS140670 | 70 | 851 | BA_peak_9011 |
| DS140735 | 245 | 682 | BA_peak_9013 |
| DS140742 | 217 | 1007 | BA_peak_9014 |
| DS140815 | 131 | 1055 | BA_peak_9015 |
| DS140824 | 211 | 637 | BA_peak_9016 |
| DS140838 | 225 | 999 | BA_peak_9017 |
| DS140891 | 119 | 508 | BA_peak_9018 |
| DS140902 | 66 | 1029 | BA_peak_9019 |
| DS140910 | 199 | 922 | BA_peak_9020 |
| DS140941 | 341 | 915 | BA_peak_9021 |
| DS140944 | 92 | 420 | BA_peak_9022 |
| DS140957 | 238 | 741 | BA_peak_9023 |
| DS140961 | 213 | 874 | BA_peak_9024 |
| DS140965 | 4 | 371 | BA_peak_9026 |
| DS140967 | 80 | 921 | BA_peak_9027 |
| DS141005 | 275 | 822 | BA_peak_9028 |
| DS141033 | 220 | 767 | BA_peak_9029 |
| DS141077 | 560 | 1054 | BA_peak_9032 |
| DS141078 | 470 | 948 | BA_peak_9033 |
| DS141091 | 144 | 799 | BA_peak_9034 |
| DS141100 | 525 | 1055 | BA_peak_9035 |
| DS141125 | 105 | 1051 | BA_peak_9036 |
| DS141126 | 618 | 1013 | BA_peak_9037 |
| DS141143 | 378 | 1009 | BA_peak_9038 |
| DS141168 | 612 | 1022 | BA_peak_9039 |
| DS141198 | 139 | 1018 | BA_peak_9040 |
| DS141206 | 529 | 979 | BA_peak_9041 |
| DS141219 | 386 | 894 | BA_peak_9043 |
| DS141236 | 56 | 578 | BA_peak_9045 |
| DS141240 | 433 | 889 | BA_peak_9046 |
| DS141245 | 76 | 1040 | BA_peak_9047 |
| DS141275 | 66 | 1026 | BA_peak_9048 |
| DS141276 | 195 | 633 | BA_peak_9049 |
| DS141292 | 236 | 996 | BA_peak_9050 |
| DS141298 | 117 | 987 | BA_peak_9051 |
| DS141328 | 204 | 967 | BA_peak_9052 |
| DS141334 | 62 | 644 | BA_peak_9053 |
| DS141341 | 80 | 991 | BA_peak_9054 |
| DS141368 | 594 | 955 | BA_peak_9055 |
| DS141388 | 46 | 559 | BA_peak_9056 |
| DS141417 | 581 | 942 | BA_peak_9057 |
| DS141436 | 240 | 711 | BA_peak_9059 |
| DS141464 | 497 | 863 | BA_peak_9060 |
| DS141474 | 84 | 850 | BA_peak_9061 |

| | | | |
|---|---|---|---|
| DS141483 | 166 | 861 | BA_peak_9062 |
| DS141507 | 79 | 1009 | BA_peak_9063 |
| DS141516 | 198 | 658 | BA_peak_9064 |
| DS141518 | 147 | 650 | BA_peak_9065 |
| DS141529 | 161 | 873 | BA_peak_9066 |
| DS141535 | 376 | 837 | BA_peak_9067 |
| DS141537 | 57 | 547 | BA_peak_9068 |
| DS141547 | 384 | 1043 | BA_peak_9069 |
| DS141557 | 285 | 868 | BA_peak_9071 |
| DS141580 | 270 | 912 | BA_peak_9072 |
| DS141582 | 587 | 916 | BA_peak_9073 |
| DS141589 | 317 | 771 | BA_peak_9074 |
| DS141595 | 239 | 641 | BA_peak_9075 |
| DS141631 | 183 | 981 | BA_peak_9077 |
| DS141646 | 88 | 661 | BA_peak_9078 |
| DS141680 | 27 | 595 | BA_peak_9079 |
| DS141686 | 366 | 993 | BA_peak_9080 |
| DS141687 | 173 | 1010 | BA_peak_9081 |
| DS141689 | 153 | 483 | BA_peak_9082 |
| DS141697 | 271 | 1010 | BA_peak_9083 |
| DS141726 | 37 | 945 | BA_peak_9084 |
| DS141780 | 184 | 572 | BA_peak_9086 |
| DS141804 | 122 | 846 | BA_peak_9087 |
| DS141847 | 189 | 496 | BA_peak_9089 |
| DS141860 | 53 | 1024 | BA_peak_9091 |
| DS141881 | 90 | 966 | BA_peak_9092 |
| DS141885 | 433 | 735 | BA_peak_9093 |
| DS141903 | 202 | 578 | BA_peak_9094 |
| DS141917 | 162 | 903 | BA_peak_9095 |
| DS141932 | 20 | 657 | BA_peak_9096 |
| DS141941 | 421 | 934 | BA_peak_9097 |
| DS141949 | 176 | 956 | BA_peak_9098 |
| DS141960 | 237 | 619 | BA_peak_9099 |
| DS141966 | 451 | 964 | BA_peak_9100 |
| DS141993 | 225 | 1010 | BA_peak_9101 |
| DS141997 | 52 | 493 | BA_peak_9102 |
| DS142022 | 274 | 841 | BA_peak_9103 |
| DS142033 | 422 | 724 | BA_peak_9104 |
| DS142050 | 87 | 795 | BA_peak_9105 |
| DS142051 | 123 | 431 | BA_peak_9106 |
| DS142066 | 73 | 812 | BA_peak_9107 |
| DS142079 | 305 | 889 | BA_peak_9108 |
| DS142124 | 276 | 874 | BA_peak_9109 |
| DS142130 | 663 | 1058 | BA_peak_9110 |
| DS142145 | 357 | 898 | BA_peak_9111 |
| DS142157 | 104 | 1021 | BA_peak_9113 |
| DS142185 | 475 | 981 | BA_peak_9114 |

| | | | |
|---|---|---|---|
| DS142188 | 285 | 865 | BA_peak_9115 |
| DS142192 | 138 | 780 | BA_peak_9116 |
| DS142204 | 258 | 809 | BA_peak_9117 |
| DS142219 | 201 | 740 | BA_peak_9118 |
| DS142252 | 77 | 1014 | BA_peak_9119 |
| DS142253 | 31 | 560 | BA_peak_9120 |
| DS142255 | 509 | 797 | BA_peak_9121 |
| DS142270 | 46 | 487 | BA_peak_9123 |
| DS142310 | 278 | 988 | BA_peak_9124 |
| DS142317 | 39 | 1041 | BA_peak_9126 |
| DS142325 | 309 | 748 | BA_peak_9127 |
| DS142333 | 96 | 658 | BA_peak_9128 |
| DS142343 | 25 | 649 | BA_peak_9129 |
| DS142348 | 112 | 969 | BA_peak_9130 |
| DS142365 | 60 | 873 | BA_peak_9131 |
| DS142375 | 622 | 1038 | BA_peak_9132 |
| DS142385 | 81 | 598 | BA_peak_9133 |
| DS142388 | 363 | 950 | BA_peak_9134 |
| DS142411 | 103 | 643 | BA_peak_9135 |
| DS142462 | 524 | 936 | BA_peak_9136 |
| DS142464 | 35 | 1041 | BA_peak_9137 |
| DS142465 | 406 | 1026 | BA_peak_9138 |
| DS142471 | 102 | 931 | BA_peak_9139 |
| DS142492 | 680 | 1019 | BA_peak_9140 |
| DS142501 | 80 | 1012 | BA_peak_9141 |
| DS142526 | 107 | 872 | BA_peak_9143 |
| DS142528 | 410 | 899 | BA_peak_9144 |
| DS142546 | 474 | 917 | BA_peak_9145 |
| DS142563 | 181 | 857 | BA_peak_9146 |
| DS142591 | 47 | 474 | BA_peak_9147 |
| DS142599 | 587 | 1017 | BA_peak_9148 |
| DS142616 | 133 | 471 | BA_peak_9149 |
| DS142644 | 111 | 738 | BA_peak_9150 |
| DS142665 | 154 | 524 | BA_peak_9151 |
| DS142727 | 173 | 712 | BA_peak_9152 |
| DS142732 | 335 | 892 | BA_peak_9153 |
| DS142738 | 96 | 585 | BA_peak_9154 |
| DS142748 | 484 | 975 | BA_peak_9155 |
| DS142762 | 376 | 775 | BA_peak_9156 |
| DS142770 | 623 | 1016 | BA_peak_9157 |
| DS142802 | 49 | 816 | BA_peak_9158 |
| DS142856 | 159 | 935 | BA_peak_9159 |
| DS142860 | 147 | 596 | BA_peak_9160 |
| DS142861 | 415 | 967 | BA_peak_9161 |
| DS142882 | 220 | 758 | BA_peak_9162 |
| DS142884 | 107 | 558 | BA_peak_9163 |
| DS142899 | 104 | 996 | BA_peak_9164 |

| | | | |
|---|---|---|---|
| DS142905 | 108 | 704 | BA_peak_9165 |
| DS142906 | 106 | 1026 | BA_peak_9166 |
| DS142921 | 219 | 867 | BA_peak_9167 |
| DS142935 | 227 | 574 | BA_peak_9168 |
| DS142968 | 58 | 1015 | BA_peak_9169 |
| DS142970 | 139 | 603 | BA_peak_9170 |
| DS142972 | 116 | 590 | BA_peak_9171 |
| DS142991 | 54 | 408 | BA_peak_9172 |
| DS142994 | 82 | 1019 | BA_peak_9173 |
| DS143001 | 196 | 683 | BA_peak_9174 |
| DS143034 | 422 | 788 | BA_peak_9175 |
| DS143070 | 80 | 652 | BA_peak_9176 |
| DS143085 | 64 | 561 | BA_peak_9177 |
| DS143110 | 133 | 777 | BA_peak_9178 |
| DS143119 | 21 | 1030 | BA_peak_9179 |
| DS143120 | 218 | 948 | BA_peak_9180 |
| DS143141 | 42 | 959 | BA_peak_9181 |
| DS143145 | 172 | 944 | BA_peak_9182 |
| DS143171 | 544 | 1021 | BA_peak_9184 |
| DS143226 | 132 | 950 | BA_peak_9185 |
| DS143227 | 212 | 888 | BA_peak_9186 |
| DS143245 | 458 | 786 | BA_peak_9187 |
| DS143247 | 513 | 854 | BA_peak_9188 |
| DS143252 | 256 | 640 | BA_peak_9189 |
| DS143256 | 157 | 1002 | BA_peak_9190 |
| DS143313 | 41 | 1049 | BA_peak_9191 |
| DS143316 | 156 | 816 | BA_peak_9192 |
| DS143327 | 124 | 788 | BA_peak_9193 |
| DS143358 | 106 | 980 | BA_peak_9194 |
| DS143375 | 193 | 614 | BA_peak_9195 |
| DS143465 | 80 | 960 | BA_peak_9196 |
| DS143477 | 497 | 1010 | BA_peak_9197 |
| DS143484 | 377 | 950 | BA_peak_9198 |
| DS143499 | 584 | 1021 | BA_peak_9199 |
| DS143506 | 33 | 430 | BA_peak_9200 |
| DS143518 | 480 | 801 | BA_peak_9201 |
| DS143530 | 17 | 395 | BA_peak_9202 |
| DS143538 | 484 | 879 | BA_peak_9204 |
| DS143552 | 197 | 772 | BA_peak_9205 |
| DS143585 | 155 | 980 | BA_peak_9206 |
| DS143586 | 14 | 795 | BA_peak_9207 |
| DS143606 | 413 | 982 | BA_peak_9208 |
| DS143613 | 458 | 920 | BA_peak_9209 |
| DS143653 | 296 | 608 | BA_peak_9210 |
| DS143690 | 36 | 801 | BA_peak_9211 |
| DS143716 | 227 | 941 | BA_peak_9213 |
| DS143721 | 356 | 1027 | BA_peak_9214 |

| | | | |
|---|---|---|---|
| DS143731 | 373 | 888 | BA_peak_9215 |
| DS143764 | 93 | 820 | BA_peak_9216 |
| DS143823 | 441 | 827 | BA_peak_9217 |
| DS143842 | 130 | 804 | BA_peak_9218 |
| DS143878 | 392 | 821 | BA_peak_9220 |
| DS143902 | 509 | 914 | BA_peak_9221 |
| DS143941 | 178 | 734 | BA_peak_9222 |
| DS143969 | 142 | 594 | BA_peak_9223 |
| DS143971 | 125 | 782 | BA_peak_9224 |
| DS143986 | 350 | 973 | BA_peak_9225 |
| DS143993 | 59 | 721 | BA_peak_9226 |
| DS144035 | 39 | 1024 | BA_peak_9227 |
| DS144049 | 81 | 912 | BA_peak_9228 |
| DS144056 | 27 | 399 | BA_peak_9229 |
| DS144059 | 74 | 663 | BA_peak_9230 |
| DS144109 | 458 | 919 | BA_peak_9232 |
| DS144116 | 263 | 802 | BA_peak_9233 |
| DS144121 | 54 | 1037 | BA_peak_9234 |
| DS144149 | 80 | 491 | BA_peak_9235 |
| DS144177 | 345 | 946 | BA_peak_9236 |
| DS144179 | 67 | 950 | BA_peak_9237 |
| DS144190 | 173 | 938 | BA_peak_9238 |
| DS144226 | 410 | 981 | BA_peak_9239 |
| DS144230 | 151 | 963 | BA_peak_9241 |
| DS144299 | 426 | 962 | BA_peak_9242 |
| DS144301 | 155 | 785 | BA_peak_9243 |
| DS144306 | 466 | 876 | BA_peak_9244 |
| DS144309 | 332 | 856 | BA_peak_9245 |
| DS144330 | 170 | 680 | BA_peak_9246 |
| DS144338 | 509 | 1002 | BA_peak_9247 |
| DS144347 | 118 | 791 | BA_peak_9248 |
| DS144366 | 494 | 960 | BA_peak_9249 |
| DS144367 | 391 | 806 | BA_peak_9250 |
| DS144376 | 583 | 1013 | BA_peak_9251 |
| DS144398 | 181 | 638 | BA_peak_9252 |
| DS144401 | 169 | 1001 | BA_peak_9253 |
| DS144417 | 40 | 885 | BA_peak_9254 |
| DS144420 | 226 | 614 | BA_peak_9255 |
| DS144494 | 183 | 585 | BA_peak_9257 |
| DS144516 | 481 | 969 | BA_peak_9258 |
| DS144524 | 171 | 637 | BA_peak_9259 |
| DS144570 | 377 | 773 | BA_peak_9260 |
| DS144576 | 69 | 496 | BA_peak_9261 |
| DS144595 | 100 | 748 | BA_peak_9262 |
| DS144605 | 120 | 961 | BA_peak_9263 |
| DS144606 | 351 | 619 | BA_peak_9264 |
| DS144615 | 51 | 763 | BA_peak_9265 |

| | | | |
|---|---|---|---|
| DS144617 | 389 | 932 | BA_peak_9266 |
| DS144638 | 606 | 1003 | BA_peak_9267 |
| DS144653 | 151 | 740 | BA_peak_9268 |
| DS144668 | 55 | 985 | BA_peak_9269 |
| DS144750 | 170 | 907 | BA_peak_9270 |
| DS144761 | 340 | 651 | BA_peak_9271 |
| DS144773 | 35 | 462 | BA_peak_9272 |
| DS144781 | 121 | 1008 | BA_peak_9273 |
| DS144791 | 116 | 639 | BA_peak_9274 |
| DS144804 | 208 | 1024 | BA_peak_9275 |
| DS144848 | 409 | 944 | BA_peak_9276 |
| DS144879 | 18 | 1011 | BA_peak_9278 |
| DS144909 | 68 | 912 | BA_peak_9279 |
| DS144916 | 330 | 782 | BA_peak_9280 |
| DS144917 | 380 | 803 | BA_peak_9281 |
| DS144936 | 222 | 814 | BA_peak_9283 |
| DS144946 | 17 | 1008 | BA_peak_9284 |
| DS144948 | 136 | 671 | BA_peak_9285 |
| DS144956 | 31 | 972 | BA_peak_9286 |
| DS144957 | 273 | 604 | BA_peak_9287 |
| DS144965 | 94 | 930 | BA_peak_9288 |
| DS144992 | 87 | 483 | BA_peak_9290 |
| DS145028 | 325 | 953 | BA_peak_9291 |
| DS145035 | 25 | 961 | BA_peak_9292 |
| DS145036 | 442 | 860 | BA_peak_9293 |
| DS145038 | 283 | 630 | BA_peak_9294 |
| DS145042 | 94 | 968 | BA_peak_9295 |
| DS145085 | 55 | 742 | BA_peak_9297 |
| DS145089 | 75 | 995 | BA_peak_9298 |
| DS145120 | 386 | 946 | BA_peak_9301 |
| DS145122 | 66 | 995 | BA_peak_9302 |
| DS145131 | 155 | 917 | BA_peak_9303 |
| DS145187 | 89 | 423 | BA_peak_9304 |
| DS145201 | 49 | 687 | BA_peak_9305 |
| DS145253 | 396 | 747 | BA_peak_9307 |
| DS145255 | 98 | 538 | BA_peak_9308 |
| DS145272 | 485 | 884 | BA_peak_9309 |
| DS145275 | 145 | 871 | BA_peak_9310 |
| DS145280 | 398 | 703 | BA_peak_9311 |
| DS145287 | 362 | 1009 | BA_peak_9312 |
| DS145330 | 125 | 846 | BA_peak_9313 |
| DS145378 | 64 | 750 | BA_peak_9314 |
| DS145388 | 637 | 994 | BA_peak_9316 |
| DS145406 | 125 | 989 | BA_peak_9317 |
| DS145418 | 137 | 803 | BA_peak_9319 |
| DS145477 | 50 | 734 | BA_peak_9320 |
| DS145508 | 177 | 842 | BA_peak_9321 |

| | | | |
|---|---|---|---|
| DS145520 | 81 | 768 | BA_peak_9322 |
| DS145531 | 85 | 992 | BA_peak_9323 |
| DS145534 | 103 | 849 | BA_peak_9324 |
| DS145538 | 539 | 852 | BA_peak_9325 |
| DS145549 | 23 | 997 | BA_peak_9326 |
| DS145552 | 124 | 905 | BA_peak_9327 |
| DS145560 | 39 | 944 | BA_peak_9328 |
| DS145588 | 331 | 773 | BA_peak_9329 |
| DS145622 | 241 | 636 | BA_peak_9331 |
| DS145628 | 84 | 454 | BA_peak_9332 |
| DS145650 | 171 | 684 | BA_peak_9333 |
| DS145662 | 180 | 915 | BA_peak_9335 |
| DS145736 | 61 | 865 | BA_peak_9337 |
| DS145737 | 43 | 974 | BA_peak_9338 |
| DS145802 | 48 | 1032 | BA_peak_9339 |
| DS145830 | 570 | 963 | BA_peak_9340 |
| DS145840 | 112 | 908 | BA_peak_9341 |
| DS145877 | 67 | 971 | BA_peak_9342 |
| DS145897 | 186 | 994 | BA_peak_9343 |
| DS145922 | 167 | 803 | BA_peak_9344 |
| DS145991 | 37 | 847 | BA_peak_9345 |
| DS146023 | 32 | 953 | BA_peak_9347 |
| DS146044 | 76 | 571 | BA_peak_9348 |
| DS146051 | 65 | 898 | BA_peak_9349 |
| DS146081 | 92 | 959 | BA_peak_9350 |
| DS146091 | 237 | 890 | BA_peak_9351 |
| DS146096 | 421 | 896 | BA_peak_9352 |
| DS146109 | 57 | 722 | BA_peak_9353 |
| DS146119 | 66 | 681 | BA_peak_9354 |
| DS146140 | 216 | 884 | BA_peak_9355 |
| DS146159 | 74 | 413 | BA_peak_9357 |
| DS146160 | 323 | 896 | BA_peak_9358 |
| DS146168 | 180 | 978 | BA_peak_9359 |
| DS146188 | 119 | 869 | BA_peak_9360 |
| DS146200 | 92 | 962 | BA_peak_9361 |
| DS146217 | 159 | 874 | BA_peak_9362 |
| DS146228 | 229 | 616 | BA_peak_9364 |
| DS146281 | 110 | 928 | BA_peak_9365 |
| DS146284 | 589 | 971 | BA_peak_9366 |
| DS146285 | 260 | 820 | BA_peak_9367 |
| DS146315 | 150 | 782 | BA_peak_9368 |
| DS146338 | 439 | 769 | BA_peak_9369 |
| DS146372 | 47 | 1011 | BA_peak_9370 |
| DS146389 | 100 | 959 | BA_peak_9371 |
| DS146391 | 112 | 918 | BA_peak_9372 |
| DS146395 | 239 | 669 | BA_peak_9373 |
| DS146402 | 509 | 850 | BA_peak_9374 |

| | | | |
|---|---|---|---|
| DS146404 | 211 | 951 | BA_peak_9375 |
| DS146432 | 37 | 1006 | BA_peak_9376 |
| DS146470 | 235 | 689 | BA_peak_9377 |
| DS146497 | 24 | 1028 | BA_peak_9378 |
| DS146498 | 37 | 718 | BA_peak_9379 |
| DS146499 | 276 | 592 | BA_peak_9380 |
| DS146510 | 158 | 832 | BA_peak_9381 |
| DS146522 | 41 | 894 | BA_peak_9382 |
| DS146547 | 42 | 946 | BA_peak_9383 |
| DS146556 | 98 | 936 | BA_peak_9384 |
| DS146559 | 94 | 600 | BA_peak_9385 |
| DS146572 | 194 | 689 | BA_peak_9386 |
| DS146575 | 127 | 942 | BA_peak_9387 |
| DS146627 | 63 | 427 | BA_peak_9389 |
| DS146642 | 499 | 960 | BA_peak_9390 |
| DS146649 | 69 | 1000 | BA_peak_9391 |
| DS146682 | 633 | 988 | BA_peak_9392 |
| DS146727 | 318 | 662 | BA_peak_9393 |
| DS146731 | 75 | 947 | BA_peak_9394 |
| DS146742 | 43 | 883 | BA_peak_9395 |
| DS146766 | 402 | 833 | BA_peak_9396 |
| DS146789 | 261 | 735 | BA_peak_9397 |
| DS146795 | 62 | 996 | BA_peak_9398 |
| DS146811 | 243 | 994 | BA_peak_9399 |
| DS146835 | 47 | 838 | BA_peak_9400 |
| DS146848 | 292 | 754 | BA_peak_9401 |
| DS146851 | 127 | 863 | BA_peak_9402 |
| DS146854 | 15 | 978 | BA_peak_9403 |
| DS146868 | 254 | 670 | BA_peak_9404 |
| DS146873 | 342 | 938 | BA_peak_9405 |
| DS146915 | 219 | 626 | BA_peak_9406 |
| DS146933 | 228 | 931 | BA_peak_9407 |
| DS146948 | 483 | 897 | BA_peak_9408 |
| DS146951 | 491 | 955 | BA_peak_9409 |
| DS146975 | 441 | 878 | BA_peak_9410 |
| DS146982 | 193 | 820 | BA_peak_9412 |
| DS146993 | 376 | 934 | BA_peak_9413 |
| DS147010 | 253 | 606 | BA_peak_9414 |
| DS147022 | 489 | 819 | BA_peak_9415 |
| DS147038 | 96 | 941 | BA_peak_9416 |
| DS147068 | 170 | 515 | BA_peak_9417 |
| DS147084 | 125 | 1016 | BA_peak_9418 |
| DS147095 | 140 | 810 | BA_peak_9419 |
| DS147101 | 118 | 774 | BA_peak_9420 |
| DS147108 | 315 | 984 | BA_peak_9421 |
| DS147159 | 30 | 449 | BA_peak_9422 |
| DS147177 | 79 | 988 | BA_peak_9423 |

| | | | |
|---|---|---|---|
| DS147190 | 82 | 495 | BA_peak_9424 |
| DS147192 | 176 | 789 | BA_peak_9425 |
| DS147194 | 352 | 954 | BA_peak_9426 |
| DS147278 | 274 | 963 | BA_peak_9427 |
| DS147282 | 242 | 945 | BA_peak_9428 |
| DS147297 | 152 | 461 | BA_peak_9429 |
| DS147302 | 38 | 974 | BA_peak_9430 |
| DS147333 | 11 | 923 | BA_peak_9431 |
| DS147350 | 525 | 971 | BA_peak_9432 |
| DS147367 | 121 | 827 | BA_peak_9433 |
| DS147379 | 320 | 870 | BA_peak_9434 |
| DS147380 | 144 | 822 | BA_peak_9435 |
| DS147398 | 98 | 680 | BA_peak_9436 |
| DS147441 | 87 | 908 | BA_peak_9437 |
| DS147459 | 260 | 765 | BA_peak_9438 |
| DS147541 | 336 | 836 | BA_peak_9440 |
| DS147547 | 316 | 906 | BA_peak_9441 |
| DS147577 | 326 | 767 | BA_peak_9443 |
| DS147597 | 253 | 1013 | BA_peak_9444 |
| DS147600 | 139 | 547 | BA_peak_9445 |
| DS147644 | 228 | 978 | BA_peak_9446 |
| DS147671 | 446 | 779 | BA_peak_9447 |
| DS147681 | 192 | 969 | BA_peak_9449 |
| DS147684 | 108 | 491 | BA_peak_9450 |
| DS147757 | 48 | 919 | BA_peak_9451 |
| DS147773 | 208 | 903 | BA_peak_9452 |
| DS147783 | 83 | 848 | BA_peak_9453 |
| DS147791 | 169 | 509 | BA_peak_9454 |
| DS147803 | 156 | 748 | BA_peak_9455 |
| DS147804 | 383 | 917 | BA_peak_9456 |
| DS147815 | 28 | 900 | BA_peak_9457 |
| DS147836 | 148 | 995 | BA_peak_9458 |
| DS147865 | 133 | 838 | BA_peak_9459 |
| DS147881 | 53 | 988 | BA_peak_9460 |
| DS147910 | 429 | 868 | BA_peak_9461 |
| DS147919 | 614 | 958 | BA_peak_9462 |
| DS147992 | 51 | 974 | BA_peak_9464 |
| DS148007 | 131 | 924 | BA_peak_9465 |
| DS148012 | 147 | 736 | BA_peak_9466 |
| DS148050 | 45 | 864 | BA_peak_9467 |
| DS148067 | 188 | 742 | BA_peak_9468 |
| DS148069 | 19 | 986 | BA_peak_9469 |
| DS148074 | 184 | 656 | BA_peak_9470 |
| DS148077 | 367 | 731 | BA_peak_9471 |
| DS148093 | 37 | 837 | BA_peak_9472 |
| DS148134 | 154 | 547 | BA_peak_9473 |
| DS148135 | 24 | 657 | BA_peak_9474 |

| | | | |
|---|---|---|---|
| DS148153 | 236 | 633 | BA_peak_9475 |
| DS148163 | 226 | 742 | BA_peak_9476 |
| DS148179 | 530 | 982 | BA_peak_9477 |
| DS148201 | 398 | 823 | BA_peak_9478 |
| DS148206 | 24 | 935 | BA_peak_9479 |
| DS148225 | 29 | 1010 | BA_peak_9480 |
| DS148258 | 338 | 817 | BA_peak_9481 |
| DS148274 | 11 | 614 | BA_peak_9482 |
| DS148288 | 393 | 1001 | BA_peak_9483 |
| DS148336 | 582 | 981 | BA_peak_9484 |
| DS148339 | 309 | 911 | BA_peak_9485 |
| DS148359 | 46 | 947 | BA_peak_9486 |
| DS148413 | 319 | 692 | BA_peak_9487 |
| DS148418 | 32 | 981 | BA_peak_9489 |
| DS148434 | 139 | 536 | BA_peak_9490 |
| DS148439 | 401 | 937 | BA_peak_9491 |
| DS148467 | 286 | 715 | BA_peak_9492 |
| DS148530 | 533 | 906 | BA_peak_9494 |
| DS148542 | 424 | 730 | BA_peak_9495 |
| DS148549 | 101 | 775 | BA_peak_9496 |
| DS148573 | 180 | 997 | BA_peak_9497 |
| DS148592 | 176 | 576 | BA_peak_9498 |
| DS148596 | 484 | 892 | BA_peak_9499 |
| DS148597 | 197 | 635 | BA_peak_9500 |
| DS148613 | 639 | 980 | BA_peak_9501 |
| DS148617 | 464 | 945 | BA_peak_9502 |
| DS148641 | 161 | 711 | BA_peak_9503 |
| DS148657 | 81 | 457 | BA_peak_9505 |
| DS148661 | 360 | 865 | BA_peak_9506 |
| DS148677 | 23 | 880 | BA_peak_9507 |
| DS148688 | 125 | 834 | BA_peak_9508 |
| DS148690 | 16 | 857 | BA_peak_9509 |
| DS148699 | 199 | 900 | BA_peak_9510 |
| DS148700 | 481 | 824 | BA_peak_9511 |
| DS148733 | 107 | 888 | BA_peak_9512 |
| DS148738 | 513 | 846 | BA_peak_9513 |
| DS148742 | 64 | 905 | BA_peak_9514 |
| DS148780 | 67 | 918 | BA_peak_9516 |
| DS148800 | 117 | 682 | BA_peak_9517 |
| DS148816 | 28 | 453 | BA_peak_9518 |
| DS148822 | 52 | 562 | BA_peak_9519 |
| DS148835 | 22 | 895 | BA_peak_9520 |
| DS148846 | 522 | 883 | BA_peak_9521 |
| DS148901 | 57 | 926 | BA_peak_9522 |
| DS148909 | 180 | 973 | BA_peak_9523 |
| DS148934 | 107 | 919 | BA_peak_9525 |
| DS148955 | 221 | 808 | BA_peak_9526 |

| | | | |
|---|---|---|---|
| DS148965 | 80 | 950 | BA_peak_9527 |
| DS148981 | 426 | 836 | BA_peak_9528 |
| DS148997 | 258 | 869 | BA_peak_9529 |
| DS149053 | 179 | 931 | BA_peak_9532 |
| DS149070 | 209 | 791 | BA_peak_9533 |
| DS149091 | 88 | 845 | BA_peak_9534 |
| DS149103 | 60 | 782 | BA_peak_9535 |
| DS149135 | 418 | 693 | BA_peak_9536 |
| DS149140 | 238 | 921 | BA_peak_9537 |
| DS149147 | 35 | 374 | BA_peak_9538 |
| DS149156 | 71 | 840 | BA_peak_9539 |
| DS149162 | 366 | 781 | BA_peak_9540 |
| DS149166 | 250 | 676 | BA_peak_9541 |
| DS149176 | 124 | 682 | BA_peak_9542 |
| DS149187 | 294 | 800 | BA_peak_9543 |
| DS149235 | 142 | 725 | BA_peak_9544 |
| DS149248 | 175 | 830 | BA_peak_9545 |
| DS149249 | 162 | 516 | BA_peak_9546 |
| DS149250 | 354 | 918 | BA_peak_9547 |
| DS149280 | 296 | 930 | BA_peak_9548 |
| DS149291 | 118 | 877 | BA_peak_9549 |
| DS149322 | 260 | 898 | BA_peak_9550 |
| DS149340 | 100 | 621 | BA_peak_9551 |
| DS149398 | 267 | 642 | BA_peak_9552 |
| DS149420 | 37 | 907 | BA_peak_9553 |
| DS149464 | 484 | 888 | BA_peak_9556 |
| DS149480 | 512 | 946 | BA_peak_9557 |
| DS149484 | 393 | 671 | BA_peak_9558 |
| DS149537 | 432 | 973 | BA_peak_9559 |
| DS149546 | 33 | 656 | BA_peak_9560 |
| DS149557 | 26 | 873 | BA_peak_9561 |
| DS149558 | 192 | 845 | BA_peak_9562 |
| DS149576 | 199 | 517 | BA_peak_9563 |
| DS149627 | 113 | 716 | BA_peak_9564 |
| DS149631 | 101 | 501 | BA_peak_9565 |
| DS149643 | 107 | 923 | BA_peak_9566 |
| DS149652 | 305 | 638 | BA_peak_9567 |
| DS149665 | 60 | 514 | BA_peak_9568 |
| DS149667 | 203 | 815 | BA_peak_9569 |
| DS149702 | 191 | 787 | BA_peak_9570 |
| DS149709 | 287 | 829 | BA_peak_9571 |
| DS149757 | 380 | 856 | BA_peak_9572 |
| DS149766 | 40 | 1003 | BA_peak_9573 |
| DS149817 | 113 | 421 | BA_peak_9574 |
| DS149852 | 17 | 983 | BA_peak_9576 |
| DS149864 | 388 | 837 | BA_peak_9577 |
| DS149871 | 126 | 947 | BA_peak_9578 |

| | | | |
|---|---|---|---|
| DS149883 | 144 | 844 | BA_peak_9579 |
| DS149931 | 545 | 848 | BA_peak_9580 |
| DS149933 | 188 | 490 | BA_peak_9581 |
| DS149960 | 111 | 881 | BA_peak_9583 |
| DS149985 | 426 | 873 | BA_peak_9584 |
| DS150061 | 61 | 769 | BA_peak_9585 |
| DS150119 | 53 | 375 | BA_peak_9586 |
| DS150126 | 17 | 485 | BA_peak_9587 |
| DS150152 | 134 | 754 | BA_peak_9588 |
| DS150173 | 109 | 785 | BA_peak_9589 |
| DS150212 | 85 | 1001 | BA_peak_9591 |
| DS150244 | 72 | 862 | BA_peak_9592 |
| DS150314 | 226 | 961 | BA_peak_9593 |
| DS150323 | 81 | 797 | BA_peak_9594 |
| DS150343 | 105 | 777 | BA_peak_9595 |
| DS150393 | 224 | 853 | BA_peak_9597 |
| DS150414 | 68 | 979 | BA_peak_9598 |
| DS150416 | 379 | 846 | BA_peak_9599 |
| DS150417 | 53 | 926 | BA_peak_9600 |
| DS150419 | 28 | 930 | BA_peak_9601 |
| DS150433 | 336 | 877 | BA_peak_9602 |
| DS150436 | 152 | 694 | BA_peak_9603 |
| DS150447 | 107 | 914 | BA_peak_9604 |
| DS150466 | 82 | 916 | BA_peak_9605 |
| DS150478 | 416 | 863 | BA_peak_9606 |
| DS150513 | 62 | 906 | BA_peak_9607 |
| DS150537 | 230 | 867 | BA_peak_9608 |
| DS150553 | 75 | 790 | BA_peak_9609 |
| DS150583 | 108 | 554 | BA_peak_9610 |
| DS150605 | 59 | 972 | BA_peak_9611 |
| DS150614 | 120 | 733 | BA_peak_9612 |
| DS150620 | 50 | 959 | BA_peak_9613 |
| DS150630 | 33 | 461 | BA_peak_9615 |
| DS150644 | 538 | 959 | BA_peak_9617 |
| DS150655 | 99 | 799 | BA_peak_9618 |
| DS150688 | 123 | 544 | BA_peak_9619 |
| DS150697 | 203 | 966 | BA_peak_9620 |
| DS150704 | 373 | 807 | BA_peak_9621 |
| DS150728 | 121 | 908 | BA_peak_9622 |
| DS150748 | 131 | 920 | BA_peak_9623 |
| DS150770 | 37 | 381 | BA_peak_9625 |
| DS150775 | 441 | 897 | BA_peak_9626 |
| DS150800 | 117 | 950 | BA_peak_9627 |
| DS150819 | 136 | 791 | BA_peak_9628 |
| DS150823 | 633 | 984 | BA_peak_9629 |
| DS150845 | 168 | 923 | BA_peak_9631 |
| DS150864 | 146 | 880 | BA_peak_9632 |

| | | | |
|---|---|---|---|
| DS150879 | 341 | 909 | BA_peak_9633 |
| DS150890 | 226 | 963 | BA_peak_9634 |
| DS150894 | 75 | 504 | BA_peak_9635 |
| DS150898 | 50 | 963 | BA_peak_9636 |
| DS150924 | 86 | 598 | BA_peak_9637 |
| DS150932 | 259 | 715 | BA_peak_9638 |
| DS150948 | 616 | 931 | BA_peak_9639 |
| DS150962 | 80 | 975 | BA_peak_9640 |
| DS151003 | 243 | 654 | BA_peak_9641 |
| DS151020 | 128 | 846 | BA_peak_9642 |
| DS151022 | 166 | 573 | BA_peak_9643 |
| DS151034 | 247 | 831 | BA_peak_9644 |
| DS151044 | 213 | 541 | BA_peak_9645 |
| DS151077 | 349 | 898 | BA_peak_9647 |
| DS151104 | 90 | 609 | BA_peak_9648 |
| DS151111 | 530 | 900 | BA_peak_9649 |
| DS151123 | 441 | 836 | BA_peak_9650 |
| DS151133 | 249 | 733 | BA_peak_9651 |
| DS151177 | 590 | 935 | BA_peak_9652 |
| DS151199 | 103 | 942 | BA_peak_9653 |
| DS151227 | 511 | 982 | BA_peak_9654 |
| DS151234 | 385 | 836 | BA_peak_9655 |
| DS151244 | 40 | 997 | BA_peak_9656 |
| DS151248 | 72 | 825 | BA_peak_9657 |
| DS151251 | 76 | 639 | BA_peak_9658 |
| DS151255 | 224 | 583 | BA_peak_9659 |
| DS151259 | 204 | 934 | BA_peak_9660 |
| DS151262 | 84 | 816 | BA_peak_9661 |
| DS151268 | 325 | 808 | BA_peak_9663 |
| DS151276 | 63 | 947 | BA_peak_9664 |
| DS151281 | 261 | 989 | BA_peak_9665 |
| DS151283 | 100 | 895 | BA_peak_9666 |
| DS151324 | 158 | 899 | BA_peak_9668 |
| DS151325 | 95 | 852 | BA_peak_9669 |
| DS151343 | 308 | 726 | BA_peak_9670 |
| DS151349 | 107 | 899 | BA_peak_9671 |
| DS151371 | 144 | 668 | BA_peak_9674 |
| DS151373 | 84 | 851 | BA_peak_9675 |
| DS151393 | 336 | 769 | BA_peak_9676 |
| DS151415 | 183 | 946 | BA_peak_9677 |
| DS151421 | 214 | 838 | BA_peak_9679 |
| DS151429 | 278 | 936 | BA_peak_9681 |
| DS151437 | 245 | 921 | BA_peak_9682 |
| DS151455 | 83 | 546 | BA_peak_9683 |
| DS151462 | 87 | 812 | BA_peak_9684 |
| DS151473 | 178 | 753 | BA_peak_9685 |
| DS151476 | 284 | 940 | BA_peak_9686 |

| | | | |
|---|---|---|---|
| DS151496 | 251 | 821 | BA_peak_9688 |
| DS151502 | 93 | 873 | BA_peak_9689 |
| DS151510 | 384 | 887 | BA_peak_9690 |
| DS151517 | 34 | 816 | BA_peak_9691 |
| DS151526 | 136 | 772 | BA_peak_9692 |
| DS151534 | 59 | 920 | BA_peak_9693 |
| DS151543 | 150 | 777 | BA_peak_9695 |
| DS151551 | 17 | 474 | BA_peak_9696 |
| DS151577 | 433 | 889 | BA_peak_9697 |
| DS151586 | 228 | 564 | BA_peak_9698 |
| DS151591 | 489 | 855 | BA_peak_9699 |
| DS151614 | 79 | 916 | BA_peak_9701 |
| DS151632 | 204 | 753 | BA_peak_9702 |
| DS151660 | 287 | 879 | BA_peak_9704 |
| DS151666 | 499 | 953 | BA_peak_9705 |
| DS151668 | 49 | 904 | BA_peak_9706 |
| DS151678 | 133 | 530 | BA_peak_9708 |
| DS151679 | 126 | 673 | BA_peak_9709 |
| DS151691 | 203 | 786 | BA_peak_9710 |
| DS151716 | 284 | 839 | BA_peak_9711 |
| DS151718 | 152 | 914 | BA_peak_9712 |
| DS151751 | 113 | 834 | BA_peak_9714 |
| DS151768 | 17 | 861 | BA_peak_9715 |
| DS151849 | 50 | 787 | BA_peak_9717 |
| DS151882 | 241 | 587 | BA_peak_9720 |
| DS151912 | 78 | 971 | BA_peak_9721 |
| DS151914 | 138 | 936 | BA_peak_9722 |
| DS151968 | 41 | 960 | BA_peak_9723 |
| DS151997 | 114 | 885 | BA_peak_9725 |
| DS151999 | 76 | 665 | BA_peak_9726 |
| DS152028 | 271 | 924 | BA_peak_9728 |
| DS152067 | 69 | 973 | BA_peak_9730 |
| DS152071 | 438 | 755 | BA_peak_9731 |
| DS152084 | 212 | 924 | BA_peak_9732 |
| DS152182 | 191 | 497 | BA_peak_9733 |
| DS152189 | 440 | 845 | BA_peak_9734 |
| DS152193 | 358 | 878 | BA_peak_9735 |
| DS152196 | 113 | 729 | BA_peak_9736 |
| DS152197 | 131 | 933 | BA_peak_9737 |
| DS152236 | 56 | 596 | BA_peak_9740 |
| DS152248 | 19 | 419 | BA_peak_9741 |
| DS152267 | 46 | 925 | BA_peak_9742 |
| DS152350 | 82 | 407 | BA_peak_9744 |
| DS152351 | 81 | 870 | BA_peak_9745 |
| DS152359 | 90 | 924 | BA_peak_9746 |
| DS152371 | 61 | 564 | BA_peak_9747 |
| DS152391 | 99 | 946 | BA_peak_9748 |

| | | | |
|---|---|---|---|
| DS152407 | 13 | 351 | BA_peak_9749 |
| DS152414 | 117 | 470 | BA_peak_9750 |
| DS152462 | 124 | 968 | BA_peak_9751 |
| DS152463 | 242 | 672 | BA_peak_9752 |
| DS152475 | 248 | 560 | BA_peak_9753 |
| DS152480 | 79 | 618 | BA_peak_9754 |
| DS152483 | 11 | 982 | BA_peak_9756 |
| DS152487 | 76 | 679 | BA_peak_9757 |
| DS152501 | 29 | 458 | BA_peak_9758 |
| DS152524 | 437 | 878 | BA_peak_9759 |
| DS152538 | 241 | 954 | BA_peak_9760 |
| DS152545 | 191 | 867 | BA_peak_9761 |
| DS152577 | 32 | 861 | BA_peak_9762 |
| DS152587 | 196 | 955 | BA_peak_9763 |
| DS152689 | 258 | 789 | BA_peak_9766 |
| DS152706 | 489 | 753 | BA_peak_9767 |
| DS152734 | 182 | 494 | BA_peak_9768 |
| DS152738 | 381 | 898 | BA_peak_9769 |
| DS152741 | 80 | 876 | BA_peak_9770 |
| DS152754 | 193 | 855 | BA_peak_9771 |
| DS152760 | 339 | 668 | BA_peak_9772 |
| DS152763 | 44 | 710 | BA_peak_9773 |
| DS152792 | 531 | 969 | BA_peak_9775 |
| DS152821 | 518 | 830 | BA_peak_9776 |
| DS152852 | 287 | 730 | BA_peak_9779 |
| DS152860 | 257 | 967 | BA_peak_9780 |
| DS152870 | 47 | 912 | BA_peak_9781 |
| DS152878 | 54 | 561 | BA_peak_9782 |
| DS152884 | 67 | 855 | BA_peak_9783 |
| DS152886 | 88 | 977 | BA_peak_9784 |
| DS152930 | 414 | 933 | BA_peak_9785 |
| DS152967 | 61 | 841 | BA_peak_9788 |
| DS152988 | 219 | 894 | BA_peak_9789 |
| DS153018 | 91 | 617 | BA_peak_9790 |
| DS153085 | 213 | 633 | BA_peak_9791 |
| DS153129 | 12 | 419 | BA_peak_9792 |
| DS153154 | 139 | 907 | BA_peak_9793 |
| DS153168 | 121 | 502 | BA_peak_9794 |
| DS153198 | 296 | 935 | BA_peak_9796 |
| DS153199 | 367 | 837 | BA_peak_9797 |
| DS153214 | 141 | 734 | BA_peak_9798 |
| DS153215 | 173 | 698 | BA_peak_9799 |
| DS153241 | 91 | 429 | BA_peak_9800 |
| DS153242 | 162 | 826 | BA_peak_9801 |
| DS153313 | 242 | 891 | BA_peak_9804 |
| DS153318 | 87 | 876 | BA_peak_9805 |
| DS153333 | 99 | 867 | BA_peak_9807 |

| | | | |
|---|---|---|---|
| DS153336 | 107 | 850 | BA_peak_9808 |
| DS153341 | 41 | 891 | BA_peak_9809 |
| DS153345 | 134 | 578 | BA_peak_9810 |
| DS153367 | 119 | 778 | BA_peak_9811 |
| DS153369 | 134 | 654 | BA_peak_9812 |
| DS153373 | 147 | 942 | BA_peak_9813 |
| DS153374 | 227 | 860 | BA_peak_9814 |
| DS153418 | 38 | 896 | BA_peak_9815 |
| DS153464 | 70 | 885 | BA_peak_9816 |
| DS153498 | 164 | 697 | BA_peak_9817 |
| DS153532 | 163 | 589 | BA_peak_9818 |
| DS153561 | 358 | 944 | BA_peak_9819 |
| DS153563 | 181 | 886 | BA_peak_9820 |
| DS153567 | 27 | 979 | BA_peak_9821 |
| DS153572 | 163 | 962 | BA_peak_9822 |
| DS153592 | 565 | 949 | BA_peak_9823 |
| DS153602 | 40 | 886 | BA_peak_9824 |
| DS153612 | 172 | 769 | BA_peak_9825 |
| DS153613 | 125 | 721 | BA_peak_9826 |
| DS153635 | 586 | 904 | BA_peak_9827 |
| DS153644 | 30 | 714 | BA_peak_9828 |
| DS153662 | 206 | 924 | BA_peak_9829 |
| DS153687 | 261 | 863 | BA_peak_9830 |
| DS153708 | 158 | 864 | BA_peak_9831 |
| DS153735 | 110 | 714 | BA_peak_9832 |
| DS153746 | 161 | 590 | BA_peak_9833 |
| DS153786 | 30 | 912 | BA_peak_9834 |
| DS153794 | 148 | 786 | BA_peak_9835 |
| DS153809 | 63 | 877 | BA_peak_9836 |
| DS153814 | 137 | 968 | BA_peak_9837 |
| DS153855 | 227 | 856 | BA_peak_9840 |
| DS153868 | 27 | 614 | BA_peak_9841 |
| DS153870 | 36 | 880 | BA_peak_9842 |
| DS153931 | 431 | 768 | BA_peak_9844 |
| DS153936 | 342 | 908 | BA_peak_9845 |
| DS153960 | 24 | 644 | BA_peak_9847 |
| DS153966 | 102 | 668 | BA_peak_9848 |
| DS153976 | 219 | 830 | BA_peak_9849 |
| DS153992 | 49 | 927 | BA_peak_9850 |
| DS154008 | 170 | 530 | BA_peak_9851 |
| DS154054 | 23 | 939 | BA_peak_9853 |
| DS154061 | 269 | 749 | BA_peak_9854 |
| DS154062 | 153 | 839 | BA_peak_9855 |
| DS154071 | 475 | 941 | BA_peak_9856 |
| DS154111 | 156 | 763 | BA_peak_9857 |
| DS154193 | 50 | 845 | BA_peak_9858 |
| DS154219 | 72 | 495 | BA_peak_9860 |

| | | | |
|---|---|---|---|
| DS154223 | 201 | 586 | BA_peak_9861 |
| DS154231 | 120 | 437 | BA_peak_9862 |
| DS154235 | 314 | 749 | BA_peak_9863 |
| DS154252 | 119 | 508 | BA_peak_9864 |
| DS154283 | 357 | 930 | BA_peak_9865 |
| DS154290 | 232 | 713 | BA_peak_9866 |
| DS154295 | 62 | 622 | BA_peak_9867 |
| DS154303 | 447 | 849 | BA_peak_9868 |
| DS154305 | 131 | 461 | BA_peak_9869 |
| DS154362 | 28 | 958 | BA_peak_9873 |
| DS154373 | 234 | 579 | BA_peak_9874 |
| DS154377 | 191 | 791 | BA_peak_9875 |
| DS154412 | 178 | 612 | BA_peak_9876 |
| DS154442 | 282 | 904 | BA_peak_9878 |
| DS154470 | 254 | 884 | BA_peak_9880 |
| DS154483 | 293 | 909 | BA_peak_9881 |
| DS154504 | 120 | 902 | BA_peak_9882 |
| DS154531 | 21 | 750 | BA_peak_9883 |
| DS154567 | 80 | 433 | BA_peak_9884 |
| DS154603 | 207 | 909 | BA_peak_9885 |
| DS154634 | 119 | 452 | BA_peak_9886 |
| DS154650 | 214 | 706 | BA_peak_9887 |
| DS154759 | 530 | 853 | BA_peak_9889 |
| DS154785 | 68 | 767 | BA_peak_9890 |
| DS154817 | 353 | 840 | BA_peak_9891 |
| DS154834 | 217 | 598 | BA_peak_9892 |
| DS154835 | 207 | 716 | BA_peak_9893 |
| DS154848 | 50 | 736 | BA_peak_9894 |
| DS154923 | 102 | 782 | BA_peak_9896 |
| DS154968 | 429 | 882 | BA_peak_9897 |
| DS154972 | 33 | 957 | BA_peak_9898 |
| DS154977 | 80 | 902 | BA_peak_9899 |
| DS154982 | 225 | 737 | BA_peak_9900 |
| DS154992 | 102 | 689 | BA_peak_9901 |
| DS155011 | 142 | 727 | BA_peak_9902 |
| DS155037 | 234 | 703 | BA_peak_9903 |
| DS155042 | 162 | 813 | BA_peak_9904 |
| DS155047 | 39 | 766 | BA_peak_9905 |
| DS155065 | 102 | 671 | BA_peak_9907 |
| DS155066 | 166 | 893 | BA_peak_9908 |
| DS155070 | 27 | 807 | BA_peak_9909 |
| DS155120 | 465 | 837 | BA_peak_9910 |
| DS155133 | 12 | 916 | BA_peak_9911 |
| DS155144 | 321 | 783 | BA_peak_9912 |
| DS155167 | 116 | 923 | BA_peak_9913 |
| DS155172 | 108 | 938 | BA_peak_9914 |
| DS155178 | 35 | 405 | BA_peak_9915 |

| | | | |
|---|---|---|---|
| DS155196 | 461 | 927 | BA_peak_9916 |
| DS155197 | 151 | 942 | BA_peak_9917 |
| DS155205 | 571 | 968 | BA_peak_9918 |
| DS155208 | 216 | 576 | BA_peak_9919 |
| DS155215 | 214 | 934 | BA_peak_9920 |
| DS155267 | 361 | 803 | BA_peak_9921 |
| DS155271 | 18 | 971 | BA_peak_9922 |
| DS155274 | 477 | 958 | BA_peak_9923 |
| DS155303 | 47 | 924 | BA_peak_9925 |
| DS155315 | 84 | 632 | BA_peak_9926 |
| DS155342 | 77 | 751 | BA_peak_9927 |
| DS155374 | 116 | 948 | BA_peak_9928 |
| DS155379 | 128 | 810 | BA_peak_9929 |
| DS155386 | 454 | 854 | BA_peak_9930 |
| DS155389 | 213 | 973 | BA_peak_9931 |
| DS155391 | 294 | 842 | BA_peak_9932 |
| DS155404 | 92 | 565 | BA_peak_9934 |
| DS155416 | 268 | 674 | BA_peak_9935 |
| DS155426 | 166 | 903 | BA_peak_9936 |
| DS155447 | 458 | 947 | BA_peak_9937 |
| DS155460 | 225 | 599 | BA_peak_9938 |
| DS155472 | 126 | 786 | BA_peak_9940 |
| DS155497 | 263 | 836 | BA_peak_9941 |
| DS155503 | 91 | 863 | BA_peak_9942 |
| DS155529 | 171 | 736 | BA_peak_9943 |
| DS155531 | 240 | 833 | BA_peak_9944 |
| DS155573 | 122 | 938 | BA_peak_9945 |
| DS155579 | 241 | 859 | BA_peak_9946 |
| DS155610 | 395 | 674 | BA_peak_9947 |
| DS155613 | 547 | 967 | BA_peak_9948 |
| DS155635 | 381 | 941 | BA_peak_9949 |
| DS155645 | 99 | 551 | BA_peak_9950 |
| DS155646 | 284 | 876 | BA_peak_9951 |
| DS155647 | 274 | 676 | BA_peak_9952 |
| DS155665 | 424 | 910 | BA_peak_9953 |
| DS155675 | 639 | 942 | BA_peak_9955 |
| DS155685 | 231 | 872 | BA_peak_9956 |
| DS155713 | 53 | 904 | BA_peak_9957 |
| DS155745 | 552 | 917 | BA_peak_9958 |
| DS155771 | 95 | 491 | BA_peak_9959 |
| DS155783 | 52 | 710 | BA_peak_9960 |
| DS155796 | 345 | 891 | BA_peak_9962 |
| DS155829 | 52 | 942 | BA_peak_9965 |
| DS155839 | 449 | 768 | BA_peak_9966 |
| DS155840 | 107 | 902 | BA_peak_9967 |
| DS155841 | 146 | 705 | BA_peak_9968 |
| DS155853 | 33 | 835 | BA_peak_9969 |

| | | | |
|---|---|---|---|
| DS155861 | 32 | 876 | BA_peak_9970 |
| DS155893 | 386 | 914 | BA_peak_9971 |
| DS155896 | 291 | 709 | BA_peak_9972 |
| DS155904 | 276 | 928 | BA_peak_9973 |
| DS155905 | 325 | 910 | BA_peak_9974 |
| DS155914 | 381 | 929 | BA_peak_9975 |
| DS155935 | 269 | 685 | BA_peak_9976 |
| DS155950 | 26 | 348 | BA_peak_9977 |
| DS155951 | 70 | 856 | BA_peak_9978 |
| DS155959 | 32 | 962 | BA_peak_9980 |
| DS155974 | 27 | 801 | BA_peak_9981 |
| DS155981 | 44 | 533 | BA_peak_9982 |
| DS155993 | 216 | 858 | BA_peak_9984 |
| DS156009 | 67 | 556 | BA_peak_9985 |
| DS156017 | 42 | 954 | BA_peak_9986 |
| DS156035 | 107 | 772 | BA_peak_9987 |
| DS156050 | 207 | 696 | BA_peak_9988 |
| DS156102 | 216 | 811 | BA_peak_9990 |
| DS156111 | 279 | 767 | BA_peak_9991 |
| DS156120 | 9 | 402 | BA_peak_9992 |
| DS156138 | 180 | 906 | BA_peak_9993 |
| DS156190 | 235 | 619 | BA_peak_9994 |
| DS156198 | 471 | 764 | BA_peak_9995 |
| DS156199 | 137 | 836 | BA_peak_9996 |
| DS156215 | 538 | 876 | BA_peak_9997 |
| DS156264 | 68 | 529 | BA_peak_9999 |
| DS156272 | 207 | 930 | BA_peak_10000 |
| DS156303 | 286 | 851 | BA_peak_10001 |
| DS156316 | 272 | 740 | BA_peak_10002 |
| DS156329 | 77 | 626 | BA_peak_10003 |
| DS156382 | 210 | 664 | BA_peak_10004 |
| DS156384 | 129 | 953 | BA_peak_10005 |
| DS156430 | 250 | 866 | BA_peak_10007 |
| DS156462 | 500 | 836 | BA_peak_10008 |
| DS156492 | 85 | 616 | BA_peak_10009 |
| DS156493 | 375 | 769 | BA_peak_10010 |
| DS156516 | 53 | 708 | BA_peak_10011 |
| DS156540 | 128 | 803 | BA_peak_10013 |
| DS156543 | 49 | 590 | BA_peak_10014 |
| DS156550 | 42 | 602 | BA_peak_10015 |
| DS156553 | 304 | 873 | BA_peak_10016 |
| DS156575 | 194 | 707 | BA_peak_10017 |
| DS156619 | 436 | 849 | BA_peak_10019 |
| DS156643 | 530 | 775 | BA_peak_10020 |
| DS156644 | 184 | 701 | BA_peak_10021 |
| DS156656 | 288 | 720 | BA_peak_10022 |
| DS156660 | 320 | 846 | BA_peak_10023 |

| | | | |
|---|---|---|---|
| DS156680 | 36 | 821 | BA_peak_10024 |
| DS156695 | 120 | 833 | BA_peak_10025 |
| DS156699 | 189 | 839 | BA_peak_10026 |
| DS156712 | 54 | 850 | BA_peak_10027 |
| DS156722 | 159 | 504 | BA_peak_10028 |
| DS156750 | 121 | 763 | BA_peak_10029 |
| DS156759 | 67 | 501 | BA_peak_10031 |
| DS156775 | 249 | 893 | BA_peak_10032 |
| DS156785 | 63 | 820 | BA_peak_10033 |
| DS156787 | 90 | 878 | BA_peak_10034 |
| DS156792 | 114 | 418 | BA_peak_10035 |
| DS156808 | 246 | 931 | BA_peak_10036 |
| DS156816 | 136 | 901 | BA_peak_10037 |
| DS156825 | 391 | 724 | BA_peak_10038 |
| DS156844 | 105 | 645 | BA_peak_10039 |
| DS156864 | 210 | 828 | BA_peak_10040 |
| DS156877 | 53 | 944 | BA_peak_10042 |
| DS156908 | 492 | 843 | BA_peak_10044 |
| DS156914 | 31 | 927 | BA_peak_10045 |
| DS156916 | 142 | 771 | BA_peak_10046 |
| DS156927 | 40 | 356 | BA_peak_10047 |
| DS156938 | 216 | 535 | BA_peak_10048 |
| DS156982 | 26 | 850 | BA_peak_10050 |
| DS156993 | 115 | 945 | BA_peak_10051 |
| DS157010 | 270 | 905 | BA_peak_10052 |
| DS157014 | 83 | 598 | BA_peak_10053 |
| DS157024 | 429 | 835 | BA_peak_10054 |
| DS157043 | 77 | 468 | BA_peak_10055 |
| DS157055 | 50 | 393 | BA_peak_10056 |
| DS157091 | 50 | 703 | BA_peak_10057 |
| DS157094 | 135 | 766 | BA_peak_10058 |
| DS157105 | 462 | 882 | BA_peak_10059 |
| DS157109 | 344 | 666 | BA_peak_10060 |
| DS157113 | 42 | 873 | BA_peak_10061 |
| DS157124 | 376 | 826 | BA_peak_10062 |
| DS157146 | 134 | 852 | BA_peak_10063 |
| DS157153 | 198 | 581 | BA_peak_10064 |
| DS157163 | 385 | 698 | BA_peak_10065 |
| DS157212 | 370 | 678 | BA_peak_10066 |
| DS157227 | 78 | 649 | BA_peak_10067 |
| DS157228 | 68 | 564 | BA_peak_10068 |
| DS157237 | 101 | 744 | BA_peak_10069 |
| DS157255 | 92 | 876 | BA_peak_10070 |
| DS157258 | 28 | 752 | BA_peak_10071 |
| DS157274 | 41 | 835 | BA_peak_10072 |
| DS157279 | 252 | 948 | BA_peak_10073 |
| DS157306 | 517 | 898 | BA_peak_10074 |

| | | | |
|---|---|---|---|
| DS157345 | 487 | 905 | BA_peak_10075 |
| DS157415 | 133 | 542 | BA_peak_10076 |
| DS157419 | 79 | 494 | BA_peak_10077 |
| DS157478 | 71 | 890 | BA_peak_10078 |
| DS157479 | 19 | 669 | BA_peak_10079 |
| DS157511 | 472 | 958 | BA_peak_10080 |
| DS157524 | 158 | 566 | BA_peak_10081 |
| DS157530 | 55 | 932 | BA_peak_10082 |
| DS157561 | 326 | 928 | BA_peak_10083 |
| DS157599 | 135 | 882 | BA_peak_10084 |
| DS157606 | 122 | 568 | BA_peak_10086 |
| DS157630 | 156 | 769 | BA_peak_10087 |
| DS157655 | 74 | 899 | BA_peak_10088 |
| DS157660 | 76 | 787 | BA_peak_10089 |
| DS157667 | 601 | 880 | BA_peak_10090 |
| DS157679 | 20 | 662 | BA_peak_10091 |
| DS157681 | 98 | 461 | BA_peak_10092 |
| DS157696 | 183 | 917 | BA_peak_10093 |
| DS157697 | 335 | 782 | BA_peak_10094 |
| DS157744 | 36 | 765 | BA_peak_10096 |
| DS157756 | 353 | 809 | BA_peak_10097 |
| DS157767 | 294 | 871 | BA_peak_10098 |
| DS157770 | 84 | 918 | BA_peak_10099 |
| DS157802 | 84 | 880 | BA_peak_10100 |
| DS157807 | 420 | 951 | BA_peak_10101 |
| DS157811 | 508 | 933 | BA_peak_10102 |
| DS157899 | 124 | 941 | BA_peak_10105 |
| DS157915 | 121 | 892 | BA_peak_10106 |
| DS157924 | 80 | 743 | BA_peak_10107 |
| DS157929 | 82 | 886 | BA_peak_10108 |
| DS157936 | 311 | 768 | BA_peak_10109 |
| DS157947 | 210 | 816 | BA_peak_10110 |
| DS157949 | 128 | 909 | BA_peak_10111 |
| DS157971 | 285 | 862 | BA_peak_10112 |
| DS157984 | 402 | 809 | BA_peak_10113 |
| DS158017 | 47 | 579 | BA_peak_10114 |
| DS158028 | 210 | 828 | BA_peak_10115 |
| DS158033 | 514 | 950 | BA_peak_10116 |
| DS158060 | 260 | 686 | BA_peak_10118 |
| DS158091 | 198 | 780 | BA_peak_10121 |
| DS158115 | 78 | 713 | BA_peak_10122 |
| DS158118 | 69 | 557 | BA_peak_10123 |
| DS158124 | 44 | 793 | BA_peak_10124 |
| DS158140 | 110 | 854 | BA_peak_10125 |
| DS158162 | 147 | 786 | BA_peak_10126 |
| DS158172 | 54 | 432 | BA_peak_10127 |
| DS158187 | 202 | 857 | BA_peak_10130 |

| | | | |
|---|---|---|---|
| DS158219 | 97 | 928 | BA_peak_10131 |
| DS158226 | 76 | 653 | BA_peak_10132 |
| DS158247 | 89 | 465 | BA_peak_10133 |
| DS158286 | 108 | 814 | BA_peak_10134 |
| DS158287 | 15 | 632 | BA_peak_10135 |
| DS158318 | 503 | 923 | BA_peak_10136 |
| DS158355 | 115 | 612 | BA_peak_10137 |
| DS158372 | 35 | 737 | BA_peak_10138 |
| DS158378 | 94 | 844 | BA_peak_10139 |
| DS158390 | 48 | 781 | BA_peak_10141 |
| DS158395 | 306 | 853 | BA_peak_10142 |
| DS158411 | 114 | 859 | BA_peak_10143 |
| DS158426 | 368 | 811 | BA_peak_10144 |
| DS158464 | 19 | 626 | BA_peak_10145 |
| DS158466 | 50 | 415 | BA_peak_10146 |
| DS158520 | 145 | 800 | BA_peak_10147 |
| DS158529 | 114 | 937 | BA_peak_10148 |
| DS158541 | 222 | 833 | BA_peak_10149 |
| DS158545 | 335 | 702 | BA_peak_10150 |
| DS158567 | 501 | 812 | BA_peak_10152 |
| DS158572 | 229 | 929 | BA_peak_10153 |
| DS158588 | 466 | 927 | BA_peak_10154 |
| DS158593 | 213 | 895 | BA_peak_10155 |
| DS158594 | 14 | 614 | BA_peak_10156 |
| DS158615 | 166 | 890 | BA_peak_10157 |
| DS158627 | 50 | 919 | BA_peak_10158 |
| DS158639 | 237 | 706 | BA_peak_10159 |
| DS158642 | 170 | 777 | BA_peak_10160 |
| DS158684 | 63 | 732 | BA_peak_10162 |
| DS158694 | 316 | 906 | BA_peak_10164 |
| DS158697 | 111 | 728 | BA_peak_10165 |
| DS158789 | 119 | 775 | BA_peak_10166 |
| DS158802 | 153 | 881 | BA_peak_10167 |
| DS158813 | 525 | 814 | BA_peak_10168 |
| DS158824 | 59 | 604 | BA_peak_10169 |
| DS158854 | 171 | 932 | BA_peak_10170 |
| DS158867 | 119 | 607 | BA_peak_10171 |
| DS158875 | 257 | 604 | BA_peak_10172 |
| DS158919 | 25 | 906 | BA_peak_10173 |
| DS158933 | 239 | 648 | BA_peak_10174 |
| DS158963 | 122 | 911 | BA_peak_10175 |
| DS158972 | 33 | 628 | BA_peak_10176 |
| DS158988 | 46 | 401 | BA_peak_10177 |
| DS158997 | 56 | 882 | BA_peak_10178 |
| DS159014 | 418 | 812 | BA_peak_10179 |
| DS159017 | 64 | 819 | BA_peak_10180 |
| DS159024 | 320 | 876 | BA_peak_10181 |

| | | | |
|---|---|---|---|
| DS159053 | 300 | 605 | BA_peak_10182 |
| DS159136 | 197 | 851 | BA_peak_10184 |
| DS159144 | 145 | 884 | BA_peak_10185 |
| DS159154 | 132 | 585 | BA_peak_10186 |
| DS159164 | 144 | 853 | BA_peak_10187 |
| DS159170 | 121 | 502 | BA_peak_10188 |
| DS159177 | 183 | 777 | BA_peak_10189 |
| DS159187 | 124 | 849 | BA_peak_10190 |
| DS159231 | 64 | 575 | BA_peak_10191 |
| DS159242 | 582 | 931 | BA_peak_10192 |
| DS159249 | 168 | 792 | BA_peak_10194 |
| DS159256 | 537 | 847 | BA_peak_10196 |
| DS159306 | 76 | 898 | BA_peak_10197 |
| DS159312 | 155 | 680 | BA_peak_10198 |
| DS159323 | 111 | 444 | BA_peak_10199 |
| DS159356 | 148 | 700 | BA_peak_10200 |
| DS159373 | 34 | 722 | BA_peak_10201 |
| DS159389 | 527 | 869 | BA_peak_10202 |
| DS159407 | 141 | 764 | BA_peak_10203 |
| DS159408 | 351 | 711 | BA_peak_10204 |
| DS159416 | 137 | 829 | BA_peak_10205 |
| DS159427 | 531 | 943 | BA_peak_10206 |
| DS159429 | 111 | 893 | BA_peak_10207 |
| DS159436 | 473 | 831 | BA_peak_10208 |
| DS159490 | 503 | 885 | BA_peak_10209 |
| DS159510 | 100 | 627 | BA_peak_10210 |
| DS159518 | 305 | 780 | BA_peak_10211 |
| DS159523 | 175 | 897 | BA_peak_10212 |
| DS159529 | 187 | 614 | BA_peak_10213 |
| DS159536 | 431 | 821 | BA_peak_10214 |
| DS159558 | 182 | 894 | BA_peak_10216 |
| DS159561 | 156 | 674 | BA_peak_10217 |
| DS159571 | 83 | 386 | BA_peak_10218 |
| DS159594 | 311 | 673 | BA_peak_10219 |
| DS159599 | 384 | 898 | BA_peak_10220 |
| DS159657 | 331 | 750 | BA_peak_10221 |
| DS159660 | 138 | 805 | BA_peak_10222 |
| DS159668 | 57 | 501 | BA_peak_10223 |
| DS159700 | 189 | 653 | BA_peak_10224 |
| DS159707 | 127 | 572 | BA_peak_10225 |
| DS159713 | 184 | 661 | BA_peak_10226 |
| DS159760 | 34 | 908 | BA_peak_10228 |
| DS159774 | 43 | 797 | BA_peak_10229 |
| DS159786 | 243 | 869 | BA_peak_10230 |
| DS159801 | 95 | 907 | BA_peak_10231 |
| DS159822 | 77 | 678 | BA_peak_10232 |
| DS159825 | 468 | 942 | BA_peak_10233 |

| | | | |
|---|---|---|---|
| DS159853 | 504 | 867 | BA_peak_10234 |
| DS159882 | 98 | 915 | BA_peak_10235 |
| DS159893 | 97 | 490 | BA_peak_10236 |
| DS159895 | 24 | 748 | BA_peak_10237 |
| DS159912 | 182 | 807 | BA_peak_10238 |
| DS159916 | 19 | 905 | BA_peak_10239 |
| DS159943 | 31 | 851 | BA_peak_10240 |
| DS159953 | 84 | 459 | BA_peak_10241 |
| DS159956 | 206 | 791 | BA_peak_10242 |
| DS159958 | 99 | 502 | BA_peak_10243 |
| DS159990 | 233 | 807 | BA_peak_10244 |
| DS160003 | 83 | 507 | BA_peak_10245 |
| DS160004 | 36 | 933 | BA_peak_10246 |
| DS160024 | 340 | 743 | BA_peak_10247 |
| DS160031 | 81 | 443 | BA_peak_10248 |
| DS160050 | 240 | 890 | BA_peak_10249 |
| DS160088 | 305 | 876 | BA_peak_10251 |
| DS160101 | 141 | 783 | BA_peak_10252 |
| DS160113 | 67 | 927 | BA_peak_10254 |
| DS160133 | 113 | 800 | BA_peak_10255 |
| DS160141 | 146 | 592 | BA_peak_10256 |
| DS160155 | 298 | 928 | BA_peak_10257 |
| DS160176 | 98 | 950 | BA_peak_10258 |
| DS160222 | 318 | 717 | BA_peak_10259 |
| DS160227 | 422 | 791 | BA_peak_10260 |
| DS160254 | 494 | 860 | BA_peak_10261 |
| DS160280 | 115 | 614 | BA_peak_10262 |
| DS160282 | 164 | 787 | BA_peak_10263 |
| DS160288 | 70 | 867 | BA_peak_10264 |
| DS160297 | 97 | 542 | BA_peak_10265 |
| DS160301 | 64 | 462 | BA_peak_10266 |
| DS160326 | 307 | 922 | BA_peak_10267 |
| DS160329 | 60 | 902 | BA_peak_10268 |
| DS160337 | 489 | 944 | BA_peak_10269 |
| DS160363 | 24 | 955 | BA_peak_10270 |
| DS160373 | 25 | 925 | BA_peak_10271 |
| DS160387 | 481 | 868 | BA_peak_10272 |
| DS160388 | 200 | 893 | BA_peak_10273 |
| DS160389 | 359 | 727 | BA_peak_10274 |
| DS160405 | 151 | 746 | BA_peak_10275 |
| DS160441 | 90 | 575 | BA_peak_10279 |
| DS160446 | 435 | 874 | BA_peak_10280 |
| DS160465 | 26 | 707 | BA_peak_10281 |
| DS160470 | 429 | 906 | BA_peak_10282 |
| DS160472 | 191 | 714 | BA_peak_10283 |
| DS160480 | 100 | 651 | BA_peak_10284 |
| DS160484 | 132 | 889 | BA_peak_10285 |

| | | | |
|---|---|---|---|
| DS160504 | 410 | 818 | BA_peak_10286 |
| DS160560 | 57 | 840 | BA_peak_10288 |
| DS160561 | 33 | 610 | BA_peak_10289 |
| DS160585 | 96 | 765 | BA_peak_10290 |
| DS160661 | 105 | 598 | BA_peak_10291 |
| DS160663 | 17 | 949 | BA_peak_10292 |
| DS160689 | 44 | 881 | BA_peak_10294 |
| DS160706 | 262 | 742 | BA_peak_10295 |
| DS160717 | 26 | 941 | BA_peak_10296 |
| DS160726 | 135 | 767 | BA_peak_10297 |
| DS160733 | 44 | 789 | BA_peak_10298 |
| DS160750 | 248 | 896 | BA_peak_10299 |
| DS160770 | 144 | 906 | BA_peak_10300 |
| DS160775 | 191 | 930 | BA_peak_10301 |
| DS160779 | 121 | 695 | BA_peak_10302 |
| DS160785 | 74 | 741 | BA_peak_10303 |
| DS160803 | 17 | 438 | BA_peak_10304 |
| DS160816 | 248 | 643 | BA_peak_10305 |
| DS160859 | 47 | 382 | BA_peak_10306 |
| DS160881 | 87 | 892 | BA_peak_10307 |
| DS160927 | 190 | 837 | BA_peak_10311 |
| DS160928 | 170 | 632 | BA_peak_10312 |
| DS160958 | 66 | 843 | BA_peak_10313 |
| DS160962 | 58 | 805 | BA_peak_10314 |
| DS160964 | 73 | 591 | BA_peak_10315 |
| DS161015 | 86 | 838 | BA_peak_10316 |
| DS161079 | 37 | 429 | BA_peak_10317 |
| DS161157 | 18 | 926 | BA_peak_10319 |
| DS161197 | 74 | 690 | BA_peak_10320 |
| DS161198 | 105 | 897 | BA_peak_10321 |
| DS161199 | 269 | 612 | BA_peak_10322 |
| DS161207 | 431 | 871 | BA_peak_10323 |
| DS161232 | 528 | 832 | BA_peak_10325 |
| DS161271 | 94 | 858 | BA_peak_10327 |
| DS161347 | 271 | 548 | BA_peak_10329 |
| DS161357 | 107 | 825 | BA_peak_10330 |
| DS161360 | 77 | 767 | BA_peak_10331 |
| DS161362 | 26 | 928 | BA_peak_10332 |
| DS161394 | 202 | 824 | BA_peak_10333 |
| DS161432 | 125 | 755 | BA_peak_10334 |
| DS161449 | 107 | 717 | BA_peak_10335 |
| DS161491 | 32 | 451 | BA_peak_10336 |
| DS161492 | 198 | 802 | BA_peak_10337 |
| DS161508 | 365 | 901 | BA_peak_10338 |
| DS161518 | 447 | 834 | BA_peak_10339 |
| DS161632 | 50 | 929 | BA_peak_10340 |
| DS161641 | 48 | 701 | BA_peak_10341 |

| | | | |
|---|---|---|---|
| DS161674 | 29 | 512 | BA_peak_10342 |
| DS161685 | 20 | 424 | BA_peak_10343 |
| DS161709 | 92 | 821 | BA_peak_10346 |
| DS161725 | 383 | 796 | BA_peak_10348 |
| DS161737 | 48 | 543 | BA_peak_10350 |
| DS161782 | 417 | 882 | BA_peak_10351 |
| DS161820 | 349 | 917 | BA_peak_10352 |
| DS161842 | 82 | 710 | BA_peak_10354 |
| DS161884 | 130 | 752 | BA_peak_10355 |
| DS161905 | 104 | 406 | BA_peak_10356 |
| DS161911 | 54 | 561 | BA_peak_10357 |
| DS161921 | 539 | 888 | BA_peak_10358 |
| DS161953 | 49 | 887 | BA_peak_10359 |
| DS161962 | 429 | 813 | BA_peak_10362 |
| DS161964 | 341 | 887 | BA_peak_10363 |
| DS161970 | 122 | 802 | BA_peak_10364 |
| DS162019 | 27 | 902 | BA_peak_10365 |
| DS162067 | 131 | 868 | BA_peak_10366 |
| DS162131 | 55 | 863 | BA_peak_10367 |
| DS162139 | 372 | 818 | BA_peak_10368 |
| DS162142 | 81 | 858 | BA_peak_10369 |
| DS162156 | 38 | 828 | BA_peak_10370 |
| DS162164 | 228 | 730 | BA_peak_10371 |
| DS162171 | 105 | 699 | BA_peak_10372 |
| DS162183 | 30 | 707 | BA_peak_10373 |
| DS162222 | 223 | 802 | BA_peak_10374 |
| DS162246 | 64 | 885 | BA_peak_10376 |
| DS162258 | 105 | 503 | BA_peak_10377 |
| DS162271 | 54 | 411 | BA_peak_10378 |
| DS162275 | 70 | 702 | BA_peak_10379 |
| DS162401 | 484 | 902 | BA_peak_10380 |
| DS162423 | 129 | 710 | BA_peak_10381 |
| DS162425 | 210 | 623 | BA_peak_10382 |
| DS162430 | 85 | 909 | BA_peak_10383 |
| DS162437 | 199 | 803 | BA_peak_10384 |
| DS162481 | 72 | 709 | BA_peak_10385 |
| DS162494 | 122 | 532 | BA_peak_10386 |
| DS162518 | 39 | 910 | BA_peak_10387 |
| DS162521 | 284 | 655 | BA_peak_10388 |
| DS162527 | 215 | 765 | BA_peak_10389 |
| DS162534 | 458 | 829 | BA_peak_10390 |
| DS162545 | 25 | 896 | BA_peak_10391 |
| DS162548 | 390 | 908 | BA_peak_10392 |
| DS162585 | 175 | 736 | BA_peak_10394 |
| DS162643 | 178 | 893 | BA_peak_10395 |
| DS162650 | 285 | 829 | BA_peak_10396 |
| DS162660 | 349 | 666 | BA_peak_10397 |

| | | | |
|---|---|---|---|
| DS162663 | 90 | 881 | BA_peak_10398 |
| DS162673 | 275 | 676 | BA_peak_10399 |
| DS162677 | 400 | 822 | BA_peak_10400 |
| DS162680 | 371 | 891 | BA_peak_10401 |
| DS162692 | 130 | 893 | BA_peak_10402 |
| DS162717 | 15 | 502 | BA_peak_10404 |
| DS162718 | 373 | 898 | BA_peak_10405 |
| DS162727 | 102 | 757 | BA_peak_10406 |
| DS162753 | 127 | 623 | BA_peak_10407 |
| DS162773 | 55 | 839 | BA_peak_10408 |
| DS162796 | 460 | 826 | BA_peak_10409 |
| DS162806 | 449 | 789 | BA_peak_10410 |
| DS162825 | 77 | 912 | BA_peak_10412 |
| DS162876 | 90 | 610 | BA_peak_10414 |
| DS162904 | 109 | 755 | BA_peak_10415 |
| DS162978 | 191 | 829 | BA_peak_10417 |
| DS162994 | 26 | 748 | BA_peak_10418 |
| DS163013 | 34 | 573 | BA_peak_10420 |
| DS163024 | 138 | 616 | BA_peak_10421 |
| DS163031 | 49 | 547 | BA_peak_10422 |
| DS163041 | 80 | 904 | BA_peak_10423 |
| DS163043 | 103 | 781 | BA_peak_10424 |
| DS163050 | 53 | 713 | BA_peak_10425 |
| DS163103 | 121 | 485 | BA_peak_10427 |
| DS163105 | 368 | 819 | BA_peak_10428 |
| DS163129 | 233 | 864 | BA_peak_10429 |
| DS163130 | 43 | 888 | BA_peak_10430 |
| DS163131 | 158 | 514 | BA_peak_10431 |
| DS163150 | 143 | 452 | BA_peak_10433 |
| DS163159 | 293 | 708 | BA_peak_10434 |
| DS163176 | 328 | 671 | BA_peak_10435 |
| DS163202 | 37 | 826 | BA_peak_10437 |
| DS163207 | 143 | 763 | BA_peak_10438 |
| DS163223 | 143 | 459 | BA_peak_10439 |
| DS163229 | 68 | 639 | BA_peak_10440 |
| DS163261 | 297 | 680 | BA_peak_10441 |
| DS163279 | 61 | 890 | BA_peak_10442 |
| DS163282 | 39 | 777 | BA_peak_10443 |
| DS163372 | 358 | 742 | BA_peak_10444 |
| DS163399 | 43 | 471 | BA_peak_10446 |
| DS163407 | 123 | 686 | BA_peak_10447 |
| DS163412 | 247 | 848 | BA_peak_10449 |
| DS163424 | 193 | 815 | BA_peak_10450 |
| DS163452 | 131 | 651 | BA_peak_10452 |
| DS163470 | 193 | 872 | BA_peak_10453 |
| DS163503 | 41 | 904 | BA_peak_10454 |
| DS163504 | 93 | 617 | BA_peak_10455 |

| | | | |
|---|---|---|---|
| DS163541 | 144 | 569 | BA_peak_10456 |
| DS163549 | 303 | 754 | BA_peak_10457 |
| DS163576 | 65 | 807 | BA_peak_10458 |
| DS163627 | 252 | 901 | BA_peak_10460 |
| DS163637 | 212 | 623 | BA_peak_10461 |
| DS163642 | 359 | 811 | BA_peak_10462 |
| DS163644 | 177 | 765 | BA_peak_10463 |
| DS163665 | 14 | 778 | BA_peak_10464 |
| DS163726 | 117 | 759 | BA_peak_10465 |
| DS163735 | 483 | 868 | BA_peak_10466 |
| DS163737 | 230 | 659 | BA_peak_10467 |
| DS163743 | 398 | 725 | BA_peak_10468 |
| DS163755 | 232 | 580 | BA_peak_10469 |
| DS163759 | 64 | 538 | BA_peak_10470 |
| DS163782 | 278 | 612 | BA_peak_10471 |
| DS163812 | 314 | 768 | BA_peak_10472 |
| DS163841 | 33 | 884 | BA_peak_10473 |
| DS163862 | 382 | 711 | BA_peak_10474 |
| DS163870 | 233 | 816 | BA_peak_10475 |
| DS163919 | 165 | 902 | BA_peak_10476 |
| DS163945 | 43 | 495 | BA_peak_10477 |
| DS163956 | 71 | 602 | BA_peak_10478 |
| DS163958 | 207 | 691 | BA_peak_10479 |
| DS163964 | 88 | 550 | BA_peak_10481 |
| DS163970 | 87 | 882 | BA_peak_10482 |
| DS163974 | 189 | 842 | BA_peak_10483 |
| DS164014 | 169 | 897 | BA_peak_10484 |
| DS164065 | 112 | 736 | BA_peak_10486 |
| DS164090 | 413 | 856 | BA_peak_10487 |
| DS164102 | 121 | 701 | BA_peak_10488 |
| DS164132 | 98 | 554 | BA_peak_10490 |
| DS164140 | 210 | 817 | BA_peak_10491 |
| DS164166 | 316 | 764 | BA_peak_10492 |
| DS164178 | 237 | 795 | BA_peak_10493 |
| DS164231 | 141 | 717 | BA_peak_10495 |
| DS164238 | 140 | 692 | BA_peak_10496 |
| DS164257 | 29 | 786 | BA_peak_10497 |
| DS164262 | 307 | 858 | BA_peak_10498 |
| DS164267 | 295 | 677 | BA_peak_10499 |
| DS164268 | 184 | 901 | BA_peak_10500 |
| DS164277 | 113 | 782 | BA_peak_10502 |
| DS164334 | 393 | 885 | BA_peak_10503 |
| DS164386 | 215 | 814 | BA_peak_10504 |
| DS164389 | 28 | 870 | BA_peak_10505 |
| DS164391 | 215 | 687 | BA_peak_10506 |
| DS164400 | 140 | 777 | BA_peak_10507 |
| DS164405 | 58 | 403 | BA_peak_10508 |

| | | | |
|---|---|---|---|
| DS164408 | 155 | 697 | BA_peak_10509 |
| DS164411 | 106 | 721 | BA_peak_10510 |
| DS164442 | 231 | 910 | BA_peak_10512 |
| DS164454 | 117 | 779 | BA_peak_10513 |
| DS164458 | 175 | 459 | BA_peak_10514 |
| DS164462 | 103 | 770 | BA_peak_10515 |
| DS164467 | 132 | 846 | BA_peak_10516 |
| DS164484 | 352 | 929 | BA_peak_10517 |
| DS164492 | 217 | 492 | BA_peak_10518 |
| DS164497 | 54 | 844 | BA_peak_10519 |
| DS164573 | 46 | 381 | BA_peak_10521 |
| DS164575 | 162 | 609 | BA_peak_10522 |
| DS164579 | 170 | 728 | BA_peak_10523 |
| DS164602 | 53 | 560 | BA_peak_10524 |
| DS164637 | 189 | 805 | BA_peak_10525 |
| DS164651 | 24 | 779 | BA_peak_10526 |
| DS164678 | 381 | 775 | BA_peak_10527 |
| DS164703 | 176 | 762 | BA_peak_10528 |
| DS164760 | 151 | 528 | BA_peak_10530 |
| DS164762 | 507 | 874 | BA_peak_10531 |
| DS164781 | 36 | 876 | BA_peak_10533 |
| DS164782 | 18 | 850 | BA_peak_10534 |
| DS164839 | 163 | 902 | BA_peak_10536 |
| DS164852 | 67 | 670 | BA_peak_10537 |
| DS164875 | 167 | 612 | BA_peak_10538 |
| DS164884 | 187 | 806 | BA_peak_10539 |
| DS164908 | 92 | 521 | BA_peak_10541 |
| DS164909 | 308 | 838 | BA_peak_10542 |
| DS164922 | 401 | 805 | BA_peak_10544 |
| DS164939 | 37 | 917 | BA_peak_10545 |
| DS164943 | 97 | 568 | BA_peak_10546 |
| DS164944 | 38 | 889 | BA_peak_10547 |
| DS164946 | 263 | 615 | BA_peak_10548 |
| DS164964 | 198 | 638 | BA_peak_10550 |
| DS164977 | 131 | 660 | BA_peak_10551 |
| DS164978 | 239 | 705 | BA_peak_10552 |
| DS164980 | 73 | 537 | BA_peak_10553 |
| DS164983 | 324 | 799 | BA_peak_10554 |
| DS165019 | 140 | 593 | BA_peak_10555 |
| DS165030 | 263 | 850 | BA_peak_10556 |
| DS165038 | 102 | 910 | BA_peak_10557 |
| DS165091 | 105 | 476 | BA_peak_10558 |
| DS165096 | 88 | 776 | BA_peak_10559 |
| DS165108 | 166 | 795 | BA_peak_10560 |
| DS165111 | 414 | 711 | BA_peak_10561 |
| DS165167 | 538 | 860 | BA_peak_10563 |
| DS165197 | 490 | 866 | BA_peak_10564 |

| | | | |
|---|---|---|---|
| DS165222 | 37 | 483 | BA_peak_10565 |
| DS165227 | 480 | 905 | BA_peak_10566 |
| DS165230 | 93 | 822 | BA_peak_10567 |
| DS165252 | 27 | 677 | BA_peak_10568 |
| DS165263 | 77 | 618 | BA_peak_10569 |
| DS165296 | 171 | 817 | BA_peak_10570 |
| DS165300 | 210 | 879 | BA_peak_10571 |
| DS165317 | 47 | 741 | BA_peak_10572 |
| DS165324 | 246 | 765 | BA_peak_10573 |
| DS165328 | 47 | 828 | BA_peak_10574 |
| DS165358 | 104 | 782 | BA_peak_10575 |
| DS165373 | 179 | 876 | BA_peak_10577 |
| DS165374 | 88 | 754 | BA_peak_10578 |
| DS165376 | 184 | 868 | BA_peak_10579 |
| DS165402 | 21 | 810 | BA_peak_10580 |
| DS165406 | 119 | 702 | BA_peak_10581 |
| DS165434 | 450 | 907 | BA_peak_10582 |
| DS165435 | 201 | 881 | BA_peak_10583 |
| DS165479 | 54 | 885 | BA_peak_10584 |
| DS165497 | 35 | 799 | BA_peak_10585 |
| DS165519 | 140 | 861 | BA_peak_10586 |
| DS165554 | 347 | 702 | BA_peak_10587 |
| DS165555 | 135 | 501 | BA_peak_10588 |
| DS165566 | 191 | 855 | BA_peak_10589 |
| DS165583 | 184 | 870 | BA_peak_10590 |
| DS165614 | 16 | 409 | BA_peak_10591 |
| DS165629 | 41 | 910 | BA_peak_10592 |
| DS165666 | 268 | 668 | BA_peak_10593 |
| DS165673 | 47 | 795 | BA_peak_10594 |
| DS165720 | 51 | 408 | BA_peak_10596 |
| DS165731 | 140 | 839 | BA_peak_10597 |
| DS165738 | 88 | 513 | BA_peak_10599 |
| DS165752 | 28 | 516 | BA_peak_10602 |
| DS165762 | 290 | 757 | BA_peak_10603 |
| DS165768 | 129 | 848 | BA_peak_10604 |
| DS165776 | 174 | 560 | BA_peak_10605 |
| DS165780 | 150 | 525 | BA_peak_10606 |
| DS165795 | 282 | 653 | BA_peak_10607 |
| DS165798 | 186 | 905 | BA_peak_10608 |
| DS165807 | 90 | 430 | BA_peak_10609 |
| DS165810 | 36 | 866 | BA_peak_10610 |
| DS165832 | 53 | 741 | BA_peak_10612 |
| DS165837 | 21 | 873 | BA_peak_10613 |
| DS165846 | 41 | 672 | BA_peak_10614 |
| DS165862 | 297 | 684 | BA_peak_10615 |
| DS165874 | 32 | 545 | BA_peak_10616 |
| DS165880 | 435 | 897 | BA_peak_10617 |

| | | | |
|---|---|---|---|
| DS165925 | 93 | 805 | BA_peak_10618 |
| DS165930 | 67 | 731 | BA_peak_10619 |
| DS165937 | 49 | 853 | BA_peak_10620 |
| DS165951 | 33 | 880 | BA_peak_10621 |
| DS165956 | 106 | 725 | BA_peak_10622 |
| DS165981 | 41 | 772 | BA_peak_10623 |
| DS166007 | 40 | 829 | BA_peak_10624 |
| DS166027 | 134 | 844 | BA_peak_10625 |
| DS166042 | 161 | 745 | BA_peak_10626 |
| DS166059 | 113 | 815 | BA_peak_10627 |
| DS166066 | 64 | 873 | BA_peak_10629 |
| DS166070 | 79 | 905 | BA_peak_10630 |
| DS166086 | 414 | 819 | BA_peak_10631 |
| DS166100 | 26 | 715 | BA_peak_10632 |
| DS166109 | 208 | 860 | BA_peak_10633 |
| DS166143 | 235 | 642 | BA_peak_10635 |
| DS166162 | 218 | 774 | BA_peak_10636 |
| DS166199 | 26 | 885 | BA_peak_10637 |
| DS166212 | 431 | 853 | BA_peak_10639 |
| DS166213 | 369 | 813 | BA_peak_10640 |
| DS166214 | 120 | 650 | BA_peak_10641 |
| DS166237 | 297 | 915 | BA_peak_10642 |
| DS166259 | 84 | 720 | BA_peak_10643 |
| DS166269 | 140 | 831 | BA_peak_10644 |
| DS166280 | 262 | 689 | BA_peak_10645 |
| DS166304 | 29 | 791 | BA_peak_10646 |
| DS166321 | 132 | 795 | BA_peak_10647 |
| DS166335 | 119 | 655 | BA_peak_10648 |
| DS166342 | 41 | 389 | BA_peak_10649 |
| DS166377 | 32 | 776 | BA_peak_10650 |
| DS166421 | 4 | 818 | BA_peak_10652 |
| DS166428 | 520 | 894 | BA_peak_10653 |
| DS166434 | 47 | 759 | BA_peak_10654 |
| DS166452 | 190 | 762 | BA_peak_10655 |
| DS166460 | 187 | 847 | BA_peak_10658 |
| DS166466 | 388 | 872 | BA_peak_10659 |
| DS166485 | 55 | 544 | BA_peak_10660 |
| DS166499 | 184 | 776 | BA_peak_10661 |
| DS166502 | 171 | 653 | BA_peak_10662 |
| DS166509 | 198 | 521 | BA_peak_10663 |
| DS166517 | 479 | 858 | BA_peak_10664 |
| DS166524 | 359 | 833 | BA_peak_10665 |
| DS166525 | 100 | 893 | BA_peak_10666 |
| DS166538 | 117 | 719 | BA_peak_10667 |
| DS166562 | 215 | 792 | BA_peak_10669 |
| DS166569 | 231 | 661 | BA_peak_10670 |
| DS166589 | 73 | 590 | BA_peak_10671 |

| | | | |
|---|---|---|---|
| DS166605 | 39 | 849 | BA_peak_10672 |
| DS166626 | 56 | 863 | BA_peak_10673 |
| DS166628 | 234 | 864 | BA_peak_10674 |
| DS166654 | 396 | 797 | BA_peak_10675 |
| DS166655 | 291 | 675 | BA_peak_10676 |
| DS166661 | 217 | 836 | BA_peak_10677 |
| DS166687 | 94 | 544 | BA_peak_10678 |
| DS166706 | 209 | 855 | BA_peak_10679 |
| DS166740 | 42 | 525 | BA_peak_10680 |
| DS166741 | 153 | 836 | BA_peak_10681 |
| DS166757 | 73 | 704 | BA_peak_10682 |
| DS166796 | 131 | 799 | BA_peak_10683 |
| DS166818 | 70 | 833 | BA_peak_10684 |
| DS166820 | 110 | 653 | BA_peak_10685 |
| DS166839 | 103 | 502 | BA_peak_10686 |
| DS166848 | 105 | 867 | BA_peak_10687 |
| DS166872 | 140 | 565 | BA_peak_10688 |
| DS166884 | 100 | 743 | BA_peak_10689 |
| DS166918 | 159 | 625 | BA_peak_10691 |
| DS166919 | 116 | 889 | BA_peak_10692 |
| DS166922 | 64 | 806 | BA_peak_10693 |
| DS166930 | 360 | 882 | BA_peak_10694 |
| DS166945 | 125 | 451 | BA_peak_10695 |
| DS166980 | 412 | 855 | BA_peak_10696 |
| DS166998 | 82 | 758 | BA_peak_10697 |
| DS167029 | 25 | 490 | BA_peak_10698 |
| DS167053 | 242 | 831 | BA_peak_10699 |
| DS167158 | 161 | 780 | BA_peak_10701 |
| DS167170 | 95 | 492 | BA_peak_10702 |
| DS167174 | 74 | 883 | BA_peak_10703 |
| DS167178 | 157 | 879 | BA_peak_10704 |
| DS167203 | 62 | 545 | BA_peak_10705 |
| DS167211 | 487 | 886 | BA_peak_10706 |
| DS167259 | 200 | 817 | BA_peak_10708 |
| DS167272 | 186 | 794 | BA_peak_10709 |
| DS167289 | 176 | 867 | BA_peak_10710 |
| DS167293 | 34 | 879 | BA_peak_10711 |
| DS167305 | 278 | 766 | BA_peak_10712 |
| DS167307 | 272 | 735 | BA_peak_10713 |
| DS167311 | 16 | 872 | BA_peak_10714 |
| DS167315 | 60 | 862 | BA_peak_10715 |
| DS167318 | 66 | 902 | BA_peak_10716 |
| DS167384 | 44 | 573 | BA_peak_10717 |
| DS167401 | 134 | 534 | BA_peak_10718 |
| DS167407 | 185 | 798 | BA_peak_10719 |
| DS167471 | 227 | 699 | BA_peak_10723 |
| DS167478 | 56 | 797 | BA_peak_10724 |

| | | | |
|---|---|---|---|
| DS167512 | 244 | 600 | BA_peak_10725 |
| DS167520 | 207 | 765 | BA_peak_10726 |
| DS167556 | 142 | 609 | BA_peak_10727 |
| DS167565 | 396 | 853 | BA_peak_10728 |
| DS167566 | 138 | 591 | BA_peak_10729 |
| DS167591 | 60 | 700 | BA_peak_10730 |
| DS167595 | 394 | 841 | BA_peak_10731 |
| DS167598 | 350 | 669 | BA_peak_10732 |
| DS167606 | 59 | 784 | BA_peak_10733 |
| DS167646 | 137 | 857 | BA_peak_10735 |
| DS167696 | 155 | 639 | BA_peak_10737 |
| DS167703 | 142 | 817 | BA_peak_10738 |
| DS167722 | 446 | 813 | BA_peak_10739 |
| DS167794 | 261 | 844 | BA_peak_10740 |
| DS167799 | 237 | 773 | BA_peak_10741 |
| DS167801 | 186 | 759 | BA_peak_10742 |
| DS167836 | 50 | 833 | BA_peak_10743 |
| DS167837 | 65 | 444 | BA_peak_10744 |
| DS167841 | 106 | 881 | BA_peak_10745 |
| DS167846 | 307 | 897 | BA_peak_10746 |
| DS167973 | 138 | 581 | BA_peak_10748 |
| DS167975 | 56 | 851 | BA_peak_10749 |
| DS167994 | 143 | 454 | BA_peak_10751 |
| DS168050 | 67 | 868 | BA_peak_10752 |
| DS168074 | 124 | 604 | BA_peak_10753 |
| DS168099 | 124 | 697 | BA_peak_10754 |
| DS168101 | 173 | 759 | BA_peak_10755 |
| DS168103 | 232 | 850 | BA_peak_10756 |
| DS168123 | 146 | 853 | BA_peak_10757 |
| DS168138 | 54 | 392 | BA_peak_10758 |
| DS168140 | 62 | 885 | BA_peak_10759 |
| DS168169 | 307 | 779 | BA_peak_10760 |
| DS168183 | 379 | 739 | BA_peak_10762 |
| DS168191 | 69 | 762 | BA_peak_10763 |
| DS168209 | 13 | 849 | BA_peak_10764 |
| DS168234 | 96 | 596 | BA_peak_10765 |
| DS168241 | 242 | 867 | BA_peak_10766 |
| DS168250 | 148 | 879 | BA_peak_10767 |
| DS168269 | 171 | 720 | BA_peak_10768 |
| DS168273 | 331 | 736 | BA_peak_10769 |
| DS168276 | 64 | 805 | BA_peak_10770 |
| DS168282 | 27 | 837 | BA_peak_10771 |
| DS168286 | 63 | 878 | BA_peak_10772 |
| DS168339 | 301 | 768 | BA_peak_10775 |
| DS168346 | 369 | 820 | BA_peak_10776 |
| DS168351 | 86 | 565 | BA_peak_10777 |
| DS168367 | 93 | 827 | BA_peak_10778 |

| | | | |
|---|---|---|---|
| DS168382 | 268 | 660 | BA_peak_10779 |
| DS168385 | 187 | 881 | BA_peak_10780 |
| DS168387 | 420 | 866 | BA_peak_10781 |
| DS168390 | 151 | 859 | BA_peak_10782 |
| DS168393 | 44 | 865 | BA_peak_10783 |
| DS168394 | 24 | 522 | BA_peak_10784 |
| DS168402 | 124 | 808 | BA_peak_10785 |
| DS168425 | 50 | 688 | BA_peak_10786 |
| DS168468 | 91 | 863 | BA_peak_10787 |
| DS168490 | 36 | 650 | BA_peak_10788 |
| DS168491 | 167 | 835 | BA_peak_10789 |
| DS168506 | 128 | 560 | BA_peak_10790 |
| DS168521 | 145 | 637 | BA_peak_10791 |
| DS168532 | 131 | 692 | BA_peak_10792 |
| DS168534 | 461 | 791 | BA_peak_10793 |
| DS168547 | 190 | 777 | BA_peak_10794 |
| DS168571 | 61 | 739 | BA_peak_10795 |
| DS168574 | 165 | 494 | BA_peak_10796 |
| DS168593 | 79 | 661 | BA_peak_10797 |
| DS168607 | 46 | 752 | BA_peak_10798 |
| DS168610 | 193 | 609 | BA_peak_10799 |
| DS168619 | 141 | 767 | BA_peak_10800 |
| DS168623 | 38 | 748 | BA_peak_10801 |
| DS168630 | 87 | 871 | BA_peak_10802 |
| DS168689 | 481 | 858 | BA_peak_10803 |
| DS168710 | 346 | 731 | BA_peak_10804 |
| DS168717 | 273 | 591 | BA_peak_10805 |
| DS168718 | 333 | 716 | BA_peak_10806 |
| DS168727 | 177 | 564 | BA_peak_10807 |
| DS168728 | 25 | 605 | BA_peak_10808 |
| DS168736 | 73 | 827 | BA_peak_10809 |
| DS168774 | 73 | 756 | BA_peak_10811 |
| DS168828 | 359 | 838 | BA_peak_10812 |
| DS168832 | 17 | 488 | BA_peak_10813 |
| DS168863 | 144 | 807 | BA_peak_10814 |
| DS168942 | 391 | 713 | BA_peak_10816 |
| DS168966 | 539 | 838 | BA_peak_10818 |
| DS168993 | 114 | 789 | BA_peak_10819 |
| DS169002 | 61 | 886 | BA_peak_10820 |
| DS169020 | 49 | 731 | BA_peak_10821 |
| DS169023 | 52 | 624 | BA_peak_10822 |
| DS169042 | 211 | 710 | BA_peak_10823 |
| DS169062 | 357 | 755 | BA_peak_10824 |
| DS169100 | 423 | 744 | BA_peak_10825 |
| DS169136 | 45 | 703 | BA_peak_10826 |
| DS169145 | 63 | 539 | BA_peak_10827 |
| DS169160 | 127 | 661 | BA_peak_10828 |

| | | | |
|---|---|---|---|
| DS169167 | 27 | 848 | BA_peak_10829 |
| DS169170 | 144 | 558 | BA_peak_10830 |
| DS169190 | 153 | 783 | BA_peak_10831 |
| DS169219 | 132 | 518 | BA_peak_10832 |
| DS169232 | 162 | 853 | BA_peak_10833 |
| DS169235 | 114 | 885 | BA_peak_10834 |
| DS169251 | 388 | 825 | BA_peak_10835 |
| DS169286 | 40 | 786 | BA_peak_10836 |
| DS169289 | 111 | 623 | BA_peak_10837 |
| DS169291 | 101 | 771 | BA_peak_10838 |
| DS169305 | 166 | 643 | BA_peak_10839 |
| DS169333 | 162 | 524 | BA_peak_10840 |
| DS169365 | 252 | 756 | BA_peak_10841 |
| DS169378 | 138 | 774 | BA_peak_10842 |
| DS169423 | 184 | 724 | BA_peak_10843 |
| DS169449 | 117 | 847 | BA_peak_10844 |
| DS169454 | 240 | 669 | BA_peak_10845 |
| DS169488 | 277 | 786 | BA_peak_10846 |
| DS169495 | 271 | 801 | BA_peak_10847 |
| DS169496 | 249 | 832 | BA_peak_10848 |
| DS169511 | 35 | 873 | BA_peak_10849 |
| DS169514 | 318 | 642 | BA_peak_10850 |
| DS169537 | 37 | 678 | BA_peak_10851 |
| DS169574 | 46 | 646 | BA_peak_10853 |
| DS169583 | 328 | 817 | BA_peak_10854 |
| DS169591 | 32 | 817 | BA_peak_10855 |
| DS169598 | 224 | 618 | BA_peak_10856 |
| DS169609 | 193 | 793 | BA_peak_10857 |
| DS169645 | 53 | 534 | BA_peak_10858 |
| DS169692 | 76 | 698 | BA_peak_10860 |
| DS169747 | 50 | 745 | BA_peak_10861 |
| DS169758 | 265 | 845 | BA_peak_10862 |
| DS169770 | 376 | 821 | BA_peak_10865 |
| DS169783 | 93 | 846 | BA_peak_10866 |
| DS169805 | 15 | 873 | BA_peak_10868 |
| DS169807 | 23 | 700 | BA_peak_10869 |
| DS169835 | 379 | 835 | BA_peak_10870 |
| DS169853 | 302 | 654 | BA_peak_10871 |
| DS169854 | 80 | 747 | BA_peak_10872 |
| DS169856 | 400 | 748 | BA_peak_10873 |
| DS169860 | 88 | 656 | BA_peak_10874 |
| DS169865 | 369 | 823 | BA_peak_10875 |
| DS169877 | 203 | 586 | BA_peak_10876 |
| DS169905 | 177 | 818 | BA_peak_10877 |
| DS169931 | 242 | 697 | BA_peak_10878 |
| DS169956 | 144 | 488 | BA_peak_10880 |
| DS169964 | 29 | 843 | BA_peak_10881 |

| | | | |
|---|---|---|---|
| DS169977 | 83 | 679 | BA_peak_10882 |
| DS169980 | 162 | 602 | BA_peak_10883 |
| DS169994 | 96 | 803 | BA_peak_10884 |
| DS170011 | 144 | 674 | BA_peak_10885 |
| DS170012 | 382 | 746 | BA_peak_10886 |
| DS170027 | 55 | 870 | BA_peak_10888 |
| DS170053 | 275 | 762 | BA_peak_10890 |
| DS170074 | 32 | 615 | BA_peak_10891 |
| DS170078 | 93 | 817 | BA_peak_10892 |
| DS170108 | 83 | 827 | BA_peak_10893 |
| DS170113 | 182 | 522 | BA_peak_10894 |
| DS170115 | 136 | 878 | BA_peak_10895 |
| DS170158 | 59 | 728 | BA_peak_10897 |
| DS170161 | 214 | 852 | BA_peak_10898 |
| DS170180 | 103 | 449 | BA_peak_10899 |
| DS170185 | 95 | 781 | BA_peak_10900 |
| DS170221 | 21 | 773 | BA_peak_10901 |
| DS170249 | 378 | 786 | BA_peak_10902 |
| DS170265 | 88 | 642 | BA_peak_10903 |
| DS170284 | 160 | 762 | BA_peak_10905 |
| DS170288 | 210 | 717 | BA_peak_10906 |
| DS170322 | 283 | 839 | BA_peak_10907 |
| DS170352 | 85 | 826 | BA_peak_10908 |
| DS170388 | 110 | 771 | BA_peak_10909 |
| DS170401 | 31 | 760 | BA_peak_10910 |
| DS170417 | 168 | 722 | BA_peak_10911 |
| DS170442 | 60 | 692 | BA_peak_10912 |
| DS170486 | 44 | 553 | BA_peak_10915 |
| DS170488 | 14 | 432 | BA_peak_10916 |
| DS170508 | 48 | 814 | BA_peak_10917 |
| DS170516 | 9 | 873 | BA_peak_10918 |
| DS170526 | 47 | 771 | BA_peak_10919 |
| DS170532 | 107 | 855 | BA_peak_10920 |
| DS170558 | 335 | 801 | BA_peak_10921 |
| DS170582 | 411 | 757 | BA_peak_10922 |
| DS170606 | 58 | 431 | BA_peak_10923 |
| DS170621 | 100 | 514 | BA_peak_10924 |
| DS170631 | 126 | 824 | BA_peak_10926 |
| DS170681 | 144 | 829 | BA_peak_10928 |
| DS170698 | 140 | 786 | BA_peak_10929 |
| DS170721 | 139 | 651 | BA_peak_10930 |
| DS170727 | 221 | 847 | BA_peak_10931 |
| DS170856 | 248 | 688 | BA_peak_10933 |
| DS170863 | 35 | 828 | BA_peak_10934 |
| DS170887 | 58 | 801 | BA_peak_10935 |
| DS170915 | 204 | 547 | BA_peak_10936 |
| DS170938 | 101 | 805 | BA_peak_10937 |

| | | | |
|---|---|---|---|
| DS170941 | 99 | 601 | BA_peak_10938 |
| DS170954 | 75 | 745 | BA_peak_10939 |
| DS170962 | 438 | 831 | BA_peak_10940 |
| DS170984 | 246 | 763 | BA_peak_10941 |
| DS171001 | 189 | 496 | BA_peak_10942 |
| DS171011 | 317 | 818 | BA_peak_10943 |
| DS171047 | 21 | 857 | BA_peak_10944 |
| DS171074 | 112 | 832 | BA_peak_10945 |
| DS171115 | 198 | 751 | BA_peak_10946 |
| DS171156 | 218 | 526 | BA_peak_10947 |
| DS171172 | 70 | 552 | BA_peak_10948 |
| DS171175 | 32 | 688 | BA_peak_10949 |
| DS171205 | 92 | 847 | BA_peak_10950 |
| DS171206 | 127 | 746 | BA_peak_10951 |
| DS171209 | 73 | 835 | BA_peak_10952 |
| DS171213 | 227 | 808 | BA_peak_10953 |
| DS171219 | 195 | 636 | BA_peak_10954 |
| DS171224 | 458 | 863 | BA_peak_10955 |
| DS171259 | 363 | 764 | BA_peak_10956 |
| DS171273 | 120 | 674 | BA_peak_10957 |
| DS171297 | 398 | 799 | BA_peak_10958 |
| DS171298 | 69 | 442 | BA_peak_10959 |
| DS171303 | 210 | 693 | BA_peak_10960 |
| DS171318 | 61 | 599 | BA_peak_10961 |
| DS171329 | 132 | 568 | BA_peak_10963 |
| DS171334 | 313 | 823 | BA_peak_10964 |
| DS171349 | 37 | 800 | BA_peak_10965 |
| DS171356 | 168 | 677 | BA_peak_10966 |
| DS171357 | 98 | 817 | BA_peak_10967 |
| DS171378 | 470 | 779 | BA_peak_10968 |
| DS171393 | 182 | 512 | BA_peak_10969 |
| DS171406 | 90 | 835 | BA_peak_10970 |
| DS171409 | 223 | 783 | BA_peak_10971 |
| DS171431 | 34 | 833 | BA_peak_10972 |
| DS171475 | 294 | 691 | BA_peak_10973 |
| DS171484 | 63 | 782 | BA_peak_10974 |
| DS171513 | 274 | 854 | BA_peak_10975 |
| DS171524 | 146 | 787 | BA_peak_10976 |
| DS171557 | 52 | 808 | BA_peak_10977 |
| DS171595 | 22 | 488 | BA_peak_10979 |
| DS171622 | 231 | 837 | BA_peak_10980 |
| DS171625 | 200 | 721 | BA_peak_10981 |
| DS171629 | 37 | 736 | BA_peak_10982 |
| DS171652 | 277 | 771 | BA_peak_10983 |
| DS171687 | 196 | 590 | BA_peak_10984 |
| DS171733 | 283 | 853 | BA_peak_10985 |
| DS171801 | 25 | 817 | BA_peak_10987 |

| | | | |
|---|---|---|---|
| DS171820 | 268 | 610 | BA_peak_10988 |
| DS171823 | 210 | 787 | BA_peak_10989 |
| DS171828 | 288 | 719 | BA_peak_10990 |
| DS171838 | 113 | 816 | BA_peak_10991 |
| DS171872 | 93 | 684 | BA_peak_10992 |
| DS171883 | 26 | 694 | BA_peak_10993 |
| DS171894 | 143 | 828 | BA_peak_10995 |
| DS171907 | 114 | 703 | BA_peak_10996 |
| DS171918 | 338 | 678 | BA_peak_10998 |
| DS171925 | 192 | 526 | BA_peak_11000 |
| DS171942 | 199 | 760 | BA_peak_11001 |
| DS171955 | 294 | 720 | BA_peak_11002 |
| DS171960 | 33 | 817 | BA_peak_11003 |
| DS171965 | 81 | 783 | BA_peak_11004 |
| DS171970 | 262 | 698 | BA_peak_11005 |
| DS171971 | 307 | 657 | BA_peak_11006 |
| DS171978 | 56 | 518 | BA_peak_11007 |
| DS171984 | 120 | 701 | BA_peak_11008 |
| DS172004 | 395 | 826 | BA_peak_11010 |
| DS172008 | 85 | 659 | BA_peak_11011 |
| DS172115 | 92 | 644 | BA_peak_11015 |
| DS172138 | 170 | 681 | BA_peak_11016 |
| DS172165 | 219 | 799 | BA_peak_11017 |
| DS172166 | 84 | 746 | BA_peak_11018 |
| DS172167 | 327 | 812 | BA_peak_11019 |
| DS172180 | 140 | 661 | BA_peak_11020 |
| DS172189 | 161 | 832 | BA_peak_11021 |
| DS172223 | 399 | 796 | BA_peak_11023 |
| DS172245 | 70 | 703 | BA_peak_11025 |
| DS172260 | 40 | 606 | BA_peak_11026 |
| DS172293 | 4 | 462 | BA_peak_11027 |
| DS172297 | 119 | 768 | BA_peak_11028 |
| DS172304 | 191 | 818 | BA_peak_11029 |
| DS172342 | 22 | 499 | BA_peak_11030 |
| DS172349 | 29 | 798 | BA_peak_11031 |
| DS172393 | 33 | 678 | BA_peak_11032 |
| DS172416 | 192 | 653 | BA_peak_11033 |
| DS172426 | 101 | 782 | BA_peak_11034 |
| DS172442 | 80 | 640 | BA_peak_11035 |
| DS172482 | 238 | 739 | BA_peak_11036 |
| DS172527 | 46 | 688 | BA_peak_11037 |
| DS172532 | 147 | 676 | BA_peak_11038 |
| DS172542 | 201 | 578 | BA_peak_11039 |
| DS172559 | 38 | 815 | BA_peak_11040 |
| DS172593 | 112 | 593 | BA_peak_11041 |
| DS172636 | 130 | 678 | BA_peak_11042 |
| DS172650 | 206 | 720 | BA_peak_11044 |

| | | | |
|---|---|---|---|
| DS172657 | 168 | 475 | BA_peak_11045 |
| DS172682 | 65 | 753 | BA_peak_11047 |
| DS172702 | 174 | 573 | BA_peak_11050 |
| DS172712 | 32 | 807 | BA_peak_11051 |
| DS172754 | 147 | 741 | BA_peak_11052 |
| DS172778 | 366 | 825 | BA_peak_11053 |
| DS172799 | 243 | 671 | BA_peak_11055 |
| DS172800 | 195 | 828 | BA_peak_11056 |
| DS172889 | 375 | 711 | BA_peak_11057 |
| DS172890 | 239 | 749 | BA_peak_11058 |
| DS172891 | 172 | 778 | BA_peak_11059 |
| DS172929 | 109 | 482 | BA_peak_11060 |
| DS172933 | 48 | 727 | BA_peak_11061 |
| DS172946 | 112 | 647 | BA_peak_11062 |
| DS172965 | 180 | 767 | BA_peak_11065 |
| DS172989 | 97 | 428 | BA_peak_11066 |
| DS173000 | 189 | 688 | BA_peak_11067 |
| DS173006 | 353 | 692 | BA_peak_11068 |
| DS173012 | 73 | 736 | BA_peak_11069 |
| DS173043 | 201 | 672 | BA_peak_11070 |
| DS173058 | 26 | 788 | BA_peak_11071 |
| DS173059 | 130 | 456 | BA_peak_11072 |
| DS173063 | 83 | 501 | BA_peak_11073 |
| DS173086 | 78 | 772 | BA_peak_11074 |
| DS173099 | 217 | 785 | BA_peak_11075 |
| DS173102 | 20 | 836 | BA_peak_11077 |
| DS173117 | 30 | 722 | BA_peak_11078 |
| DS173125 | 126 | 660 | BA_peak_11079 |
| DS173134 | 102 | 725 | BA_peak_11080 |
| DS173135 | 54 | 560 | BA_peak_11081 |
| DS173145 | 18 | 380 | BA_peak_11082 |
| DS173177 | 9 | 488 | BA_peak_11083 |
| DS173184 | 51 | 800 | BA_peak_11084 |
| DS173190 | 94 | 758 | BA_peak_11085 |
| DS173244 | 33 | 821 | BA_peak_11086 |
| DS173293 | 332 | 756 | BA_peak_11087 |
| DS173310 | 101 | 644 | BA_peak_11088 |
| DS173333 | 90 | 613 | BA_peak_11089 |
| DS173336 | 173 | 516 | BA_peak_11090 |
| DS173373 | 79 | 578 | BA_peak_11093 |
| DS173398 | 50 | 752 | BA_peak_11094 |
| DS173411 | 306 | 688 | BA_peak_11095 |
| DS173421 | 182 | 660 | BA_peak_11096 |
| DS173422 | 25 | 770 | BA_peak_11097 |
| DS173463 | 91 | 612 | BA_peak_11098 |
| DS173496 | 99 | 798 | BA_peak_11099 |
| DS173498 | 360 | 731 | BA_peak_11100 |

| | | | |
|---|---|---|---|
| DS173514 | 187 | 717 | BA_peak_11103 |
| DS173522 | 292 | 603 | BA_peak_11104 |
| DS173533 | 9 | 818 | BA_peak_11106 |
| DS173538 | 230 | 722 | BA_peak_11107 |
| DS173565 | 279 | 686 | BA_peak_11109 |
| DS173571 | 58 | 407 | BA_peak_11110 |
| DS173578 | 213 | 757 | BA_peak_11111 |
| DS173594 | 65 | 751 | BA_peak_11112 |
| DS173617 | 63 | 736 | BA_peak_11113 |
| DS173623 | 440 | 778 | BA_peak_11114 |
| DS173649 | 111 | 720 | BA_peak_11115 |
| DS173674 | 334 | 790 | BA_peak_11116 |
| DS173698 | 400 | 738 | BA_peak_11117 |
| DS173724 | 104 | 544 | BA_peak_11118 |
| DS173785 | 235 | 558 | BA_peak_11120 |
| DS173790 | 40 | 679 | BA_peak_11121 |
| DS173798 | 254 | 608 | BA_peak_11122 |
| DS173806 | 96 | 749 | BA_peak_11123 |
| DS173826 | 46 | 631 | BA_peak_11124 |
| DS173830 | 244 | 716 | BA_peak_11125 |
| DS173851 | 37 | 747 | BA_peak_11126 |
| DS173905 | 30 | 499 | BA_peak_11127 |
| DS173924 | 42 | 758 | BA_peak_11129 |
| DS173952 | 272 | 698 | BA_peak_11130 |
| DS173968 | 59 | 778 | BA_peak_11131 |
| DS173984 | 369 | 706 | BA_peak_11133 |
| DS174011 | 105 | 478 | BA_peak_11134 |
| DS174017 | 174 | 753 | BA_peak_11135 |
| DS174045 | 207 | 663 | BA_peak_11136 |
| DS174049 | 281 | 638 | BA_peak_11137 |
| DS174063 | 36 | 677 | BA_peak_11139 |
| DS174064 | 291 | 671 | BA_peak_11140 |
| DS174080 | 46 | 651 | BA_peak_11141 |
| DS174129 | 167 | 601 | BA_peak_11143 |
| DS174145 | 48 | 775 | BA_peak_11144 |
| DS174169 | 41 | 470 | BA_peak_11145 |
| DS174184 | 184 | 782 | BA_peak_11147 |
| DS174192 | 10 | 734 | BA_peak_11148 |
| DS174195 | 164 | 721 | BA_peak_11149 |
| DS174199 | 121 | 782 | BA_peak_11150 |
| DS174262 | 16 | 752 | BA_peak_11153 |
| DS174269 | 443 | 753 | BA_peak_11154 |
| DS174279 | 162 | 637 | BA_peak_11155 |
| DS174286 | 20 | 423 | BA_peak_11156 |
| DS174299 | 349 | 781 | BA_peak_11157 |
| DS174300 | 24 | 782 | BA_peak_11158 |
| DS174302 | 140 | 644 | BA_peak_11159 |

| | | | |
|---|---|---|---|
| DS174338 | 75 | 759 | BA_peak_11160 |
| DS174341 | 219 | 612 | BA_peak_11161 |
| DS174397 | 41 | 759 | BA_peak_11163 |
| DS174411 | 85 | 666 | BA_peak_11164 |
| DS174413 | 94 | 569 | BA_peak_11165 |
| DS174416 | 61 | 699 | BA_peak_11166 |
| DS174466 | 104 | 774 | BA_peak_11168 |
| DS174489 | 104 | 726 | BA_peak_11170 |
| DS174596 | 17 | 560 | BA_peak_11172 |
| DS174603 | 77 | 772 | BA_peak_11173 |
| DS174605 | 50 | 508 | BA_peak_11174 |
| DS174619 | 119 | 740 | BA_peak_11176 |
| DS174659 | 330 | 693 | BA_peak_11178 |
| DS174665 | 137 | 578 | BA_peak_11179 |
| DS174666 | 162 | 779 | BA_peak_11180 |
| DS174696 | 101 | 787 | BA_peak_11181 |
| DS174722 | 135 | 438 | BA_peak_11182 |
| DS174733 | 96 | 572 | BA_peak_11183 |
| DS174743 | 156 | 743 | BA_peak_11184 |
| DS174751 | 197 | 646 | BA_peak_11185 |
| DS174754 | 28 | 684 | BA_peak_11186 |
| DS174779 | 283 | 630 | BA_peak_11187 |
| DS174802 | 62 | 751 | BA_peak_11188 |
| DS174803 | 203 | 616 | BA_peak_11189 |
| DS174814 | 12 | 445 | BA_peak_11190 |
| DS174833 | 69 | 780 | BA_peak_11191 |
| DS174853 | 31 | 681 | BA_peak_11192 |
| DS174873 | 132 | 653 | BA_peak_11193 |
| DS174875 | 109 | 675 | BA_peak_11194 |
| DS174901 | 205 | 717 | BA_peak_11195 |
| DS174942 | 72 | 739 | BA_peak_11198 |
| DS174978 | 97 | 599 | BA_peak_11199 |
| DS175003 | 160 | 732 | BA_peak_11200 |
| DS175014 | 189 | 667 | BA_peak_11201 |
| DS175057 | 359 | 643 | BA_peak_11204 |
| DS175098 | 151 | 591 | BA_peak_11205 |
| DS175143 | 42 | 776 | BA_peak_11206 |
| DS175158 | 34 | 553 | BA_peak_11208 |
| DS175161 | 147 | 590 | BA_peak_11209 |
| DS175162 | 52 | 745 | BA_peak_11210 |
| DS175181 | 47 | 700 | BA_peak_11211 |
| DS175190 | 64 | 652 | BA_peak_11212 |
| DS175191 | 67 | 472 | BA_peak_11213 |
| DS175223 | 33 | 731 | BA_peak_11214 |
| DS175225 | 86 | 647 | BA_peak_11215 |
| DS175227 | 66 | 757 | BA_peak_11216 |
| DS175242 | 21 | 706 | BA_peak_11217 |

| | | | |
|---|---|---|---|
| DS175245 | 208 | 740 | BA_peak_11218 |
| DS175257 | 29 | 755 | BA_peak_11219 |
| DS175268 | 126 | 674 | BA_peak_11220 |
| DS175284 | 99 | 757 | BA_peak_11221 |
| DS175296 | 186 | 651 | BA_peak_11222 |
| DS175336 | 156 | 756 | BA_peak_11223 |
| DS175381 | 59 | 643 | BA_peak_11225 |
| DS175419 | 81 | 710 | BA_peak_11226 |
| DS175453 | 181 | 563 | BA_peak_11227 |
| DS175467 | 193 | 698 | BA_peak_11228 |
| DS175510 | 193 | 653 | BA_peak_11229 |
| DS175511 | 7 | 448 | BA_peak_11230 |
| DS175512 | 203 | 514 | BA_peak_11231 |
| DS175516 | 180 | 647 | BA_peak_11232 |
| DS175522 | 305 | 710 | BA_peak_11233 |
| DS175529 | 22 | 600 | BA_peak_11234 |
| DS175548 | 67 | 487 | BA_peak_11235 |
| DS175554 | 71 | 663 | BA_peak_11236 |
| DS175571 | 82 | 632 | BA_peak_11237 |
| DS175573 | 309 | 701 | BA_peak_11238 |
| DS175581 | 27 | 713 | BA_peak_11239 |
| DS175603 | 241 | 682 | BA_peak_11240 |
| DS175605 | 40 | 746 | BA_peak_11241 |
| DS175615 | 162 | 539 | BA_peak_11242 |
| DS175618 | 14 | 631 | BA_peak_11243 |
| DS175619 | 421 | 730 | BA_peak_11244 |
| DS175669 | 99 | 564 | BA_peak_11245 |
| DS175679 | 302 | 646 | BA_peak_11246 |
| DS175684 | 113 | 478 | BA_peak_11247 |
| DS175686 | 50 | 655 | BA_peak_11248 |
| DS175718 | 50 | 520 | BA_peak_11249 |
| DS175720 | 20 | 662 | BA_peak_11250 |
| DS175742 | 17 | 687 | BA_peak_11251 |
| DS175772 | 32 | 664 | BA_peak_11252 |
| DS175846 | 200 | 601 | BA_peak_11254 |
| DS175887 | 197 | 627 | BA_peak_11256 |
| DS175902 | 39 | 523 | BA_peak_11257 |
| DS175913 | 16 | 416 | BA_peak_11258 |
| DS175929 | 57 | 695 | BA_peak_11259 |
| DS175968 | 176 | 565 | BA_peak_11261 |
| DS175977 | 200 | 715 | BA_peak_11262 |
| DS175979 | 68 | 464 | BA_peak_11263 |
| DS175999 | 247 | 552 | BA_peak_11264 |
| DS176005 | 283 | 684 | BA_peak_11265 |
| DS176023 | 28 | 670 | BA_peak_11266 |
| DS176041 | 197 | 627 | BA_peak_11267 |
| DS176072 | 213 | 685 | BA_peak_11268 |

| | | | |
|---|---|---|---|
| DS176078 | 33 | 577 | BA_peak_11269 |
| DS176084 | 56 | 479 | BA_peak_11270 |
| DS176101 | 308 | 643 | BA_peak_11271 |
| DS176112 | 81 | 648 | BA_peak_11272 |
| DS176116 | 29 | 613 | BA_peak_11273 |
| DS176122 | 172 | 628 | BA_peak_11274 |
| DS176136 | 285 | 648 | BA_peak_11275 |
| DS176154 | 91 | 496 | BA_peak_11276 |
| DS176203 | 28 | 451 | BA_peak_11277 |
| DS176221 | 138 | 545 | BA_peak_11278 |
| DS176259 | 78 | 536 | BA_peak_11280 |
| DS176287 | 43 | 674 | BA_peak_11281 |
| DS176292 | 252 | 591 | BA_peak_11282 |
| DS176298 | 95 | 498 | BA_peak_11283 |
| DS176332 | 56 | 556 | BA_peak_11284 |
| DS176333 | 90 | 490 | BA_peak_11285 |
| DS176336 | 44 | 469 | BA_peak_11286 |
| DS176353 | 84 | 543 | BA_peak_11287 |
| DS176416 | 160 | 573 | BA_peak_11289 |
| DS176490 | 173 | 584 | BA_peak_11291 |
| DS176496 | 284 | 615 | BA_peak_11292 |
| DS176525 | 72 | 481 | BA_peak_11294 |
| DS176527 | 135 | 559 | BA_peak_11295 |
| DS176546 | 83 | 535 | BA_peak_11296 |
| DS176547 | 98 | 402 | BA_peak_11297 |
| DS176575 | 281 | 658 | BA_peak_11298 |
| DS176704 | 104 | 629 | BA_peak_11302 |
| DS176793 | 71 | 472 | BA_peak_11303 |
| DS176794 | 98 | 566 | BA_peak_11304 |
| DS176811 | 228 | 609 | BA_peak_11305 |
| DS176812 | 206 | 595 | BA_peak_11306 |
| DS176829 | 106 | 501 | BA_peak_11307 |
| DS176837 | 37 | 440 | BA_peak_11309 |
| DS176936 | 30 | 405 | BA_peak_11312 |
| DS176941 | 56 | 582 | BA_peak_11314 |
| DS176958 | 98 | 483 | BA_peak_11316 |
| DS176977 | 157 | 523 | BA_peak_11317 |
| DS177007 | 25 | 499 | BA_peak_11318 |
| DS177016 | 140 | 454 | BA_peak_11319 |
| DS177019 | 71 | 533 | BA_peak_11320 |
| DS177031 | 198 | 575 | BA_peak_11321 |
| DS177048 | 64 | 548 | BA_peak_11322 |
| DS177068 | 96 | 519 | BA_peak_11324 |
| DS177084 | 9 | 346 | BA_peak_11325 |
| DS177143 | 41 | 416 | BA_peak_11326 |
| DS177159 | 30 | 485 | BA_peak_11328 |
| DS177308 | 56 | 461 | BA_peak_11331 |

| | | | |
|---|---|---|---|
| DS177317 | 27 | 432 | BA_peak_11332 |
| DS177388 | 104 | 412 | BA_peak_11334 |
| DS177544 | 25 | 370 | BA_peak_11337 |
| DS177618 | 26 | 339 | BA_peak_11338 |

**Supplementary Table 2. Methylated genes IDs**

| Gene ID | Genomic Location | Product Description | Repbase TE classification |
|---|---|---|---|
| TVAG_249310 | DS113187:141,903..142,548(-) | conserved hypothetical protein | DNA/Sola |
| TVAG_423440 | DS113244:103,132..103,640(-) | conserved hypothetical protein | DNA/Sola |
| TVAG_051620 | DS113549:73,931..74,101(-) | conserved hypothetical protein | Harbinger-1_TV |
| TVAG_126000 | DS113357:13,751..15,835(+) | Clan CA, family C19, ubiquitin hydrolase-like cysteine peptidase | Harbinger-1_TV |
| TVAG_525300 | DS140214:524..1,090(-) | conserved hypothetical protein | Harbinger-1_TV |
| TVAG_528100 | DS151751:726..896(-) | conserved hypothetical protein | Harbinger-1_TV |
| TVAG_559990 | DS137739:743..1,073(-) | conserved hypothetical protein | Harbinger-1_TV |
| TVAG_564990 | DS116453:127..297(+) | conserved hypothetical protein | Harbinger-1_TV |
| TVAG_TEG_DS113561_1_1 | DS113561:19,002..21,086(-) | Harbinger-1_Harbinger/PIFA2 transposable element protein | Harbinger-1_TV |
| TVAG_TEG_DS113729_1_1 | DS113729:20,316..22,400(-) | Harbinger-1_Harbinger/PIFA2 transposable element protein | Harbinger-1_TV |
| TVAG_091460 | DS114373:11,439..11,714(+) | conserved hypothetical protein | Harbinger-2_TV |
| TVAG_274810 | DS113344:92,222..92,964(+) | conserved hypothetical protein | hAT-2N1_TV |
| TVAG_104800 | DS114199:7,151..7,402(+) | conserved hypothetical protein | hAT-3N1_TV |
| TVAG_129370 | DS126186:398..649(-) | conserved hypothetical protein | hAT-3N1_TV |
| TVAG_133860 | DS113739:15,122..15,373(-) | conserved hypothetical protein | hAT-3N1_TV |
| TVAG_154830 | DS113582:65,799..66,050(+) | conserved hypothetical protein | hAT-3N1_TV |
| TVAG_239880 | DS113374:50,732..50,983(-) | conserved hypothetical protein | hAT-3N1_TV |
| TVAG_278820 | DS114610:5,146..5,397(+) | conserved hypothetical protein | hAT-3N1_TV |
| TVAG_279260 | DS120332:1,183..1,434(-) | conserved hypothetical protein | hAT-3N1_TV |
| TVAG_294660 | DS113214:8,685..9,282(-) | conserved hypothetical protein | hAT-3N1_TV |
| TVAG_306890 | DS125915:95..346(+) | conserved hypothetical protein | hAT-3N1_TV |
| TVAG_307440 | DS114369:351..974(-) | conserved hypothetical protein | hAT-3N1_TV |
| TVAG_331290 | DS113966:2,314..2,565(-) | conserved hypothetical protein | hAT-3N1_TV |
| TVAG_334530 | DS113949:33,742..33,993(-) | conserved hypothetical protein | hAT-3N1_TV |
| TVAG_345380 | DS113513:64,374..64,625(+) | conserved hypothetical protein | hAT-3N1_TV |
| TVAG_403430 | DS113667:57,670..57,921(+) | conserved hypothetical protein | hAT-3N1_TV |
| TVAG_470560 | DS113429:32,901..33,152(+) | conserved hypothetical protein | hAT-3N1_TV |
| TVAG_529090 | DS132834:111..362(+) | conserved hypothetical protein | hAT-3N1_TV |
| TVAG_023390 | DS113749:5,745..5,996(+) | conserved hypothetical protein | hAT-3N1_TV |
| TVAG_074430 | DS113298:29,640..29,891(+) | conserved hypothetical protein | hAT-3N1_TV |
| TVAG_092010 | DS113899:35,630..35,881(+) | conserved hypothetical protein | hAT-3N1_TV |
| TVAG_161090 | DS113304:58,355..58,606(-) | conserved hypothetical protein | hAT-3N1_TV |
| TVAG_230260 | DS113568:66,800..67,051(+) | conserved hypothetical protein | hAT-3N1_TV |
| TVAG_329920 | DS113950:32,088..32,339(-) | conserved hypothetical protein | hAT-3N1_TV |
| TVAG_TEG_DS113239_2_2 | DS113239:124,643..124,894(+) | Maverick transposable element conserved hypothetical protein | hAT-3N1_TV |
| TVAG_488900 | DS113862:20,571..21,283(-) | conserved hypothetical protein | hAT-3N1_TV/Kolobok-N1_TV |
| TVAG_320740 | DS118472:411..827(+) | conserved hypothetical protein | Kolobok-1_TV |
| TVAG_474820 | DS113468:45,295..46,587(+) | conserved hypothetical protein | Kolobok-1_TV |
| TVAG_073510 | DS116210:2,004..3,296(-) | conserved hypothetical protein | Kolobok-3_TV |
| TVAG_248080 | DS116026:109..642(+) | conserved hypothetical protein | Kolobok-3_TV |
| TVAG_415260 | DS113515:4,009..4,362(-) | conserved hypothetical protein | Kolobok-3_TV |
| TVAG_452830 | DS114560:10,217..10,750(+) | conserved hypothetical protein | Kolobok-3_TV |
| TVAG_511120 | DS135412:598..951(-) | conserved hypothetical protein | Kolobok-3_TV |
| TVAG_527230 | DS135453:33..863(+) | conserved hypothetical protein | Kolobok-3_TV |
| TVAG_601900 | DS144149:131..664(-) | conserved hypothetical protein | Kolobok-3_TV |
| TVAG_TEG_DS113338_1_1 | DS113338:12,681..13,984(+) | Kolobok-3_Kolobok/PhamA4 transposable element protein | Kolobok-3_TV |
| TVAG_TEG_DS113446_1_2 | DS113446:17,554..17,907(+) | Kolobok-3_Kolobok/PhamA4 transposable element protein | Kolobok-3_TV |
| TVAG_TEG_DS113461_2_1 | DS113461:65,426..66,385(+) | Kolobok-3_Kolobok/PhamA4 transposable element protein | Kolobok-3_TV |
| TVAG_TEG_DS113492_1_1 | DS113492:36,818..37,606(-) | Kolobok-3_Kolobok/PhamA4 transposable element protein | Kolobok-3_TV |
| TVAG_TEG_DS113512_2_1 | DS113512:70,063..70,596(-) | Kolobok-3_Kolobok/PhamA4 transposable element protein | Kolobok-3_TV |
| TVAG_TEG_DS113536_3_2 | DS113536:65,165..65,953(+) | Kolobok-3_Kolobok/PhamA4 transposable element protein | Kolobok-3_TV |
| TVAG_TEG_DS113617_1_1 | DS113617:30,290..30,643(-) | Kolobok-3_Kolobok/PhamA4 transposable element protein | Kolobok-3_TV |
| TVAG_TEG_DS113667_1_1 | DS113667:14,906..16,132(-) | Kolobok-3_Kolobok/PhamA4 transposable element protein | Kolobok-3_TV |
| TVAG_TEG_DS113838_2_1 | DS113838:26,179..27,405(+) | Kolobok-3_Kolobok/PhamA4 transposable element protein | Kolobok-3_TV |
| TVAG_TEG_DS113916_1_1 | DS113916:15,178..16,404(-) | Kolobok-3_Kolobok/PhamA4 transposable element protein | Kolobok-3_TV |
| TVAG_TEG_DS114155_1_2 | DS114155:18,918..19,358(+) | Kolobok-3_Kolobok/PhamA4 transposable element protein | Kolobok-3_TV |
| TVAG_TEG_DS114174_1_1 | DS114174:6,212..7,438(+) | Kolobok-3_Kolobok/PhamA4 transposable element protein | Kolobok-3_TV |
| TVAG_TEG_DS114434_3_2 | DS114434:13,096..13,896(-) | Kolobok-3_Kolobok/PhamA4 transposable element protein | Kolobok-3_TV |
| TVAG_005220 | DS116770:18..782(+) | conserved hypothetical protein | Kolobok-3_TV |
| TVAG_037760 | DS119025:262..1,269(+) | conserved hypothetical protein | Kolobok-3_TV |
| TVAG_049880 | DS113512:67,505..68,041(+) | conserved hypothetical protein | Kolobok-3_TV |
| TVAG_064870 | DS113389:96,360..97,556(+) | conserved hypothetical protein | Kolobok-3_TV |
| TVAG_079980 | DS113702:1,230..2,189(-) | conserved hypothetical protein | Kolobok-3_TV |
| TVAG_085550 | DS113591:30,889..31,653(-) | conserved hypothetical protein | Kolobok-3_TV |
| TVAG_138230 | DS113440:208..1,434(+) | conserved hypothetical protein | Kolobok-3_TV |
| TVAG_179450 | DS129583:976..1,272(-) | conserved hypothetical protein | Kolobok-3_TV |
| TVAG_253970 | DS113220:87,955..88,719(-) | conserved hypothetical protein | Kolobok-3_TV |
| TVAG_296470 | DS129462:1,038..1,286(-) | conserved hypothetical protein | Kolobok-3_TV |
| TVAG_314240 | DS134781:846..1,142(-) | conserved hypothetical protein | Kolobok-3_TV |
| TVAG_317970 | DS134224:360..1,124(-) | conserved hypothetical protein | Kolobok-3_TV |
| TVAG_322130 | DS122689:1,248..1,496(-) | conserved hypothetical protein | Kolobok-3_TV |

| | | | |
|---|---|---|---|
| TVAG_374260 | DS114333:20,931..21,371(-) | conserved hypothetical protein | Kolobok-3_TV |
| TVAG_400030 | DS122729:1,237..1,533(-) | conserved hypothetical protein | Kolobok-3_TV |
| TVAG_415240 | DS113515:75..863(+) | conserved hypothetical protein | Kolobok-3_TV |
| TVAG_484520 | DS113579:26,265..27,491(+) | conserved hypothetical protein | Kolobok-3_TV |
| TVAG_506690 | DS127975:907..1,260(-) | conserved hypothetical protein | Kolobok-3_TV |
| TVAG_507350 | DS132269:28..381(+) | conserved hypothetical protein | Kolobok-3_TV |
| TVAG_526210 | DS119819:77..517(+) | conserved hypothetical protein | Kolobok-3_TV |
| TVAG_023350 | DS114284:21,827..22,909(-) | conserved hypothetical protein | Kolobok-4_TV |
| TVAG_034530 | DS116753:1,197..1,631(+) | conserved hypothetical protein | Kolobok-4_TV |
| TVAG_039840 | DS113461:14,777..15,904(+) | leucine-rich repeat protein, BspA family | Kolobok-4_TV |
| TVAG_177240 | DS114321:17,397..18,071(+) | conserved hypothetical protein | Kolobok-4_TV |
| TVAG_179960 | DS113606:71,210..72,577(-) | conserved hypothetical protein | Kolobok-4_TV |
| TVAG_193520 | DS113199:273,826..275,193(+) | conserved hypothetical protein | Kolobok-4_TV |
| TVAG_242580 | DS114345:696..2,063(+) | conserved hypothetical protein | Kolobok-4_TV |
| TVAG_305140 | DS114558:12,047..12,358(+) | conserved hypothetical protein | Kolobok-4_TV |
| TVAG_384690 | DS114032:29,373..29,639(-) | conserved hypothetical protein | Kolobok-4_TV |
| TVAG_440720 | DS114577:8,170..9,537(+) | conserved hypothetical protein | Kolobok-4_TV |
| TVAG_292680 | DS113561:53,396..53,662(+) | conserved hypothetical protein | Kolobok-4_TV |
| TVAG_547790 | DS147068:130..966(-) | conserved hypothetical protein | Kolobok-5_TV |
| TVAG_121030 | DS113177:540,218..541,064(+) | conserved hypothetical protein | Kolobok-N1_TV |
| TVAG_036750 | DS114114:8,008..8,523(-) | conserved hypothetical protein | Mariner-N1_TV |
| TVAG_008060 | DS113524:45,869..46,918(-) | conserved hypothetical protein | MuDr-1_TV |
| TVAG_013230 | DS113189:61,114..62,190(+) | conserved hypothetical protein | MuDr-1_TV |
| TVAG_035760 | DS113183:259,741..261,221(+) | conserved hypothetical protein | MuDr-1_TV |
| TVAG_052850 | DS114874:188..547(+) | conserved hypothetical protein | MuDr-1_TV |
| TVAG_079790 | DS120089:984..1,553(-) | conserved hypothetical protein | MuDr-1_TV |
| TVAG_079890 | DS121471:1,003..1,521(-) | conserved hypothetical protein | MuDr-1_TV |
| TVAG_088540 | DS113343:29,877..31,967(+) | conserved hypothetical protein | MuDr-1_TV |
| TVAG_114670 | DS113780:2,166..3,242(-) | conserved hypothetical protein | MuDr-1_TV |
| TVAG_134880 | DS113849:5,656..6,732(+) | conserved hypothetical protein | MuDr-1_TV |
| TVAG_144090 | DS114207:16,808..17,329(+) | conserved hypothetical protein | MuDr-1_TV |
| TVAG_147770 | DS114056:8,476..9,939(+) | conserved hypothetical protein | MuDr-1_TV |
| TVAG_167920 | DS113977:4,658..6,184(+) | conserved hypothetical protein | MuDr-1_TV |
| TVAG_172280 | DS113193:107,485..108,297(-) | conserved hypothetical protein | MuDr-1_TV |
| TVAG_178020 | DS113204:57,913..59,979(+) | conserved hypothetical protein | MuDr-1_TV |
| TVAG_205040 | DS113401:97,151..98,224(-) | conserved hypothetical protein | MuDr-1_TV |
| TVAG_208000 | DS123331:899..1,243(-) | conserved hypothetical protein | MuDr-1_TV |
| TVAG_217410 | DS113550:39,943..42,195(+) | conserved hypothetical protein | MuDr-1_TV |
| TVAG_218510 | DS117081:1,242..2,090(-) | conserved hypothetical protein | MuDr-1_TV |
| TVAG_239620 | DS113197:290,131..291,834(+) | conserved hypothetical protein | MuDr-1_TV |
| TVAG_258460 | DS113331:107,641..109,701(+) | conserved hypothetical protein | MuDr-1_TV |
| TVAG_260900 | DS113530:74,774..75,295(-) | conserved hypothetical protein | MuDr-1_TV |
| TVAG_267480 | DS150537:140..990(+) | conserved hypothetical protein | MuDr-1_TV |
| TVAG_275020 | DS113915:5,415..5,936(+) | conserved hypothetical protein | MuDr-1_TV |
| TVAG_283800 | DS113192:324,471..326,174(+) | conserved hypothetical protein | MuDr-1_TV |
| TVAG_288680 | DS113745:55,990..57,063(+) | conserved hypothetical protein | MuDr-1_TV |
| TVAG_296500 | DS127935:16..878(+) | conserved hypothetical protein | MuDr-1_TV |
| TVAG_298060 | DS113482:4,016..4,948(-) | conserved hypothetical protein | MuDr-1_TV |
| TVAG_300440 | DS113445:5,483..7,735(+) | conserved hypothetical protein | MuDr-1_TV |
| TVAG_315660 | DS125784:590..1,422(-) | conserved hypothetical protein | MuDr-1_TV |
| TVAG_336500 | DS123377:821..1,342(-) | conserved hypothetical protein | MuDr-1_TV |
| TVAG_338410 | DS114129:19,632..20,471(+) | conserved hypothetical protein | MuDr-1_TV |
| TVAG_341170 | DS113245:155,964..158,216(+) | conserved hypothetical protein | MuDr-1_TV |
| TVAG_353330 | DS113437:25,111..26,115(+) | conserved hypothetical protein | MuDr-1_TV |
| TVAG_365200 | DS114123:5,474..6,550(-) | conserved hypothetical protein | MuDr-1_TV |
| TVAG_413610 | DS134270:21..1,025(-) | conserved hypothetical protein | MuDr-1_TV |
| TVAG_441310 | DS137719:85..606(+) | conserved hypothetical protein | MuDr-1_TV |
| TVAG_443050 | DS115411:1,931..3,007(+) | conserved hypothetical protein | MuDr-1_TV |
| TVAG_459920 | DS136414:137..1,103(+) | conserved hypothetical protein | MuDr-1_TV |
| TVAG_463140 | DS113217:217,227..218,141(-) | conserved hypothetical protein | MuDr-1_TV |
| TVAG_472510 | DS114496:12,839..14,179(+) | conserved hypothetical protein | MuDr-1_TV |
| TVAG_474640 | DS139821:5..1,065(-) | conserved hypothetical protein | MuDr-1_TV |
| TVAG_477220 | DS113182:422,979..424,055(+) | conserved hypothetical protein | MuDr-1_TV |
| TVAG_491920 | DS113340:121,674..123,377(-) | conserved hypothetical protein | MuDr-1_TV |
| TVAG_500310 | DS119231:56..1,686(-) | conserved hypothetical protein | MuDr-1_TV |
| TVAG_516430 | DS141474:101..1,009(-) | conserved hypothetical protein | MuDr-1_TV |
| TVAG_517590 | DS142526:105..626(+) | conserved hypothetical protein | MuDr-1_TV |
| TVAG_528410 | DS150879:390..1,009(+) | conserved hypothetical protein | MuDr-1_TV |
| TVAG_530900 | DS132962:99..1,175(+) | conserved hypothetical protein | MuDr-1_TV |
| TVAG_540520 | DS151020:53..903(-) | conserved hypothetical protein | MuDr-1_TV |
| TVAG_562980 | DS150553:11..919(+) | conserved hypothetical protein | MuDr-1_TV |
| TVAG_572300 | DS141198:221..1,075(-) | conserved hypothetical protein | MuDr-1_TV |
| TVAG_573800 | DS140622:54..1,051(-) | conserved hypothetical protein | MuDr-1_TV |
| TVAG_578780 | DS123094:258..617(+) | conserved hypothetical protein | MuDr-1_TV |

| | | | |
|---|---|---|---|
| TVAG_579490 | DS150314:136..986(+) | conserved hypothetical protein | MuDr-1_TV |
| TVAG_583270 | DS135996:485..1,006(+) | conserved hypothetical protein | MuDr-1_TV |
| TVAG_585210 | DS144376:593..1,042(-) | conserved hypothetical protein | MuDr-1_TV |
| TVAG_587370 | DS132463:113..1,132(-) | conserved hypothetical protein | MuDr-1_TV |
| TVAG_592760 | DS152545:82..944(-) | conserved hypothetical protein | MuDr-1_TV |
| TVAG_597420 | DS116700:15..788(-) | conserved hypothetical protein | MuDr-1_TV |
| TVAG_599260 | DS130403:32..603(-) | conserved hypothetical protein | MuDr-1_TV |
| TVAG_TEG_DS114076_2_2 | DS114076:22,623..24,875(+) | MuDr-1_MuDR/Mutator_Phantom transposable element protein | MuDr-1_TV |
| TVAG_063670 | DS113492:71,633..72,739(-) | conserved hypothetical protein | MuDr-1_TV |
| TVAG_098260 | DS113352:35,135..36,241(+) | conserved hypothetical protein | MuDr-1_TV |
| TVAG_125840 | DS113484:79,043..80,149(-) | conserved hypothetical protein | MuDr-1_TV |
| TVAG_144080 | DS114207:15,461..16,567(-) | conserved hypothetical protein | MuDr-1_TV |
| TVAG_172440 | DS113193:133,044..134,150(+) | conserved hypothetical protein | MuDr-1_TV |
| TVAG_178010 | DS113204:56,472..57,578(-) | conserved hypothetical protein | MuDr-1_TV |
| TVAG_205170 | DS114743:2,147..3,253(+) | conserved hypothetical protein | MuDr-1_TV |
| TVAG_239610 | DS113197:286,837..287,943(-) | conserved hypothetical protein | MuDr-1_TV |
| TVAG_329730 | DS113345:128,147..129,253(+) | conserved hypothetical protein | MuDr-1_TV |
| TVAG_353320 | DS113437:22,494..23,600(-) | conserved hypothetical protein | MuDr-1_TV |
| TVAG_365210 | DS114123:7,299..8,405(+) | conserved hypothetical protein | MuDr-1_TV |
| TVAG_384430 | DS114015:16,855..17,961(-) | conserved hypothetical protein | MuDr-1_TV |
| TVAG_474840 | DS113468:49,278..50,384(+) | conserved hypothetical protein | MuDr-1_TV |
| TVAG_584530 | DS131970:75..1,181(+) | conserved hypothetical protein | MuDr-1_TV |
| TVAG_TEG_DS113188_4_2 | DS113188:323,958..325,064(+) | MuDr-1_MuDR/Mutator_Phantom transposable element protein | MuDr-1_TV |
| TVAG_TEG_DS113291_1_1 | DS113291:16,658..17,764(-) | MuDr-1_MuDR/Mutator_Phantom transposable element protein | MuDr-1_TV |
| TVAG_TEG_DS114076_2_1 | DS114076:21,182..22,288(-) | MuDr-1_MuDR/Mutator_Phantom transposable element protein | MuDr-1_TV |
| TVAG_012690 | DS114332:15,442..17,004(-) | conserved hypothetical protein | MuDr-1_TV |
| TVAG_031650 | DS127936:231..752(+) | conserved hypothetical protein | MuDr-1_TV |
| TVAG_031950 | DS113497:79,205..80,959(-) | conserved hypothetical protein | MuDr-1_TV |
| TVAG_041530 | DS113753:2,730..3,251(+) | conserved hypothetical protein | MuDr-1_TV |
| TVAG_063920 | DS113775:47,884..48,942(+) | conserved hypothetical protein | MuDr-1_TV |
| TVAG_096440 | DS114936:348..773(+) | conserved hypothetical protein | MuDr-1_TV |
| TVAG_126410 | DS113357:122,376..124,466(-) | conserved hypothetical protein | MuDr-1_TV |
| TVAG_132650 | DS113981:32,777..33,298(-) | conserved hypothetical protein | MuDr-1_TV |
| TVAG_215930 | DS113971:26,505..26,864(+) | conserved hypothetical protein | MuDr-1_TV |
| TVAG_216270 | DS113443:42,149..44,242(+) | conserved hypothetical protein | MuDr-1_TV |
| TVAG_246560 | DS113212:4,936..5,457(-) | conserved hypothetical protein | MuDr-1_TV |
| TVAG_264320 | DS113765:48,227..48,910(-) | conserved hypothetical protein | MuDr-1_TV |
| TVAG_301800 | DS113772:23,018..23,539(+) | conserved hypothetical protein | MuDr-1_TV |
| TVAG_385030 | DS116080:3,129..3,491(-) | conserved hypothetical protein | MuDr-1_TV |
| TVAG_405390 | DS114158:5,425..5,946(-) | conserved hypothetical protein | MuDr-1_TV |
| TVAG_417210 | DS113476:78,876..79,268(-) | conserved hypothetical protein | MuDr-1_TV |
| TVAG_420780 | DS114273:1,306..2,214(+) | conserved hypothetical protein | MuDr-1_TV |
| TVAG_521080 | DS129080:267..788(+) | conserved hypothetical protein | MuDr-1_TV |
| TVAG_527530 | DS151543:29..409(-) | conserved hypothetical protein | MuDr-1_TV |
| TVAG_528550 | DS122939:759..1,280(+) | conserved hypothetical protein | MuDr-1_TV |
| TVAG_546880 | DS146811:121..642(+) | conserved hypothetical protein | MuDr-1_TV |
| TVAG_550390 | DS134748:730..1,140(-) | conserved hypothetical protein | MuDr-1_TV |
| TVAG_589270 | DS151421:13..555(+) | conserved hypothetical protein | MuDr-1_TV |
| TVAG_602490 | DS141993:150..671(+) | conserved hypothetical protein | MuDr-1_TV |
| TVAG_605550 | DS129201:143..664(+) | conserved hypothetical protein | MuDr-1_TV |
| TVAG_TEG_DS113188_4_1 | DS113188:321,482..323,623(-) | MuDr-1_MuDR/Mutator_Phantom transposable element protein | MuDr-1_TV |
| TVAG_TEG_DS113771_1_1 | DS113771:26,759..27,658(-) | MuDr-1_MuDR/Mutator_Phantom transposable element protein | MuDr-1_TV |
| TVAG_183890 | DS113180:445,418..446,494(-) | conserved hypothetical protein | MuDr-1_TV |
| TVAG_528230 | DS128657:218..748(+) | conserved hypothetical protein | MuDr-1_TV/MAR1_TV |
| TVAG_175010 | DS117317:1,595..2,416(-) | conserved hypothetical protein | MuDr-1_TV/Polinton-4_TV |
| TVAG_438820 | DS116621:704..1,180(+) | conserved hypothetical protein | MuDr-1_TV/Polinton-4_TV |
| TVAG_059070 | DS113467:12,569..13,264(+) | conserved hypothetical protein | MuDR-3_TV |
| TVAG_124310 | DS114150:89..1,591(-) | conserved hypothetical protein | MuDR-3_TV |
| TVAG_148800 | DS114648:6,790..7,485(+) | conserved hypothetical protein | MuDR-3_TV |
| TVAG_259290 | DS113349:77,211..78,062(+) | conserved hypothetical protein | MuDR-3_TV |
| TVAG_292930 | DS113523:30,637..31,071(-) | conserved hypothetical protein | MuDR-3_TV |
| TVAG_296650 | DS124481:49..1,389(+) | conserved hypothetical protein | MuDR-3_TV |
| TVAG_332940 | DS113388:47,495..47,929(-) | conserved hypothetical protein | MuDR-3_TV |
| TVAG_372060 | DS113281:135,616..136,050(-) | conserved hypothetical protein | MuDR-3_TV |
| TVAG_386920 | DS140591:193..1,044(-) | conserved hypothetical protein | MuDR-3_TV |
| TVAG_414040 | DS113351:75,184..75,618(-) | conserved hypothetical protein | MuDR-3_TV |
| TVAG_505220 | DS116011:239..673(+) | conserved hypothetical protein | MuDR-3_TV |
| TVAG_529510 | DS149187:145..840(+) | conserved hypothetical protein | MuDR-3_TV |
| TVAG_583990 | DS124004:95..946(-) | conserved hypothetical protein | MuDR-3_TV |
| TVAG_006750 | DS113710:2,254..3,594(+) | conserved hypothetical protein | MuDR-3_TV |
| TVAG_037960 | DS114174:23,528..23,962(-) | conserved hypothetical protein | MuDR-3_TV |
| TVAG_042950 | DS113639:5,999..7,003(+) | conserved hypothetical protein | MuDR-3_TV |
| TVAG_050060 | DS113405:8,112..9,484(+) | conserved hypothetical protein | MuDR-3_TV |
| TVAG_104220 | DS114014:16,594..17,028(-) | conserved hypothetical protein | MuDR-3_TV |

| | | | |
|---|---|---|---|
| TVAG_110040 | DS113198:56,197..57,195(+) | conserved hypothetical protein | MuDR-3_TV |
| TVAG_123800 | DS113435:6,575..8,155(+) | conserved hypothetical protein | MuDR-3_TV |
| TVAG_132780 | DS115738:2,179..3,519(-) | conserved hypothetical protein | MuDR-3_TV |
| TVAG_138070 | DS113350:108,489..109,451(+) | conserved hypothetical protein | MuDR-3_TV |
| TVAG_140230 | DS113635:59,253..60,625(-) | conserved hypothetical protein | MuDR-3_TV |
| TVAG_145350 | DS113498:71,507..73,180(+) | conserved hypothetical protein | MuDR-3_TV |
| TVAG_159190 | DS115340:57..491(+) | conserved hypothetical protein | MuDR-3_TV |
| TVAG_215150 | DS113631:29,779..31,119(+) | conserved hypothetical protein | MuDR-3_TV |
| TVAG_223250 | DS114160:18,832..19,266(-) | conserved hypothetical protein | MuDR-3_TV |
| TVAG_223540 | DS113403:31,907..32,758(-) | conserved hypothetical protein | MuDR-3_TV |
| TVAG_237100 | DS113188:63,380..64,231(-) | conserved hypothetical protein | MuDR-3_TV |
| TVAG_247600 | DS116757:130..564(+) | conserved hypothetical protein | MuDR-3_TV |
| TVAG_249720 | DS113187:234,707..236,302(-) | conserved hypothetical protein | MuDR-3_TV |
| TVAG_284160 | DS113758:37,961..38,959(+) | conserved hypothetical protein | MuDR-3_TV |
| TVAG_311530 | DS113409:13,274..14,269(+) | conserved hypothetical protein | MuDR-3_TV |
| TVAG_333920 | DS113396:260..1,963(+) | conserved hypothetical protein | MuDR-3_TV |
| TVAG_365860 | DS113201:138,504..138,938(+) | conserved hypothetical protein | MuDR-3_TV |
| TVAG_414330 | DS113736:17,895..18,590(-) | conserved hypothetical protein | MuDR-3_TV |
| TVAG_470290 | DS114358:950..1,381(-) | conserved hypothetical protein | MuDR-3_TV |
| TVAG_481310 | DS113580:36,974..38,647(+) | conserved hypothetical protein | MuDR-3_TV |
| TVAG_127680 | DS114728:3,378..4,697(-) | conserved hypothetical protein | MuDR-3_TV/Polinton-6_TV |
| TVAG_176880 | DS113680:6,681..8,231(+) | conserved hypothetical protein | MuDR-4_TV |
| TVAG_216490 | DS113443:89,635..91,104(+) | conserved hypothetical protein | MuDR-4_TV |
| TVAG_383250 | DS113985:2,898..4,016(+) | conserved hypothetical protein | MuDR-4_TV |
| TVAG_461230 | DS121162:39..419(+) | conserved hypothetical protein | MuDR-4_TV |
| TVAG_522410 | DS126929:67..839(+) | conserved hypothetical protein | MuDR-4_TV |
| TVAG_493010 | DS114043:27,024..27,203(+) | conserved hypothetical protein | MuDR-4N1_TV |
| TVAG_196870 | DS113731:1,383..2,000(+) | conserved hypothetical protein | MuDR-6_TV |
| TVAG_291100 | DS113603:60,017..60,676(-) | conserved hypothetical protein | MuDR-6_TV |
| TVAG_335720 | DS114870:7,683..8,438(-) | conserved hypothetical protein | MuDR-7_TV |
| TVAG_577950 | DS117991:1,565..2,130(-) | conserved hypothetical protein | MuDR-7_TV |
| TVAG_068980 | DS113553:80,123..81,028(+) | conserved hypothetical protein | MuDRx-2_TV |
| TVAG_TEG_DS113626_1_1 | DS113626:51,595..54,489(+) | NeSL-1_NeSL/ transposable element protein | NeSL-1_TV |
| TVAG_008960 | DS113784:40,473..41,273(-) | conserved hypothetical protein | P-2_TV |
| TVAG_078690 | DS114502:2,773..3,213(+) | conserved hypothetical protein | P-2_TV |
| TVAG_526440 | DS143145:708..1,049(-) | conserved hypothetical protein | P-2_TV |
| TVAG_518140 | DS139031:164..964(-) | conserved hypothetical protein | P-2_TV |
| TVAG_429840 | DS113852:45,664..46,335(+) | conserved hypothetical protein | Polinton-1_TV |
| TVAG_522800 | DS151691:28..696(+) | conserved hypothetical protein | Polinton-1_TV |
| TVAG_569900 | DS146848:1..653(+) | conserved hypothetical protein | Polinton-1_TV |
| TVAG_199310 | DS113190:282,084..282,674(+) | conserved hypothetical protein | Polinton-1_TV |
| TVAG_443330 | DS113600:16,073..16,888(+) | conserved hypothetical protein | Polinton-1_TV |
| TVAG_041670 | DS113753:38,637..39,308(+) | conserved hypothetical protein | Polinton-1_TV |
| TVAG_085710 | DS113591:75,109..75,780(-) | conserved hypothetical protein | Polinton-1_TV |
| TVAG_109670 | DS113339:94,139..94,351(-) | conserved hypothetical protein | Polinton-1_TV |
| TVAG_132060 | DS128021:444..1,142(-) | conserved hypothetical protein | Polinton-1_TV |
| TVAG_238950 | DS113197:121,606..122,277(+) | conserved hypothetical protein | Polinton-1_TV |
| TVAG_267770 | DS113215:62,467..63,138(-) | conserved hypothetical protein | Polinton-1_TV |
| TVAG_284870 | DS114642:7,988..8,575(-) | conserved hypothetical protein | Polinton-1_TV |
| TVAG_315570 | DS147671:1..661(+) | conserved hypothetical protein | Polinton-1_TV |
| TVAG_323290 | DS114024:191..859(+) | conserved hypothetical protein | Polinton-1_TV |
| TVAG_323770 | DS113607:2,347..3,015(-) | conserved hypothetical protein | Polinton-1_TV |
| TVAG_398320 | DS115399:2,817..3,515(-) | conserved hypothetical protein | Polinton-1_TV |
| TVAG_416810 | DS115555:1,817..2,488(-) | conserved hypothetical protein | Polinton-1_TV |
| TVAG_493150 | DS113584:25,851..26,438(+) | conserved hypothetical protein | Polinton-1_TV |
| TVAG_513230 | DS132017:412..1,083(-) | conserved hypothetical protein | Polinton-1_TV |
| TVAG_529270 | DS130217:335..1,033(+) | conserved hypothetical protein | Polinton-1_TV |
| TVAG_541090 | DS132415:223..921(+) | conserved hypothetical protein | Polinton-1_TV |
| TVAG_551240 | DS122512:662..1,300(+) | conserved hypothetical protein | Polinton-1_TV |
| TVAG_558030 | DS141368:156..854(+) | conserved hypothetical protein | Polinton-1_TV |
| TVAG_559970 | DS142616:238..936(-) | conserved hypothetical protein | Polinton-1_TV |
| TVAG_563540 | DS151255:354..1,005(-) | conserved hypothetical protein | Polinton-1_TV |
| TVAG_029890 | DS114886:6,842..7,057(-) | conserved hypothetical protein | Polinton-1_TV |
| TVAG_053380 | DS113434:29,848..30,063(-) | conserved hypothetical protein | Polinton-1_TV |
| TVAG_109830 | DS113339:127,072..127,287(-) | conserved hypothetical protein | Polinton-1_TV |
| TVAG_301580 | DS113307:142,654..142,869(-) | conserved hypothetical protein | Polinton-1_TV |
| TVAG_005800 | DS113799:45,847..46,518(+) | conserved hypothetical protein | Polinton-1_TV |
| TVAG_154860 | DS113582:74,229..74,900(+) | conserved hypothetical protein | Polinton-1_TV |
| TVAG_256080 | DS113271:170,077..170,775(+) | conserved hypothetical protein | Polinton-1_TV |
| TVAG_269210 | DS113379:67,352..68,020(+) | conserved hypothetical protein | Polinton-1_TV |
| TVAG_290570 | DS113460:67,522..68,190(-) | conserved hypothetical protein | Polinton-1_TV |
| TVAG_246280 | DS115056:4,096..4,656(+) | conserved hypothetical protein | Polinton-1_TV |
| TVAG_397410 | DS114967:2,057..2,626(+) | conserved hypothetical protein | Polinton-1_TV |
| TVAG_250680 | DS113479:68,571..69,050(-) | conserved hypothetical protein | Polinton-1B_TV |

| | | | |
|---|---|---|---|
| TVAG_255280 | DS113933:38,763..39,206(+) | conserved hypothetical protein | Polinton-1B_TV |
| TVAG_269470 | DS132340:30..473(-) | conserved hypothetical protein | Polinton-1B_TV |
| TVAG_275580 | DS116956:40..519(-) | conserved hypothetical protein | Polinton-1B_TV |
| TVAG_308680 | DS114058:30,047..30,490(+) | conserved hypothetical protein | Polinton-1B_TV |
| TVAG_332020 | DS114780:1,465..1,908(-) | conserved hypothetical protein | Polinton-1B_TV |
| TVAG_373230 | DS113274:92,306..92,749(-) | conserved hypothetical protein | Polinton-1B_TV |
| TVAG_430610 | DS113294:94,133..94,576(-) | conserved hypothetical protein | Polinton-1B_TV |
| TVAG_438280 | DS113542:35,033..35,512(-) | conserved hypothetical protein | Polinton-1B_TV |
| TVAG_443240 | DS117077:143..586(-) | conserved hypothetical protein | Polinton-1B_TV |
| TVAG_451730 | DS145534:397..840(+) | conserved hypothetical protein | Polinton-1B_TV |
| TVAG_548650 | DS139877:387..830(+) | conserved hypothetical protein | Polinton-1B_TV |
| TVAG_568480 | DS120007:1,234..1,677(+) | conserved hypothetical protein | Polinton-1B_TV |
| TVAG_572330 | DS152193:393..836(-) | conserved hypothetical protein | Polinton-1B_TV |
| TVAG_591530 | DS126080:819..1,262(+) | conserved hypothetical protein | Polinton-1B_TV |
| TVAG_078850 | DS114940:4,553..5,032(-) | conserved hypothetical protein | Polinton-1B_TV |
| TVAG_079360 | DS113373:45,312..45,968(+) | conserved hypothetical protein | Polinton-1B_TV |
| TVAG_092330 | DS117777:1,358..1,801(+) | conserved hypothetical protein | Polinton-1B_TV |
| TVAG_123280 | DS113791:53,338..53,817(+) | conserved hypothetical protein | Polinton-1B_TV |
| TVAG_128500 | DS114498:14,783..15,262(+) | conserved hypothetical protein | Polinton-1B_TV |
| TVAG_129340 | DS118993:142..621(-) | conserved hypothetical protein | Polinton-1B_TV |
| TVAG_133480 | DS113361:118,212..118,655(+) | conserved hypothetical protein | Polinton-1B_TV |
| TVAG_140580 | DS113408:64,201..64,644(+) | conserved hypothetical protein | Polinton-1B_TV |
| TVAG_150820 | DS117611:847..1,290(+) | conserved hypothetical protein | Polinton-1B_TV |
| TVAG_153870 | DS113451:80,194..80,637(-) | conserved hypothetical protein | Polinton-1B_TV |
| TVAG_154540 | DS125320:951..1,394(-) | conserved hypothetical protein | Polinton-1B_TV |
| TVAG_185000 | DS113179:74,223..74,666(+) | conserved hypothetical protein | Polinton-1B_TV |
| TVAG_251520 | DS113372:18,228..18,671(-) | conserved hypothetical protein | Polinton-1B_TV |
| TVAG_270030 | DS113672:35,247..35,690(-) | conserved hypothetical protein | Polinton-1B_TV |
| TVAG_272990 | DS113562:76,925..77,368(+) | conserved hypothetical protein | Polinton-1B_TV |
| TVAG_275910 | DS113540:55,135..55,590(+) | conserved hypothetical protein | Polinton-1B_TV |
| TVAG_279450 | DS114438:16,531..16,974(+) | conserved hypothetical protein | Polinton-1B_TV |
| TVAG_297230 | DS113235:88,887..89,330(-) | conserved hypothetical protein | Polinton-1B_TV |
| TVAG_312700 | DS132098:685..1,128(+) | conserved hypothetical protein | Polinton-1B_TV |
| TVAG_340810 | DS113245:40,237..40,680(+) | conserved hypothetical protein | Polinton-1B_TV |
| TVAG_341840 | DS114811:2,201..2,644(-) | conserved hypothetical protein | Polinton-1B_TV |
| TVAG_345020 | DS113566:64,554..65,222(-) | conserved hypothetical protein | Polinton-1B_TV |
| TVAG_365070 | DS124020:699..1,142(+) | conserved hypothetical protein | Polinton-1B_TV |
| TVAG_380550 | DS115261:3,249..3,692(+) | conserved hypothetical protein | Polinton-1B_TV |
| TVAG_439440 | DS113688:15,652..16,095(+) | conserved hypothetical protein | Polinton-1B_TV |
| TVAG_440800 | DS113527:27,700..28,143(-) | conserved hypothetical protein | Polinton-1B_TV |
| TVAG_443360 | DS113600:29,582..30,025(-) | conserved hypothetical protein | Polinton-1B_TV |
| TVAG_459420 | DS113797:290..733(-) | conserved hypothetical protein | Polinton-1B_TV |
| TVAG_459720 | DS114283:3,358..3,837(-) | conserved hypothetical protein | Polinton-1B_TV |
| TVAG_460560 | DS113267:10,842..11,285(-) | conserved hypothetical protein | Polinton-1B_TV |
| TVAG_526750 | DS144605:97..540(-) | conserved hypothetical protein | Polinton-1B_TV |
| TVAG_537820 | DS137146:67..510(-) | conserved hypothetical protein | Polinton-1B_TV |
| TVAG_538690 | DS150890:526..969(-) | conserved hypothetical protein | Polinton-1B_TV |
| TVAG_550830 | DS144401:155..634(+) | conserved hypothetical protein | Polinton-1B_TV |
| TVAG_558720 | DS133229:562..1,005(-) | conserved hypothetical protein | Polinton-1B_TV |
| TVAG_572530 | DS119438:641..1,300(+) | conserved hypothetical protein | Polinton-1B_TV |
| TVAG_076180 | DS113181:202,848..203,291(+) | conserved hypothetical protein | Polinton-1B_TV |
| TVAG_094120 | DS113185:259,081..259,524(+) | conserved hypothetical protein | Polinton-1B_TV |
| TVAG_144600 | DS114239:24,386..24,829(+) | conserved hypothetical protein | Polinton-1B_TV |
| TVAG_147990 | DS113952:563..1,006(-) | conserved hypothetical protein | Polinton-1B_TV |
| TVAG_192250 | DS113490:91,709..92,152(+) | conserved hypothetical protein | Polinton-1B_TV |
| TVAG_202550 | DS113439:8,326..8,805(-) | conserved hypothetical protein | Polinton-1B_TV |
| TVAG_207450 | DS113587:26,036..26,479(+) | conserved hypothetical protein | Polinton-1B_TV |
| TVAG_218530 | DS125764:873..1,352(+) | conserved hypothetical protein | Polinton-1B_TV |
| TVAG_233740 | DS113875:32,460..33,126(-) | conserved hypothetical protein | Polinton-1B_TV |
| TVAG_014370 | DS117682:614..1,075(-) | conserved hypothetical protein | Polinton-2_TV |
| TVAG_274960 | DS115169:1,020..1,550(+) | conserved hypothetical protein | Polinton-2_TV |
| TVAG_111940 | DS114372:9,402..10,100(+) | conserved hypothetical protein | Polinton-2B_TV |
| TVAG_RG_DS117241_1 | DS117241:1,423..2,187(-) | conserved hypothetical protein | Polinton-2B_TV |
| TVAG_TEG_DS113593_1_1 | DS113593:11,828..12,016(-) | Maverick transposable element conserved hypothetical protein | Polinton-3_TV |
| TVAG_TEG_DS114353_1_4 | DS114353:8,115..8,516(+) | Maverick transposable element conserved hypothetical protein | Polinton-3_TV |
| TVAG_103900 | DS113898:1,259..1,702(+) | conserved hypothetical protein | Polinton-4_TV |
| TVAG_252830 | DS115625:1,143..1,463(+) | conserved hypothetical protein | Polinton-4_TV |
| TVAG_295910 | DS119958:1,316..1,558(-) | conserved hypothetical protein | Polinton-4_TV |
| TVAG_338310 | DS114564:11,097..11,417(-) | conserved hypothetical protein | Polinton-4_TV |
| TVAG_596940 | DS125001:169..495(+) | conserved hypothetical protein | Polinton-4_TV |
| TVAG_TEG_DS113458_4_8 | DS113460:5,974..6,210(+) | Maverick transposable element conserved hypothetical protein | Polinton-4_TV |
| TVAG_TEG_DS113500_2_1 | DS113500:20,230..20,763(-) | Maverick transposable element hypothetical protein | Polinton-4_TV |
| TVAG_350440 | DS113477:41,059..41,343(+) | conserved hypothetical protein | Poseidon-2B_PM |
| TVAG_415490 | DS114307:1,672..1,995(-) | conserved hypothetical protein | SOLA/Cas |

| | | | |
|---|---|---|---|
| TVAG_143020 | DS114568:1..339(+) | conserved hypothetical protein | VANDAL6 |
| TVAG_042320 | DS113501:70,722..70,979(-) | conserved hypothetical protein | VANDAL6 |
| TVAG_087690 | DS113737:56,968..57,510(-) | conserved hypothetical protein | VANDAL6 |
| TVAG_097870 | DS113289:88,579..89,304(-) | conserved hypothetical protein | VANDAL6 |
| TVAG_104020 | DS113898:26,698..27,147(+) | conserved hypothetical protein | VANDAL6 |
| TVAG_110900 | DS113198:278,957..279,301(-) | conserved hypothetical protein | VANDAL6 |
| TVAG_169190 | DS123187:20..919(+) | conserved hypothetical protein | VANDAL6 |
| TVAG_232340 | DS117325:1..989(+) | conserved hypothetical protein | VANDAL6 |
| TVAG_250670 | DS113479:56,682..57,311(+) | conserved hypothetical protein | VANDAL6 |
| TVAG_306580 | DS113222:169,903..170,460(-) | conserved hypothetical protein | VANDAL6 |
| TVAG_326250 | DS115011:5,677..5,988(-) | conserved hypothetical protein | VANDAL6 |
| TVAG_372760 | DS113546:68,280..68,489(+) | conserved hypothetical protein | VANDAL6 |
| TVAG_399260 | DS113310:28,878..29,249(+) | conserved hypothetical protein | VANDAL6 |
| TVAG_399800 | DS126184:249..1,408(-) | conserved hypothetical protein | VANDAL6 |
| TVAG_402770 | DS113202:241,823..241,978(+) | conserved hypothetical protein | VANDAL6 |
| TVAG_500630 | DS127112:8..571(+) | conserved hypothetical protein | VANDAL6 |
| TVAG_516970 | DS149250:1..802(+) | conserved hypothetical protein | VANDAL6 |
| TVAG_547940 | DS119867:830..1,228(+) | conserved hypothetical protein | VANDAL6 |
| TVAG_098470 | DS113352:106,007..106,555(-) | conserved hypothetical protein | VANDAL6 |
| TVAG_127770 | DS115690:812..1,048(-) | conserved hypothetical protein | VANDAL6 |
| TVAG_132230 | DS113908:9,312..9,464(-) | conserved hypothetical protein | VANDAL6 |
| TVAG_133370 | DS113361:93,972..94,346(+) | conserved hypothetical protein | VANDAL6 |
| TVAG_151660 | DS113425:22,929..23,912(-) | conserved hypothetical protein | VANDAL6 |
| TVAG_158700 | DS113315:37,386..37,643(+) | conserved hypothetical protein | VANDAL6 |
| TVAG_166540 | DS113191:140,378..140,683(+) | conserved hypothetical protein | VANDAL6 |
| TVAG_166570 | DS113191:146,347..146,865(+) | conserved hypothetical protein | VANDAL6 |
| TVAG_177660 | DS128356:892..1,329(-) | conserved hypothetical protein | VANDAL6 |
| TVAG_189580 | DS113576:60,310..60,486(+) | conserved hypothetical protein | VANDAL6 |
| TVAG_189610 | DS113576:61,525..61,956(+) | conserved hypothetical protein | VANDAL6 |
| TVAG_243890 | DS127559:555..707(-) | conserved hypothetical protein | VANDAL6 |
| TVAG_243900 | DS121484:1..340(+) | conserved hypothetical protein | VANDAL6 |
| TVAG_255650 | DS113271:47,163..47,342(+) | conserved hypothetical protein | VANDAL6 |
| TVAG_264080 | DS113765:2,728..3,027(+) | conserved hypothetical protein | VANDAL6 |
| TVAG_275140 | DS113915:33,244..33,405(+) | conserved hypothetical protein | VANDAL6 |
| TVAG_291640 | DS113233:95,875..96,030(-) | conserved hypothetical protein | VANDAL6 |
| TVAG_380290 | DS113263:129,231..129,785(+) | conserved hypothetical protein | VANDAL6 |
| TVAG_380670 | DS114083:10,833..11,102(+) | conserved hypothetical protein | VANDAL6 |
| TVAG_398000 | DS116470:1,666..2,010(-) | conserved hypothetical protein | VANDAL6 |
| TVAG_501090 | DS143313:1..399(+) | conserved hypothetical protein | VANDAL6 |
| TVAG_341270 | DS113245:175,491..176,279(-) | coiled-coil domain-containing protein, putative | VANDAL6 |
| TVAG_317190 | DS113954:14,099..14,275(+) | conserved hypothetical protein | VANDAL6 |
| TVAG_010500 | DS113225:29,526..29,912(-) | conserved hypothetical protein | VANDAL6 |
| TVAG_284540 | DS113438:81,485..82,199(-) | conserved hypothetical protein | VANDAL6 |
| TVAG_345530 | DS114066:9,903..10,787(+) | conserved hypothetical protein | VANDAL6 |
| TVAG_391660 | DS113195:225,182..226,297(-) | hypothetical protein | VANDAL6 |
| TVAG_468410 | DS135198:19..715(+) | conserved hypothetical protein | VANDAL6 |
| TVAG_496810 | DS113917:32,320..32,934(-) | conserved hypothetical protein | VANDAL6 |
| TVAG_503660 | DS122153:696..1,601(-) | conserved hypothetical protein | VANDAL6 |
| TVAG_095990 | DS115470:16..684(+) | conserved hypothetical protein | |
| TVAG_416150 | DS113938:16..684(+) | conserved hypothetical protein | |
| TVAG_573900 | DS115087:227..1,954(-) | conserved hypothetical protein | |
| TVAG_116940 | DS113297:4,685..5,415(+) | conserved hypothetical protein | |
| TVAG_203290 | DS113906:4,759..5,284(-) | conserved hypothetical protein | |
| TVAG_233920 | DS125492:492..1,223(-) | conserved hypothetical protein | |
| TVAG_388050 | DS113282:125,276..125,883(+) | conserved hypothetical protein | |
| TVAG_397960 | DS114045:28,972..29,481(+) | conserved hypothetical protein | |
| TVAG_565730 | DS139282:347..837(+) | conserved hypothetical protein | |
| TVAG_138600 | DS113440:93,881..94,324(+) | conserved hypothetical protein | |
| TVAG_138610 | DS113440:97,878..99,080(+) | conserved hypothetical protein | |
| TVAG_564940 | DS145253:1..814(+) | conserved hypothetical protein | |
| TVAG_262740 | DS113976:18,933..20,426(-) | ankyrin 2,3/unc44, putative | |
| TVAG_387640 | DS113282:9,845..10,219(+) | centaurin beta, putative | |
| TVAG_369830 | DS113980:32,161..34,535(+) | conserved hypothetical protein | |
| TVAG_525630 | DS135889:11..807(-) | conserved hypothetical protein | |
| TVAG_453400 | DS119476:880..1,697(-) | conserved hypothetical protein | |
| TVAG_555820 | DS141100:45..935(-) | conserved hypothetical protein | |
| TVAG_511550 | DS150845:181..874(-) | conserved hypothetical protein | |
| TVAG_582580 | DS130050:579..839(+) | conserved hypothetical protein | |
| TVAG_283960 | DS115762:246..986(-) | conserved hypothetical protein | |
| TVAG_353180 | DS116385:749..2,323(-) | conserved hypothetical protein | |
| TVAG_414760 | DS114272:407..2,827(+) | conserved hypothetical protein | |
| TVAG_499200 | DS128850:532..1,259(-) | conserved hypothetical protein | |
| TVAG_515580 | DS151437:23..653(-) | conserved hypothetical protein | |
| TVAG_518410 | DS145552:391..1,031(-) | conserved hypothetical protein | |

| | | |
|---|---|---|
| TVAG_520870 | DS141949:1..1,051(-) | conserved hypothetical protein |
| TVAG_551960 | DS118738:457..945(+) | conserved hypothetical protein |
| TVAG_588030 | DS133715:23..605(-) | conserved hypothetical protein |
| TVAG_080520 | DS149162:17..1,019(+) | conserved hypothetical protein |
| TVAG_086650 | DS127697:40..904(-) | conserved hypothetical protein |
| TVAG_286190 | DS120565:50..1,130(-) | conserved hypothetical protein |
| TVAG_408220 | DS146572:79..977(+) | conserved hypothetical protein |
| TVAG_001120 | DS114414:838..2,774(+) | conserved hypothetical protein |
| TVAG_001140 | DS114414:4,784..5,281(-) | conserved hypothetical protein |
| TVAG_001660 | DS113997:18,792..19,380(+) | conserved hypothetical protein |
| TVAG_004510 | DS113242:47,633..48,766(+) | lipase containing protein, putative |
| TVAG_008680 | DS113653:19,701..21,090(+) | tubulin epsilon chain, putative |
| TVAG_012750 | DS114545:3,413..3,985(+) | conserved hypothetical protein |
| TVAG_014190 | DS113189:331,645..332,181(-) | conserved hypothetical protein |
| TVAG_014570 | DS114623:10,533..11,087(+) | C-MYB, putative |
| TVAG_023490 | DS113749:34,104..34,460(-) | conserved hypothetical protein |
| TVAG_029390 | DS114679:1,553..3,937(-) | conserved hypothetical protein |
| TVAG_034880 | DS113183:30,120..30,434(-) | Clan IH, family I25, phytocystatin-like peptidase inhibitor |
| TVAG_043330 | DS114339:14,467..14,853(+) | tubulin, putative |
| TVAG_047950 | DS113552:30,560..31,880(-) | CAMK family protein kinase |
| TVAG_052540 | DS113779:37,315..38,271(-) | ankyrin repeat-containing protein, putative |
| TVAG_054330 | DS113539:9,985..10,848(-) | conserved hypothetical protein |
| TVAG_056780 | DS113412:32..619(+) | conserved hypothetical protein |
| TVAG_056810 | DS113412:5,282..6,008(+) | conserved hypothetical protein |
| TVAG_068370 | DS113861:12,144..14,018(-) | conserved hypothetical protein |
| TVAG_072170 | DS133551:861..1,196(-) | conserved hypothetical protein |
| TVAG_072390 | DS113495:40,505..40,990(+) | conserved hypothetical protein |
| TVAG_080440 | DS113874:30,309..31,808(+) | CMGC family protein kinase |
| TVAG_085990 | DS114125:8,188..11,886(+) | Basic proline-rich protein precursor |
| TVAG_087030 | DS113436:2,287..3,966(+) | conserved hypothetical protein |
| TVAG_092170 | DS114085:14,006..15,460(-) | preprotein translocase secy subunit, putative |
| TVAG_095260 | DS114718:5,324..6,226(+) | AGC family protein kinase |
| TVAG_098430 | DS113352:95,802..96,050(-) | 30S/40S ribosomal protein S4, putative |
| TVAG_100000 | DS113411:61,523..61,720(+) | hypothetical protein |
| TVAG_103780 | DS113836:21,545..22,308(-) | Family T1, proteasome alpha subunit, threonine peptidase |
| TVAG_104690 | DS113913:25,236..26,432(-) | conserved hypothetical protein |
| TVAG_107240 | DS116257:842..1,243(-) | conserved hypothetical protein |
| TVAG_114120 | DS113602:9,002..9,853(-) | conserved hypothetical protein |
| TVAG_124330 | DS114150:11,644..12,309(-) | conserved hypothetical protein |
| TVAG_125640 | DS113484:17,922..18,782(+) | MYB-1, putative |
| TVAG_136580 | DS113207:199,392..202,022(+) | heat shock protein, putative |
| TVAG_136630 | DS113207:206,570..208,811(+) | ankyrin repeat-containing protein, putative |
| TVAG_140090 | DS113635:31,244..31,609(+) | conserved hypothetical protein |
| TVAG_152490 | DS113927:11,778..12,179(+) | hypothetical protein |
| TVAG_157150 | DS113334:15,918..16,845(-) | CAMK family protein kinase |
| TVAG_157220 | DS113334:29,739..30,743(+) | conserved hypothetical protein |
| TVAG_157290 | DS113334:48,513..49,919(-) | conserved hypothetical protein |
| TVAG_157320 | DS113334:56,556..57,098(+) | conserved hypothetical protein |
| TVAG_157480 | DS113334:104,521..106,089(-) | conserved hypothetical protein |
| TVAG_158440 | DS113919:36,770..42,822(+) | centromeric protein E, putative |
| TVAG_159260 | DS113620:7,046..7,761(+) | conserved hypothetical protein |
| TVAG_161980 | DS114779:5,988..8,003(+) | conserved hypothetical protein |
| TVAG_173240 | DS113193:314,957..315,109(+) | hypothetical protein |
| TVAG_178150 | DS113204:98,675..99,991(-) | conserved hypothetical protein |
| TVAG_182070 | DS113641:61,603..62,037(+) | conserved hypothetical protein |
| TVAG_190450 | DS113211:170,005..173,261(+) | kakapo, putative |
| TVAG_195270 | DS113877:6,029..7,715(+) | AGC family protein kinase |
| TVAG_204750 | DS113401:38,561..39,826(-) | conserved hypothetical protein |
| TVAG_205470 | DS113767:56,046..56,633(-) | conserved hypothetical protein |
| TVAG_216380 | DS113443:70,679..71,648(+) | conserved hypothetical protein |
| TVAG_222740 | DS113803:3,003..5,435(-) | conserved hypothetical protein |
| TVAG_224480 | DS113257:116,068..117,486(+) | CAMK family protein kinase |
| TVAG_226520 | DS113464:65,912..67,220(-) | conserved hypothetical protein |
| TVAG_230010 | DS127057:1,092..1,382(-) | conserved hypothetical protein |
| TVAG_232620 | DS113494:58,685..59,128(+) | conserved hypothetical protein |
| TVAG_233640 | DS113875:11,407..12,109(-) | conserved hypothetical protein |
| TVAG_233680 | DS113875:17,952..18,680(+) | conserved hypothetical protein |
| TVAG_237910 | DS113188:234,621..238,370(-) | conserved hypothetical protein |
| TVAG_239310 | DS113197:216,802..220,473(-) | bollus pemphigoid antigen, putative |
| TVAG_241540 | DS113859:32,101..32,298(-) | conserved hypothetical protein |
| TVAG_248270 | DS113318:17,119..17,688(+) | conserved hypothetical protein |
| TVAG_254150 | DS113220:132,617..132,985(-) | serine/threonine-protein kinase chk2, putative |
| TVAG_254160 | DS113220:136,637..138,547(+) | conserved hypothetical protein |
| TVAG_256090 | DS114137:7,789..10,182(-) | conserved hypothetical protein |

| | | |
|---|---|---|
| TVAG_257540 | DS113422:101,554..103,239(+) | ankyrin, putative |
| TVAG_259370 | DS113349:102,516..102,728(-) | conserved hypothetical protein |
| TVAG_266660 | DS113232:89,538..92,102(-) | conserved hypothetical protein |
| TVAG_274030 | DS113991:23,148..23,528(+) | MYB3r3, putative |
| TVAG_275370 | DS113691:41,222..42,352(-) | tankyrase, putative |
| TVAG_276140 | DS113355:28,113..28,370(-) | conserved hypothetical protein |
| TVAG_277610 | DS113756:8,534..10,477(+) | hyaluronan mediated motility receptor, putative |
| TVAG_278640 | DS114942:3,864..4,112(-) | conserved hypothetical protein |
| TVAG_282800 | DS113192:58,579..59,880(+) | conserved hypothetical protein |
| TVAG_293350 | DS114384:4,255..4,602(-) | conserved hypothetical protein |
| TVAG_298330 | DS113482:63,702..64,763(-) | conserved hypothetical protein |
| TVAG_299950 | DS113687:38,352..43,082(-) | adenylate cyclase, putative |
| TVAG_301190 | DS113307:41,414..44,122(-) | conserved hypothetical protein |
| TVAG_303020 | DS114159:11,788..15,935(+) | conserved hypothetical protein |
| TVAG_306880 | DS117621:33..353(+) | conserved hypothetical protein |
| TVAG_307360 | DS138932:106..348(+) | conserved hypothetical protein |
| TVAG_308060 | DS113383:45,284..46,038(-) | conserved hypothetical protein |
| TVAG_312970 | DS114037:16,484..17,338(-) | conserved hypothetical protein |
| TVAG_318200 | DS114356:16,409..16,864(-) | r2r3-MYB transcription factor, putative |
| TVAG_332930 | DS113388:45,246..46,352(+) | conserved hypothetical protein |
| TVAG_333220 | DS113388:110,252..111,952(-) | conserved hypothetical protein |
| TVAG_345100 | DS114470:7,984..9,114(-) | tip60, putative |
| TVAG_352010 | DS113275:167,319..167,777(+) | conserved hypothetical protein |
| TVAG_361420 | DS114185:4,325..5,005(-) | small GTPase RAB, putative |
| TVAG_363120 | DS113944:13,196..13,576(-) | conserved hypothetical protein |
| TVAG_369900 | DS113525:22,379..22,678(-) | conserved hypothetical protein |
| TVAG_371930 | DS113281:92,627..96,439(+) | TPR Domain containing protein |
| TVAG_373190 | DS113274:76,778..79,267(+) | conserved hypothetical protein |
| TVAG_386640 | DS114535:381..1,061(+) | conserved hypothetical protein |
| TVAG_386690 | DS114535:8,554..9,594(+) | conserved hypothetical protein |
| TVAG_399260 | DS113310:28,878..29,249(+) | conserved hypothetical protein |
| TVAG_404020 | DS113382:14,350..15,890(+) | conserved hypothetical protein |
| TVAG_405500 | DS113978:9,393..15,785(-) | PIKK family atypical protein kinase |
| TVAG_415700 | DS114095:7,944..8,468(-) | conserved hypothetical protein |
| TVAG_416160 | DS113938:3,194..3,913(+) | conserved hypothetical protein |
| TVAG_416210 | DS113938:13,942..14,892(-) | conserved hypothetical protein |
| TVAG_416330 | DS113938:41,705..41,965(+) | conserved hypothetical protein |
| TVAG_419630 | DS113400:52,391..53,131(+) | conserved hypothetical protein |
| TVAG_420820 | DS114273:10,313..19,876(-) | neurobeachin, putative |
| TVAG_424980 | DS113622:15,362..16,039(+) | conserved hypothetical protein |
| TVAG_431850 | DS113655:21,980..22,759(+) | adenylate and guanylate cyclases, putative |
| TVAG_434730 | DS113240:174,638..175,561(+) | conserved hypothetical protein |
| TVAG_440010 | DS115640:566..1,939(+) | cGMP-inhibited 3,5-cyclic phosphodiesterase, putative |
| TVAG_441130 | DS113994:34,874..35,980(+) | leucine-rich repeat protein, BspA family |
| TVAG_441540 | DS113863:16,280..17,552(-) | conserved hypothetical protein |
| TVAG_446480 | DS115049:2,582..3,768(+) | AGC family protein kinase |
| TVAG_447330 | DS113238:57,328..57,696(+) | conserved hypothetical protein |
| TVAG_448940 | DS145388:313..796(-) | conserved hypothetical protein |
| TVAG_454260 | DS113229:200,491..200,772(-) | 60S ribosomal protein L37, putative |
| TVAG_458090 | DS113855:20,126..25,286(+) | centromeric protein E, putative |
| TVAG_458930 | DS113312:84,548..86,203(-) | conserved hypothetical protein |
| TVAG_459310 | DS114009:11,156..11,305(+) | hypothetical protein |
| TVAG_464340 | DS116159:909..1,592(+) | conserved hypothetical protein |
| TVAG_464660 | DS113368:89,475..90,737(-) | conserved hypothetical protein |
| TVAG_465300 | DS113246:100,721..102,295(+) | leucine-rich repeat protein, BspA family |
| TVAG_469850 | DS114073:11,554..12,042(-) | conserved hypothetical protein |
| TVAG_470750 | DS113429:97,733..98,520(+) | conserved hypothetical protein |
| TVAG_483480 | DS113485:60,636..61,889(-) | AGC family protein kinase |
| TVAG_484330 | DS114036:1,068..1,397(+) | hypothetical protein |
| TVAG_487670 | DS113376:86,639..88,045(-) | conserved hypothetical protein |
| TVAG_504730 | DS145036:74..614(-) | conserved hypothetical protein |
| TVAG_505650 | DS150630:288..653(+) | conserved hypothetical protein |
| TVAG_514380 | DS125181:75..1,459(+) | ankyrin repeat-containing protein, putative |
| TVAG_516690 | DS136433:111..1,134(-) | conserved hypothetical protein |
| TVAG_524430 | DS121661:813..1,635(+) | conserved hypothetical protein |
| TVAG_527100 | DS132024:680..1,099(-) | conserved hypothetical protein |
| TVAG_542550 | DS140230:329..589(+) | conserved hypothetical protein |
| TVAG_554060 | DS118368:733..1,092(+) | conserved hypothetical protein |
| TVAG_561190 | DS144791:193..366(-) | conserved hypothetical protein |
| TVAG_580850 | DS124706:318..1,049(-) | conserved hypothetical protein |
| TVAG_605780 | DS150478:53..593(-) | conserved hypothetical protein |
| TVAG_RG_DS113361_14 | DS113361:80,198..81,238(-) | conserved hypothetical protein |
| TVAG_RG_DS113405_5 | DS113405:103,048..103,668(+) | conserved hypothetical protein |
| TVAG_RG_DS113755_7 | DS113755:31,620..32,594(-) | conserved hypothetical protein |

| | | |
|---|---|---|
| TVAG_RG_DS114977_3 | DS114977:3,696..4,151(+) | conserved hypothetical protein |

**Supplementary Table 3.  GO Enrichment of methylated genes**

| ID | Name | Fold enrichment | P-value |
|---|---|---|---|
| GO:0006796 | phosphate-containing compound metabolic process | 2.42 | 4.83E-03 |
| GO:0006793 | phosphorus metabolic process | 2.41 | 5.00E-03 |
| GO:0006468 | protein phosphorylation | 2.56 | 7.68E-03 |
| GO:0016310 | phosphorylation | 2.39 | 1.18E-02 |
| GO:0019538 | protein metabolic process | 1.88 | 1.31E-02 |
| GO:0006464 | cellular protein modification process | 2.13 | 1.70E-02 |
| GO:0036211 | protein modification process | 2.13 | 1.70E-02 |
| GO:0043412 | macromolecule modification | 2.08 | 1.98E-02 |
| GO:0044267 | cellular protein metabolic process | 1.9 | 2.09E-02 |
| GO:1901564 | organonitrogen compound metabolic process | 1.75 | 2.40E-02 |
| GO:0016573 | histone acetylation | 24.08 | 4.08E-02 |
| GO:0006473 | protein acetylation | 20.64 | 4.74E-02 |
| GO:0006475 | internal protein amino acid acetylation | 20.64 | 4.74E-02 |
| GO:0018393 | internal peptidyl-lysine acetylation | 20.64 | 4.74E-02 |
| GO:0018394 | peptidyl-lysine acetylation | 20.64 | 4.74E-02 |

**Supplementary Table 4. List of *T. vaginalis* TEs**

| RE superfamily | n° in genome | % methylated |
|---|---|---|
| Polinton | 55782 | 2.7 |
| MuDR | 4758 | 28.5 |
| Kolobok | 2461 | 35.8 |
| Harbinger | 430 | 18.6 |
| Mariner | 1838 | 15.7 |
| hAT | 1926 | 24.0 |
| NeSL | 11 | 0.0 |
| P | 579 | 75.3 |
| Repeat Simple-1 | 4216 | 0.0 |
| Repeat DNA7 | 856 | 65.5 |

**Supplementary Table 5. Relative expression of genes with known expression in strain B7268**

| Gene | Expression level | RPKM | log2 Fold change |
|---|---|---|---|
| FR1 | low | 0 | 0.006024261 |
| PFOR | moderate | 8 | 1 |
| TVAG_248270 | high | 24 | 5.548123877 |
| BAP2 | high | 243 | 20.67764529 |

**Supplementary Table 6. Expression levels of non TE methylated genes**

| Gene ID | Expression level |
|---|---|
| TVAG_001120 | High |
| TVAG_001660 | High |
| TVAG_008680 | High |
| TVAG_072390 | High |
| TVAG_087030 | High |
| TVAG_092170 | High |
| TVAG_125640 | High |
| TVAG_136580 | High |
| TVAG_157150 | High |
| TVAG_195270 | High |
| TVAG_237910 | High |
| TVAG_248270 | High |
| TVAG_266660 | High |
| TVAG_318200 | High |
| TVAG_361420 | High |
| TVAG_373190 | High |
| TVAG_440010 | High |
| TVAG_454260 | High |
| TVAG_554060 | High |
| TVAG_095990 | Low |
| TVAG_416150 | Low |
| TVAG_116940 | Low |
| TVAG_203290 | Low |
| TVAG_233920 | Low |
| TVAG_388050 | Low |
| TVAG_397960 | Low |
| TVAG_565730 | Low |
| TVAG_138600 | Low |
| TVAG_138610 | Low |
| TVAG_564940 | Low |
| TVAG_262740 | Low |
| TVAG_525630 | Low |
| TVAG_453400 | Low |
| TVAG_555820 | Low |
| TVAG_511550 | Low |
| TVAG_582580 | Low |
| TVAG_283960 | Low |
| TVAG_414760 | Low |
| TVAG_499200 | Low |
| TVAG_515580 | Low |
| TVAG_518410 | Low |
| TVAG_520870 | Low |
| TVAG_551960 | Low |
| TVAG_080520 | Low |

| | |
|---|---|
| TVAG_086650 | Low |
| TVAG_286190 | Low |
| TVAG_408220 | Low |
| TVAG_012750 | Low |
| TVAG_014190 | Low |
| TVAG_056780 | Low |
| TVAG_068370 | Low |
| TVAG_072170 | Low |
| TVAG_100000 | Low |
| TVAG_107240 | Low |
| TVAG_124330 | Low |
| TVAG_140090 | Low |
| TVAG_152490 | Low |
| TVAG_159260 | Low |
| TVAG_204750 | Low |
| TVAG_205470 | Low |
| TVAG_233640 | Low |
| TVAG_254160 | Low |
| TVAG_259370 | Low |
| TVAG_298330 | Low |
| TVAG_306880 | Low |
| TVAG_307360 | Low |
| TVAG_312970 | Low |
| TVAG_333220 | Low |
| TVAG_369900 | Low |
| TVAG_416160 | Low |
| TVAG_419630 | Low |
| TVAG_431850 | Low |
| TVAG_434730 | Low |
| TVAG_448940 | Low |
| TVAG_464340 | Low |
| TVAG_465300 | Low |
| TVAG_469850 | Low |
| TVAG_484330 | Low |
| TVAG_504730 | Low |
| TVAG_505650 | Low |
| TVAG_514380 | Low |
| TVAG_516690 | Low |
| TVAG_524430 | Low |
| TVAG_527100 | Low |
| TVAG_542550 | Low |
| TVAG_580850 | Low |
| TVAG_605780 | Low |
| TVAG_RG_DS113361_14 | Low |
| TVAG_RG_DS113405_5 | Low |
| TVAG_RG_DS113755_7 | Low |
| TVAG_387640 | Low |

| | |
|---|---|
| TVAG_001140 | Low |
| TVAG_029390 | Low |
| TVAG_043330 | Low |
| TVAG_098430 | Low |
| TVAG_114120 | Low |
| TVAG_157320 | Low |
| TVAG_173240 | Low |
| TVAG_182070 | Low |
| TVAG_230010 | Low |
| TVAG_256090 | Low |
| TVAG_275370 | Low |
| TVAG_277610 | Low |
| TVAG_299950 | Low |
| TVAG_332930 | Low |
| TVAG_352010 | Low |
| TVAG_363120 | Low |
| TVAG_405500 | Low |
| TVAG_416330 | Low |
| TVAG_447330 | Low |
| TVAG_459310 | Low |
| TVAG_483480 | Low |
| TVAG_RG_DS114977_3 | Low |
| TVAG_573900 | Moderate |
| TVAG_369830 | Moderate |
| TVAG_353180 | Moderate |
| TVAG_588030 | Moderate |
| TVAG_004510 | Moderate |
| TVAG_014570 | Moderate |
| TVAG_023490 | Moderate |
| TVAG_034880 | Moderate |
| TVAG_047950 | Moderate |
| TVAG_052540 | Moderate |
| TVAG_054330 | Moderate |
| TVAG_056810 | Moderate |
| TVAG_080440 | Moderate |
| TVAG_085990 | Moderate |
| TVAG_095260 | Moderate |
| TVAG_103780 | Moderate |
| TVAG_104690 | Moderate |
| TVAG_136630 | Moderate |
| TVAG_157220 | Moderate |
| TVAG_157290 | Moderate |
| TVAG_157480 | Moderate |
| TVAG_158440 | Moderate |
| TVAG_161980 | Moderate |
| TVAG_178150 | Moderate |
| TVAG_190450 | Moderate |

| | |
|---|---|
| TVAG_216380 | Moderate |
| TVAG_222740 | Moderate |
| TVAG_224480 | Moderate |
| TVAG_226520 | Moderate |
| TVAG_232620 | Moderate |
| TVAG_233680 | Moderate |
| TVAG_239310 | Moderate |
| TVAG_241540 | Moderate |
| TVAG_254150 | Moderate |
| TVAG_257540 | Moderate |
| TVAG_274030 | Moderate |
| TVAG_276140 | Moderate |
| TVAG_278640 | Moderate |
| TVAG_282800 | Moderate |
| TVAG_293350 | Moderate |
| TVAG_301190 | Moderate |
| TVAG_303020 | Moderate |
| TVAG_308060 | Moderate |
| TVAG_345100 | Moderate |
| TVAG_371930 | Moderate |
| TVAG_386640 | Moderate |
| TVAG_386690 | Moderate |
| TVAG_399260 | Moderate |
| TVAG_404020 | Moderate |
| TVAG_415700 | Moderate |
| TVAG_416210 | Moderate |
| TVAG_420820 | Moderate |
| TVAG_424980 | Moderate |
| TVAG_441130 | Moderate |
| TVAG_441540 | Moderate |
| TVAG_446480 | Moderate |
| TVAG_458090 | Moderate |
| TVAG_458930 | Moderate |
| TVAG_464660 | Moderate |
| TVAG_470750 | Moderate |
| TVAG_487670 | Moderate |
| TVAG_561190 | Moderate |

**Supplementary Table 7. Intervals delimited by intergenic m6A**

| Contig | Start | End | Interval name | Type | n° of genes |
|---|---|---|---|---|---|
| DS113178 | 21470 | 37625 | BA_peak_24_BA_peak_25 | repressive | 6 |
| DS113179 | 42883 | 61444 | BA_peak_48_BA_peak_49 | repressive | 15 |
| DS113179 | 73536 | 90406 | BA_peak_50_BA_peak_51 | repressive | 9 |
| DS113179 | 293247 | 301374 | BA_peak_72_BA_peak_73 | repressive | 2 |
| DS113179 | 462207 | 469461 | BA_peak_90_BA_peak_91 | repressive | 5 |
| DS113180 | 274467 | 279110 | BA_peak_111_BA_peak_112 | repressive | 4 |
| DS113180 | 335828 | 353603 | BA_peak_114_BA_peak_115 | repressive | 15 |
| DS113181 | 137796 | 156852 | BA_peak_131_BA_peak_132 | repressive | 13 |
| DS113181 | 202833 | 214610 | BA_peak_136_BA_peak_137 | repressive | 7 |
| DS113182 | 15037 | 22610 | BA_peak_157_BA_peak_158 | repressive | 2 |
| DS113182 | 322604 | 328866 | BA_peak_175_BA_peak_176 | repressive | 2 |
| DS113182 | 418744 | 424074 | BA_peak_180_BA_peak_181 | repressive | 2 |
| DS113183 | 14049 | 29899 | BA_peak_185_BA_peak_186 | repressive | 14 |
| DS113184 | 143436 | 156434 | BA_peak_222_BA_peak_223 | repressive | 4 |
| DS113184 | 249345 | 296816 | BA_peak_228_BA_peak_230 | repressive | 37 |
| DS113184 | 308162 | 327906 | BA_peak_231_BA_peak_233 | repressive | 11 |
| DS113184 | 382376 | 390579 | BA_peak_234_BA_peak_235 | repressive | 5 |
| DS113185 | 59012 | 73929 | BA_peak_241_BA_peak_242 | repressive | 12 |
| DS113185 | 197484 | 202451 | BA_peak_247_BA_peak_248 | repressive | 2 |
| DS113185 | 256963 | 259824 | BA_peak_251_BA_peak_252 | repressive | 2 |
| DS113185 | 259013 | 273723 | BA_peak_252_BA_peak_253 | repressive | 7 |
| DS113186 | 5404 | 9673 | BA_peak_264_BA_peak_265 | repressive | 2 |
| DS113188 | 5291 | 12992 | BA_peak_317_BA_peak_318 | repressive | 2 |
| DS113188 | 320640 | 325348 | BA_peak_341_BA_peak_342 | repressive | 2 |
| DS113188 | 324059 | 329411 | BA_peak_342_BA_peak_343 | repressive | 2 |
| DS113188 | 328606 | 335389 | BA_peak_343_BA_peak_344 | repressive | 2 |
| DS113189 | 53546 | 62802 | BA_peak_349_BA_peak_350 | repressive | 7 |
| DS113189 | 136830 | 144012 | BA_peak_355_BA_peak_356 | repressive | 5 |
| DS113189 | 331045 | 334367 | BA_peak_367_BA_peak_368 | repressive | 2 |
| DS113190 | 282131 | 288625 | BA_peak_389_BA_peak_390 | repressive | 4 |
| DS113191 | 140013 | 147917 | BA_peak_401_BA_peak_403 | repressive | 4 |
| DS113191 | 182317 | 192256 | BA_peak_405_BA_peak_406 | repressive | 2 |
| DS113191 | 217963 | 222974 | BA_peak_408_BA_peak_410 | repressive | 2 |
| DS113192 | 325089 | 343460 | BA_peak_430_BA_peak_431 | repressive | 16 |
| DS113193 | 106928 | 134448 | BA_peak_441_BA_peak_442 | repressive | 26 |
| DS113193 | 301745 | 315836 | BA_peak_445_BA_peak_446 | repressive | 16 |
| DS113197 | 122031 | 130664 | BA_peak_497_BA_peak_498 | repressive | 4 |
| DS113197 | 145699 | 159977 | BA_peak_500_BA_peak_501 | repressive | 6 |
| DS113197 | 204327 | 216901 | BA_peak_502_BA_peak_503 | repressive | 10 |
| DS113197 | 286432 | 291789 | BA_peak_506_BA_peak_507 | repressive | 2 |
| DS113198 | 53347 | 57564 | BA_peak_514_BA_peak_515 | repressive | 2 |
| DS113198 | 114315 | 125206 | BA_peak_517_BA_peak_518 | repressive | 6 |
| DS113198 | 254664 | 265591 | BA_peak_525_BA_peak_526 | repressive | 9 |
| DS113199 | 73562 | 87163 | BA_peak_531_BA_peak_532 | repressive | 10 |

| | | | | | |
|---|---|---|---|---|---|
| DS113200 | 127072 | 134559 | BA_peak_550_BA_peak_551 | repressive | 2 |
| DS113200 | 222324 | 224333 | BA_peak_555_BA_peak_556 | repressive | 2 |
| DS113202 | 236139 | 242584 | BA_peak_594_BA_peak_595 | repressive | 4 |
| DS113204 | 124348 | 127538 | BA_peak_627_BA_peak_628 | repressive | 2 |
| DS113204 | 127067 | 133727 | BA_peak_628_BA_peak_629 | repressive | 4 |
| DS113206 | 16237 | 36881 | BA_peak_657_BA_peak_658 | repressive | 13 |
| DS113207 | 111207 | 119646 | BA_peak_687_BA_peak_688 | repressive | 6 |
| DS113208 | 88106 | 93064 | BA_peak_707_BA_peak_708 | repressive | 2 |
| DS113208 | 137812 | 146991 | BA_peak_713_BA_peak_714 | repressive | 6 |
| DS113208 | 179833 | 195719 | BA_peak_717_BA_peak_718 | repressive | 12 |
| DS113208 | 194892 | 199097 | BA_peak_718_BA_peak_719 | repressive | 3 |
| DS113210 | 170472 | 189343 | BA_peak_750_BA_peak_751 | repressive | 10 |
| DS113213 | 124378 | 142018 | BA_peak_786_BA_peak_787 | repressive | 10 |
| DS113214 | 142997 | 160115 | BA_peak_802_BA_peak_803 | repressive | 12 |
| DS113215 | 54697 | 62685 | BA_peak_808_BA_peak_809 | repressive | 5 |
| DS113215 | 61625 | 69729 | BA_peak_809_BA_peak_810 | repressive | 6 |
| DS113217 | 45389 | 49433 | BA_peak_832_BA_peak_833 | repressive | 3 |
| DS113217 | 49056 | 54699 | BA_peak_833_BA_peak_834 | repressive | 2 |
| DS113217 | 111249 | 121273 | BA_peak_837_BA_peak_838 | repressive | 6 |
| DS113217 | 142343 | 152050 | BA_peak_840_BA_peak_841 | repressive | 6 |
| DS113219 | 18771 | 32245 | BA_peak_858_BA_peak_859 | repressive | 8 |
| DS113220 | 175874 | 186605 | BA_peak_881_BA_peak_882 | repressive | 9 |
| DS113221 | 66424 | 77612 | BA_peak_886_BA_peak_887 | repressive | 5 |
| DS113221 | 77110 | 81651 | BA_peak_887_BA_peak_888 | repressive | 2 |
| DS113221 | 108448 | 127647 | BA_peak_890_BA_peak_891 | repressive | 12 |
| DS113225 | 29443 | 37343 | BA_peak_928_BA_peak_929 | repressive | 2 |
| DS113226 | 35257 | 47014 | BA_peak_938_BA_peak_939 | repressive | 5 |
| DS113228 | 197975 | 203779 | BA_peak_963_BA_peak_964 | repressive | 4 |
| DS113229 | 196860 | 199908 | BA_peak_979_BA_peak_980 | repressive | 2 |
| DS113230 | 101170 | 111790 | BA_peak_990_BA_peak_991 | repressive | 7 |
| DS113232 | 143767 | 154779 | BA_peak_1010_BA_peak_1011 | repressive | 7 |
| DS113234 | 81012 | 89505 | BA_peak_1020_BA_peak_1021 | repressive | 4 |
| DS113234 | 89121 | 93514 | BA_peak_1021_BA_peak_1022 | repressive | 3 |
| DS113234 | 135005 | 141465 | BA_peak_1024_BA_peak_1025 | repressive | 3 |
| DS113234 | 140318 | 143417 | BA_peak_1025_BA_peak_1026 | repressive | 2 |
| DS113234 | 154111 | 160560 | BA_peak_1028_BA_peak_1029 | repressive | 4 |
| DS113235 | 81984 | 90065 | BA_peak_1038_BA_peak_1039 | repressive | 6 |
| DS113236 | 78876 | 80847 | BA_peak_1051_BA_peak_1052 | repressive | 2 |
| DS113238 | 52241 | 58564 | BA_peak_1080_BA_peak_1081 | repressive | 2 |
| DS113238 | 58252 | 66119 | BA_peak_1081_BA_peak_1082 | repressive | 2 |
| DS113239 | 59556 | 69752 | BA_peak_1095_BA_peak_1096 | repressive | 3 |
| DS113239 | 121434 | 124891 | BA_peak_1099_BA_peak_1100 | repressive | 2 |
| DS113239 | 124588 | 131479 | BA_peak_1100_BA_peak_1101 | repressive | 8 |
| DS113239 | 139479 | 162437 | BA_peak_1102_BA_peak_1103 | repressive | 18 |
| DS113240 | 140611 | 147327 | BA_peak_1112_BA_peak_1113 | repressive | 3 |
| DS113242 | 22264 | 36492 | BA_peak_1127_BA_peak_1128 | repressive | 8 |
| DS113242 | 36057 | 38930 | BA_peak_1128_BA_peak_1129 | repressive | 2 |

| | | | | | |
|---|---|---|---|---|---|
| DS113242 | 49333 | 54593 | BA_peak_1130_BA_peak_1131 | repressive | 2 |
| DS113242 | 54290 | 66507 | BA_peak_1131_BA_peak_1132 | repressive | 7 |
| DS113242 | 151439 | 175333 | BA_peak_1144_BA_peak_1145 | repressive | 15 |
| DS113243 | 75891 | 88236 | BA_peak_1149_BA_peak_1150 | repressive | 8 |
| DS113245 | 39502 | 53978 | BA_peak_1169_BA_peak_1170 | repressive | 8 |
| DS113245 | 144005 | 154554 | BA_peak_1176_BA_peak_1177 | repressive | 7 |
| DS113247 | 49249 | 53282 | BA_peak_1197_BA_peak_1198 | repressive | 2 |
| DS113247 | 52967 | 70995 | BA_peak_1198_BA_peak_1199 | repressive | 17 |
| DS113248 | 109448 | 122236 | BA_peak_1208_BA_peak_1209 | repressive | 3 |
| DS113249 | 35026 | 41101 | BA_peak_1217_BA_peak_1218 | repressive | 2 |
| DS113252 | 21423 | 37793 | BA_peak_1245_BA_peak_1246 | repressive | 9 |
| DS113252 | 125159 | 136257 | BA_peak_1254_BA_peak_1255 | repressive | 7 |
| DS113253 | 155843 | 165054 | BA_peak_1269_BA_peak_1270 | repressive | 5 |
| DS113256 | 137243 | 143594 | BA_peak_1293_BA_peak_1294 | repressive | 5 |
| DS113258 | 70304 | 89720 | BA_peak_1315_BA_peak_1316 | repressive | 18 |
| DS113259 | 169126 | 182314 | BA_peak_1334_BA_peak_1336 | repressive | 8 |
| DS113260 | 24272 | 38113 | BA_peak_1337_BA_peak_1338 | repressive | 11 |
| DS113262 | 91291 | 105542 | BA_peak_1356_BA_peak_1357 | repressive | 5 |
| DS113263 | 70011 | 82719 | BA_peak_1364_BA_peak_1365 | repressive | 9 |
| DS113265 | 62093 | 64763 | BA_peak_1385_BA_peak_1386 | repressive | 2 |
| DS113271 | 21035 | 28829 | BA_peak_1451_BA_peak_1452 | repressive | 3 |
| DS113271 | 165871 | 170803 | BA_peak_1463_BA_peak_1464 | repressive | 3 |
| DS113272 | 165697 | 170289 | BA_peak_1480_BA_peak_1481 | repressive | 2 |
| DS113273 | 39492 | 45115 | BA_peak_1486_BA_peak_1487 | repressive | 5 |
| DS113274 | 79773 | 92849 | BA_peak_1498_BA_peak_1499 | repressive | 8 |
| DS113274 | 139281 | 148763 | BA_peak_1504_BA_peak_1505 | repressive | 6 |
| DS113275 | 67 | 7587 | BA_peak_1508_BA_peak_1509 | repressive | 7 |
| DS113275 | 161495 | 168152 | BA_peak_1514_BA_peak_1515 | repressive | 4 |
| DS113279 | 47587 | 64290 | BA_peak_1538_BA_peak_1539 | repressive | 11 |
| DS113279 | 86097 | 94016 | BA_peak_1545_BA_peak_1546 | repressive | 2 |
| DS113280 | 81969 | 88936 | BA_peak_1557_BA_peak_1558 | repressive | 4 |
| DS113283 | 50753 | 65436 | BA_peak_1592_BA_peak_1593 | repressive | 9 |
| DS113283 | 65051 | 69894 | BA_peak_1593_BA_peak_1594 | repressive | 2 |
| DS113283 | 136673 | 146765 | BA_peak_1595_BA_peak_1596 | repressive | 6 |
| DS113284 | 91191 | 96873 | BA_peak_1601_BA_peak_1602 | repressive | 3 |
| DS113285 | 102469 | 122771 | BA_peak_1608_BA_peak_1611 | repressive | 10 |
| DS113287 | 132272 | 142546 | BA_peak_1627_BA_peak_1628 | repressive | 7 |
| DS113288 | 78 | 6661 | BA_peak_1631_BA_peak_1632 | repressive | 4 |
| DS113288 | 78305 | 88609 | BA_peak_1636_BA_peak_1637 | repressive | 3 |
| DS113290 | 23766 | 34033 | BA_peak_1654_BA_peak_1655 | repressive | 5 |
| DS113290 | 117407 | 126205 | BA_peak_1662_BA_peak_1663 | repressive | 3 |
| DS113291 | 16312 | 21065 | BA_peak_1669_BA_peak_1670 | repressive | 2 |
| DS113291 | 92178 | 96244 | BA_peak_1676_BA_peak_1677 | repressive | 2 |
| DS113291 | 95938 | 98783 | BA_peak_1677_BA_peak_1678 | repressive | 4 |
| DS113294 | 89571 | 94617 | BA_peak_1702_BA_peak_1703 | repressive | 4 |
| DS113295 | 43081 | 54141 | BA_peak_1711_BA_peak_1712 | repressive | 7 |
| DS113296 | 49498 | 54833 | BA_peak_1721_BA_peak_1722 | repressive | 2 |

| | | | | | |
|---|---|---|---|---|---|
| DS113296 | 77497 | 94119 | BA_peak_1723_BA_peak_1724 | repressive | 11 |
| DS113297 | 146684 | 150295 | BA_peak_1739_BA_peak_1740 | repressive | 2 |
| DS113299 | 3672 | 26632 | BA_peak_1751_BA_peak_1752 | repressive | 10 |
| DS113300 | 89135 | 105042 | BA_peak_1761_BA_peak_1763 | repressive | 11 |
| DS113302 | 76283 | 94236 | BA_peak_1768_BA_peak_1770 | repressive | 11 |
| DS113304 | 81744 | 85853 | BA_peak_1784_BA_peak_1785 | repressive | 3 |
| DS113305 | 63261 | 70567 | BA_peak_1794_BA_peak_1795 | repressive | 2 |
| DS113305 | 69415 | 73834 | BA_peak_1795_BA_peak_1796 | repressive | 2 |
| DS113305 | 90534 | 100933 | BA_peak_1798_BA_peak_1799 | repressive | 6 |
| DS113308 | 105062 | 110869 | BA_peak_1827_BA_peak_1828 | repressive | 5 |
| DS113308 | 110532 | 124087 | BA_peak_1828_BA_peak_1829 | repressive | 6 |
| DS113311 | 76403 | 84852 | BA_peak_1850_BA_peak_1851 | repressive | 7 |
| DS113313 | 100054 | 108682 | BA_peak_1862_BA_peak_1863 | repressive | 4 |
| DS113314 | 90921 | 125591 | BA_peak_1871_BA_peak_1873 | repressive | 30 |
| DS113318 | 41024 | 45511 | BA_peak_1911_BA_peak_1912 | repressive | 2 |
| DS113321 | 111775 | 128289 | BA_peak_1934_BA_peak_1935 | repressive | 14 |
| DS113323 | 125012 | 131176 | BA_peak_1950_BA_peak_1951 | repressive | 2 |
| DS113325 | 18150 | 37057 | BA_peak_1962_BA_peak_1963 | repressive | 14 |
| DS113325 | 119519 | 126382 | BA_peak_1968_BA_peak_1969 | repressive | 2 |
| DS113330 | 40185 | 47759 | BA_peak_2016_BA_peak_2017 | repressive | 4 |
| DS113332 | 64850 | 80830 | BA_peak_2040_BA_peak_2041 | repressive | 10 |
| DS113334 | 57800 | 72645 | BA_peak_2056_BA_peak_2059 | repressive | 10 |
| DS113335 | 12904 | 28787 | BA_peak_2073_BA_peak_2074 | repressive | 10 |
| DS113339 | 93018 | 104119 | BA_peak_2105_BA_peak_2106 | repressive | 8 |
| DS113341 | 56068 | 69523 | BA_peak_2119_BA_peak_2120 | repressive | 8 |
| DS113342 | 8799 | 17323 | BA_peak_2124_BA_peak_2125 | repressive | 2 |
| DS113342 | 22752 | 30081 | BA_peak_2126_BA_peak_2127 | repressive | 3 |
| DS113343 | 93616 | 103907 | BA_peak_2138_BA_peak_2139 | repressive | 7 |
| DS113344 | 64871 | 79592 | BA_peak_2145_BA_peak_2146 | repressive | 6 |
| DS113344 | 121348 | 123486 | BA_peak_2151_BA_peak_2152 | repressive | 2 |
| DS113346 | 10441 | 17842 | BA_peak_2161_BA_peak_2162 | repressive | 3 |
| DS113346 | 17004 | 30391 | BA_peak_2162_BA_peak_2163 | repressive | 8 |
| DS113347 | 48228 | 53776 | BA_peak_2171_BA_peak_2172 | repressive | 2 |
| DS113347 | 52265 | 61747 | BA_peak_2172_BA_peak_2173 | repressive | 4 |
| DS113348 | 36761 | 62050 | BA_peak_2180_BA_peak_2181 | repressive | 25 |
| DS113349 | 116978 | 126815 | BA_peak_2193_BA_peak_2194 | repressive | 5 |
| DS113351 | 116735 | 123284 | BA_peak_2219_BA_peak_2220 | repressive | 3 |
| DS113352 | 26697 | 32930 | BA_peak_2222_BA_peak_2223 | repressive | 4 |
| DS113354 | 30219 | 50697 | BA_peak_2243_BA_peak_2244 | repressive | 11 |
| DS113355 | 2123 | 10164 | BA_peak_2249_BA_peak_2250 | repressive | 3 |
| DS113357 | 5427 | 12498 | BA_peak_2260_BA_peak_2261 | repressive | 3 |
| DS113358 | 68148 | 71481 | BA_peak_2272_BA_peak_2273 | repressive | 2 |
| DS113360 | 9148 | 15385 | BA_peak_2282_BA_peak_2283 | repressive | 2 |
| DS113361 | 81235 | 93173 | BA_peak_2296_BA_peak_2297 | repressive | 6 |
| DS113361 | 93700 | 99869 | BA_peak_2298_BA_peak_2299 | repressive | 3 |
| DS113362 | 19669 | 31544 | BA_peak_2303_BA_peak_2304 | repressive | 8 |
| DS113366 | 106192 | 117185 | BA_peak_2339_BA_peak_2340 | repressive | 3 |

| | | | | | |
|---|---|---|---|---|---|
| DS113368 | 68829 | 89549 | BA_peak_2353_BA_peak_2354 | repressive | 17 |
| DS113370 | 87284 | 102078 | BA_peak_2368_BA_peak_2369 | repressive | 8 |
| DS113371 | 29631 | 39150 | BA_peak_2374_BA_peak_2376 | repressive | 6 |
| DS113373 | 44848 | 56416 | BA_peak_2385_BA_peak_2387 | repressive | 5 |
| DS113373 | 68277 | 74552 | BA_peak_2389_BA_peak_2390 | repressive | 3 |
| DS113375 | 14522 | 41601 | BA_peak_2398_BA_peak_2400 | repressive | 15 |
| DS113377 | 51589 | 55359 | BA_peak_2410_BA_peak_2411 | repressive | 3 |
| DS113377 | 85120 | 101672 | BA_peak_2414_BA_peak_2415 | repressive | 12 |
| DS113379 | 67883 | 78831 | BA_peak_2425_BA_peak_2426 | repressive | 6 |
| DS113380 | 29606 | 35538 | BA_peak_2438_BA_peak_2439 | repressive | 2 |
| DS113380 | 51499 | 63175 | BA_peak_2442_BA_peak_2443 | repressive | 10 |
| DS113380 | 62622 | 69050 | BA_peak_2443_BA_peak_2444 | repressive | 4 |
| DS113380 | 81187 | 87505 | BA_peak_2445_BA_peak_2446 | repressive | 4 |
| DS113382 | 80026 | 90796 | BA_peak_2467_BA_peak_2468 | repressive | 9 |
| DS113387 | 29259 | 50095 | BA_peak_2504_BA_peak_2505 | repressive | 11 |
| DS113387 | 87161 | 110687 | BA_peak_2510_BA_peak_2511 | repressive | 19 |
| DS113388 | 68137 | 71923 | BA_peak_2522_BA_peak_2523 | repressive | 2 |
| DS113390 | 57220 | 73688 | BA_peak_2538_BA_peak_2539 | repressive | 12 |
| DS113390 | 89319 | 102422 | BA_peak_2542_BA_peak_2543 | repressive | 3 |
| DS113391 | 37581 | 55108 | BA_peak_2546_BA_peak_2547 | repressive | 10 |
| DS113396 | 27342 | 37355 | BA_peak_2585_BA_peak_2586 | repressive | 6 |
| DS113397 | 65006 | 70939 | BA_peak_2592_BA_peak_2593 | repressive | 3 |
| DS113401 | 30184 | 38591 | BA_peak_2621_BA_peak_2622 | repressive | 2 |
| DS113405 | 96886 | 104011 | BA_peak_2656_BA_peak_2657 | repressive | 4 |
| DS113408 | 63408 | 79726 | BA_peak_2666_BA_peak_2667 | repressive | 8 |
| DS113412 | 28722 | 40399 | BA_peak_2705_BA_peak_2706 | repressive | 4 |
| DS113414 | 42871 | 55781 | BA_peak_2716_BA_peak_2717 | repressive | 6 |
| DS113416 | 73846 | 82733 | BA_peak_2725_BA_peak_2726 | repressive | 6 |
| DS113417 | 454 | 14147 | BA_peak_2727_BA_peak_2728 | repressive | 9 |
| DS113422 | 43162 | 50694 | BA_peak_2752_BA_peak_2753 | repressive | 2 |
| DS113424 | 552 | 15781 | BA_peak_2758_BA_peak_2759 | repressive | 9 |
| DS113425 | 22848 | 32750 | BA_peak_2762_BA_peak_2763 | repressive | 5 |
| DS113429 | 73267 | 84316 | BA_peak_2788_BA_peak_2789 | repressive | 6 |
| DS113430 | 9314 | 21623 | BA_peak_2794_BA_peak_2795 | repressive | 6 |
| DS113430 | 22191 | 34859 | BA_peak_2796_BA_peak_2798 | repressive | 7 |
| DS113430 | 57812 | 65277 | BA_peak_2802_BA_peak_2803 | repressive | 2 |
| DS113430 | 70160 | 78485 | BA_peak_2805_BA_peak_2806 | repressive | 9 |
| DS113432 | 46304 | 58068 | BA_peak_2817_BA_peak_2818 | repressive | 7 |
| DS113433 | 3216 | 9979 | BA_peak_2819_BA_peak_2820 | repressive | 5 |
| DS113435 | 23880 | 27850 | BA_peak_2839_BA_peak_2840 | repressive | 2 |
| DS113437 | 94868 | 100573 | BA_peak_2859_BA_peak_2860 | repressive | 2 |
| DS113439 | 88 | 8978 | BA_peak_2866_BA_peak_2867 | repressive | 10 |
| DS113440 | 25772 | 36082 | BA_peak_2875_BA_peak_2876 | repressive | 6 |
| DS113441 | 60526 | 65640 | BA_peak_2883_BA_peak_2884 | repressive | 3 |
| DS113446 | 11640 | 18056 | BA_peak_2916_BA_peak_2917 | repressive | 2 |
| DS113449 | 55383 | 63117 | BA_peak_2937_BA_peak_2938 | repressive | 2 |
| DS113450 | 54021 | 63895 | BA_peak_2947_BA_peak_2948 | repressive | 4 |

| | | | | | |
|---|---|---|---|---|---|
| DS113451 | 65975 | 81323 | BA_peak_2952_BA_peak_2953 | repressive | 11 |
| DS113455 | 33079 | 56736 | BA_peak_2978_BA_peak_2979 | repressive | 17 |
| DS113456 | 55780 | 71941 | BA_peak_2987_BA_peak_2988 | repressive | 10 |
| DS113459 | 6730 | 8984 | BA_peak_2998_BA_peak_2999 | repressive | 2 |
| DS113460 | 6985 | 22292 | BA_peak_3004_BA_peak_3006 | repressive | 9 |
| DS113460 | 57483 | 67657 | BA_peak_3008_BA_peak_3009 | repressive | 5 |
| DS113460 | 66829 | 72261 | BA_peak_3009_BA_peak_3010 | repressive | 4 |
| DS113461 | 66024 | 76588 | BA_peak_3017_BA_peak_3018 | repressive | 8 |
| DS113468 | 46587 | 50398 | BA_peak_3056_BA_peak_3057 | repressive | 2 |
| DS113470 | 32725 | 52481 | BA_peak_3069_BA_peak_3070 | repressive | 12 |
| DS113472 | 8256 | 25239 | BA_peak_3080_BA_peak_3082 | repressive | 13 |
| DS113474 | 4892 | 14586 | BA_peak_3087_BA_peak_3088 | repressive | 2 |
| DS113475 | 28261 | 42173 | BA_peak_3094_BA_peak_3095 | repressive | 8 |
| DS113477 | 7879 | 40547 | BA_peak_3103_BA_peak_3104 | repressive | 17 |
| DS113479 | 2930 | 7069 | BA_peak_3111_BA_peak_3112 | repressive | 2 |
| DS113479 | 56138 | 60839 | BA_peak_3114_BA_peak_3115 | repressive | 2 |
| DS113479 | 59890 | 69726 | BA_peak_3115_BA_peak_3116 | repressive | 4 |
| DS113484 | 41630 | 49095 | BA_peak_3135_BA_peak_3136 | repressive | 7 |
| DS113484 | 69680 | 75224 | BA_peak_3139_BA_peak_3140 | repressive | 4 |
| DS113488 | 6206 | 13014 | BA_peak_3163_BA_peak_3164 | repressive | 3 |
| DS113489 | 4539 | 11236 | BA_peak_3168_BA_peak_3169 | repressive | 3 |
| DS113492 | 32 | 8776 | BA_peak_3180_BA_peak_3181 | repressive | 2 |
| DS113492 | 71295 | 75045 | BA_peak_3183_BA_peak_3184 | repressive | 2 |
| DS113494 | 19995 | 24661 | BA_peak_3193_BA_peak_3194 | repressive | 4 |
| DS113494 | 40099 | 54618 | BA_peak_3197_BA_peak_3198 | repressive | 9 |
| DS113495 | 65455 | 74019 | BA_peak_3207_BA_peak_3210 | repressive | 3 |
| DS113497 | 56409 | 64193 | BA_peak_3229_BA_peak_3230 | repressive | 4 |
| DS113498 | 3111 | 17035 | BA_peak_3233_BA_peak_3234 | repressive | 9 |
| DS113498 | 23018 | 38551 | BA_peak_3235_BA_peak_3237 | repressive | 10 |
| DS113500 | 12807 | 20279 | BA_peak_3245_BA_peak_3246 | repressive | 2 |
| DS113500 | 19822 | 32705 | BA_peak_3246_BA_peak_3247 | repressive | 10 |
| DS113500 | 56574 | 66639 | BA_peak_3251_BA_peak_3252 | repressive | 4 |
| DS113504 | 15780 | 31028 | BA_peak_3272_BA_peak_3273 | repressive | 8 |
| DS113507 | 43570 | 56929 | BA_peak_3289_BA_peak_3290 | repressive | 7 |
| DS113509 | 24740 | 32164 | BA_peak_3304_BA_peak_3305 | repressive | 3 |
| DS113510 | 44592 | 49776 | BA_peak_3310_BA_peak_3311 | repressive | 2 |
| DS113514 | 7734 | 23435 | BA_peak_3333_BA_peak_3334 | repressive | 10 |
| DS113514 | 39189 | 66426 | BA_peak_3338_BA_peak_3339 | repressive | 20 |
| DS113517 | 44741 | 65017 | BA_peak_3356_BA_peak_3358 | repressive | 12 |
| DS113518 | 176 | 8586 | BA_peak_3360_BA_peak_3361 | repressive | 5 |
| DS113520 | 25920 | 39792 | BA_peak_3372_BA_peak_3373 | repressive | 9 |
| DS113523 | 30164 | 32712 | BA_peak_3393_BA_peak_3394 | repressive | 3 |
| DS113523 | 76679 | 84866 | BA_peak_3398_BA_peak_3399 | repressive | 5 |
| DS113527 | 13896 | 28918 | BA_peak_3415_BA_peak_3416 | repressive | 8 |
| DS113528 | 38998 | 57032 | BA_peak_3421_BA_peak_3422 | repressive | 14 |
| DS113542 | 34813 | 45045 | BA_peak_3495_BA_peak_3496 | repressive | 5 |
| DS113544 | 30330 | 35942 | BA_peak_3506_BA_peak_3507 | repressive | 2 |

| | | | | | |
|---|---|---|---|---|---|
| DS113545 | 21550 | 67579 | BA_peak_3516_BA_peak_3518 | repressive | 26 |
| DS113546 | 23 | 11741 | BA_peak_3519_BA_peak_3520 | repressive | 8 |
| DS113546 | 32755 | 52687 | BA_peak_3524_BA_peak_3525 | repressive | 18 |
| DS113547 | 16168 | 22576 | BA_peak_3530_BA_peak_3531 | repressive | 2 |
| DS113550 | 50060 | 53474 | BA_peak_3560_BA_peak_3561 | repressive | 2 |
| DS113562 | 67702 | 77653 | BA_peak_3636_BA_peak_3637 | repressive | 5 |
| DS113564 | 7870 | 15772 | BA_peak_3640_BA_peak_3641 | repressive | 4 |
| DS113564 | 29785 | 38876 | BA_peak_3642_BA_peak_3643 | repressive | 9 |
| DS113566 | 55287 | 64725 | BA_peak_3656_BA_peak_3657 | repressive | 5 |
| DS113569 | 47 | 10811 | BA_peak_3667_BA_peak_3668 | repressive | 6 |
| DS113572 | 101 | 9325 | BA_peak_3681_BA_peak_3683 | repressive | 5 |
| DS113572 | 38136 | 48096 | BA_peak_3684_BA_peak_3685 | repressive | 6 |
| DS113576 | 59973 | 62230 | BA_peak_3708_BA_peak_3709 | repressive | 4 |
| DS113579 | 9713 | 28057 | BA_peak_3721_BA_peak_3722 | repressive | 5 |
| DS113579 | 27153 | 36731 | BA_peak_3722_BA_peak_3724 | repressive | 3 |
| DS113579 | 36256 | 54131 | BA_peak_3724_BA_peak_3725 | repressive | 3 |
| DS113580 | 53840 | 64614 | BA_peak_3732_BA_peak_3733 | repressive | 6 |
| DS113581 | 60849 | 72251 | BA_peak_3736_BA_peak_3737 | repressive | 12 |
| DS113583 | 10046 | 24129 | BA_peak_3746_BA_peak_3747 | repressive | 11 |
| DS113584 | 25823 | 40556 | BA_peak_3750_BA_peak_3752 | repressive | 10 |
| DS113587 | 26021 | 50398 | BA_peak_3761_BA_peak_3762 | repressive | 14 |
| DS113591 | 62837 | 75333 | BA_peak_3776_BA_peak_3777 | repressive | 7 |
| DS113593 | 11392 | 30214 | BA_peak_3781_BA_peak_3782 | repressive | 20 |
| DS113598 | 69276 | 74598 | BA_peak_3805_BA_peak_3807 | repressive | 2 |
| DS113600 | 204 | 16523 | BA_peak_3812_BA_peak_3813 | repressive | 5 |
| DS113600 | 16048 | 30781 | BA_peak_3813_BA_peak_3814 | repressive | 7 |
| DS113600 | 29443 | 49031 | BA_peak_3814_BA_peak_3815 | repressive | 6 |
| DS113602 | 39208 | 50927 | BA_peak_3820_BA_peak_3821 | repressive | 8 |
| DS113605 | 18025 | 33768 | BA_peak_3840_BA_peak_3841 | repressive | 9 |
| DS113607 | 1926 | 5049 | BA_peak_3849_BA_peak_3850 | repressive | 2 |
| DS113615 | 9410 | 18198 | BA_peak_3881_BA_peak_3882 | repressive | 6 |
| DS113617 | 30138 | 33931 | BA_peak_3889_BA_peak_3890 | repressive | 2 |
| DS113618 | 39007 | 52183 | BA_peak_3895_BA_peak_3897 | repressive | 7 |
| DS113619 | 19044 | 37206 | BA_peak_3900_BA_peak_3901 | repressive | 16 |
| DS113620 | 6723 | 14045 | BA_peak_3902_BA_peak_3903 | repressive | 3 |
| DS113623 | 46859 | 57382 | BA_peak_3920_BA_peak_3921 | repressive | 6 |
| DS113629 | 35334 | 51963 | BA_peak_3939_BA_peak_3940 | repressive | 5 |
| DS113639 | 764 | 7654 | BA_peak_3983_BA_peak_3984 | repressive | 6 |
| DS113640 | 15314 | 23269 | BA_peak_3993_BA_peak_3994 | repressive | 7 |
| DS113641 | 62389 | 66484 | BA_peak_4000_BA_peak_4001 | repressive | 4 |
| DS113649 | 6237 | 44737 | BA_peak_4018_BA_peak_4019 | repressive | 26 |
| DS113650 | 27911 | 36982 | BA_peak_4023_BA_peak_4024 | repressive | 6 |
| DS113654 | 9762 | 18690 | BA_peak_4036_BA_peak_4037 | repressive | 2 |
| DS113658 | 26739 | 39378 | BA_peak_4055_BA_peak_4056 | repressive | 5 |
| DS113667 | 14619 | 20070 | BA_peak_4096_BA_peak_4097 | repressive | 2 |
| DS113672 | 22033 | 36454 | BA_peak_4112_BA_peak_4113 | repressive | 9 |
| DS113688 | 14937 | 32039 | BA_peak_4177_BA_peak_4178 | repressive | 9 |

| | | | | | |
|---|---|---|---|---|---|
| DS113688 | 58330 | 63784 | BA_peak_4180_BA_peak_4181 | repressive | 3 |
| DS113701 | 13777 | 21705 | BA_peak_4239_BA_peak_4240 | repressive | 2 |
| DS113702 | 1008 | 6350 | BA_peak_4244_BA_peak_4245 | repressive | 2 |
| DS113704 | 43687 | 58027 | BA_peak_4251_BA_peak_4252 | repressive | 13 |
| DS113707 | 50332 | 62586 | BA_peak_4260_BA_peak_4261 | repressive | 10 |
| DS113709 | 4087 | 10512 | BA_peak_4271_BA_peak_4272 | repressive | 3 |
| DS113711 | 16625 | 39755 | BA_peak_4280_BA_peak_4281 | repressive | 17 |
| DS113714 | 18449 | 30384 | BA_peak_4301_BA_peak_4302 | repressive | 5 |
| DS113729 | 19660 | 41635 | BA_peak_4353_BA_peak_4354 | repressive | 16 |
| DS113731 | 33940 | 38905 | BA_peak_4361_BA_peak_4362 | repressive | 2 |
| DS113738 | 43406 | 57664 | BA_peak_4391_BA_peak_4392 | repressive | 5 |
| DS113739 | 14055 | 18082 | BA_peak_4395_BA_peak_4396 | repressive | 2 |
| DS113746 | 20518 | 36948 | BA_peak_4427_BA_peak_4428 | repressive | 11 |
| DS113747 | 3479 | 10379 | BA_peak_4432_BA_peak_4433 | repressive | 7 |
| DS113749 | 5704 | 10039 | BA_peak_4438_BA_peak_4439 | repressive | 3 |
| DS113752 | 32805 | 44728 | BA_peak_4453_BA_peak_4454 | repressive | 10 |
| DS113758 | 30533 | 34337 | BA_peak_4483_BA_peak_4484 | repressive | 3 |
| DS113758 | 33600 | 39515 | BA_peak_4484_BA_peak_4485 | repressive | 3 |
| DS113765 | 20020 | 23117 | BA_peak_4505_BA_peak_4506 | repressive | 2 |
| DS113766 | 27705 | 33544 | BA_peak_4509_BA_peak_4510 | repressive | 3 |
| DS113770 | 26112 | 30119 | BA_peak_4520_BA_peak_4521 | repressive | 2 |
| DS113771 | 24994 | 30555 | BA_peak_4524_BA_peak_4525 | repressive | 2 |
| DS113775 | 44762 | 49687 | BA_peak_4543_BA_peak_4545 | repressive | 2 |
| DS113777 | 5457 | 28461 | BA_peak_4551_BA_peak_4552 | repressive | 19 |
| DS113778 | 14710 | 28801 | BA_peak_4554_BA_peak_4555 | repressive | 9 |
| DS113779 | 24441 | 36839 | BA_peak_4560_BA_peak_4561 | repressive | 8 |
| DS113782 | 29889 | 39745 | BA_peak_4572_BA_peak_4573 | repressive | 4 |
| DS113783 | 260 | 7836 | BA_peak_4577_BA_peak_4578 | repressive | 3 |
| DS113799 | 40207 | 46583 | BA_peak_4637_BA_peak_4638 | repressive | 5 |
| DS113802 | 32115 | 53917 | BA_peak_4647_BA_peak_4648 | repressive | 7 |
| DS113803 | 48325 | 53754 | BA_peak_4655_BA_peak_4656 | repressive | 2 |
| DS113818 | 25404 | 43838 | BA_peak_4695_BA_peak_4696 | repressive | 14 |
| DS113827 | 985 | 11015 | BA_peak_4722_BA_peak_4723 | repressive | 5 |
| DS113836 | 12763 | 17330 | BA_peak_4742_BA_peak_4743 | repressive | 3 |
| DS113836 | 16372 | 21116 | BA_peak_4743_BA_peak_4744 | repressive | 2 |
| DS113837 | 32009 | 36513 | BA_peak_4752_BA_peak_4753 | repressive | 2 |
| DS113843 | 26353 | 44778 | BA_peak_4773_BA_peak_4774 | repressive | 11 |
| DS113847 | 41481 | 45773 | BA_peak_4791_BA_peak_4792 | repressive | 2 |
| DS113848 | 7396 | 10745 | BA_peak_4793_BA_peak_4794 | repressive | 2 |
| DS113852 | 43530 | 46422 | BA_peak_4813_BA_peak_4814 | repressive | 3 |
| DS113853 | 6063 | 11172 | BA_peak_4815_BA_peak_4816 | repressive | 4 |
| DS113859 | 21245 | 31640 | BA_peak_4829_BA_peak_4830 | repressive | 6 |
| DS113867 | 12218 | 29470 | BA_peak_4860_BA_peak_4861 | repressive | 13 |
| DS113870 | 24344 | 31384 | BA_peak_4869_BA_peak_4870 | repressive | 2 |
| DS113875 | 18888 | 32940 | BA_peak_4894_BA_peak_4895 | repressive | 10 |
| DS113877 | 18673 | 28572 | BA_peak_4900_BA_peak_4901 | repressive | 6 |
| DS113880 | 961 | 14322 | BA_peak_4906_BA_peak_4907 | repressive | 4 |

| | | | | | |
|---|---|---|---|---|---|
| DS113880 | 14801 | 28680 | BA_peak_4908_BA_peak_4909 | repressive | 5 |
| DS113882 | 39174 | 44723 | BA_peak_4915_BA_peak_4916 | repressive | 3 |
| DS113888 | 38093 | 43755 | BA_peak_4937_BA_peak_4938 | repressive | 2 |
| DS113889 | 37318 | 41394 | BA_peak_4944_BA_peak_4945 | repressive | 2 |
| DS113906 | 38 | 5450 | BA_peak_4990_BA_peak_4991 | repressive | 2 |
| DS113907 | 5331 | 9020 | BA_peak_4999_BA_peak_5000 | repressive | 2 |
| DS113914 | 22982 | 29694 | BA_peak_5019_BA_peak_5020 | repressive | 3 |
| DS113915 | 1728 | 6611 | BA_peak_5021_BA_peak_5022 | repressive | 2 |
| DS113915 | 12966 | 33622 | BA_peak_5024_BA_peak_5025 | repressive | 13 |
| DS113927 | 22048 | 42061 | BA_peak_5054_BA_peak_5055 | repressive | 23 |
| DS113930 | 4071 | 15890 | BA_peak_5057_BA_peak_5058 | repressive | 4 |
| DS113937 | 10840 | 13916 | BA_peak_5073_BA_peak_5074 | repressive | 2 |
| DS113945 | 4360 | 15552 | BA_peak_5096_BA_peak_5097 | repressive | 9 |
| DS113948 | 431 | 9150 | BA_peak_5106_BA_peak_5107 | repressive | 5 |
| DS113951 | 16108 | 23045 | BA_peak_5120_BA_peak_5121 | repressive | 4 |
| DS113954 | 948 | 10290 | BA_peak_5130_BA_peak_5131 | repressive | 6 |
| DS113966 | 18045 | 34914 | BA_peak_5166_BA_peak_5167 | repressive | 13 |
| DS113968 | 6663 | 12204 | BA_peak_5169_BA_peak_5170 | repressive | 2 |
| DS113976 | 7091 | 11195 | BA_peak_5197_BA_peak_5198 | repressive | 2 |
| DS113980 | 23577 | 31414 | BA_peak_5221_BA_peak_5222 | repressive | 3 |
| DS113981 | 22550 | 27389 | BA_peak_5225_BA_peak_5226 | repressive | 5 |
| DS113983 | 16833 | 26033 | BA_peak_5233_BA_peak_5234 | repressive | 2 |
| DS113984 | 5151 | 15483 | BA_peak_5236_BA_peak_5237 | repressive | 4 |
| DS113992 | 25551 | 28167 | BA_peak_5264_BA_peak_5265 | repressive | 2 |
| DS113994 | 25316 | 33810 | BA_peak_5269_BA_peak_5270 | repressive | 5 |
| DS113998 | 3453 | 8503 | BA_peak_5281_BA_peak_5282 | repressive | 3 |
| DS114002 | 14931 | 34905 | BA_peak_5288_BA_peak_5289 | repressive | 19 |
| DS114004 | 13923 | 19985 | BA_peak_5290_BA_peak_5291 | repressive | 3 |
| DS114004 | 29163 | 36632 | BA_peak_5294_BA_peak_5295 | repressive | 4 |
| DS114015 | 16594 | 24346 | BA_peak_5321_BA_peak_5322 | repressive | 5 |
| DS114019 | 733 | 11905 | BA_peak_5331_BA_peak_5332 | repressive | 5 |
| DS114032 | 29167 | 33037 | BA_peak_5364_BA_peak_5365 | repressive | 2 |
| DS114058 | 9465 | 14990 | BA_peak_5423_BA_peak_5424 | repressive | 2 |
| DS114061 | 19666 | 27212 | BA_peak_5432_BA_peak_5433 | repressive | 10 |
| DS114069 | 6671 | 11399 | BA_peak_5454_BA_peak_5455 | repressive | 3 |
| DS114071 | 1189 | 26970 | BA_peak_5460_BA_peak_5461 | repressive | 22 |
| DS114074 | 22842 | 31933 | BA_peak_5467_BA_peak_5468 | repressive | 2 |
| DS114076 | 20724 | 25637 | BA_peak_5473_BA_peak_5474 | repressive | 2 |
| DS114076 | 26366 | 31734 | BA_peak_5475_BA_peak_5476 | repressive | 4 |
| DS114079 | 5959 | 20226 | BA_peak_5481_BA_peak_5483 | repressive | 8 |
| DS114093 | 2932 | 19929 | BA_peak_5512_BA_peak_5513 | repressive | 11 |
| DS114115 | 1938 | 26540 | BA_peak_5544_BA_peak_5545 | repressive | 9 |
| DS114123 | 5017 | 8725 | BA_peak_5559_BA_peak_5560 | repressive | 2 |
| DS114127 | 3372 | 9240 | BA_peak_5565_BA_peak_5566 | repressive | 3 |
| DS114128 | 147 | 13997 | BA_peak_5569_BA_peak_5570 | repressive | 6 |
| DS114134 | 7917 | 21375 | BA_peak_5583_BA_peak_5584 | repressive | 6 |
| DS114161 | 1358 | 13060 | BA_peak_5633_BA_peak_5634 | repressive | 9 |

| | | | | | |
|---|---|---|---|---|---|
| DS114165 | 21027 | 25201 | BA_peak_5644_BA_peak_5645 | repressive | 3 |
| DS114174 | 939 | 7631 | BA_peak_5667_BA_peak_5668 | repressive | 4 |
| DS114176 | 6043 | 20641 | BA_peak_5672_BA_peak_5673 | repressive | 9 |
| DS114178 | 13636 | 27271 | BA_peak_5677_BA_peak_5678 | repressive | 10 |
| DS114192 | 2891 | 22962 | BA_peak_5706_BA_peak_5707 | repressive | 16 |
| DS114196 | 2528 | 8814 | BA_peak_5710_BA_peak_5711 | repressive | 6 |
| DS114200 | 19699 | 26750 | BA_peak_5716_BA_peak_5717 | repressive | 2 |
| DS114202 | 16137 | 22283 | BA_peak_5720_BA_peak_5721 | repressive | 5 |
| DS114207 | 15061 | 17704 | BA_peak_5728_BA_peak_5728 | repressive | 2 |
| DS114230 | 3091 | 9980 | BA_peak_5766_BA_peak_5767 | repressive | 3 |
| DS114232 | 6589 | 18383 | BA_peak_5774_BA_peak_5775 | repressive | 4 |
| DS114270 | 725 | 12265 | BA_peak_5847_BA_peak_5848 | repressive | 7 |
| DS114273 | 1368 | 16802 | BA_peak_5855_BA_peak_5856 | repressive | 4 |
| DS114333 | 6570 | 21269 | BA_peak_5942_BA_peak_5943 | repressive | 10 |
| DS114358 | 268 | 6407 | BA_peak_5983_BA_peak_5984 | repressive | 3 |
| DS114372 | 2165 | 10827 | BA_peak_5999_BA_peak_6000 | repressive | 5 |
| DS114407 | 235 | 9949 | BA_peak_6048_BA_peak_6049 | repressive | 4 |
| DS114430 | 11099 | 15568 | BA_peak_6080_BA_peak_6081 | repressive | 2 |
| DS114431 | 840 | 9495 | BA_peak_6083_BA_peak_6084 | repressive | 7 |
| DS114455 | 26 | 4926 | BA_peak_6111_BA_peak_6112 | repressive | 2 |
| DS114477 | 12969 | 15461 | BA_peak_6129_BA_peak_6130 | repressive | 2 |
| DS114481 | 4147 | 10890 | BA_peak_6137_BA_peak_6138 | repressive | 4 |
| DS114495 | 8755 | 15649 | BA_peak_6150_BA_peak_6151 | repressive | 2 |
| DS114497 | 1945 | 6776 | BA_peak_6154_BA_peak_6155 | repressive | 2 |
| DS114501 | 1125 | 14836 | BA_peak_6162_BA_peak_6163 | repressive | 4 |
| DS114545 | 3201 | 7333 | BA_peak_6214_BA_peak_6215 | repressive | 2 |
| DS114564 | 1277 | 10535 | BA_peak_6239_BA_peak_6240 | repressive | 4 |
| DS114642 | 5055 | 8639 | BA_peak_6303_BA_peak_6304 | repressive | 2 |
| DS114642 | 7497 | 11872 | BA_peak_6304_BA_peak_6305 | repressive | 3 |
| DS114936 | 441 | 6358 | BA_peak_6507_BA_peak_6508 | repressive | 3 |
| DS114946 | 789 | 3843 | BA_peak_6514_BA_peak_6515 | repressive | 2 |
| DS116470 | 164 | 2402 | BA_peak_6942_BA_peak_6943 | repressive | 2 |
| DS113177 | 260812 | 270312 | BA_peak_9_BA_peak_10 | active | 5 |
| DS113177 | 456517 | 480926 | BA_peak_16_BA_peak_18 | active | 8 |
| DS113178 | 2093 | 7672 | BA_peak_22_BA_peak_23 | active | 2 |
| DS113179 | 186575 | 202163 | BA_peak_59_BA_peak_60 | active | 6 |
| DS113179 | 260776 | 272801 | BA_peak_66_BA_peak_67 | active | 5 |
| DS113179 | 282272 | 288852 | BA_peak_69_BA_peak_70 | active | 2 |
| DS113179 | 340168 | 345091 | BA_peak_76_BA_peak_77 | active | 2 |
| DS113179 | 468930 | 476916 | BA_peak_91_BA_peak_92 | active | 4 |
| DS113180 | 102625 | 112986 | BA_peak_100_BA_peak_101 | active | 4 |
| DS113180 | 124577 | 135620 | BA_peak_103_BA_peak_104 | active | 4 |
| DS113180 | 202424 | 248088 | BA_peak_108_BA_peak_109 | active | 19 |
| DS113180 | 250564 | 274802 | BA_peak_110_BA_peak_111 | active | 7 |
| DS113181 | 220292 | 225619 | BA_peak_138_BA_peak_139 | active | 2 |
| DS113181 | 225142 | 235429 | BA_peak_139_BA_peak_140 | active | 5 |
| DS113181 | 238940 | 244484 | BA_peak_142_BA_peak_143 | active | 2 |

| | | | | | |
|---|---|---|---|---|---|
| DS113181 | 341678 | 346111 | BA_peak_147_BA_peak_148 | active | 2 |
| DS113181 | 426653 | 432659 | BA_peak_152_BA_peak_153 | active | 2 |
| DS113182 | 236918 | 252370 | BA_peak_166_BA_peak_167 | active | 4 |
| DS113182 | 309474 | 313490 | BA_peak_170_BA_peak_171 | active | 2 |
| DS113182 | 312799 | 318934 | BA_peak_171_BA_peak_172 | active | 3 |
| DS113183 | 197988 | 206711 | BA_peak_189_BA_peak_190 | active | 3 |
| DS113183 | 227048 | 232936 | BA_peak_191_BA_peak_192 | active | 2 |
| DS113184 | 23548 | 29869 | BA_peak_206_BA_peak_207 | active | 2 |
| DS113184 | 32382 | 47612 | BA_peak_208_BA_peak_209 | active | 3 |
| DS113184 | 126839 | 138662 | BA_peak_220_BA_peak_221 | active | 2 |
| DS113185 | 238020 | 257887 | BA_peak_250_BA_peak_251 | active | 8 |
| DS113186 | 118481 | 135910 | BA_peak_271_BA_peak_272 | active | 6 |
| DS113186 | 217773 | 228329 | BA_peak_276_BA_peak_277 | active | 4 |
| DS113186 | 378791 | 396786 | BA_peak_287_BA_peak_288 | active | 4 |
| DS113187 | 67598 | 72387 | BA_peak_296_BA_peak_297 | active | 2 |
| DS113187 | 106005 | 120749 | BA_peak_299_BA_peak_300 | active | 5 |
| DS113187 | 120351 | 131952 | BA_peak_300_BA_peak_301 | active | 3 |
| DS113187 | 131231 | 141140 | BA_peak_301_BA_peak_302 | active | 4 |
| DS113187 | 244056 | 250939 | BA_peak_309_BA_peak_310 | active | 2 |
| DS113187 | 250500 | 266407 | BA_peak_310_BA_peak_311 | active | 6 |
| DS113189 | 318335 | 331860 | BA_peak_366_BA_peak_367 | active | 2 |
| DS113190 | 14033 | 19722 | BA_peak_376_BA_peak_377 | active | 2 |
| DS113190 | 170131 | 176857 | BA_peak_383_BA_peak_384 | active | 2 |
| DS113190 | 208793 | 225875 | BA_peak_387_BA_peak_388 | active | 6 |
| DS113191 | 83644 | 106108 | BA_peak_394_BA_peak_395 | active | 11 |
| DS113191 | 105418 | 111432 | BA_peak_395_BA_peak_396 | active | 2 |
| DS113191 | 111030 | 118766 | BA_peak_396_BA_peak_398 | active | 2 |
| DS113191 | 311695 | 321112 | BA_peak_415_BA_peak_416 | active | 2 |
| DS113193 | 45105 | 66326 | BA_peak_435_BA_peak_438 | active | 4 |
| DS113193 | 65228 | 77364 | BA_peak_438_BA_peak_439 | active | 3 |
| DS113194 | 177316 | 188530 | BA_peak_453_BA_peak_454 | active | 2 |
| DS113194 | 238773 | 264312 | BA_peak_456_BA_peak_457 | active | 6 |
| DS113195 | 14566 | 27286 | BA_peak_462_BA_peak_463 | active | 4 |
| DS113195 | 206232 | 226290 | BA_peak_469_BA_peak_470 | active | 6 |
| DS113195 | 242926 | 251627 | BA_peak_472_BA_peak_473 | active | 4 |
| DS113195 | 272690 | 284435 | BA_peak_475_BA_peak_476 | active | 5 |
| DS113196 | 191989 | 197819 | BA_peak_489_BA_peak_490 | active | 2 |
| DS113197 | 216540 | 235584 | BA_peak_503_BA_peak_504 | active | 6 |
| DS113198 | 40055 | 55130 | BA_peak_512_BA_peak_514 | active | 3 |
| DS113198 | 239093 | 246457 | BA_peak_523_BA_peak_524 | active | 2 |
| DS113198 | 265049 | 279462 | BA_peak_526_BA_peak_527 | active | 8 |
| DS113199 | 161987 | 173675 | BA_peak_535_BA_peak_536 | active | 5 |
| DS113200 | 28880 | 36267 | BA_peak_545_BA_peak_546 | active | 4 |
| DS113200 | 93960 | 99369 | BA_peak_548_BA_peak_549 | active | 3 |
| DS113200 | 223704 | 251541 | BA_peak_556_BA_peak_557 | active | 9 |
| DS113201 | 72793 | 112325 | BA_peak_560_BA_peak_561 | active | 13 |
| DS113201 | 163039 | 173885 | BA_peak_563_BA_peak_564 | active | 3 |

| | | | | | |
|---|---|---|---|---|---|
| DS113201 | 173191 | 195236 | BA_peak_564_BA_peak_565 | active | 13 |
| DS113201 | 216221 | 223962 | BA_peak_568_BA_peak_569 | active | 4 |
| DS113201 | 224330 | 244853 | BA_peak_570_BA_peak_572 | active | 9 |
| DS113202 | 29507 | 47664 | BA_peak_576_BA_peak_577 | active | 6 |
| DS113202 | 87520 | 100546 | BA_peak_580_BA_peak_581 | active | 5 |
| DS113202 | 158594 | 167582 | BA_peak_585_BA_peak_586 | active | 5 |
| DS113203 | 1234 | 12711 | BA_peak_599_BA_peak_600 | active | 5 |
| DS113203 | 120844 | 139642 | BA_peak_605_BA_peak_606 | active | 10 |
| DS113203 | 138866 | 145653 | BA_peak_606_BA_peak_607 | active | 2 |
| DS113205 | 178689 | 188652 | BA_peak_648_BA_peak_649 | active | 6 |
| DS113206 | 12653 | 16701 | BA_peak_656_BA_peak_657 | active | 2 |
| DS113206 | 174396 | 186362 | BA_peak_673_BA_peak_674 | active | 4 |
| DS113207 | 77425 | 101173 | BA_peak_683_BA_peak_684 | active | 6 |
| DS113207 | 150691 | 173539 | BA_peak_690_BA_peak_692 | active | 11 |
| DS113207 | 179409 | 200969 | BA_peak_695_BA_peak_696 | active | 8 |
| DS113208 | 8696 | 25620 | BA_peak_700_BA_peak_701 | active | 3 |
| DS113208 | 24318 | 35260 | BA_peak_701_BA_peak_702 | active | 5 |
| DS113208 | 126579 | 133484 | BA_peak_711_BA_peak_712 | active | 3 |
| DS113208 | 131362 | 138811 | BA_peak_712_BA_peak_713 | active | 2 |
| DS113208 | 163626 | 173674 | BA_peak_715_BA_peak_716 | active | 4 |
| DS113208 | 198706 | 213279 | BA_peak_719_BA_peak_720 | active | 4 |
| DS113209 | 60139 | 80600 | BA_peak_728_BA_peak_729 | active | 4 |
| DS113212 | 40394 | 48770 | BA_peak_762_BA_peak_763 | active | 7 |
| DS113212 | 116587 | 125433 | BA_peak_769_BA_peak_770 | active | 2 |
| DS113213 | 1723 | 5827 | BA_peak_777_BA_peak_778 | active | 2 |
| DS113213 | 62209 | 71548 | BA_peak_779_BA_peak_780 | active | 3 |
| DS113213 | 86004 | 95423 | BA_peak_783_BA_peak_784 | active | 2 |
| DS113213 | 172118 | 178230 | BA_peak_789_BA_peak_790 | active | 2 |
| DS113214 | 108404 | 120658 | BA_peak_800_BA_peak_801 | active | 2 |
| DS113215 | 15673 | 35204 | BA_peak_806_BA_peak_807 | active | 8 |
| DS113215 | 168409 | 176489 | BA_peak_815_BA_peak_816 | active | 3 |
| DS113216 | 194903 | 218329 | BA_peak_825_BA_peak_826 | active | 8 |
| DS113217 | 40597 | 46167 | BA_peak_831_BA_peak_832 | active | 2 |
| DS113217 | 98542 | 106751 | BA_peak_835_BA_peak_836 | active | 5 |
| DS113217 | 106095 | 111938 | BA_peak_836_BA_peak_837 | active | 2 |
| DS113218 | 81409 | 90558 | BA_peak_847_BA_peak_848 | active | 2 |
| DS113218 | 110746 | 123518 | BA_peak_849_BA_peak_850 | active | 4 |
| DS113220 | 129188 | 133557 | BA_peak_874_BA_peak_875 | active | 2 |
| DS113222 | 170854 | 178223 | BA_peak_901_BA_peak_902 | active | 4 |
| DS113222 | 188843 | 196890 | BA_peak_905_BA_peak_906 | active | 3 |
| DS113224 | 101311 | 109614 | BA_peak_917_BA_peak_918 | active | 2 |
| DS113224 | 171066 | 180458 | BA_peak_922_BA_peak_923 | active | 4 |
| DS113225 | 36991 | 50104 | BA_peak_929_BA_peak_930 | active | 9 |
| DS113227 | 118426 | 137438 | BA_peak_951_BA_peak_952 | active | 4 |
| DS113227 | 136991 | 147292 | BA_peak_952_BA_peak_953 | active | 7 |
| DS113229 | 96929 | 117219 | BA_peak_971_BA_peak_972 | active | 11 |
| DS113229 | 128955 | 135473 | BA_peak_973_BA_peak_974 | active | 2 |

| | | | | | |
|---|---|---|---|---|---|
| DS113230 | 85921 | 102145 | BA_peak_989_BA_peak_990 | active | 9 |
| DS113231 | 135621 | 144697 | BA_peak_1001_BA_peak_1002 | active | 4 |
| DS113233 | 86505 | 96477 | BA_peak_1015_BA_peak_1016 | active | 4 |
| DS113235 | 30165 | 40566 | BA_peak_1036_BA_peak_1037 | active | 5 |
| DS113235 | 94020 | 97559 | BA_peak_1040_BA_peak_1041 | active | 2 |
| DS113236 | 71975 | 79239 | BA_peak_1050_BA_peak_1051 | active | 3 |
| DS113236 | 80158 | 89632 | BA_peak_1052_BA_peak_1053 | active | 2 |
| DS113236 | 196963 | 201691 | BA_peak_1063_BA_peak_1064 | active | 2 |
| DS113237 | 16931 | 38979 | BA_peak_1065_BA_peak_1066 | active | 6 |
| DS113237 | 56538 | 63530 | BA_peak_1070_BA_peak_1071 | active | 2 |
| DS113239 | 131053 | 140024 | BA_peak_1101_BA_peak_1102 | active | 4 |
| DS113239 | 161567 | 174594 | BA_peak_1103_BA_peak_1104 | active | 7 |
| DS113241 | 22171 | 30661 | BA_peak_1117_BA_peak_1118 | active | 3 |
| DS113244 | 161275 | 169455 | BA_peak_1165_BA_peak_1166 | active | 2 |
| DS113245 | 52299 | 67961 | BA_peak_1170_BA_peak_1171 | active | 5 |
| DS113245 | 135652 | 144827 | BA_peak_1175_BA_peak_1176 | active | 4 |
| DS113245 | 170647 | 176123 | BA_peak_1180_BA_peak_1182 | active | 3 |
| DS113245 | 175567 | 191323 | BA_peak_1182_BA_peak_1183 | active | 10 |
| DS113245 | 191013 | 196377 | BA_peak_1183_BA_peak_1184 | active | 3 |
| DS113248 | 67408 | 72980 | BA_peak_1206_BA_peak_1207 | active | 3 |
| DS113248 | 161428 | 176798 | BA_peak_1211_BA_peak_1212 | active | 5 |
| DS113249 | 113853 | 122048 | BA_peak_1220_BA_peak_1221 | active | 4 |
| DS113249 | 127208 | 132252 | BA_peak_1222_BA_peak_1223 | active | 2 |
| DS113250 | 41981 | 62869 | BA_peak_1228_BA_peak_1229 | active | 9 |
| DS113252 | 37429 | 47214 | BA_peak_1246_BA_peak_1247 | active | 3 |
| DS113253 | 1321 | 19689 | BA_peak_1257_BA_peak_1258 | active | 3 |
| DS113254 | 92559 | 112899 | BA_peak_1277_BA_peak_1278 | active | 6 |
| DS113256 | 78735 | 86650 | BA_peak_1291_BA_peak_1292 | active | 4 |
| DS113256 | 142806 | 148005 | BA_peak_1294_BA_peak_1295 | active | 2 |
| DS113256 | 147644 | 155732 | BA_peak_1295_BA_peak_1296 | active | 4 |
| DS113257 | 74416 | 80609 | BA_peak_1304_BA_peak_1305 | active | 3 |
| DS113258 | 135633 | 142351 | BA_peak_1317_BA_peak_1318 | active | 2 |
| DS113259 | 117972 | 127201 | BA_peak_1326_BA_peak_1327 | active | 3 |
| DS113259 | 133168 | 141607 | BA_peak_1330_BA_peak_1331 | active | 3 |
| DS113261 | 25993 | 44183 | BA_peak_1346_BA_peak_1347 | active | 9 |
| DS113262 | 8736 | 14392 | BA_peak_1353_BA_peak_1354 | active | 2 |
| DS113263 | 9284 | 21489 | BA_peak_1362_BA_peak_1363 | active | 4 |
| DS113263 | 81908 | 99767 | BA_peak_1365_BA_peak_1366 | active | 6 |
| DS113263 | 128335 | 131266 | BA_peak_1368_BA_peak_1369 | active | 2 |
| DS113264 | 171107 | 177107 | BA_peak_1373_BA_peak_1374 | active | 3 |
| DS113266 | 112331 | 139588 | BA_peak_1399_BA_peak_1400 | active | 7 |
| DS113266 | 138730 | 158304 | BA_peak_1400_BA_peak_1401 | active | 7 |
| DS113267 | 112983 | 118683 | BA_peak_1412_BA_peak_1413 | active | 2 |
| DS113268 | 52495 | 59929 | BA_peak_1425_BA_peak_1426 | active | 4 |
| DS113268 | 59398 | 85382 | BA_peak_1426_BA_peak_1427 | active | 11 |
| DS113268 | 85069 | 96276 | BA_peak_1427_BA_peak_1428 | active | 3 |
| DS113271 | 145135 | 166647 | BA_peak_1462_BA_peak_1463 | active | 10 |

| | | | | | |
|---|---|---|---|---|---|
| DS113272 | 50225 | 59786 | BA_peak_1468_BA_peak_1469 | active | 4 |
| DS113272 | 110604 | 117612 | BA_peak_1471_BA_peak_1472 | active | 3 |
| DS113272 | 129411 | 139157 | BA_peak_1474_BA_peak_1475 | active | 4 |
| DS113274 | 2979 | 16322 | BA_peak_1493_BA_peak_1494 | active | 4 |
| DS113274 | 42395 | 53204 | BA_peak_1495_BA_peak_1496 | active | 2 |
| DS113274 | 120327 | 139648 | BA_peak_1503_BA_peak_1504 | active | 5 |
| DS113275 | 6630 | 13834 | BA_peak_1509_BA_peak_1510 | active | 2 |
| DS113275 | 141937 | 162568 | BA_peak_1513_BA_peak_1514 | active | 9 |
| DS113276 | 38529 | 52540 | BA_peak_1518_BA_peak_1519 | active | 8 |
| DS113277 | 52684 | 56753 | BA_peak_1523_BA_peak_1524 | active | 2 |
| DS113277 | 76904 | 92851 | BA_peak_1525_BA_peak_1526 | active | 6 |
| DS113278 | 62063 | 90618 | BA_peak_1528_BA_peak_1529 | active | 7 |
| DS113279 | 10882 | 21333 | BA_peak_1534_BA_peak_1535 | active | 3 |
| DS113279 | 23153 | 28896 | BA_peak_1536_BA_peak_1537 | active | 2 |
| DS113279 | 26938 | 49522 | BA_peak_1537_BA_peak_1538 | active | 9 |
| DS113279 | 160681 | 168030 | BA_peak_1552_BA_peak_1553 | active | 2 |
| DS113280 | 107515 | 119873 | BA_peak_1562_BA_peak_1563 | active | 8 |
| DS113280 | 120616 | 130463 | BA_peak_1564_BA_peak_1565 | active | 5 |
| DS113280 | 129968 | 136362 | BA_peak_1565_BA_peak_1566 | active | 3 |
| DS113280 | 141245 | 145276 | BA_peak_1567_BA_peak_1568 | active | 2 |
| DS113281 | 75301 | 93854 | BA_peak_1574_BA_peak_1575 | active | 7 |
| DS113282 | 10299 | 17604 | BA_peak_1581_BA_peak_1582 | active | 2 |
| DS113282 | 35124 | 43987 | BA_peak_1583_BA_peak_1584 | active | 2 |
| DS113282 | 116756 | 124559 | BA_peak_1587_BA_peak_1588 | active | 2 |
| DS113284 | 95684 | 108493 | BA_peak_1602_BA_peak_1603 | active | 4 |
| DS113287 | 106948 | 132952 | BA_peak_1625_BA_peak_1627 | active | 14 |
| DS113287 | 144641 | 152842 | BA_peak_1629_BA_peak_1630 | active | 4 |
| DS113288 | 4871 | 17957 | BA_peak_1632_BA_peak_1633 | active | 2 |
| DS113291 | 75625 | 81431 | BA_peak_1674_BA_peak_1675 | active | 2 |
| DS113296 | 117524 | 122734 | BA_peak_1726_BA_peak_1727 | active | 2 |
| DS113297 | 126399 | 133499 | BA_peak_1737_BA_peak_1738 | active | 2 |
| DS113297 | 130138 | 147594 | BA_peak_1738_BA_peak_1739 | active | 5 |
| DS113298 | 2935 | 11175 | BA_peak_1743_BA_peak_1744 | active | 4 |
| DS113298 | 56798 | 79145 | BA_peak_1747_BA_peak_1748 | active | 11 |
| DS113300 | 102654 | 113422 | BA_peak_1763_BA_peak_1764 | active | 3 |
| DS113305 | 73150 | 90038 | BA_peak_1796_BA_peak_1797 | active | 6 |
| DS113306 | 41112 | 54568 | BA_peak_1802_BA_peak_1803 | active | 4 |
| DS113306 | 53782 | 64934 | BA_peak_1803_BA_peak_1804 | active | 3 |
| DS113309 | 13158 | 28995 | BA_peak_1832_BA_peak_1833 | active | 6 |
| DS113309 | 25739 | 44576 | BA_peak_1833_BA_peak_1834 | active | 3 |
| DS113310 | 29944 | 38069 | BA_peak_1837_BA_peak_1838 | active | 4 |
| DS113311 | 88398 | 95043 | BA_peak_1852_BA_peak_1853 | active | 3 |
| DS113313 | 20204 | 25939 | BA_peak_1860_BA_peak_1861 | active | 3 |
| DS113314 | 70440 | 78833 | BA_peak_1868_BA_peak_1869 | active | 2 |
| DS113315 | 102489 | 112029 | BA_peak_1883_BA_peak_1884 | active | 2 |
| DS113316 | 16807 | 29203 | BA_peak_1891_BA_peak_1892 | active | 6 |
| DS113316 | 31853 | 40863 | BA_peak_1894_BA_peak_1895 | active | 4 |

| | | | | | |
|---|---|---|---|---|---|
| DS113316 | 40118 | 45743 | BA_peak_1895_BA_peak_1896 | active | 2 |
| DS113319 | 122183 | 127869 | BA_peak_1920_BA_peak_1921 | active | 3 |
| DS113320 | 120560 | 127459 | BA_peak_1927_BA_peak_1928 | active | 4 |
| DS113320 | 126562 | 134661 | BA_peak_1928_BA_peak_1929 | active | 2 |
| DS113322 | 67267 | 79725 | BA_peak_1940_BA_peak_1941 | active | 2 |
| DS113324 | 41959 | 52492 | BA_peak_1954_BA_peak_1955 | active | 4 |
| DS113324 | 52099 | 75410 | BA_peak_1955_BA_peak_1956 | active | 6 |
| DS113326 | 21060 | 27054 | BA_peak_1974_BA_peak_1975 | active | 2 |
| DS113328 | 81566 | 89485 | BA_peak_1998_BA_peak_1999 | active | 2 |
| DS113330 | 82923 | 95477 | BA_peak_2018_BA_peak_2019 | active | 2 |
| DS113333 | 52949 | 64313 | BA_peak_2045_BA_peak_2046 | active | 2 |
| DS113333 | 63587 | 69700 | BA_peak_2046_BA_peak_2048 | active | 2 |
| DS113333 | 85156 | 99165 | BA_peak_2049_BA_peak_2050 | active | 6 |
| DS113333 | 98237 | 120574 | BA_peak_2050_BA_peak_2051 | active | 13 |
| DS113334 | 16194 | 29821 | BA_peak_2053_BA_peak_2054 | active | 6 |
| DS113334 | 75397 | 81478 | BA_peak_2061_BA_peak_2062 | active | 2 |
| DS113334 | 95857 | 105663 | BA_peak_2065_BA_peak_2066 | active | 3 |
| DS113334 | 105119 | 113251 | BA_peak_2066_BA_peak_2067 | active | 4 |
| DS113336 | 80665 | 85098 | BA_peak_2080_BA_peak_2081 | active | 2 |
| DS113337 | 20190 | 25534 | BA_peak_2086_BA_peak_2087 | active | 2 |
| DS113337 | 24917 | 41063 | BA_peak_2087_BA_peak_2088 | active | 8 |
| DS113337 | 83184 | 92263 | BA_peak_2090_BA_peak_2091 | active | 4 |
| DS113337 | 91811 | 108639 | BA_peak_2091_BA_peak_2092 | active | 8 |
| DS113338 | 78643 | 93526 | BA_peak_2101_BA_peak_2102 | active | 3 |
| DS113340 | 95194 | 111664 | BA_peak_2113_BA_peak_2114 | active | 5 |
| DS113344 | 21671 | 27415 | BA_peak_2141_BA_peak_2142 | active | 2 |
| DS113344 | 92666 | 103574 | BA_peak_2147_BA_peak_2148 | active | 2 |
| DS113344 | 102978 | 107966 | BA_peak_2148_BA_peak_2149 | active | 2 |
| DS113345 | 114581 | 126480 | BA_peak_2158_BA_peak_2159 | active | 3 |
| DS113348 | 60 | 4539 | BA_peak_2176_BA_peak_2177 | active | 2 |
| DS113348 | 28849 | 36218 | BA_peak_2178_BA_peak_2179 | active | 3 |
| DS113350 | 70371 | 82627 | BA_peak_2199_BA_peak_2200 | active | 2 |
| DS113350 | 109271 | 124720 | BA_peak_2205_BA_peak_2206 | active | 3 |
| DS113354 | 99056 | 123849 | BA_peak_2247_BA_peak_2248 | active | 11 |
| DS113356 | 19832 | 25451 | BA_peak_2255_BA_peak_2256 | active | 2 |
| DS113362 | 30844 | 46518 | BA_peak_2304_BA_peak_2305 | active | 8 |
| DS113362 | 87215 | 100419 | BA_peak_2309_BA_peak_2310 | active | 2 |
| DS113363 | 1352 | 6537 | BA_peak_2313_BA_peak_2314 | active | 2 |
| DS113363 | 6154 | 20815 | BA_peak_2314_BA_peak_2315 | active | 9 |
| DS113363 | 93958 | 101290 | BA_peak_2318_BA_peak_2319 | active | 3 |
| DS113366 | 38033 | 48123 | BA_peak_2331_BA_peak_2332 | active | 2 |
| DS113366 | 80597 | 90352 | BA_peak_2336_BA_peak_2337 | active | 2 |
| DS113366 | 87034 | 105597 | BA_peak_2337_BA_peak_2338 | active | 4 |
| DS113369 | 1884 | 17015 | BA_peak_2358_BA_peak_2359 | active | 6 |
| DS113370 | 24103 | 29588 | BA_peak_2365_BA_peak_2366 | active | 3 |
| DS113371 | 76023 | 95012 | BA_peak_2379_BA_peak_2380 | active | 3 |
| DS113373 | 56776 | 69048 | BA_peak_2388_BA_peak_2389 | active | 7 |

| | | | | | |
|---|---|---|---|---|---|
| DS113374 | 41792 | 50946 | BA_peak_2395_BA_peak_2396 | active | 4 |
| DS113379 | 87645 | 103327 | BA_peak_2427_BA_peak_2428 | active | 3 |
| DS113380 | 21692 | 31309 | BA_peak_2437_BA_peak_2438 | active | 5 |
| DS113382 | 6937 | 16183 | BA_peak_2456_BA_peak_2457 | active | 3 |
| DS113382 | 46817 | 59026 | BA_peak_2463_BA_peak_2464 | active | 4 |
| DS113382 | 100481 | 108403 | BA_peak_2469_BA_peak_2470 | active | 3 |
| DS113386 | 9843 | 28280 | BA_peak_2493_BA_peak_2494 | active | 2 |
| DS113386 | 102663 | 110468 | BA_peak_2500_BA_peak_2501 | active | 3 |
| DS113387 | 19402 | 29644 | BA_peak_2503_BA_peak_2504 | active | 3 |
| DS113387 | 81907 | 87596 | BA_peak_2509_BA_peak_2510 | active | 2 |
| DS113388 | 47530 | 58887 | BA_peak_2519_BA_peak_2520 | active | 6 |
| DS113390 | 31860 | 48622 | BA_peak_2536_BA_peak_2537 | active | 5 |
| DS113393 | 54825 | 68772 | BA_peak_2561_BA_peak_2562 | active | 6 |
| DS113395 | 28126 | 39596 | BA_peak_2573_BA_peak_2574 | active | 4 |
| DS113395 | 61174 | 71805 | BA_peak_2575_BA_peak_2576 | active | 3 |
| DS113395 | 72519 | 79778 | BA_peak_2577_BA_peak_2578 | active | 5 |
| DS113397 | 1716 | 12320 | BA_peak_2588_BA_peak_2589 | active | 2 |
| DS113397 | 70551 | 83184 | BA_peak_2593_BA_peak_2594 | active | 7 |
| DS113398 | 29530 | 43896 | BA_peak_2596_BA_peak_2597 | active | 2 |
| DS113400 | 12807 | 29285 | BA_peak_2607_BA_peak_2608 | active | 5 |
| DS113400 | 28912 | 48688 | BA_peak_2608_BA_peak_2609 | active | 7 |
| DS113400 | 52556 | 63281 | BA_peak_2611_BA_peak_2612 | active | 4 |
| DS113402 | 44886 | 54405 | BA_peak_2636_BA_peak_2637 | active | 3 |
| DS113402 | 105960 | 112243 | BA_peak_2638_BA_peak_2639 | active | 2 |
| DS113404 | 95309 | 108067 | BA_peak_2651_BA_peak_2652 | active | 5 |
| DS113409 | 93495 | 104370 | BA_peak_2677_BA_peak_2678 | active | 5 |
| DS113410 | 13364 | 27911 | BA_peak_2684_BA_peak_2685 | active | 6 |
| DS113410 | 47102 | 56135 | BA_peak_2686_BA_peak_2687 | active | 4 |
| DS113411 | 51351 | 58725 | BA_peak_2690_BA_peak_2691 | active | 4 |
| DS113411 | 67906 | 80096 | BA_peak_2697_BA_peak_2698 | active | 4 |
| DS113411 | 77300 | 86079 | BA_peak_2698_BA_peak_2699 | active | 3 |
| DS113414 | 54369 | 64424 | BA_peak_2717_BA_peak_2718 | active | 4 |
| DS113414 | 63431 | 71978 | BA_peak_2718_BA_peak_2719 | active | 6 |
| DS113419 | 36685 | 51108 | BA_peak_2738_BA_peak_2739 | active | 5 |
| DS113419 | 53683 | 74388 | BA_peak_2740_BA_peak_2741 | active | 8 |
| DS113422 | 7723 | 24145 | BA_peak_2750_BA_peak_2751 | active | 6 |
| DS113427 | 20542 | 39830 | BA_peak_2769_BA_peak_2770 | active | 6 |
| DS113427 | 40550 | 47490 | BA_peak_2771_BA_peak_2772 | active | 3 |
| DS113427 | 46671 | 60472 | BA_peak_2772_BA_peak_2773 | active | 2 |
| DS113427 | 93718 | 100704 | BA_peak_2777_BA_peak_2778 | active | 4 |
| DS113428 | 73334 | 81364 | BA_peak_2782_BA_peak_2783 | active | 2 |
| DS113430 | 37876 | 55090 | BA_peak_2799_BA_peak_2800 | active | 7 |
| DS113434 | 20377 | 29429 | BA_peak_2829_BA_peak_2830 | active | 5 |
| DS113434 | 94502 | 102841 | BA_peak_2834_BA_peak_2835 | active | 2 |
| DS113435 | 7709 | 21758 | BA_peak_2836_BA_peak_2837 | active | 5 |
| DS113437 | 15321 | 23513 | BA_peak_2852_BA_peak_2853 | active | 3 |
| DS113437 | 42102 | 69473 | BA_peak_2856_BA_peak_2857 | active | 13 |

| | | | | | |
|---|---|---|---|---|---|
| DS113440 | 80471 | 94180 | BA_peak_2878_BA_peak_2879 | active | 4 |
| DS113442 | 86171 | 97512 | BA_peak_2890_BA_peak_2891 | active | 3 |
| DS113443 | 37314 | 41315 | BA_peak_2896_BA_peak_2897 | active | 2 |
| DS113443 | 47297 | 54634 | BA_peak_2899_BA_peak_2900 | active | 2 |
| DS113445 | 50736 | 62695 | BA_peak_2912_BA_peak_2913 | active | 3 |
| DS113446 | 17568 | 24093 | BA_peak_2917_BA_peak_2918 | active | 4 |
| DS113448 | 4118 | 10744 | BA_peak_2925_BA_peak_2926 | active | 3 |
| DS113448 | 76959 | 83883 | BA_peak_2933_BA_peak_2934 | active | 2 |
| DS113450 | 23302 | 28724 | BA_peak_2944_BA_peak_2945 | active | 2 |
| DS113452 | 74819 | 82543 | BA_peak_2962_BA_peak_2963 | active | 2 |
| DS113452 | 80916 | 87957 | BA_peak_2963_BA_peak_2964 | active | 2 |
| DS113454 | 13061 | 18846 | BA_peak_2970_BA_peak_2971 | active | 2 |
| DS113454 | 18503 | 23226 | BA_peak_2971_BA_peak_2972 | active | 2 |
| DS113455 | 85761 | 95338 | BA_peak_2982_BA_peak_2983 | active | 3 |
| DS113456 | 71556 | 94397 | BA_peak_2988_BA_peak_2989 | active | 5 |
| DS113457 | 53779 | 60924 | BA_peak_2991_BA_peak_2992 | active | 2 |
| DS113459 | 12427 | 21867 | BA_peak_3000_BA_peak_3001 | active | 4 |
| DS113461 | 4324 | 11750 | BA_peak_3012_BA_peak_3013 | active | 2 |
| DS113463 | 79712 | 88644 | BA_peak_3026_BA_peak_3027 | active | 2 |
| DS113464 | 59256 | 66010 | BA_peak_3029_BA_peak_3030 | active | 3 |
| DS113469 | 73103 | 81059 | BA_peak_3066_BA_peak_3067 | active | 2 |
| DS113471 | 80241 | 92319 | BA_peak_3077_BA_peak_3078 | active | 6 |
| DS113472 | 3333 | 8585 | BA_peak_3079_BA_peak_3080 | active | 2 |
| DS113475 | 42665 | 63421 | BA_peak_3096_BA_peak_3097 | active | 6 |
| DS113481 | 63416 | 81939 | BA_peak_3122_BA_peak_3123 | active | 5 |
| DS113482 | 49010 | 64060 | BA_peak_3126_BA_peak_3127 | active | 3 |
| DS113484 | 50644 | 65035 | BA_peak_3137_BA_peak_3138 | active | 5 |
| DS113484 | 64271 | 70943 | BA_peak_3138_BA_peak_3139 | active | 2 |
| DS113485 | 43526 | 53402 | BA_peak_3146_BA_peak_3147 | active | 5 |
| DS113485 | 70730 | 79850 | BA_peak_3151_BA_peak_3152 | active | 4 |
| DS113485 | 79449 | 87671 | BA_peak_3152_BA_peak_3153 | active | 3 |
| DS113486 | 13126 | 21247 | BA_peak_3156_BA_peak_3157 | active | 2 |
| DS113486 | 64402 | 73082 | BA_peak_3159_BA_peak_3160 | active | 3 |
| DS113491 | 6426 | 15498 | BA_peak_3176_BA_peak_3177 | active | 3 |
| DS113492 | 81919 | 90427 | BA_peak_3187_BA_peak_3188 | active | 2 |
| DS113494 | 55078 | 60217 | BA_peak_3199_BA_peak_3200 | active | 2 |
| DS113495 | 59130 | 67071 | BA_peak_3206_BA_peak_3207 | active | 2 |
| DS113496 | 22592 | 30104 | BA_peak_3216_BA_peak_3217 | active | 3 |
| DS113496 | 37930 | 47663 | BA_peak_3219_BA_peak_3220 | active | 5 |
| DS113497 | 62929 | 79806 | BA_peak_3230_BA_peak_3231 | active | 5 |
| DS113498 | 16568 | 23588 | BA_peak_3234_BA_peak_3235 | active | 2 |
| DS113500 | 45815 | 56907 | BA_peak_3248_BA_peak_3251 | active | 2 |
| DS113501 | 71703 | 84658 | BA_peak_3258_BA_peak_3259 | active | 3 |
| DS113502 | 64326 | 74798 | BA_peak_3265_BA_peak_3266 | active | 3 |
| DS113506 | 11437 | 16277 | BA_peak_3282_BA_peak_3283 | active | 2 |
| DS113507 | 58624 | 66443 | BA_peak_3291_BA_peak_3292 | active | 2 |
| DS113512 | 46042 | 55561 | BA_peak_3317_BA_peak_3318 | active | 2 |

| | | | | | |
|---|---|---|---|---|---|
| DS113514 | 64487 | 74467 | BA_peak_3339_BA_peak_3340 | active | 4 |
| DS113516 | 71817 | 81640 | BA_peak_3352_BA_peak_3353 | active | 3 |
| DS113517 | 63274 | 86015 | BA_peak_3358_BA_peak_3359 | active | 10 |
| DS113520 | 38565 | 46398 | BA_peak_3373_BA_peak_3374 | active | 2 |
| DS113520 | 64915 | 77550 | BA_peak_3376_BA_peak_3377 | active | 7 |
| DS113522 | 3420 | 13993 | BA_peak_3387_BA_peak_3388 | active | 4 |
| DS113522 | 38492 | 54601 | BA_peak_3391_BA_peak_3392 | active | 7 |
| DS113523 | 32282 | 43685 | BA_peak_3394_BA_peak_3395 | active | 3 |
| DS113523 | 69262 | 76248 | BA_peak_3396_BA_peak_3397 | active | 2 |
| DS113525 | 6009 | 18727 | BA_peak_3404_BA_peak_3405 | active | 2 |
| DS113531 | 12101 | 38552 | BA_peak_3434_BA_peak_3436 | active | 9 |
| DS113532 | 31208 | 42411 | BA_peak_3439_BA_peak_3440 | active | 5 |
| DS113534 | 28474 | 42286 | BA_peak_3447_BA_peak_3448 | active | 8 |
| DS113535 | 25669 | 33154 | BA_peak_3450_BA_peak_3451 | active | 4 |
| DS113535 | 32373 | 44378 | BA_peak_3451_BA_peak_3452 | active | 3 |
| DS113539 | 8596 | 21926 | BA_peak_3479_BA_peak_3480 | active | 5 |
| DS113539 | 21115 | 26594 | BA_peak_3480_BA_peak_3481 | active | 3 |
| DS113541 | 10646 | 27587 | BA_peak_3491_BA_peak_3492 | active | 8 |
| DS113542 | 46119 | 53908 | BA_peak_3497_BA_peak_3499 | active | 2 |
| DS113547 | 24072 | 29814 | BA_peak_3532_BA_peak_3533 | active | 2 |
| DS113549 | 38788 | 55243 | BA_peak_3548_BA_peak_3549 | active | 3 |
| DS113549 | 55588 | 66487 | BA_peak_3549_BA_peak_3551 | active | 4 |
| DS113549 | 65770 | 73972 | BA_peak_3551_BA_peak_3552 | active | 2 |
| DS113550 | 19997 | 29287 | BA_peak_3554_BA_peak_3555 | active | 6 |
| DS113551 | 2326 | 16693 | BA_peak_3564_BA_peak_3565 | active | 5 |
| DS113552 | 11129 | 20036 | BA_peak_3566_BA_peak_3567 | active | 3 |
| DS113554 | 19503 | 33838 | BA_peak_3581_BA_peak_3582 | active | 6 |
| DS113554 | 60799 | 75689 | BA_peak_3588_BA_peak_3589 | active | 3 |
| DS113555 | 8390 | 14577 | BA_peak_3591_BA_peak_3592 | active | 3 |
| DS113558 | 6141 | 13111 | BA_peak_3605_BA_peak_3606 | active | 2 |
| DS113558 | 11456 | 18066 | BA_peak_3606_BA_peak_3607 | active | 3 |
| DS113558 | 42240 | 63588 | BA_peak_3611_BA_peak_3612 | active | 7 |
| DS113558 | 64968 | 74466 | BA_peak_3614_BA_peak_3615 | active | 3 |
| DS113565 | 37757 | 45348 | BA_peak_3650_BA_peak_3651 | active | 2 |
| DS113565 | 51728 | 65474 | BA_peak_3652_BA_peak_3653 | active | 5 |
| DS113569 | 53348 | 65032 | BA_peak_3670_BA_peak_3672 | active | 3 |
| DS113572 | 46989 | 74198 | BA_peak_3685_BA_peak_3686 | active | 14 |
| DS113573 | 4724 | 9883 | BA_peak_3689_BA_peak_3690 | active | 2 |
| DS113573 | 8923 | 20063 | BA_peak_3690_BA_peak_3691 | active | 8 |
| DS113574 | 41958 | 50338 | BA_peak_3696_BA_peak_3697 | active | 2 |
| DS113580 | 47637 | 55405 | BA_peak_3731_BA_peak_3732 | active | 3 |
| DS113586 | 14691 | 36421 | BA_peak_3756_BA_peak_3757 | active | 9 |
| DS113586 | 52096 | 65808 | BA_peak_3758_BA_peak_3759 | active | 6 |
| DS113593 | 44954 | 52628 | BA_peak_3785_BA_peak_3786 | active | 3 |
| DS113594 | 17193 | 22035 | BA_peak_3788_BA_peak_3789 | active | 2 |
| DS113596 | 53537 | 61183 | BA_peak_3796_BA_peak_3797 | active | 3 |
| DS113602 | 58383 | 64743 | BA_peak_3824_BA_peak_3825 | active | 4 |

| | | | | | |
|---|---|---|---|---|---|
| DS113605 | 4282 | 17709 | BA_peak_3838_BA_peak_3839 | active | 2 |
| DS113607 | 3997 | 12334 | BA_peak_3850_BA_peak_3851 | active | 5 |
| DS113608 | 26103 | 36308 | BA_peak_3853_BA_peak_3854 | active | 2 |
| DS113608 | 35497 | 60122 | BA_peak_3854_BA_peak_3855 | active | 11 |
| DS113609 | 46573 | 58039 | BA_peak_3862_BA_peak_3864 | active | 2 |
| DS113615 | 31882 | 40269 | BA_peak_3883_BA_peak_3884 | active | 5 |
| DS113617 | 32836 | 48862 | BA_peak_3890_BA_peak_3892 | active | 11 |
| DS113621 | 13903 | 21641 | BA_peak_3906_BA_peak_3907 | active | 3 |
| DS113621 | 26542 | 32717 | BA_peak_3908_BA_peak_3909 | active | 2 |
| DS113621 | 31967 | 46377 | BA_peak_3909_BA_peak_3910 | active | 6 |
| DS113622 | 13673 | 17060 | BA_peak_3915_BA_peak_3916 | active | 2 |
| DS113625 | 65934 | 71983 | BA_peak_3928_BA_peak_3929 | active | 2 |
| DS113630 | 6083 | 26600 | BA_peak_3945_BA_peak_3946 | active | 12 |
| DS113631 | 12747 | 20469 | BA_peak_3952_BA_peak_3953 | active | 4 |
| DS113632 | 18238 | 25718 | BA_peak_3960_BA_peak_3961 | active | 2 |
| DS113633 | 20394 | 35939 | BA_peak_3964_BA_peak_3965 | active | 4 |
| DS113641 | 9580 | 26534 | BA_peak_3997_BA_peak_3998 | active | 4 |
| DS113644 | 11213 | 21732 | BA_peak_4004_BA_peak_4005 | active | 4 |
| DS113650 | 42710 | 58715 | BA_peak_4025_BA_peak_4026 | active | 6 |
| DS113654 | 20679 | 43327 | BA_peak_4038_BA_peak_4039 | active | 7 |
| DS113657 | 6056 | 15055 | BA_peak_4049_BA_peak_4050 | active | 3 |
| DS113659 | 49292 | 59361 | BA_peak_4065_BA_peak_4066 | active | 3 |
| DS113659 | 57624 | 64021 | BA_peak_4066_BA_peak_4067 | active | 2 |
| DS113664 | 5821 | 35318 | BA_peak_4086_BA_peak_4087 | active | 10 |
| DS113667 | 1351 | 15237 | BA_peak_4095_BA_peak_4096 | active | 6 |
| DS113674 | 53436 | 64906 | BA_peak_4120_BA_peak_4121 | active | 3 |
| DS113677 | 50938 | 62874 | BA_peak_4130_BA_peak_4131 | active | 7 |
| DS113680 | 21567 | 32927 | BA_peak_4145_BA_peak_4146 | active | 6 |
| DS113681 | 39027 | 57686 | BA_peak_4151_BA_peak_4152 | active | 10 |
| DS113691 | 58216 | 63781 | BA_peak_4191_BA_peak_4192 | active | 3 |
| DS113694 | 47822 | 57239 | BA_peak_4207_BA_peak_4209 | active | 3 |
| DS113695 | 43083 | 49355 | BA_peak_4214_BA_peak_4215 | active | 2 |
| DS113699 | 6492 | 15317 | BA_peak_4235_BA_peak_4236 | active | 4 |
| DS113701 | 20899 | 26417 | BA_peak_4240_BA_peak_4241 | active | 2 |
| DS113704 | 11490 | 36757 | BA_peak_4249_BA_peak_4250 | active | 13 |
| DS113704 | 32912 | 44184 | BA_peak_4250_BA_peak_4251 | active | 2 |
| DS113708 | 34345 | 44371 | BA_peak_4268_BA_peak_4269 | active | 4 |
| DS113712 | 13179 | 19059 | BA_peak_4284_BA_peak_4285 | active | 3 |
| DS113712 | 20771 | 33033 | BA_peak_4287_BA_peak_4288 | active | 3 |
| DS113712 | 29840 | 40004 | BA_peak_4288_BA_peak_4289 | active | 2 |
| DS113713 | 44458 | 52641 | BA_peak_4297_BA_peak_4298 | active | 2 |
| DS113714 | 13915 | 18961 | BA_peak_4300_BA_peak_4301 | active | 2 |
| DS113714 | 34448 | 40458 | BA_peak_4305_BA_peak_4306 | active | 3 |
| DS113714 | 42352 | 55394 | BA_peak_4308_BA_peak_4309 | active | 6 |
| DS113718 | 18743 | 23684 | BA_peak_4313_BA_peak_4314 | active | 2 |
| DS113721 | 25304 | 39890 | BA_peak_4322_BA_peak_4323 | active | 5 |
| DS113721 | 50131 | 59820 | BA_peak_4324_BA_peak_4325 | active | 4 |

| | | | | | |
|---|---|---|---|---|---|
| DS113723 | 42719 | 55601 | BA_peak_4332_BA_peak_4333 | active | 6 |
| DS113725 | 3056 | 12615 | BA_peak_4338_BA_peak_4339 | active | 4 |
| DS113725 | 12233 | 21650 | BA_peak_4339_BA_peak_4340 | active | 2 |
| DS113729 | 11337 | 20415 | BA_peak_4352_BA_peak_4353 | active | 5 |
| DS113732 | 38551 | 46109 | BA_peak_4366_BA_peak_4367 | active | 3 |
| DS113733 | 50860 | 55334 | BA_peak_4373_BA_peak_4374 | active | 2 |
| DS113734 | 5039 | 36517 | BA_peak_4375_BA_peak_4376 | active | 8 |
| DS113734 | 44287 | 56268 | BA_peak_4377_BA_peak_4378 | active | 4 |
| DS113739 | 17353 | 25612 | BA_peak_4396_BA_peak_4397 | active | 3 |
| DS113740 | 3003 | 14144 | BA_peak_4402_BA_peak_4403 | active | 4 |
| DS113744 | 28182 | 37474 | BA_peak_4419_BA_peak_4420 | active | 2 |
| DS113744 | 36272 | 49223 | BA_peak_4420_BA_peak_4421 | active | 5 |
| DS113744 | 48484 | 56401 | BA_peak_4421_BA_peak_4422 | active | 2 |
| DS113745 | 47154 | 55354 | BA_peak_4424_BA_peak_4425 | active | 2 |
| DS113748 | 29383 | 43944 | BA_peak_4436_BA_peak_4437 | active | 3 |
| DS113751 | 345 | 6357 | BA_peak_4445_BA_peak_4446 | active | 2 |
| DS113752 | 24150 | 34048 | BA_peak_4452_BA_peak_4453 | active | 3 |
| DS113758 | 38524 | 45430 | BA_peak_4485_BA_peak_4486 | active | 3 |
| DS113763 | 18195 | 38237 | BA_peak_4496_BA_peak_4497 | active | 10 |
| DS113763 | 37733 | 55467 | BA_peak_4497_BA_peak_4498 | active | 8 |
| DS113771 | 29258 | 39700 | BA_peak_4525_BA_peak_4526 | active | 2 |
| DS113776 | 31319 | 42900 | BA_peak_4546_BA_peak_4547 | active | 5 |
| DS113776 | 43638 | 52614 | BA_peak_4548_BA_peak_4549 | active | 4 |
| DS113777 | 1069 | 5883 | BA_peak_4550_BA_peak_4551 | active | 2 |
| DS113782 | 44330 | 49279 | BA_peak_4575_BA_peak_4576 | active | 2 |
| DS113787 | 6195 | 15716 | BA_peak_4589_BA_peak_4590 | active | 2 |
| DS113787 | 15098 | 29768 | BA_peak_4590_BA_peak_4591 | active | 5 |
| DS113789 | 30587 | 47581 | BA_peak_4598_BA_peak_4599 | active | 4 |
| DS113792 | 6163 | 17487 | BA_peak_4607_BA_peak_4608 | active | 3 |
| DS113792 | 20032 | 31551 | BA_peak_4610_BA_peak_4611 | active | 6 |
| DS113795 | 11862 | 22035 | BA_peak_4614_BA_peak_4615 | active | 3 |
| DS113795 | 20000 | 30285 | BA_peak_4615_BA_peak_4616 | active | 2 |
| DS113796 | 26044 | 38468 | BA_peak_4623_BA_peak_4624 | active | 6 |
| DS113797 | 9140 | 18316 | BA_peak_4627_BA_peak_4628 | active | 4 |
| DS113799 | 32531 | 41568 | BA_peak_4636_BA_peak_4637 | active | 3 |
| DS113808 | 25169 | 37953 | BA_peak_4661_BA_peak_4662 | active | 4 |
| DS113814 | 23542 | 43217 | BA_peak_4682_BA_peak_4683 | active | 4 |
| DS113819 | 30738 | 41617 | BA_peak_4699_BA_peak_4700 | active | 3 |
| DS113830 | 6535 | 21243 | BA_peak_4729_BA_peak_4730 | active | 5 |
| DS113837 | 4876 | 22530 | BA_peak_4749_BA_peak_4750 | active | 3 |
| DS113842 | 7003 | 37345 | BA_peak_4769_BA_peak_4770 | active | 11 |
| DS113846 | 9411 | 18913 | BA_peak_4785_BA_peak_4786 | active | 3 |
| DS113847 | 33381 | 42751 | BA_peak_4790_BA_peak_4791 | active | 2 |
| DS113848 | 31249 | 39351 | BA_peak_4797_BA_peak_4798 | active | 2 |
| DS113851 | 6506 | 12493 | BA_peak_4809_BA_peak_4810 | active | 3 |
| DS113858 | 12489 | 20018 | BA_peak_4826_BA_peak_4827 | active | 4 |
| DS113863 | 8073 | 16269 | BA_peak_4841_BA_peak_4842 | active | 5 |

| | | | | | |
|---|---|---|---|---|---|
| DS113863 | 15883 | 22639 | BA_peak_4842_BA_peak_4843 | active | 3 |
| DS113865 | 30724 | 38960 | BA_peak_4850_BA_peak_4851 | active | 2 |
| DS113869 | 38140 | 46987 | BA_peak_4867_BA_peak_4868 | active | 2 |
| DS113873 | 1143 | 8334 | BA_peak_4880_BA_peak_4881 | active | 2 |
| DS113874 | 33966 | 39677 | BA_peak_4890_BA_peak_4891 | active | 2 |
| DS113879 | 11638 | 27561 | BA_peak_4904_BA_peak_4905 | active | 4 |
| DS113883 | 1181 | 5974 | BA_peak_4918_BA_peak_4919 | active | 2 |
| DS113888 | 26917 | 39461 | BA_peak_4936_BA_peak_4937 | active | 2 |
| DS113893 | 11450 | 20023 | BA_peak_4951_BA_peak_4952 | active | 4 |
| DS113898 | 26355 | 32288 | BA_peak_4964_BA_peak_4965 | active | 2 |
| DS113899 | 13719 | 26471 | BA_peak_4968_BA_peak_4969 | active | 8 |
| DS113904 | 5245 | 10505 | BA_peak_4984_BA_peak_4985 | active | 2 |
| DS113908 | 8857 | 23575 | BA_peak_5002_BA_peak_5003 | active | 6 |
| DS113910 | 7707 | 12862 | BA_peak_5005_BA_peak_5006 | active | 2 |
| DS113910 | 35428 | 41465 | BA_peak_5008_BA_peak_5009 | active | 2 |
| DS113911 | 11963 | 22526 | BA_peak_5012_BA_peak_5013 | active | 3 |
| DS113913 | 24314 | 29818 | BA_peak_5015_BA_peak_5016 | active | 3 |
| DS113915 | 7454 | 14182 | BA_peak_5023_BA_peak_5024 | active | 3 |
| DS113920 | 15779 | 22761 | BA_peak_5032_BA_peak_5033 | active | 3 |
| DS113921 | 18989 | 32898 | BA_peak_5038_BA_peak_5039 | active | 6 |
| DS113923 | 772 | 28848 | BA_peak_5041_BA_peak_5042 | active | 11 |
| DS113923 | 28325 | 35676 | BA_peak_5042_BA_peak_5043 | active | 3 |
| DS113923 | 35339 | 42833 | BA_peak_5043_BA_peak_5044 | active | 3 |
| DS113931 | 18698 | 34575 | BA_peak_5061_BA_peak_5062 | active | 2 |
| DS113932 | 10741 | 21617 | BA_peak_5063_BA_peak_5064 | active | 6 |
| DS113940 | 21067 | 33121 | BA_peak_5084_BA_peak_5085 | active | 6 |
| DS113941 | 15596 | 33194 | BA_peak_5088_BA_peak_5089 | active | 7 |
| DS113948 | 27991 | 38280 | BA_peak_5109_BA_peak_5110 | active | 4 |
| DS113958 | 4628 | 15399 | BA_peak_5140_BA_peak_5141 | active | 5 |
| DS113964 | 8519 | 18559 | BA_peak_5159_BA_peak_5160 | active | 4 |
| DS113972 | 11326 | 21559 | BA_peak_5180_BA_peak_5181 | active | 3 |
| DS113972 | 30013 | 38846 | BA_peak_5183_BA_peak_5185 | active | 4 |
| DS113976 | 10834 | 20279 | BA_peak_5198_BA_peak_5199 | active | 4 |
| DS113983 | 10601 | 17640 | BA_peak_5232_BA_peak_5233 | active | 2 |
| DS113985 | 6633 | 15379 | BA_peak_5239_BA_peak_5240 | active | 2 |
| DS113985 | 14647 | 27173 | BA_peak_5240_BA_peak_5241 | active | 5 |
| DS113985 | 25631 | 34095 | BA_peak_5241_BA_peak_5242 | active | 3 |
| DS113990 | 12175 | 16289 | BA_peak_5255_BA_peak_5256 | active | 2 |
| DS113997 | 25341 | 30205 | BA_peak_5279_BA_peak_5280 | active | 3 |
| DS114002 | 5523 | 15474 | BA_peak_5287_BA_peak_5288 | active | 5 |
| DS114004 | 19283 | 26831 | BA_peak_5291_BA_peak_5292 | active | 2 |
| DS114017 | 19644 | 28316 | BA_peak_5328_BA_peak_5329 | active | 3 |
| DS114021 | 5893 | 18039 | BA_peak_5337_BA_peak_5338 | active | 5 |
| DS114026 | 609 | 11861 | BA_peak_5350_BA_peak_5351 | active | 3 |
| DS114027 | 5590 | 14357 | BA_peak_5353_BA_peak_5354 | active | 2 |
| DS114033 | 6766 | 17162 | BA_peak_5367_BA_peak_5368 | active | 3 |
| DS114033 | 20877 | 26526 | BA_peak_5370_BA_peak_5371 | active | 2 |

| | | | | | |
|---|---|---|---|---|---|
| DS114046 | 18032 | 26023 | BA_peak_5402_BA_peak_5403 | active | 2 |
| DS114049 | 9143 | 21422 | BA_peak_5406_BA_peak_5407 | active | 5 |
| DS114056 | 14245 | 26509 | BA_peak_5419_BA_peak_5420 | active | 8 |
| DS114061 | 975 | 20042 | BA_peak_5431_BA_peak_5432 | active | 7 |
| DS114069 | 15038 | 27103 | BA_peak_5457_BA_peak_5458 | active | 5 |
| DS114071 | 27643 | 32212 | BA_peak_5462_BA_peak_5463 | active | 2 |
| DS114076 | 5930 | 22201 | BA_peak_5472_BA_peak_5473 | active | 5 |
| DS114088 | 11805 | 30633 | BA_peak_5501_BA_peak_5502 | active | 6 |
| DS114090 | 5656 | 18198 | BA_peak_5505_BA_peak_5506 | active | 7 |
| DS114092 | 7538 | 21105 | BA_peak_5510_BA_peak_5511 | active | 3 |
| DS114105 | 101 | 8654 | BA_peak_5529_BA_peak_5530 | active | 2 |
| DS114114 | 7164 | 18341 | BA_peak_5541_BA_peak_5542 | active | 5 |
| DS114127 | 8079 | 21494 | BA_peak_5566_BA_peak_5567 | active | 8 |
| DS114138 | 13002 | 25192 | BA_peak_5592_BA_peak_5593 | active | 5 |
| DS114146 | 12806 | 19150 | BA_peak_5603_BA_peak_5604 | active | 3 |
| DS114149 | 13073 | 22281 | BA_peak_5607_BA_peak_5608 | active | 5 |
| DS114159 | 15897 | 26333 | BA_peak_5629_BA_peak_5630 | active | 5 |
| DS114161 | 12542 | 25897 | BA_peak_5634_BA_peak_5635 | active | 5 |
| DS114162 | 351 | 13344 | BA_peak_5636_BA_peak_5637 | active | 3 |
| DS114162 | 8722 | 20983 | BA_peak_5637_BA_peak_5638 | active | 2 |
| DS114164 | 7724 | 20383 | BA_peak_5640_BA_peak_5641 | active | 4 |
| DS114165 | 4081 | 14075 | BA_peak_5642_BA_peak_5643 | active | 5 |
| DS114165 | 13313 | 21707 | BA_peak_5643_BA_peak_5644 | active | 3 |
| DS114169 | 7193 | 22574 | BA_peak_5652_BA_peak_5653 | active | 5 |
| DS114186 | 94 | 17719 | BA_peak_5694_BA_peak_5695 | active | 9 |
| DS114215 | 2749 | 11803 | BA_peak_5740_BA_peak_5741 | active | 4 |
| DS114223 | 1707 | 9385 | BA_peak_5754_BA_peak_5755 | active | 4 |
| DS114239 | 4124 | 23149 | BA_peak_5786_BA_peak_5787 | active | 8 |
| DS114240 | 15408 | 22705 | BA_peak_5790_BA_peak_5791 | active | 3 |
| DS114244 | 3482 | 15949 | BA_peak_5796_BA_peak_5798 | active | 5 |
| DS114248 | 13160 | 18709 | BA_peak_5811_BA_peak_5812 | active | 2 |
| DS114251 | 8036 | 20169 | BA_peak_5820_BA_peak_5821 | active | 3 |
| DS114257 | 2660 | 13382 | BA_peak_5827_BA_peak_5828 | active | 2 |
| DS114262 | 3390 | 10845 | BA_peak_5835_BA_peak_5836 | active | 3 |
| DS114278 | 7380 | 15219 | BA_peak_5863_BA_peak_5864 | active | 3 |
| DS114279 | 6768 | 20745 | BA_peak_5865_BA_peak_5866 | active | 4 |
| DS114285 | 8270 | 20969 | BA_peak_5874_BA_peak_5875 | active | 8 |
| DS114333 | 712 | 8217 | BA_peak_5941_BA_peak_5942 | active | 2 |
| DS114335 | 6726 | 17352 | BA_peak_5947_BA_peak_5948 | active | 3 |
| DS114355 | 6807 | 15956 | BA_peak_5974_BA_peak_5975 | active | 3 |
| DS114383 | 8907 | 15660 | BA_peak_6015_BA_peak_6016 | active | 3 |
| DS114400 | 67 | 8947 | BA_peak_6037_BA_peak_6038 | active | 2 |
| DS114426 | 8597 | 17735 | BA_peak_6073_BA_peak_6074 | active | 3 |
| DS114429 | 4357 | 12823 | BA_peak_6076_BA_peak_6077 | active | 4 |
| DS114441 | 6437 | 15612 | BA_peak_6096_BA_peak_6097 | active | 2 |
| DS114454 | 5038 | 12306 | BA_peak_6109_BA_peak_6110 | active | 4 |
| DS114473 | 7629 | 15459 | BA_peak_6126_BA_peak_6127 | active | 4 |

| | | | | | |
|---|---|---|---|---|---|
| DS114496 | 1878 | 13895 | BA_peak_6152_BA_peak_6153 | active | 2 |
| DS114535 | 786 | 10345 | BA_peak_6200_BA_peak_6201 | active | 5 |
| DS114537 | 41 | 9243 | BA_peak_6204_BA_peak_6205 | active | 3 |
| DS114552 | 6685 | 13735 | BA_peak_6223_BA_peak_6224 | active | 2 |
| DS114604 | 1018 | 11217 | BA_peak_6273_BA_peak_6274 | active | 3 |
| DS114636 | 70 | 6223 | BA_peak_6300_BA_peak_6301 | active | 2 |
| DS114718 | 132 | 3888 | BA_peak_6359_BA_peak_6360 | active | 2 |
| DS114743 | 2204 | 8644 | BA_peak_6376_BA_peak_6377 | active | 3 |
| DS114759 | 75 | 4741 | BA_peak_6389_BA_peak_6390 | active | 2 |
| DS114773 | 1063 | 9515 | BA_peak_6396_BA_peak_6398 | active | 3 |
| DS114783 | 30 | 8331 | BA_peak_6402_BA_peak_6403 | active | 2 |
| DS114818 | 2802 | 8973 | BA_peak_6430_BA_peak_6431 | active | 3 |
| DS114908 | 1035 | 7658 | BA_peak_6483_BA_peak_6484 | active | 2 |
| DS114931 | 841 | 7504 | BA_peak_6500_BA_peak_6501 | active | 3 |
| DS115094 | 343 | 6231 | BA_peak_6573_BA_peak_6574 | active | 2 |

**Supplementary Table 8. BspA containing intervals**

| Interval name | Genes IDs | Product description | Type of interval |
|---|---|---|---|
| p5512-5513 | TVAG_437640 | conserved hypothetical protein | repressive |
| | **TVAG_437650** | **leucine-rich repeat protein BspA family** | |
| | **TVAG_437660** | **leucine-rich repeat protein BspA family** | |
| | **TVAG_437670** | **leucine-rich repeat protein BspA family** | |
| | **TVAG_437680** | **leucine-rich repeat protein BspA family** | |
| | TVAG_437690 | conserved hypothetical protein | |
| | TVAG_437700 | conserved hypothetical protein | |
| | TVAG_437710 | conserved hypothetical protein | |
| | TVAG_437720 | conserved hypothetical protein | |
| | TVAG_437730 | conserved hypothetical protein | |
| | TVAG_TEG_DS114093_1_1 | mariner transposase putative | |
| p4130-4131 | TVAG_125490 | conserved hypothetical protein | active |
| | TVAG_125500 | thioredoxin, putative | |
| | TVAG_125510 | thioredoxin, putative | |
| | TVAG_125520 | CAMK family protein kinase | |
| | **TVAG_125530** | **leucine-rich repeat protein, BspA family** | |
| | TVAG_125540 | high mobility group protein, putative | |
| | TVAG_125550 | ribosomal protein L3p, putative | |
| | TVAG_176930 | conserved hypothetical protein | |
| p2296-2297 | **TVAG_133310** | **leucine-rich repeat protein BspA family** | repressive |
| | TVAG_133320 | conserved hypothetical protein | |
| | **TVAG_133330** | **leucine-rich repeat protein BspA family** | |
| | **TVAG_133340** | **leucine-rich repeat protein BspA family** | |
| | TVAG_133350 | conserved hypothetical protein | |
| | **TVAG_133360** | **leucine-rich repeat protein BspA family** | |
| p560-561 | **TVAG_365610** | **leucine-rich repeat protein, BspA family** | active |
| | TVAG_365620 | ankyrin, putative | |
| | TVAG_365630 | hypothetical protein | |
| | TVAG_365640 | cytoplasmic dynein heavy chain, putative | |
| | TVAG_365650 | conserved hypothetical protein | |
| | TVAG_365660 | ctl transporter, putative | |
| | TVAG_365670 | hypothetical protein | |
| | TVAG_365680 | conserved hypothetical protein | |
| | TVAG_365690 | conserved hypothetical protein | |
| | TVAG_365700 | casein kinase, putative | |
| | TVAG_365710 | hypothetical protein | |
| | TVAG_365720 | casein kinase II beta chain, putative | |
| | TVAG_365730 | conserved hypothetical protein | |
| p271-272 | TVAG_456730 | 2-deoxyglucose-6-phosphate phosphatase, putative | active |
| | **TVAG_456740** | **leucine-rich repeat protein, BspA family** | |
| | TVAG_456750 | adenylate cyclase type vIII, putative | |
| | TVAG_456760 | conserved hypothetical protein | |
| | TVAG_456770 | iron-sulfur cluster assembly protein, putative | |
| | TVAG_456780 | conserved hypothetical protein | |
| p3434-3436 | TVAG_254810 | conserved hypothetical protein | active |
| | TVAG_254820 | conserved hypothetical protein | |
| | TVAG_254830 | conserved hypothetical protein | |
| | TVAG_254840 | 60S ribosomal protein L12, putative | |
| | **TVAG_254850** | **leucine-rich repeat protein, BspA family** | |
| | TVAG_254860 | conserved hypothetical protein | |
| | TVAG_254870 | conserved hypothetical protein | |
| | TVAG_254880 | ubiquitin-conjugating enzyme, putative | |
| | TVAG_254890 | pyruvate-flavodoxin oxidoreductase, putative | |
| p556-557 | TVAG_021830 | conserved hypothetical protein | active |
| | TVAG_021840 | adenylate cyclase, type VII, putative | |
| | TVAG_021850 | conserved hypothetical protein | |
| | **TVAG_021860** | **leucine-rich repeat protein, BspA family** | |
| | TVAG_021870 | myosin-4, putative | |
| | TVAG_021880 | nucleoprotein TPR, putative | |
| | TVAG_021890 | vacuolar ATP synthase subunit E, putative | |
| | TVAG_021900 | conserved hypothetical protein | |
| | TVAG_021910 | conserved hypothetical protein | |
| p605-606 | TVAG_129870 | ankyrin repeat-containing protein, putative | active |
| | TVAG_129880 | amino acid transporter, putative | |
| | TVAG_129890 | amino acid transporter, putative | |
| | TVAG_129900 | phosphatidate phosphatase, putative | |
| | TVAG_129910 | conserved hypothetical protein | |
| | TVAG_129920 | conserved hypothetical protein | |
| | TVAG_129930 | hypothetical protein | |
| | TVAG_129940 | conserved hypothetical protein | |
| | TVAG_129950 | myosin heavy chain, clone, putative | |
| | **TVAG_129960** | **leucine-rich repeat protein, BspA family** | |

**Supplementary Table 9. Intervals considered for 3C assay**

| Interval name | n° of genes | Type | Interval size | PCR product size |
|---|---|---|---|---|
| p271-272 | 6 | active | 17,4kbp | 290 |
| p456-457 | 6 | active | 25kbp | 208 |
| p560-561 | 13 | active | 39,7kbp | 259 |
| p690-692 | 11 | active | 23kbp | 109 |
| p806-807 | 8 | active | 19,8kbp | 117 |
| p1065-1066 | 6 | active | 28,3kbp | 113 |
| p4130-4131 | 8 | active | 12kbp | 184 |
| p228-230 | 30 | repressive | 47kbp | 158 |
| p657-658 | 13 | repressive | 19,2kbp | 209 |
| p786-787 | 10 | repressive | 18kbp | 291 |
| p1871-1873 | 22 | repressive | 34kbp | 167 |
| p2296-2297 | 6 | repressive | 12kbp | 285 |
| p55125-513 | 11 | repressive | 17kbp | 249 |

**Supplementary Table 10. Localization of 6mA peaks in TE genes**

| Peak position | n° of methylated TE genes | % of total methylated TE genes |
|---|---|---|
| TSS | 39 | 9.4 |
| CDS | 207 | 49.9 |
| TTS | 169 | 40.7 |

**Supplementary Table 11. 3C PCR primers**

| Interval name | Fw | Rv |
|---|---|---|
| p16-18 | ACTCTCTCCAGACACACTTGA | CTTGGGGCTGTGCAATTATTG |
| p271-272 | TGTTCGATTATTTCTAGAATACAATGTCCT | GTTTGTGTCTTGATAAAATTGGAACA |
| p456-457 | CAGCAGCGTCATCAAGAGTC | GGCTGCAAATACATCCTGGG |
| alternative p456-457 region | AGCCTTCAGAGTCATCACGA | GGCTGCAAATACATCCTGGG |
| p560-561 | AACAATTTCTGCCAACGACGTT | TCCAGAAGACTCATCAATAAGCAGA |
| p690-692 | AGCAAAACGAAACCCTTACACA | TGCGTTTTGTGTATCGAAGCA |
| alternative p690-692 region | AGCAAAACGAAACCCTTACACA | TGTTGAACGCGATGTCCTTG |
| p806-807 | TCCCTTAGTTATCCCAATCTGA | AGCACTTCACATACGGAAATAGC |
| alternative p806-807 region | TCCCTTAGTTATCCCAATCTGA | TCTGCCTTCTTCATTATGTCGT |
| p1065-1066 | AACAGGTCTTCTTTTGGGCT | GCATGCACTATATGTCGAAATGC |
| p4130-4131 | TTGTTACCCAATCCGAGGAGTTT | GGATTTCCTTATGGCGTGGTTATG |
| alternative p4130-4131 region | CACGCGGAATCATACATTTC | GGATTTCCTTATGGCGTGGTTATG |
| p228-230 | GCACAGCAACAACCACAAAC | AGGTTTGGCGATGAATGTGT |
| p657-658 | TCATAATCAAAGACCTCATAAACGA | CGAGCTCGCTATACAGTACTG |
| alternative p657-658 region | TCATAATCAAAGACCTCATAAACGA | CCGAAAACACGTGGGTCTTT |
| p786-787 | TGTATGAGCTGAGTGTAAATGTCT | GGAACAATTCAACCATGTACAA |
| alternative p786-787 region | GGAACGTTTATCACTGTGCC | GGAACAATTCAACCATGTACAA |
| p1102-1103 | ATCGAGCCAATTCACACCAT | CCGTGTGGCTGTGTGATAAC |
| alternative p1102-1103 region | TGAGTCATTTGCGTCAATGT | CCGTGTGGCTGTGTGATAAC |
| p1871-1873 | ACCACCCTCACACGTATGAA | GGTAATCTGATTGCTATTGATGC |
| alternative p1871-1873 region | ATAACAGCCGGATTTGCCCT | GGTAATCTGATTGCTATTGATGC |
| p2296-2297 | CTGCGTTCGCTTATCATCTGT | TGAAACGATACAAAACGATGAGAA |
| p55125-513 | TCTGATTTCTTTTACTCTCTCGTATATACT | GCCCAACCAGAAAGATCCAA |
| alternative p5512-5513 region | TCTGATTTCTTTTACTCTCTCGTATATACT | GCAAAATTCGAATGCTGATG |
